# Decoupling membrane permeabilization and antimicrobial activity in amphipathic peptides

**DOI:** 10.1101/2025.05.22.655621

**Authors:** John S. Albin, Bradley L. Pentelute

## Abstract

The prevailing model of cathelicidin function holds that peptide helicity leads to membrane permeabilization, which in turn leads to killing of gram-negative bacteria. Using a paired Ala-Aib mutagenesis approach to isolate sidechain and structure-dependent functions, we demonstrate here that the effect of helicity on gram-negative killing in a model cathelicidin-derived scaffold is species-dependent. We then leverage this insight to derive lead compounds with up to 32-fold improvements in selectivity for bacterial over mammalian cells. Further interrogation of the mechanistic basis for selectivity demonstrates that, while helicity may predict permeabilization, permeabilization does not predict killing of gram-negative bacteria. Thus, neither helicity nor permeabilization is a universal requirement for cathelicidin-mediated killing.

## Introduction

Antimicrobial resistance is a growing source of global morbidity and mortality, accounting for more than 1 million deaths per year in recent years with the potential to become the single largest cause of death worldwide in the coming decades^1–3^. One strategy to counteract the growing threat posed by antimicrobial resistance is to derive new chemical entities by mining and developing natural product host defense peptides (HDPs), which despite their potential have yet to reach clinical use^4–11^.

Humans express a variety of structurally diverse HDPs, including α defensins, β defensins, histatins, neuropeptides, and a number of miscellaneous protein cleavage products that have varying degrees of reported antimicrobial activity against different microbes^12–14^.

Humans also express a single cathelicidin, initially as a pro-peptide packaged within neutrophil granules, which undergoes cleavage to form the active peptide LL-37^15–17^. In its active form, LL-37 is a prototypical amphipathic helix that is thought to act through membrane disruption, though we have recently reported results to suggest that other mechanisms may contribute to its antimicrobial activity, consistent with an emerging literature on the non-membranolytic mechanisms of HDP action^18–21^. Regardless, the antimicrobial activity of LL-37 is thought to be dependent upon its helical structure, rendering this a valuable model in which to study associations between helical structure and functional phenotypes^22–25^. Thus, in the prevailing model of human cathelicidin activity against gram-negative bacteria, helicity leads to membrane permeabilization, which in turn leads to gram-negative cell killing.

Many prior reports have described the activities of diverse fragments of LL-37. Among these fragments, KR-12 spanning residues 18-29 of full-length LL-37 is generally regarded as the minimal unit of LL-37 activity^23,24,26–33^. Building on this work, we have recently used rational design principles involving the transposition of an LL-37 N-terminal biphenyl motif onto the KR-12 core to yield a short cathelicidin called FF-14 with >16-fold greater activity against gram-negative bacteria^34^. In related work, we have also identified mutations in the LL-37 oligomerization interface, I24 and L28, that drive LL-37 hemolytic activity^20^. Each of these oligomerization drivers is also found in FF-14 at positions I9 and L13, and thus may also contribute to the hemolytic activity of this short cathelicidin derivative.

Using a paired Ala-Aib mutagenesis approach to isolate structure- and sidechain-dependent functions in a model cathelicidin scaffold, we find that, although Aib can promote helical structure^35–38^, the effects of helicity on downstream activity against different cell types is species-dependent. We then leverage this distinction to develop short cathelicidin derivatives with up to 32-fold increases in selectivity for bacterial over mammalian cells. On further interrogating the mechanism underlying this selectivity, however, we find that efficient permeabilization does not predict efficient killing of gram-negative bacteria. Our work thus calls into question the premise that helicity and membrane permeabilization are universal requirements for cathelicidin-mediated killing of gram-negative bacteria.

## Results

### Antimicrobial and hemolytic activity correlate in grouped Ala mutants of FF-14

Our prior structure-activity studies of full-length LL-37 use grouped Ala mutagenesis to isolate the functions of distinct surfaces in LL-37 while retaining global helical structure. An important finding of this work is that derivatives with substitutions at key residues in the LL-37 oligomerization interface diminish activity against mammalian membranes, improving selectivity for bacterial over mammalian cells^20,39,40^.

To determine whether it would be possible to use this same surface mutagenesis approach to separate functions in the FF-14 template, we first made a series of grouped Ala mutants corresponding to those initially described in full-length LL-37^20^. As shown in **Figure 1A-B**, activity of these grouped Ala mutants against *E. coli* and *P. aeruginosa* was ablated, though substitution with a critical biphenyl motif [FF]^34^ rescued antimicrobial activity at a cost of increased hemolytic activity (**Figure 1C**). Importantly, however, m4 substitutions, which include the oligomerization-associated residues I9 and L13, lacked both antimicrobial and hemolytic activity. Thus, in contrast with the ability of LL-37 m4 and colistin to separate antibacterial and hemolytic functions, antimicrobial and hemolytic activity profiles in FF-14 derivatives remain coupled. Mutants tested are shown in **Figure 1D**.

**Figure 1.**
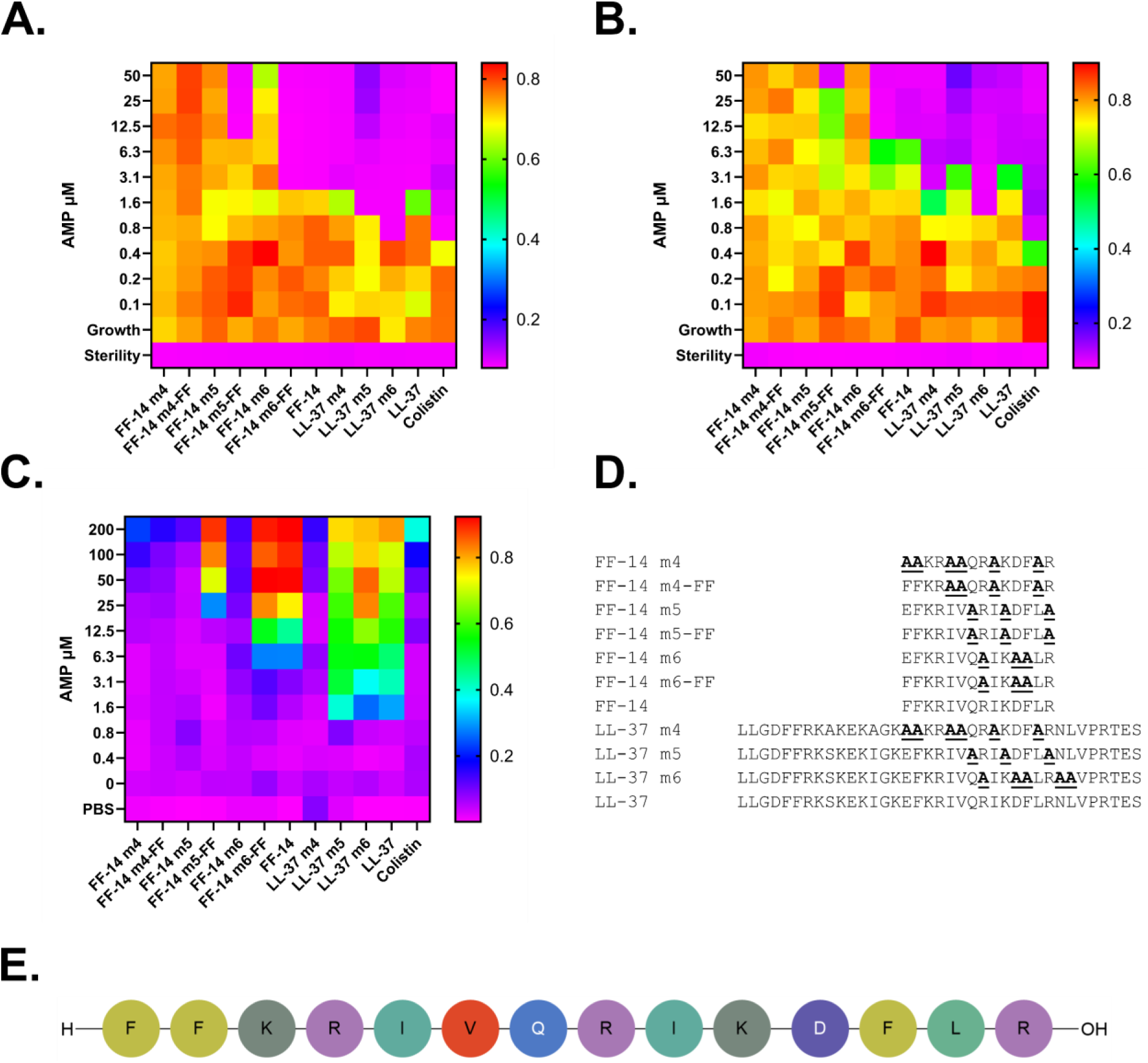
Antimicrobial and hemolytic activities correlate in grouped Ala mutants of FF-14. A-B. Antimicrobial susceptibility testing of *E. coli* (**A**) and *P. aeruginosa* (**B**) was carried out using a series of grouped Ala mutants based on those previously used in full-length LL-37. In contrast with the separation of antibacterial and hemolytic functions in LL-37 m4 as well as in the control antibiotic colistin, antibacterial and hemolytic functions remain coupled among FF-14 grouped Ala mutant derivatives. Data show the raw OD600 after 18 hours of incubation at 37 °C; warm colors indicate bacterial growth, while cool colors are low or absent growth. **C.** Hemolytic activity of each of the tested FF-14 derivatives as assessed by hemoglobin release after 18 hours of incubation at 37 °C. Cool colors indicate low signal at 414 nm, while warm colors are high signal; values are scaled to a 100% freeze-thaw hemolysis control. **D.** Sequence alignment showing each of the FF-14 mutants in **A-C** as well as the parent LL-37 mutants used as controls. **E.** Schematic depiction of the FF-14 scaffold showing the parent sequence.

### Both helicity and sidechain identity contribute to FF-14 function

Based on the results of **Figure 1**, we hypothesized that loss of function in FF-14 derivatives might be due to an associated loss of helicity in the context of multiple amino acid substitutions. To test this, we analyzed these mutants by circular dichroism (CD) spectroscopy. Spectra for each mutant are shown in **Figure S1A-G**. Helicity as estimated using BeStSel^41^ is summarized in **Figure S1H**. Although some mutants such as FF-14 m4 and FF-14 m4-FF appear to lose structure, the remaining grouped mutants retain at least intermediate helical character, with activity profiles driven more by the biphenyl motif at the first two positions than by helicity itself, consistent with our initial description of this scaffold^34^. Thus, both sidechain identities and helicity drive function in this template, complicating our ability to distinguish the effects of perturbation of each parameter on downstream functions. We therefore proceeded to a more systematic evaluation of individual positions in the FF-14 scaffold.

Figure S2 shows the data from Figure 1 in histogram format including error bars. The kinetic curves associated with the endpoint data provided in Figure 1 are shown in **Figures S3-14** (antimicrobial susceptibility testing) and **Figures S15-26** (hemolysis assays). A standardized, quantitative presentation of Figure 1 data is provided in **Table S1**.

### Antimicrobial and hemolytic activity correlate in single Ala mutants of FF-14

Although LL-37 tolerates up to at least eight simultaneous Ala mutants out of the 37-mer sequence without substantial loss of helical structure^20^, FF-14 appeared less tolerant of such alterations in Figure 1. We therefore hypothesized that FF-14 might better tolerate single Ala mutations to separate antimicrobial and hemolytic activities and proceeded to make a series of single Ala mutants covering the FF-14 sequence.

Antimicrobial susceptibility testing with these mutants against *E. coli* as shown in Figure 2A showed modest improvement in activity with mutants R8A and D11A^31^. While mutant D11A also showed improved activity against *P. aeruginosa* in Figure 2B, R8A demonstrated an increased MIC against this distinct comparator species. As shown in Figure 2C, hemolytic activity generally tracked with activity against *E. coli*. For example, the greatest losses in antimicrobial activity against *E. coli* in Figure 2A localized to I9A and L13A, corresponding to the oligomerization interface residues primarily responsible for separation of antibacterial and hemolytic activity in full-length LL-37, I24 and L28^20^. Thus, individual Ala mutations did not reveal a means by which to separate antimicrobial and hemolytic functions in FF-14.

**Figure 2.**
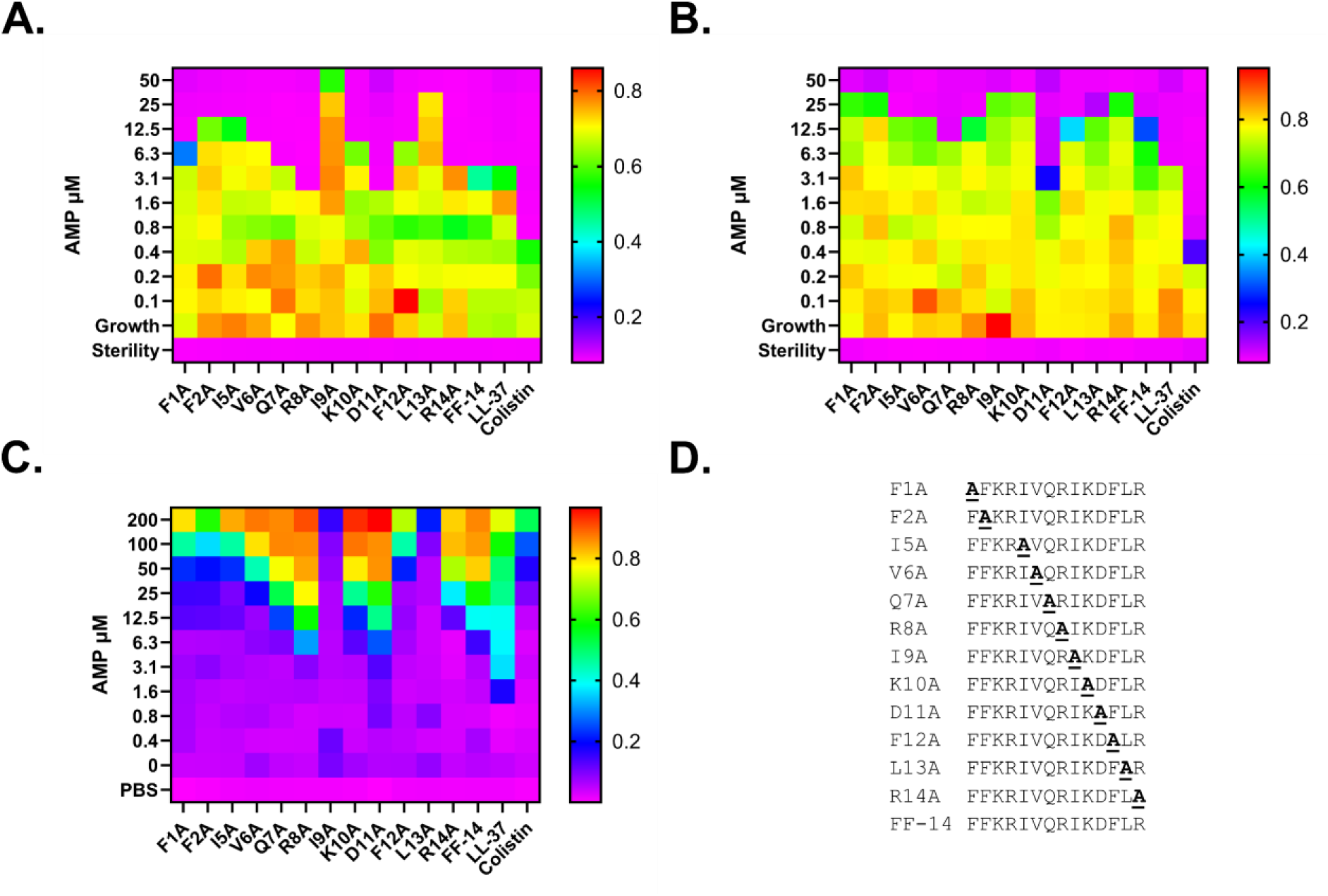
Antimicrobial and hemolytic activities of FF-14 single Ala mutants correlate with each other. A-B. Antimicrobial susceptibility testing of *E. coli* (**A**) and *P. aeruginosa* (**B**) was carried out using a series of single Ala mutants in FF-14. Data show the raw OD600 after 18 hours of incubation at 37 °C; warm colors indicate high signal representing bacterial growth, while cool colors are low signal and growth. **C.** Hemolytic activity of each of the tested FF-14 derivatives as assessed by hemoglobin release after 18 hours of incubation at 37 °C. Cool colors indicate low signal at 414 nm, while warm colors are high signal; values are scaled to a 100% freeze-thaw hemolysis control. **D.** Sequence alignment showing each of the FF-14 mutants in **A-C**.

Figure S27 shows the data from Figure 2 in histogram format including error bars. The kinetic curves associated with the endpoint data provided in Figure 2 are shown in **Figures S28-42** (antimicrobial susceptibility testing) and **Figures S43-57** (hemolysis assays). A standardized, quantitative presentation of Figure 2 data is provided in **Table S2**.

### Binary correlation between antimicrobial activity and FF-14 Ala mutant helicity

Based on the loss of helicity associated with FF-14 m4 grouped mutants in Figure 1 and **Figure S1**, we hypothesized that helicity might correlate with antimicrobial activity among single Ala mutants. To assess this, we again collected CD spectra from each of the single Ala mutants used in Figure 2. As shown in **Figure S58**, strong or absent helicity was generally associated with strong or weak antimicrobial activity, respectively. For example, the R8A and D11A mutants are strongly helical and more active than wild-type FF-14 against *E. coli*, while the I9A and L13A mutants are disordered and largely inactive. Correlation at intermediate levels of helicity, however, was modest. Thus, helicity appears predictive of activity primarily at the extremes. Overall, these results are consistent with prior studies demonstrating the importance of helical structure for LL-37 activity^17,22^. Individual spectra are provided in **Figure S59**.

### Aminoisobutyric acid (Aib) incorporation improves cathelicidin activity against *E. coli* but not *P. aeruginosa*

One method of better supporting structure in helical peptides is to incorporate α,α-disubstituted amino acids such as Aib (*e.g.*, ^35,42^), which is also a prominent feature in helical fungal natural product peptaibols^43–46^. We therefore reasoned that generating a single mutant Aib dataset corresponding to the Ala mutants in Figure 2 would allow us to isolate the structure-specific requirements underlying FF-14 functions independent of sidechain identity, which may in turn highlight a pathway to rationally incorporating modifications at residues I9 and L13 to separate antimicrobial and hemolytic activity as in full-length LL-37^20^.

To test this hypothesis, we completed a structural scan of FF-14 in which we used Aib to replace each residue in FF-14 rather than Ala, thus maintaining sidechain equivalence while introducing conformational restriction. As shown in Figures 3A-B and **Table S3**, antimicrobial activity with Aib substitution was generally improved against *E. coli* at all residues relative to Ala. This was particularly true of F2, I5, and I9, where substitution of Aib in place of Ala resulted in 8-fold improvements in MIC against *E. coli*. Despite these improvements against *E. coli*, the effect of Aib substitution on MICs against *P. aeruginosa* was neutral, while effects on hemolytic activity were intermediate between these extremes (Figure 3C). Thus, the sidechain-neutral conformational constraints introduced by Aib exert their primary effects at the level of FF-14 activity against *E. coli*, with lesser effects on hemolytic activity and no substantial effect on antimicrobial activity against *P. aeruginosa*. A summary of the mutants used in these studies is shown in Figure 3D.

**Figure 3.**
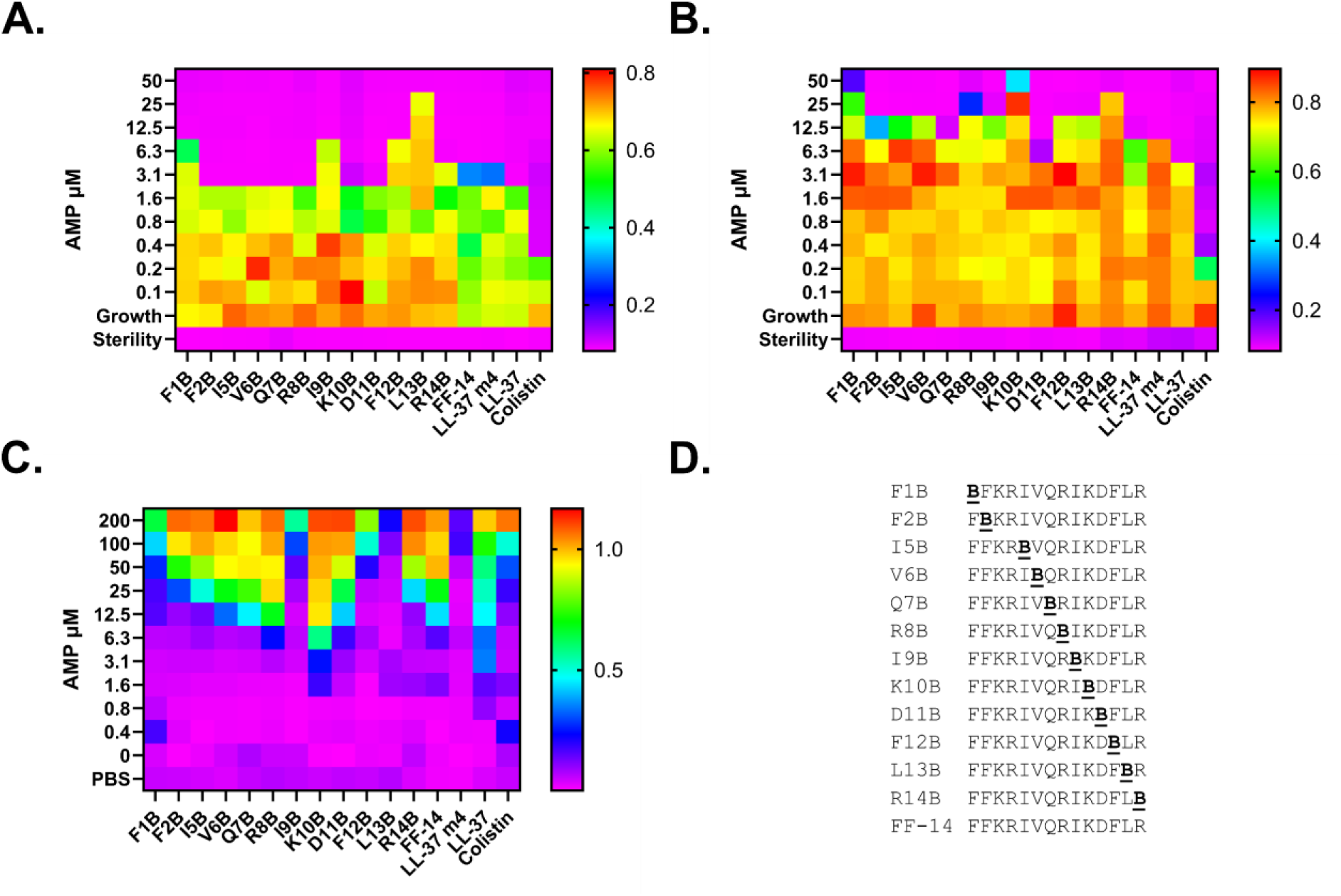
Aib substitution improves FF-14 antimicrobial activity against *E. coli*, but not *P. aeruginosa*. Although antimicrobial frequently improved against *E. coli* on substitution with Aib along the length of FF-14, this effect was not observed on MICs against *P. aeruginosa*. **A-B.** Antimicrobial susceptibility testing of *E. coli* (**A**) and *P. aeruginosa* (**B**) was carried out using a series of single Aib mutants in FF-14. Data show the raw OD600 after 18 hours of incubation at 37 °C; warm colors indicate high signal representing bacterial growth, while cool colors are low signal and growth. **C.** Hemolytic activity of each of the tested FF-14 derivatives as assessed by hemoglobin release after 18 hours of incubation at 37 °C. Cool colors indicate low signal at 414 nm, while warm colors are high signal; values are scaled to a 100% freeze-thaw hemolysis control. **D.** Sequence alignment showing each of the FF-14 mutants in **A-C**.

Figure S60 shows the data from Figure 3 in histogram format including error bars. The kinetic curves associated with the endpoint data provided in Figure 3 are shown in **Figures S61-76** (antimicrobial susceptibility testing) and **Figures S77-92** (hemolysis assays). A standardized, quantitative presentation of Figure 3 data is provided in **Table S3**.

### Paired Ala-Aib scanning distinguishes species-specific effects of helicity on FF-14 functions

Building on the comprehensive dataset generated in the course of Figures 2-3, we next sought to derive general principles about the relationships between FF-14 structure and associated functions. As shown in **Table 1**, overall helicity increased an average of 3.2-fold with Aib substitution, which was mirrored by an average 3.3-fold improvement in antimicrobial activity against *E. coli* among Aib mutants. This effect on helicity was not evenly distributed throughout the scaffold, however, but rather appeared partially dependent on local sequence context, with a range of fold-changes from 0.7-6.7 and strong effects at both the hydrophobic (*e.g.*, I9 6.7-fold and V6 5.3-fold) and hydrophilic (*e.g.*, K10 5.6-fold and R14 4.6-fold) surfaces. Cathelicidin activity against *E. coli* therefore tracks closely with helicity, but the ability to induce helicity is influenced by Aib positioning within the scaffold.

**Table 1.** Paired Ala-Aib helix scanning distinguishes the effects of helical content on FF-14 functional activities. This table summarizes the quantitative effects of replacing a given residue in FF-14 with Aib in place of Ala. Antimicrobial activity against *E. coli* tracks closely with increased helical content, while antimicrobial activity against *P. aeruginosa* does not. Hemolytic activity demonstrates intermediate dependence, allowing for a modest increase in cell type selectivity as measured against the *E. coli* MIC throughout this paper. % Helix at 37 °C shows the percentage of helical content in each peptide as estimated by BeStSel based on the spectra shown in **Supplementary Figures 58** and **92**. % Possible divides the absolute % helix change observed by the total possible % helix change (to a maximum of 100%) to account for different baseline helical content among peptides. Fold-change is derived by dividing the Aib percentage by the Ala percentage. MIC indicates the minimum inhibitory concentrations of each peptide against *E. coli* or *P. aeruginosa* as shown in **Figures 2** and **3** and associated **Supplementary Figures**, with the fold change derived by dividing the Ala MIC by the Aib MIC, such that larger numbers indicate more potent antimicrobial activity in Aib mutants. 20% Hemolysis indicates the highest peptide concentration with less than 20% hemolysis as shown in **Figures 2** and **3** and associated **Supplementary Figures**, with the fold change derived by dividing the Ala concentration by the Aib concentration, such that larger numbers indicate more toxic activity in Aib-containing derivatives. Fold Hemolysis / MIC indicates the ratio of the 20% Hemolysis threshold for each peptide divided by the MIC for each peptide against *E. coli*. The fold change is derived by dividing the Aib ratio by the Ala ratio, such that larger numbers indicate improved *in vitro* therapeutic indices.

|  | % Helix at 37 °C |  |  |  | MIC *E. coli* |  |  | MIC *Pseudomonas* |  |  | 20% Hemolysis |  |  | Fold Hemolysis / MIC |  |  |
| --- | --- | --- | --- | --- | --- | --- | --- | --- | --- | --- | --- | --- | --- | --- | --- | --- |
|  | Ala | Aib | % Possible | Fold Δ B:A | Ala | Aib | Fold Δ A:B | Ala | Aib | Fold Δ A:B | Ala | Aib | Fold Δ A:B | Ala | Aib | Fold Δ B:A |
| **F1** | 39.6 | 51.7 | 0.20 | 1.3 | 12.5 | 12.5 | 1.0 | 50 | 100 | 0.5 | 25 | 25 | 1.0 | 2.0 | 2.0 | 1.0 |
| **F2** | 16.7 | 74.2 | 0.69 | 4.4 | 25 | 3.1 | 8.0 | 50 | 25 | 2.0 | 25 | 12.5 | 2.0 | 1.0 | 4.0 | 4.0 |
| **I5** | 55.3 | 91.6 | 0.81 | 1.7 | 25 | 3.1 | 8.0 | 25 | 25 | 1.0 | 25 | 12.5 | 2.0 | 1.0 | 4.0 | 4.0 |
| **V6** | 18.7 | 100 | 1.00 | 5.3 | 12.5 | 3.1 | 4.0 | 25 | 25 | 1.0 | 25 | 6.3 | 4.0 | 2.0 | 2.0 | 1.0 |
| **Q7** | 26.5 | 96.5 | 0.95 | 3.6 | 6.3 | 3.1 | 2.0 | 12.5 | 12.5 | 1.0 | 6.3 | 6.3 | 1.0 | 1.0 | 2.0 | 2.0 |
| **R8** | 100 | 100 | n/a | 1.0 | 3.1 | 3.1 | 1.0 | 25 | 50 | 0.5 | 6.3 | 3.1 | 2.0 | 2.0 | 1.0 | 0.5 |
| **I9** | 8.1 | 54 | 0.50 | 6.7 | 100 | 12.5 | 8.0 | 50 | 25 | 2.0 | 200 | 50 | 4.0 | 2.0 | 4.0 | 2.0 |
| **K10** | 16.2 | 90.3 | 0.88 | 5.6 | 12.5 | 3.1 | 4.0 | 50 | 100 | 0.5 | 6.3 | 1.6 | 4.0 | 0.5 | 0.5 | 1.0 |
| **D11** | 92.7 | 68.7 | n/a | 0.7 | 3.1 | 3.1 | 1.0 | 6.3 | 6.3 | 1.0 | 3.1 | 6.3 | 0.5 | 1.0 | 2.0 | 2.0 |
| **F12** | 75.9 | 84.7 | 0.37 | 1.1 | 12.5 | 12.5 | 1.0 | 25 | 25 | 1.0 | 50 | 25 | 2.0 | 4.0 | 2.0 | 0.5 |
| **L13** | 10.3 | 18.2 | 0.09 | 1.8 | 50 | 50 | 1.0 | 25 | 25 | 1.0 | 200 | 100 | 2.0 | 4.0 | 2.0 | 0.5 |
| **R14** | 16 | 74.1 | 0.69 | 4.6 | 6.3 | 6.3 | 1.0 | 50 | 50 | 1.0 | 12.5 | 12.5 | 1.0 | 2.0 | 2.0 | 1.0 |
| **Avg** | 39.7 | 75.3 | 0.62 |  | 22.4 | 9.6 |  | 32.8 | 39.1 |  | 48.7 | 21.7 |  | 1.9 | 2.3 |  |
|  |  |  | Avg Δ | 3.2 |  | Avg Δ | 3.3 |  | Avg Δ | 1.0 |  | Avg Δ | 2.1 |  | Avg Δ | 1.6 |

Despite the three-fold changes in helicity and antimicrobial activity against *E. coli* effected by substitution with Aib over Ala, the average effect of Aib substitution on antimicrobial activity against *P. aeruginosa* was neutral. Consistent with differential effects of helicity on different target cell types, hemolytic activity among Aib substituted mutants increased 2-fold compared with Ala, occupying an intermediate position between effects on *E. coli* and effects on *P. aeruginosa*. The net effect of Aib substitution on selectivity for activity against bacterial over mammalian cells is thus a modest 1.6-fold average improvement. CD spectra for Aib-substituted mutants are shown in **Figure S93**.

### Improved cell type selectivity of combinatorial Aib substitutions

Given the ability of single Aib mutants to promote helical structure, we reasoned that double mutants might yield further improvements, particularly where each individual residue was found in **Table 1** to promote helicity. To test this hypothesis, we made a series of four dual Aib substitutions at *i,i+4* positions covering all possible combinations with this spacing along the helical hydrophobic surface (1+5, 2+6, 5+9, and 9+13). Among these, all except for I9B+L13B retained appreciable activity against *E. coli* (Figure 4A), with F2B+V6B being the most active.

**Figure 4.**
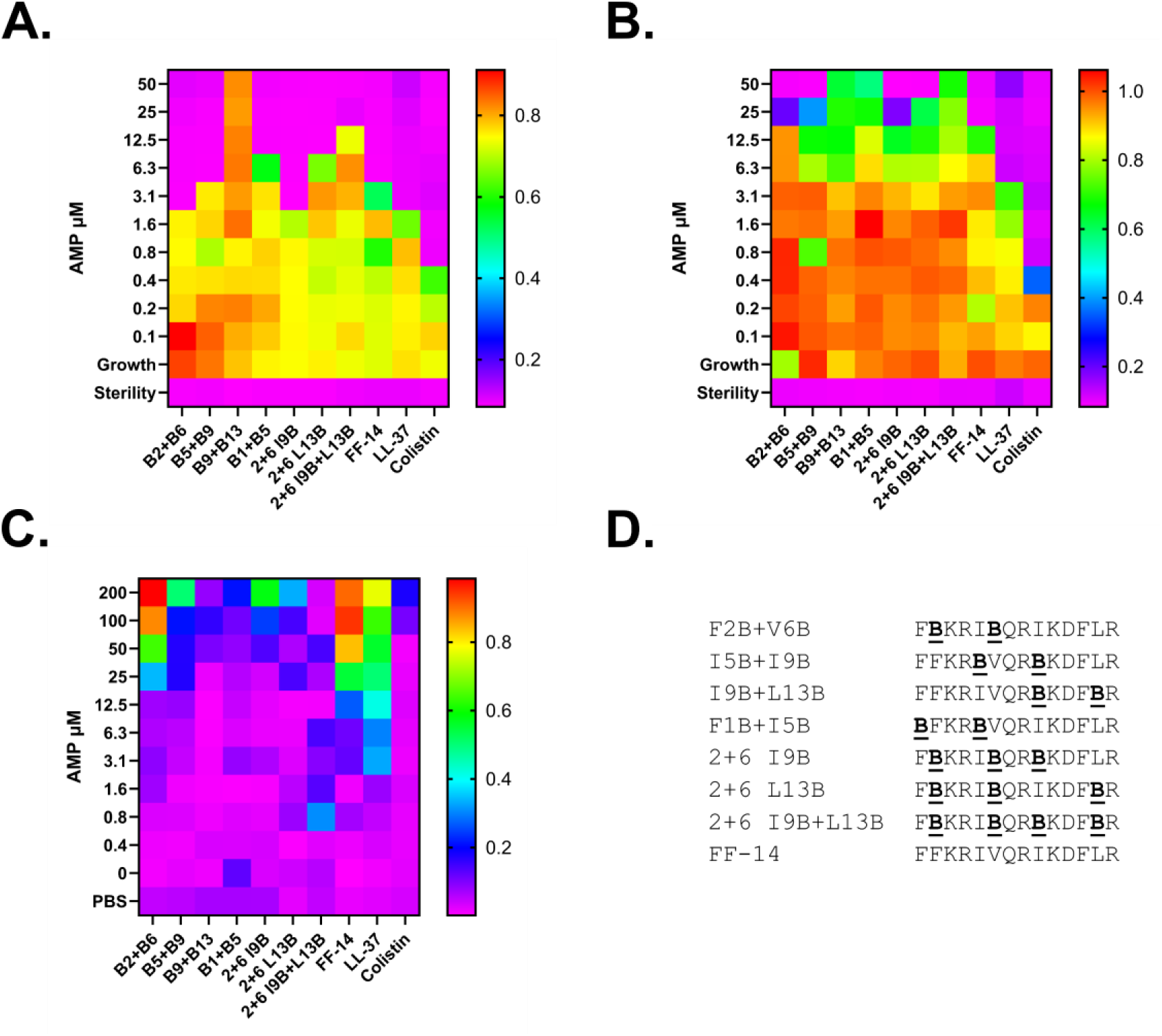
A modified FF-14 scaffold F2B+V6B with an additional I9B modification retains antimicrobial activity against *E. coli* while demonstrating diminished hemolytic activity. F2B+V6B with I9B retains a low-micromolar MIC against *E. coli* while demonstrating diminished hemolytic activity, leading to an overall 32-fold increase in cell type selectivity over the parent LL-37 peptide (see also **Supplementary Table 4**). Activity of multiply-Aib substituted peptides against *P. aeruginosa*, however, was generally poor. **A-B.** Antimicrobial susceptibility testing of *E. coli* (**A**) and *P. aeruginosa* (**B**) was carried out using a series of mutants with 2-4 Aib substitutions. Data show the raw OD600 after 18 hours of incubation at 37 °C; warm colors indicate higher bacterial growth, while cool colors are low / no growth. **C.** Hemolytic activity of each of the tested FF-14 derivatives as assessed by hemoglobin release after 18 hours of incubation at 37 °C. Cool colors indicate low signal at 414 nm, while warm colors are high signal; values are scaled to a 100% freeze-thaw hemolysis control. **D.** Sequence alignment showing each of the FF-14 Aib mutants in **A-C**.

These also generally retained strong helical character, again with the exception of I9B+L13B (**Figure S94**). Reinforcing the context dependence of these substitutions, however, we noted that the effects of multiple Aib substitutions on helicity were not additive. That is, the helicity of some double mutants was diminished relative to the best of their individual substitutions – *e.g.*, I5B is 91.6% helical, but I5B+I9B is 72.3% helical; I9B is 54% helical while I9B+L13B is 32.8% helical.

In contrast with its effect on activity against *E. coli*, double Aib substitution was generally detrimental to FF-14 activity against *P. aeruginosa* in Figure 4B despite strong helical character in some of these mutants. This echoed the results of **Table 1**, in which helicity was associated with activity against *E. coli* but not *P. aeruginosa*, again suggesting distinct structural requirements for FF-14 killing of different gram-negative species beyond helical structure alone.

To determine whether it would be possible to substitute additional Aib mutations that might contribute to improved selectivity of the parent template, we made three additional mutants adding LL-37 I9B and/or L13B to the F2B+V6B scaffold. Among these, activity remained equal to that of parent F2B+V6B when adding I9B, but with an increase of two dilutions in the threshold hemolysis level to 50 µM, thus overall yielding a 32-fold improvement in cell type selectivity over parent LL-37 (Figure 4C**, Table S4**). To a lesser extent, this was also true of L13B and I9B+L13B mutants within the F2B+V6B scaffold, albeit at a loss of antimicrobial potency against *E. coli* as quantified in **Table S4**. Despite these improvements against *E. coli*, however, the Aib mutants tested again showed diminished activity against *P. aeruginosa* (Figure 4B).

Thus, a combination of three Aib residues within the FF-14 scaffold can improve antimicrobial activity against *E. coli* while diminishing hemolytic activity, yielding up to 32-fold improvements in cell type selectivity. Application of these same substitutions to *P. aeruginosa*, however, paradoxically decreases antimicrobial potency. Understanding such distinctions in antimicrobial killing may enable rational development of narrower spectrum antibiotics based on naturally evolved HDP templates.

Figure S95 shows the data from Figure 4 in histogram format. **Figures S96-105** (antimicrobial susceptibility testing) and **S106-115** (hemolysis) show the kinetic curves associated with the endpoint measures in Figure 4. A standardized, quantitative presentation of Figure 4 data is provided in **Table S4**.

### Emergent effects limit further substitution in Aib-induced helices

Given the ability of Aib to improve helicity, we next asked whether pairing Aib residues in the F2B+V6B scaffold with canonical amino acid substitutions might improve upon the effects of the equivalent substitutions in FF-14. Reasoning that such effects might best focus on features less likely to impact intrinsic helical character, we focused our efforts on the hydrophilic face of FF-14 as well as I9 and L13.

As shown in Figure 5, substitution of selected positions with Ala in the F2B+V6B scaffold yields overall improvement for certain residues. For example, while I9A ablates activity in the FF-14 template (Figure 2), activity of this mutant in the context of F2B+V6B is 25 µM with a neutral effect on overall cell type selectivity relative to full-length LL-37 (Figure 5, **Tables S5-6**). For most Ala mutations within the F2B+V6B scaffold, however, this overall change in selectivity decreases, driven primarily by diminished antimicrobial activity of mutants in the F2B+V6B scaffold compared with FF-14 – for example, at residues R8 and D11 (**Table S6**). Thus, despite the ability of Aib substitutions in FF-14 to support helical structure and, in the case of *E. coli*, antimicrobial activity, combinatorial dynamics may limit further substitutions. Engineering of cathelicidin and other helical amphipathic templates may therefore benefit from approaches capable of directly assaying large combinatorial sequence spaces^47–49^.

**Figure 5.**
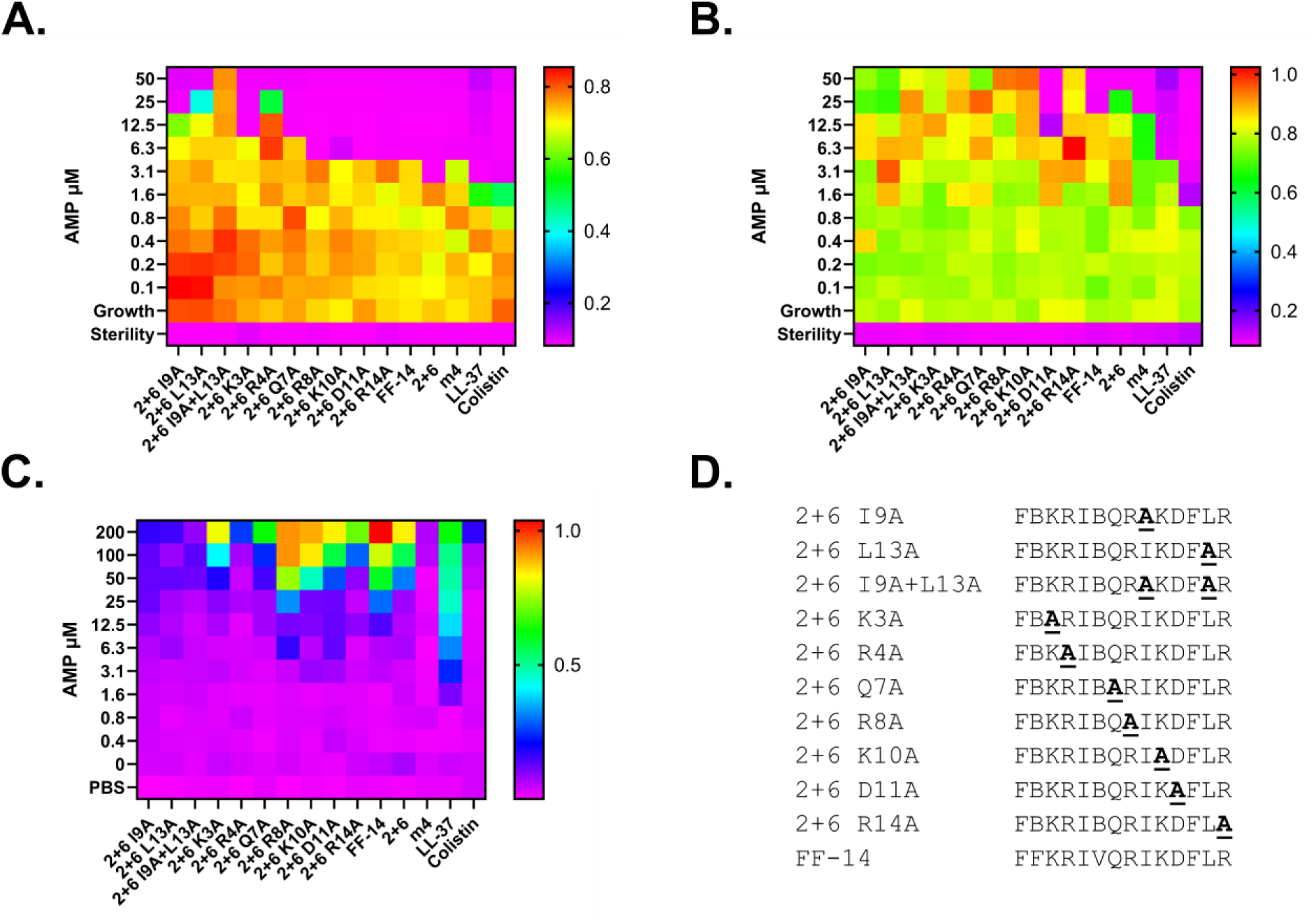
Emergent effects limit further substitution in Aib-induced helices. While activity of some derivatives such as F2B+V6B + I9A is improved against *E. coli* over the equivalent FF-14 I9A mutant, other substitutions such as R8A and D11A no longer improve upon the antimicrobial activity of the parent scaffold and activity against *P. aeruginosa* was poor across the mutants tested. **A-B.** Antimicrobial susceptibility testing of *E. coli* (**A**) and *P. aeruginosa* (**B**) was carried out using a series of single Ala mutations in FF-14 F2B+V6B. Data show the raw OD600 after 18 hours of incubation at 37 °C; warm colors indicate high signal representing bacterial growth, while cool colors are low signal and growth. **C.** Hemolytic activity of each of the tested FF-14 derivatives as assessed by hemoglobin release after 18 hours of incubation at 37 °C. Cool colors indicate low signal at 414 nm, while warm colors are high signal; values are scaled to a 100% freeze-thaw hemolysis control. **D.** Sequence alignment showing each of the mutants in **A-C**.

Figure S116 shows the data from Figure 5 in histogram format. **Figures S117-131** (antimicrobial susceptibility testing) and **S132-146** (hemolysis) show the kinetic curves associated with the endpoint measures in Figure 5.

### Increased helicity results in increased outer and inner membrane permeabilization

To better understand the mechanistic basis for distinct effects of helicity on activity against *E. coli* and *P. aeruginosa*, we carried out dual membrane permeabilization assays in which each cell type was combined with N-phenyl-naphthylamine (NPN) as a marker of outer membrane (OM) permeabilization and propidium iodide (PI) as a marker of inner membrane (IM) permeabilization. We then incubated these mixtures with selected peptides over a range of concentrations for 240 minutes with monitoring of fluorescence at 15-minute intervals. Specifically, we chose the Q7A / Q7B and K10A / K10B pairs as two instances in which Aib substitution led to the greatest increase in helicity with a given peptide – with a corresponding effect on activity against *E. coli* but not *P. aeruginosa*. We further included the F2B+V6B scaffold and F2B+V6B with I9B as examples of a pair with improved cell type selectivity as well as FF-14 and LL-37 controls.

Figure 6 shows a single point permeabilization index that represents the product of the maximum permeabilization achieved by each mutant and the proportion of the incubation required to achieve the maximum permeabilization activity of each individual mutant – thus incorporating both intensity and kinetic components into a single measure of permeabilization. By this metric, substitution of Aib at both Q7 and K10 results in increased permeabilization of both the IM and OM in both *E. coli* and *P. aeruginosa*, while F2B+V6B and F2B+V6B with I9B display permeabilization comparable to the parent FF-14 scaffold. Thus, increased helicity generally results in increased inner and outer membrane permeabilization in both species tested. Kinetic curves showing all time points and concentrations tested are provided in **Figures S147-150**.

**Figure 6.**
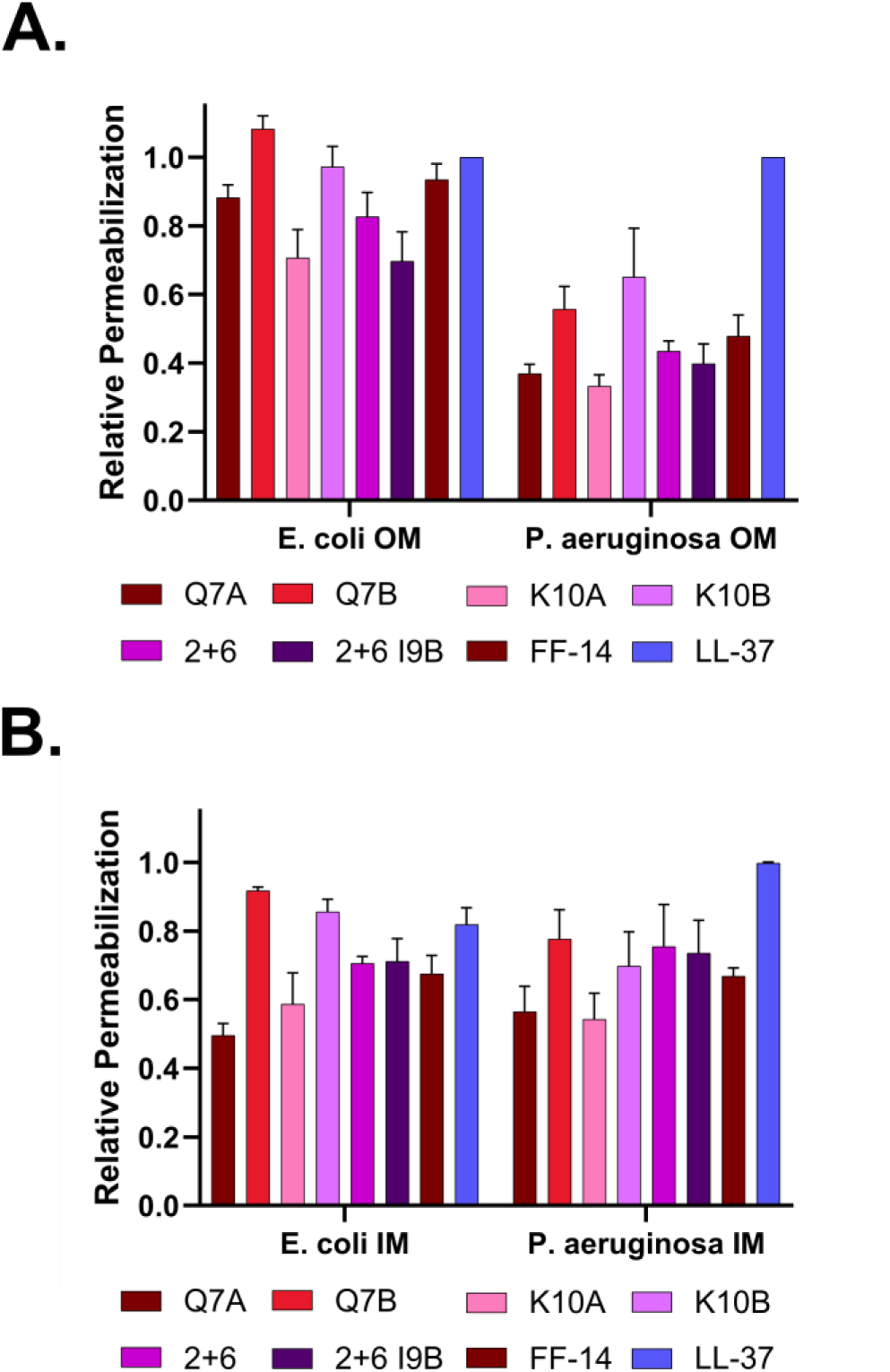
Increased helical content in short cathelicidin derivatives results in increased inner and outer membrane permeabilization in gram-negative bacteria. Dual membrane permeabilization assays were carried out using N-phenyl-naphthylamine (NPN) as a reporter of outer membrane (OM) permeabilization and propidium iodide (PI) as a reporter of inner membrane (IM) permeabilization. *E. coli* (ATCC 25922) or *P. aeruginosa* (ATCC 27853) cells were mixed with NPN and PI and then incubated with a range of concentrations of the indicated cathelicidin derivatives for 240 minutes, with monitoring of fluorescence at 15-minute intervals. The full 240-minute incubation curves at all concentrations are shown in **Figures S147-S150**. The data in **A** (OM permeabilization) and **B** (IM permeabilization) show a single-point index of overall permeabilization activity for each mutant that represents the product of the maximum permeabilization of each mutant scaled to LL-37 set to 100% and the proportion of the incubation required to achieve at least 80% of the maximum signal observed for each individual peptide. This index thus incorporates components measuring both peak permeabilization activity and kinetics. Increased helical content in Aib-containing derivatives Q7B and K10B results in increased IM and OM permeabilization. Mutants F2B+V6B and F2B+V6B with I9B demonstrate overall permeabilization activity comparable to the starting template FF-14, thus reinforcing the improved cell type selectivity observed in our antimicrobial susceptibility and hemolysis assays.

### Membrane permeabilization does not predict short cathelicidin killing of gram-negative bacteria

The metric used in Figure 6 is effectively an estimate of the maximum permeabilization potential of a given mutant, typically deriving from high peptide concentration conditions. To better assess the associations between membrane permeabilization and antimicrobial activity at lower peptide concentrations, we determined the lowest concentration of each peptide capable of achieving at least 10% of the permeabilization level observed with control LL-37 set to 100%. This threshold was chosen based on the fact that LL-37 permeabilization of both the OM and IM is detectable at a concentration less than or equal to the LL-37 MIC, thus providing a means of internally calibrating permeabilization against observed killing activity in distinct assays.

**Table 2** summarizes the OM and IM 10% permeabilization thresholds as well as representative MIC measures for each peptide against *E. coli* and *P. aeruginosa*. In both species, the ratio of the MIC to the OM permeabilization threshold was generally consistent. Despite the association between Aib substitution and increased membrane permeabilization, however, efficient permeabilization of *P. aeruginosa* did not result in cell killing commensurate with membrane permeabilization activity. That is, efficient permeabilization was observed even for peptides with MICs of ≥ 50 µM (*e.g.*, K10A, K10B, F2B+V6B, F2B+V6B with I9B). Consistent with the lack of association between permeabilization and killing of *P. aeruginosa*, equivalent permeabilization activity was also observed among mutants with lower MICs (*e.g.*, Q7A and Q7B). Conversely, we noted that despite similar or identical MICs against *E. coli*, permeabilization was detectable for most FF-14 derivatives at concentrations 4-8-fold higher than were observed with LL-37. Together, our results suggest that membrane permeabilization alone does not predict FF-14 killing of gram-negative bacteria.

**Table 2.**
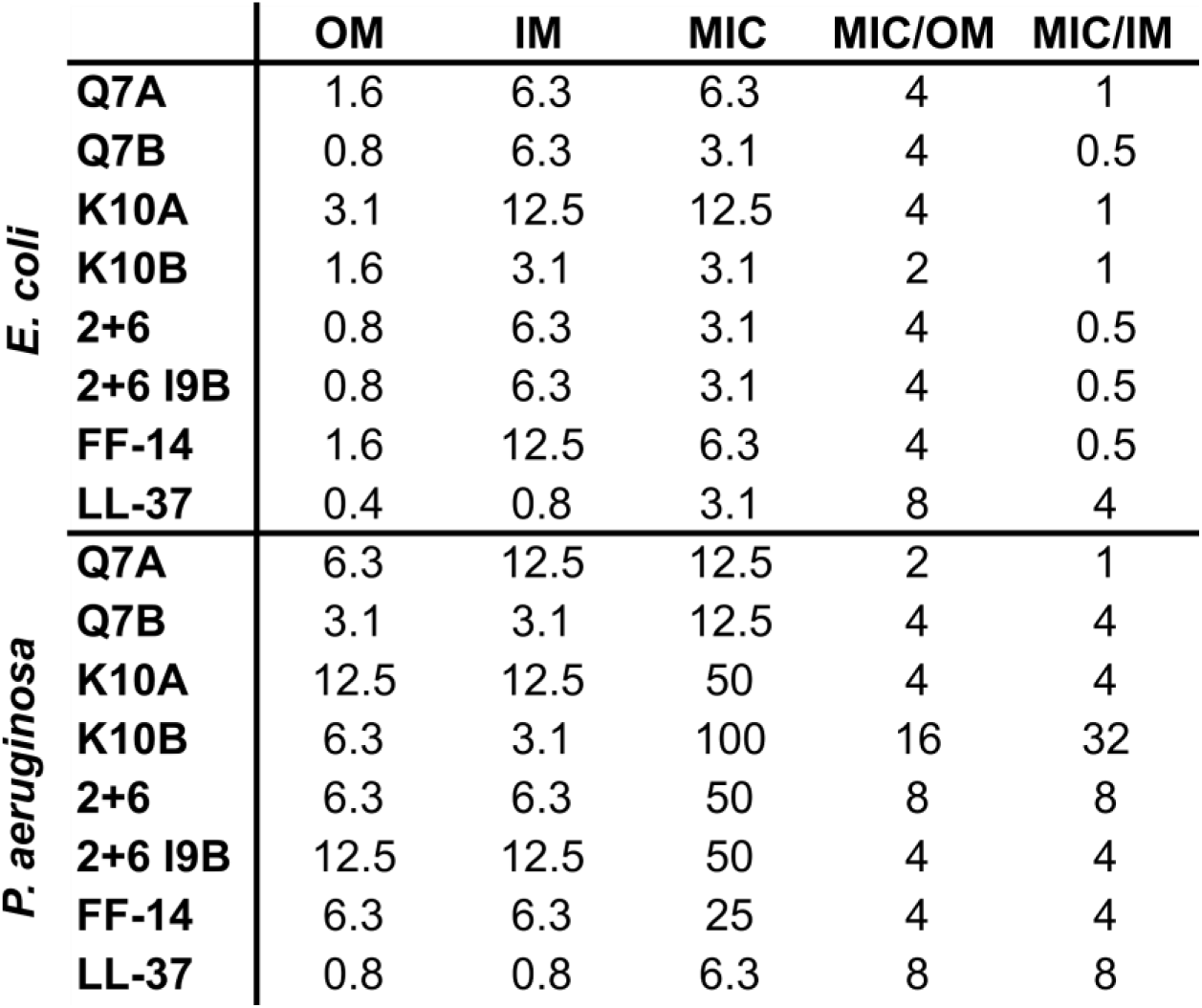
Efficient permeabilization by short cathelicidin derivatives does not correlate with efficient killing in *P. aeruginosa*. NPN and PI were mixed with each of *E. coli* (ATCC 25922) or *P. aeruginosa* (ATCC 27853) in M9+glucose with divalent cation concentrations adjusted to approximate those in the unadjusted MHB used for antimicrobial susceptibility testing in these studies. Mixtures were then incubated with a range of concentrations of each peptide over 240 minutes, with monitoring of fluorescence at 15-minute intervals. This table shows the lowest concentration at which each peptide yields at least 10% of the permeabilization activity of LL-37 set to 100%, which was chosen as a conservative estimate of the lower limits of assay detection. These concentrations were then compared with the MICs for each peptide. While the ratio of the MIC to OM permeabilization concentrations was generally consistent both between species and in comparison with wild-type LL-37, the ratio of the MIC to IM permeabilization was higher in *P. aeruginosa* in comparison with *E. coli*. Efficient IM permeabilization was further noted to occur even for peptides with MICs of ≥ 50 µM (*e.g.*, K10A, K10B, F2B+V6B, F2B+V6B with I9B), and at levels equivalent to those observed with peptides demonstrating lower MICs (*e.g.*, Q7A and Q7B). Moreover, within the *E. coli* conditions tested, we observed a distinction of 4-8 fold between the levels of IM permeabilization achieved by each peptide and those achieved by wild-type LL-37 despite comparable MICs. Gram-negative killing by short cathelicidin derivatives is thus not predicted by permeabilization activity alone.

## Discussion

Here we introduce a paired Ala-Aib mutagenesis approach to probing the fundamental biology of a model helical amphipathic scaffold. Our approach reveals two major insights into cathelicidin biology: i) The effects of helicity on activity against different cell types are species dependent, a point that we then leverage to derive mutants with improved cell type selectivity; and ii) Membrane permeabilization does not predict cathelicidin killing of gram-negative bacteria.

The observed correlation between helicity and activity against *E. coli* is consistent with prior literature on full-length LL-37^22^ and serves as internal validation of the paired mutagenesis approach. The association between increased helicity and increased membrane permeabilization is also expected. In *P. aeruginosa*, however, there is no discernable association between increased helicity and increased antimicrobial activity, which demonstrates that the effects of helicity on antimicrobial activity differ by species.

This principle also extends to mammalian cells, which here show effects intermediate between those observed with *E. coli* and *P. aeruginosa*. Leveraging this insight, we develop here a lead sequence that preserves activity against *E. coli* while decreasing activity against mammalian membranes 32-fold relative to LL-37. While substantial optimization lies ahead, this combines our prior work on increasing the potency of short cathelicidin derivatives into the low micromolar range typical of approved antibiotics^34^ with our prior work improving the cell type selectivity of full-length LL-37^20^ to yield a meaningful and rational improvement along the path to viable short cathelicidin therapeutics. Reliance on Aib to support structure where needed may prove a chemically simpler method than alternatives such as chemical stapling^50,51^.

In addition to the dissociation between helicity and activity against distinct cell types, our data demonstrate no apparent relationship between membrane permeabilization and killing of *P. aeruginosa* (**Table 2**). Even for *E. coli*, the detection of IM permeabilization only at concentrations 4-8-fold higher than would be expected based on the behavior of LL-37 in these same systems is suggestive of a non-membranolytic mechanism of action occurring at lower concentrations. Based on the similar effects of all peptides on OM permeabilization relative to MIC in both species combined with divergence in IM permeabilization phenotypes, we hypothesize that LL-37 may have antimicrobial activities localized to the periplasm.

In summary, we demonstrate here that neither helical structure nor membrane permeabilization is a universal determinant of killing in distinct gram-negative species. Our data thus point toward unexpected complexity in cathelicidin biology while providing a generalizable means of finding similar complexity in a broader range of helical peptides.

## Methods

### Peptide synthesis and purification

Peptide synthesis was completed as previously described on third generation automated flow peptide synthesizers^52–54^. Aib residues were coupled as Ala. 4-(4-hydroxymethyl-3-methoxyphenoxy)butyric acid (HMPB) resin was used throughout (ChemMatrix). Following synthesis, peptides were cleaved with Reagent K (82.5% trifluoroacetic acid (TFA), 5% water, 5% phenol, 2.5% ethanedithiol) for 2 hours at room temperature followed by peptide precipitation in ether chilled on dry ice. Following precipitation, crude peptide was resuspended in 50% water / 50% acetonitrile with 0.1% TFA, flash frozen in liquid nitrogen, and lyophilized.

Peptide purification was carried out on reverse phase columns with a Biotage Selekt flash chromatography system, with subsequent confirmation of the peptide identity and purity by LCMS and HPLC. Analytical data associated with the peptides synthesized for this study are shown in the following **Figures S151-161** (associated with Figure 1), **S162-173** (associated with Figure 2), **S174-185** (associated with Figure 3), **S186-192** (associated with Figure 4), and **S193-202** (associated with Figure 5).

### Circular dichroism spectroscopy

Peptides were dissolved in water at 0.5 mg/mL. Trifluoroethanol was then added to 20% v/v, and spectra were collected on an Aviv 420 spectrometer at 37 °C and 20 °C in 0.1 cm pathlength cuvettes. Temperatures were chosen to mimic distinct physiological contexts in which LL-37 or proteolytic fragments thereof may occur – 37 °C for most body sites and 20 °C to mimic ambient-exposed surfaces such as skin. Spectra were analyzed for secondary structural content using BeStSel^41^ for comparison of relative helicity in a uniform experimental approach.

### Antimicrobial susceptibility testing

Antimicrobial susceptibility testing was completed as per Clinical & Laboratory Standards Institute guidelines with modifications previously described for cationic peptides, including the use of polypropylene plates and unadjusted Mueller Hinton Broth^55^. Protocols were scaled down to either 35 or 50 µL in 384-well plates. After plating, wells were monitored for growth by OD600 over 18 hours at 37 °C in a Tecan Spark plate reader with a Large Humidity Cassette to minimize evaporation. Data shown are from three independent experiments unless otherwise specified.

### Hemolysis assays

Hemolysis assays were completed in 50 µL per well in 384-well plates to match the general conditions used for antimicrobial susceptibility testing above. Defibrinated sheep blood (Hardy Diagnostics) was spun at 800 rcf for 10 minutes and washed with PBS to visual clearance. Blood was then resuspended in PBS at 1% v/v, with subsequent plating of 25 µL of this into 25 µL of 2x target concentrations of peptide dissolved in PBS per well for a final concentration of 0.5% v/v. Hemolysis was then monitored kinetically by OD600 in a Tecan Spark plate reader as for antimicrobial susceptibility testing. In addition to this kinetic measure, hemolysis was monitored as an endpoint measurement by the addition of 25 µL of PBS at 18 hours, spinning for 10 minutes at 800 rcf, and then removing 25 µL of supernatant to a new 384-well plate to be read at 414 nm for free hemoglobin. Data shown are from three independent experiments unless otherwise specified.

### Dual membrane permeabilization assays

Dual membrane permeabilization assays based on prior methods^56^ were completed in M9+glucose with divalent cation concentrations adjusted to approximate those found in the unadjusted MHB used for antimicrobial susceptibility testing in these studies. In brief, *E. coli* (ATCC 25922) or *P. aeruginosa* (ATCC 27853) were grown to OD600 ∼0.5, pelleted at 10,000 rcf for 1 minute, and then resuspended in M9+glucose. These cell suspensions were then mixed with NPN to a final concentration of 10 µM and PI to a final concentration of 5 µM, and 50 µL of this mixture was added to 5 µL of a 10x peptide concentration covering the same range of dilutions used for antimicrobial susceptibility testing in 384-well black, transparent bottom plates. Fluorescence was then monitored at Ex 350 / Em 420 nm for NPN and at Ex 535 / Em 617 nm for PI at 15-minute intervals over 240 minutes of incubation at 37 °C in a Tecan Spark plate reader with large humidity cassette. Data analysis was carried out by subtracting the average cellular background fluorescence from all wells and then calculating a fold-change in the fluorescence signal observed under each condition relative to the baseline fluorescence signal in the absence of peptide. The maximum signal observed with LL-37 was then set to 100%, with scaling of remaining peptides as a percentage of this control. In rare instances, a given measure of a strongly permeabilizing peptide such as LL-37 might register under these conditions as over-assay for 1-2 timepoints. Where this occurred, the over-assay timepoint was substituted with the next observed within-assay datapoint. Data shown are the mean and standard deviation of triplicate experiments.

## Supporting information

Combined_Tables_S1-6_Figures_S1-202

## Acknowledgements

This work has been funded by the National Institute of Allergy and Infectious Diseases (T32 AI007061 and K08 AI166345 to JSA; U19 AI142780 to BLP) and by the Cystic Fibrosis Foundation (ALBIN19F0, ALBIN21Q0, ALBIN22A0-KB to JSA).

## Author Contributions

JSA designed and conducted experiments and wrote the paper. BLP guided the experiments and wrote the paper.

