## Supplementary material for "Decoupling membrane permeabilization and antimicrobial activity in amphipathic peptides": Combined_Tables_S1-6_Figures_S1-202

**Table of Contents:**

Tables S1-6 Standardized quantification of data from Figures 1-5 S3-S14

| | |
|---|---|
| Figure S1 CD spectroscopy for peptides in Figure 1 | S15-S16 |
| Figure S2 Histogram depiction of data in Figure 1 | S17-S18 |
| Figures S3-S14 Kinetic microbial growth curves associated with Figure 1 | S19-S42 |
| Figures S15-S26 Kinetic hemolysis curves associated with Figure 1 | S43-S66 |
| Figure S27 Histogram depiction of data in Figure 2 | S67-S68 |
| Figures S28-S42 Kinetic microbial growth curves associated with Figure 2 | S69-S98 |
| Figures S43-S57 Kinetic hemolysis curves associated with Figure 2 | S99-S128 |
| Figure S58 Correlation between Figure 2 helical content and MIC | S129-S130 |
| Figure S59 CD spectroscopy for peptides in Figure 2 | S131-S132 |
| Figure S60 Histogram depiction of data in Figure 3 | S133-S134 |
| Figures S61-S76 Kinetic microbial growth curves associated with Figure 3 | S135-S166 |
| Figures S77-S92 Kinetic hemolysis curves associated with Figure 3 | S167-S198 |
| Figure S93 CD spectroscopy for peptides in Figure 3 | S199-S200 |
| Figure S94 CD spectroscopy of Aib double mutants in Figure 4 | S201-S202 |
| Figure S95 Histogram depiction of data in Figure 4 | S203-S204 |
| Figures S96-S105 Kinetic microbial growth curves associated with Figure 4 | S205-S224 |
| Figures S106-S115 Kinetic hemolysis curves associated with Figure 4 | S225-S244 |
| Figure S116 Histogram depiction of data in Figure 5 | S245-S246 |
| Figures S117-S131 Kinetic microbial growth curves associated with Figure 5 | S247-S276 |
| Figures S132-S146 Kinetic hemolysis curves associated with Figure 5 | S277-S306 |
| Figures S147-S150 Kinetic permeabilization curves for Figure 6 and Table 2 | S307-S314 |
| Figures S151-S161 Analytical data for peptides in Figure 1 | S315-S336 |
| Figures S162-S173 Analytical data for peptides in Figure 2 | S337-S360 |
| Figures S174-S185 Analytical data for peptides in Figure 3 | S361-S384 |
| Figures S186-S192 Analytical data for peptides in Figure 4 | S385-S398 |
| Figures S193-S202 Analytical data for peptides in Figure 5 | S399-S418 |

| | MIC Ec | MIC Pa | < 20% Hemo | Hemo/MIC | Fold Δ |
|---|---|---|---|---|---|
| **FF-14 m4** | 100 | >50 | 100 | 1 | 4 |
| **FF-14 m4-FF** | 100 | >50 | 200 | 2 | 8 |
| **FF-14 m5** | 100 | >50 | 200 | 2 | 8 |
| **FF-14 m5-FF** | 12.5 | 50 | 12.5 | 1 | 4 |
| **FF-14 m6** | 100 | >50 | 200 | 2 | 8 |
| **FF-14 m6-FF** | 3.1 | 12.5 | 3.1 | 1 | 4 |
| **FF-14** | 3.1 | 12.5 | 3.1 | 1 | 4 |
| **LL-37 m4** | 3.1 | 3.1 | 200 | 64 | 256 |
| **LL-37 m5** | 1.6 | 6.3 | 0.8 | 0.5 | 2 |
| **LL-37 m6** | 0.8 | 1.6 | 0.8 | 1 | 4 |
| **LL-37** | 3.1 | 6.3 | 0.8 | 0.25 | 1 |
| **Colistin** | 0.8 | 0.8 | 100 | 128 | 512 |

**Table S1 – Standardized, quantitative presentation of data from Figure 1. Tables S1-5** include a standardized, quantitative presentation of data from **Figures 1-5** to aid in interpretation. In each table, the **MIC** column indicates the MIC of a given peptide for *E. coli* (Ec) or *P. aeruginosa* (Pa). The **< 20% Hemo** column indicates the highest peptide concentration at which hemolysis is less than 20%, a threshold chosen based on the level at which colistin can typically be seen as shifting between hemolysis and no/low hemolysis in histogram depictions of the data. The **Hemo/MIC** column divides the hemolysis value by the MIC Ec value to give a sense of the ability of a given peptide to separate hemolytic versus antimicrobial activities; larger numbers are better. The **Fold Δ** column divides the value in the **Hemo/MIC** column by the value obtained for LL-37 in a given set of experiments to quantify overall improvement over the LL-37 template. Colistin is used throughout as a control reference antibiotic that can be used in humans.

| | MIC Ec | MIC Pa | < 20% Hemo | Hemo/MIC | Fold Δ |
|---|---|---|---|---|---|
| **FF-14 F1A** | 12.5 | 50 | 25 | 2 | 8 |
| **FF-14 F2A** | 25 | 50 | 25 | 1 | 4 |
| **FF-14 I5A** | 25 | 25 | 25 | 1 | 4 |
| **FF-14 V6A** | 12.5 | 25 | 25 | 2 | 8 |
| **FF-14 Q7A** | 6.3 | 12.5 | 6.3 | 1 | 4 |
| **FF-14 R8A** | 3.1 | 25 | 6.3 | 2 | 8 |
| **FF-14 I9A** | 100 | 50 | 200 | 2 | 8 |
| **FF-14 K10A** | 12.5 | 50 | 6.3 | 0.5 | 2 |
| **FF-14 D11A** | 3.1 | 6.3 | 3.1 | 1 | 4 |
| **FF-14 F12A** | 12.5 | 25 | 50 | 4 | 16 |
| **FF-14 L13A** | 50 | 25 | 200 | 4 | 16 |
| **FF-14 R14A** | 6.3 | 50 | 12.5 | 2 | 8 |
| **FF-14** | 6.3 | 25 | 6.3 | 1 | 4 |
| **LL-37** | 6.3 | 6.3 | 1.6 | 0.25 | 1 |
| **Colistin** | 0.8 | 0.8 | 50 | 64 | 256 |

**Table S2 – Standardized, quantitative presentation of data from Figure 2. Tables S1-5** include a standardized, quantitative presentation of selected data from **Figures 1-5** to aid in interpretation. In each table, the **MIC** column indicates the MIC of a given peptide for *E. coli* (Ec) or *P. aeruginosa* (Pa). The **< 20% Hemo** column indicates the highest peptide concentration at which hemolysis is less than 20%, a threshold chosen based on the level at which colistin can typically be seen as shifting between hemolysis and no/low hemolysis in histogram depictions of the data. The **Hemo/MIC** column divides the hemolysis value by the MIC Ec value to give a sense of the ability of a given peptide to separate hemolytic versus antimicrobial activities; larger numbers are better. The **Fold Δ** column divides the value in the **Hemo/MIC** column by the value obtained for LL-37 in a given set of experiments to quantify overall improvement over the LL-37 template. Colistin is used throughout as a control reference antibiotic that can be used in humans.

| | MIC Ec | MIC Pa | < 20% Hemo | Hemo/MIC | Fold Δ |
|---|---|---|---|---|---|
| **FF-14 F1B** | 12.5 | 100 | 25 | 2 | 4 |
| **FF-14 F2B** | 3.1 | 25 | 12.5 | 4 | 8 |
| **FF-14 I5B** | 3.1 | 25 | 12.5 | 4 | 8 |
| **FF-14 V6B** | 3.1 | 25 | 6.3 | 2 | 4 |
| **FF-14 Q7B** | 3.1 | 12.5 | 6.3 | 2 | 4 |
| **FF-14 R8B** | 3.1 | 50 | 3.1 | 1 | 2 |
| **FF-14 I9B** | 12.5 | 25 | 50 | 4 | 8 |
| **FF-14 K10B** | 3.1 | 100 | 1.6 | 0.5 | 1 |
| **FF-14 D11B** | 3.1 | 6.3 | 6.3 | 2 | 4 |
| **FF-14 F12B** | 12.5 | 25 | 25 | 2 | 4 |
| **FF-14 L13B** | 50 | 25 | 100 | 2 | 4 |
| **FF-14 R14B** | 6.3 | 50 | 12.5 | 2 | 4 |
| **FF-14** | 6.3 | 12.5 | 6.3 | 1 | 2 |
| **LL-37** | 3.1 | 6.3 | 1.6 | 0.5 | 1 |
| **Colistin** | 0.4 | 0.4 | 25 | 64 | 128 |

**Table S3 – Standardized, quantitative presentation of data from Figure 3. Tables S1-5** include a standardized, quantitative presentation of selected data from **Figures 1-5** to aid in interpretation. In each table, the **MIC** column indicates the MIC of a given peptide for *E. coli* (Ec) or *P. aeruginosa* (Pa). The **< 20% Hemo** column indicates the highest peptide concentration at which hemolysis is less than 20%, a threshold chosen based on the level at which colistin can typically be seen as shifting between hemolysis and no/low hemolysis in histogram depictions of the data. The **Hemo/MIC** column divides the hemolysis value by the MIC Ec value to give a sense of the ability of a given peptide to separate hemolytic versus antimicrobial activities; larger numbers are better. The **Fold Δ** column divides the value in the **Hemo/MIC** column by the value obtained for LL-37 in a given set of experiments to quantify overall improvement over the LL-37 template. Colistin is used throughout as a control reference antibiotic that can be used in humans.

| | MIC Ec | MIC Pa | < 20% Hemo | Hemo/MIC | Fold Δ |
|---|---|---|---|---|---|
| **B2+B6** | 3.1 | 50 | 12.5 | 4 | 8 |
| **B5+B9** | 6.3 | 50 | 50 | 8 | 16 |
| **B9+B13** | 100 | >50 | 200 | 2 | 4 |
| **B1+B5** | 12.5 | >50 | 100 | 8 | 16 |
| **2+6 I9B** | 3.1 | 50 | 50 | 16 | 32 |
| **2+6 L13B** | 12.5 | 50 | 100 | 8 | 16 |
| **2+6 I9B+L13B** | 25 | >50 | 200 | 8 | 16 |
| **FF-14** | 6.3 | 25 | 6.3 | 1 | 2 |
| **LL-37** | 3.1 | 6.3 | 1.6 | 0.5 | 1 |
| **Colistin** | 0.8 | 0.8 | 200 | 256 | 512 |

**Table S4 – Standardized, quantitative presentation of data from Figure 4. Tables S1-5** include a standardized, quantitative presentation of selected data from **Figures 1-5** to aid in interpretation. In each table, the **MIC** column indicates the MIC of a given peptide for *E. coli* (Ec) or *P. aeruginosa* (Pa). The **< 20% Hemo** column indicates the highest peptide concentration at which hemolysis is less than 20%, a threshold chosen based on the level at which colistin can typically be seen as shifting between hemolysis and no/low hemolysis in histogram depictions of the data. The **Hemo/MIC** column divides the hemolysis value by the MIC Ec value to give a sense of the ability of a given peptide to separate hemolytic versus antimicrobial activities; larger numbers are better. The **Fold Δ** column divides the value in the **Hemo/MIC** column by the value obtained for LL-37 in a given set of experiments to quantify overall improvement over the LL-37 template. Colistin is used throughout as a control reference antibiotic that can be used in humans.

| | MIC Ec | MIC Pa | < 20% Hemo | Hemo/MIC | Fold Δ |
|---|---|---|---|---|---|
| **FF-14 2+6 I9A** | 25 | >50 | 200 | 8 | 16 |
| **FF-14 2+6 L13A** | 50 | >50 | 200 | 4 | 8 |
| **FF-14 2+6 I9A+L13A** | 100 | >50 | 200 | 2 | 4 |
| **FF-14 2+6 K3A** | 12.5 | >50 | 50 | 4 | 8 |
| **FF-14 2+6 R4A** | 25 | >50 | 100 | 4 | 8 |
| **FF-14 2+6 Q7A** | 12.5 | >50 | 50 | 4 | 8 |
| **FF-14 2+6 R8A** | 6.3 | >50 | 12.5 | 2 | 4 |
| **FF-14 2+6 K10A** | 6.3 | >50 | 25 | 4 | 8 |
| **FF-14 2+6 D11A** | 6.3 | 12.5 | 25 | 4 | 8 |
| **FF-14 2+6 R14A** | 6.3 | >50 | 50 | 8 | 16 |
| **FF-14** | 6.3 | 25 | 12.5 | 2 | 4 |
| **2+6** | 3.1 | 50 | 25 | 8 | 16 |
| **m4** | 6.3 | 25 | 200 | 32 | 64 |
| **LL-37** | 3.1 | 6.3 | 1.6 | 0.5 | 1 |
| **Colistin** | 3.1 | 1.6 | 200 | 64 | 128 |

**Table S5 – Standardized, quantitative presentation of data from Figure 5. Tables S1-5** include a standardized, quantitative presentation of selected data from **Figures 1-5** to aid in interpretation. In each table, the **MIC** column indicates the MIC of a given peptide for *E. coli* (Ec) or *P. aeruginosa* (Pa). The **< 20% Hemo** column indicates the highest peptide concentration at which hemolysis is less than 20%, a threshold chosen based on the level at which colistin can typically be seen as shifting between hemolysis and no/low hemolysis in histogram depictions of the data. The **Hemo/MIC** column divides the hemolysis value by the MIC Ec value to give a sense of the ability of a given peptide to separate hemolytic versus antimicrobial activities; larger numbers are better. The **Fold Δ** column divides the value in the **Hemo/MIC** column by the value obtained for LL-37 in a given set of experiments to quantify overall improvement over the LL-37 template. Colistin is used throughout as a control reference antibiotic that can be used in humans.

| | 2+6 | | | Ala | | | | |
|---|---|---|---|---|---|---|---|---|
| | **MIC** | **< 20% Hemc** | **Hemo/MIC** | **MIC** | **< 20% Hemo** | **Hemo/MIC** | **2+6 Fold Δ** | **FF-14 Fold Δ** |
| **I9A** | 25 | 200 | 8 | 100 | 200 | 2 | 1 | 2 |
| **L13A** | 50 | 200 | 4 | 50 | 200 | 4 | 0.5 | 4 |
| **Q7A** | 12.5 | 50 | 4 | 6.3 | 6.3 | 1 | 0.5 | 1 |
| **R8A** | 6.3 | 12.5 | 2 | 3.1 | 6.3 | 2 | 0.25 | 2 |
| **K10A** | 6.3 | 25 | 4 | 12.5 | 6.3 | 0.5 | 0.5 | 0.5 |
| **D11A** | 6.3 | 25 | 4 | 3.1 | 3.1 | 1 | 0.5 | 1 |
| **R14A** | 6.3 | 50 | 8 | 6.3 | 12.5 | 2 | 1 | 2 |
| **2+6** | 3.1 | 25 | 8 | | | | 1 | 0 |
| **FF-14** | | | | 6.3 | 6.3 | 1 | | |

**Table S6 – Comparison of phenotypic effects of shared Ala mutations between the FF-14 and FF-14 F2B+V6B templates. Table S6** recapitulates the data from the standardized formatting in **Tables S2** and **5**, except that it sets the reference peptide to either F2B+V6B for mutations in this context or to FF-14 for mutations in this context. Mutation **Fold Δ** values are typically ≤ 1 in the F2B+V6B template, indicating neutral-to-worsening *in vitro* therapeutic indices. Mutation **Fold Δ** values in the FF-14 template, by contrast, are generally ≥1, indicating neutral-to-improving therapeutic indices.

A.

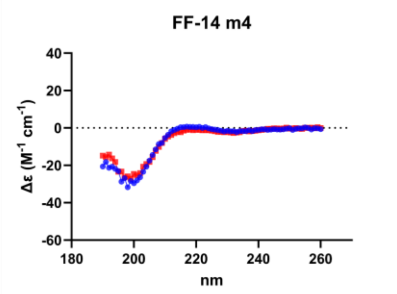


B.

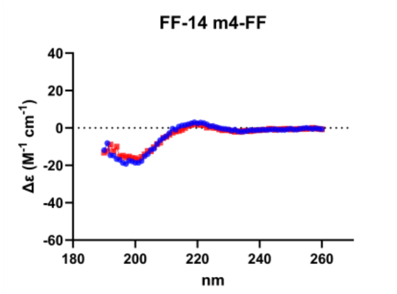


C.

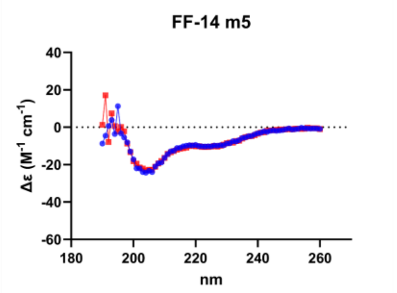


D.

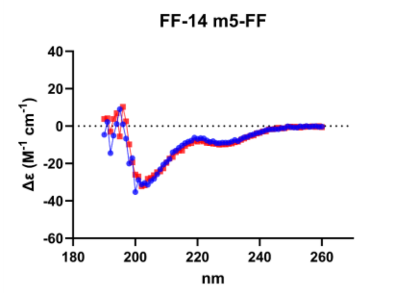


E.

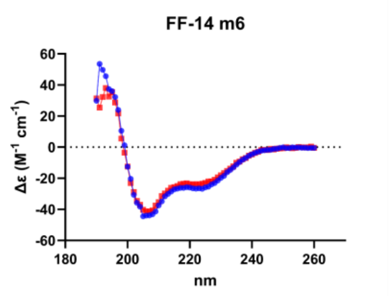


F.

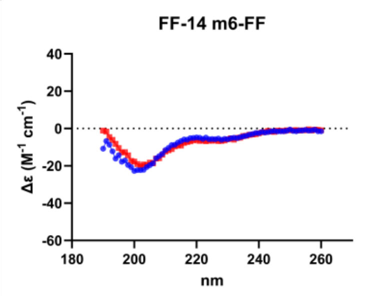


G.

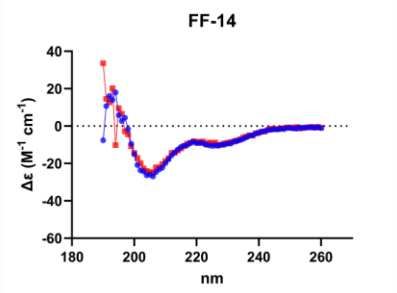


H.

| | % Helix | |
|---|---|---|
| | 20 °C | 37 °C |
| FF-14 m4 | 3 | 10.6 |
| FF-14 m4-FF | 0 | 0 |
| FF-14 m5 | 80.8 | 76.8 |
| FF-14 m5-FF | 79 | 92 |
| FF-14 m6 | 100 | 100 |
| FF-14 m6-FF | 42.6 | 49.4 |
| FF-14 | 100 | 85.8 |

**Figure S1 – Circular dichroism spectroscopy of Figure 1 mutants. A-G.** Circular dichroism spectra are displayed for each mutant collected from 190-260 nm at both 20 °C (Blue) and 37 °C (Red). **H.** Quantification of helical content as determined by BeStSel for each mutant.

A.

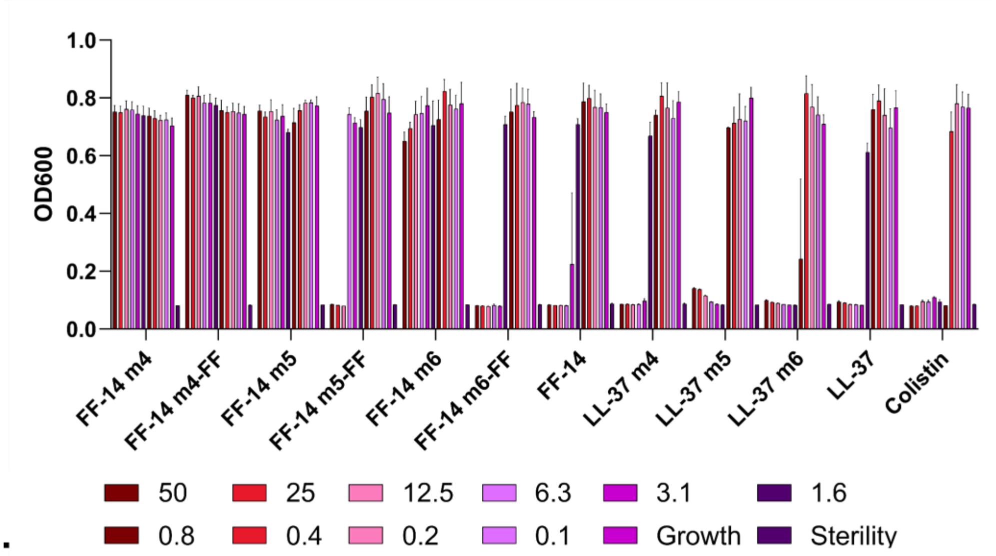
OD600
1.0
0.8
0.6
0.4
0.2
0.0
FF-14 m4
FF-14 m4-FF
FF-14 m5
FF-14 m5-FF
FF-14 m6
FF-14 m6-FF
FF-14
LL-37 m4
LL-37 m5
LL-37 m6
LL-37
Colistin
50
25
12.5
6.3
3.1
1.6
0.8
0.4
0.2
0.1
Growth
Sterility

B.

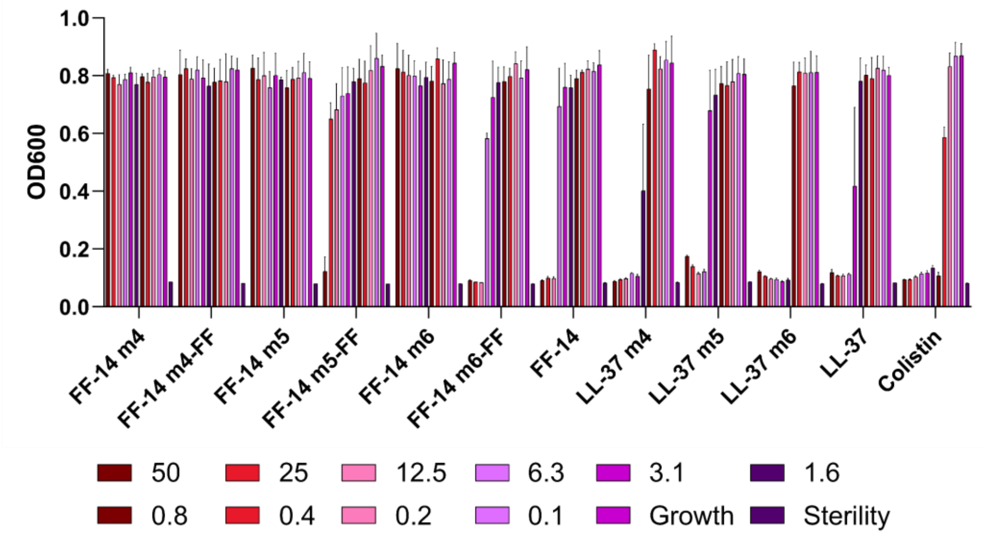
OD600
1.0
0.8
0.6
0.4
0.2
0.0
FF-14 m4
FF-14 m4-FF
FF-14 m5
FF-14 m5-FF
FF-14 m6
FF-14 m6-FF
FF-14
LL-37 m4
LL-37 m5
LL-37 m6
LL-37
Colistin
50
25
12.5
6.3
3.1
1.6
0.8
0.4
0.2
0.1
Growth
Sterility

C.

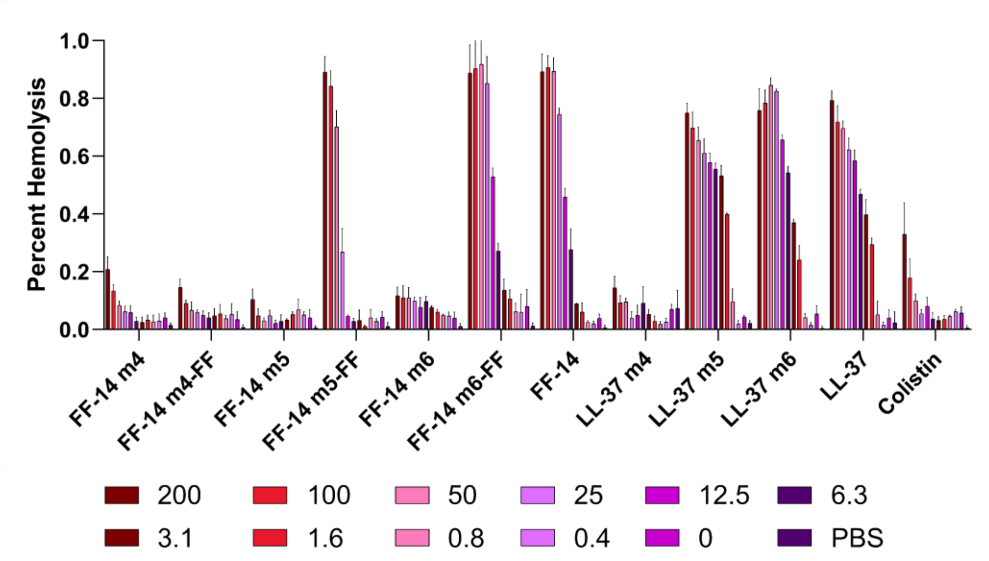
Percent Hemolysis
1.0
0.8
0.6
0.4
0.2
0.0
FF-14 m4
FF-14 m4-FF
FF-14 m5
FF-14 m5-FF
FF-14 m6
FF-14 m6-FF
FF-14
LL-37 m4
LL-37 m5
LL-37 m6
LL-37
Colistin
200
100
50
25
12.5
6.3
3.1
1.6
0.8
0.4
0
PBS

**Figure S2 – Histogram depiction of data from Figure 1. A.** Endpoint activity of each peptide against *E. coli*; measures are raw absorbance at 600 nm after 18 hours at 37 °C. **B.** Endpoint activity of each peptide against *P. aeruginosa*. **C.** Hemolytic activity of each peptide; measures are absorbance at 414 nm scaled to a thrice freeze-thawed positive control set at 100% hemolysis. Concentrations throughout are shown are in μM.

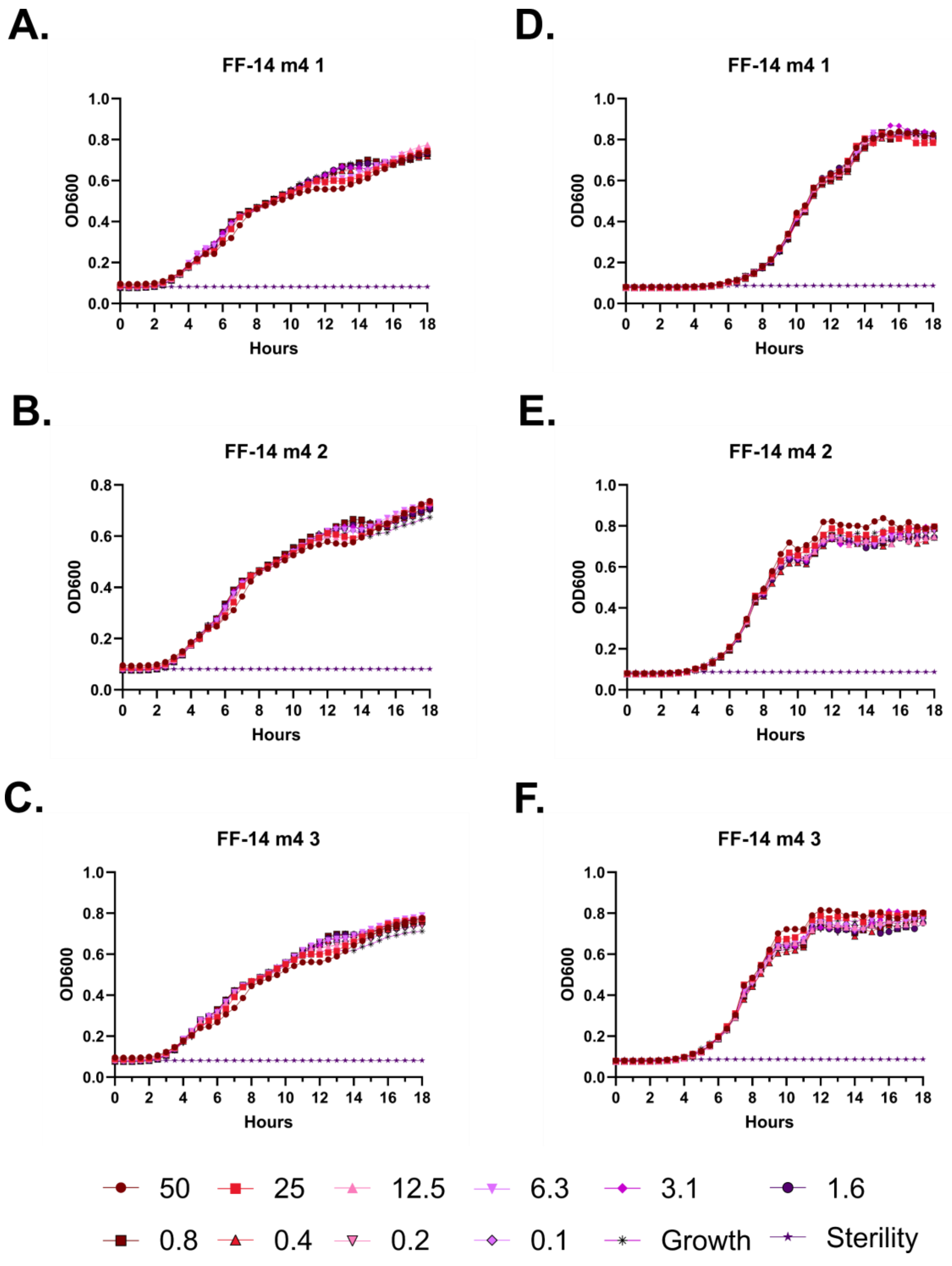
A.
FF-14 m4 1
OD600
Hours
D.
FF-14 m4 1
OD600
Hours
B.
FF-14 m4 2
OD600
Hours
E.
FF-14 m4 2
OD600
Hours
C.
FF-14 m4 3
OD600
Hours
F.
FF-14 m4 3
OD600
Hours
50
25
12.5
6.3
3.1
1.6
0.8
0.4
0.2
0.1
Growth
Sterility

**Figure S3 – Antimicrobial susceptibility testing with FF-14 m4. A-C.** Growth of *E. coli* in the presence of the indicated peptide concentrations given in μM as monitored by OD600 at 37 °C over 18 hours. Each panel indicates an independent experiment. **D-F.** As in **A-C**, but with *P. aeruginosa.*

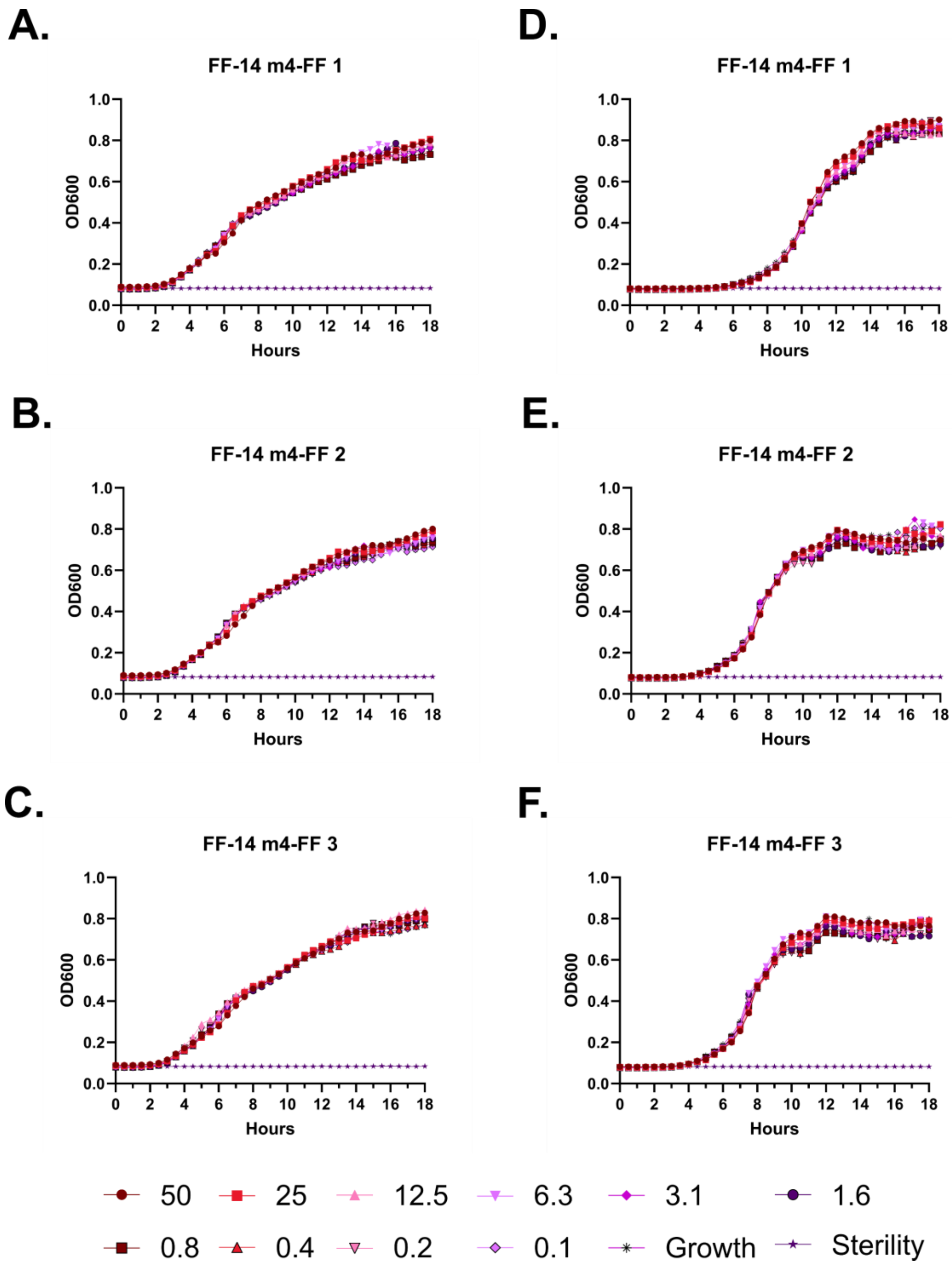
A.
FF-14 m4-FF 1
B.
FF-14 m4-FF 2
C.
FF-14 m4-FF 3
D.
FF-14 m4-FF 1
E.
FF-14 m4-FF 2
F.
FF-14 m4-FF 3
OD600
Hours
50
25
12.5
6.3
3.1
1.6
0.8
0.4
0.2
0.1
Growth
Sterility

**Figure S4 – Antimicrobial susceptibility testing with FF-14 m4-FF. A-C.** Growth of *E. coli* in the presence of the indicated peptide concentrations given in μM as monitored by OD600 at 37 °C over 18 hours. Each panel indicates an independent experiment. **D-F.** As in **A-C**, but with *P. aeruginosa.*

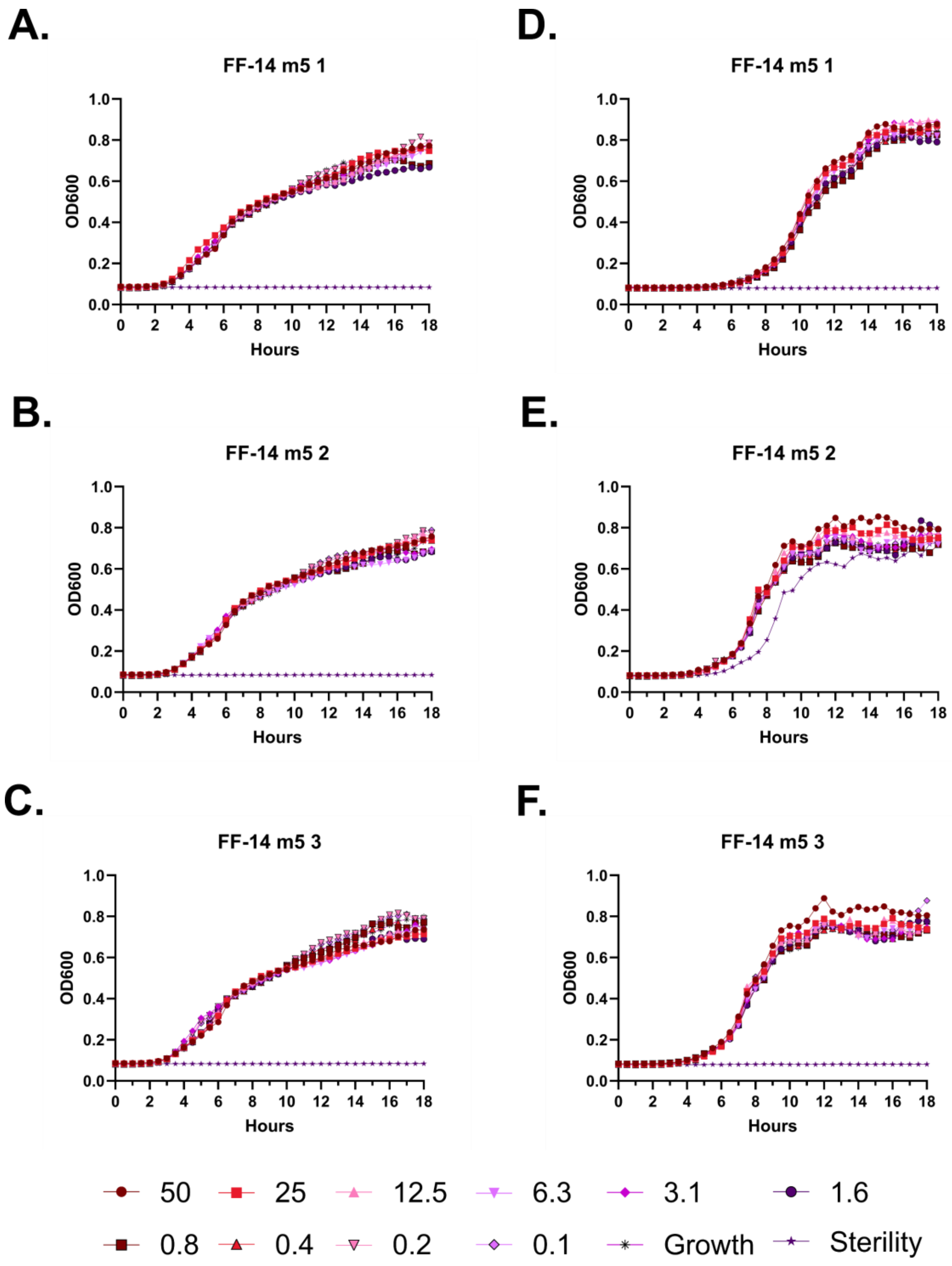
A.
FF-14 m5 1
D.
FF-14 m5 1
B.
FF-14 m5 2
E.
FF-14 m5 2
C.
FF-14 m5 3
F.
FF-14 m5 3
OD600
Hours
0.0
0.2
0.4
0.6
0.8
1.0
0
2
4
6
8
10
12
14
16
18
50
25
12.5
6.3
3.1
1.6
0.8
0.4
0.2
0.1
Growth
Sterility

**Figure S5 – Antimicrobial susceptibility testing with FF-14 m5. A-C.** Growth of *E. coli* in the presence of the indicated peptide concentrations given in μM as monitored by OD600 at 37 °C over 18 hours. Each panel indicates an independent experiment. **D-F.** As in **A-C**, but with *P. aeruginosa.*

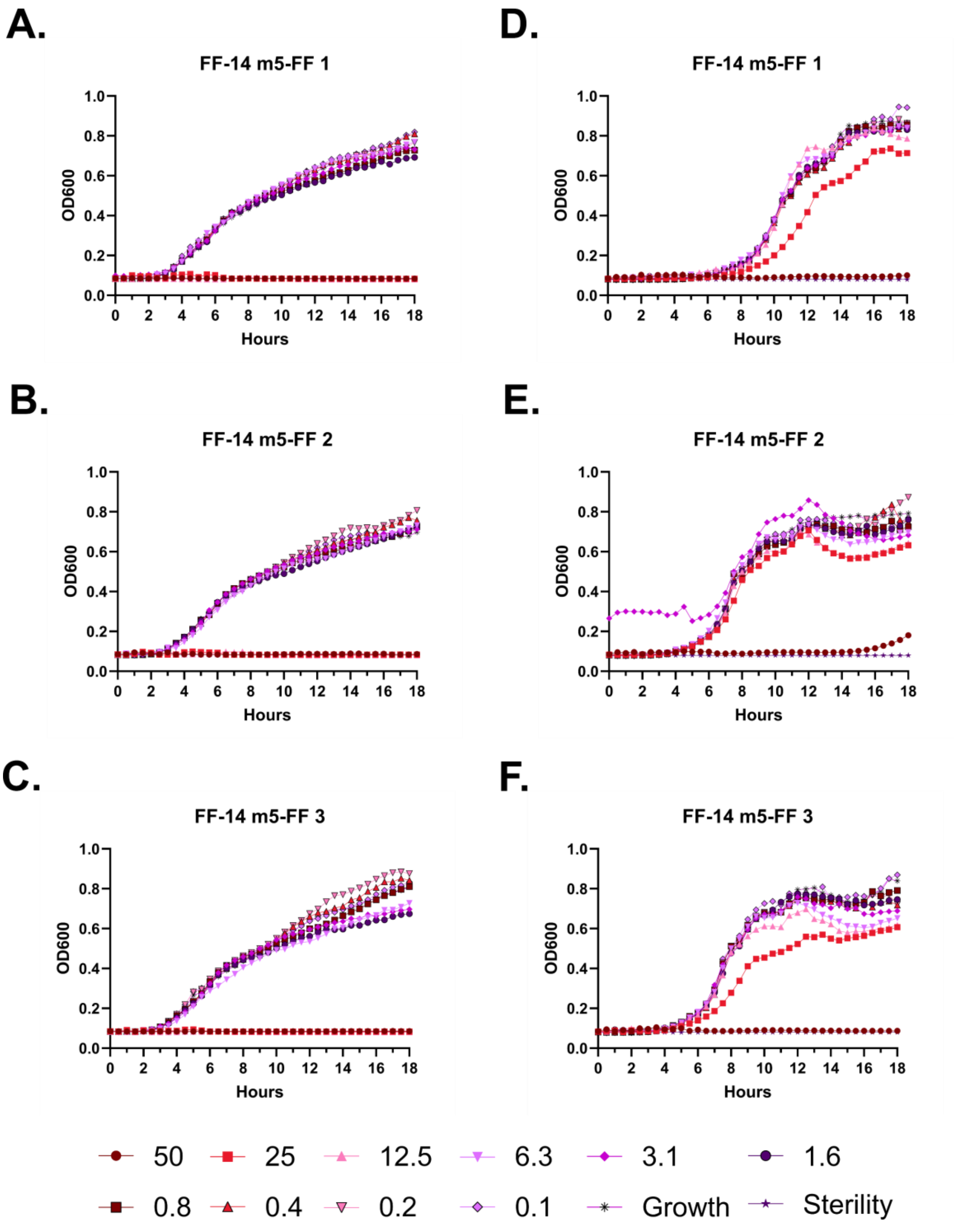
A.
FF-14 m5-FF 1
B.
FF-14 m5-FF 2
C.
FF-14 m5-FF 3
D.
FF-14 m5-FF 1
E.
FF-14 m5-FF 2
F.
FF-14 m5-FF 3
OD600
Hours
50
25
12.5
6.3
3.1
1.6
0.8
0.4
0.2
0.1
Growth
Sterility

**Figure S6 – Antimicrobial susceptibility testing with FF-14 m5-FF. A-C.** Growth of *E. coli* in the presence of the indicated peptide concentrations given in μM as monitored by OD600 at 37 °C over 18 hours. Each panel indicates an independent experiment. **D-F.** As in **A-C**, but with *P. aeruginosa.*

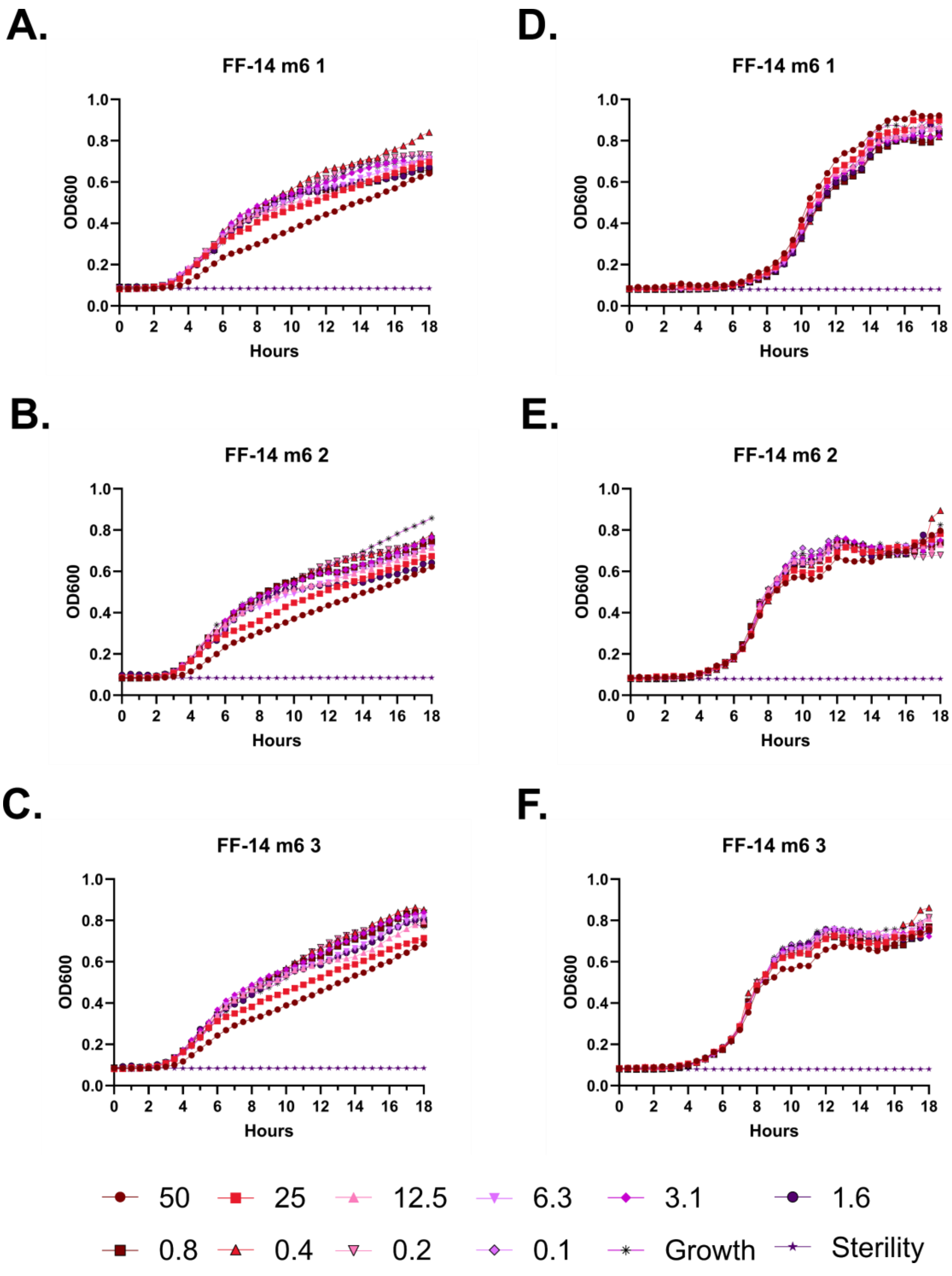

A.
FF-14 m6 1
B.
FF-14 m6 2
C.
FF-14 m6 3
D.
FF-14 m6 1
E.
FF-14 m6 2
F.
FF-14 m6 3
OD600
Hours
0.0
0.2
0.4
0.6
0.8
1.0
0 2 4 6 8 10 12 14 16 18
50
25
12.5
6.3
3.1
1.6
0.8
0.4
0.2
0.1
Growth
Sterility

**Figure S7 – Antimicrobial susceptibility testing with FF-14 m6. A-C.** Growth of *E. coli* in the presence of the indicated peptide concentrations given in μM as monitored by OD600 at 37 °C over 18 hours. Each panel indicates an independent experiment. **D-F.** As in **A-C**, but with *P. aeruginosa.*

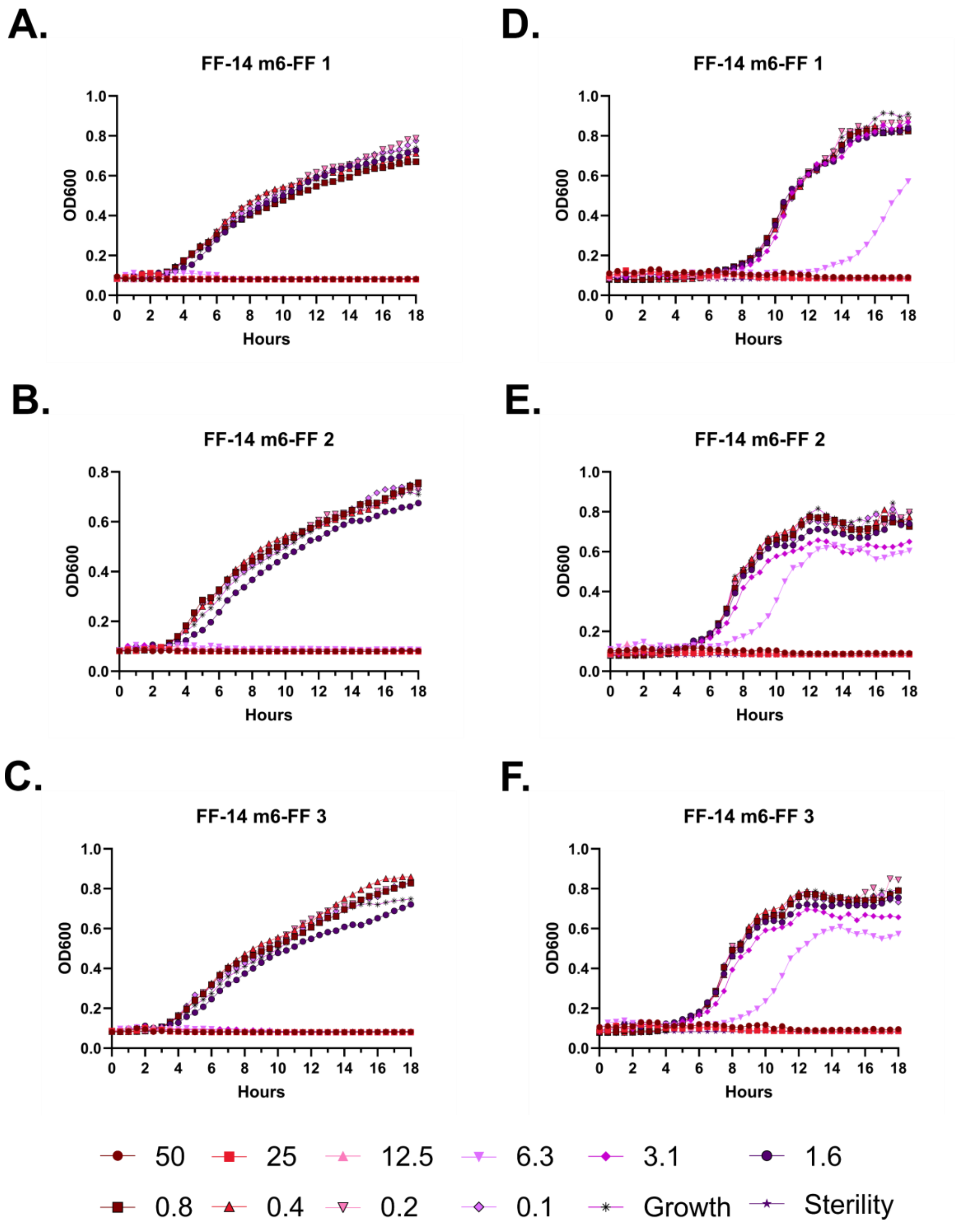
A.
FF-14 m6-FF 1
B.
FF-14 m6-FF 2
C.
FF-14 m6-FF 3
D.
FF-14 m6-FF 1
E.
FF-14 m6-FF 2
F.
FF-14 m6-FF 3
OD600
Hours
50
25
12.5
6.3
3.1
1.6
0.8
0.4
0.2
0.1
Growth
Sterility

**Figure S8 – Antimicrobial susceptibility testing with FF-14 m6-FF. A-C.** Growth of *E. coli* in the presence of the indicated peptide concentrations given in μM as monitored by OD600 at 37 °C over 18 hours. Each panel indicates an independent experiment. **D-F.** As in **A-C**, but with *P. aeruginosa.*

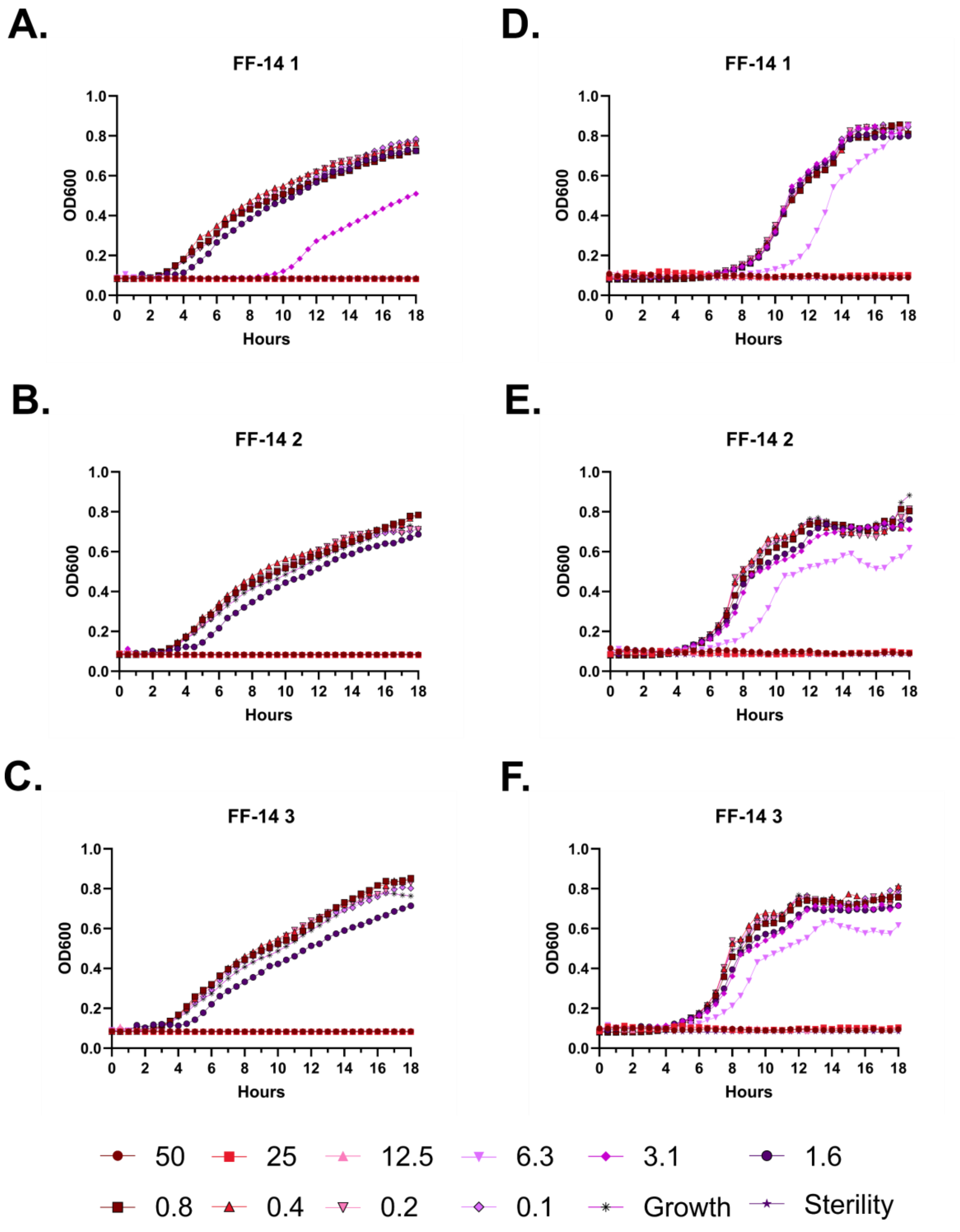
A.
FF-14 1
D.
FF-14 1
B.
FF-14 2
E.
FF-14 2
C.
FF-14 3
F.
FF-14 3
OD600
Hours
50
25
12.5
6.3
3.1
1.6
0.8
0.4
0.2
0.1
Growth
Sterility

**Figure S9 – Antimicrobial susceptibility testing with FF-14. A-C.** Growth of *E. coli* in the presence of the indicated peptide concentrations given in μM as monitored by OD600 at 37 °C over 18 hours. Each panel indicates an independent experiment. **D-F.** As in **A-C**, but with *P. aeruginosa.*

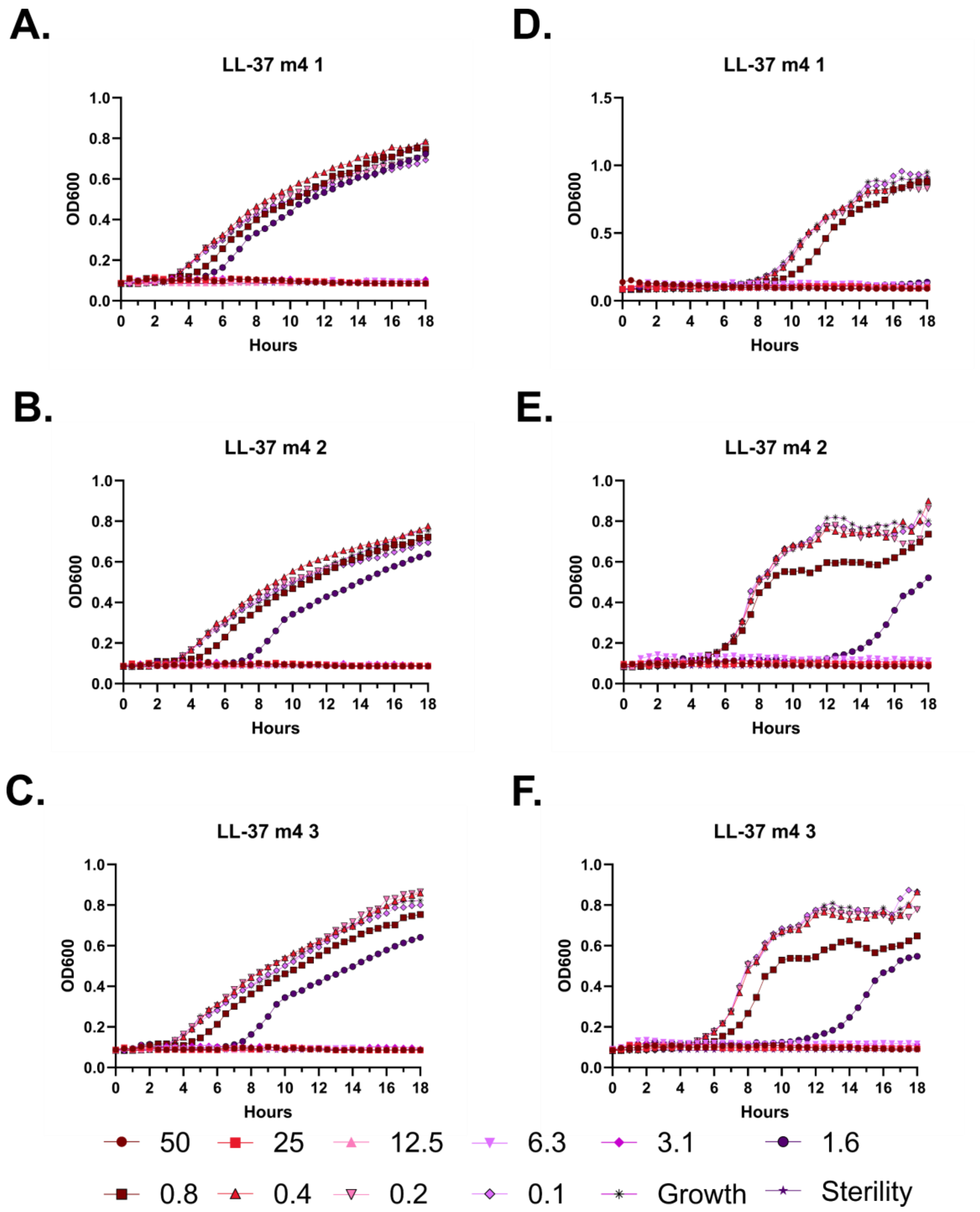
A.
LL-37 m4 1
OD600
Hours
D.
LL-37 m4 1
OD600
Hours
B.
LL-37 m4 2
OD600
Hours
E.
LL-37 m4 2
OD600
Hours
C.
LL-37 m4 3
OD600
Hours
F.
LL-37 m4 3
OD600
Hours
50
25
12.5
6.3
3.1
1.6
0.8
0.4
0.2
0.1
Growth
Sterility

**Figure S10 – Antimicrobial susceptibility testing with LL-37 m4. A-C.** Growth of *E. coli* in the presence of the indicated peptide concentrations given in μM as monitored by OD600 at 37 °C over 18 hours. Each panel indicates an independent experiment. **D-F.** As in **A-C**, but with *P. aeruginosa.*

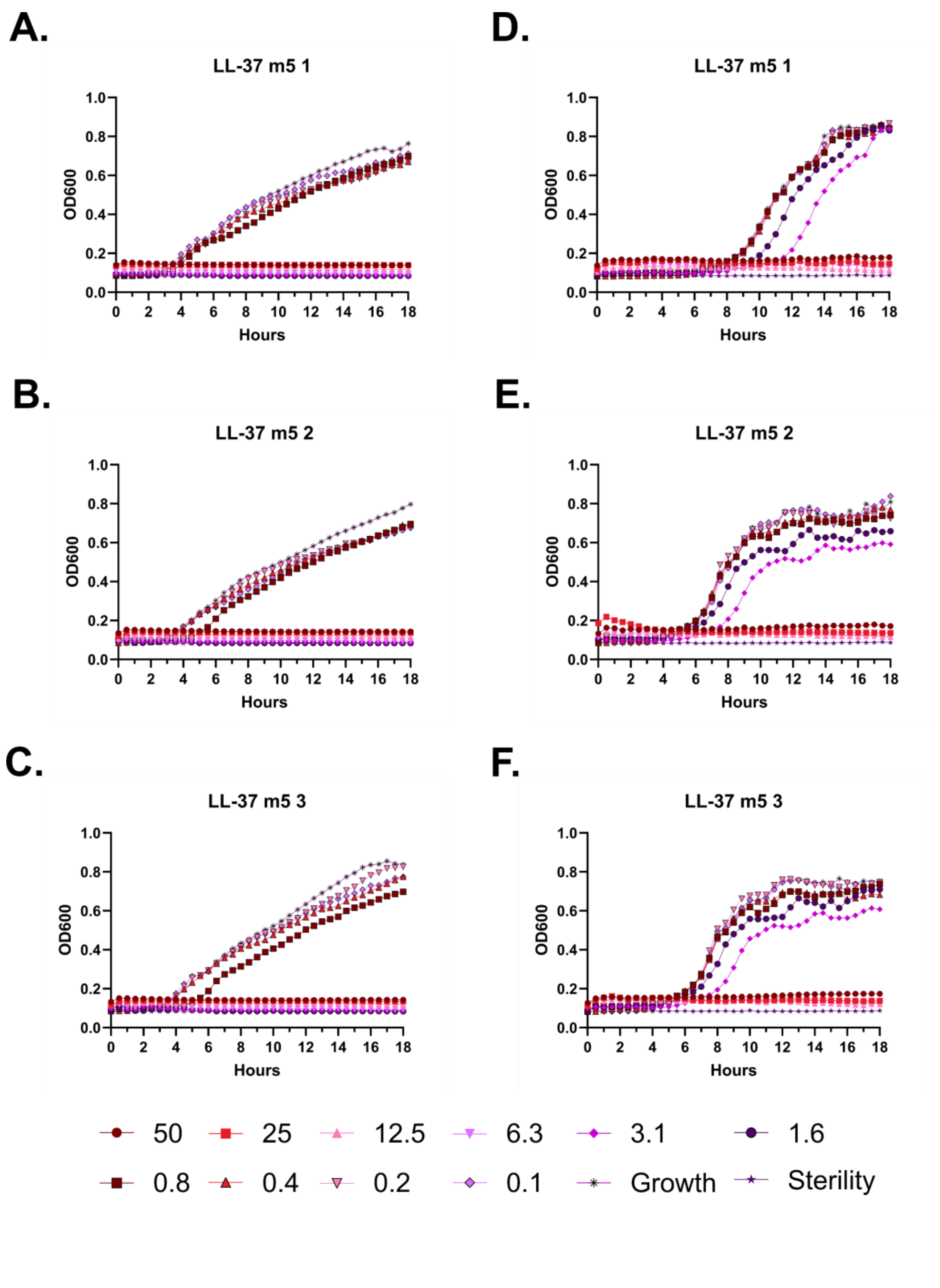
A.
LL-37 m5 1
D.
LL-37 m5 1
B.
LL-37 m5 2
E.
LL-37 m5 2
C.
LL-37 m5 3
F.
LL-37 m5 3
OD600
Hours
50
25
12.5
6.3
3.1
1.6
0.8
0.4
0.2
0.1
Growth
Sterility

**Figure S11 – Antimicrobial susceptibility testing with LL-37 m5. A-C.** Growth of *E. coli* in the presence of the indicated peptide concentrations given in μM as monitored by OD600 at 37 °C over 18 hours. Each panel indicates an independent experiment. **D-F.** As in **A-C**, but with *P. aeruginosa.*

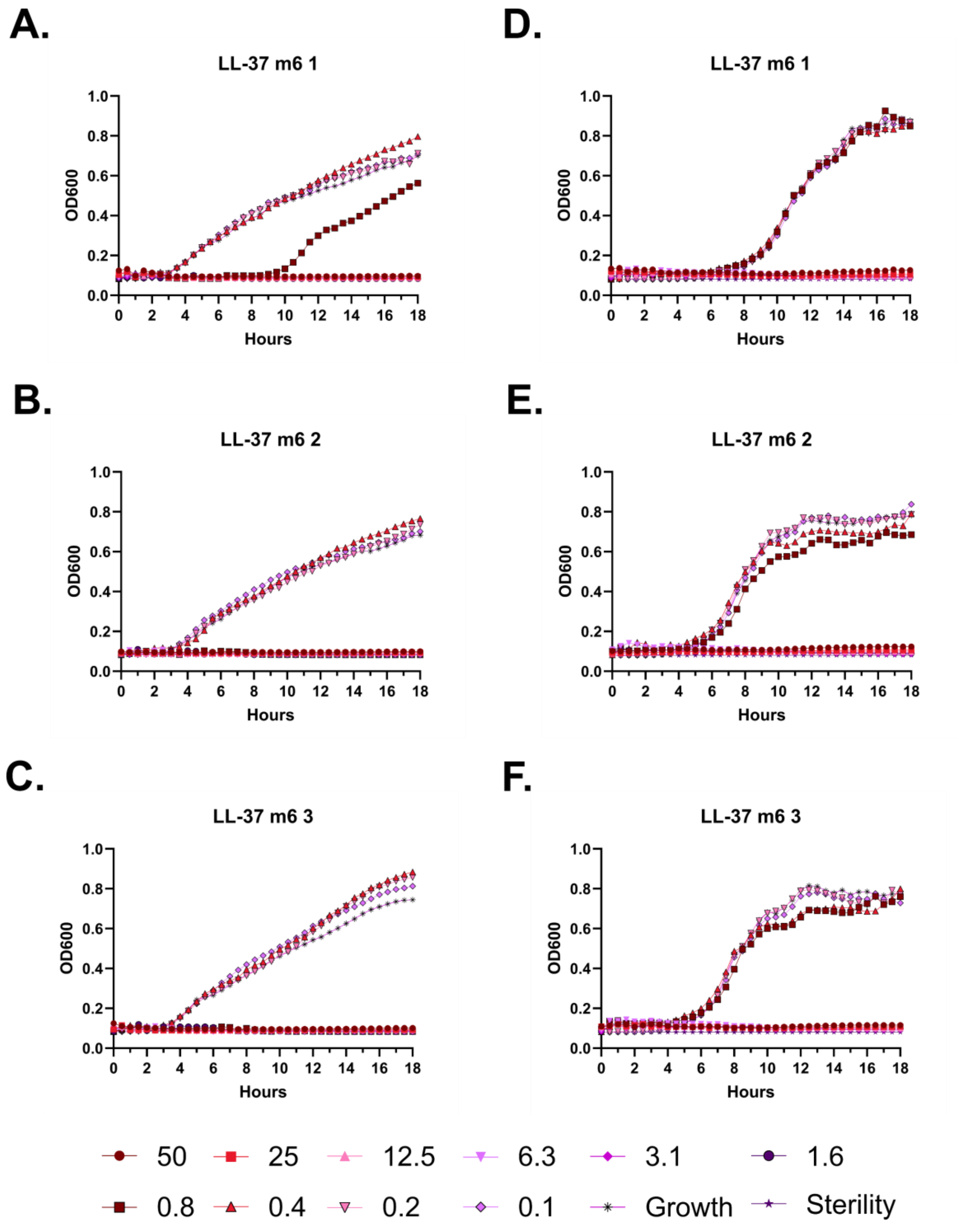
A.
LL-37 m6 1
B.
LL-37 m6 2
C.
LL-37 m6 3
D.
LL-37 m6 1
E.
LL-37 m6 2
F.
LL-37 m6 3
OD600
Hours
50
25
12.5
6.3
3.1
1.6
0.8
0.4
0.2
0.1
Growth
Sterility

**Figure S12 – Antimicrobial susceptibility testing with LL-37 m6. A-C.** Growth of *E. coli* in the presence of the indicated peptide concentrations given in μM as monitored by OD600 at 37 °C over 18 hours. Each panel indicates an independent experiment. **D-F.** As in **A-C**, but with *P. aeruginosa.*

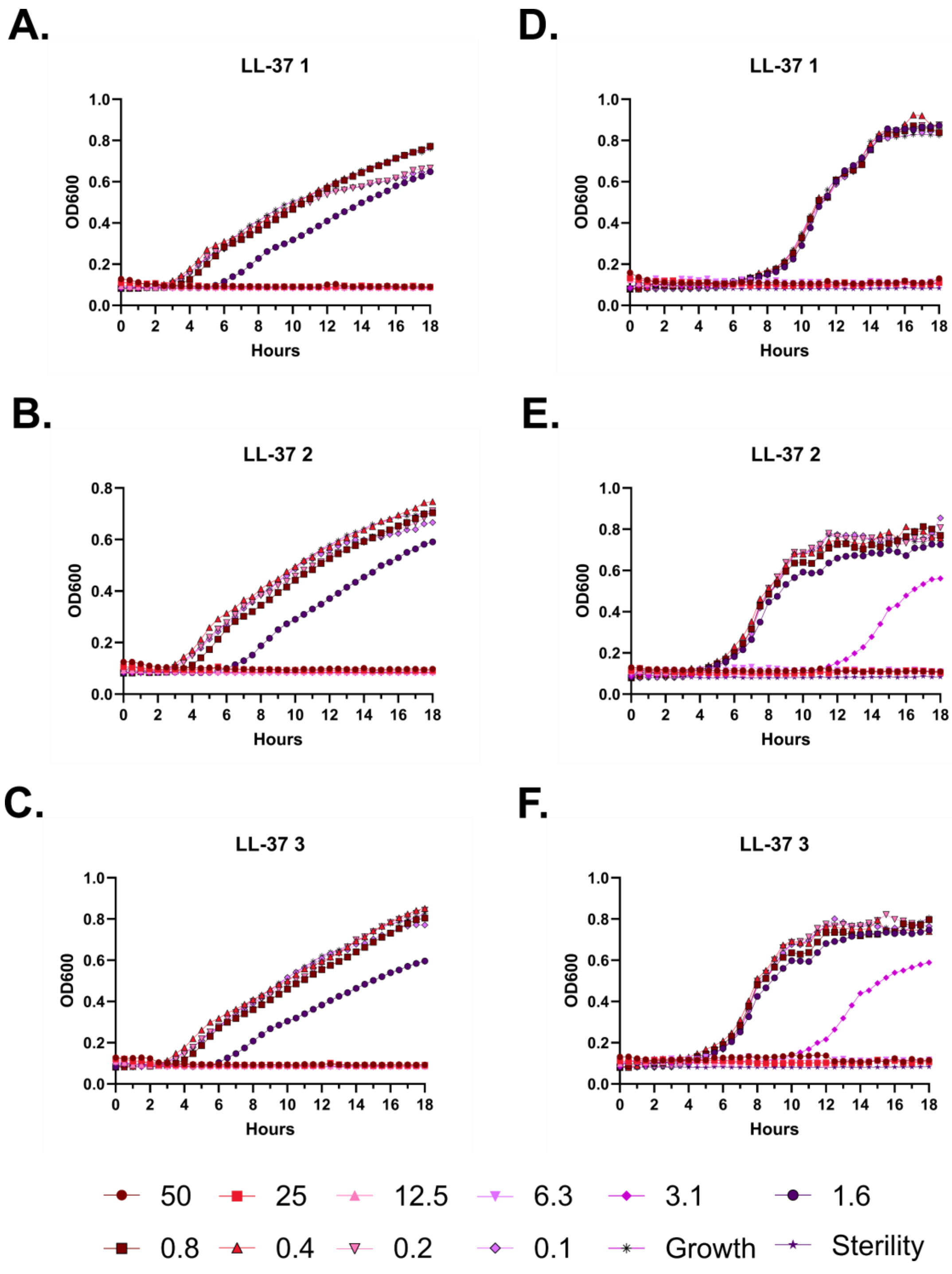
A.
LL-37 1
B.
LL-37 2
C.
LL-37 3
D.
LL-37 1
E.
LL-37 2
F.
LL-37 3
OD600
Hours
50
25
12.5
6.3
3.1
1.6
0.8
0.4
0.2
0.1
Growth
Sterility

**Figure S13 – Antimicrobial susceptibility testing with LL-37. A-C.** Growth of *E. coli* in the presence of the indicated peptide concentrations given in µM as monitored by OD600 at 37 °C over 18 hours. Each panel indicates an independent experiment. **D-F.** As in **A-C**, but with *P. aeruginosa.*

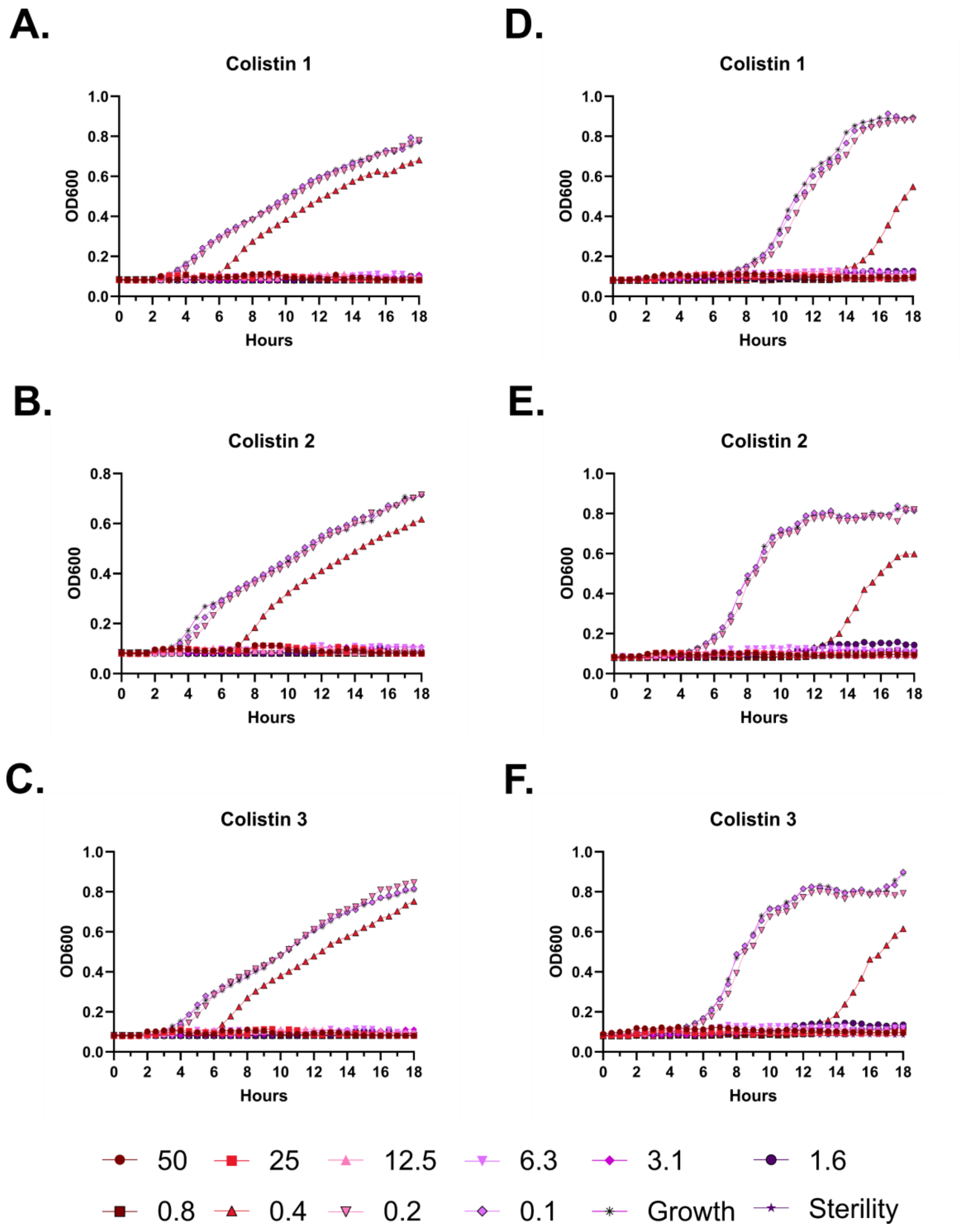
A.
Colistin 1
OD600
Hours
B.
Colistin 2
OD600
Hours
C.
Colistin 3
OD600
Hours
D.
Colistin 1
OD600
Hours
E.
Colistin 2
OD600
Hours
F.
Colistin 3
OD600
Hours
50
25
12.5
6.3
3.1
1.6
0.8
0.4
0.2
0.1
Growth
Sterility

**Figure S14 – Antimicrobial susceptibility testing with colistin. A-C.** Growth of *E. coli* in the presence of the indicated peptide concentrations given in μM as monitored by OD600 at 37 °C over 18 hours. Each panel indicates an independent experiment. **D-F.** As in **A-C**, but with *P. aeruginosa.*

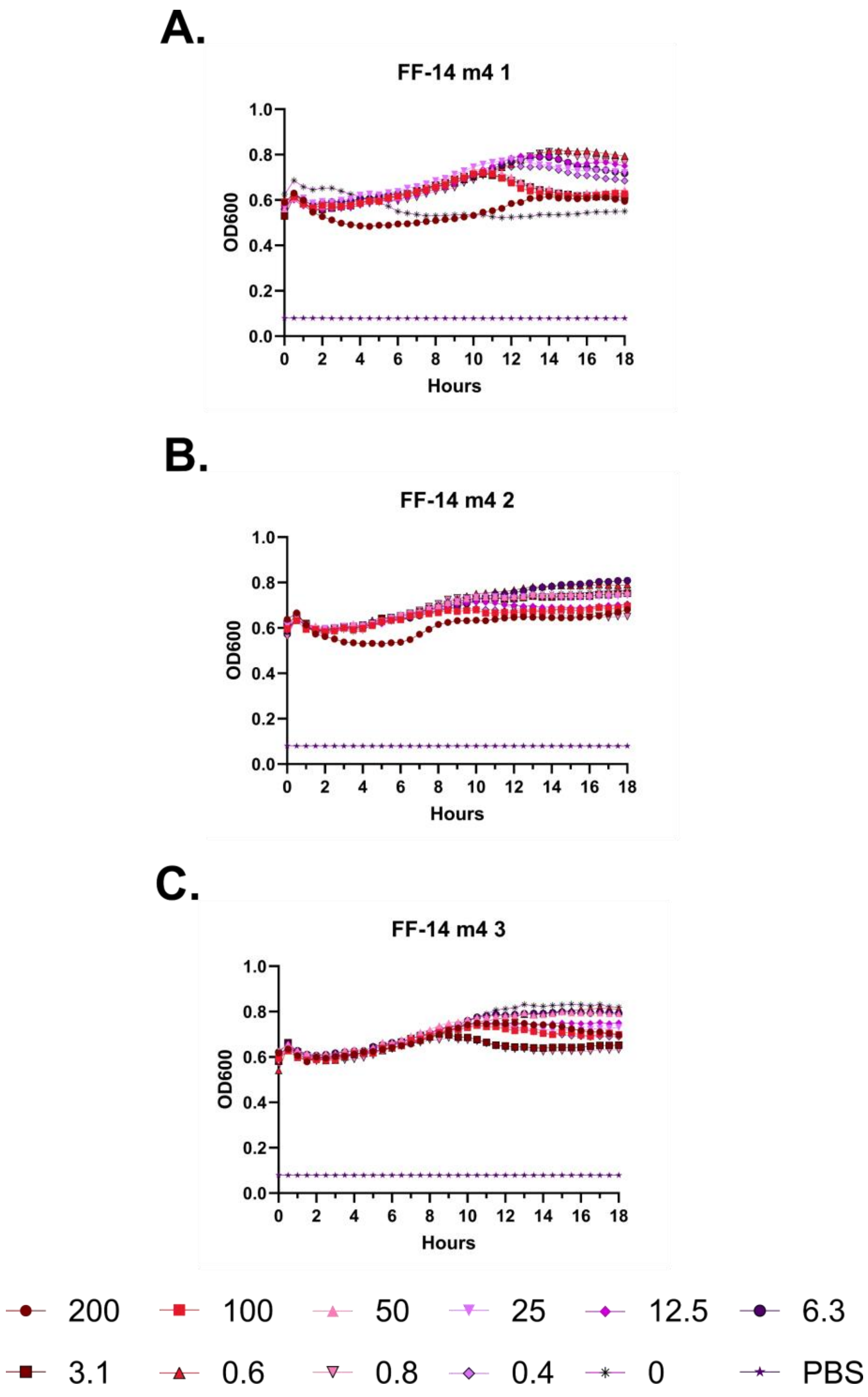
A.
FF-14 m4 1
OD600
Hours
B.
FF-14 m4 2
OD600
Hours
C.
FF-14 m4 3
OD600
Hours
200
100
50
25
12.5
6.3
3.1
0.6
0.8
0.4
0
PBS

**Figure S15 – Hemolysis over time with FF-14 m4. A-C.** Each of three independent experiments showing trends in OD600 over time on incubation of defibrinated sheep blood with the indicated concentrations of peptide in μM at 37 °C. Decreasing OD600 indicates increasing hemolysis.

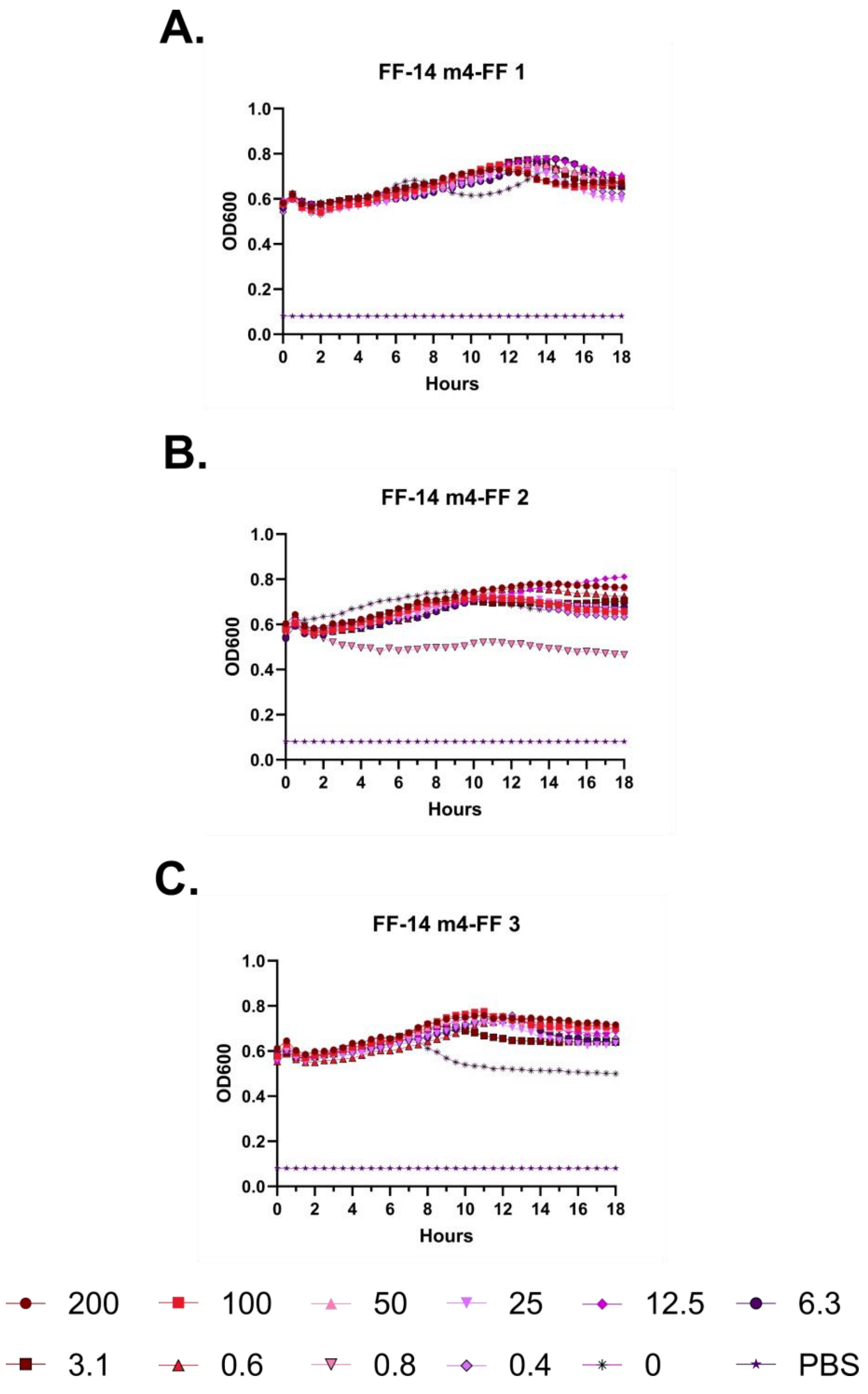
A.
FF-14 m4-FF 1
OD600
Hours
B.
FF-14 m4-FF 2
OD600
Hours
C.
FF-14 m4-FF 3
OD600
Hours
200
100
50
25
12.5
6.3
3.1
0.6
0.8
0.4
0
PBS

**Figure S16 – Hemolysis over time with FF-14 m4-FF. A-C.** Each of three independent experiments showing trends in OD600 over time on incubation of defibrinated sheep blood with the indicated concentrations of peptide in μM at 37 °C. Decreasing OD600 indicates increasing hemolysis.

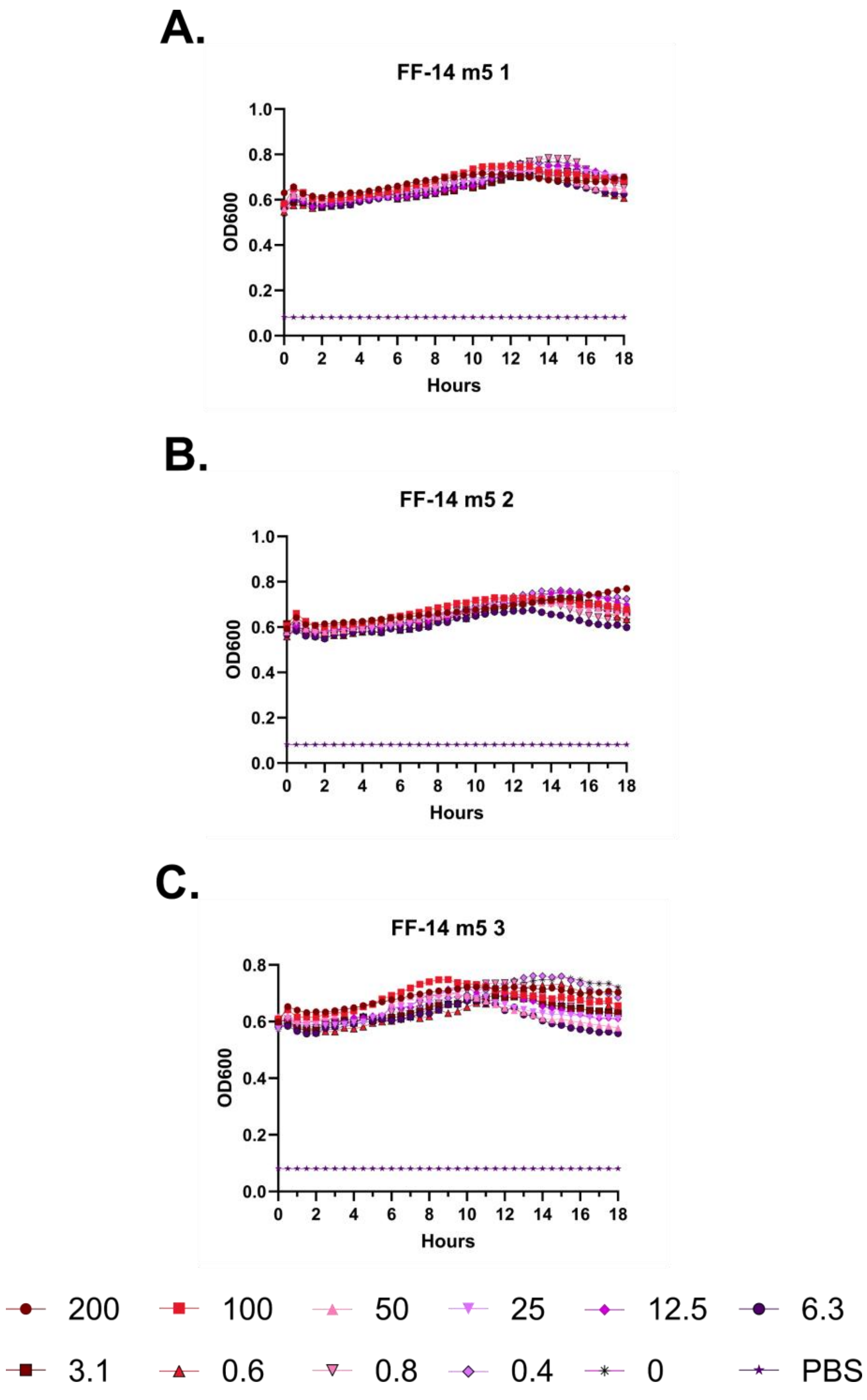
A.
FF-14 m5 1
OD600
Hours
B.
FF-14 m5 2
OD600
Hours
C.
FF-14 m5 3
OD600
Hours
200
100
50
25
12.5
6.3
3.1
0.6
0.8
0.4
0
PBS

**Figure S17 – Hemolysis over time with FF-14 m5. A-C.** Each of three independent experiments showing trends in OD600 over time on incubation of defibrinated sheep blood with the indicated concentrations of peptide in μM at 37 °C. Decreasing OD600 indicates increasing hemolysis.

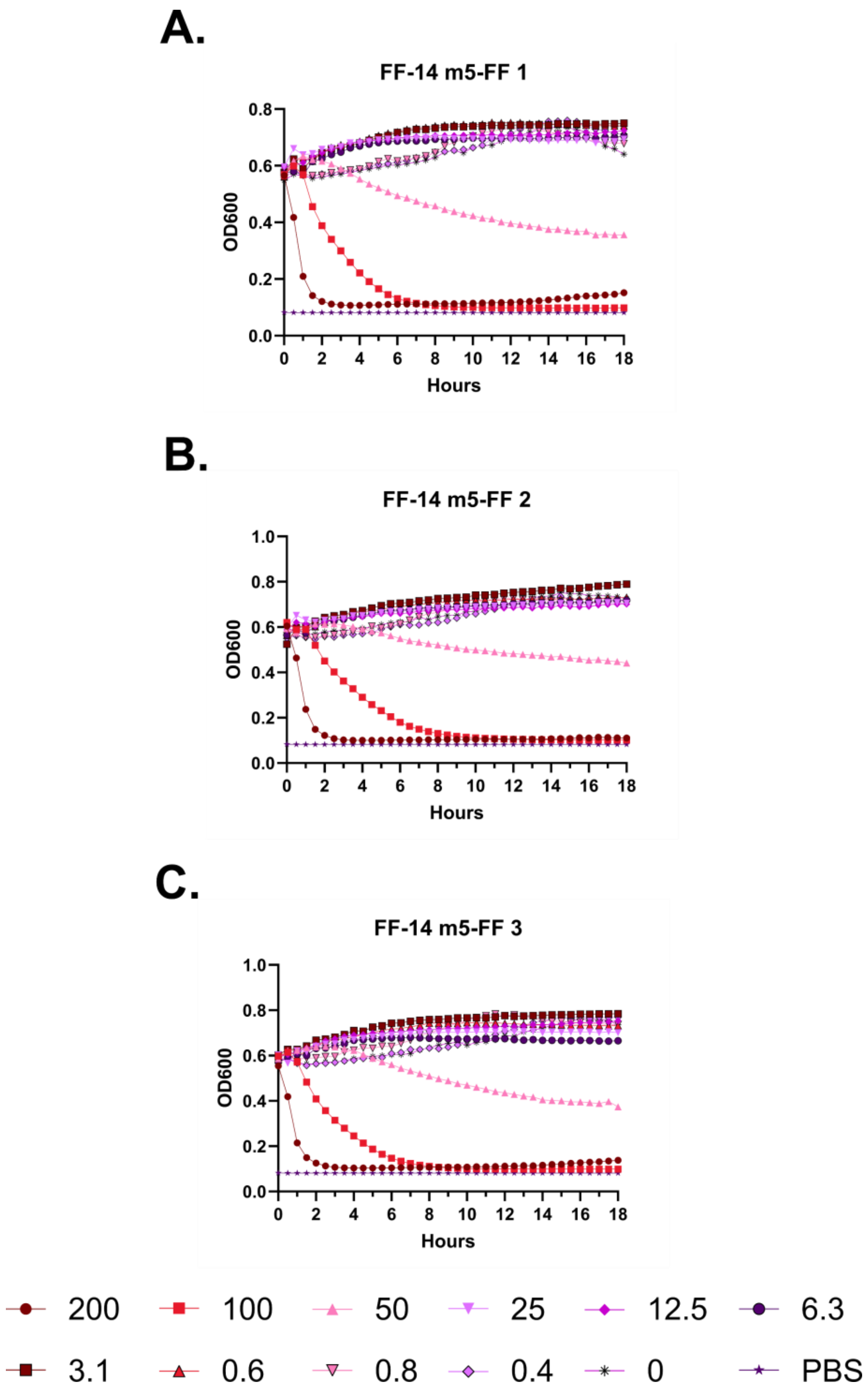

A.
FF-14 m5-FF 1
OD600
Hours
B.
FF-14 m5-FF 2
OD600
Hours
C.
FF-14 m5-FF 3
OD600
Hours
200
100
50
25
12.5
6.3
3.1
0.6
0.8
0.4
0
PBS

**Figure S18 – Hemolysis over time with FF-14 m5-FF. A-C.** Each of three independent experiments showing trends in OD600 over time on incubation of defibrinated sheep blood with the indicated concentrations of peptide in μM at 37 °C. Decreasing OD600 indicates increasing hemolysis.

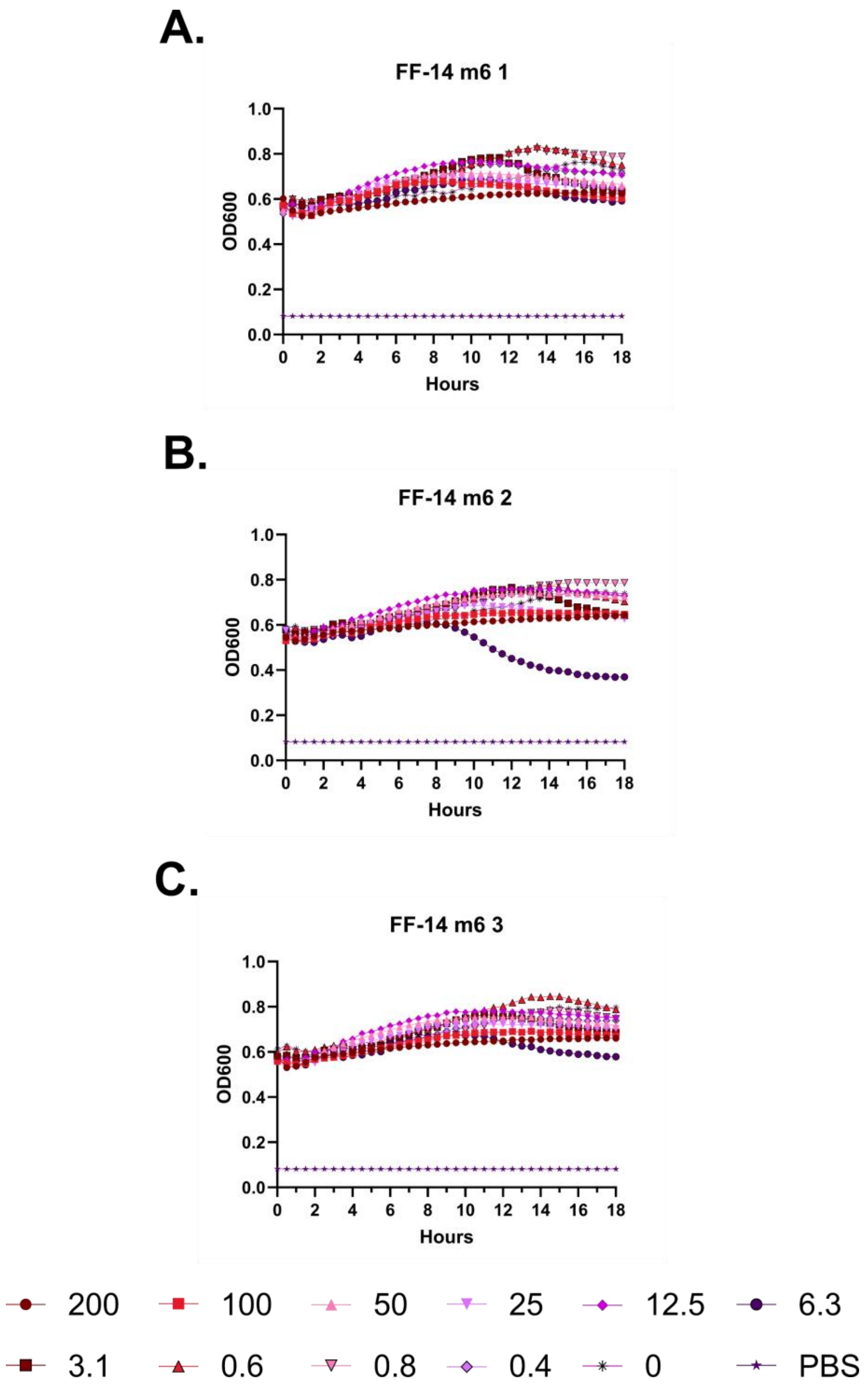
A.
FF-14 m6 1
OD600
Hours
B.
FF-14 m6 2
OD600
Hours
C.
FF-14 m6 3
OD600
Hours
200
100
50
25
12.5
6.3
3.1
0.6
0.8
0.4
0
PBS

**Figure S19 – Hemolysis over time with FF-14 m6. A-C.** Each of three independent experiments showing trends in OD600 over time on incubation of defibrinated sheep blood with the indicated concentrations of peptide in µM at 37 °C. Decreasing OD600 indicates increasing hemolysis.

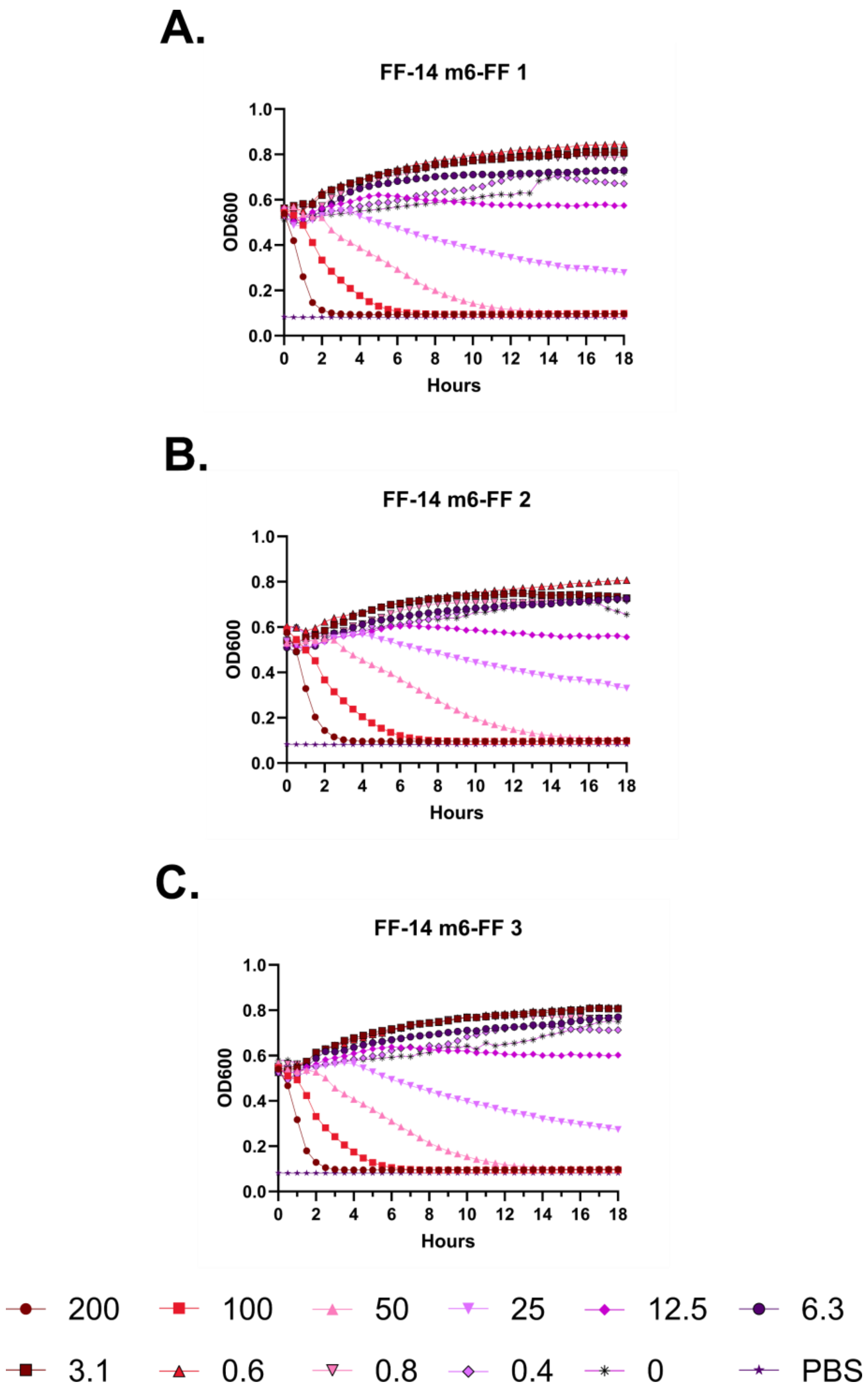
A.
FF-14 m6-FF 1
OD600
Hours
B.
FF-14 m6-FF 2
OD600
Hours
C.
FF-14 m6-FF 3
OD600
Hours
200
100
50
25
12.5
6.3
3.1
0.6
0.8
0.4
0
PBS

**Figure S20 – Hemolysis over time with FF-14 m6-FF. A-C.** Each of three independent experiments showing trends in OD600 over time on incubation of defibrinated sheep blood with the indicated concentrations of peptide in μM at 37 °C. Decreasing OD600 indicates increasing hemolysis.

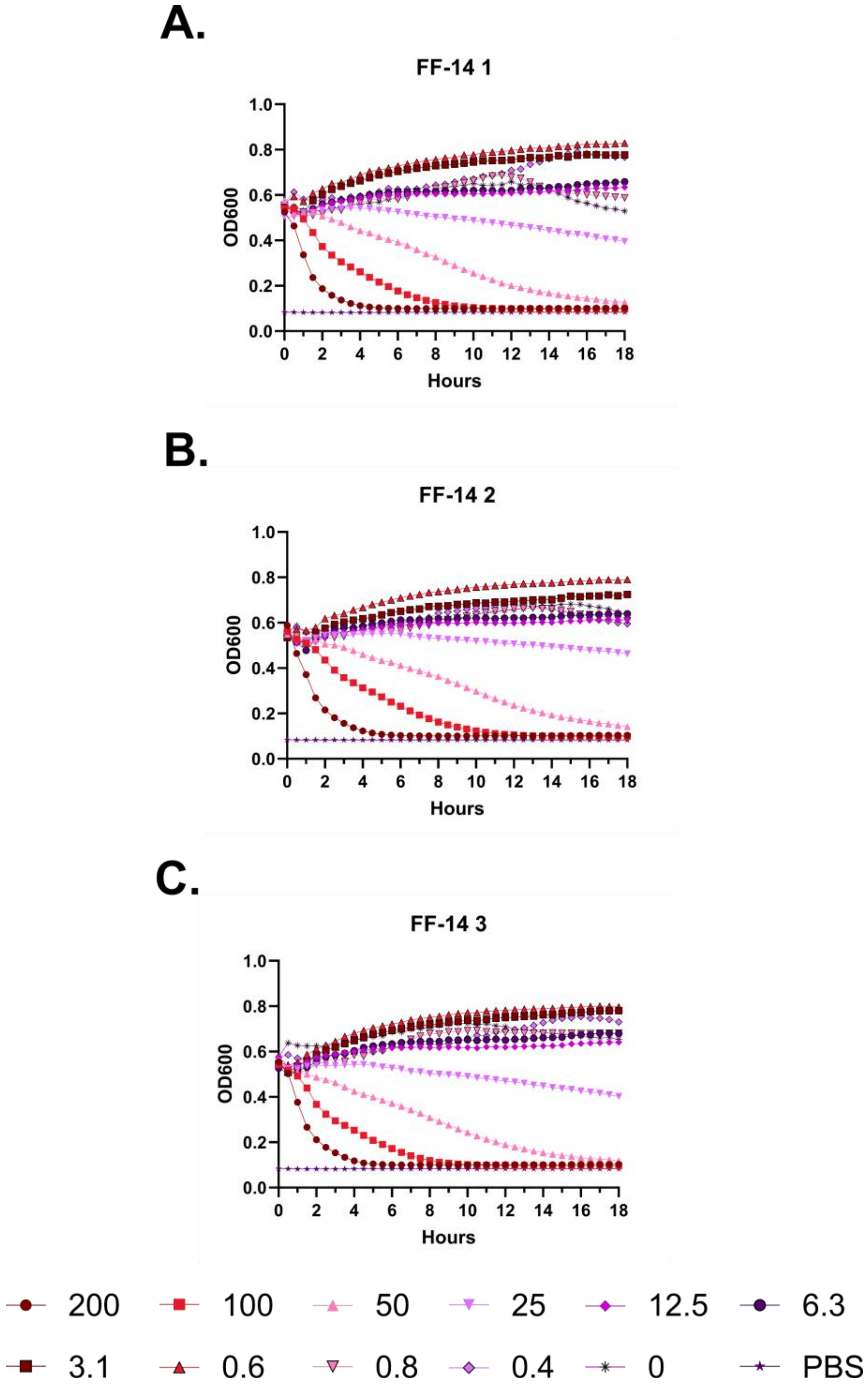
A.
FF-14 1
OD600
Hours
B.
FF-14 2
OD600
Hours
C.
FF-14 3
OD600
Hours
200
100
50
25
12.5
6.3
3.1
0.6
0.8
0.4
0
PBS

**Figure S21 – Hemolysis over time with FF-14. A-C.** Each of three independent experiments showing trends in OD600 over time on incubation of defibrinated sheep blood with the indicated concentrations of peptide in μM at 37 °C. Decreasing OD600 indicates increasing hemolysis.

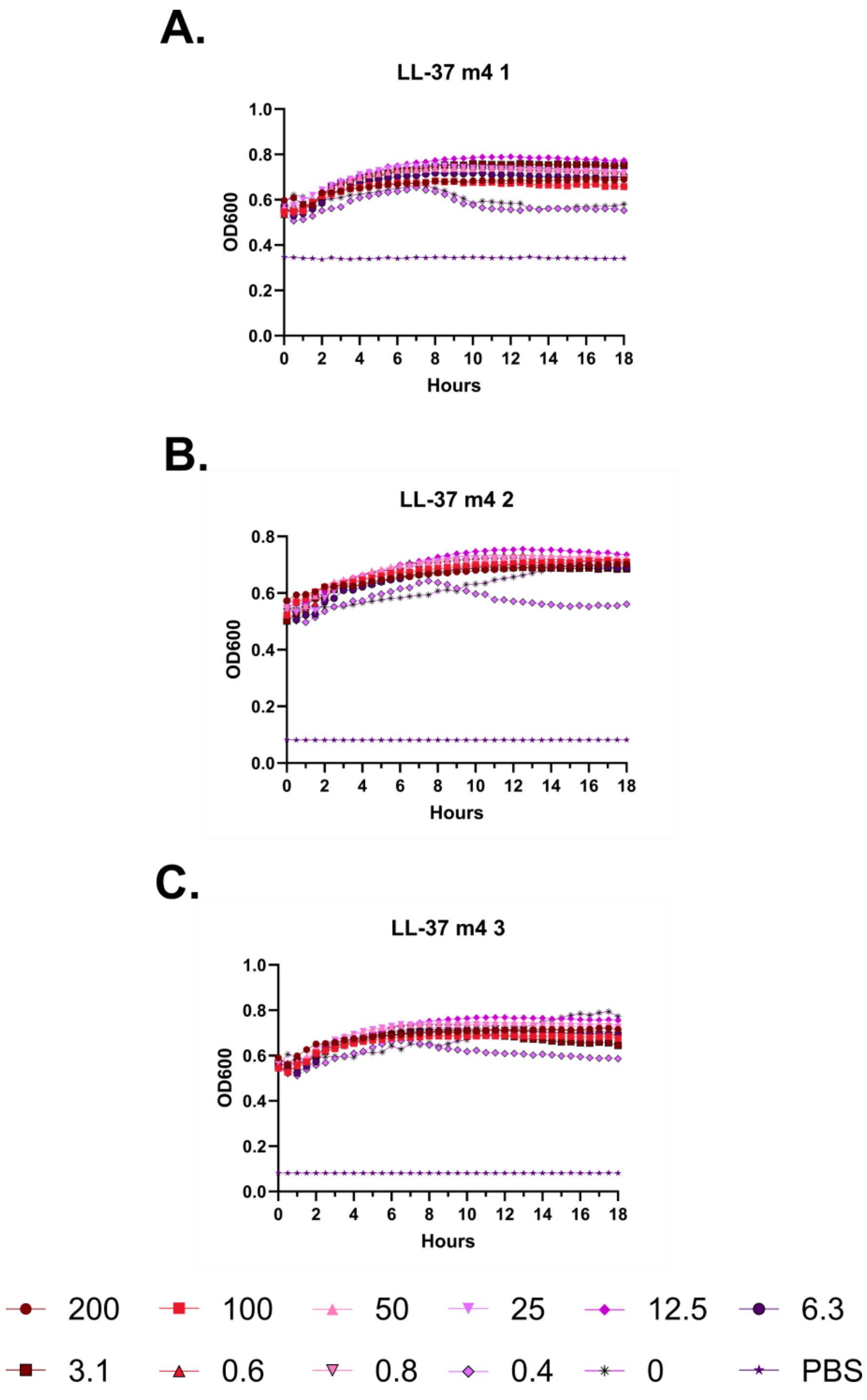
A.
LL-37 m4 1
OD600
Hours
B.
LL-37 m4 2
OD600
Hours
C.
LL-37 m4 3
OD600
Hours
200
100
50
25
12.5
6.3
3.1
0.6
0.8
0.4
0
PBS

**Figure S22 – Hemolysis over time with LL-37 m4 A-C.** Each of three independent experiments showing trends in OD600 over time on incubation of defibrinated sheep blood with the indicated concentrations of peptide in μM at 37 °C. Decreasing OD600 indicates increasing hemolysis.

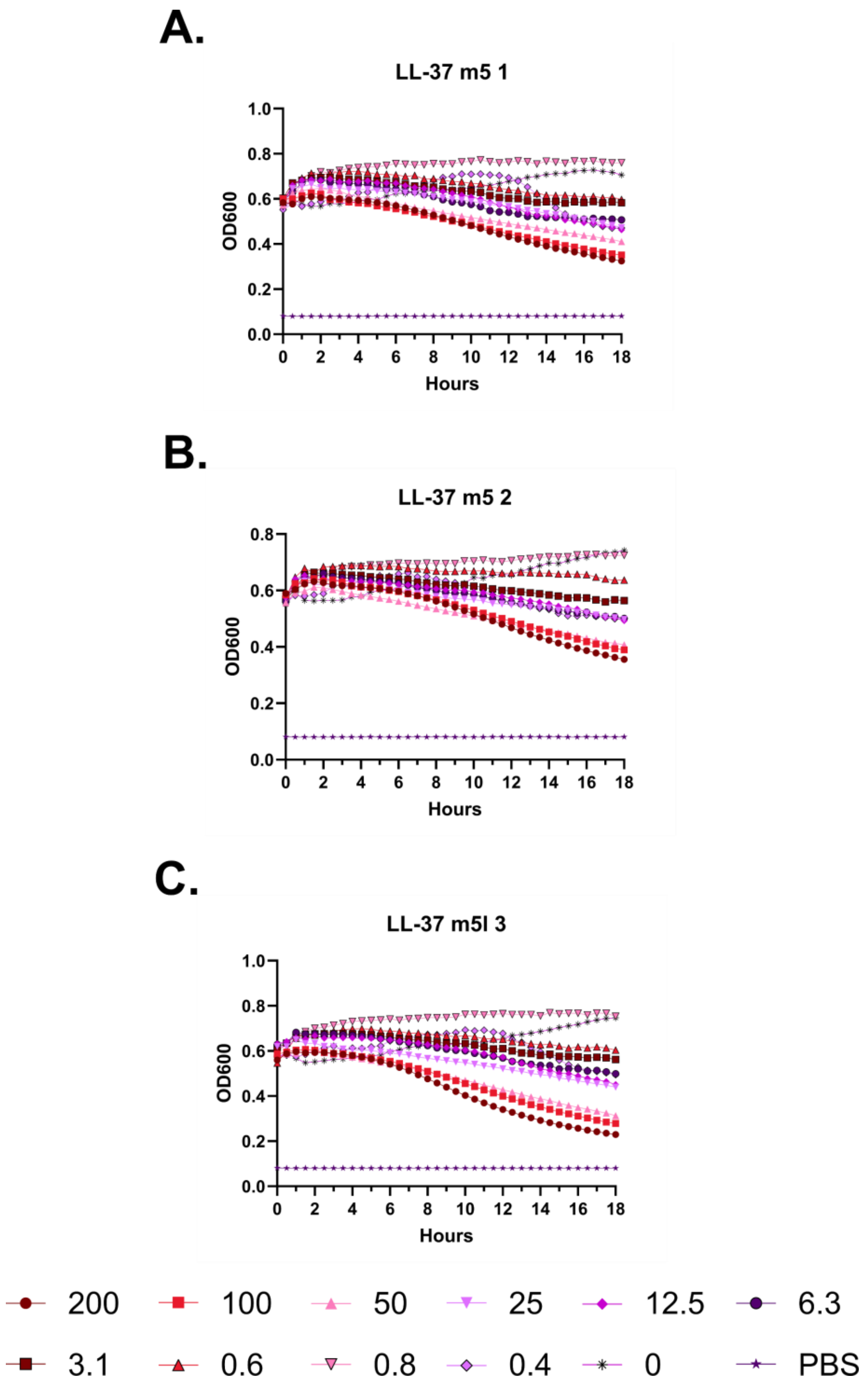
A.
LL-37 m5 1
OD600
Hours
B.
LL-37 m5 2
OD600
Hours
C.
LL-37 m5l 3
OD600
Hours
200
100
50
25
12.5
6.3
3.1
0.6
0.8
0.4
0
PBS

**Figure S23 – Hemolysis over time with LL-37 m5. A-C.** Each of three independent experiments showing trends in OD600 over time on incubation of defibrinated sheep blood with the indicated concentrations of peptide in μM at 37 °C. Decreasing OD600 indicates increasing hemolysis.

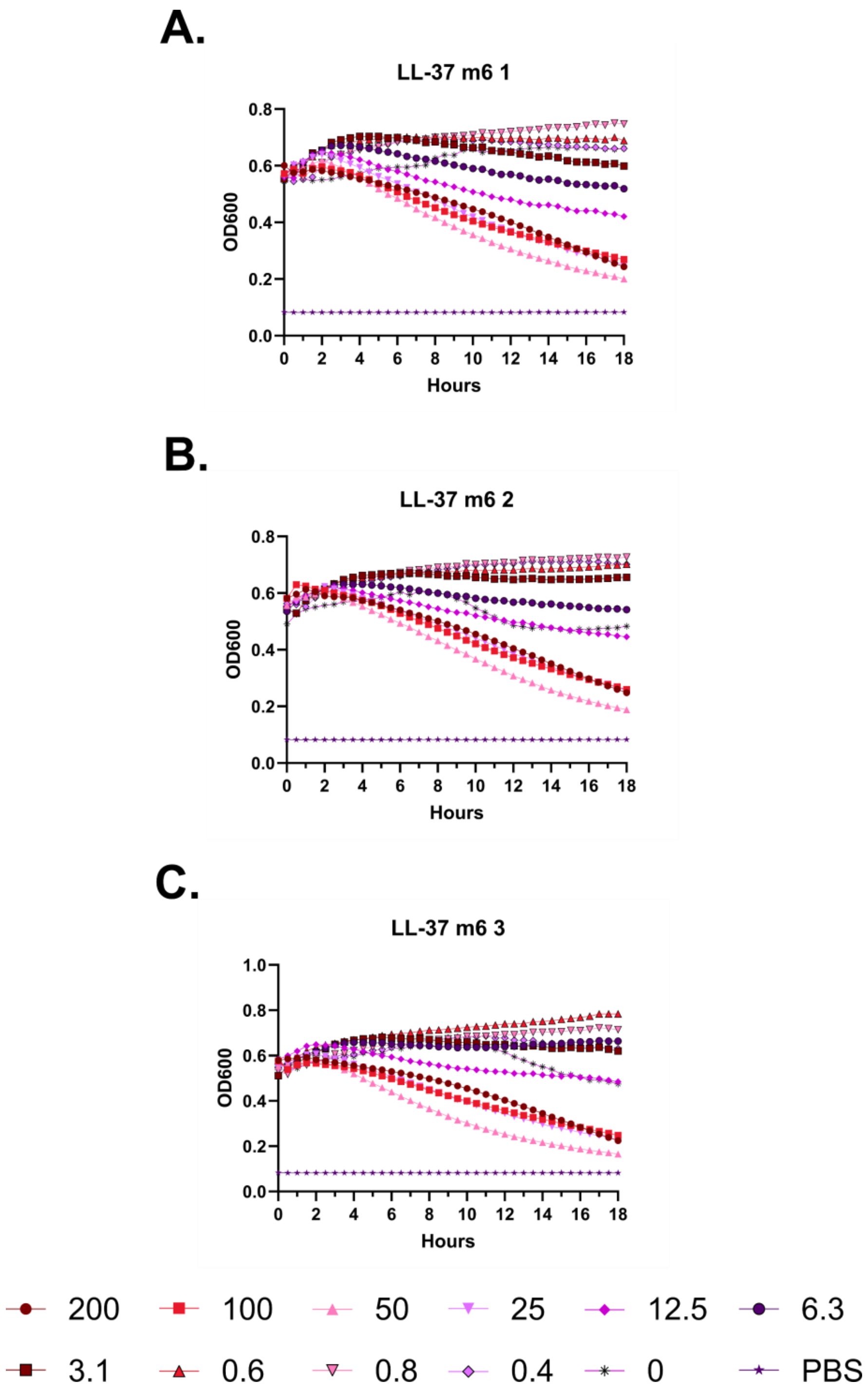
A.
LL-37 m6 1
OD600
Hours
0.0
0.2
0.4
0.6
0.8
0 2 4 6 8 10 12 14 16 18
B.
LL-37 m6 2
OD600
Hours
0.0
0.2
0.4
0.6
0.8
0 2 4 6 8 10 12 14 16 18
C.
LL-37 m6 3
OD600
Hours
0.0
0.2
0.4
0.6
0.8
1.0
0 2 4 6 8 10 12 14 16 18
200
100
50
25
12.5
6.3
3.1
0.6
0.8
0.4
0
PBS

**Figure S24 – Hemolysis over time with LL-37 m6. A-C.** Each of three independent experiments showing trends in OD600 over time on incubation of defibrinated sheep blood with the indicated concentrations of peptide in μM at 37 °C. Decreasing OD600 indicates increasing hemolysis.

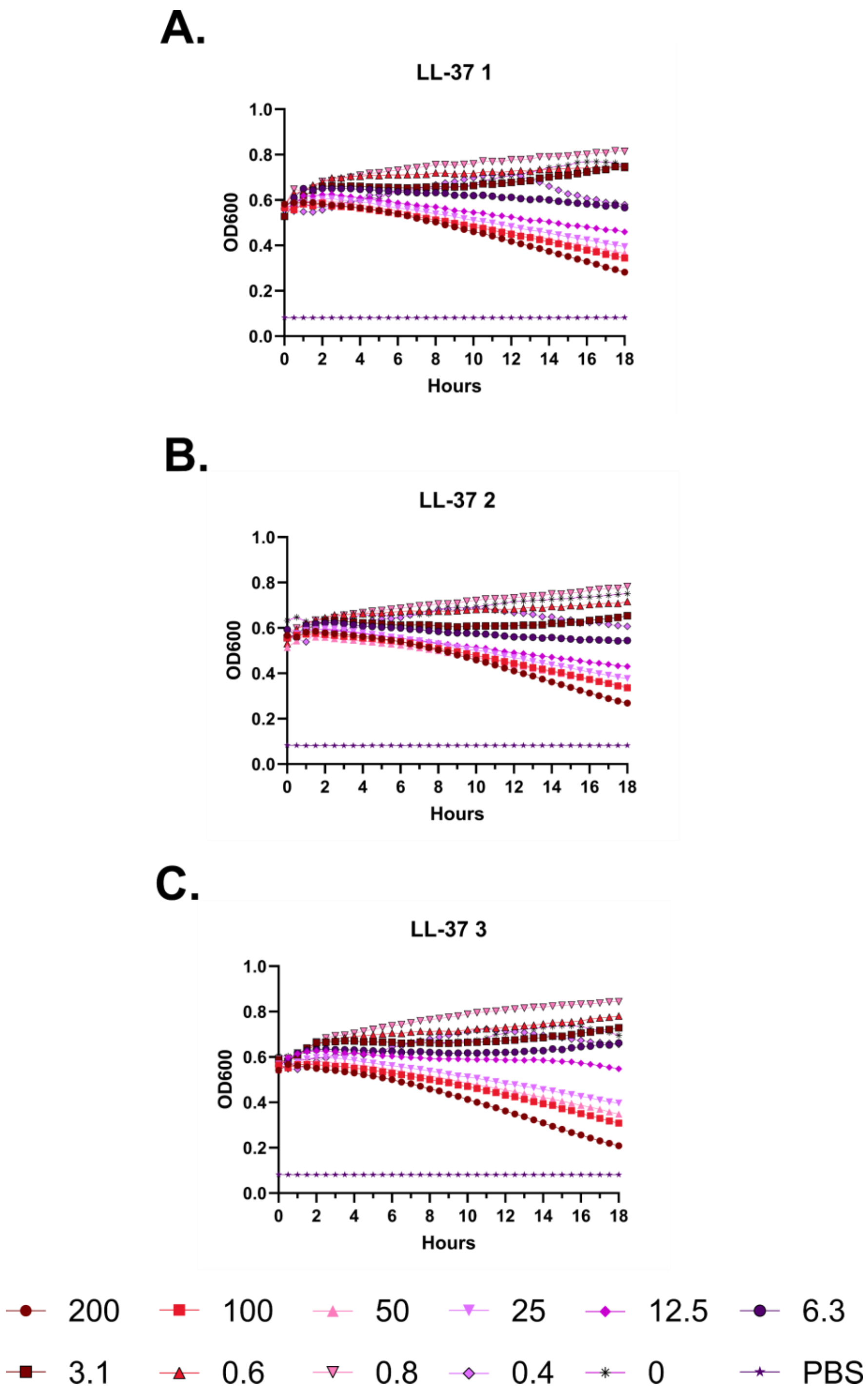
A.
LL-37 1
OD600
Hours
B.
LL-37 2
OD600
Hours
C.
LL-37 3
OD600
Hours
200
100
50
25
12.5
6.3
3.1
0.6
0.8
0.4
0
PBS

**Figure S25 – Hemolysis over time with LL-37. A-C.** Each of three independent experiments showing trends in OD600 over time on incubation of defibrinated sheep blood with the indicated concentrations of peptide in μM at 37 °C. Decreasing OD600 indicates increasing hemolysis.

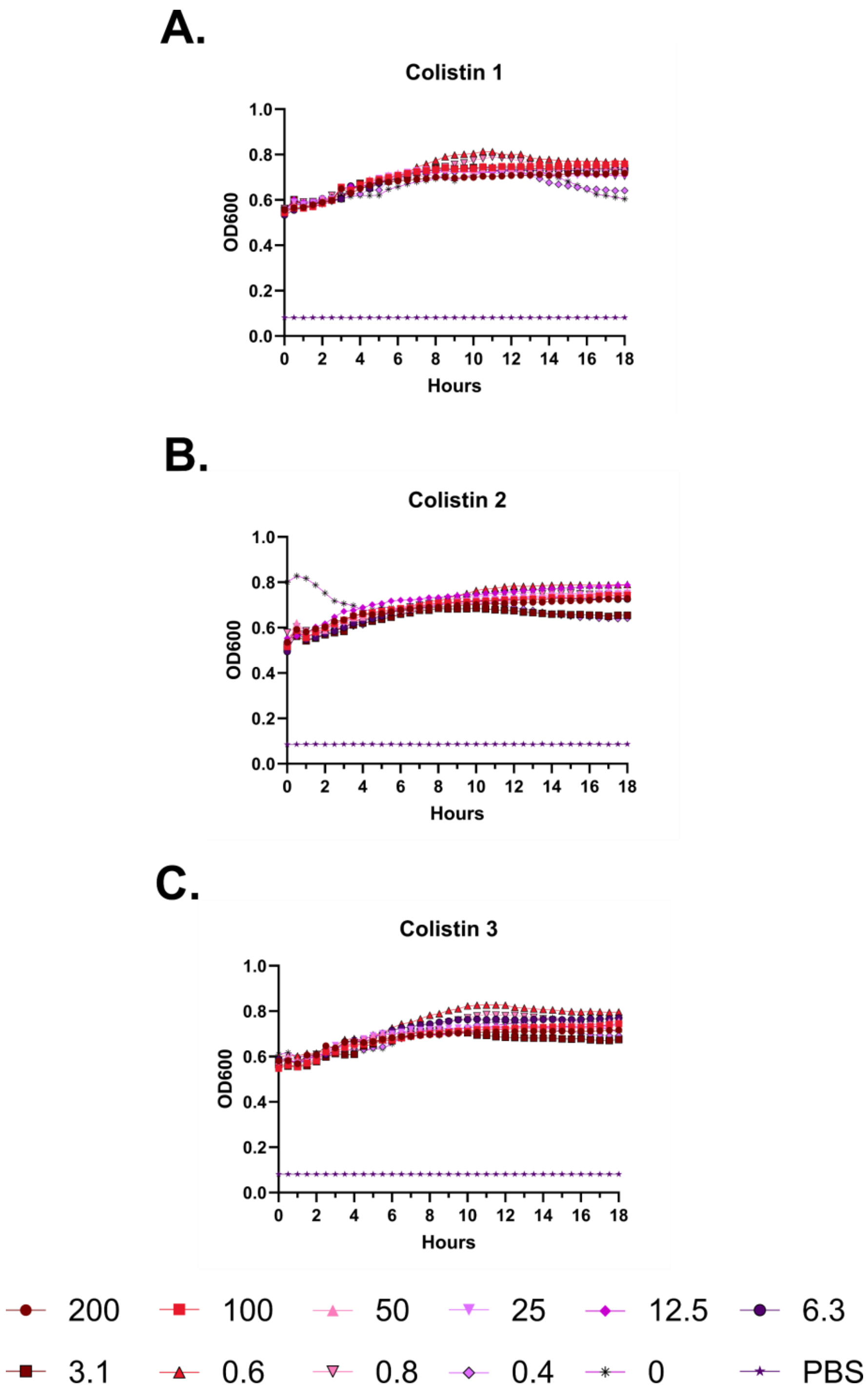
A.
Colistin 1
OD600
Hours
B.
Colistin 2
OD600
Hours
C.
Colistin 3
OD600
Hours
200
100
50
25
12.5
6.3
3.1
0.6
0.8
0.4
0
PBS

**Figure S26 – Hemolysis over time with colistin. A-C.** Each of three independent experiments showing trends in OD600 over time on incubation of defibrinated sheep blood with the indicated concentrations of peptide in μM at 37 °C. Decreasing OD600 indicates increasing hemolysis.

**A.**

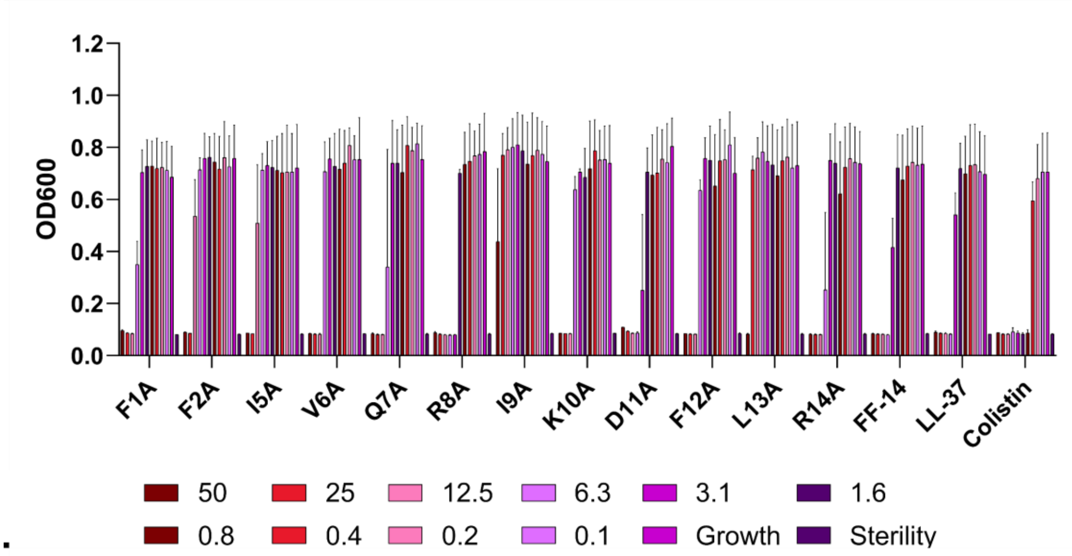

OD600
1.2
1.0
0.8
0.6
0.4
0.2
0.0
F1A
F2A
I5A
V6A
Q7A
R8A
I9A
K10A
D11A
F12A
L13A
R14A
FF-14
LL-37
Colistin
50
25
12.5
6.3
3.1
1.6
0.8
0.4
0.2
0.1
Growth
Sterility


**B.**

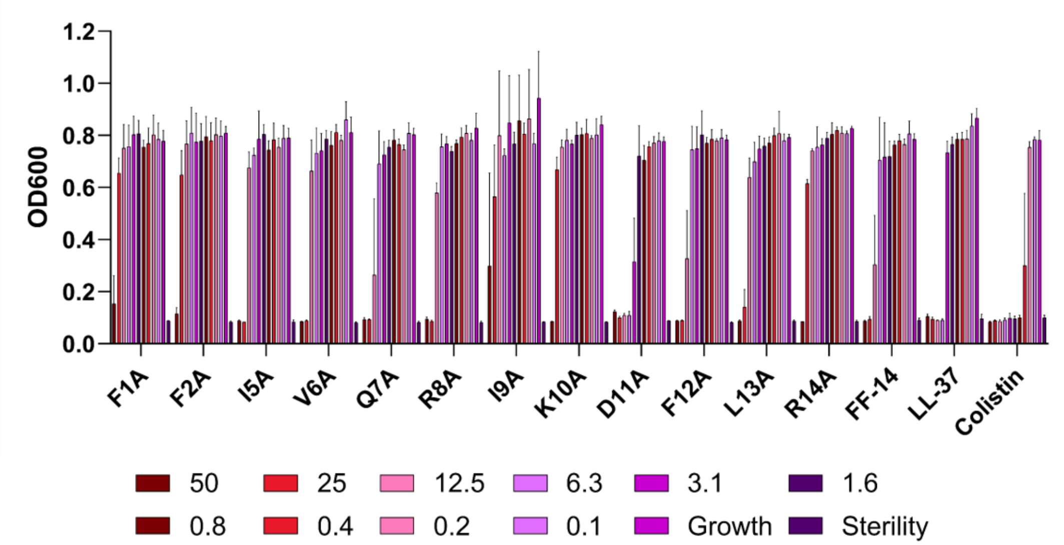

OD600
1.2
1.0
0.8
0.6
0.4
0.2
0.0
F1A
F2A
I5A
V6A
Q7A
R8A
I9A
K10A
D11A
F12A
L13A
R14A
FF-14
LL-37
Colistin
50
25
12.5
6.3
3.1
1.6
0.8
0.4
0.2
0.1
Growth
Sterility


**C.**

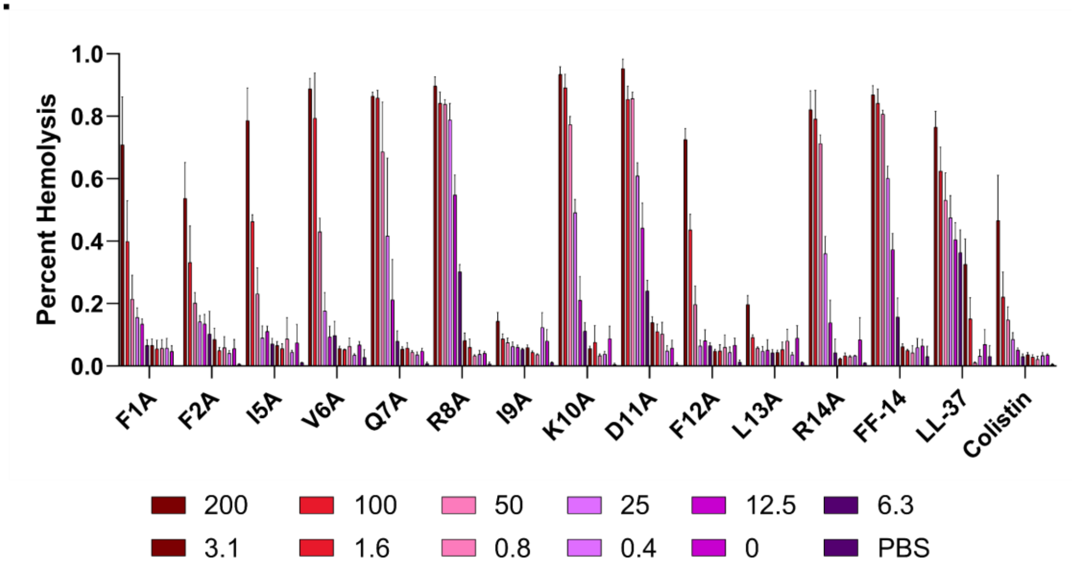

Percent Hemolysis
1.0
0.8
0.6
0.4
0.2
0.0
F1A
F2A
I5A
V6A
Q7A
R8A
I9A
K10A
D11A
F12A
L13A
R14A
FF-14
LL-37
Colistin
200
100
50
25
12.5
6.3
3.1
1.6
0.8
0.4
0
PBS

**Figure S27 – Histogram depiction of data from Figure 2. A.** Endpoint activity of each peptide against *E. coli*; measures are raw absorbance at 600 nm at 18 hours. **B.** Endpoint activity of each peptide against *P. aeruginosa*. **C.** Hemolytic activity of each peptide; measures are absorbance at 414 nm scaled to a freeze-thaw positive control set at 100% hemolysis. Concentrations throughout are shown are in μM.

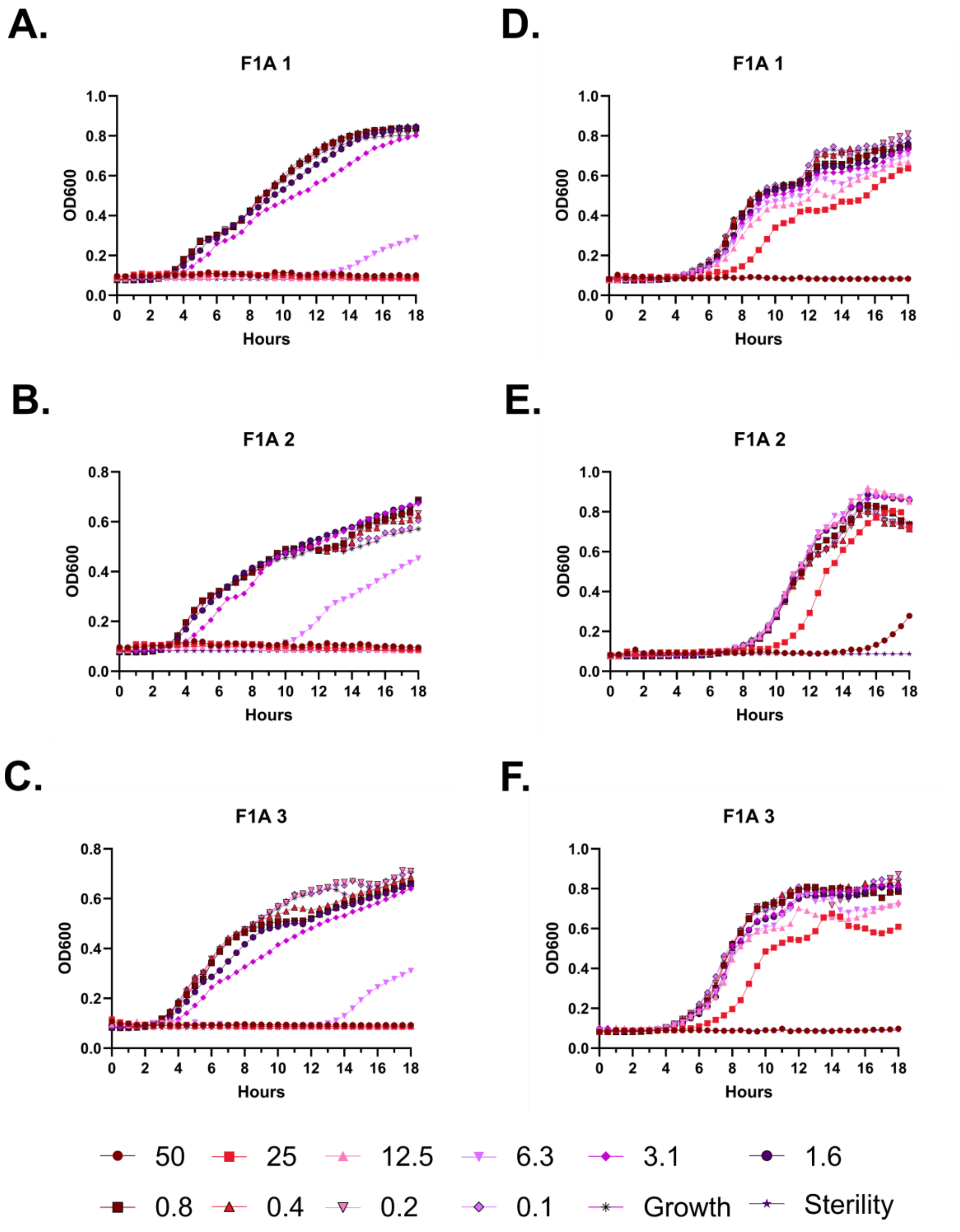
A.
F1A 1
OD600
Hours
B.
F1A 2
OD600
Hours
C.
F1A 3
OD600
Hours
D.
F1A 1
OD600
Hours
E.
F1A 2
OD600
Hours
F.
F1A 3
OD600
Hours
50
25
12.5
6.3
3.1
1.6
0.8
0.4
0.2
0.1
Growth
Sterility

**Figure S28 – Antimicrobial susceptibility testing with FF-14 F1A. A-C.** Growth of *E. coli* in the presence of the indicated peptide concentrations given in µM as monitored by OD600 at 37 °C over 18 hours. Each panel indicates an independent experiment. **D-F.** As in **A-C**, but with *P. aeruginosa.*

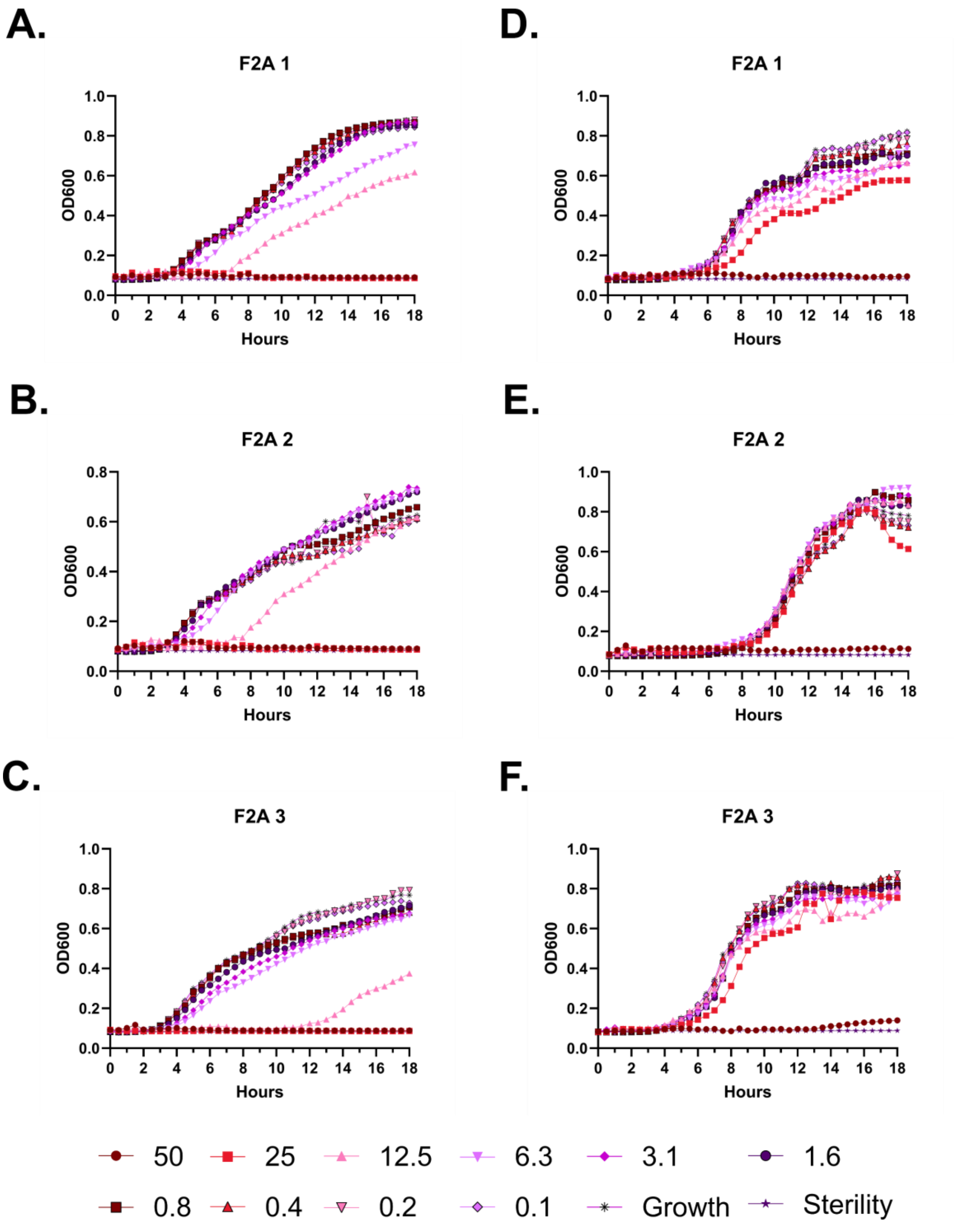
A.
F2A 1
OD600
Hours
D.
F2A 1
OD600
Hours
B.
F2A 2
OD600
Hours
E.
F2A 2
OD600
Hours
C.
F2A 3
OD600
Hours
F.
F2A 3
OD600
Hours
50
25
12.5
6.3
3.1
1.6
0.8
0.4
0.2
0.1
Growth
Sterility

**Figure S29 – Antimicrobial susceptibility testing with FF-14 F2A. A-C.** Growth of *E. coli* in the presence of the indicated peptide concentrations given in μM as monitored by OD600 at 37 °C over 18 hours. Each panel indicates an independent experiment. **D-F.** As in **A-C**, but with *P. aeruginosa.*

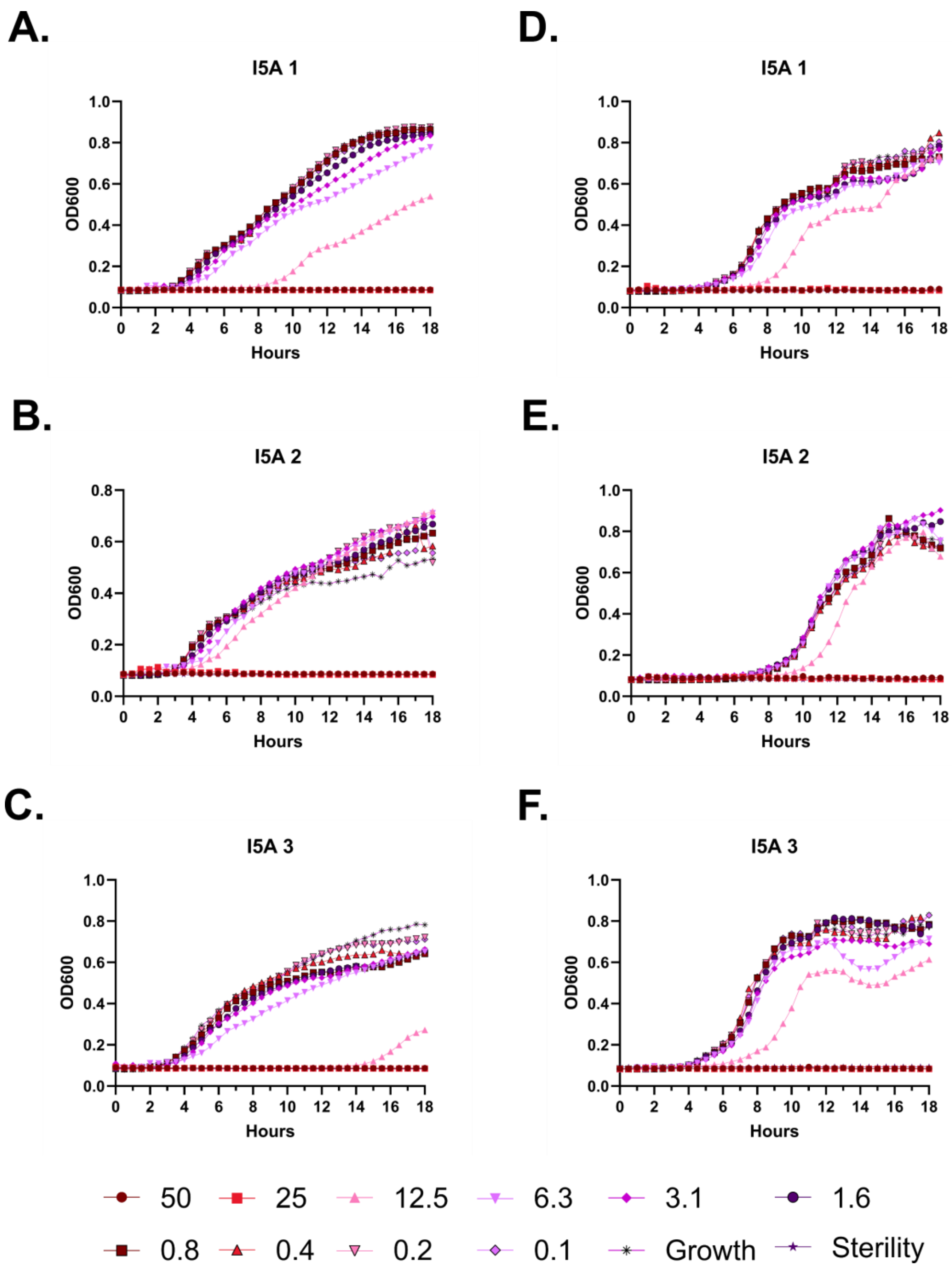
A.
I5A 1
OD600
Hours
B.
I5A 2
OD600
Hours
C.
I5A 3
OD600
Hours
D.
I5A 1
OD600
Hours
E.
I5A 2
OD600
Hours
F.
I5A 3
OD600
Hours
50
25
12.5
6.3
3.1
1.6
0.8
0.4
0.2
0.1
Growth
Sterility

**Figure S30 – Antimicrobial susceptibility testing with FF-14 I5A. A-C.** Growth of *E. coli* in the presence of the indicated peptide concentrations given in μM as monitored by OD600 at 37 °C over 18 hours. Each panel indicates an independent experiment. **D-F.** As in **A-C**, but with *P. aeruginosa.*

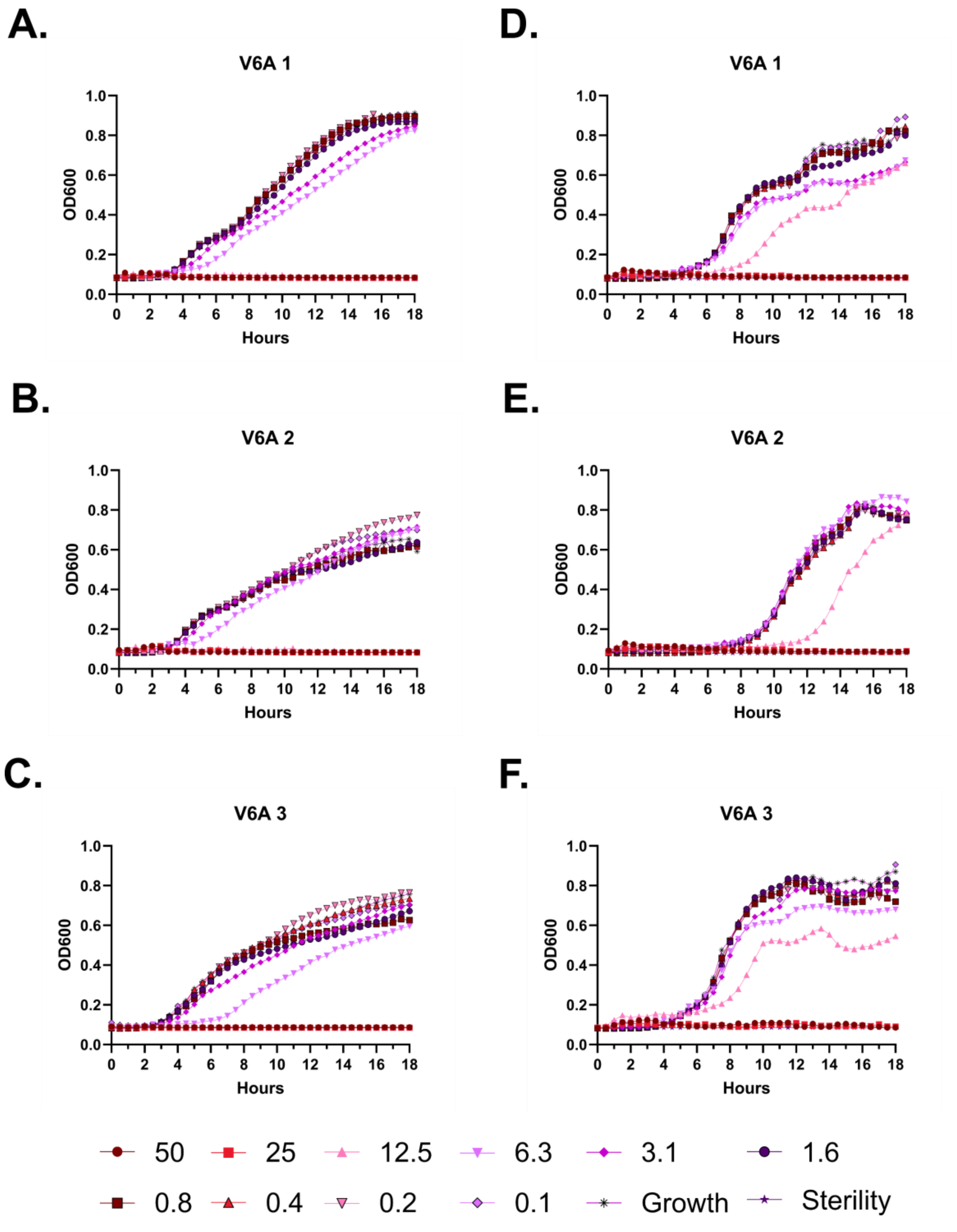
A.
V6A 1
D.
V6A 1
B.
V6A 2
E.
V6A 2
C.
V6A 3
F.
V6A 3
OD600
Hours
50
25
12.5
6.3
3.1
1.6
0.8
0.4
0.2
0.1
Growth
Sterility

**Figure S31 – Antimicrobial susceptibility testing with FF-14 V6A. A-C.** Growth of *E. coli* in the presence of the indicated peptide concentrations given in µM as monitored by OD600 at 37 °C over 18 hours. Each panel indicates an independent experiment. **D-F.** As in **A-C**, but with *P. aeruginosa.*

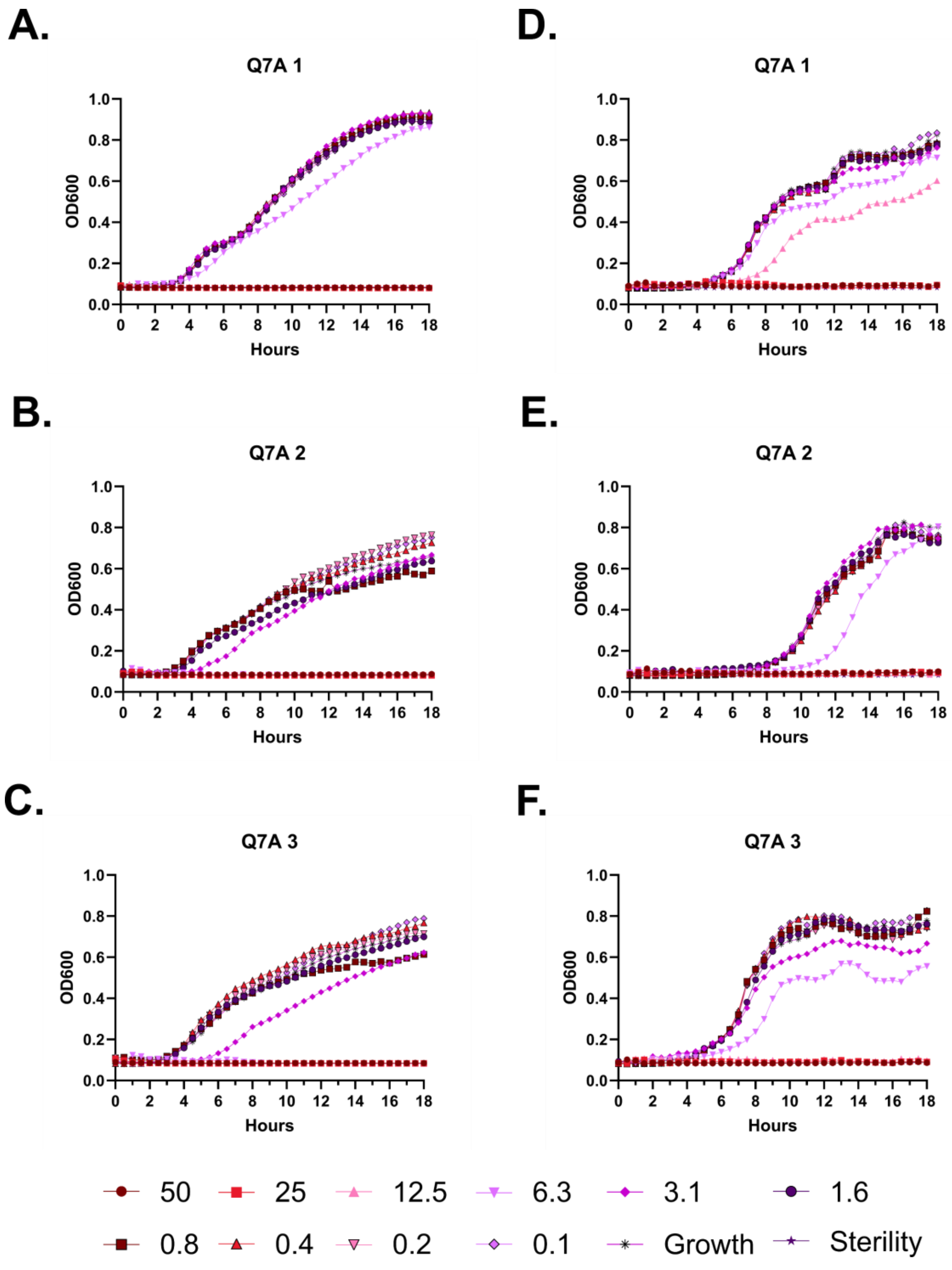

A.
Q7A 1
D.
Q7A 1
B.
Q7A 2
E.
Q7A 2
C.
Q7A 3
F.
Q7A 3
OD600
Hours
50
25
12.5
6.3
3.1
1.6
0.8
0.4
0.2
0.1
Growth
Sterility

**Figure S32 – Antimicrobial susceptibility testing with FF-14 Q7A. A-C.** Growth of *E. coli* in the presence of the indicated peptide concentrations given in µM as monitored by OD600 at 37 °C over 18 hours. Each panel indicates an independent experiment. **D-F.** As in **A-C**, but with *P. aeruginosa.*

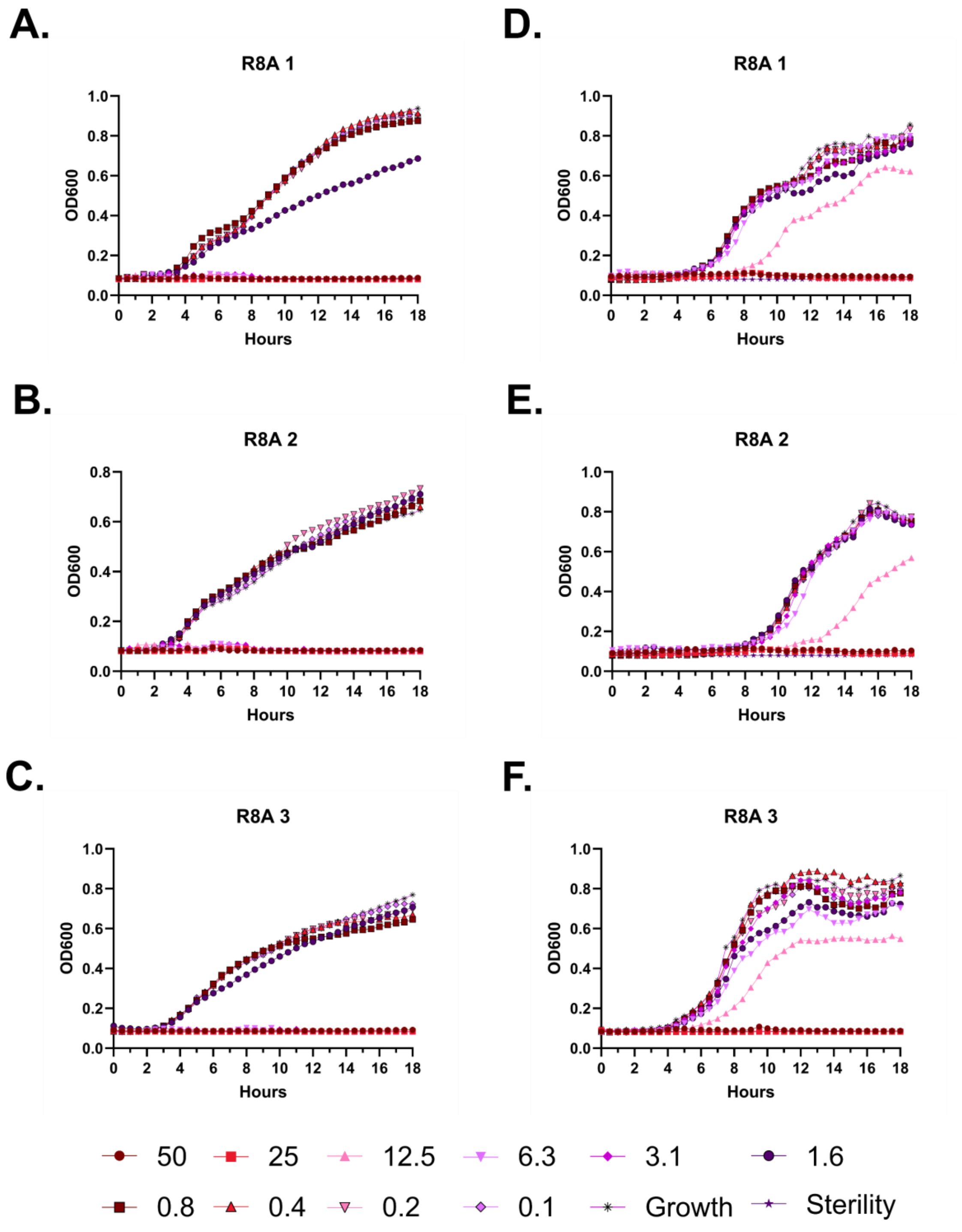
A.
R8A 1
D.
R8A 1
B.
R8A 2
E.
R8A 2
C.
R8A 3
F.
R8A 3
OD600
Hours
50
25
12.5
6.3
3.1
1.6
0.8
0.4
0.2
0.1
Growth
Sterility

**Figure S33 – Antimicrobial susceptibility testing with FF-14 R8A. A-C.** Growth of *E. coli* in the presence of the indicated peptide concentrations given in μM as monitored by OD600 at 37 °C over 18 hours. Each panel indicates an independent experiment. **D-F.** As in **A-C**, but with *P. aeruginosa.*

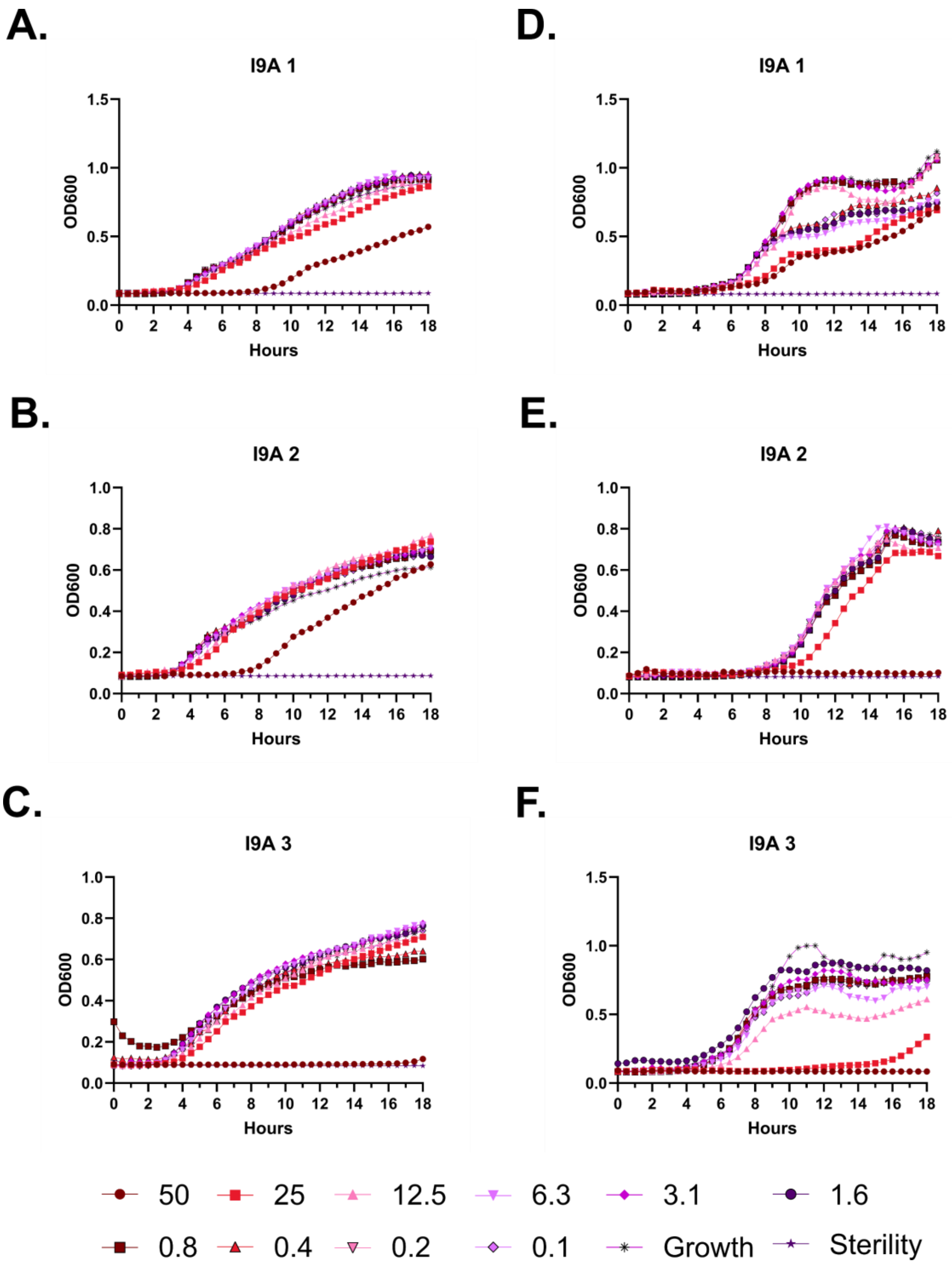
A.
I9A 1
OD600
Hours
D.
I9A 1
OD600
Hours
B.
I9A 2
OD600
Hours
E.
I9A 2
OD600
Hours
C.
I9A 3
OD600
Hours
F.
I9A 3
OD600
Hours
50
25
12.5
6.3
3.1
1.6
0.8
0.4
0.2
0.1
Growth
Sterility

**Figure S34 – Antimicrobial susceptibility testing with FF-14 I9A. A-C.** Growth of *E. coli* in the presence of the indicated peptide concentrations given in μM as monitored by OD600 at 37 °C over 18 hours. Each panel indicates an independent experiment. **D-F.** As in **A-C**, but with *P. aeruginosa.*

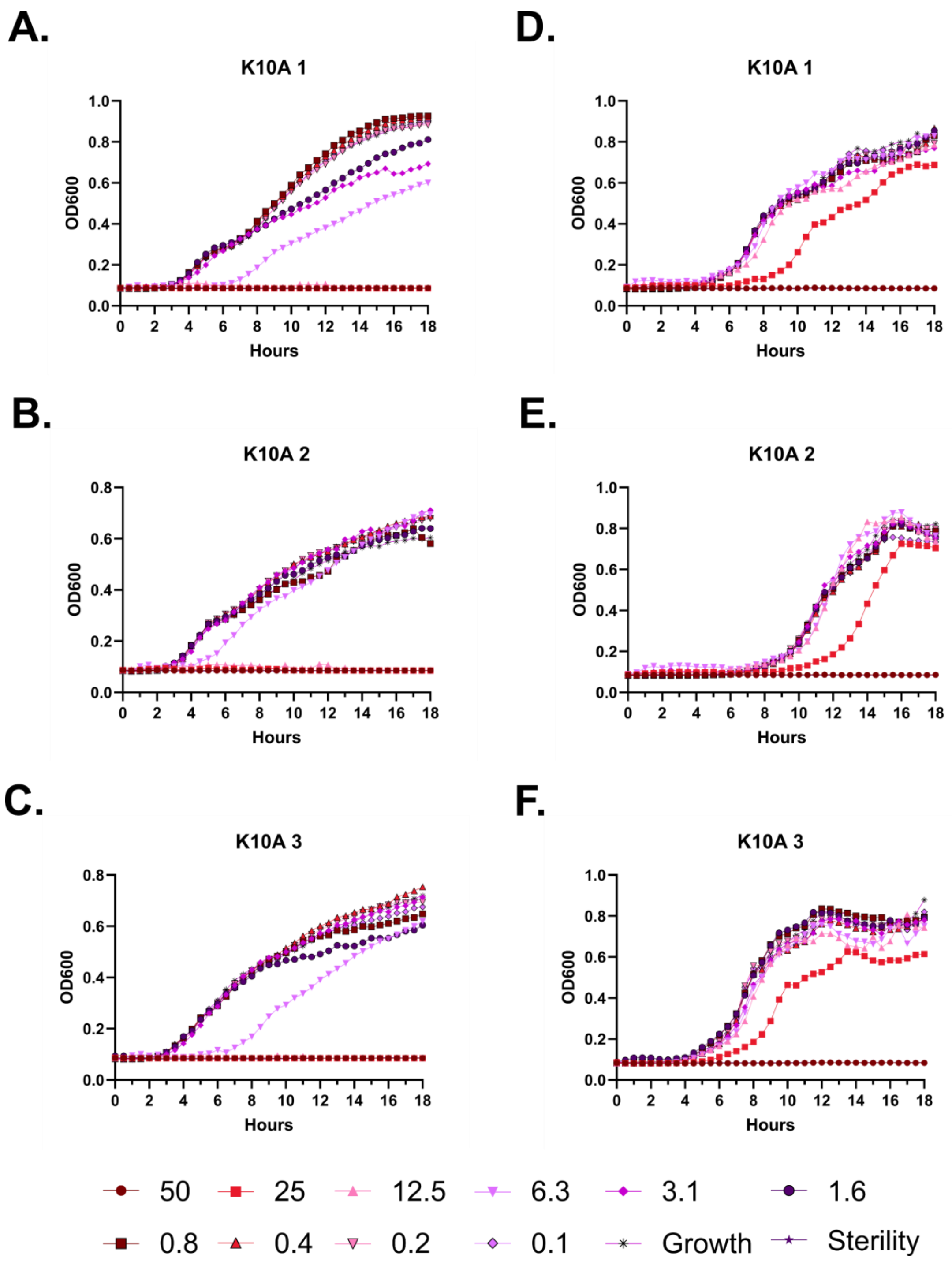
A.
K10A 1
B.
K10A 2
C.
K10A 3
D.
K10A 1
E.
K10A 2
F.
K10A 3
OD600
Hours
0 2 4 6 8 10 12 14 16 18
0.0 0.2 0.4 0.6 0.8 1.0
50
25
12.5
6.3
3.1
1.6
0.8
0.4
0.2
0.1
Growth
Sterility

**Figure S35 – Antimicrobial susceptibility testing with FF-14 K10A. A-C.** Growth of *E. coli* in the presence of the indicated peptide concentrations given in μM as monitored by OD600 at 37 °C over 18 hours. Each panel indicates an independent experiment. **D-F.** As in **A-C**, but with *P. aeruginosa.*

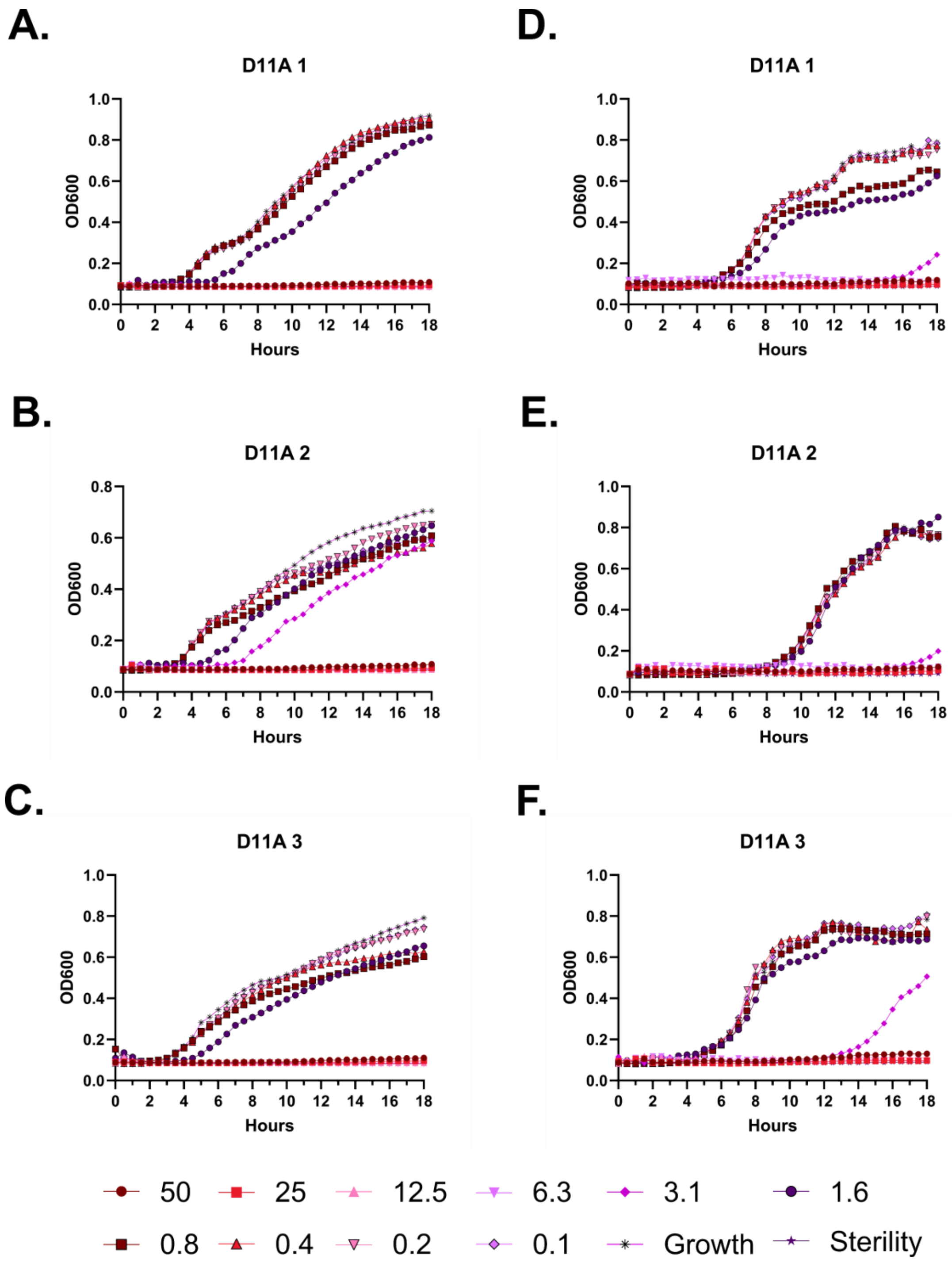
A.
D11A 1
B.
D11A 2
C.
D11A 3
D.
D11A 1
E.
D11A 2
F.
D11A 3
OD600
Hours
50
25
12.5
6.3
3.1
1.6
0.8
0.4
0.2
0.1
Growth
Sterility

**Figure S36 – Antimicrobial susceptibility testing with FF-14 D11A. A-C.** Growth of *E. coli* in the presence of the indicated peptide concentrations given in μM as monitored by OD600 at 37 °C over 18 hours. Each panel indicates an independent experiment. **D-F.** As in **A-C**, but with *P. aeruginosa.*

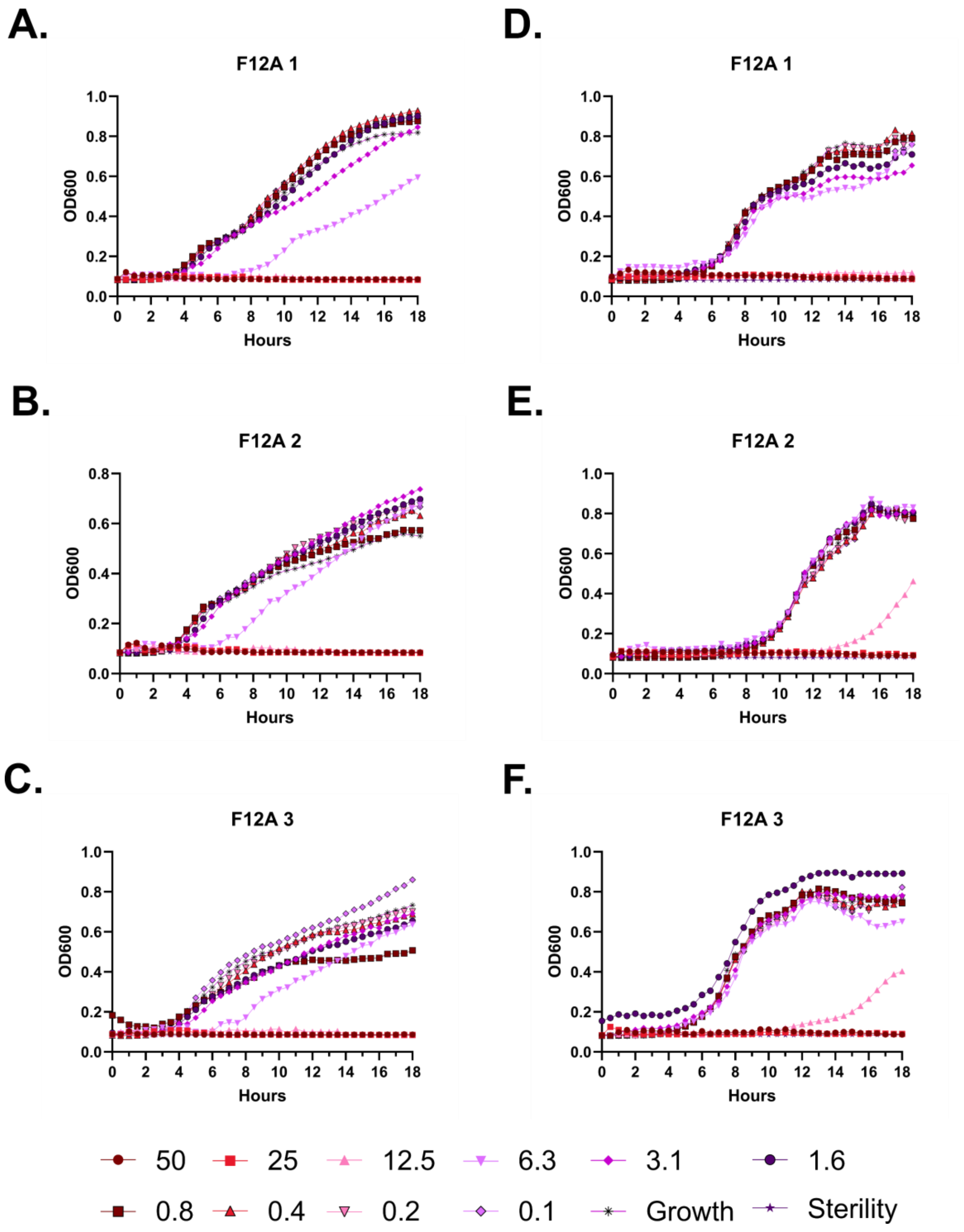

A.
F12A 1
D.
F12A 1
B.
F12A 2
E.
F12A 2
C.
F12A 3
F.
F12A 3
OD600
Hours
50
25
12.5
6.3
3.1
1.6
0.8
0.4
0.2
0.1
Growth
Sterility

**Figure S37 – Antimicrobial susceptibility testing with FF-14 F12A. A-C.** Growth of *E. coli* in the presence of the indicated peptide concentrations given in μM as monitored by OD600 at 37 °C over 18 hours. Each panel indicates an independent experiment. **D-F.** As in **A-C**, but with *P. aeruginosa.*

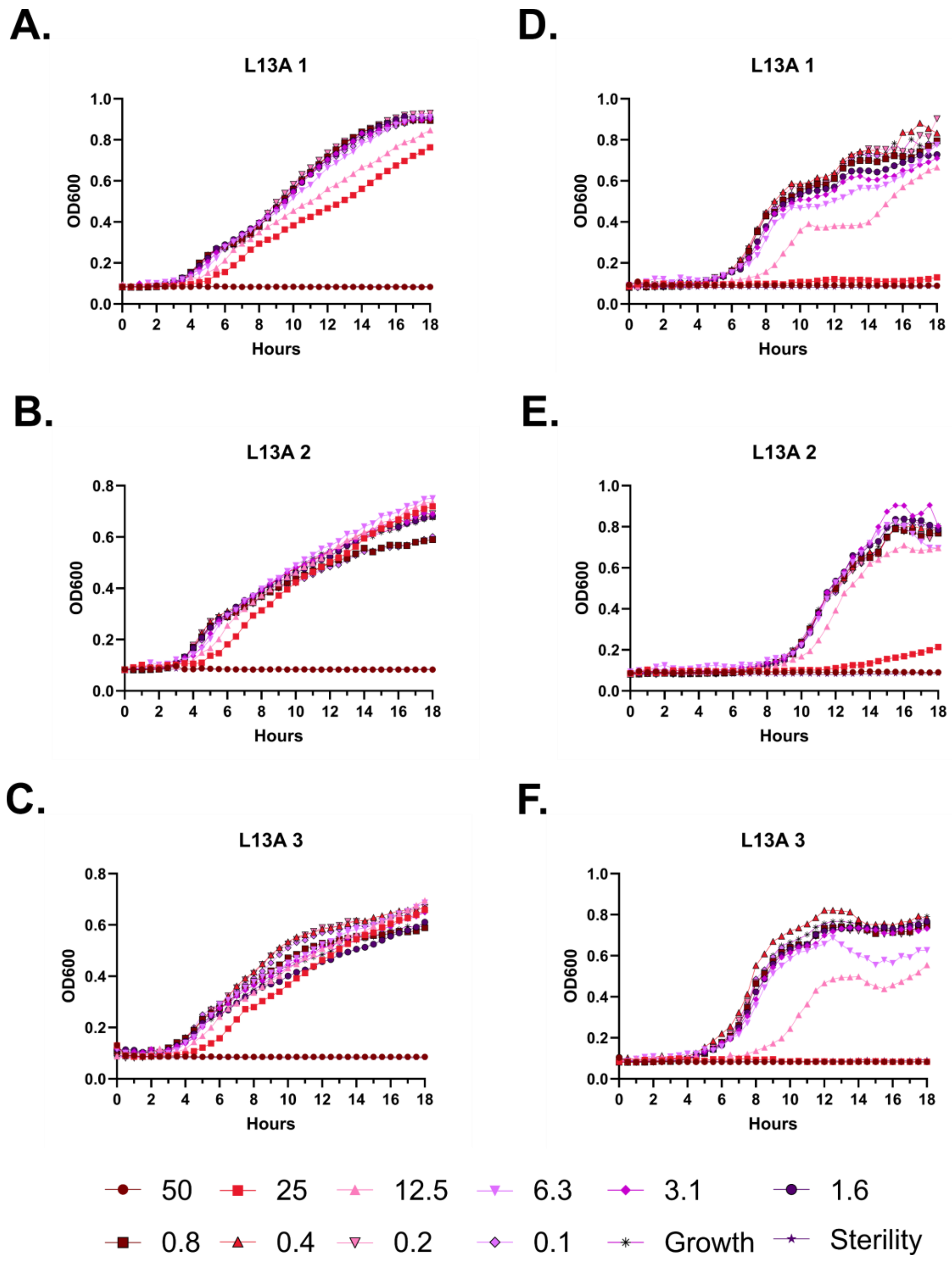
A.
L13A 1
OD600
Hours
D.
L13A 1
OD600
Hours
B.
L13A 2
OD600
Hours
E.
L13A 2
OD600
Hours
C.
L13A 3
OD600
Hours
F.
L13A 3
OD600
Hours
50
25
12.5
6.3
3.1
1.6
0.8
0.4
0.2
0.1
Growth
Sterility

**Figure S38 – Antimicrobial susceptibility testing with FF-14 L13A. A-C.** Growth of *E. coli* in the presence of the indicated peptide concentrations given in μM as monitored by OD600 at 37 °C over 18 hours. Each panel indicates an independent experiment. **D-F.** As in **A-C**, but with *P. aeruginosa.*

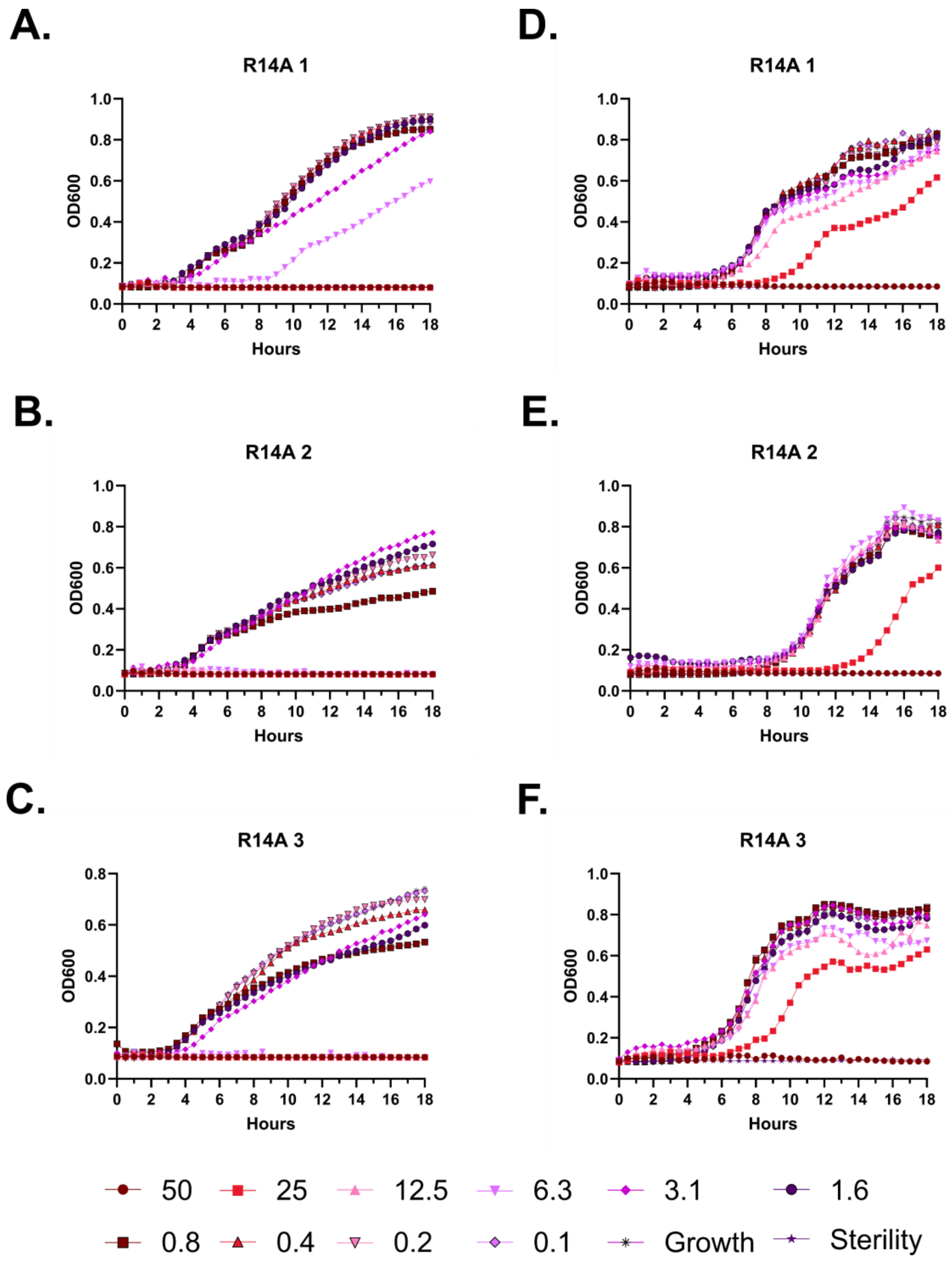
A.
R14A 1
D.
R14A 1
B.
R14A 2
E.
R14A 2
C.
R14A 3
F.
R14A 3
OD600
Hours
50
25
12.5
6.3
3.1
1.6
0.8
0.4
0.2
0.1
Growth
Sterility

**Figure S39 – Antimicrobial susceptibility testing with FF-14 R14A. A-C.** Growth of *E. coli* in the presence of the indicated peptide concentrations given in μM as monitored by OD600 at 37 °C over 18 hours. Each panel indicates an independent experiment. **D-F.** As in **A-C**, but with *P. aeruginosa.*

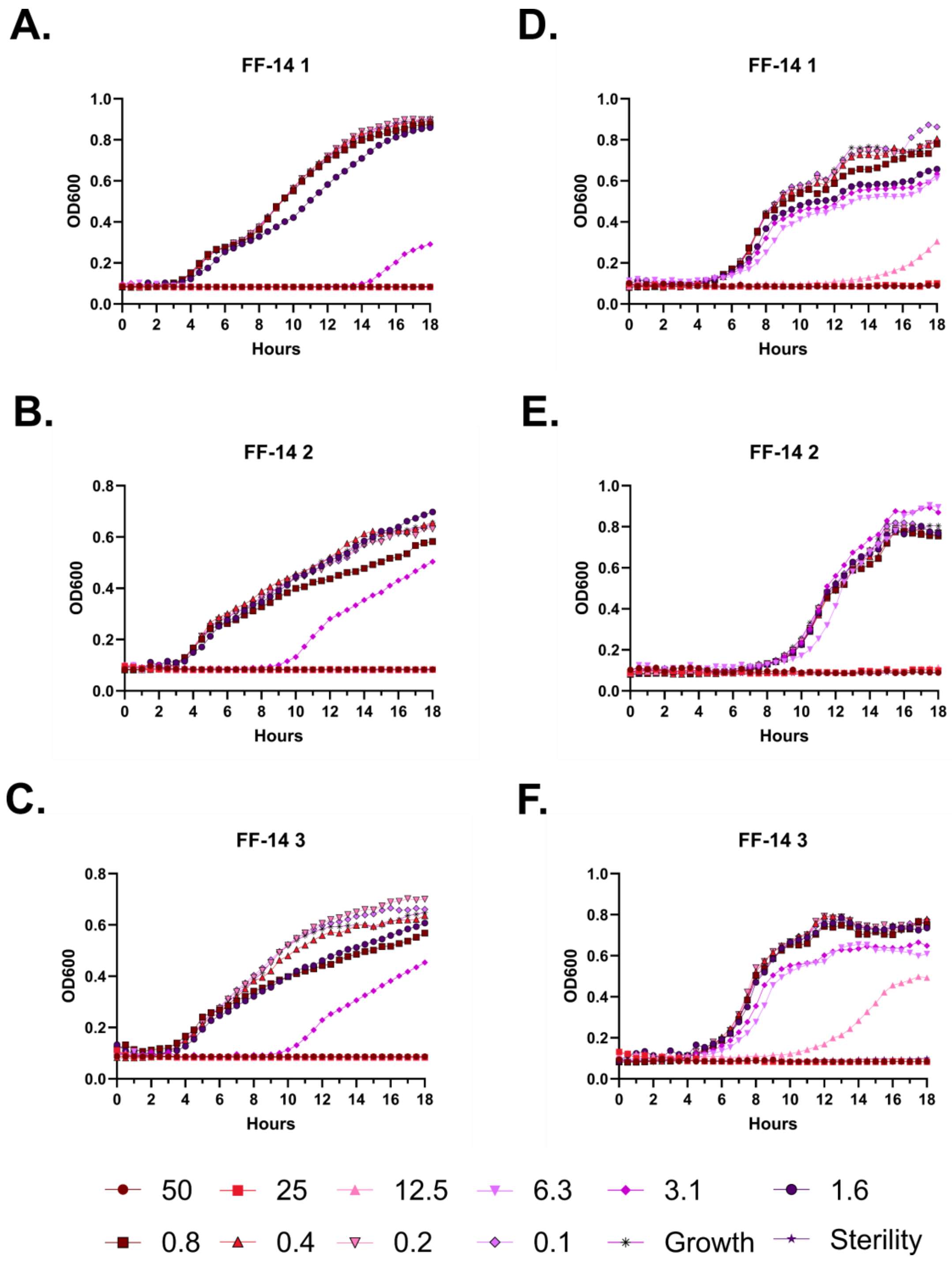
A.
FF-14 1
B.
FF-14 2
C.
FF-14 3
D.
FF-14 1
E.
FF-14 2
F.
FF-14 3
OD600
Hours
50
25
12.5
6.3
3.1
1.6
0.8
0.4
0.2
0.1
Growth
Sterility

**Figure S40 – Antimicrobial susceptibility testing with FF-14. A-C.** Growth of *E. coli* in the presence of the indicated peptide concentrations given in μM as monitored by OD600 at 37 °C over 18 hours. Each panel indicates an independent experiment. **D-F.** As in **A-C**, but with *P. aeruginosa.*

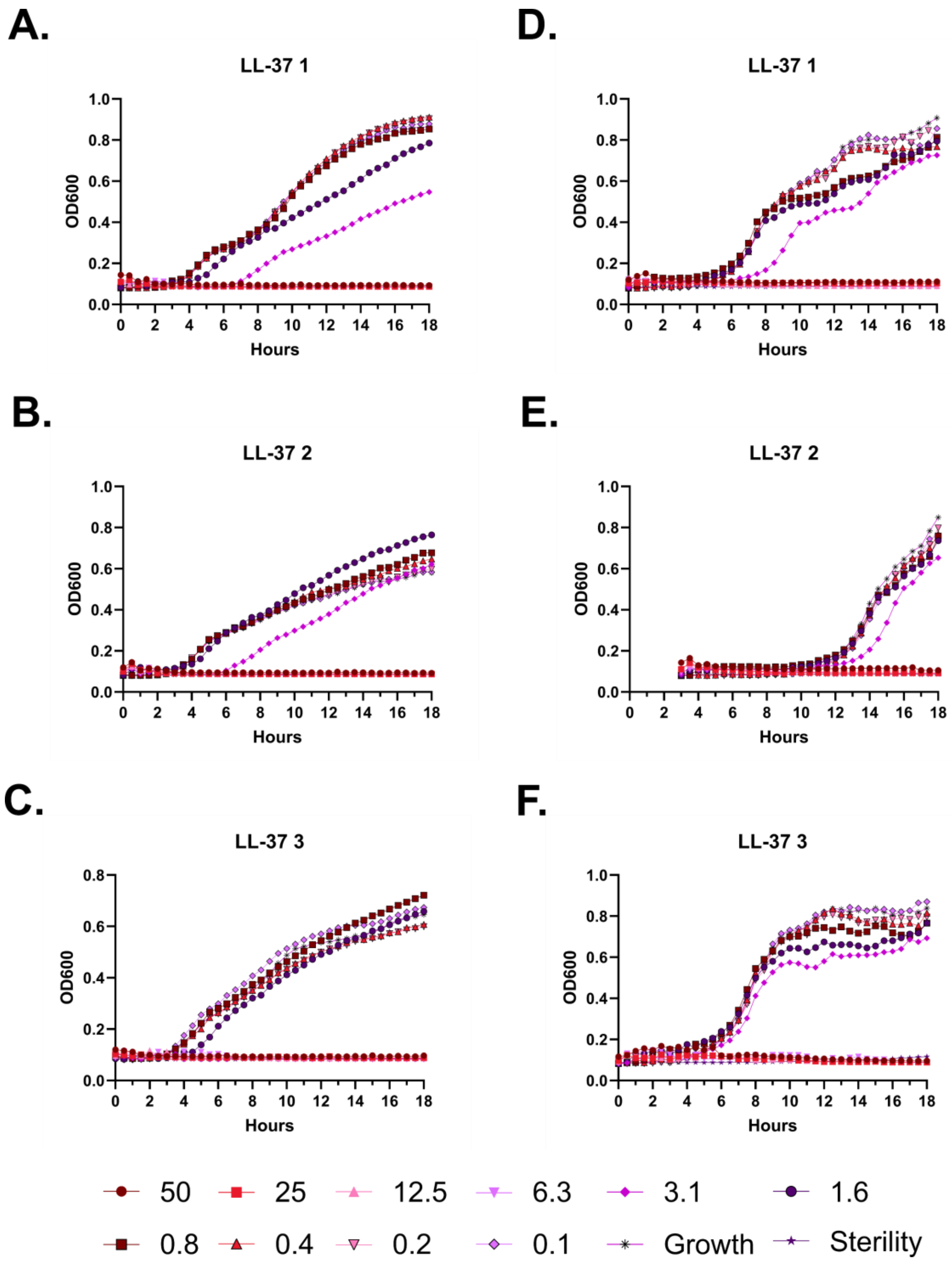
A.
LL-37 1
OD600
Hours
B.
LL-37 2
C.
LL-37 3
D.
LL-37 1
E.
LL-37 2
F.
LL-37 3
50
25
12.5
6.3
3.1
1.6
0.8
0.4
0.2
0.1
Growth
Sterility

**Figure S41 – Antimicrobial susceptibility testing with LL-37. A-C.** Growth of *E. coli* in the presence of the indicated peptide concentrations given in μM as monitored by OD600 at 37 °C over 18 hours. Each panel indicates an independent experiment. **D-F.** As in **A-C**, but with *P. aeruginosa.*

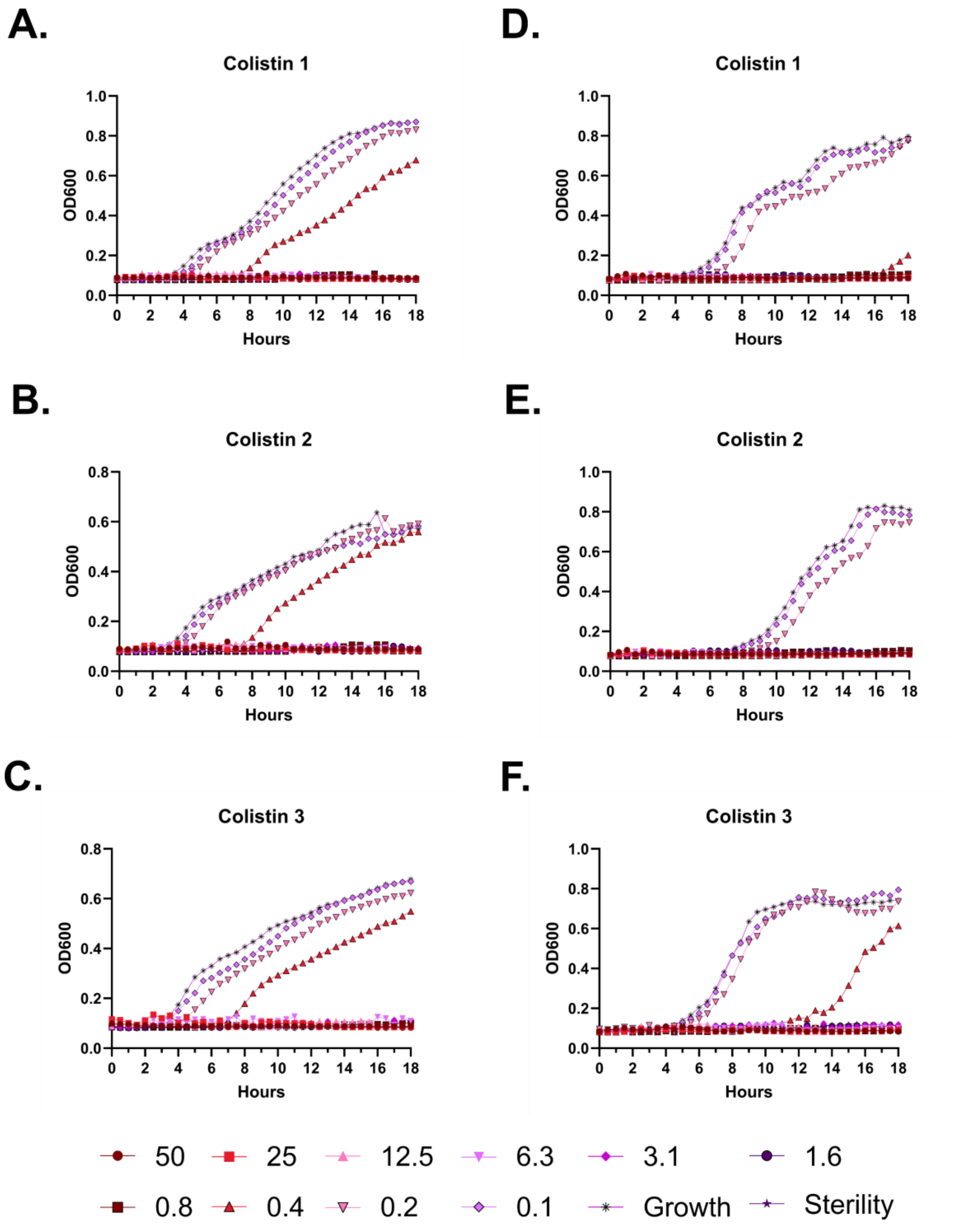

A.
Colistin 1
OD600
Hours
D.
Colistin 1
OD600
Hours
B.
Colistin 2
OD600
Hours
E.
Colistin 2
OD600
Hours
C.
Colistin 3
OD600
Hours
F.
Colistin 3
OD600
Hours
50
25
12.5
6.3
3.1
1.6
0.8
0.4
0.2
0.1
Growth
Sterility

**Figure S42 – Antimicrobial susceptibility testing with colistin. A-C.** Growth of *E. coli* in the presence of the indicated peptide concentrations given in μM as monitored by OD600 at 37 °C over 18 hours. Each panel indicates an independent experiment. **D-F.** As in **A-C**, but with *P. aeruginosa.*

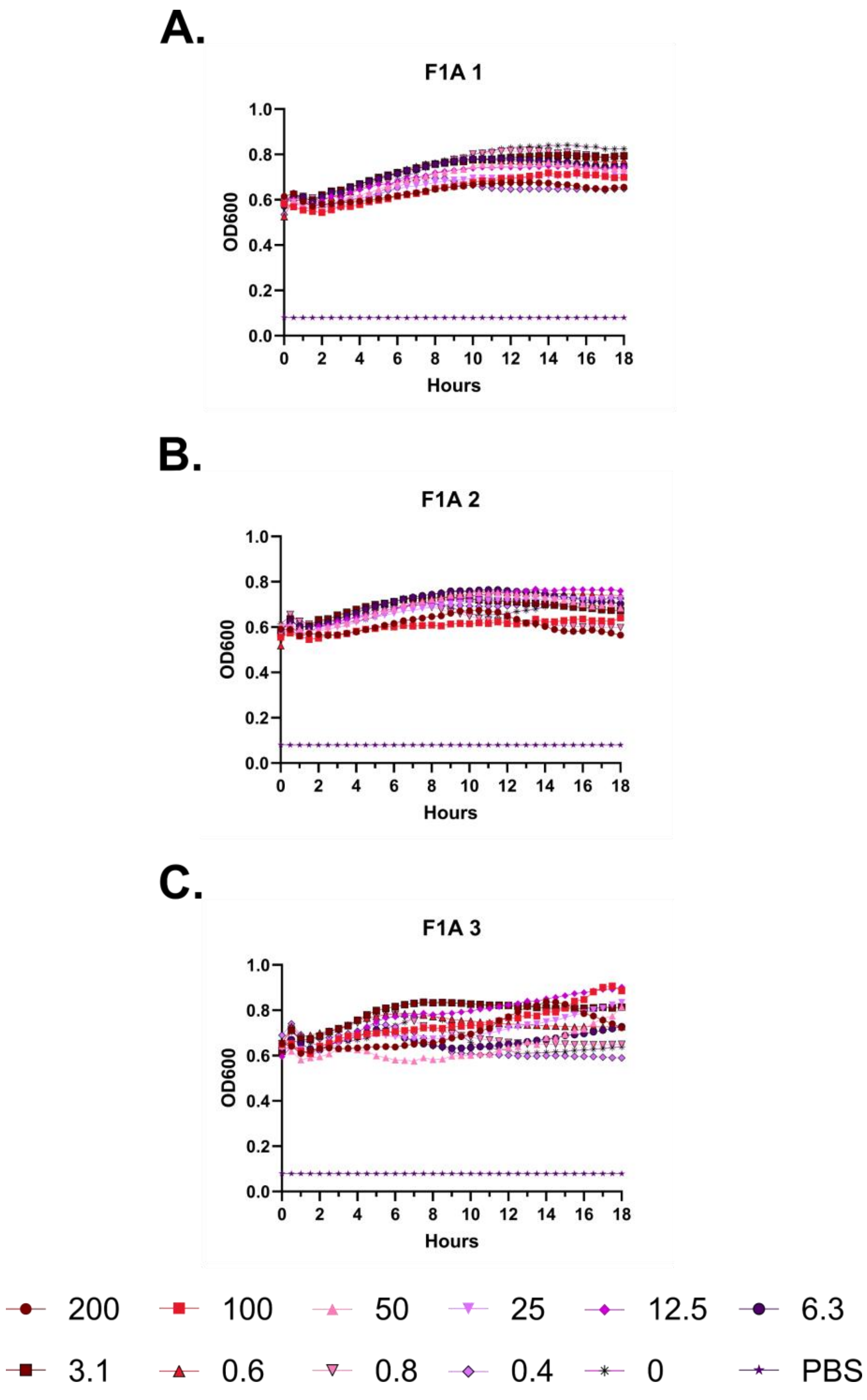

A.
F1A 1
OD600
Hours
B.
F1A 2
OD600
Hours
C.
F1A 3
OD600
Hours
200
100
50
25
12.5
6.3
3.1
0.6
0.8
0.4
0
PBS

**Figure S43 – Hemolysis over time with FF-14 F1A. A-C.** Each of three independent experiments showing trends in OD600 over time on incubation of defibrinated sheep blood with the indicated concentrations of peptide in μM at 37 °C. Decreasing OD600 indicates increasing hemolysis.

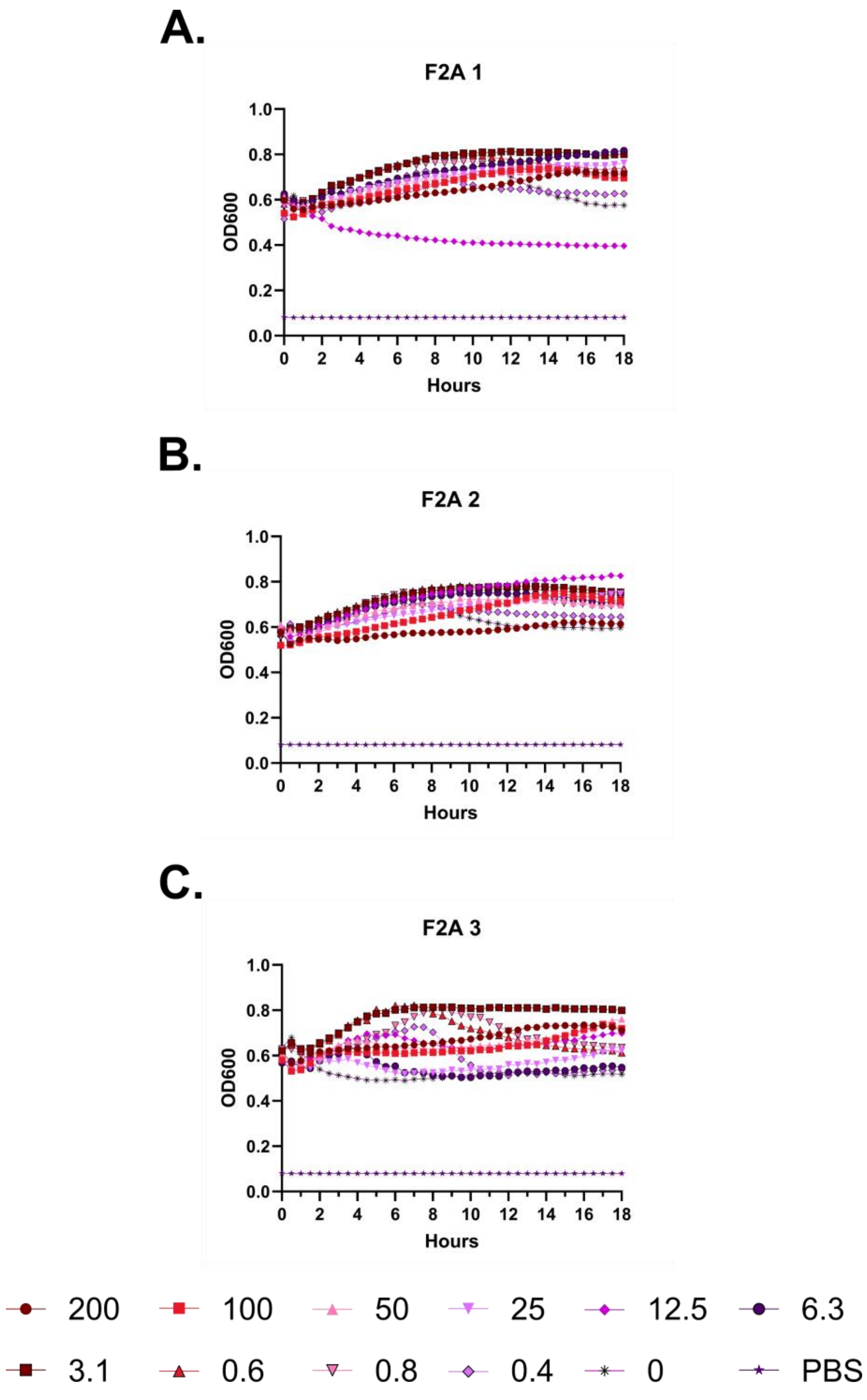
A.
F2A 1
OD600
Hours
B.
F2A 2
OD600
Hours
C.
F2A 3
OD600
Hours
200
100
50
25
12.5
6.3
3.1
0.6
0.8
0.4
0
PBS

**Figure S44 – Hemolysis over time with FF-14 F2A. A-C.** Each of three independent experiments showing trends in OD600 over time on incubation of defibrinated sheep blood with the indicated concentrations of peptide in μM at 37 °C. Decreasing OD600 indicates increasing hemolysis.

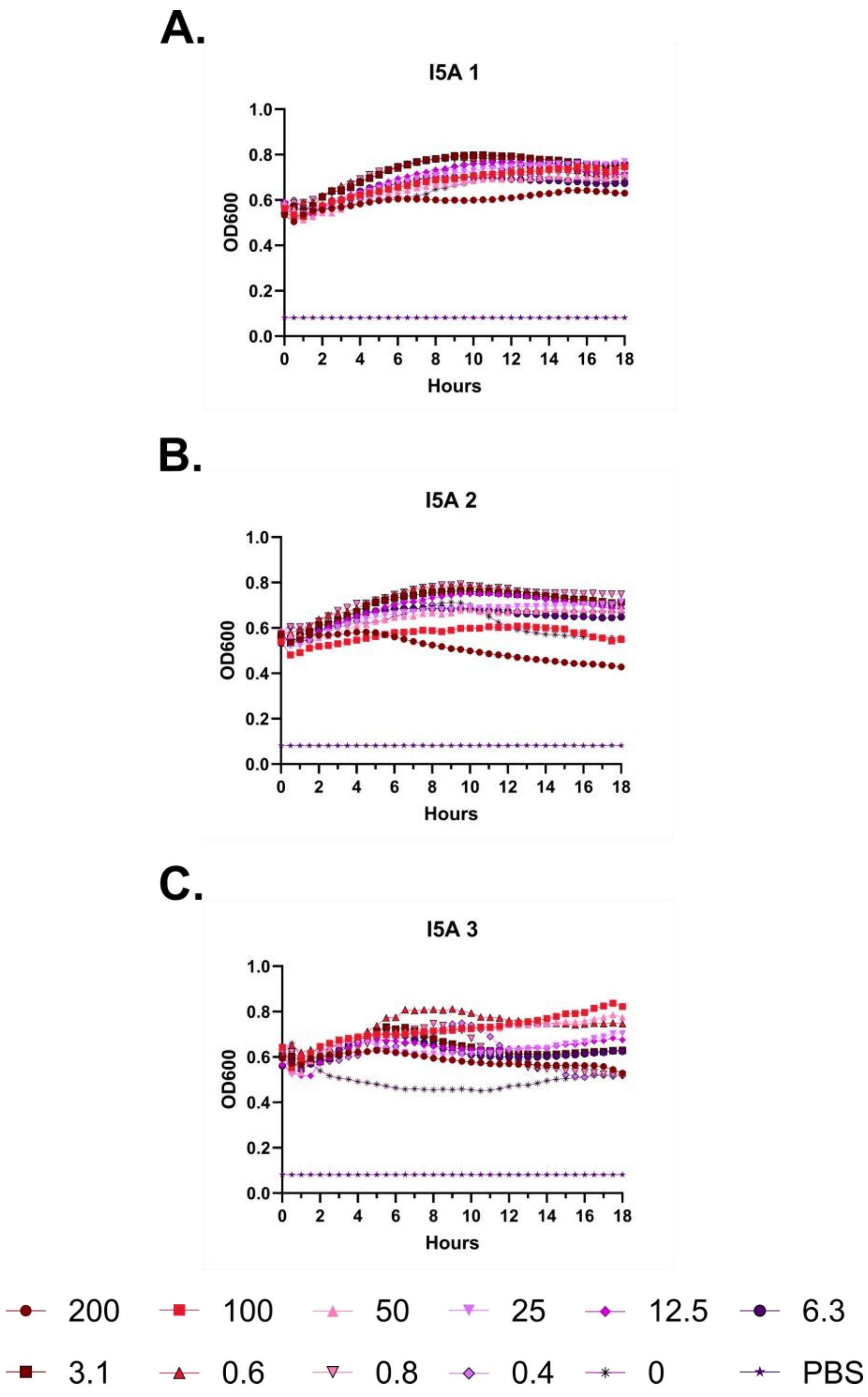
A.
I5A 1
B.
I5A 2
C.
I5A 3
OD600
Hours
1.0
0.8
0.6
0.4
0.2
0.0
0
2
4
6
8
10
12
14
16
18
200
100
50
25
12.5
6.3
3.1
0.6
0.8
0.4
0
PBS

**Figure S45 – Hemolysis over time with FF-14 I5A. A-C.** Each of three independent experiments showing trends in OD600 over time on incubation of defibrinated sheep blood with the indicated concentrations of peptide in μM at 37 °C. Decreasing OD600 indicates increasing hemolysis.

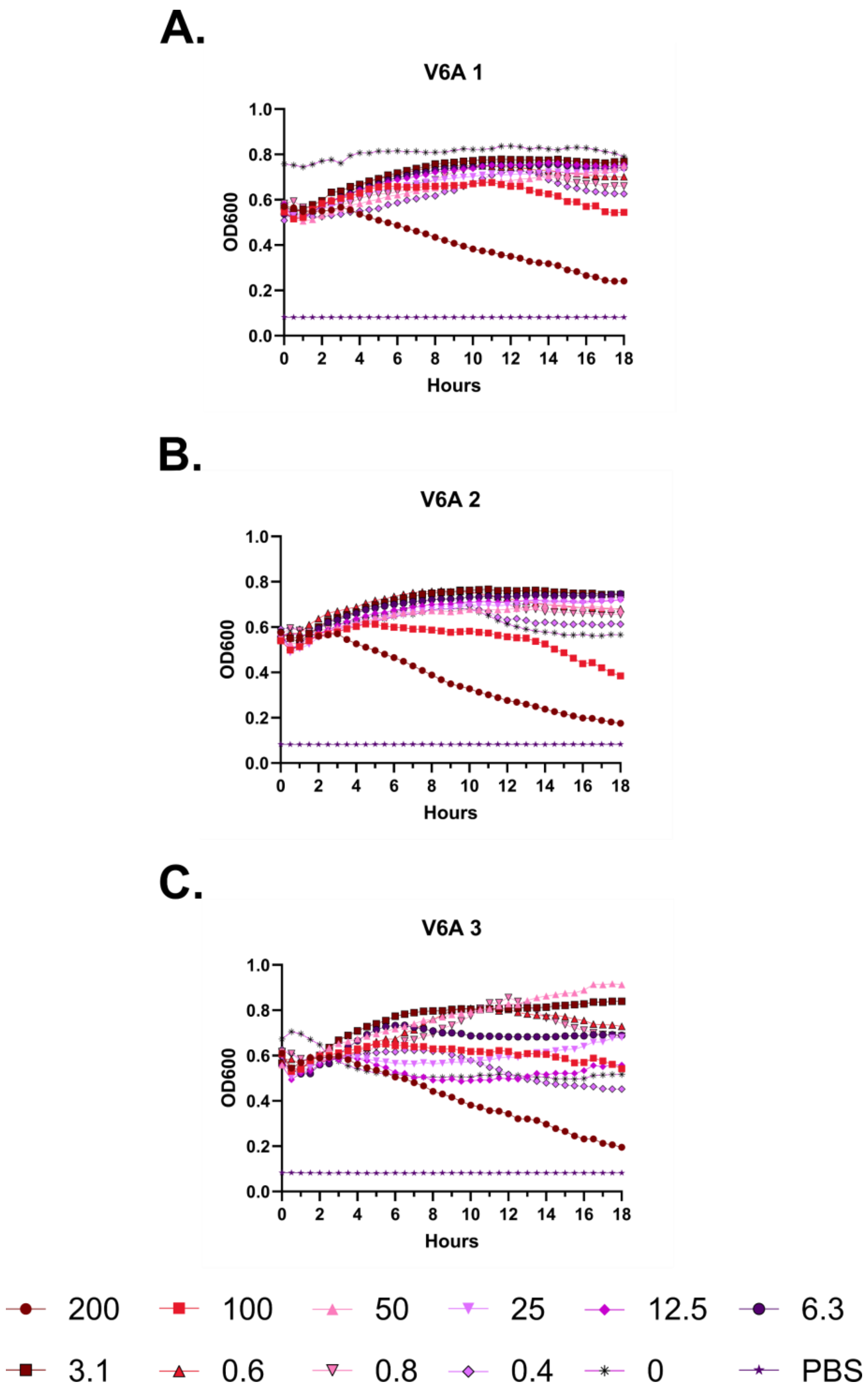
A.
V6A 1
OD600
Hours
B.
V6A 2
OD600
Hours
C.
V6A 3
OD600
Hours
200
100
50
25
12.5
6.3
3.1
0.6
0.8
0.4
0
PBS

**Figure S46 – Hemolysis over time with FF-14 V6A. A-C.** Each of three independent experiments showing trends in OD600 over time on incubation of defibrinated sheep blood with the indicated concentrations of peptide in μM at 37 °C. Decreasing OD600 indicates increasing hemolysis.

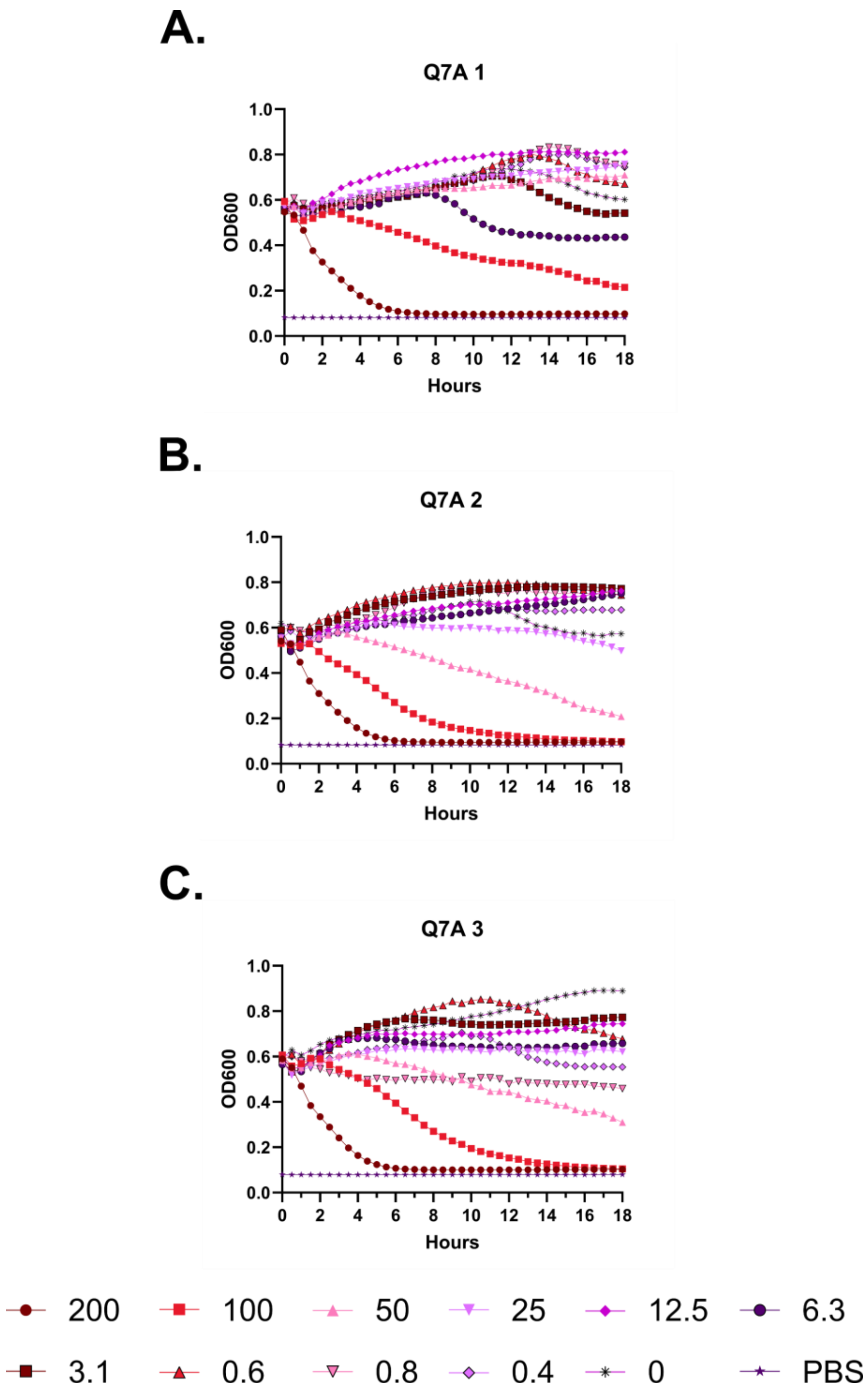
A.
Q7A 1
B.
Q7A 2
C.
Q7A 3
OD600
Hours
200
100
50
25
12.5
6.3
3.1
0.6
0.8
0.4
0
PBS

**Figure S47 – Hemolysis over time with FF-14 Q7A. A-C.** Each of three independent experiments showing trends in OD600 over time on incubation of defibrinated sheep blood with the indicated concentrations of peptide in μM at 37 °C. Decreasing OD600 indicates increasing hemolysis.

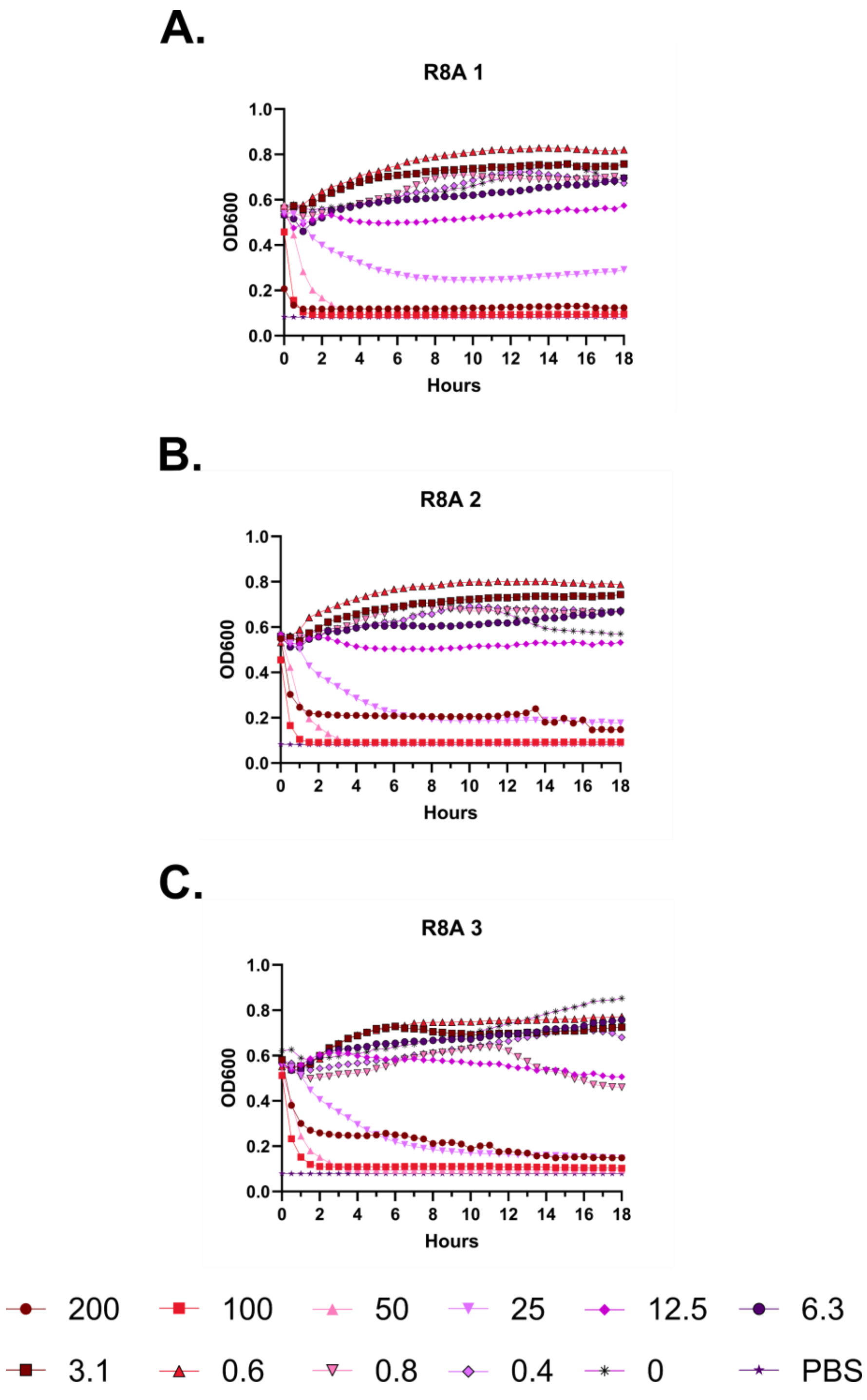
A.
R8A 1
OD600
Hours
B.
R8A 2
OD600
Hours
C.
R8A 3
OD600
Hours
200
100
50
25
12.5
6.3
3.1
0.6
0.8
0.4
0
PBS

**Figure S48 – Hemolysis over time with FF-14 R8A. A-C.** Each of three independent experiments showing trends in OD600 over time on incubation of defibrinated sheep blood with the indicated concentrations of peptide in μM at 37 °C. Decreasing OD600 indicates increasing hemolysis.

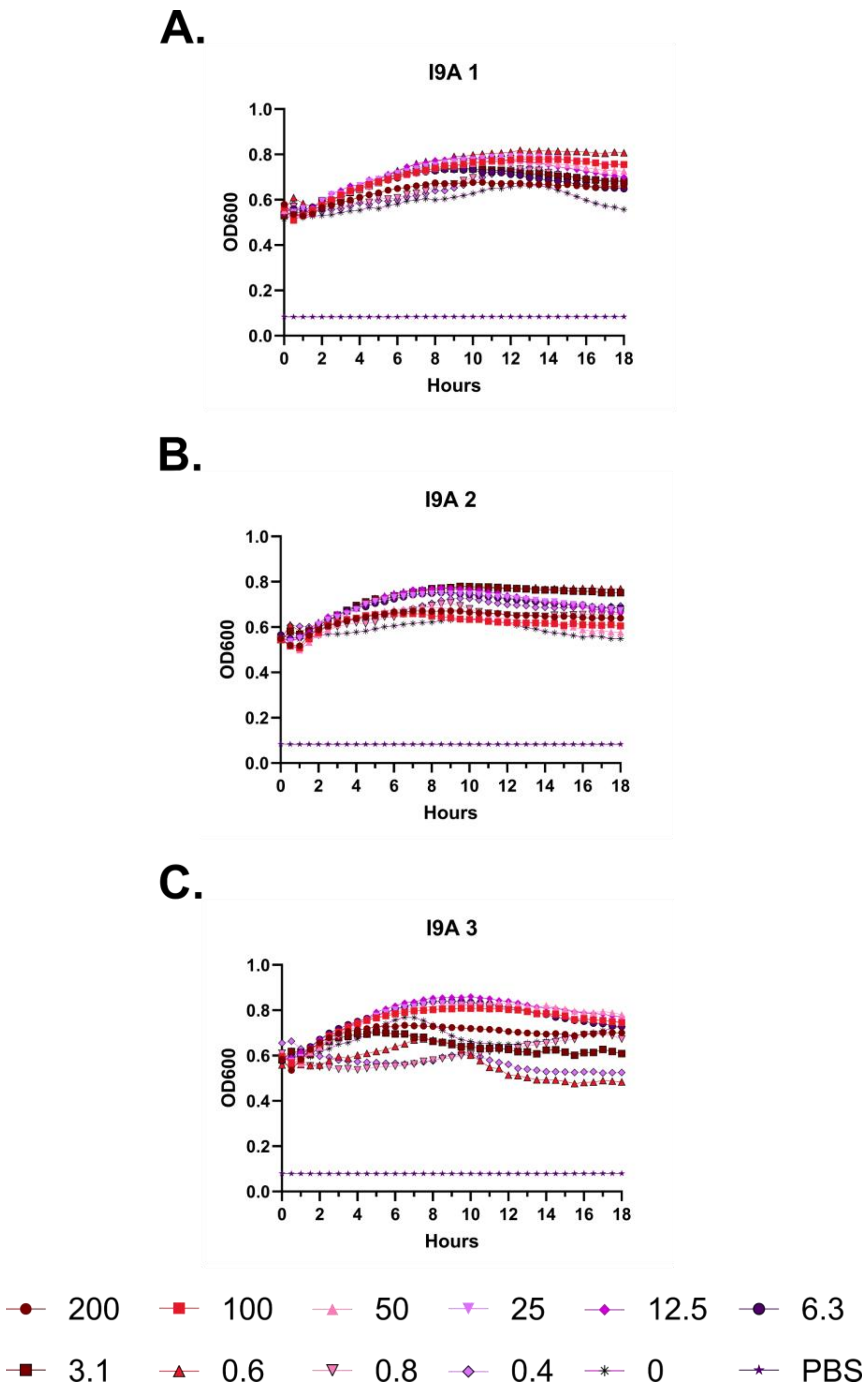
A.
I9A 1
OD600
Hours
B.
I9A 2
OD600
Hours
C.
I9A 3
OD600
Hours
200
100
50
25
12.5
6.3
3.1
0.6
0.8
0.4
0
PBS

**Figure S49 – Hemolysis over time with FF-14 I9A. A-C.** Each of three independent experiments showing trends in OD600 over time on incubation of defibrinated sheep blood with the indicated concentrations of peptide in μM at 37 °C. Decreasing OD600 indicates increasing hemolysis.

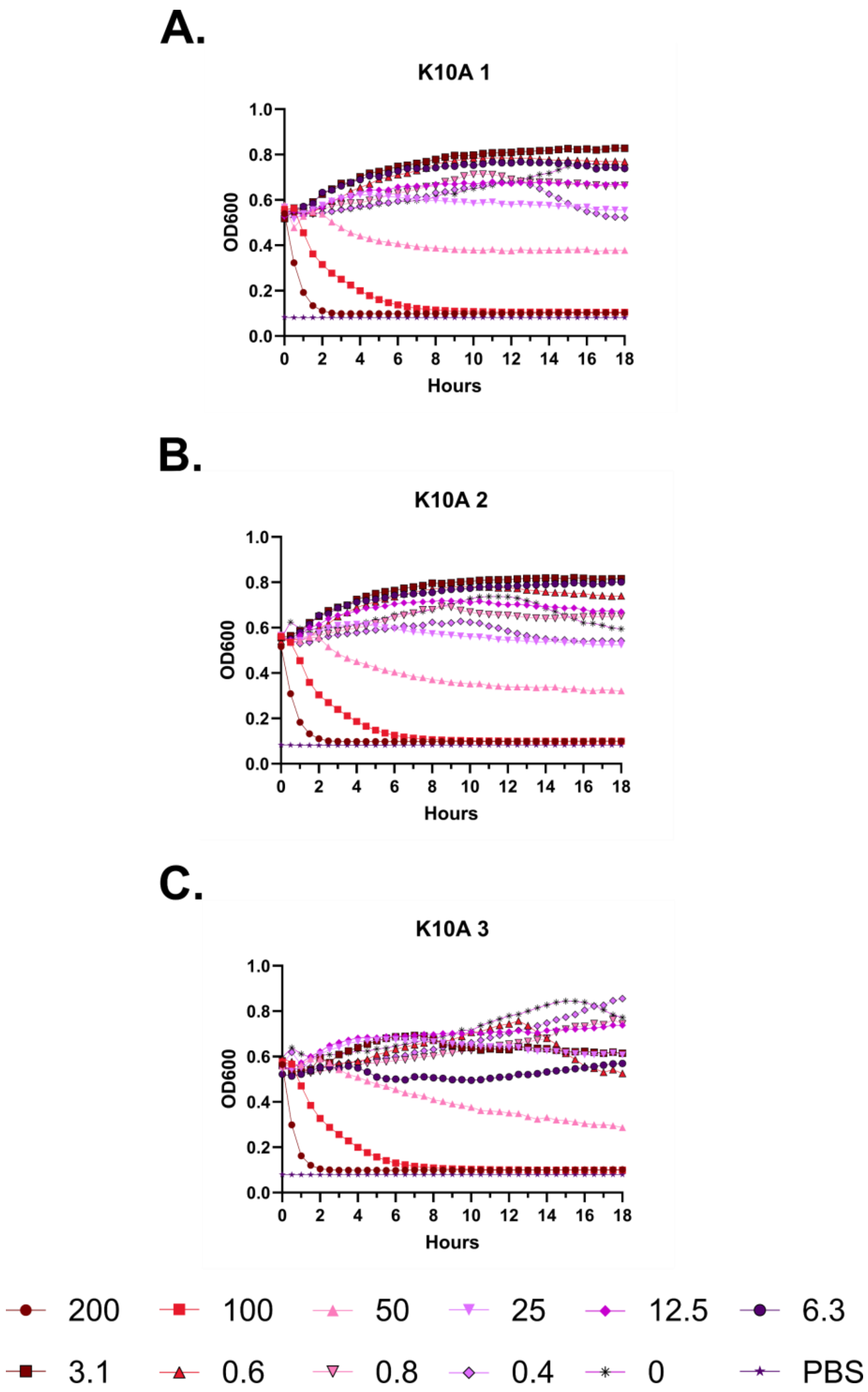
A.
K10A 1
OD600
Hours
B.
K10A 2
OD600
Hours
C.
K10A 3
OD600
Hours
200
100
50
25
12.5
6.3
3.1
0.6
0.8
0.4
0
PBS

**Figure S50 – Hemolysis over time with FF-14 K10A. A-C.** Each of three independent experiments showing trends in OD600 over time on incubation of defibrinated sheep blood with the indicated concentrations of peptide in μM at 37 °C. Decreasing OD600 indicates increasing hemolysis.

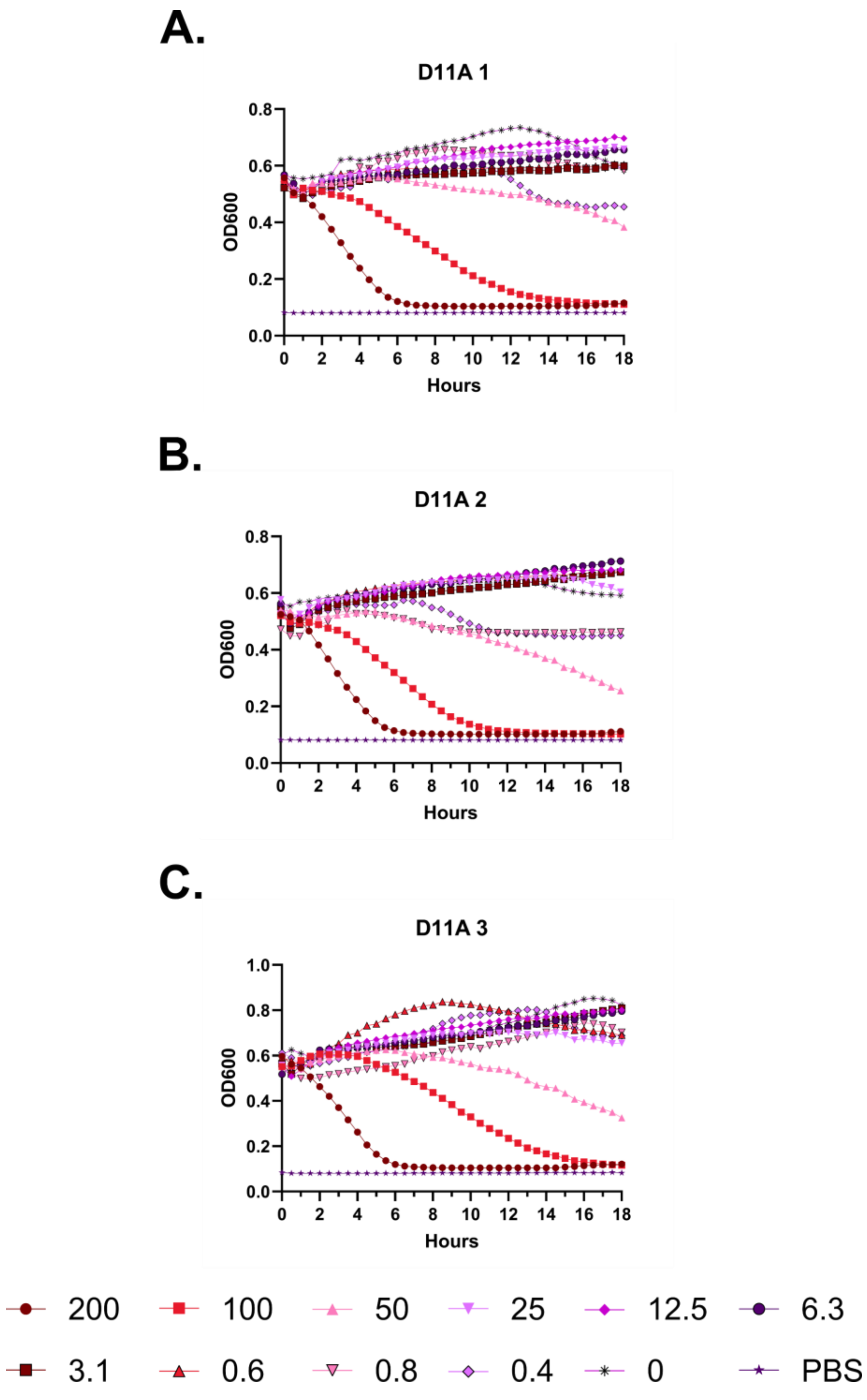
A.
D11A 1
OD600
Hours
B.
D11A 2
OD600
Hours
C.
D11A 3
OD600
Hours
200
100
50
25
12.5
6.3
3.1
0.6
0.8
0.4
0
PBS

**Figure S51 – Hemolysis over time with FF-14 D11A. A-C.** Each of three independent experiments showing trends in OD600 over time on incubation of defibrinated sheep blood with the indicated concentrations of peptide in μM at 37 °C. Decreasing OD600 indicates increasing hemolysis.

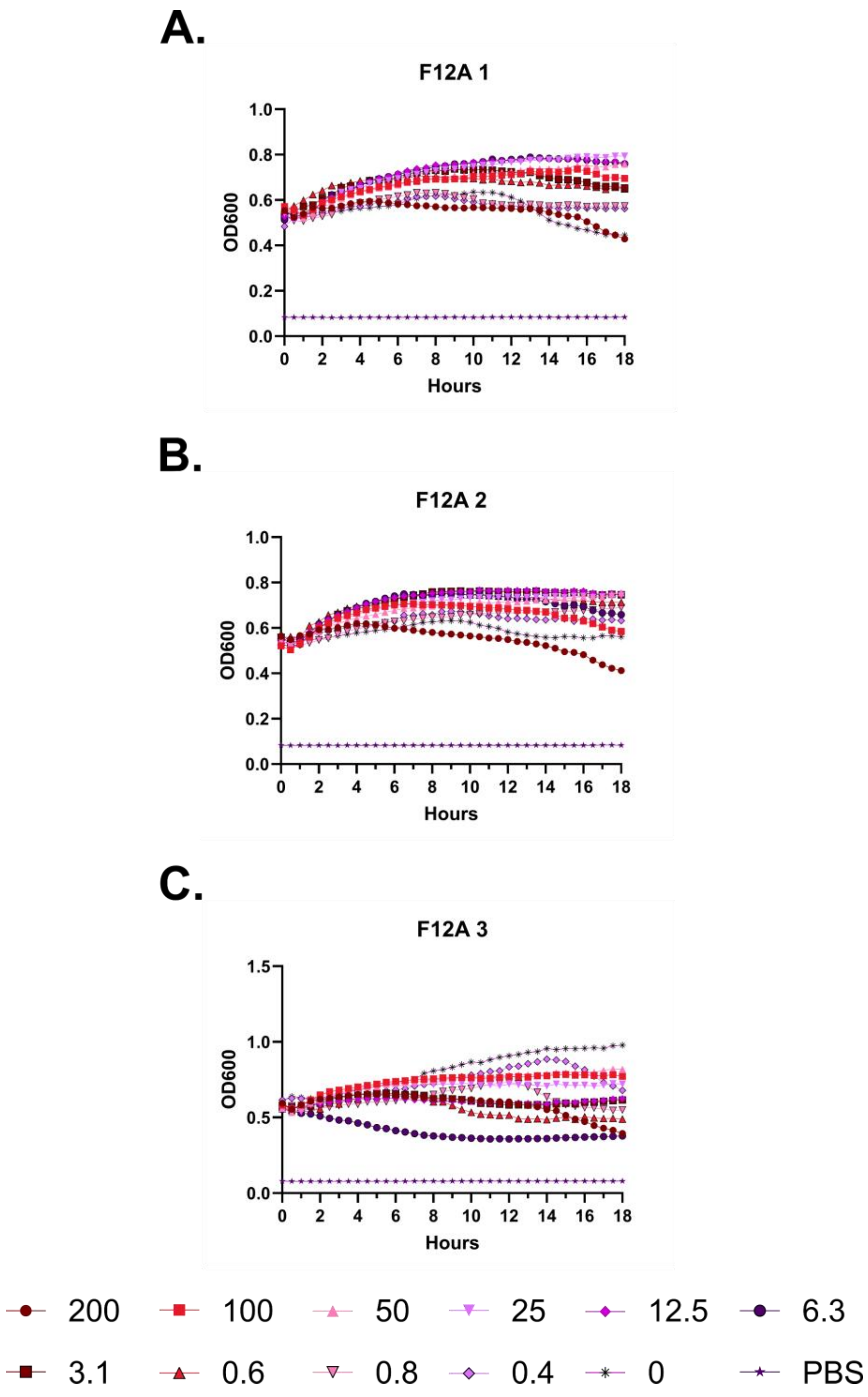

A.
F12A 1
OD600
Hours
B.
F12A 2
OD600
Hours
C.
F12A 3
OD600
Hours
200
100
50
25
12.5
6.3
3.1
0.6
0.8
0.4
0
PBS

**Figure S52 – Hemolysis over time with FF-14 F12A. A-C.** Each of three independent experiments showing trends in OD600 over time on incubation of defibrinated sheep blood with the indicated concentrations of peptide in μM at 37 °C. Decreasing OD600 indicates increasing hemolysis.

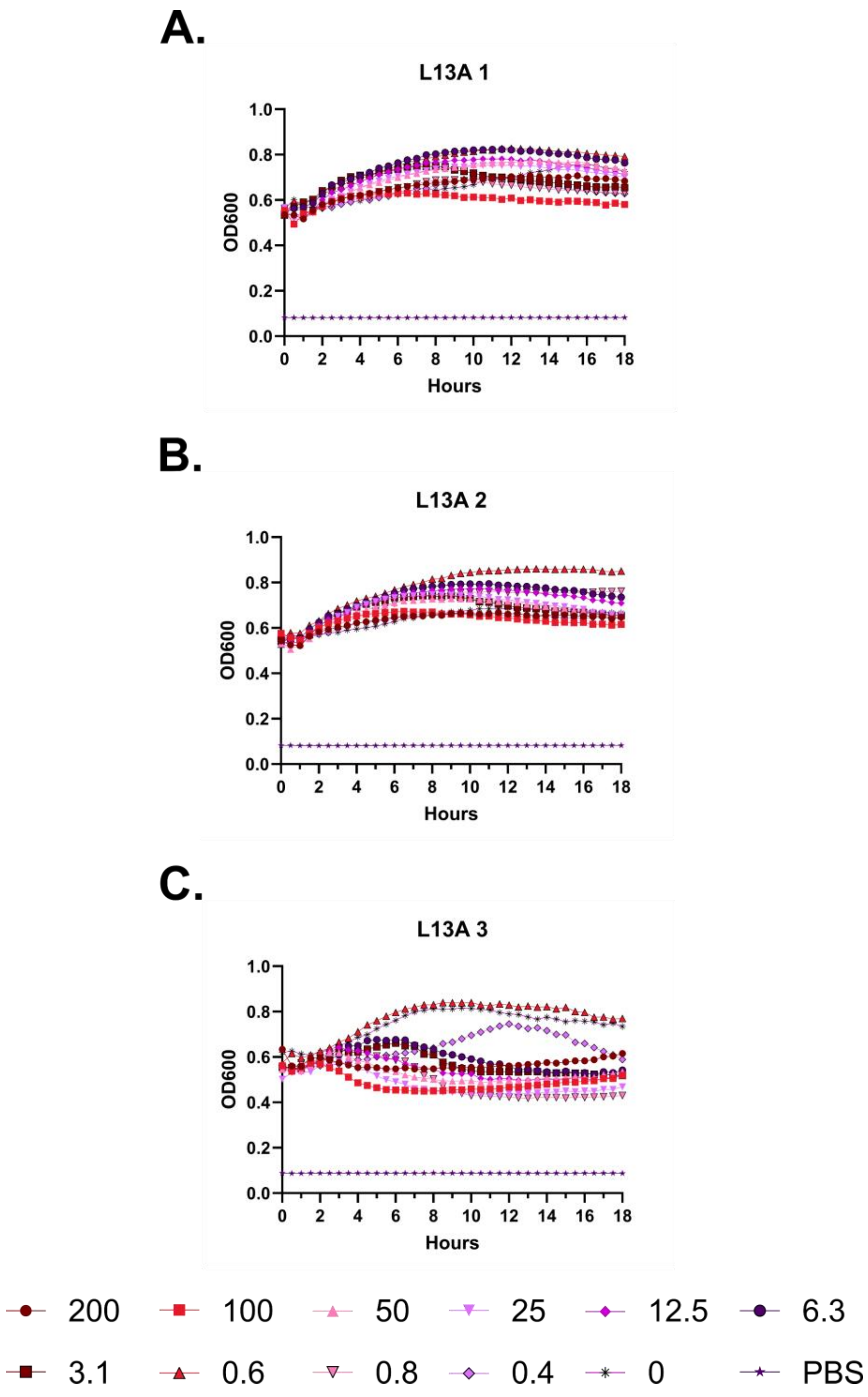
A.
L13A 1
OD600
Hours
B.
L13A 2
OD600
Hours
C.
L13A 3
OD600
Hours
200
100
50
25
12.5
6.3
3.1
0.6
0.8
0.4
0
PBS

**Figure S53 – Hemolysis over time with FF-14 L13A. A-C.** Each of three independent experiments showing trends in OD600 over time on incubation of defibrinated sheep blood with the indicated concentrations of peptide in μM at 37 °C. Decreasing OD600 indicates increasing hemolysis.

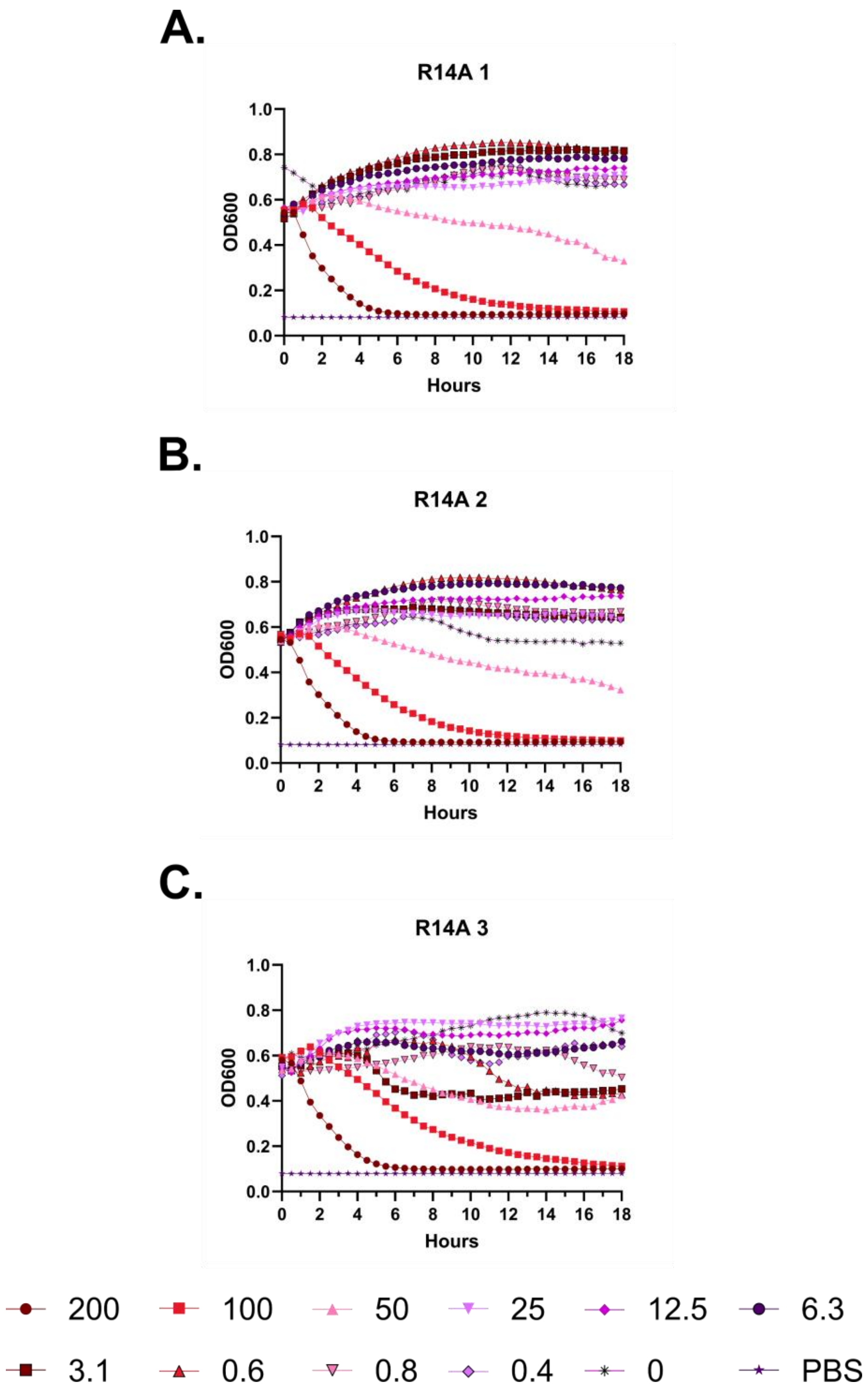
A.
R14A 1
OD600
Hours
B.
R14A 2
OD600
Hours
C.
R14A 3
OD600
Hours
200
100
50
25
12.5
6.3
3.1
0.6
0.8
0.4
0
PBS

**Figure S54 – Hemolysis over time with FF-14 R14A. A-C.** Each of three independent experiments showing trends in OD600 over time on incubation of defibrinated sheep blood with the indicated concentrations of peptide in μM at 37 °C. Decreasing OD600 indicates increasing hemolysis.

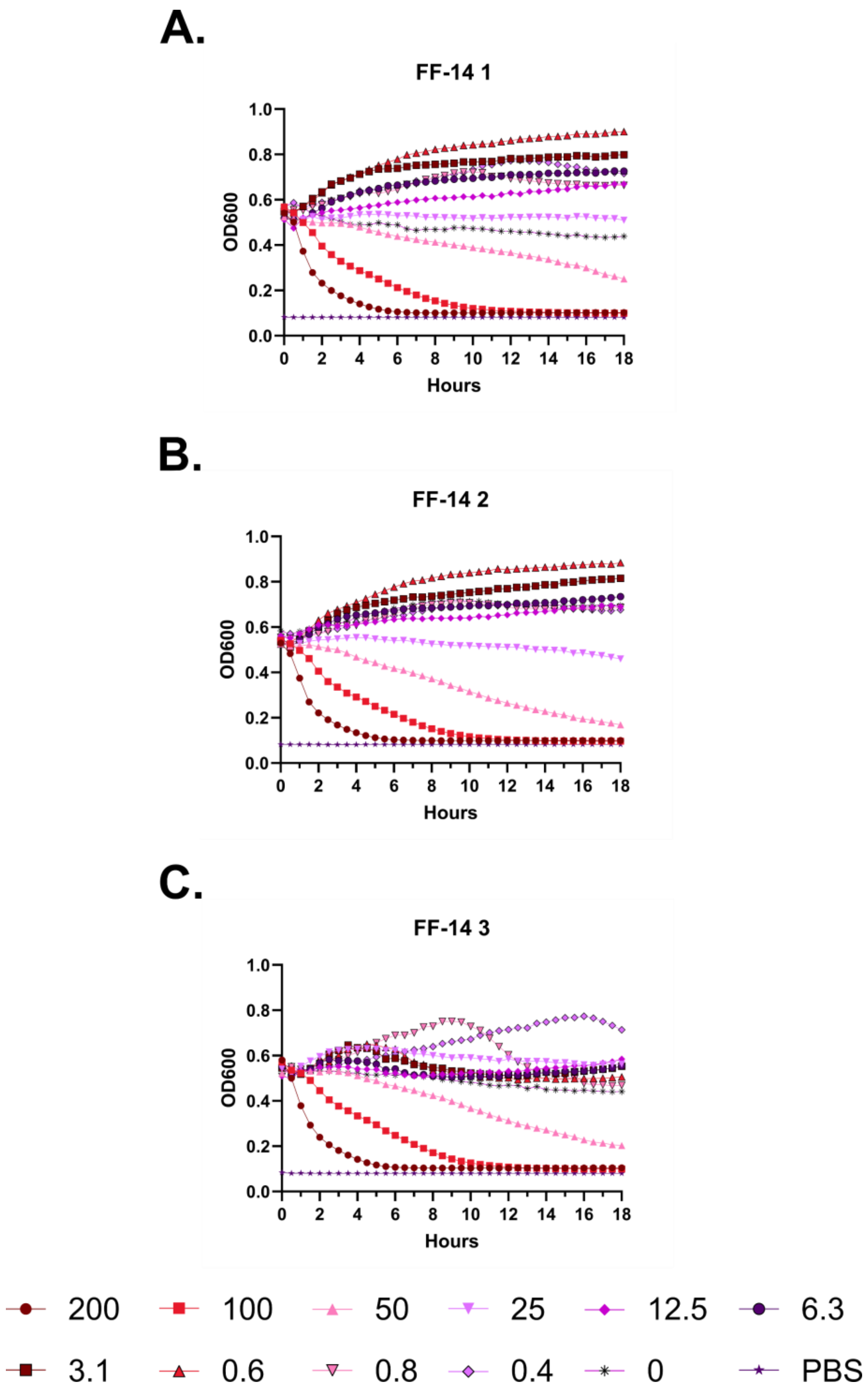
A.
FF-14 1
B.
FF-14 2
C.
FF-14 3
OD600
Hours
0.0
0.2
0.4
0.6
0.8
1.0
0 2 4 6 8 10 12 14 16 18
200
100
50
25
12.5
6.3
3.1
0.6
0.8
0.4
0
PBS

**Figure S55 – Hemolysis over time with FF-14. A-C.** Each of three independent experiments showing trends in OD600 over time on incubation of defibrinated sheep blood with the indicated concentrations of peptide in μM at 37 °C. Decreasing OD600 indicates increasing hemolysis.

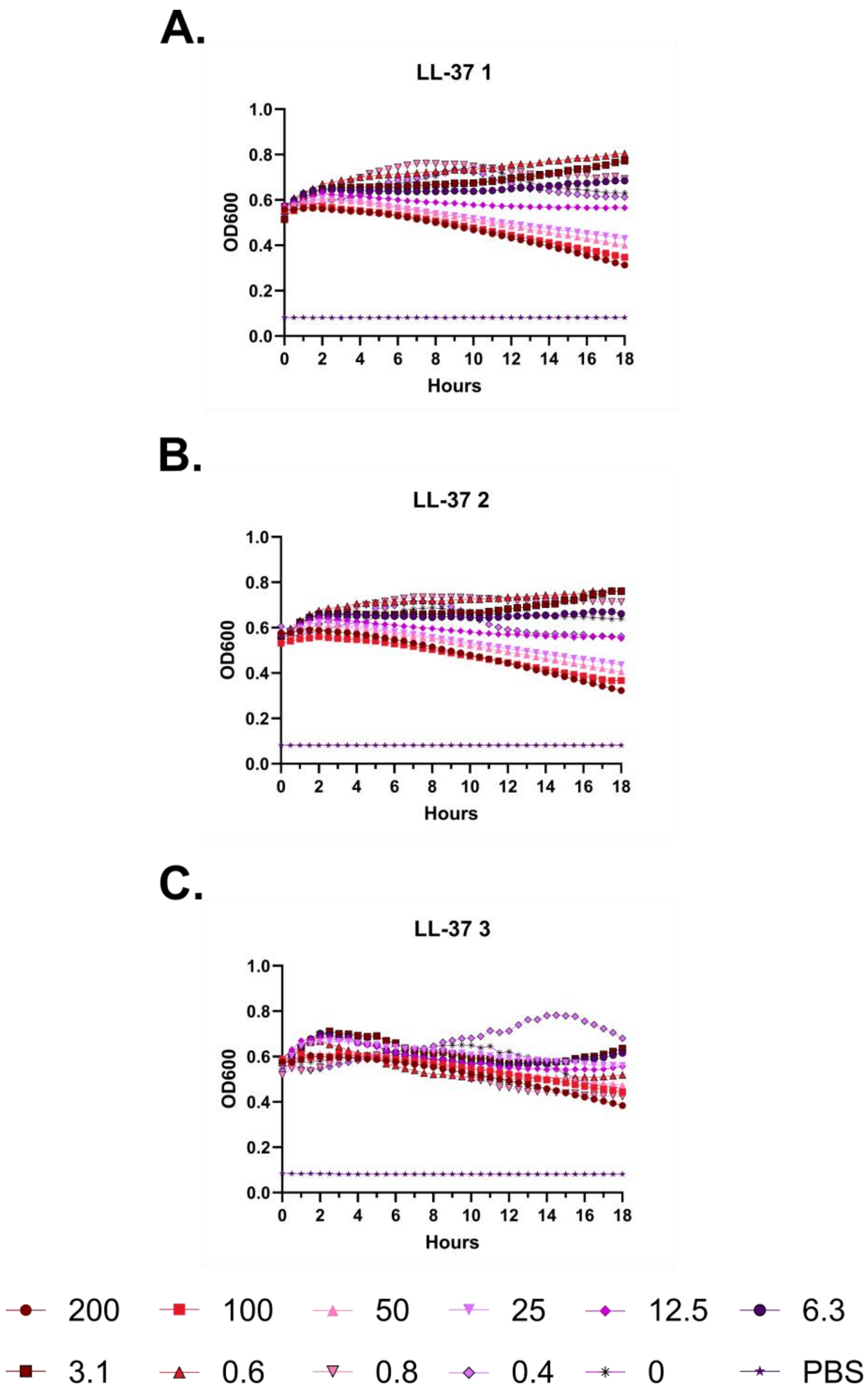
A.
LL-37 1
OD600
Hours
B.
LL-37 2
OD600
Hours
C.
LL-37 3
OD600
Hours
200
100
50
25
12.5
6.3
3.1
0.6
0.8
0.4
0
PBS

**Figure S56 – Hemolysis over time with LL-37. A-C.** Each of three independent experiments showing trends in OD600 over time on incubation of defibrinated sheep blood with the indicated concentrations of peptide in μM at 37 °C. Decreasing OD600 indicates increasing hemolysis.

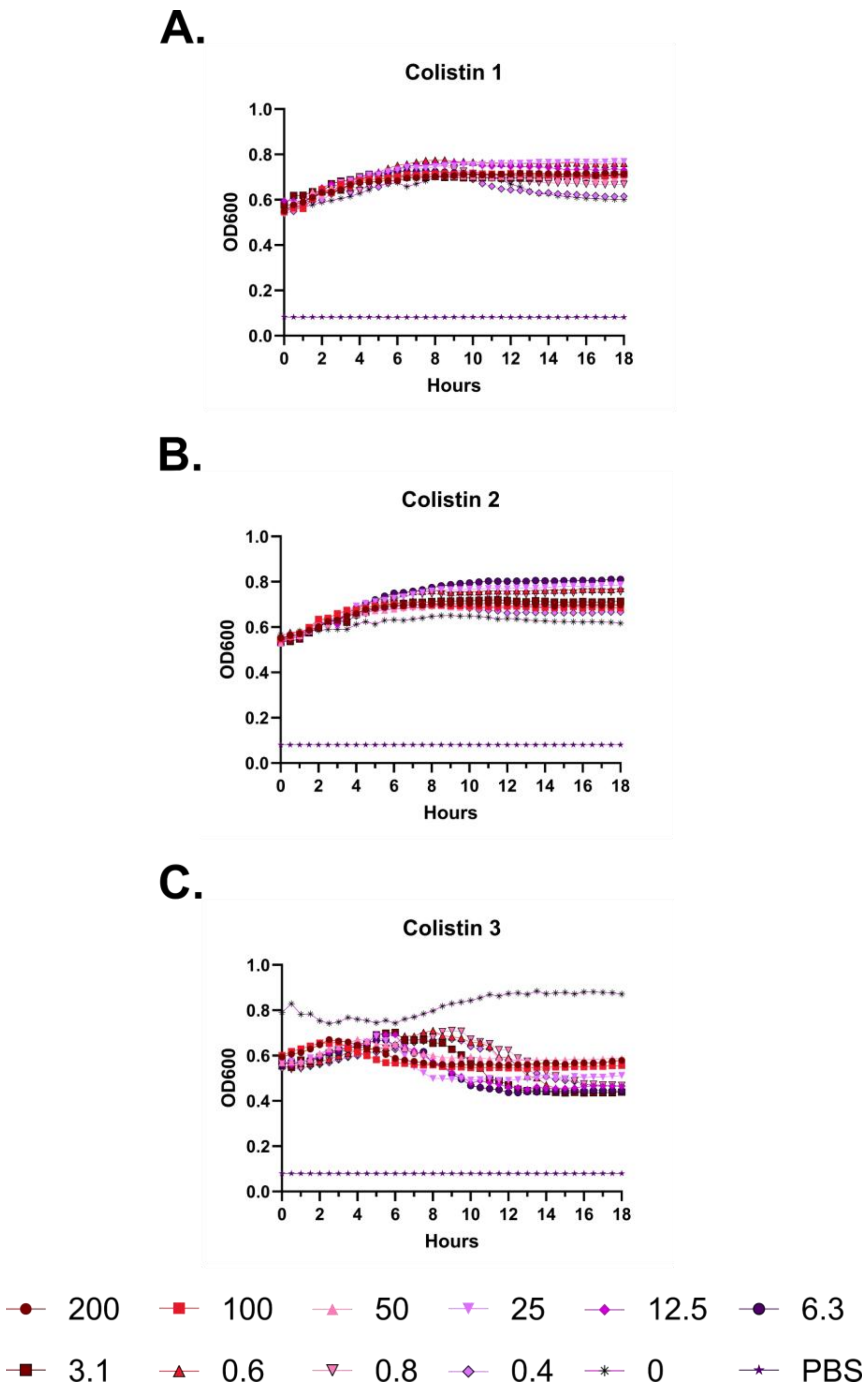
A.
Colistin 1
OD600
Hours
B.
Colistin 2
OD600
Hours
C.
Colistin 3
OD600
Hours
200
100
50
25
12.5
6.3
3.1
0.6
0.8
0.4
0
PBS

**Figure S57 – Hemolysis over time with colistin. A-C.** Each of three independent experiments showing trends in OD600 over time on incubation of defibrinated sheep blood with the indicated concentrations of peptide in μM at 37 °C. Decreasing OD600 indicates increasing hemolysis.

**A.**

| | % Helix | MIC |
|---|---|---|
| **F1** | 39.6 | 12.5 |
| **F2** | 16.7 | 25 |
| **I5** | 55.3 | 25 |
| **V6** | 18.7 | 12.5 |
| **Q7** | 26.5 | 6.3 |
| **R8** | 100 | 3.1 |
| **I9** | 8.1 | 100 |
| **K10** | 16.2 | 12.5 |
| **D11** | 92.7 | 3.1 |
| **F12** | 75.9 | 12.5 |
| **L13** | 10.3 | 50 |
| **R14** | 16 | 6.3 |

**B**

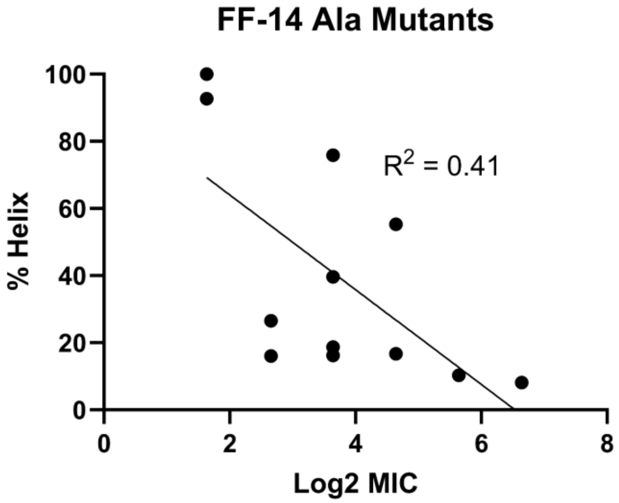

**Figure S58 – Modest overall correlation between helical content and MIC. A.** Table showing the calculated helical content for each FF-14 Ala mutant and the corresponding MIC against *E. coli*. **B.** Scatter plot with linear regression showing modest correlation between helical content and MIC among FF-14 Ala mutants.

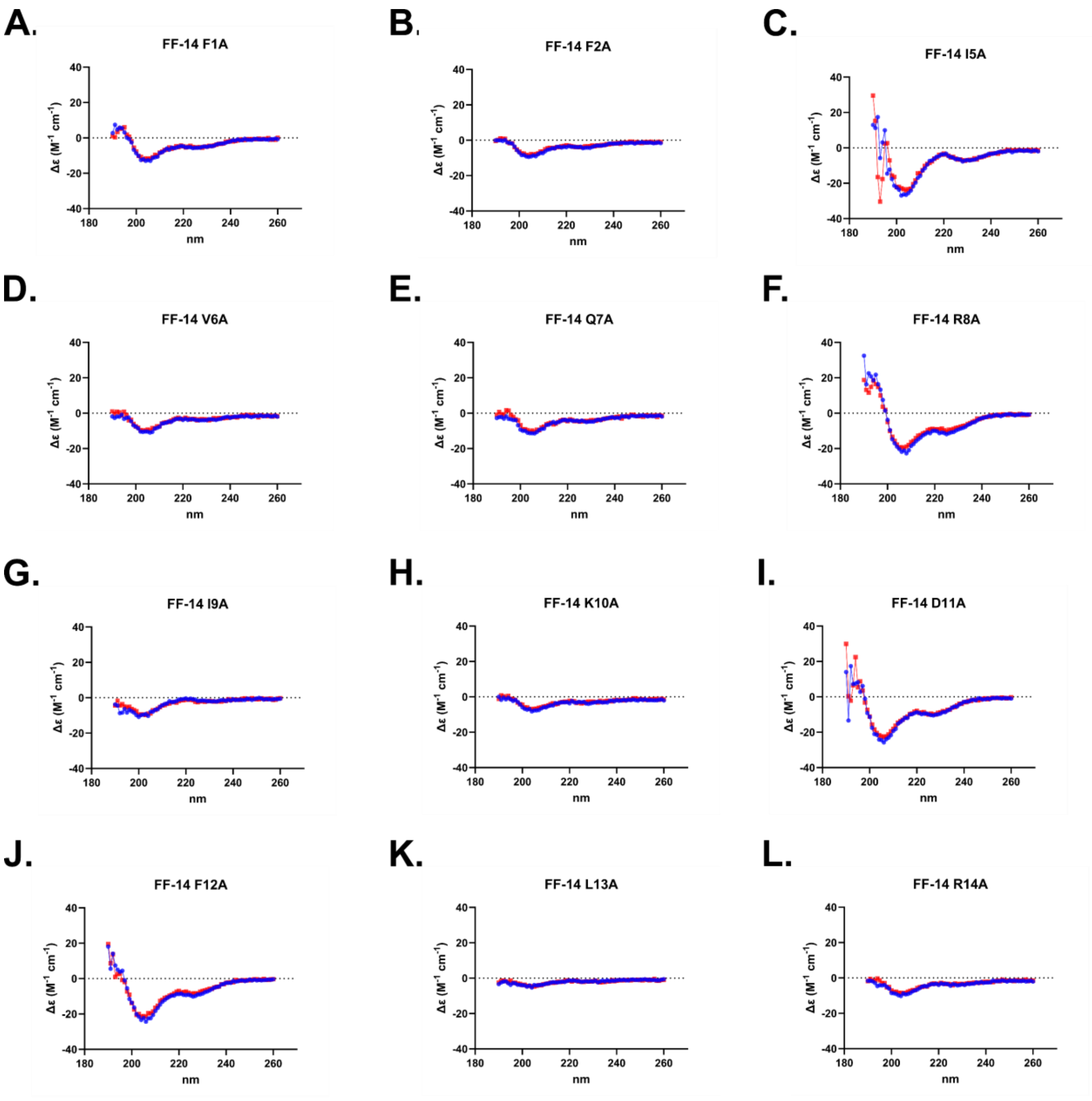


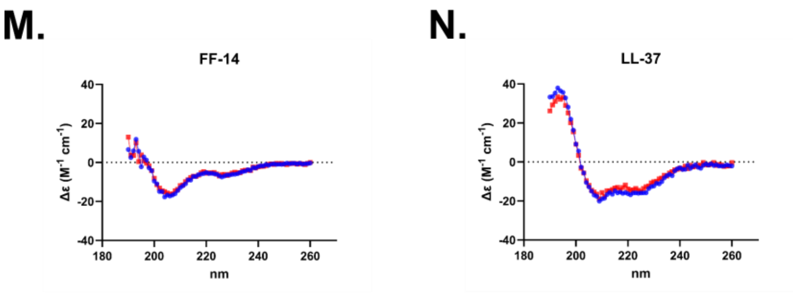


**O.**

| | % Helix | |
|---|---|---|
| | 20 °C | 37 °C |
| FF-14 F1A | 40.2 | 39.6 |
| FF-14 F2A | 20.6 | 16.7 |
| FF-14 I5A | 76.5 | 55.3 |
| FF-14 V6A | 22.3 | 18.7 |
| FF-14 Q7A | 27.8 | 26.5 |
| FF-14 R8A | 100 | 100 |
| FF-14 I9A | 5 | 8.1 |
| FF-14 K10A | 16 | 16.2 |
| FF-14 D11A | 90.6 | 92.7 |
| FF-14 F12A | 84.4 | 75.9 |
| FF-14 L13A | 10.1 | 10.3 |
| FF-14 R14A | 18.1 | 16 |
| FF-14 | 66.4 | 62.2 |
| LL-37 | 100 | 100 |

**Figure S59 – Circular dichroism spectroscopy of Figure 2 mutants. A-N.** CD spectra are displayed for each mutant collected from 190-260 nm at both 20 °C (Blue) and 37 °C (Red). **O.** Quantification of helical content as determined by BeStSel for each mutant.

A.

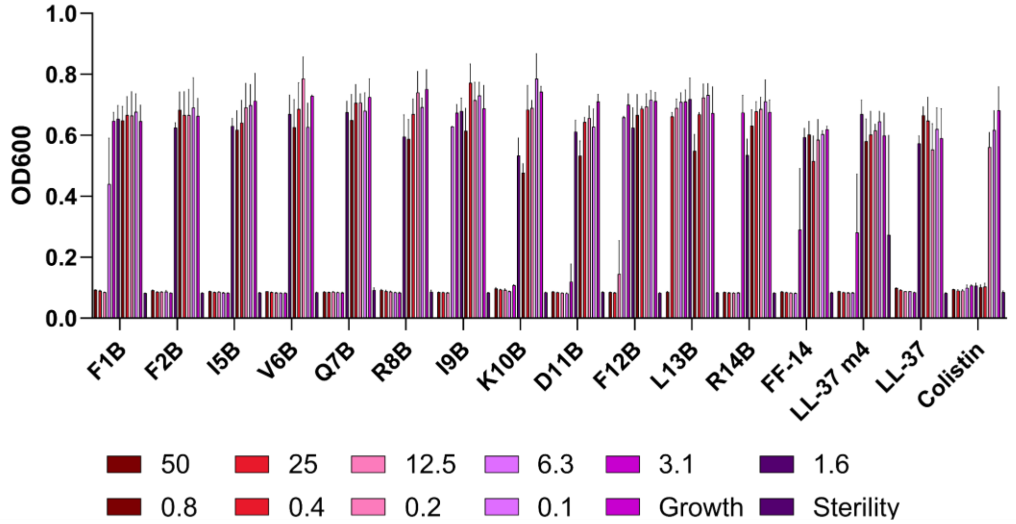

OD600
1.0
0.8
0.6
0.4
0.2
0.0
F1B
F2B
I5B
V6B
Q7B
R8B
I9B
K10B
D11B
F12B
L13B
R14B
FF-14
LL-37 m4
LL-37
Colistin
50
25
12.5
6.3
3.1
1.6
0.8
0.4
0.2
0.1
Growth
Sterility


B.

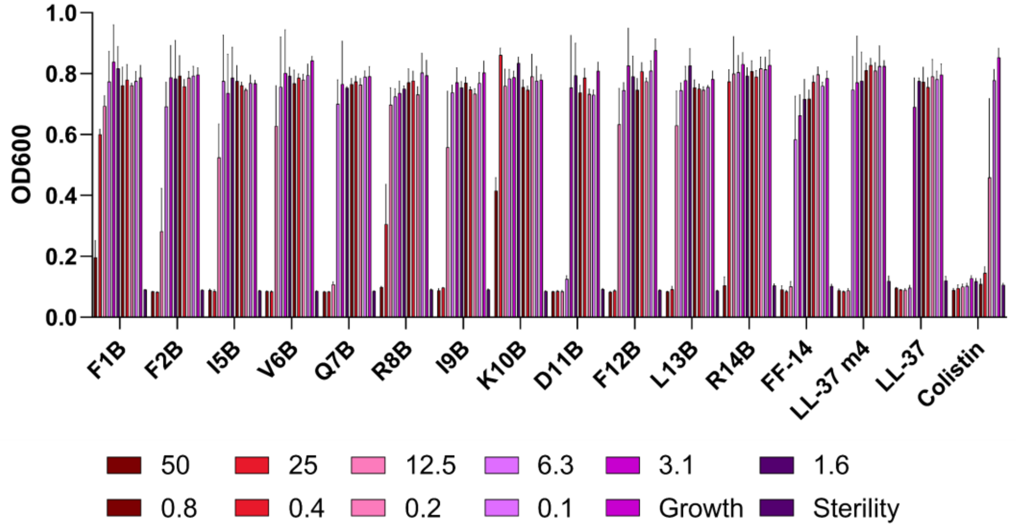

OD600
1.0
0.8
0.6
0.4
0.2
0.0
F1B
F2B
I5B
V6B
Q7B
R8B
I9B
K10B
D11B
F12B
L13B
R14B
FF-14
LL-37 m4
LL-37
Colistin
50
25
12.5
6.3
3.1
1.6
0.8
0.4
0.2
0.1
Growth
Sterility


C.

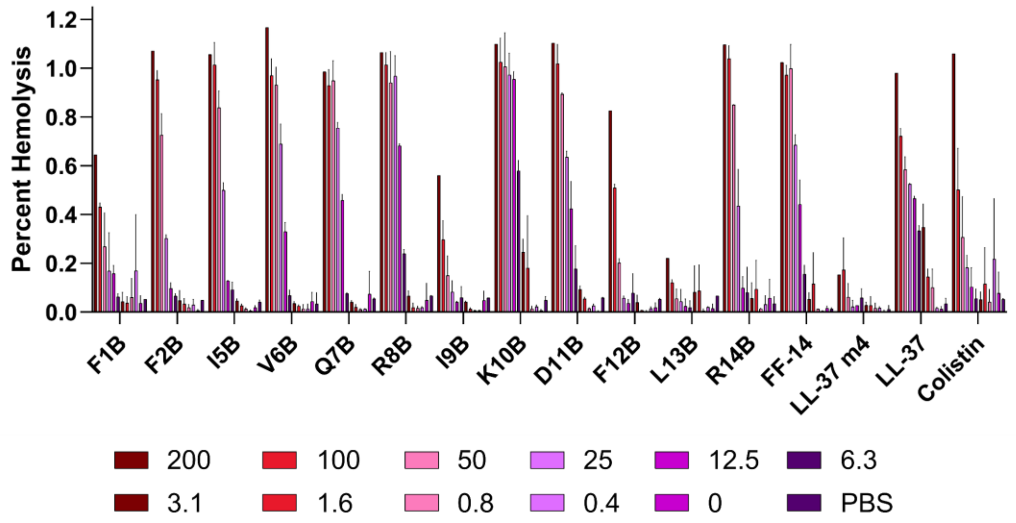

Percent Hemolysis
1.2
1.0
0.8
0.6
0.4
0.2
0.0
F1B
F2B
I5B
V6B
Q7B
R8B
I9B
K10B
D11B
F12B
L13B
R14B
FF-14
LL-37 m4
LL-37
Colistin
200
100
50
25
12.5
6.3
3.1
1.6
0.8
0.4
0
PBS

**Figure S60 – Histogram depiction of data from Figure 3. A.** Endpoint activity of each peptide against *E. coli*; measures are raw absorbance at 600 nm at 18 hours. Data show the mean and standard deviation of three independent experiments. **B.** Endpoint activity of each peptide against *P. aeruginosa*. **C.** Hemolytic activity of each peptide; measures are absorbance at 414 nm scaled to a freeze-thaw positive control set at 100% hemolysis. Data show the mean and standard deviation of two independent experiments; the 200 μM concentration is omitted from one of these experiments due to a dilution error. Concentrations throughout are shown in μM.

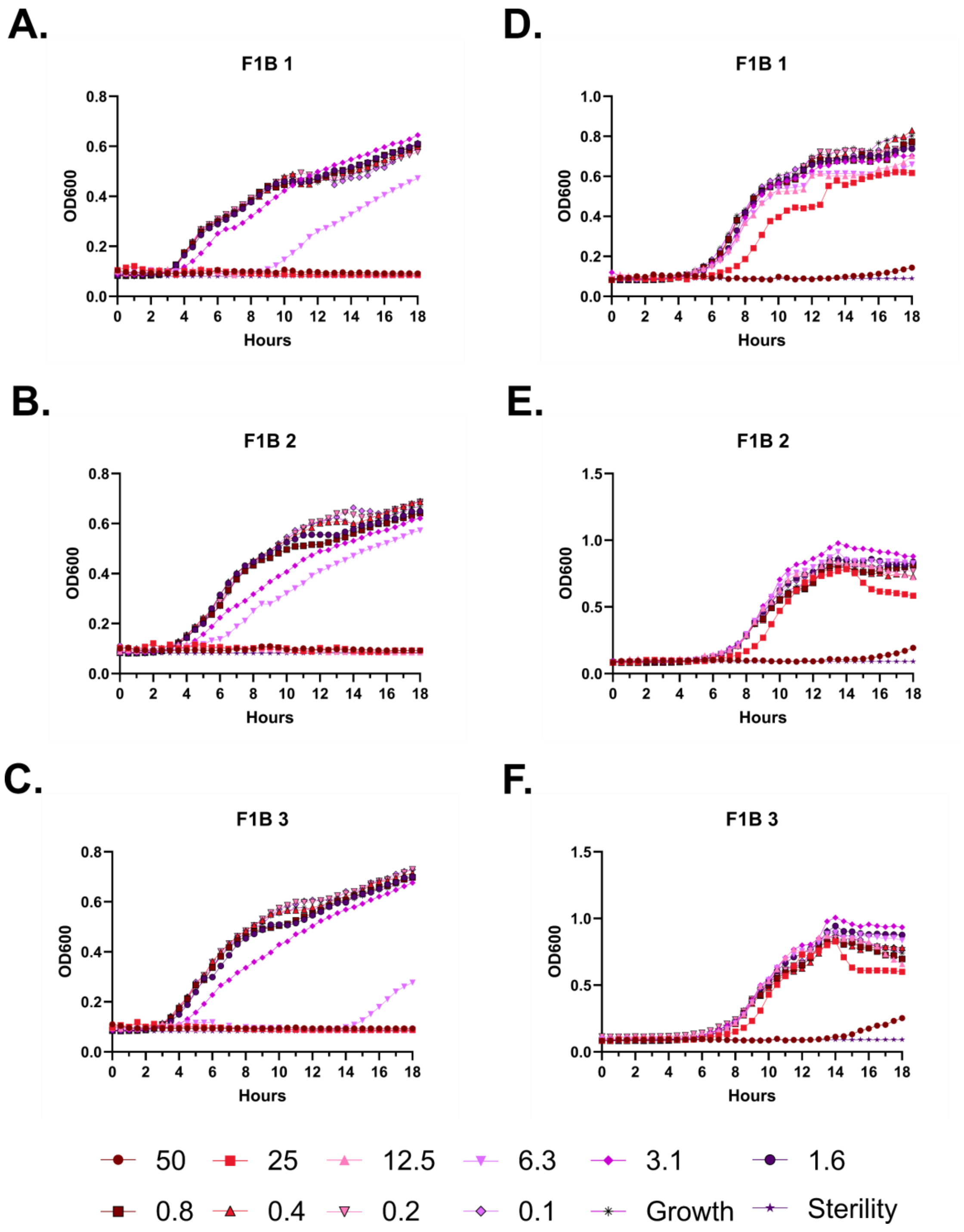
A.
F1B 1
OD600
Hours
D.
F1B 1
OD600
Hours
B.
F1B 2
OD600
Hours
E.
F1B 2
OD600
Hours
C.
F1B 3
OD600
Hours
F.
F1B 3
OD600
Hours
50
25
12.5
6.3
3.1
1.6
0.8
0.4
0.2
0.1
Growth
Sterility

**Figure S61 – Antimicrobial susceptibility testing with FF-14 F1B. A-C.** Growth of *E. coli* in the presence of the indicated peptide concentrations given in μM as monitored by OD600 at 37 °C over 18 hours. Each panel indicates an independent experiment. **D-F.** As in **A-C**, but with *P. aeruginosa.*

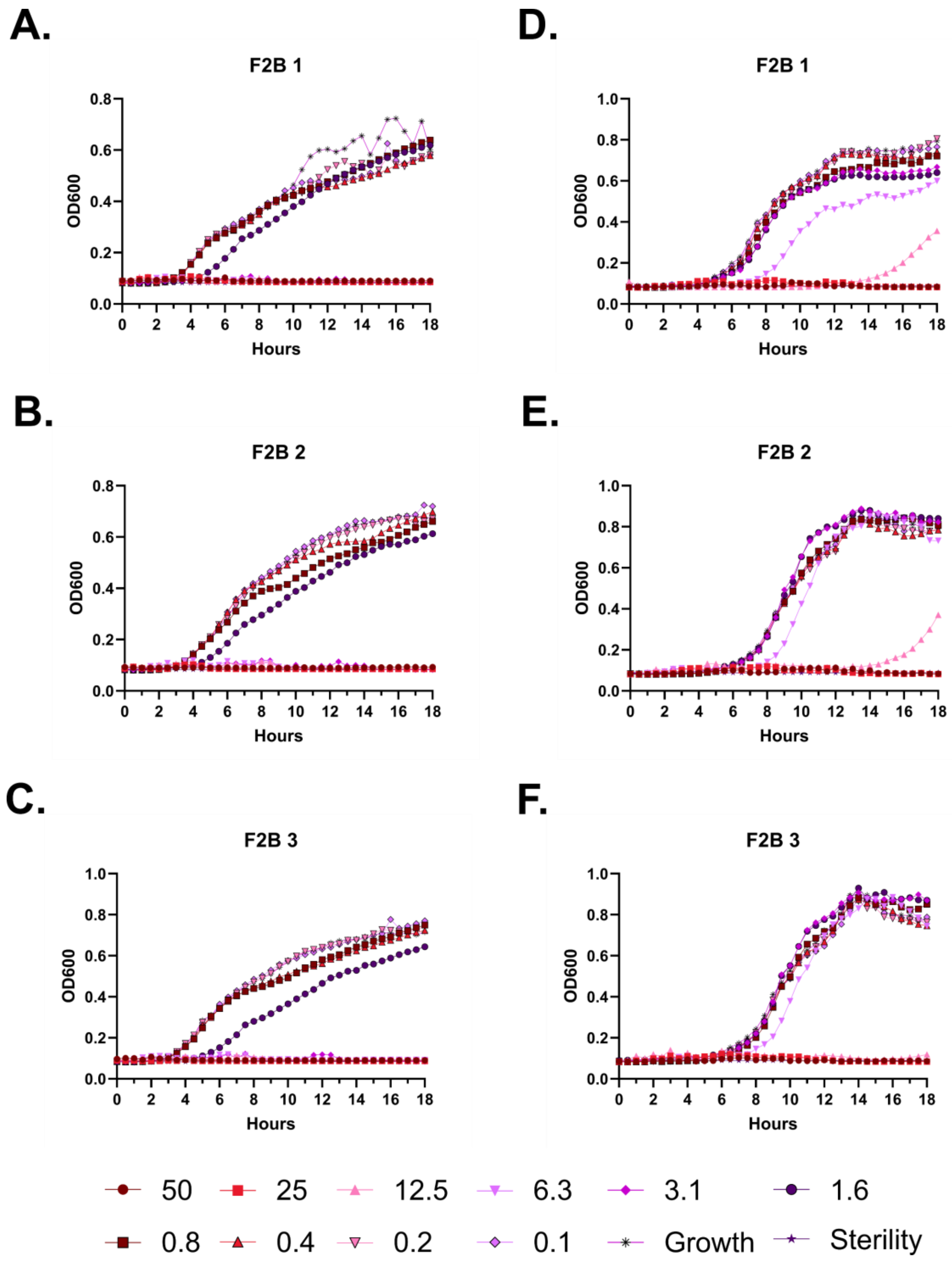
A.
F2B 1
OD600
Hours
D.
F2B 1
OD600
Hours
B.
F2B 2
OD600
Hours
E.
F2B 2
OD600
Hours
C.
F2B 3
OD600
Hours
F.
F2B 3
OD600
Hours
50
25
12.5
6.3
3.1
1.6
0.8
0.4
0.2
0.1
Growth
Sterility

**Figure S62 – Antimicrobial susceptibility testing with FF-14 F2B. A-C.** Growth of *E. coli* in the presence of the indicated peptide concentrations given in μM as monitored by OD600 at 37 °C over 18 hours. Each panel indicates an independent experiment. **D-F.** As in **A-C**, but with *P. aeruginosa.*

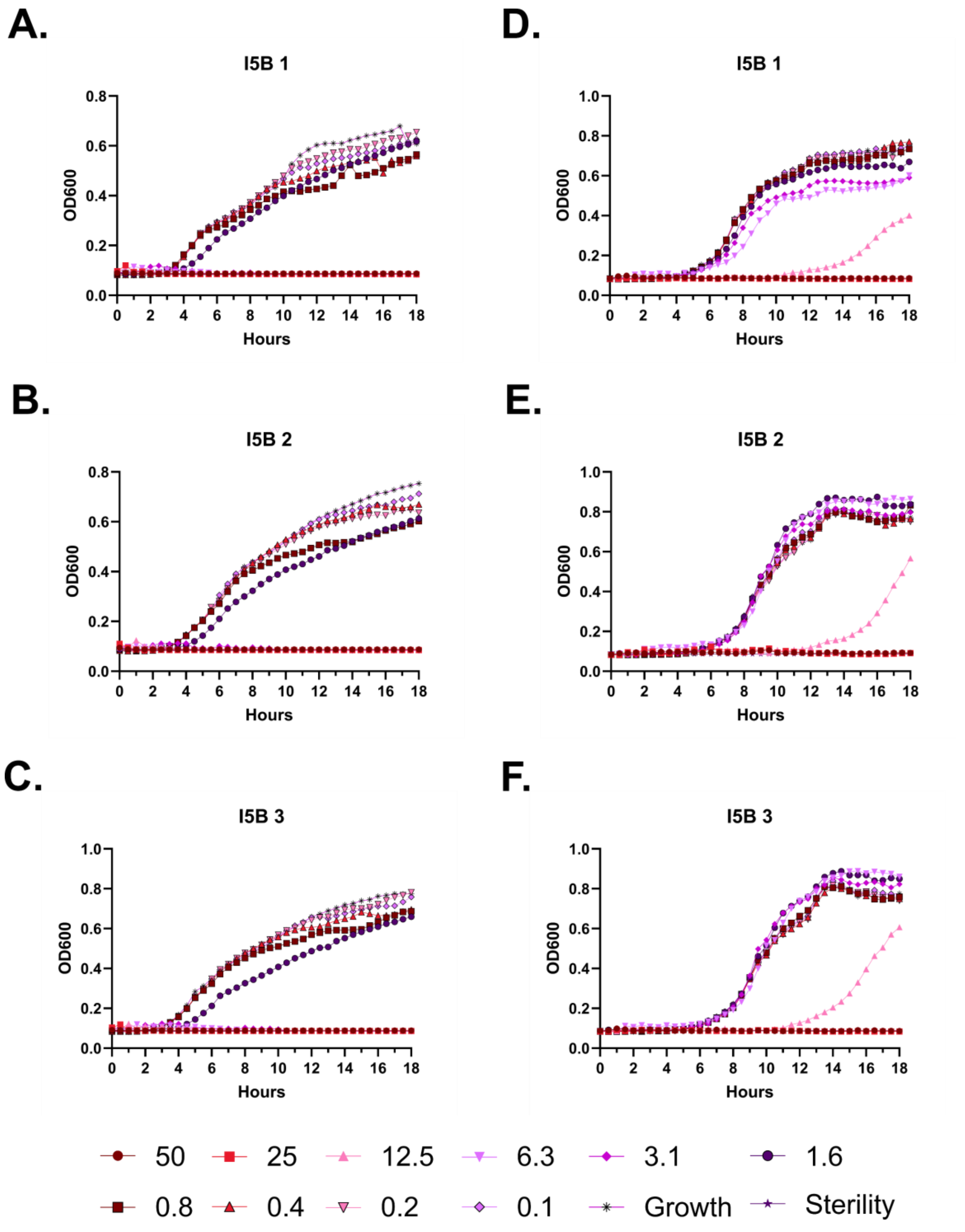

A.
I5B 1
B.
I5B 2
C.
I5B 3
D.
I5B 1
E.
I5B 2
F.
I5B 3
OD600
Hours
50
25
12.5
6.3
3.1
1.6
0.8
0.4
0.2
0.1
Growth
Sterility

**Figure S63 – Antimicrobial susceptibility testing with FF-14 I5B. A-C.** Growth of *E. coli* in the presence of the indicated peptide concentrations given in μM as monitored by OD600 at 37 °C over 18 hours. Each panel indicates an independent experiment. **D-F.** As in **A-C**, but with *P. aeruginosa.*

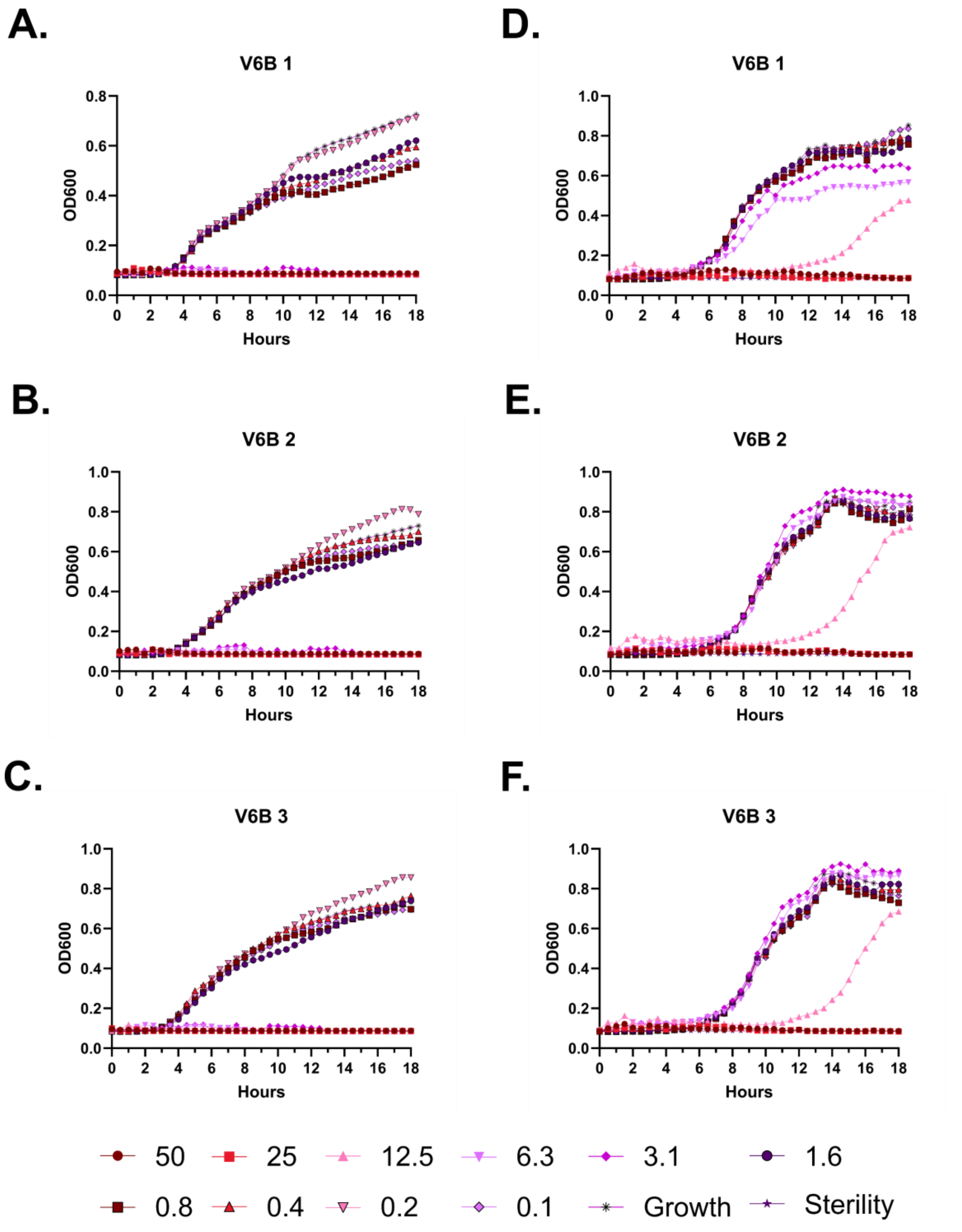
A.
V6B 1
OD600
Hours
D.
V6B 1
OD600
Hours
B.
V6B 2
OD600
Hours
E.
V6B 2
OD600
Hours
C.
V6B 3
OD600
Hours
F.
V6B 3
OD600
Hours
50
25
12.5
6.3
3.1
1.6
0.8
0.4
0.2
0.1
Growth
Sterility

**Figure S64 – Antimicrobial susceptibility testing with FF-14 V6B. A-C.** Growth of *E. coli* in the presence of the indicated peptide concentrations given in μM as monitored by OD600 at 37 °C over 18 hours. Each panel indicates an independent experiment. **D-F.** As in **A-C**, but with *P. aeruginosa.*

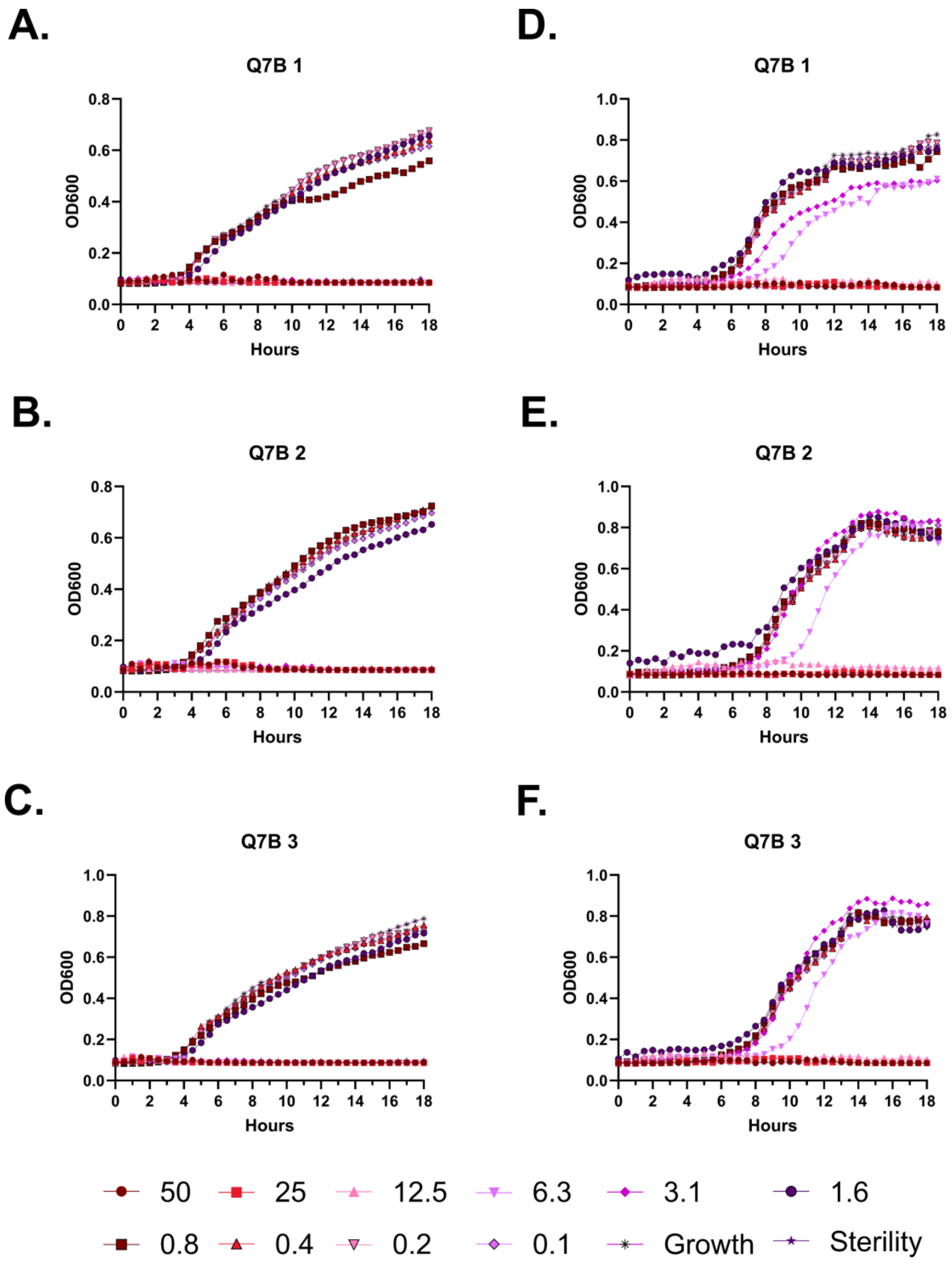
A.
Q7B 1
D.
Q7B 1
B.
Q7B 2
E.
Q7B 2
C.
Q7B 3
F.
Q7B 3
OD600
Hours
50
25
12.5
6.3
3.1
1.6
0.8
0.4
0.2
0.1
Growth
Sterility

**Figure S65 – Antimicrobial susceptibility testing with FF-14 Q7B. A-C.** Growth of *E. coli* in the presence of the indicated peptide concentrations given in μM as monitored by OD600 at 37 °C over 18 hours. Each panel indicates an independent experiment. **D-F.** As in **A-C**, but with *P. aeruginosa.*

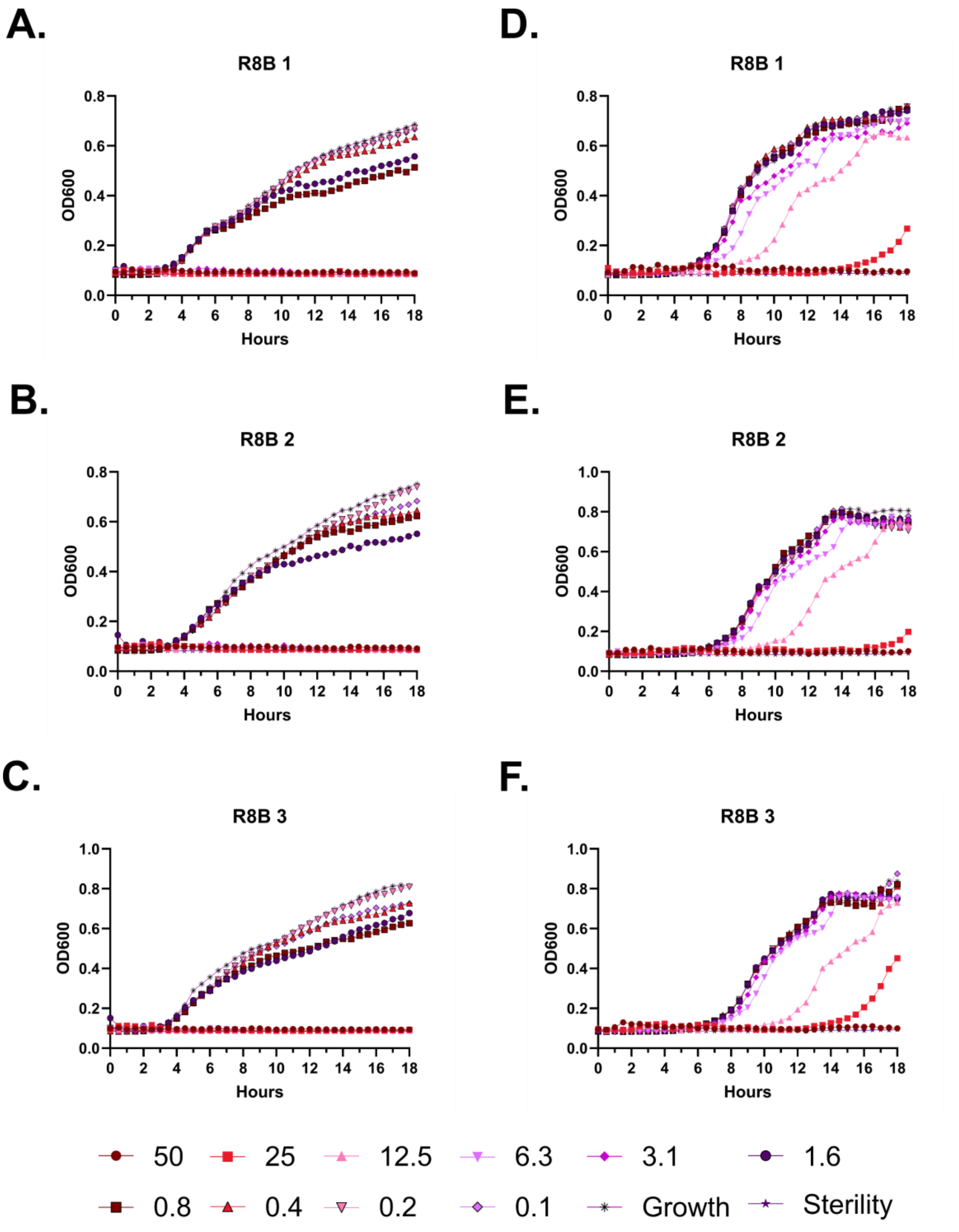
A.
R8B 1
D.
R8B 1
B.
R8B 2
E.
R8B 2
C.
R8B 3
F.
R8B 3
OD600
Hours
50
25
12.5
6.3
3.1
1.6
0.8
0.4
0.2
0.1
Growth
Sterility

**Figure S66 – Antimicrobial susceptibility testing with FF-14 R8B. A-C.** Growth of *E. coli* in the presence of the indicated peptide concentrations given in μM as monitored by OD600 at 37 °C over 18 hours. Each panel indicates an independent experiment. **D-F.** As in **A-C**, but with *P. aeruginosa.*

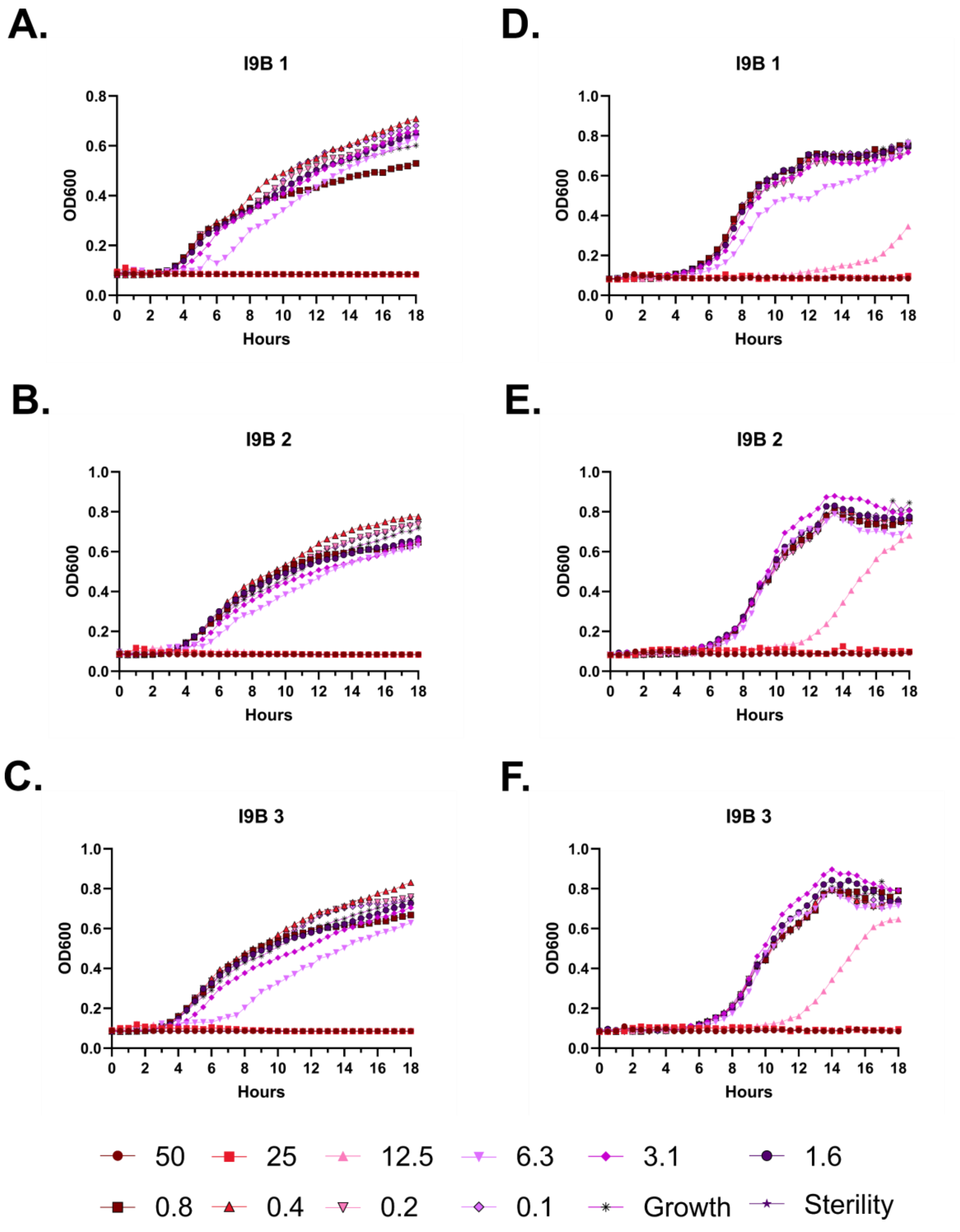
A.
I9B 1
D.
I9B 1
B.
I9B 2
E.
I9B 2
C.
I9B 3
F.
I9B 3
OD600
Hours
50
25
12.5
6.3
3.1
1.6
0.8
0.4
0.2
0.1
Growth
Sterility

**Figure S67 – Antimicrobial susceptibility testing with FF-14 I9B. A-C.** Growth of *E. coli* in the presence of the indicated peptide concentrations given in μM as monitored by OD600 at 37 °C over 18 hours. Each panel indicates an independent experiment. **D-F.** As in **A-C**, but with *P. aeruginosa.*

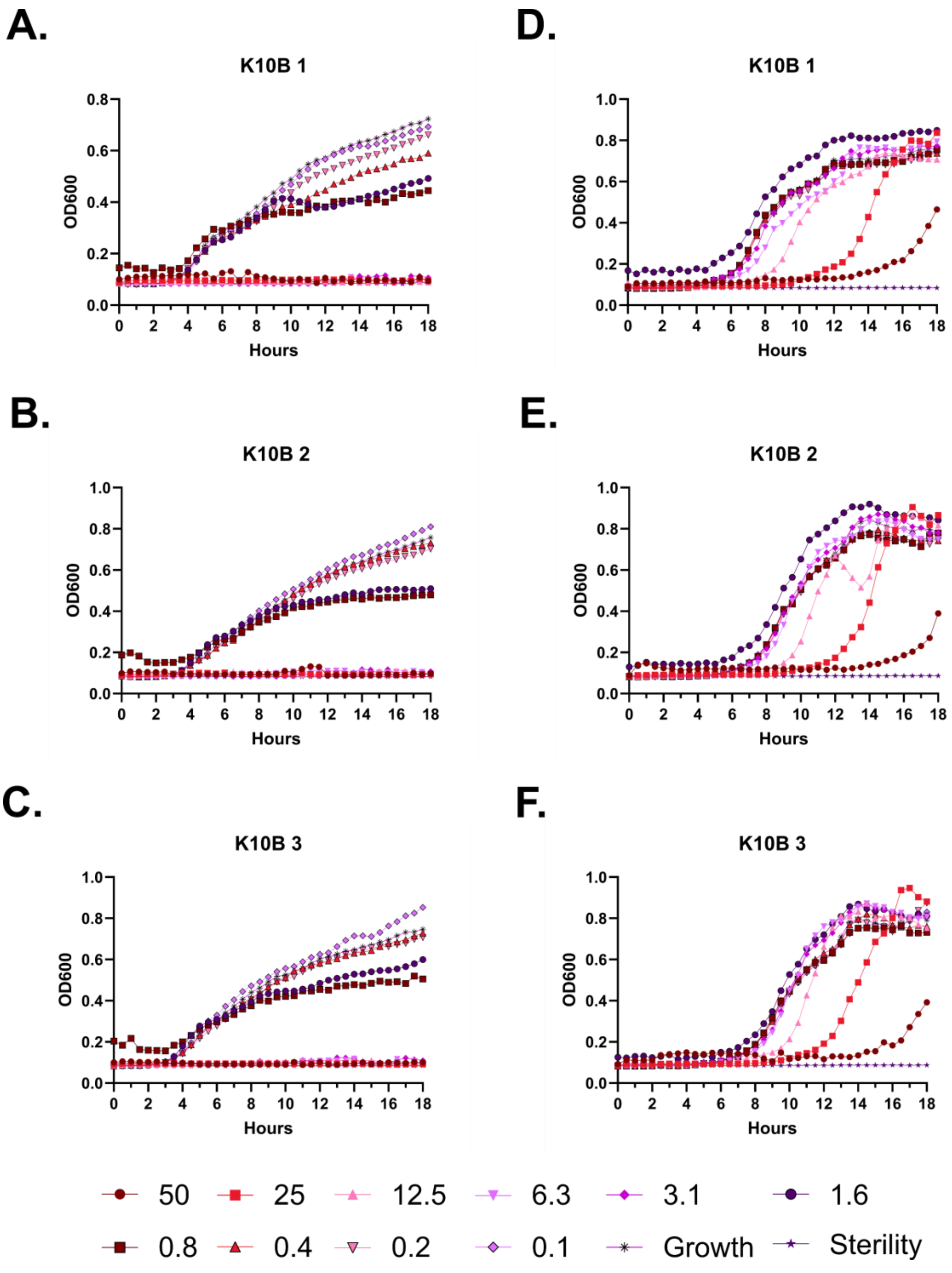
A.
K10B 1
OD600
Hours
B.
K10B 2
OD600
Hours
C.
K10B 3
OD600
Hours
D.
K10B 1
OD600
Hours
E.
K10B 2
OD600
Hours
F.
K10B 3
OD600
Hours
50
25
12.5
6.3
3.1
1.6
0.8
0.4
0.2
0.1
Growth
Sterility

**Figure S68 – Antimicrobial susceptibility testing with FF-14 K10B. A-C.** Growth of *E. coli* in the presence of the indicated peptide concentrations given in μM as monitored by OD600 at 37 °C over 18 hours. Each panel indicates an independent experiment. **D-F.** As in **A-C**, but with *P. aeruginosa.*

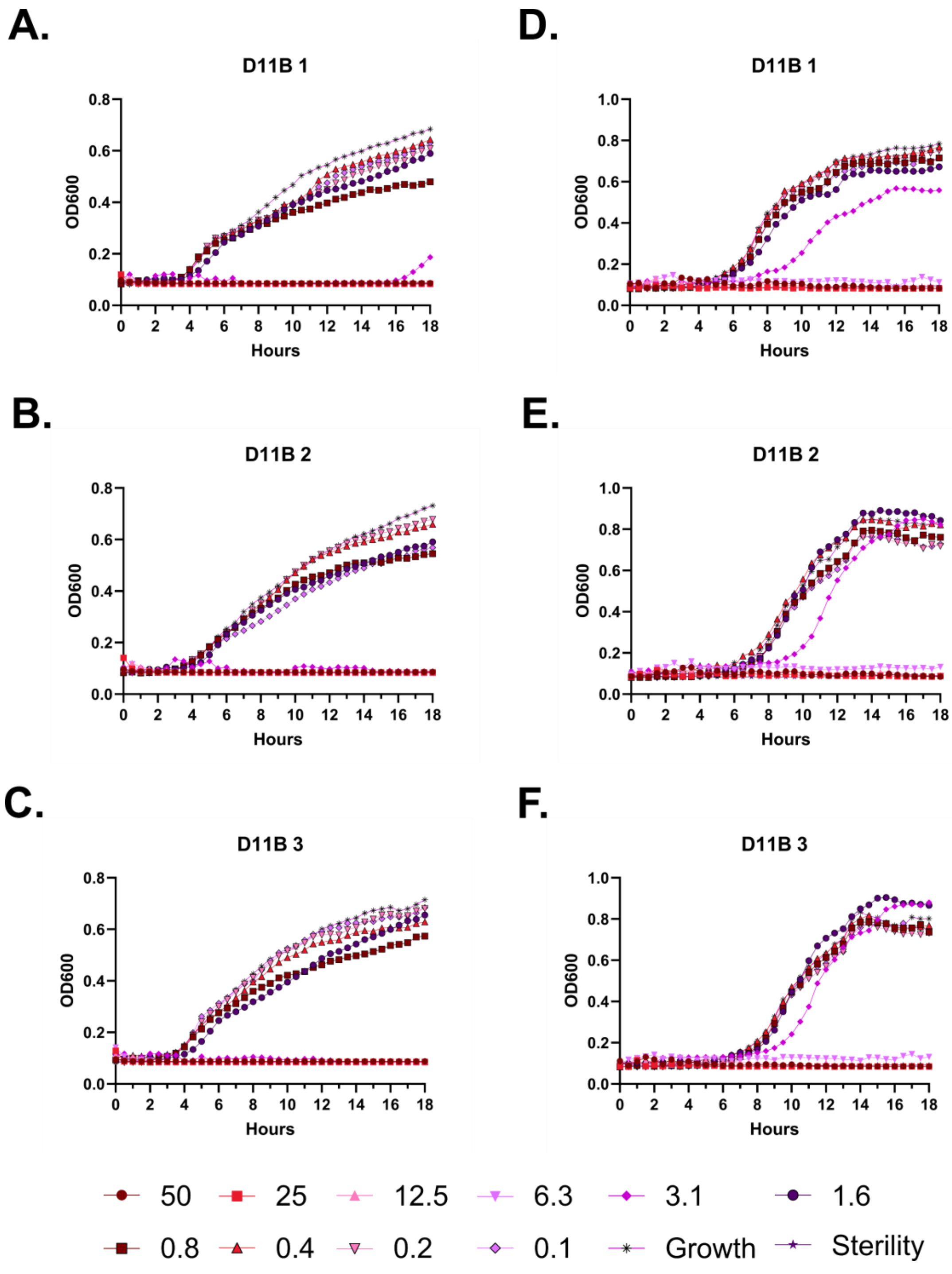
A.
D11B 1
OD600
Hours
D.
D11B 1
OD600
Hours
B.
D11B 2
OD600
Hours
E.
D11B 2
OD600
Hours
C.
D11B 3
OD600
Hours
F.
D11B 3
OD600
Hours
50
25
12.5
6.3
3.1
1.6
0.8
0.4
0.2
0.1
Growth
Sterility

**Figure S69 – Antimicrobial susceptibility testing with FF-14 D11B. A-C.** Growth of *E. coli* in the presence of the indicated peptide concentrations given in μM as monitored by OD600 at 37 °C over 18 hours. Each panel indicates an independent experiment. **D-F.** As in **A-C**, but with *P. aeruginosa.*

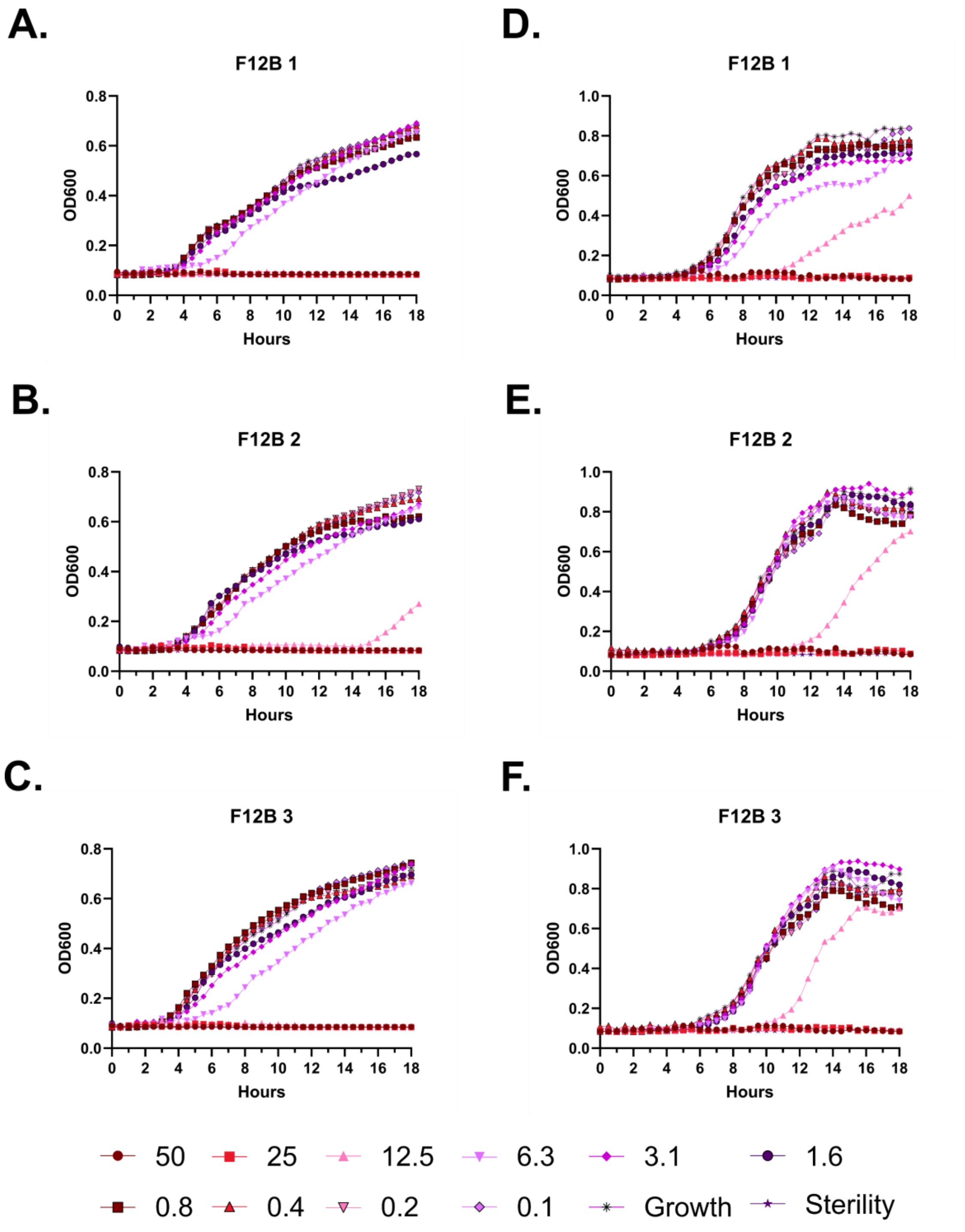

A.
F12B 1
OD600
Hours
D.
F12B 1
OD600
Hours
B.
F12B 2
OD600
Hours
E.
F12B 2
OD600
Hours
C.
F12B 3
OD600
Hours
F.
F12B 3
OD600
Hours
50
25
12.5
6.3
3.1
1.6
0.8
0.4
0.2
0.1
Growth
Sterility

**Figure S70 – Antimicrobial susceptibility testing with FF-14 F12B. A-C.** Growth of *E. coli* in the presence of the indicated peptide concentrations given in μM as monitored by OD600 at 37 °C over 18 hours. Each panel indicates an independent experiment. **D-F.** As in **A-C**, but with *P. aeruginosa.*

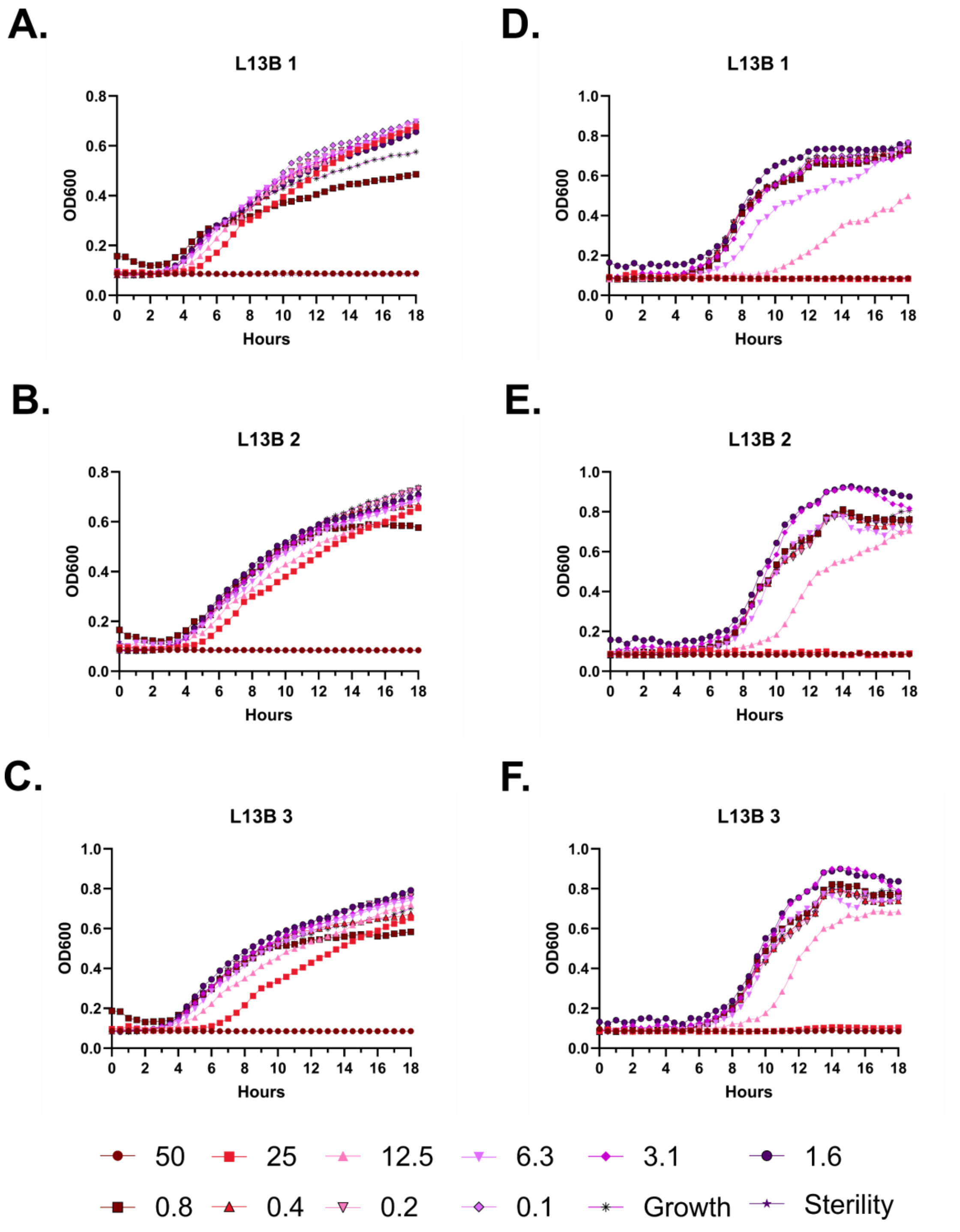
A.
L13B 1
OD600
Hours
B.
L13B 2
OD600
Hours
C.
L13B 3
OD600
Hours
D.
L13B 1
OD600
Hours
E.
L13B 2
OD600
Hours
F.
L13B 3
OD600
Hours
50
25
12.5
6.3
3.1
1.6
0.8
0.4
0.2
0.1
Growth
Sterility

**Figure S71 – Antimicrobial susceptibility testing with FF-14 L13B. A-C.** Growth of *E. coli* in the presence of the indicated peptide concentrations given in μM as monitored by OD600 at 37 °C over 18 hours. Each panel indicates an independent experiment. **D-F.** As in **A-C**, but with *P. aeruginosa.*

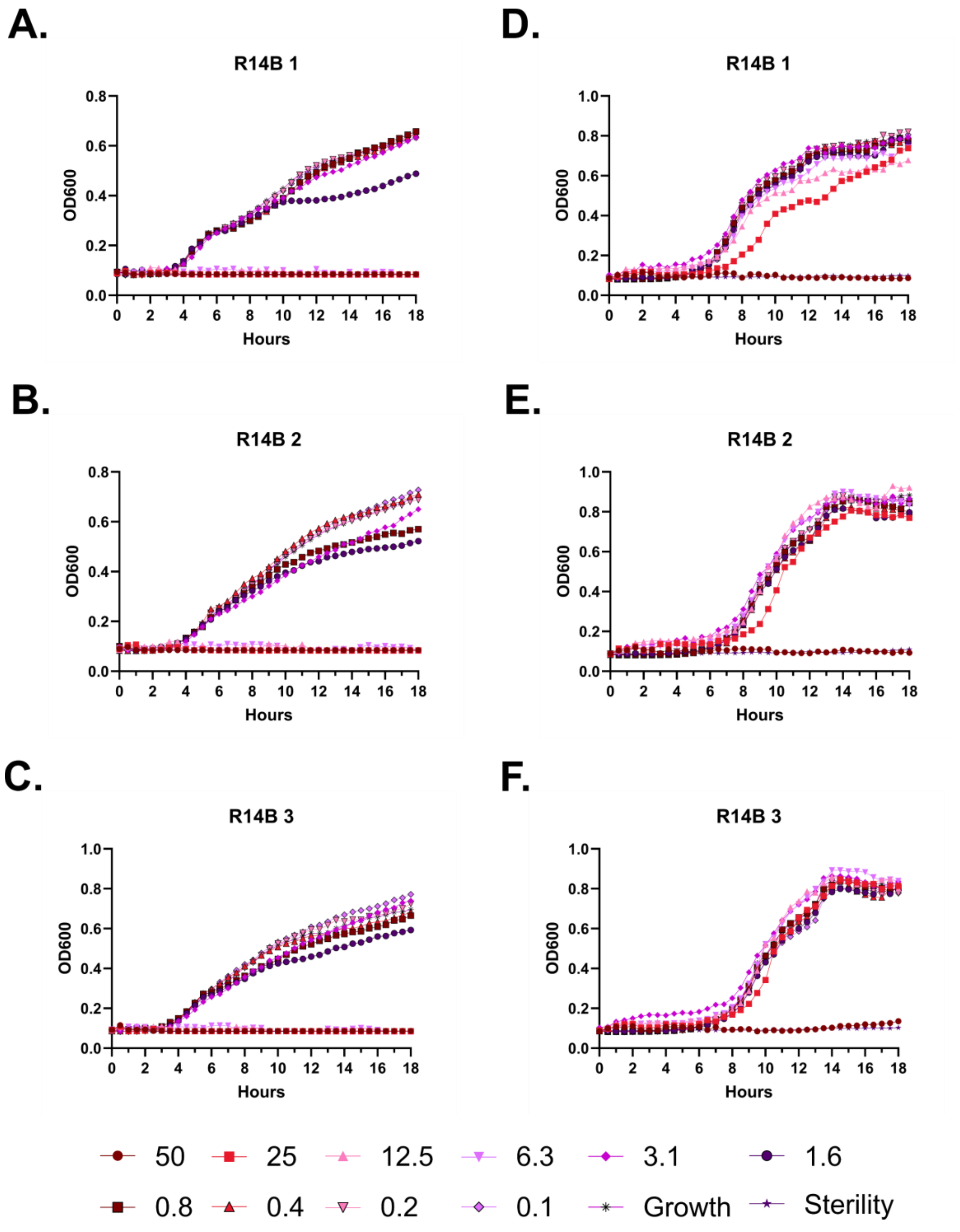
A.
R14B 1
OD600
Hours
D.
R14B 1
OD600
Hours
B.
R14B 2
OD600
Hours
E.
R14B 2
OD600
Hours
C.
R14B 3
OD600
Hours
F.
R14B 3
OD600
Hours
50
25
12.5
6.3
3.1
1.6
0.8
0.4
0.2
0.1
Growth
Sterility

**Figure S72 – Antimicrobial susceptibility testing with FF-14 R14B. A-C.** Growth of *E. coli* in the presence of the indicated peptide concentrations given in μM as monitored by OD600 at 37 °C over 18 hours. Each panel indicates an independent experiment. **D-F.** As in **A-C**, but with *P. aeruginosa.*

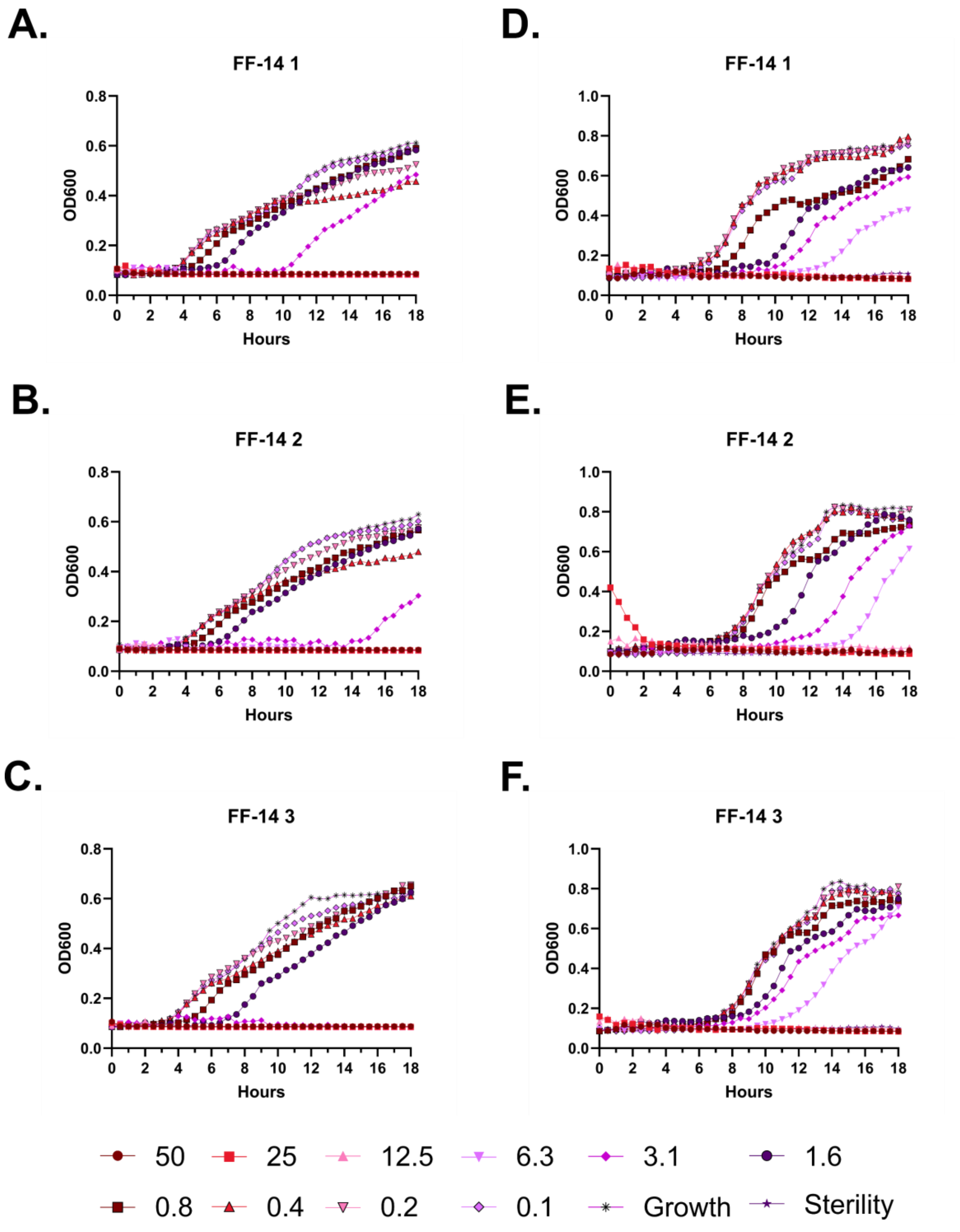
A.
FF-14 1
D.
FF-14 1
B.
FF-14 2
E.
FF-14 2
C.
FF-14 3
F.
FF-14 3
OD600
Hours
50
25
12.5
6.3
3.1
1.6
0.8
0.4
0.2
0.1
Growth
Sterility

**Figure S73 – Antimicrobial susceptibility testing with FF-14. A-C.** Growth of *E. coli* in the presence of the indicated peptide concentrations given in μM as monitored by OD600 at 37 °C over 18 hours. Each panel indicates an independent experiment. **D-F.** As in **A-C**, but with *P. aeruginosa.*

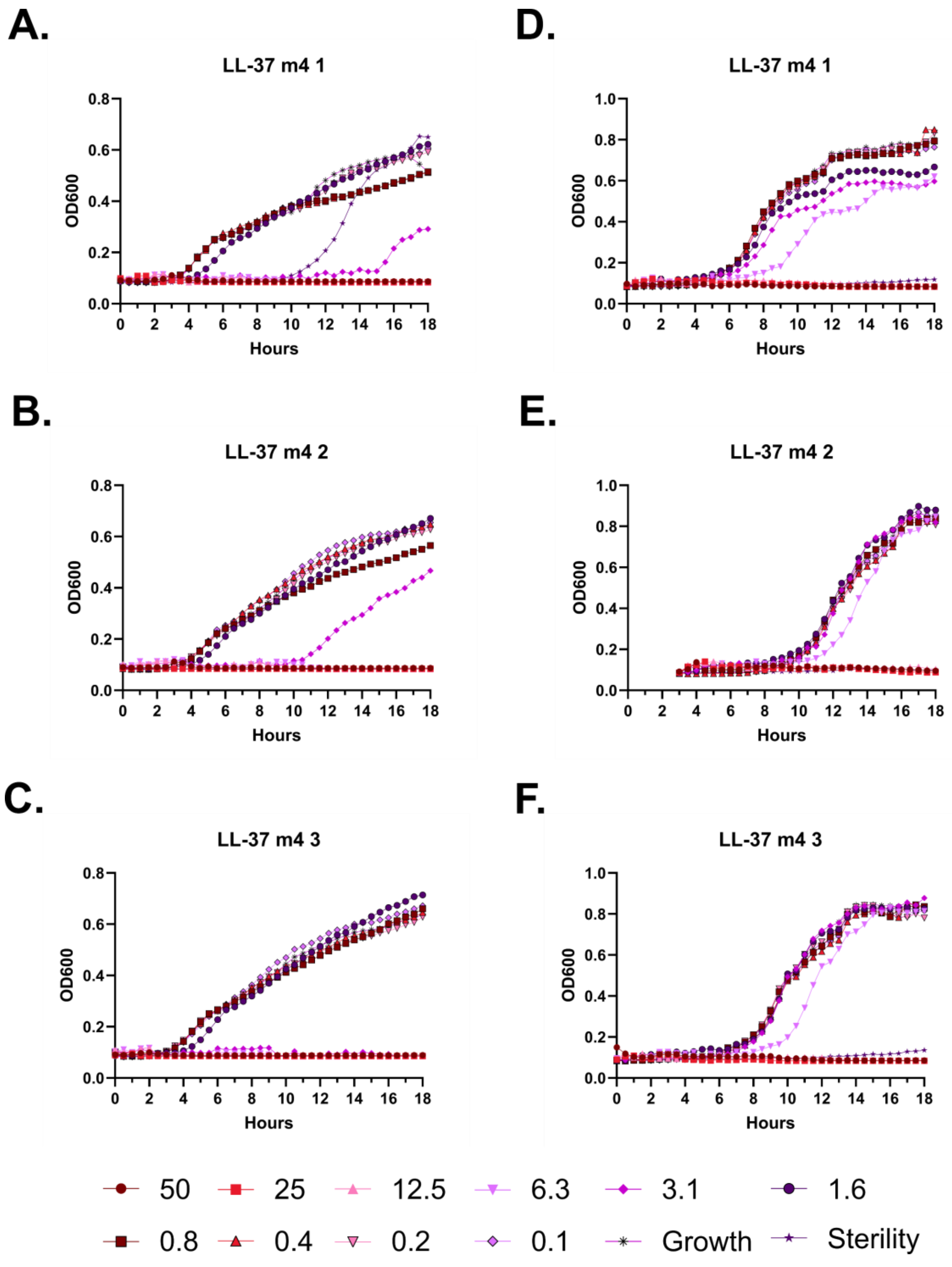
A.
LL-37 m4 1
OD600
Hours
D.
LL-37 m4 1
OD600
Hours
B.
LL-37 m4 2
OD600
Hours
E.
LL-37 m4 2
OD600
Hours
C.
LL-37 m4 3
OD600
Hours
F.
LL-37 m4 3
OD600
Hours
50
25
12.5
6.3
3.1
1.6
0.8
0.4
0.2
0.1
Growth
Sterility

**Figure S74 – Antimicrobial susceptibility testing with LL-37 m4. A-C.** Growth of *E. coli* in the presence of the indicated peptide concentrations given in μM as monitored by OD600 at 37 °C over 18 hours. Each panel indicates an independent experiment. **D-F.** As in **A-C**, but with *P. aeruginosa.*

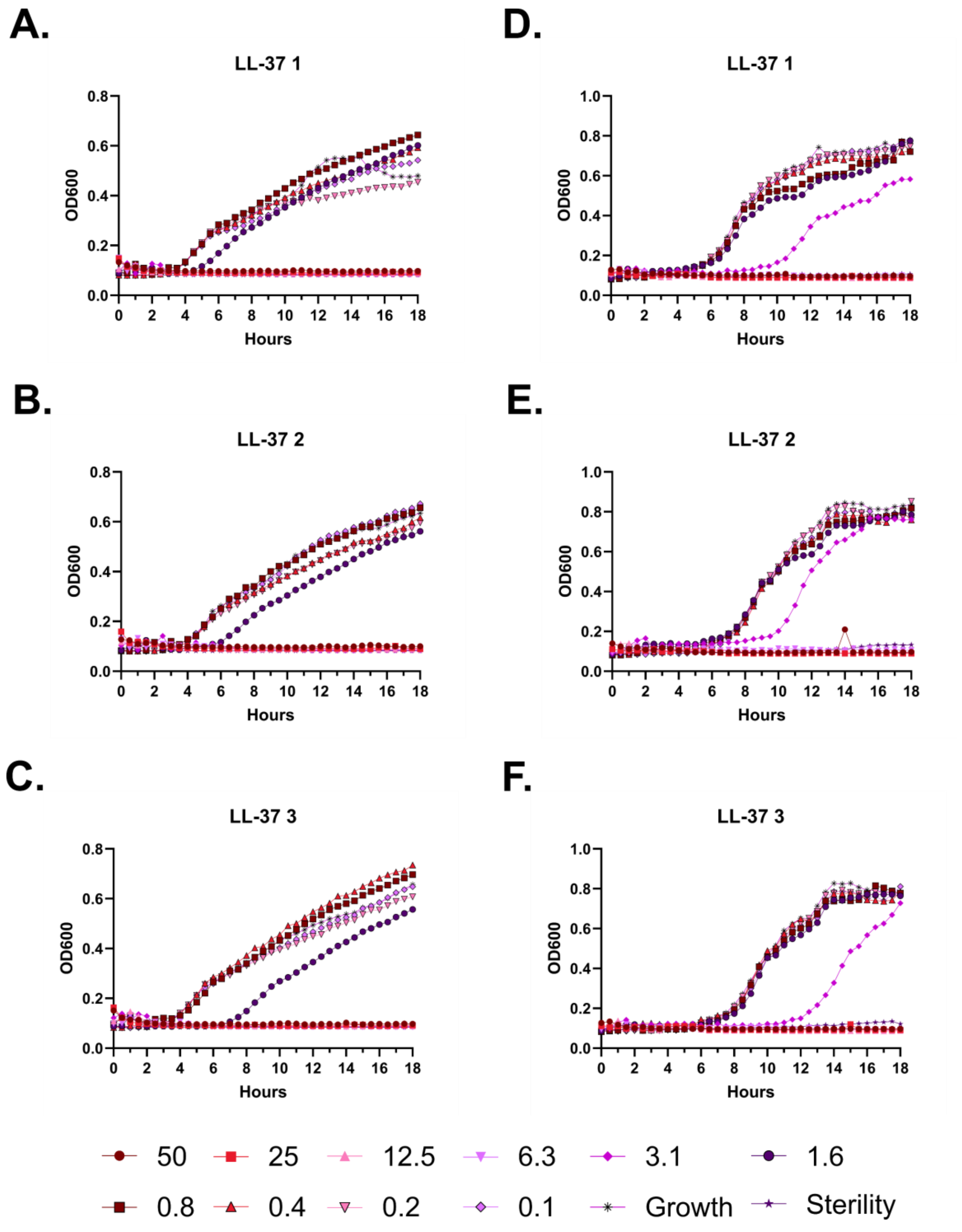

A.
LL-37 1
OD600
Hours
D.
LL-37 1
OD600
Hours
B.
LL-37 2
OD600
Hours
E.
LL-37 2
OD600
Hours
C.
LL-37 3
OD600
Hours
F.
LL-37 3
OD600
Hours
50
25
12.5
6.3
3.1
1.6
0.8
0.4
0.2
0.1
Growth
Sterility

**Figure S75 – Antimicrobial susceptibility testing with LL-37. A-C.** Growth of *E. coli* in the presence of the indicated peptide concentrations given in μM as monitored by OD600 at 37 °C over 18 hours. Each panel indicates an independent experiment. **D-F.** As in **A-C**, but with *P. aeruginosa.*

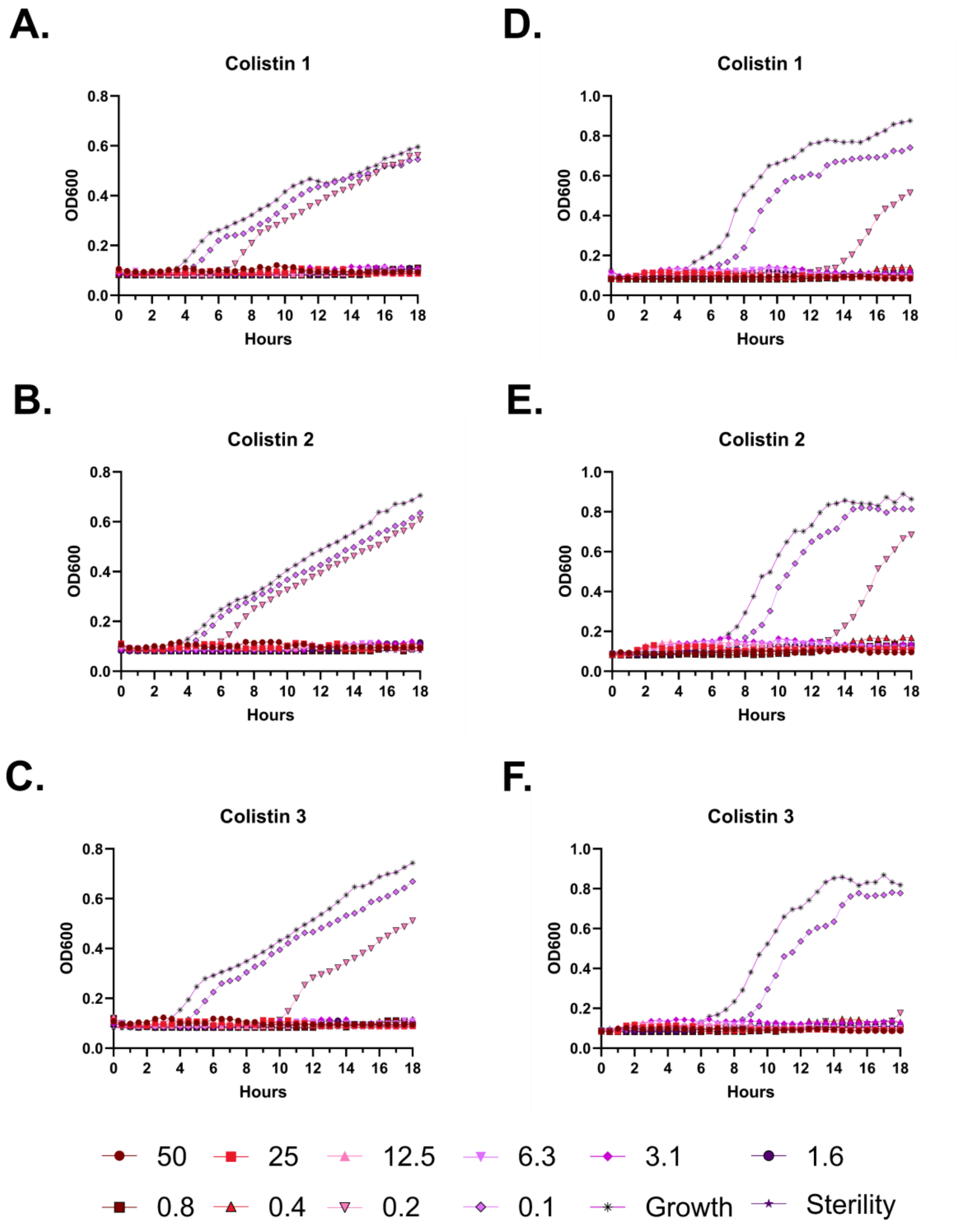
A.
Colistin 1
B.
Colistin 2
C.
Colistin 3
D.
Colistin 1
E.
Colistin 2
F.
Colistin 3
OD600
Hours
50
25
12.5
6.3
3.1
1.6
0.8
0.4
0.2
0.1
Growth
Sterility

**Figure S76 – Antimicrobial susceptibility testing with colistin. A-C.** Growth of *E. coli* in the presence of the indicated peptide concentrations given in μM as monitored by OD600 at 37 °C over 18 hours. Each panel indicates an independent experiment. **D-F.** As in **A-C**, but with *P. aeruginosa.*

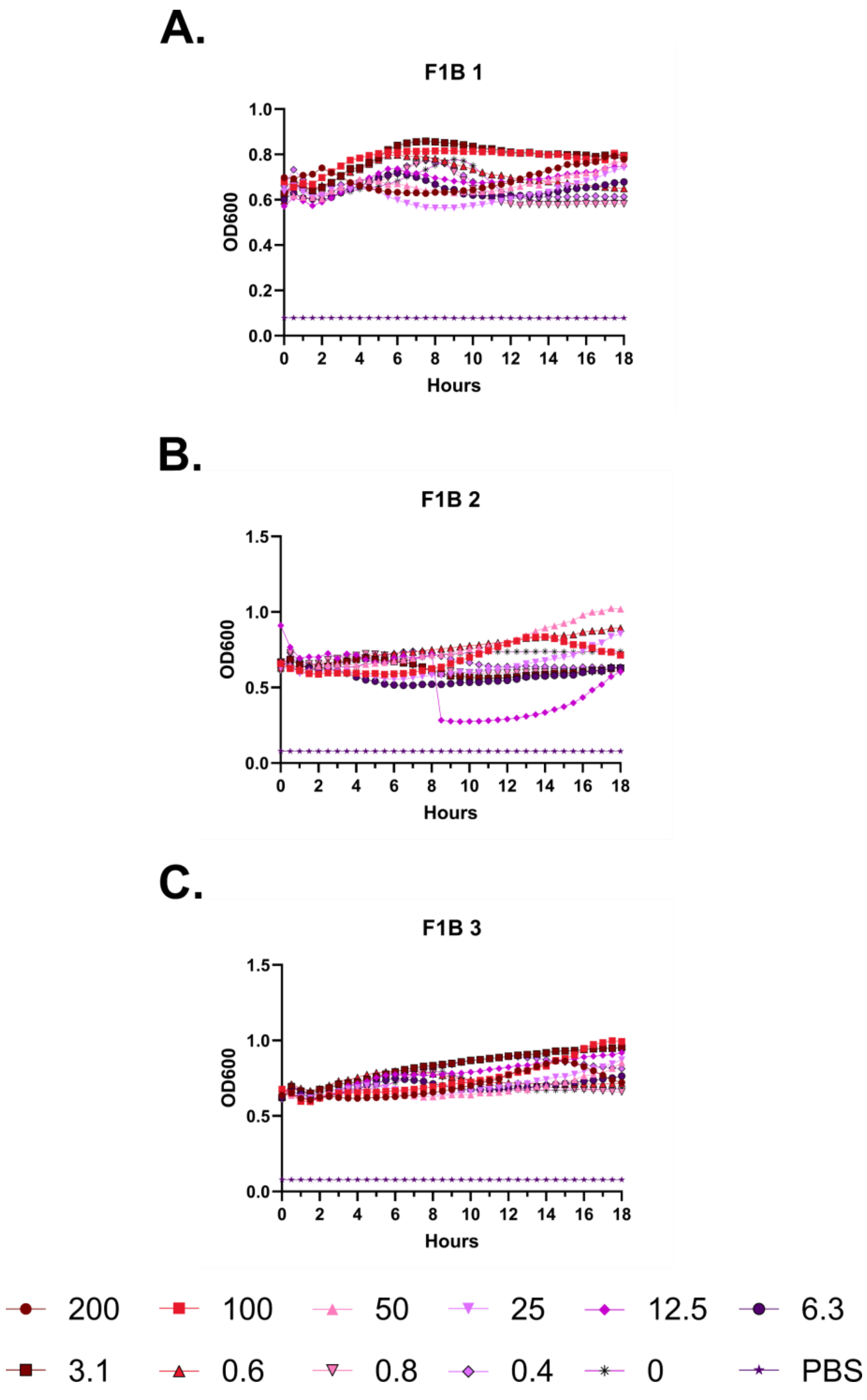
A.
F1B 1
OD600
Hours
B.
F1B 2
OD600
Hours
C.
F1B 3
OD600
Hours
200
100
50
25
12.5
6.3
3.1
0.6
0.8
0.4
0
PBS

**Figure S77 – Hemolysis over time with FF-14 F1B. A-C.** Each of three independent experiments showing trends in OD600 over time on incubation of defibrinated sheep blood with the indicated concentrations of peptide in μM at 37 °C. Decreasing OD600 indicates increasing hemolysis.

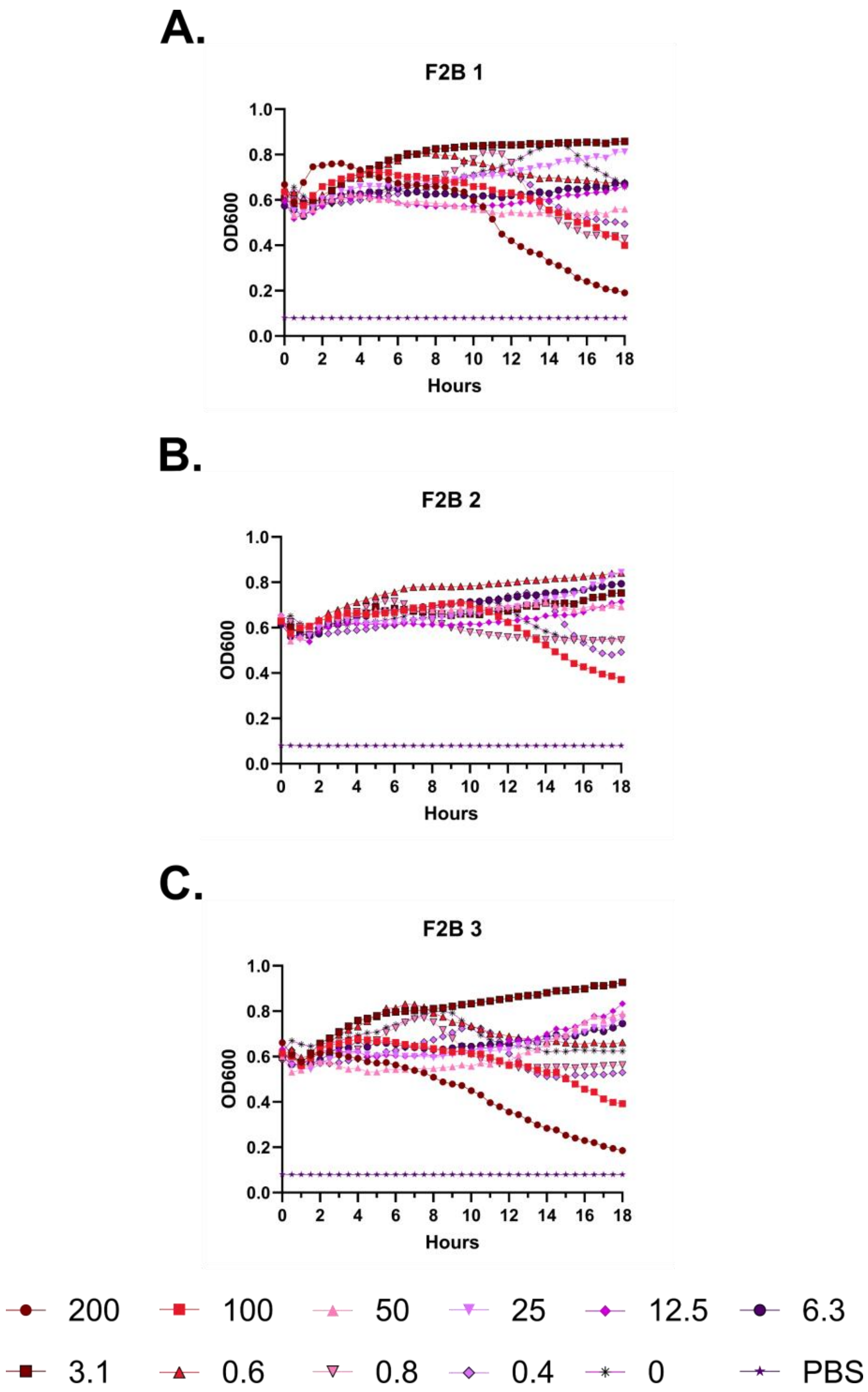
A.
F2B 1
OD600
Hours
B.
F2B 2
OD600
Hours
C.
F2B 3
OD600
Hours
200
100
50
25
12.5
6.3
3.1
0.6
0.8
0.4
0
PBS

**Figure S78 – Hemolysis over time with FF-14 F2B. A-C.** Each of three independent experiments showing trends in OD600 over time on incubation of defibrinated sheep blood with the indicated concentrations of peptide in μM at 37 °C. Decreasing OD600 indicates increasing hemolysis.

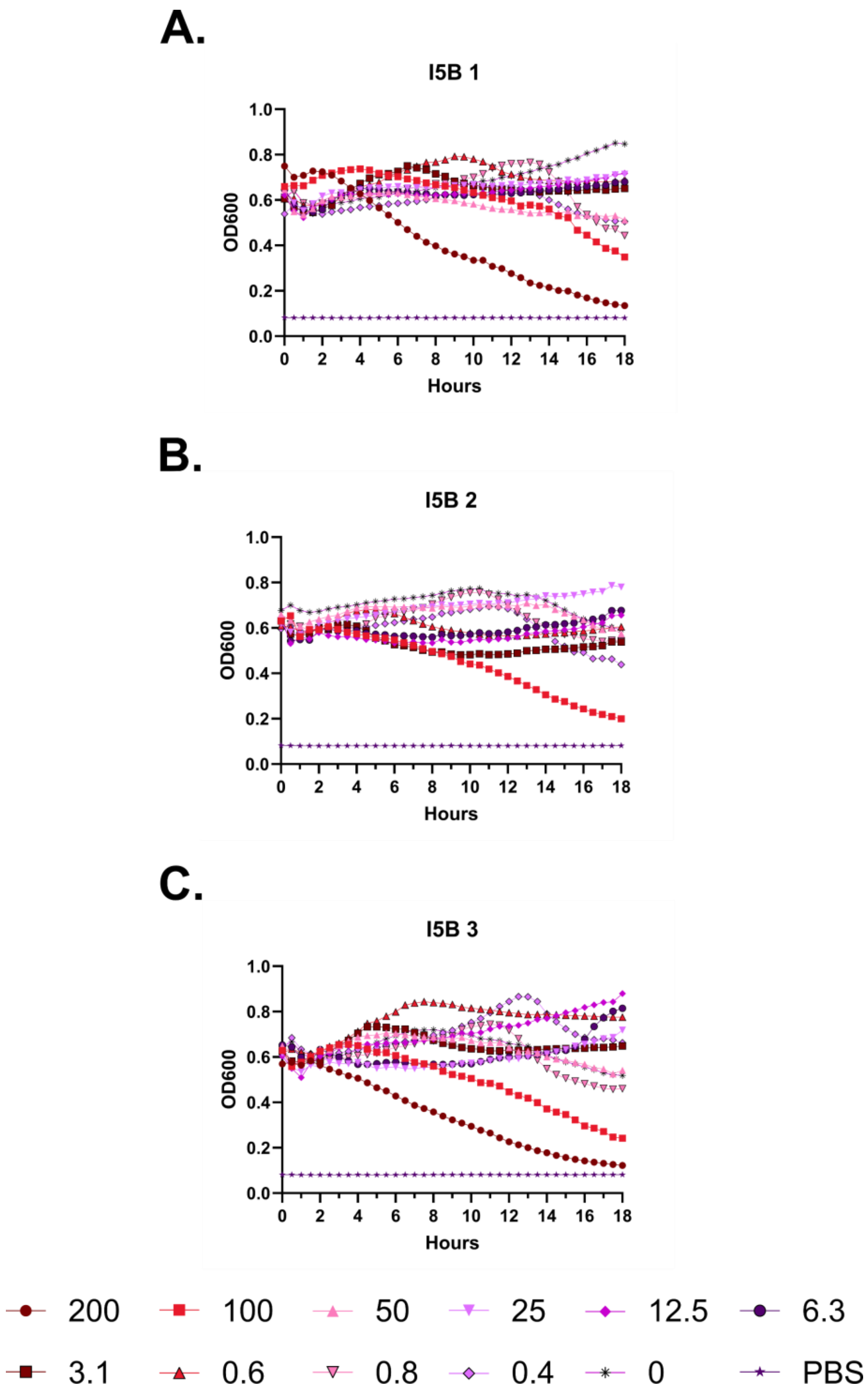
A.
I5B 1
OD600
Hours
B.
I5B 2
OD600
Hours
C.
I5B 3
OD600
Hours
200
100
50
25
12.5
6.3
3.1
0.6
0.8
0.4
0
PBS

**Figure S79 – Hemolysis over time with FF-14 I5B. A-C.** Each of three independent experiments showing trends in OD600 over time on incubation of defibrinated sheep blood with the indicated concentrations of peptide in μM at 37 °C. Decreasing OD600 indicates increasing hemolysis.

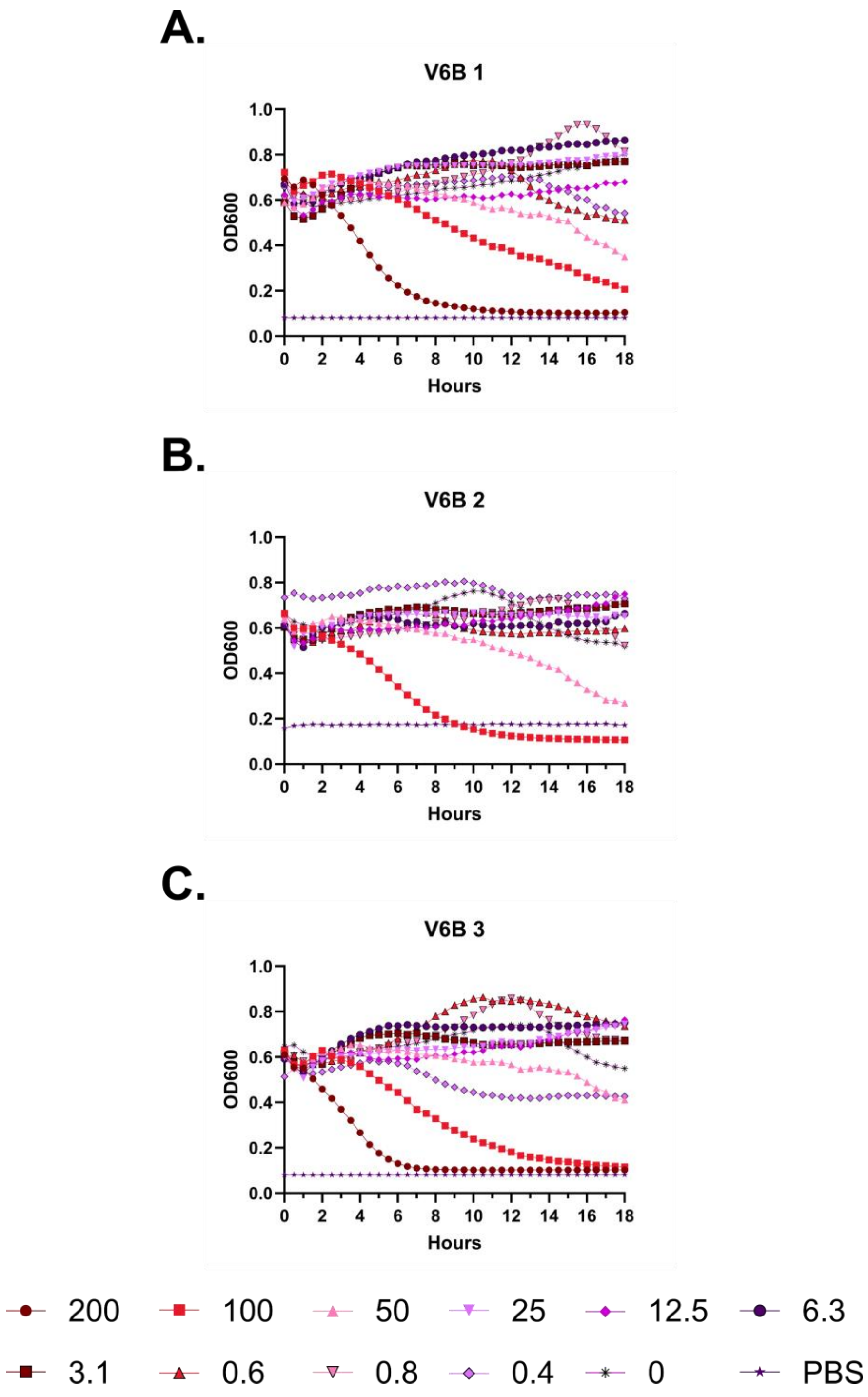
A.
V6B 1
OD600
Hours
B.
V6B 2
OD600
Hours
C.
V6B 3
OD600
Hours
200
100
50
25
12.5
6.3
3.1
0.6
0.8
0.4
0
PBS

**Figure S80 – Hemolysis over time with FF-14 V6B. A-C.** Each of three independent experiments showing trends in OD600 over time on incubation of defibrinated sheep blood with the indicated concentrations of peptide in μM at 37 °C. Decreasing OD600 indicates increasing hemolysis.

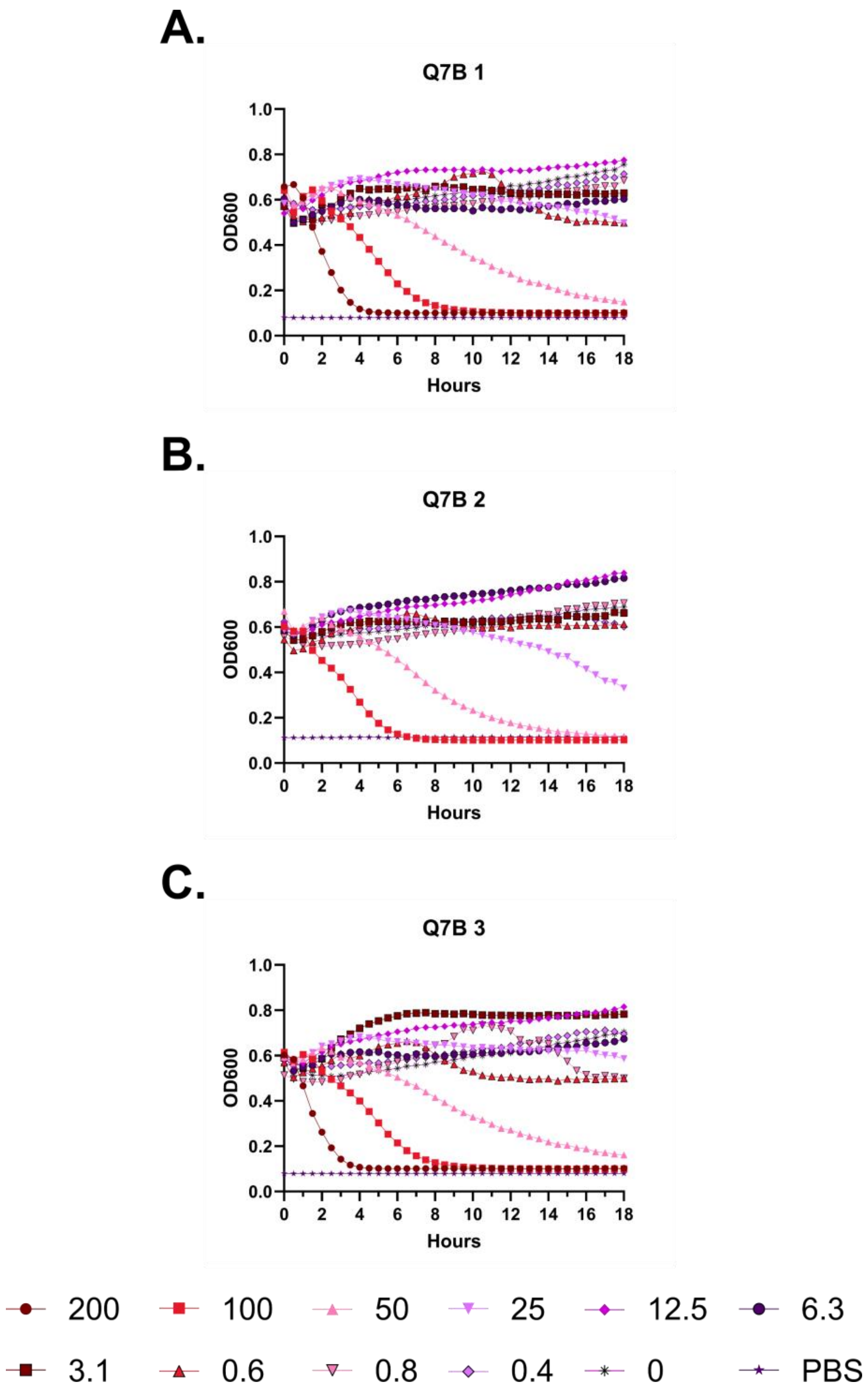
A.
Q7B 1
OD600
Hours
B.
Q7B 2
OD600
Hours
C.
Q7B 3
OD600
Hours
200
100
50
25
12.5
6.3
3.1
0.6
0.8
0.4
0
PBS

**Figure S81 – Hemolysis over time with FF-14 Q7B. A-C.** Each of three independent experiments showing trends in OD600 over time on incubation of defibrinated sheep blood with the indicated concentrations of peptide in μM at 37 °C. Decreasing OD600 indicates increasing hemolysis.

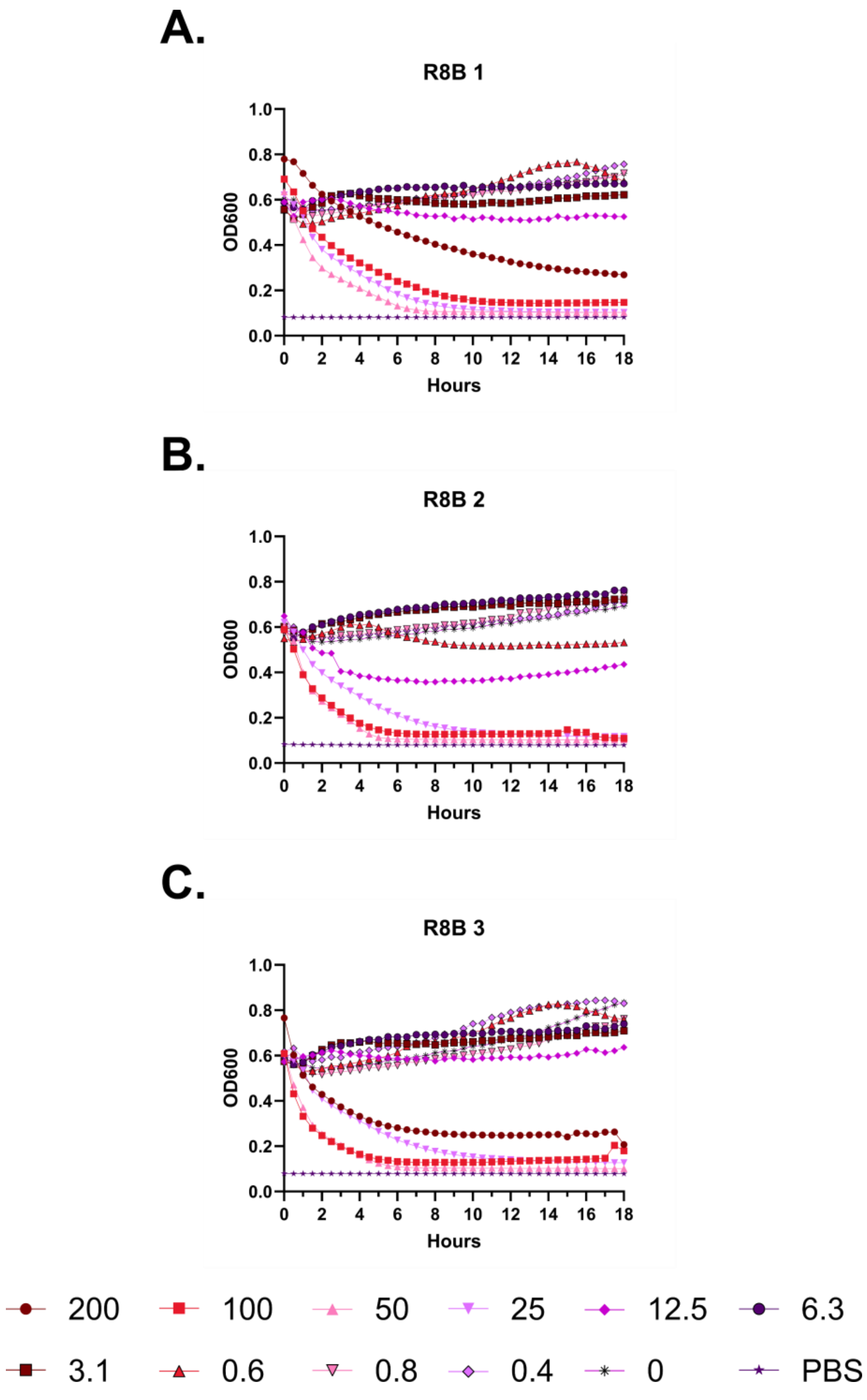
A.
R8B 1
OD600
Hours
B.
R8B 2
OD600
Hours
C.
R8B 3
OD600
Hours
200
100
50
25
12.5
6.3
3.1
0.6
0.8
0.4
0
PBS

**Figure S82 – Hemolysis over time with FF-14 R8B. A-C.** Each of three independent experiments showing trends in OD600 over time on incubation of defibrinated sheep blood with the indicated concentrations of peptide in μM at 37 °C. Decreasing OD600 indicates increasing hemolysis.

A.

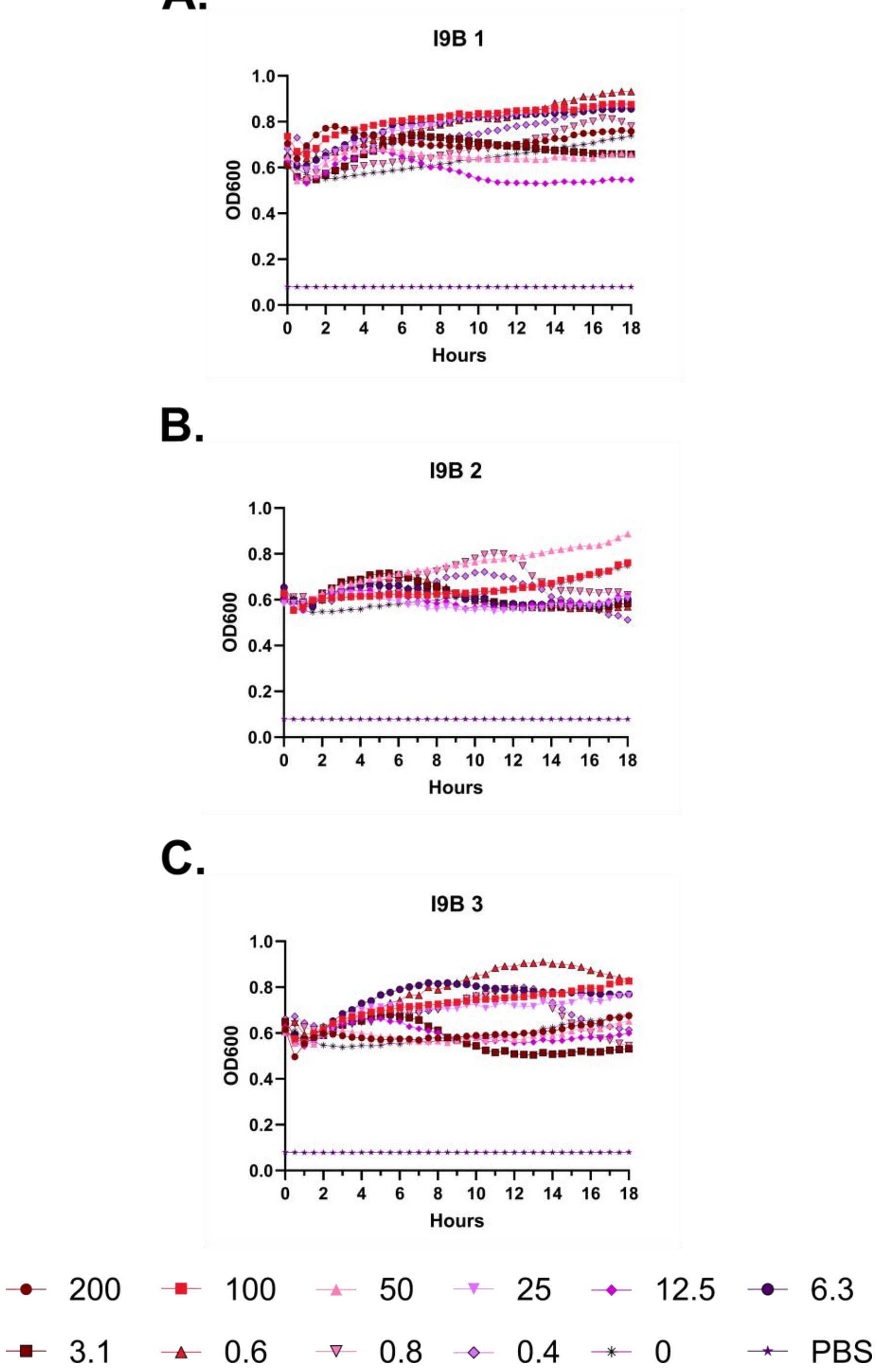
I9B 1
OD600
Hours
I9B 2
OD600
Hours
I9B 3
OD600
Hours
200
100
50
25
12.5
6.3
3.1
0.6
0.8
0.4
0
PBS

**Figure S83 – Hemolysis over time with FF-14 I9B. A-C.** Each of three independent experiments showing trends in OD600 over time on incubation of defibrinated sheep blood with the indicated concentrations of peptide in μM at 37 °C. Decreasing OD600 indicates increasing hemolysis.

A.

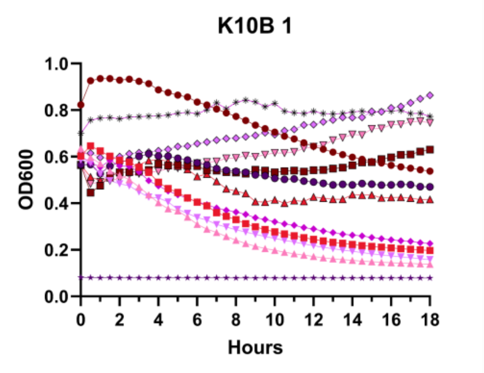
K10B 1
OD600
Hours

B.

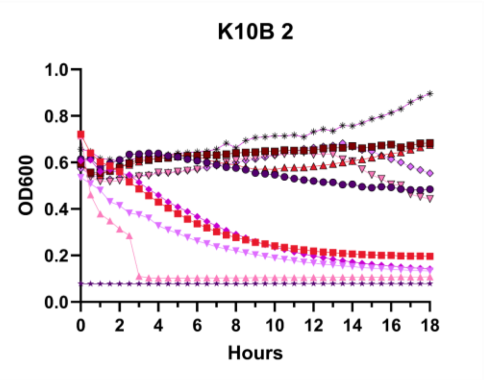
K10B 2
OD600
Hours

C.

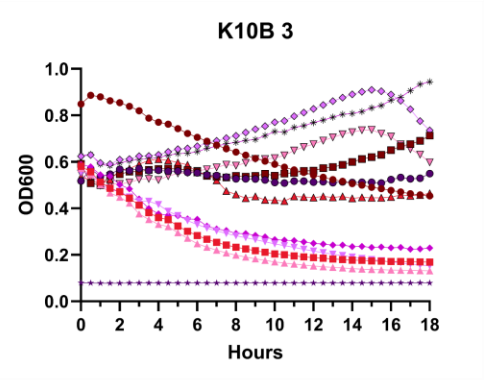
K10B 3
OD600
Hours

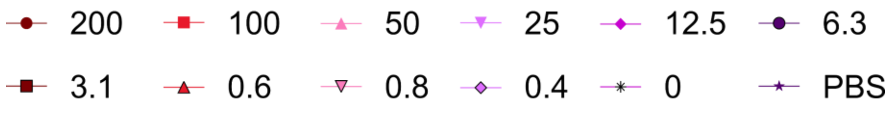
200
100
50
25
12.5
6.3
3.1
0.6
0.8
0.4
0
PBS

**Figure S84 – Hemolysis over time with FF-14 K10B. A-C.** Each of three independent experiments showing trends in OD600 over time on incubation of defibrinated sheep blood with the indicated concentrations of peptide in μM at 37 °C. Decreasing OD600 indicates increasing hemolysis.

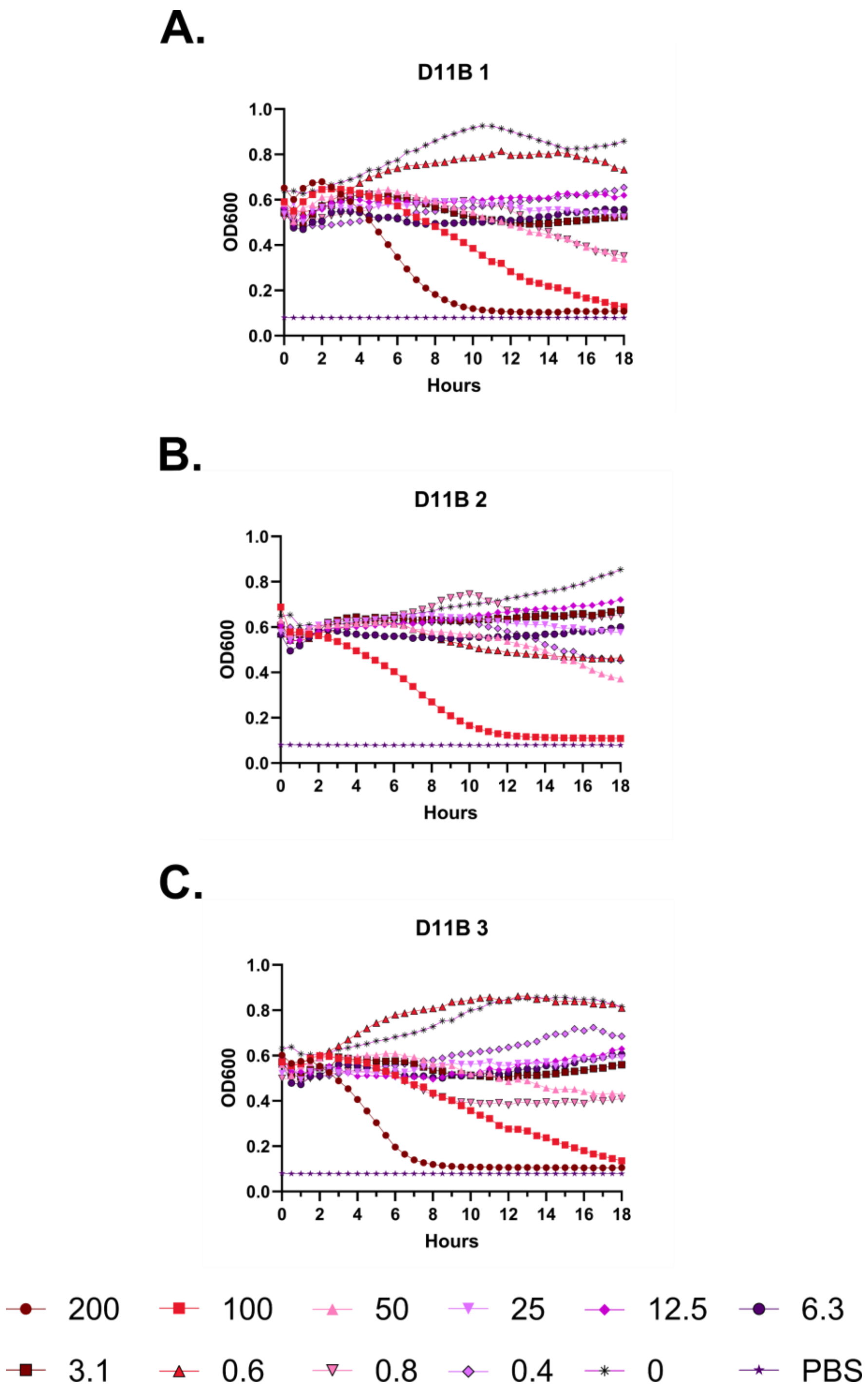
A.
D11B 1
OD600
Hours
B.
D11B 2
OD600
Hours
C.
D11B 3
OD600
Hours
200
100
50
25
12.5
6.3
3.1
0.6
0.8
0.4
0
PBS

**Figure S85 – Hemolysis over time with FF-14 D11B. A-C.** Each of three independent experiments showing trends in OD600 over time on incubation of defibrinated sheep blood with the indicated concentrations of peptide in μM at 37 °C. Decreasing OD600 indicates increasing hemolysis.

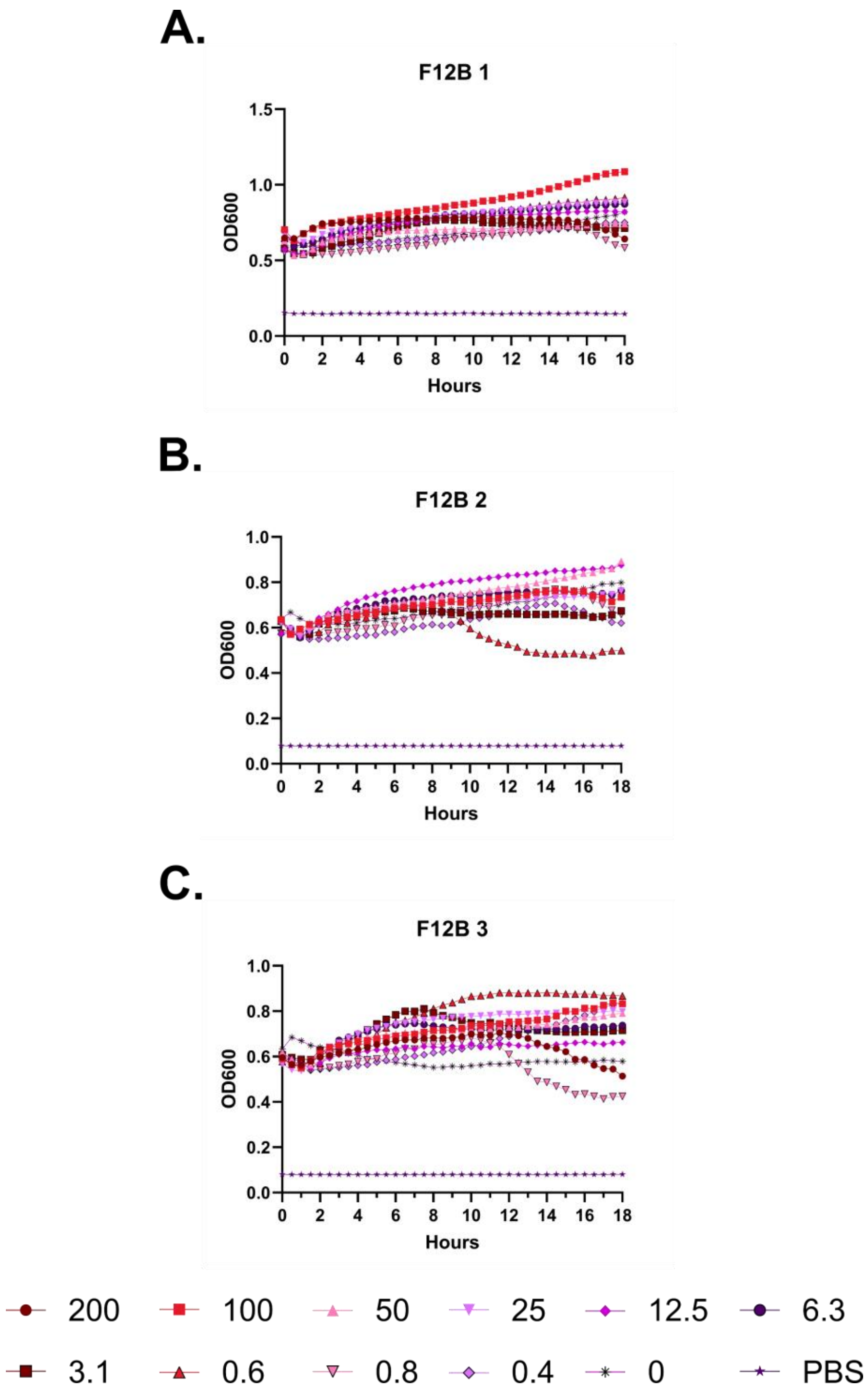
A.
F12B 1
OD600
Hours
B.
F12B 2
OD600
Hours
C.
F12B 3
OD600
Hours
200
100
50
25
12.5
6.3
3.1
0.6
0.8
0.4
0
PBS

**Figure S86 – Hemolysis over time with FF-14 F12B. A-C.** Each of three independent experiments showing trends in OD600 over time on incubation of defibrinated sheep blood with the indicated concentrations of peptide in μM at 37 °C. Decreasing OD600 indicates increasing hemolysis.

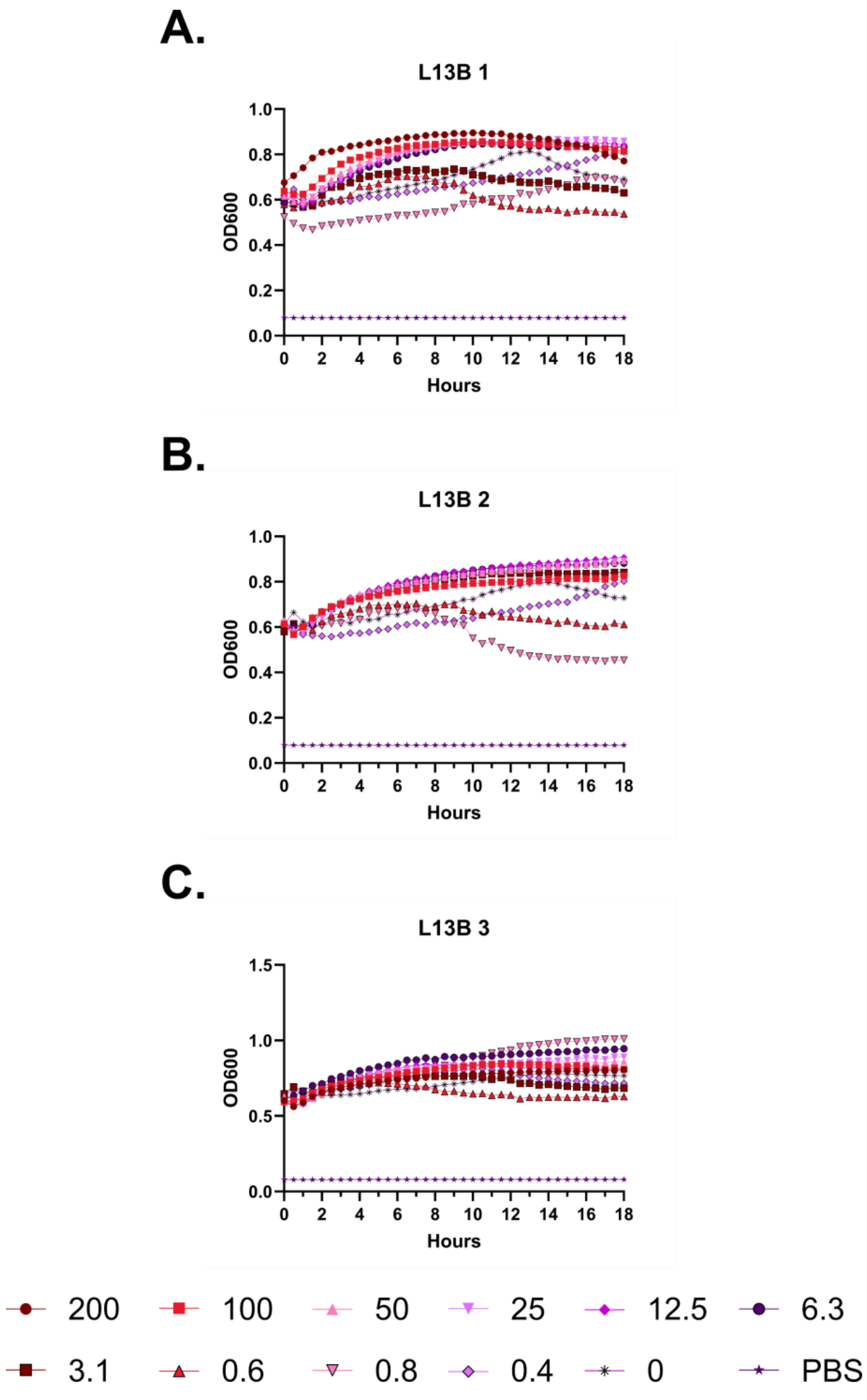
A.
L13B 1
OD600
Hours
B.
L13B 2
OD600
Hours
C.
L13B 3
OD600
Hours
200
100
50
25
12.5
6.3
3.1
0.6
0.8
0.4
0
PBS

**Figure S87 – Hemolysis over time with FF-14 L13B. A-C.** Each of three independent experiments showing trends in OD600 over time on incubation of defibrinated sheep blood with the indicated concentrations of peptide in μM at 37 °C. Decreasing OD600 indicates increasing hemolysis.

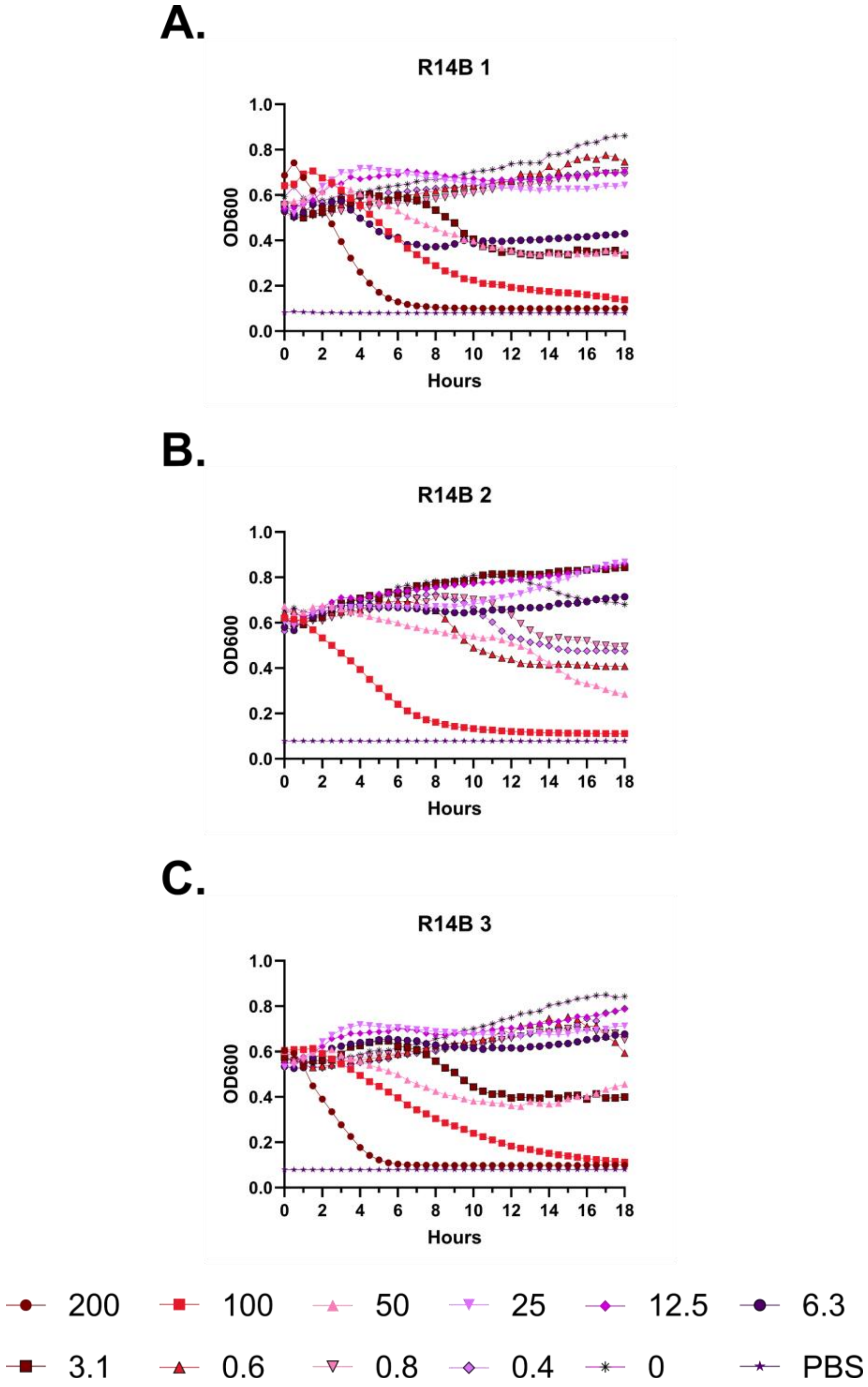
A.
R14B 1
OD600
Hours
B.
R14B 2
OD600
Hours
C.
R14B 3
OD600
Hours
200
100
50
25
12.5
6.3
3.1
0.6
0.8
0.4
0
PBS

**Figure S88 – Hemolysis over time with FF-14 R14B. A-C.** Each of three independent experiments showing trends in OD600 over time on incubation of defibrinated sheep blood with the indicated concentrations of peptide in μM at 37 °C. Decreasing OD600 indicates increasing hemolysis.

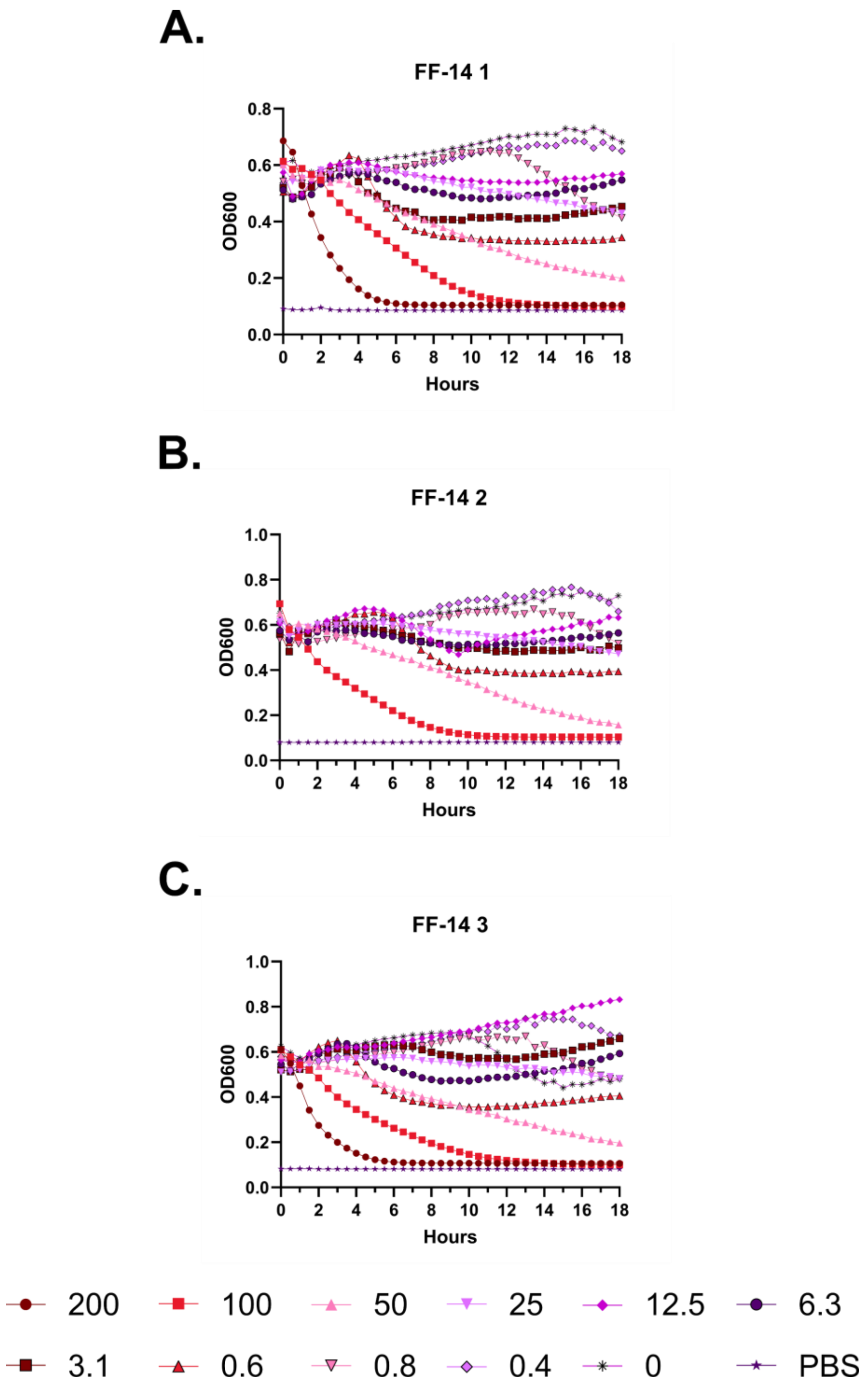
A.
FF-14 1
OD600
Hours
B.
FF-14 2
OD600
Hours
C.
FF-14 3
OD600
Hours
200
100
50
25
12.5
6.3
3.1
0.6
0.8
0.4
0
PBS

**Figure S89 – Hemolysis over time with FF-14. A-C.** Each of three independent experiments showing trends in OD600 over time on incubation of defibrinated sheep blood with the indicated concentrations of peptide in μM at 37 °C. Decreasing OD600 indicates increasing hemolysis.

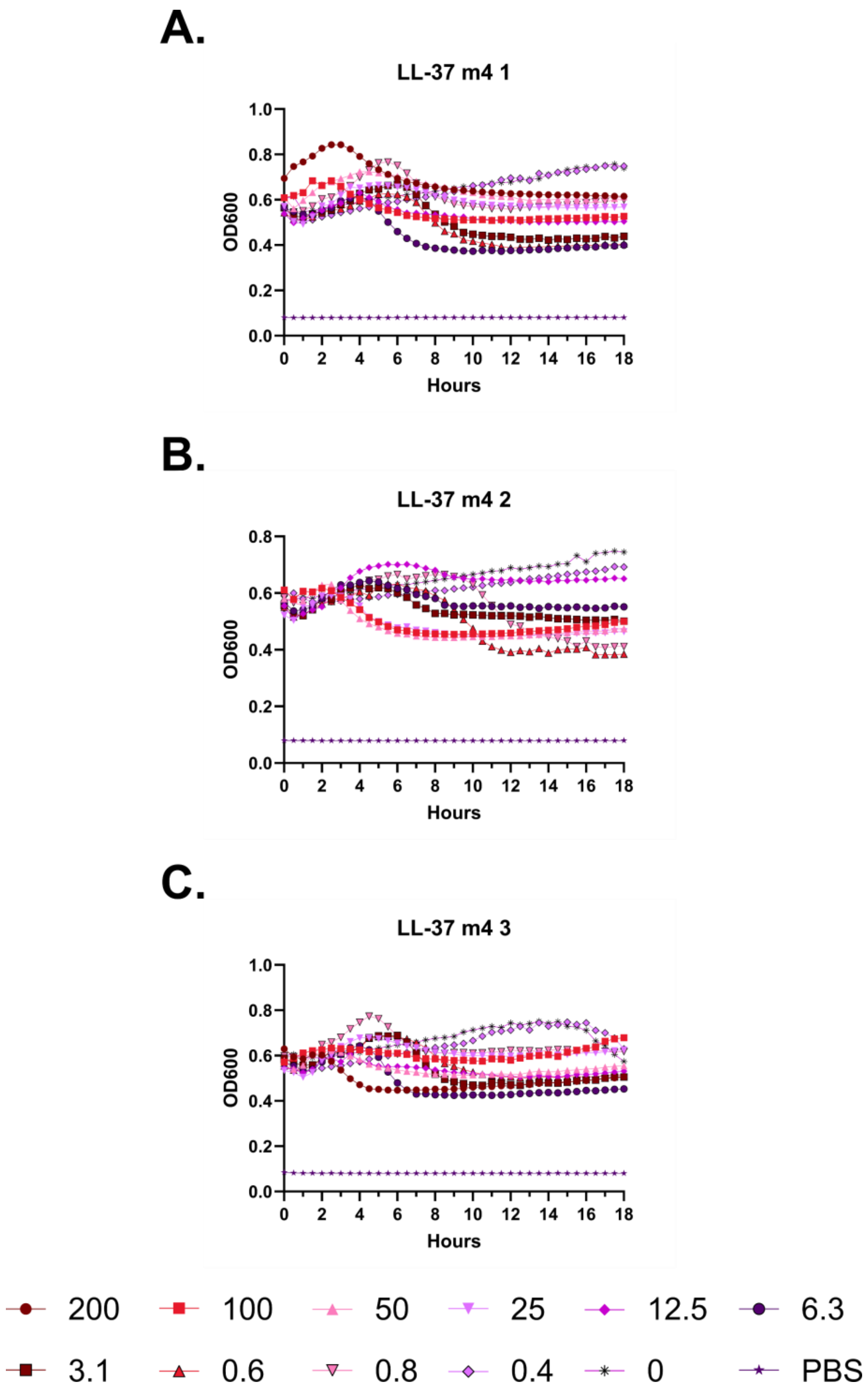
A.
LL-37 m4 1
OD600
Hours
B.
LL-37 m4 2
OD600
Hours
C.
LL-37 m4 3
OD600
Hours
200
100
50
25
12.5
6.3
3.1
0.6
0.8
0.4
0
PBS

**Figure S90 – Hemolysis over time with LL-37 m4. A-C.** Each of three independent experiments showing trends in OD600 over time on incubation of defibrinated sheep blood with the indicated concentrations of peptide in μM at 37 °C. Decreasing OD600 indicates increasing hemolysis.

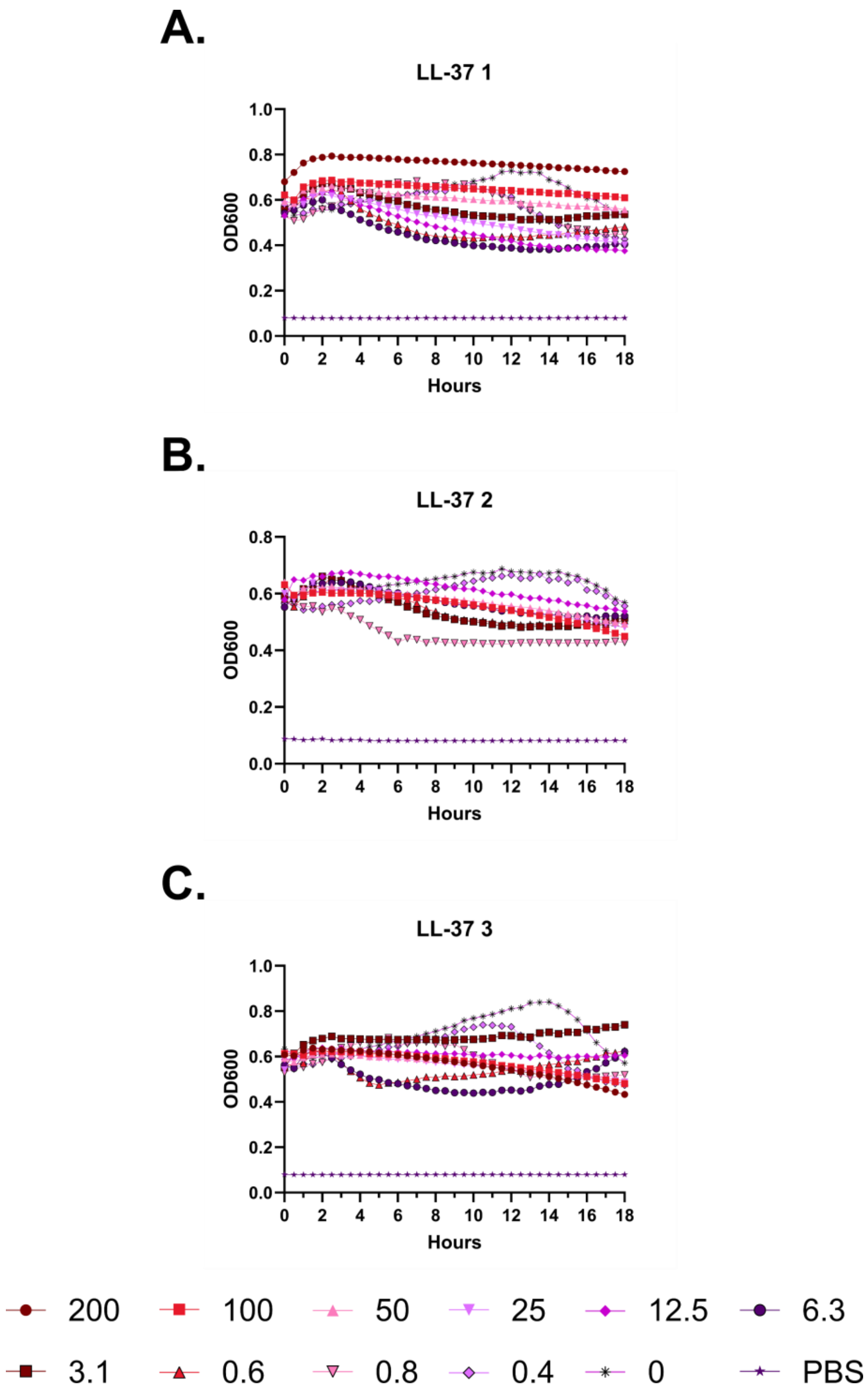
A.
LL-37 1
OD600
Hours
B.
LL-37 2
OD600
Hours
C.
LL-37 3
OD600
Hours
200
100
50
25
12.5
6.3
3.1
0.6
0.8
0.4
0
PBS

**Figure S91 – Hemolysis over time with LL-37. A-C.** Each of three independent experiments showing trends in OD600 over time on incubation of defibrinated sheep blood with the indicated concentrations of peptide in μM at 37 °C. Decreasing OD600 indicates increasing hemolysis.

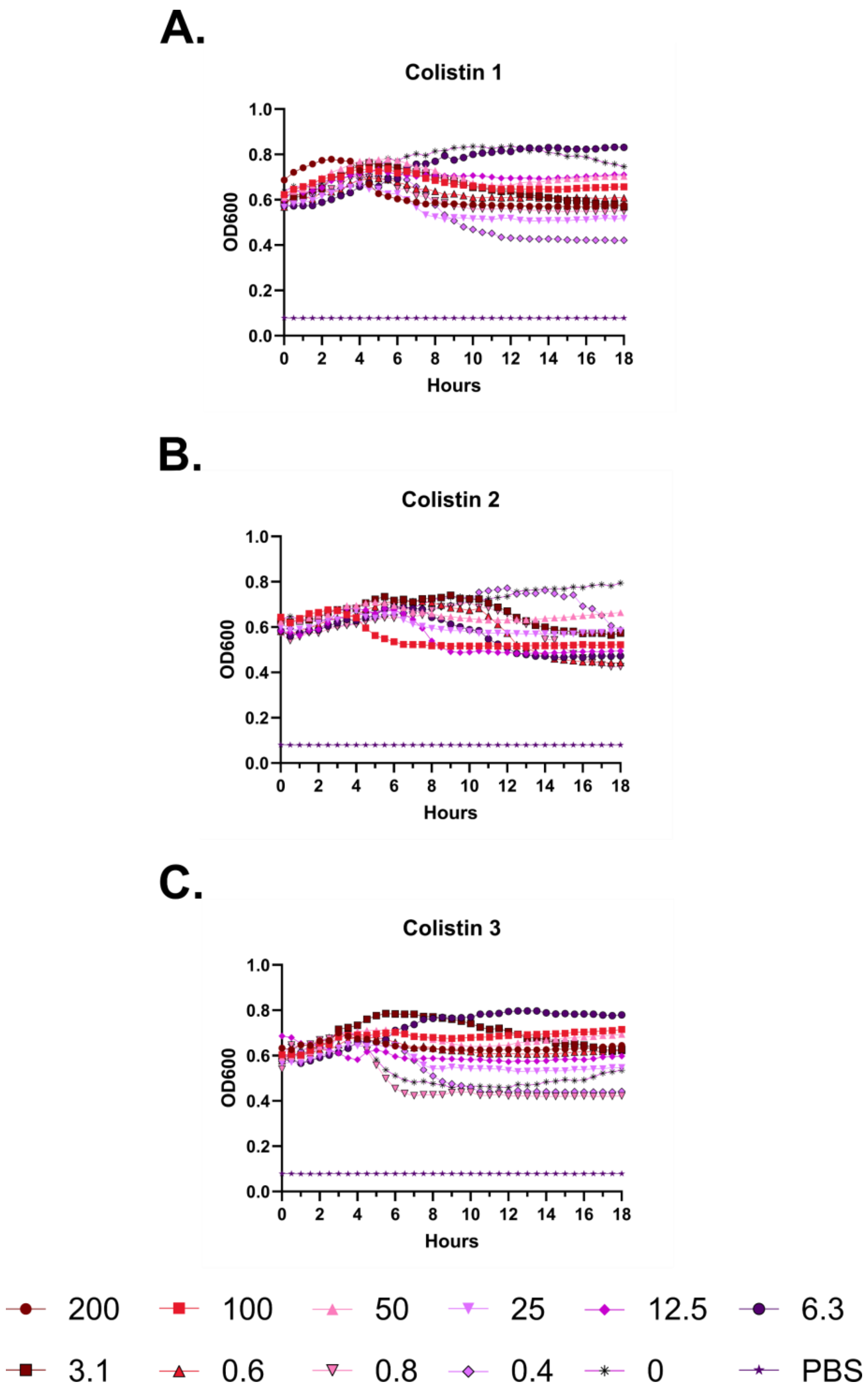
A.
Colistin 1
OD600
1.0
0.8
0.6
0.4
0.2
0.0
0
2
4
6
8
10
12
14
16
18
Hours
B.
Colistin 2
OD600
Hours
C.
Colistin 3
OD600
Hours
200
100
50
25
12.5
6.3
3.1
0.6
0.8
0.4
0
PBS

**Figure S92 – Hemolysis over time with colistin. A-C.** Each of three independent experiments showing trends in OD600 over time on incubation of defibrinated sheep blood with the indicated concentrations of peptide in μM at 37 °C. Decreasing OD600 indicates increasing hemolysis.

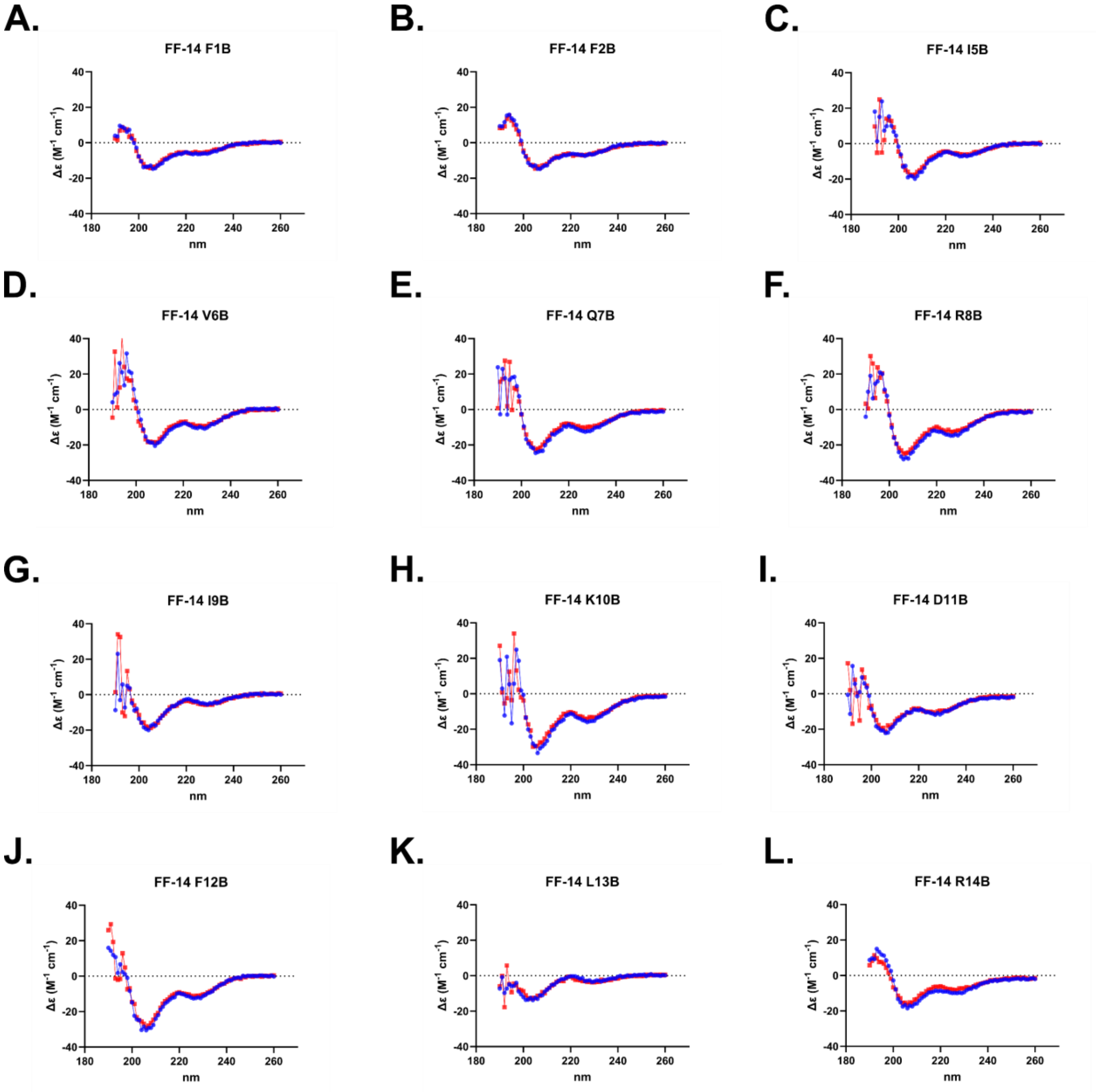


**M.**

| | % Helix | |
|---|---|---|
| | 20 °C | 37 °C |
| FF-14 F1B | 63.4 | 51.7 |
| FF-14 F2B | 74.5 | 74.2 |
| FF-14 I5B | 97.3 | 91.6 |
| FF-14 V6B | 100 | 100 |
| FF-14 Q7B | 96.1 | 96.5 |
| FF-14 R8B | 99.9 | 100 |
| FF-14 I9B | 49 | 54 |
| FF-14 K10B | 81.1 | 90.3 |
| FF-14 D11B | 77.1 | 68.7 |
| FF-14 F12B | 90.8 | 84.7 |
| FF-14 L13B | 14.4 | 18.2 |
| FF-14 R14B | 88.8 | 74.1 |

**Figure S93 – Circular dichroism spectroscopy of Figure 3 mutants. A-L.** CD spectra are displayed for each mutant collected from 190-260 nm at both 20 °C (Blue) and 37 °C (Red). **M.** Quantification of helical content as determined by BeStSel for each mutant.

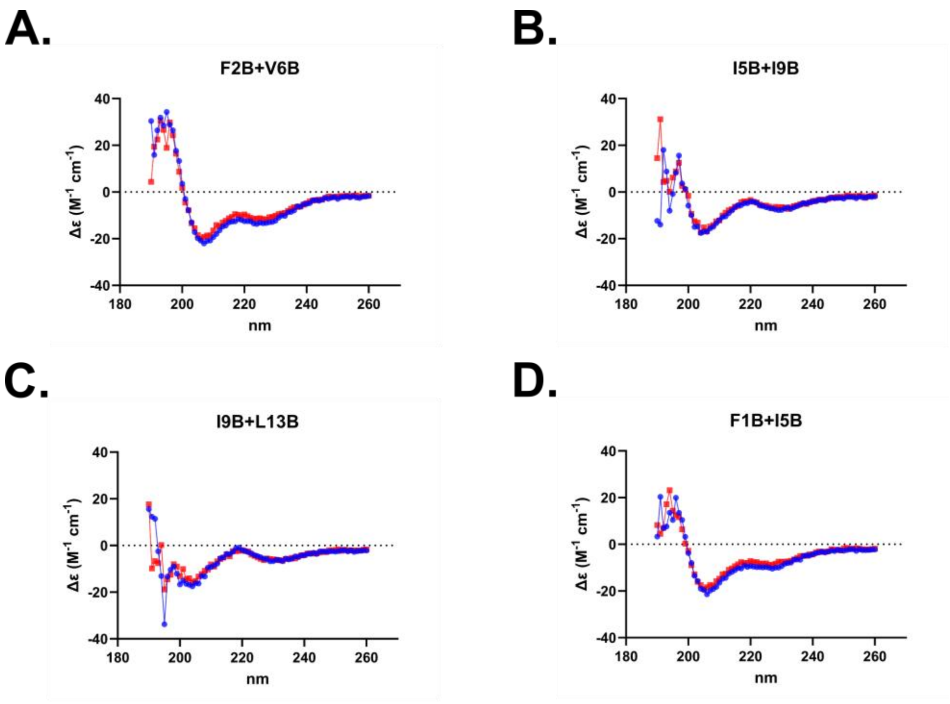


E.

| | % Helix | |
|---|---|---|
| | **20 °C** | **37 °C** |
| F2B+V6B | 100 | 100 |
| I5B+I9B | 78 | 72.3 |
| I9B+L13B | 33.1 | 32.8 |
| F1B+I5B | 98.3 | 100 |

**Figure S94 – Circular dichroism spectroscopy of double Aib mutants. A-D.** Circular dichroism spectra are displayed for each mutant collected from 190-260 nm at both 20 °C (Blue) and 37 °C (Red). **E.** Quantification of helical content as determined by BeStSel for each mutant.

A.

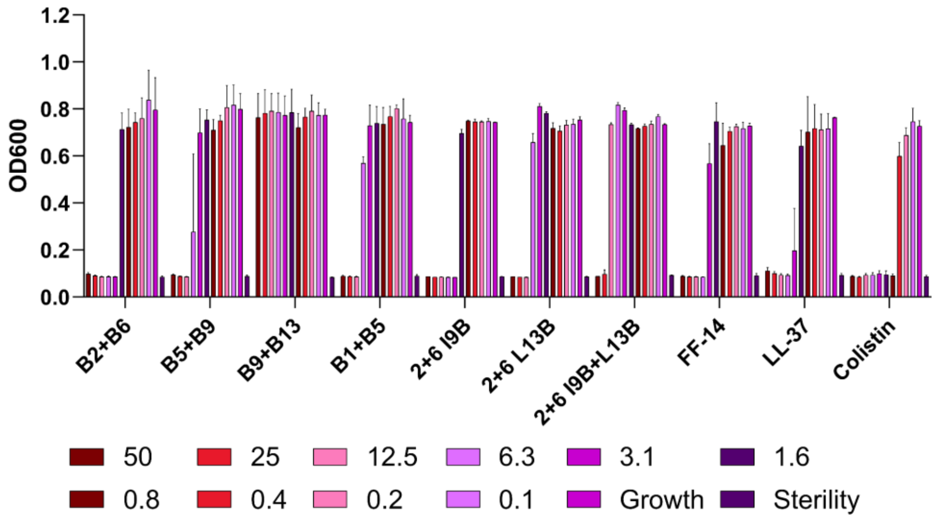
OD600
1.2
1.0
0.8
0.6
0.4
0.2
0.0
B2+B6
B5+B9
B9+B13
B1+B5
2+6 I9B
2+6 L13B
2+6 I9B+L13B
FF-14
LL-37
Colistin
50
25
12.5
6.3
3.1
1.6
0.8
0.4
0.2
0.1
Growth
Sterility

B.

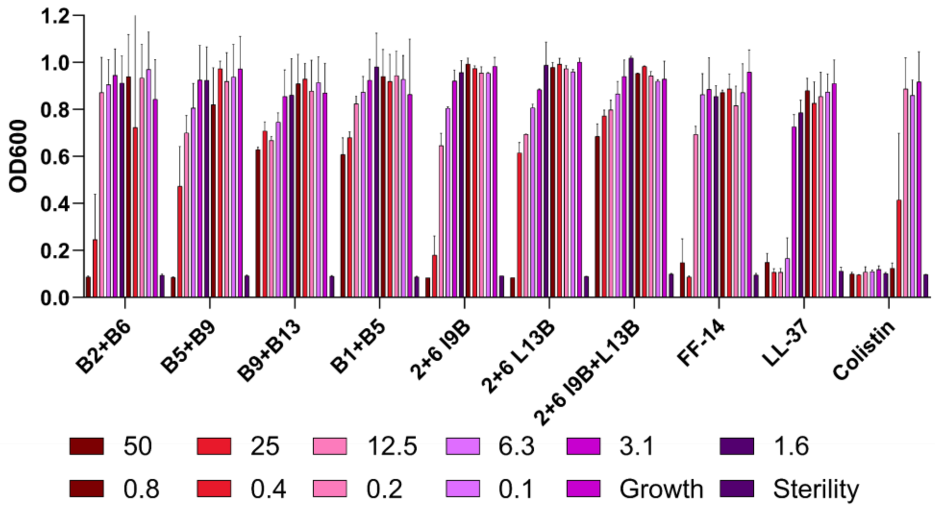
OD600
1.2
1.0
0.8
0.6
0.4
0.2
0.0
B2+B6
B5+B9
B9+B13
B1+B5
2+6 I9B
2+6 L13B
2+6 I9B+L13B
FF-14
LL-37
Colistin
50
25
12.5
6.3
3.1
1.6
0.8
0.4
0.2
0.1
Growth
Sterility

C.

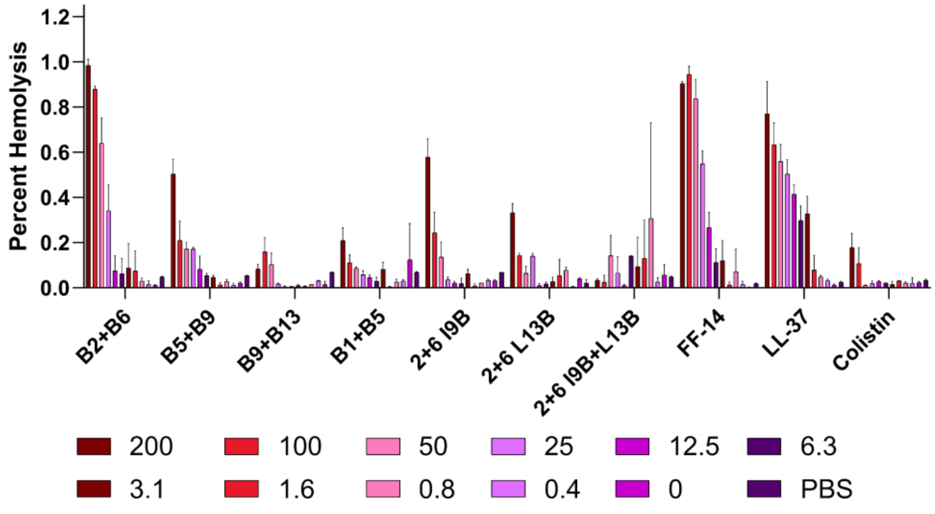
Percent Hemolysis
1.2
1.0
0.8
0.6
0.4
0.2
0.0
B2+B6
B5+B9
B9+B13
B1+B5
2+6 I9B
2+6 L13B
2+6 I9B+L13B
FF-14
LL-37
Colistin
200
100
50
25
12.5
6.3
3.1
1.6
0.8
0.4
0
PBS

**Figure S95 – Histogram depiction of data from Figure 4. A.** Endpoint activity of each peptide against *E. coli*; measures are raw absorbance at 600 nm at 18 hours. **B.** Endpoint activity of each peptide against *P. aeruginosa*. **C.** Hemolytic activity of each peptide; measures are absorbance at 414 nm scaled to a freeze-thaw positive control set at 100% hemolysis. Concentrations throughout are shown are in μM.

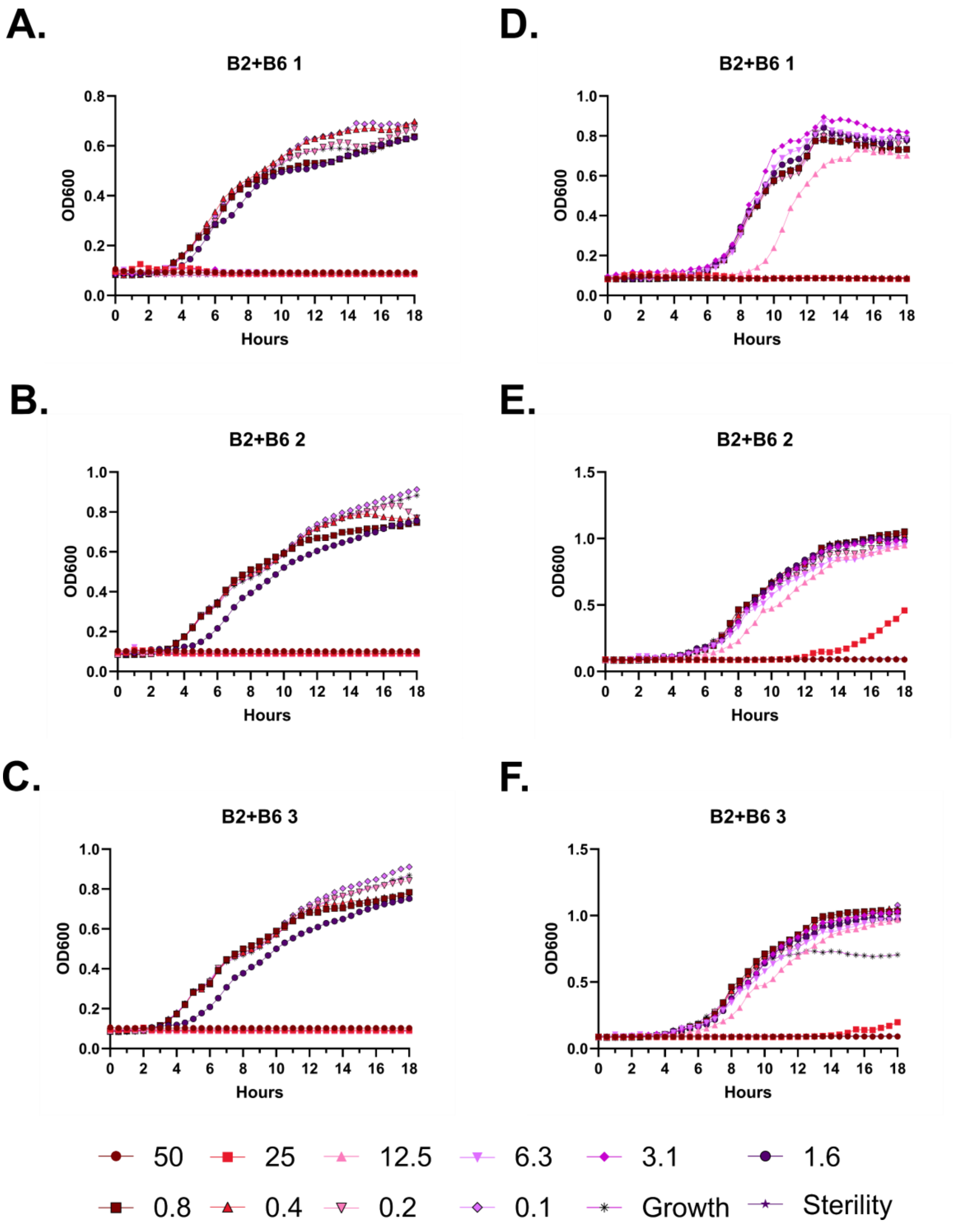
A.
B2+B6 1
OD600
Hours
D.
B2+B6 1
OD600
Hours
B.
B2+B6 2
OD600
Hours
E.
B2+B6 2
OD600
Hours
C.
B2+B6 3
OD600
Hours
F.
B2+B6 3
OD600
Hours
50
25
12.5
6.3
3.1
1.6
0.8
0.4
0.2
0.1
Growth
Sterility

**Figure S96 – Antimicrobial susceptibility testing with FF-14 B2+B6. A-C.** Growth of *E. coli* in the presence of the indicated peptide concentrations given in μM as monitored by OD600 at 37 °C over 18 hours. Each panel indicates an independent experiment. **D-F.** As in **A-C**, but with *P. aeruginosa.*

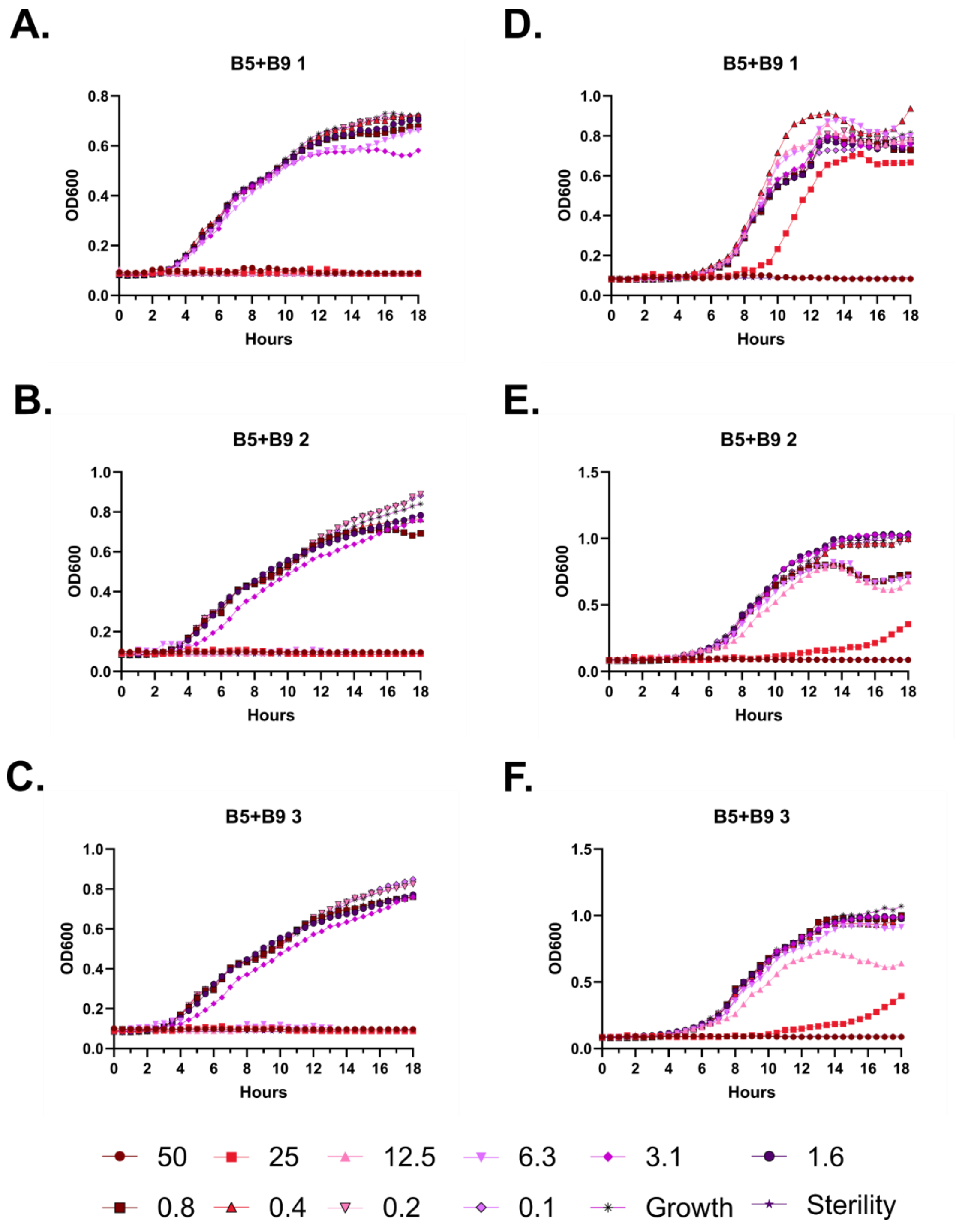

A.
B5+B9 1
B.
B5+B9 2
C.
B5+B9 3
D.
B5+B9 1
E.
B5+B9 2
F.
B5+B9 3
OD600
Hours
50
25
12.5
6.3
3.1
1.6
0.8
0.4
0.2
0.1
Growth
Sterility

**Figure S97 – Antimicrobial susceptibility testing with FF-14 B5+B9. A-C.** Growth of *E. coli* in the presence of the indicated peptide concentrations given in μM as monitored by OD600 at 37 °C over 18 hours. Each panel indicates an independent experiment. **D-F.** As in **A-C**, but with *P. aeruginosa.*

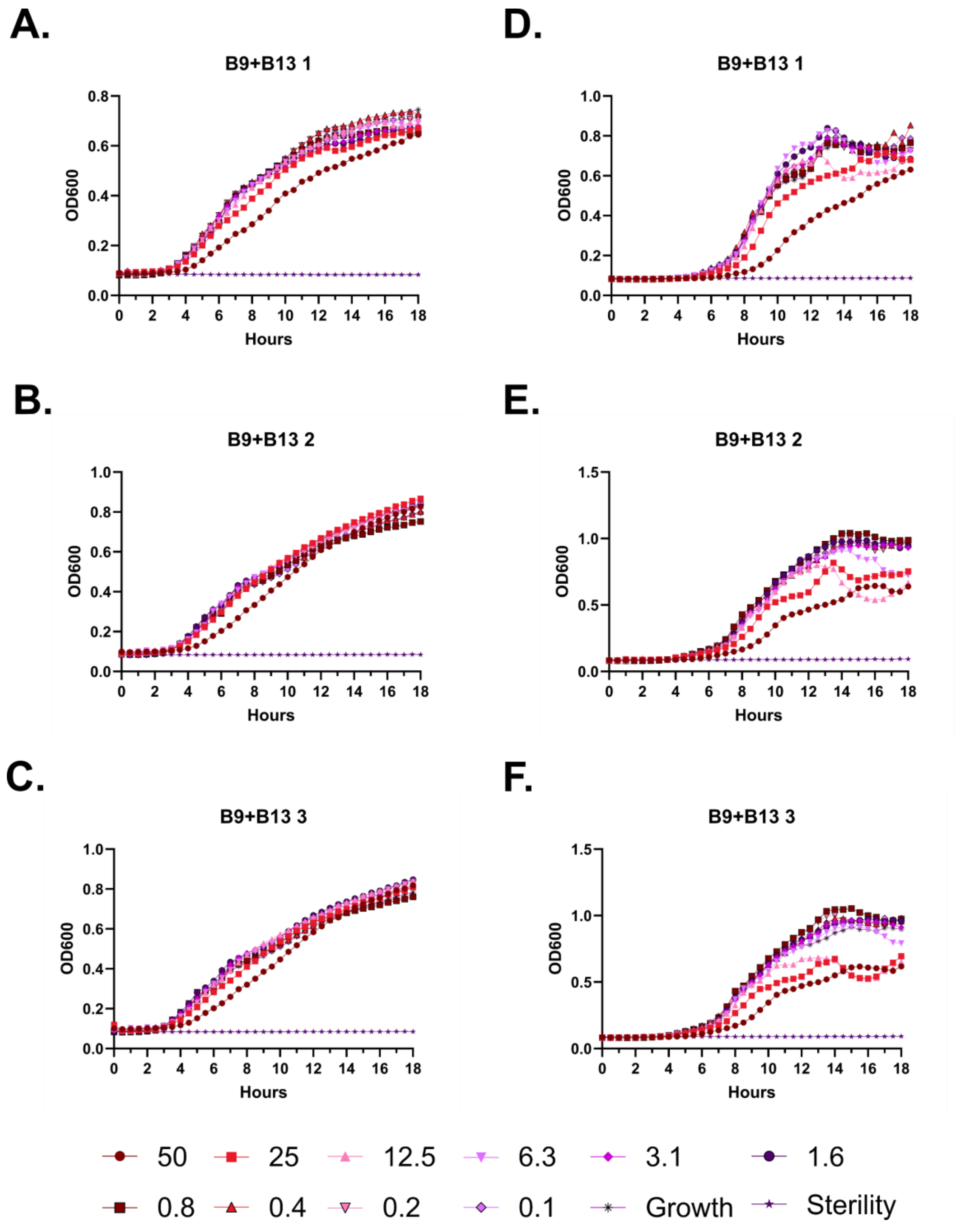
A.
B9+B13 1
D.
B9+B13 1
B.
B9+B13 2
E.
B9+B13 2
C.
B9+B13 3
F.
B9+B13 3
OD600
Hours
50
25
12.5
6.3
3.1
1.6
0.8
0.4
0.2
0.1
Growth
Sterility

**Figure S98 – Antimicrobial susceptibility testing with FF-14 B9+B13. A-C.** Growth of *E. coli* in the presence of the indicated peptide concentrations given in µM as monitored by OD600 at 37 °C over 18 hours. Each panel indicates an independent experiment. **D-F.** As in **A-C**, but with *P. aeruginosa.*

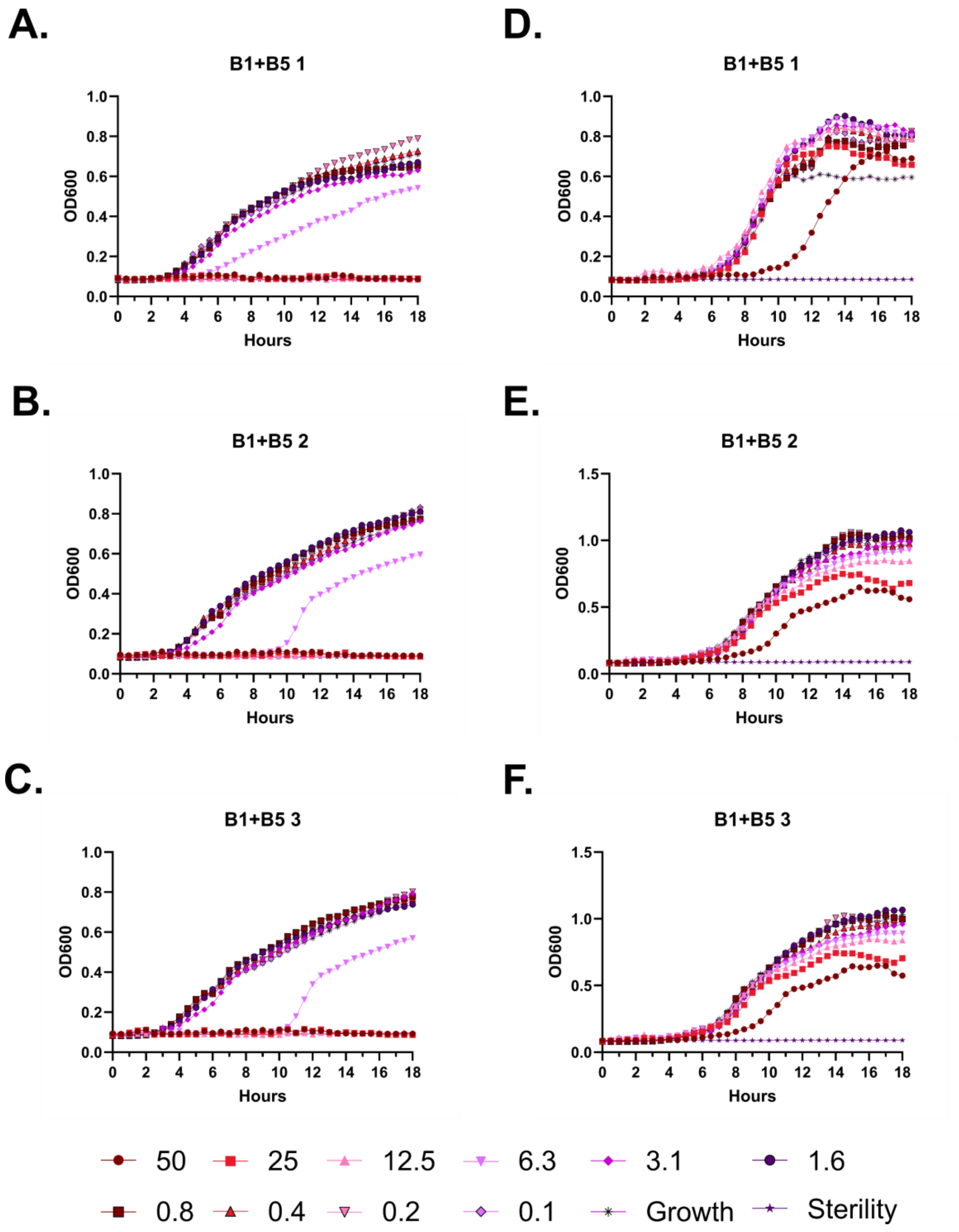

A.
B1+B5 1
OD600
Hours
D.
B1+B5 1
OD600
Hours
B.
B1+B5 2
OD600
Hours
E.
B1+B5 2
OD600
Hours
C.
B1+B5 3
OD600
Hours
F.
B1+B5 3
OD600
Hours
50
25
12.5
6.3
3.1
1.6
0.8
0.4
0.2
0.1
Growth
Sterility

**Figure S99 – Antimicrobial susceptibility testing with FF-14 B1+B5. A-C.** Growth of *E. coli* in the presence of the indicated peptide concentrations given in μM as monitored by OD600 at 37 °C over 18 hours. Each panel indicates an independent experiment. **D-F.** As in **A-C**, but with *P. aeruginosa.*

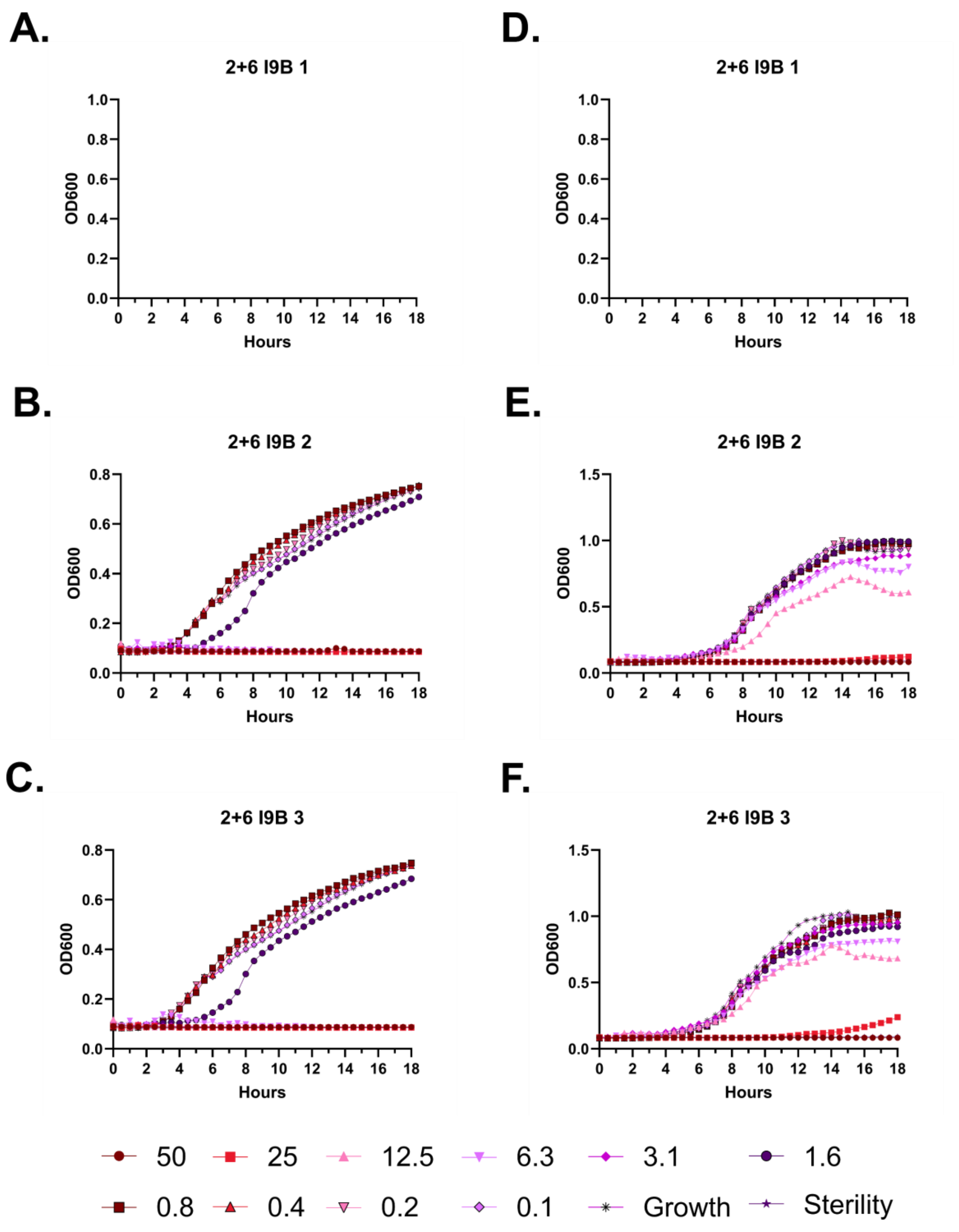
A.
2+6 I9B 1
B.
2+6 I9B 2
C.
2+6 I9B 3
D.
2+6 I9B 1
E.
2+6 I9B 2
F.
2+6 I9B 3
OD600
Hours
50
25
12.5
6.3
3.1
1.6
0.8
0.4
0.2
0.1
Growth
Sterility

**Figure S100 – Antimicrobial susceptibility testing with 2+6 I9B. A-C.** Growth of *E. coli* in the presence of the indicated peptide concentrations given in μM as monitored by OD600 at 37 °C over 18 hours. Each panel indicates an independent experiment. **D-F.** As in **A-C**, but with *P. aeruginosa.*

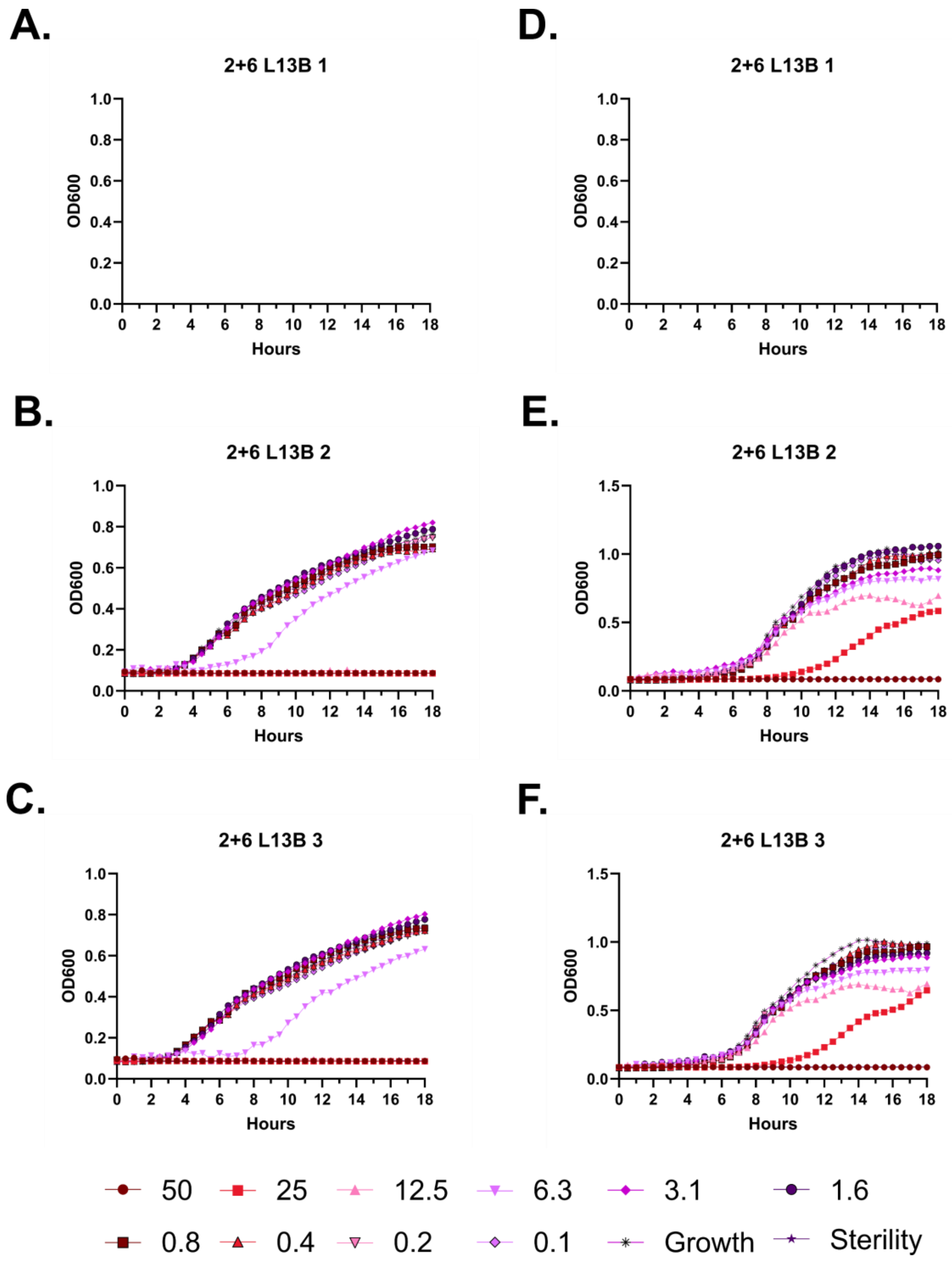
A.
2+6 L13B 1
OD600
Hours
D.
2+6 L13B 1
OD600
Hours
B.
2+6 L13B 2
OD600
Hours
E.
2+6 L13B 2
OD600
Hours
C.
2+6 L13B 3
OD600
Hours
F.
2+6 L13B 3
OD600
Hours
50
25
12.5
6.3
3.1
1.6
0.8
0.4
0.2
0.1
Growth
Sterility

**Figure S101 – Antimicrobial susceptibility testing with 2+6 L13B. A-C.** Growth of *E. coli* in the presence of the indicated peptide concentrations given in μM as monitored by OD600 at 37 °C over 18 hours. Each panel indicates an independent experiment. **D-F.** As in **A-C**, but with *P. aeruginosa.*

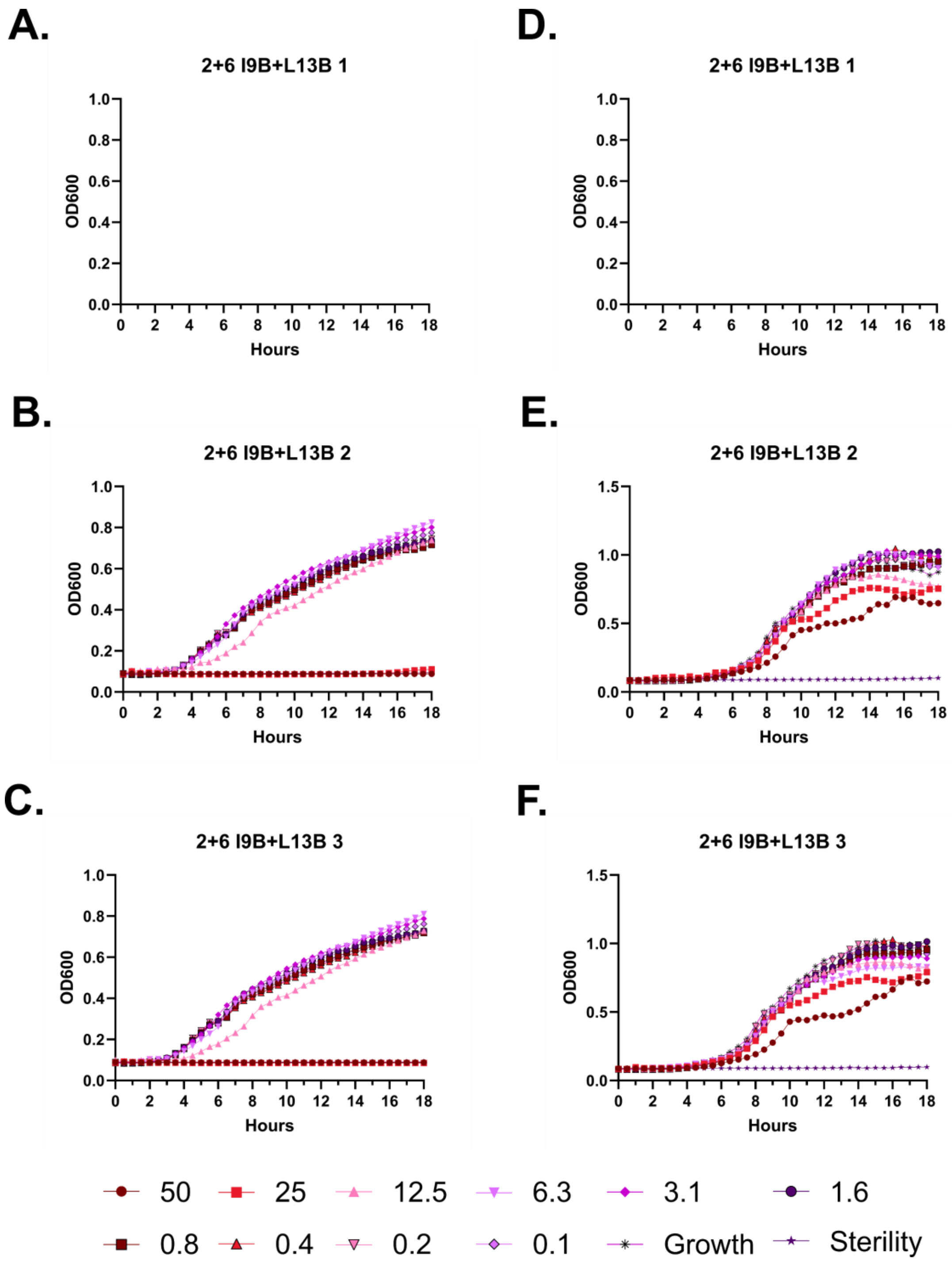
A.
2+6 I9B+L13B 1
OD600
Hours
D.
2+6 I9B+L13B 1
OD600
Hours
B.
2+6 I9B+L13B 2
OD600
Hours
E.
2+6 I9B+L13B 2
OD600
Hours
C.
2+6 I9B+L13B 3
OD600
Hours
F.
2+6 I9B+L13B 3
OD600
Hours
50
25
12.5
6.3
3.1
1.6
0.8
0.4
0.2
0.1
Growth
Sterility

**Figure S102 – Antimicrobial susceptibility testing with 2+6 I9B+L13B. A-C.** Growth of *E. coli* in the presence of the indicated peptide concentrations given in μM as monitored by OD600 at 37 °C over 18 hours. Each panel indicates an independent experiment. **D-F.** As in **A-C**, but with *P. aeruginosa*.

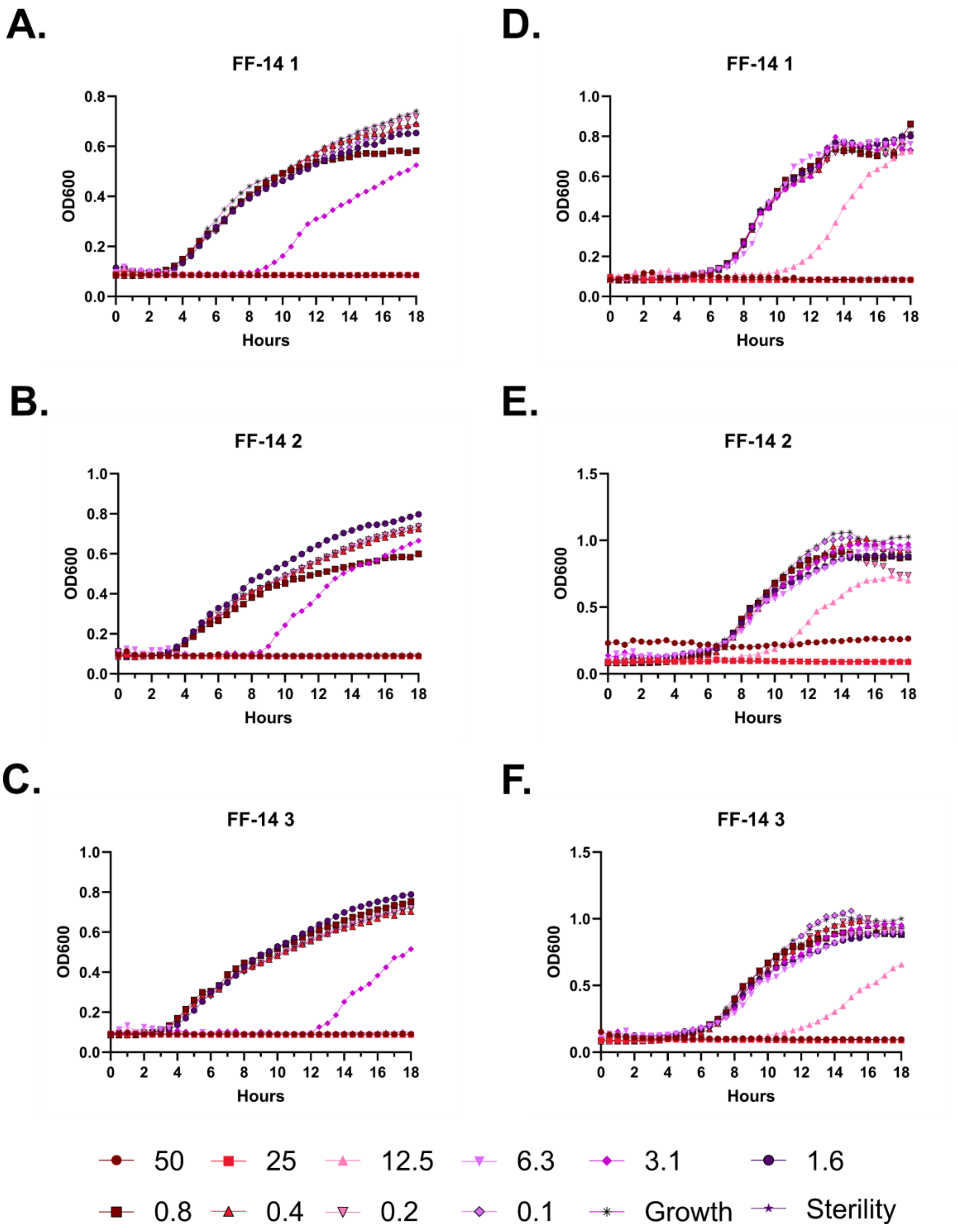
A.
FF-14 1
OD600
Hours
D.
FF-14 1
OD600
Hours
B.
FF-14 2
OD600
Hours
E.
FF-14 2
OD600
Hours
C.
FF-14 3
OD600
Hours
F.
FF-14 3
OD600
Hours
50
25
12.5
6.3
3.1
1.6
0.8
0.4
0.2
0.1
Growth
Sterility

**Figure S103 – Antimicrobial susceptibility testing with FF-14. A-C.** Growth of *E. coli* in the presence of the indicated peptide concentrations given in μM as monitored by OD600 at 37 °C over 18 hours. Each panel indicates an independent experiment. **D-F.** As in **A-C**, but with *P. aeruginosa.*

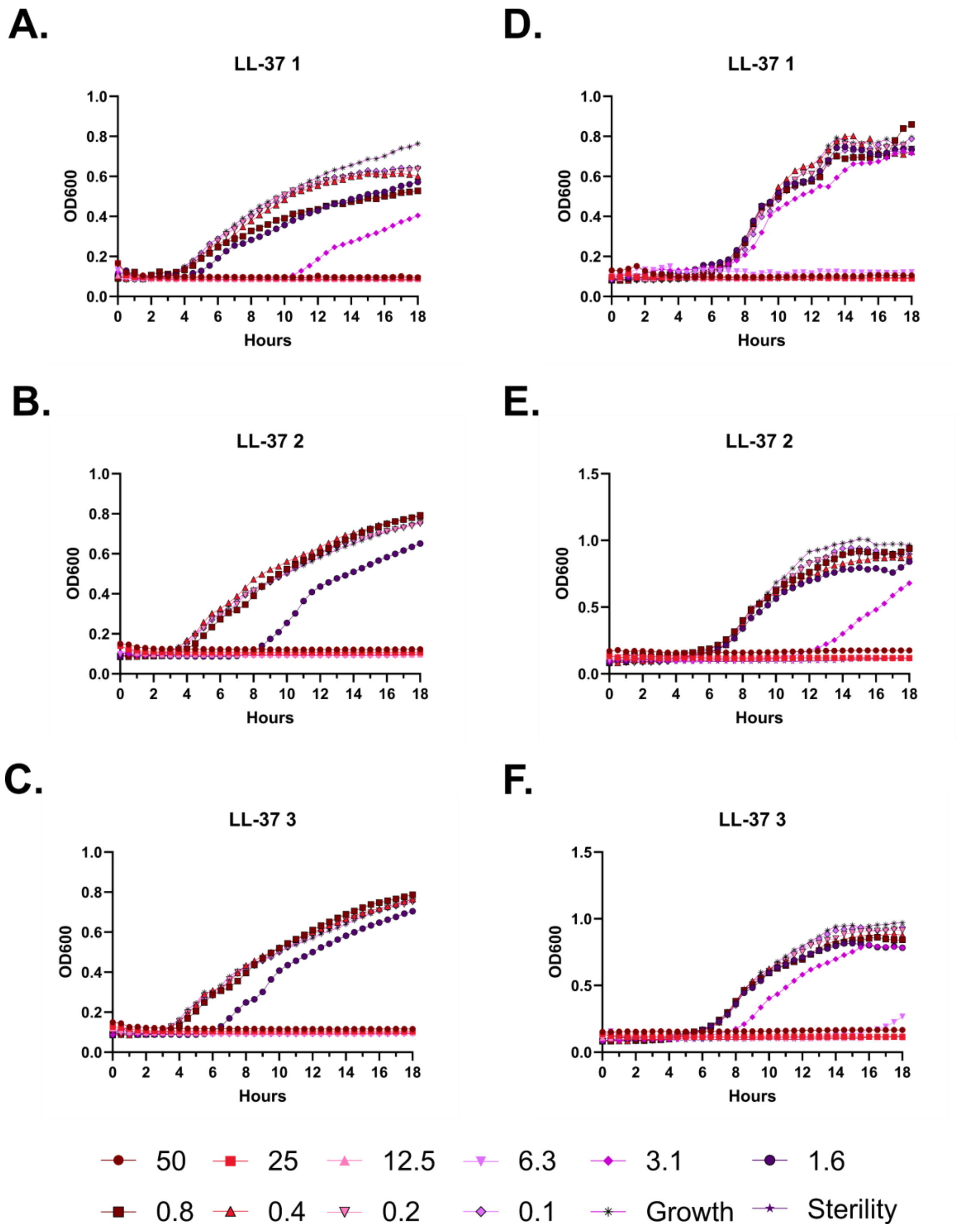

A.
LL-37 1
D.
LL-37 1
B.
LL-37 2
E.
LL-37 2
C.
LL-37 3
F.
LL-37 3
OD600
Hours
50
25
12.5
6.3
3.1
1.6
0.8
0.4
0.2
0.1
Growth
Sterility

**Figure S104 – Antimicrobial susceptibility testing with LL-37. A-C.** Growth of *E. coli* in the presence of the indicated peptide concentrations given in μM as monitored by OD600 at 37 °C over 18 hours. Each panel indicates an independent experiment. **D-F.** As in **A-C**, but with *P. aeruginosa.*

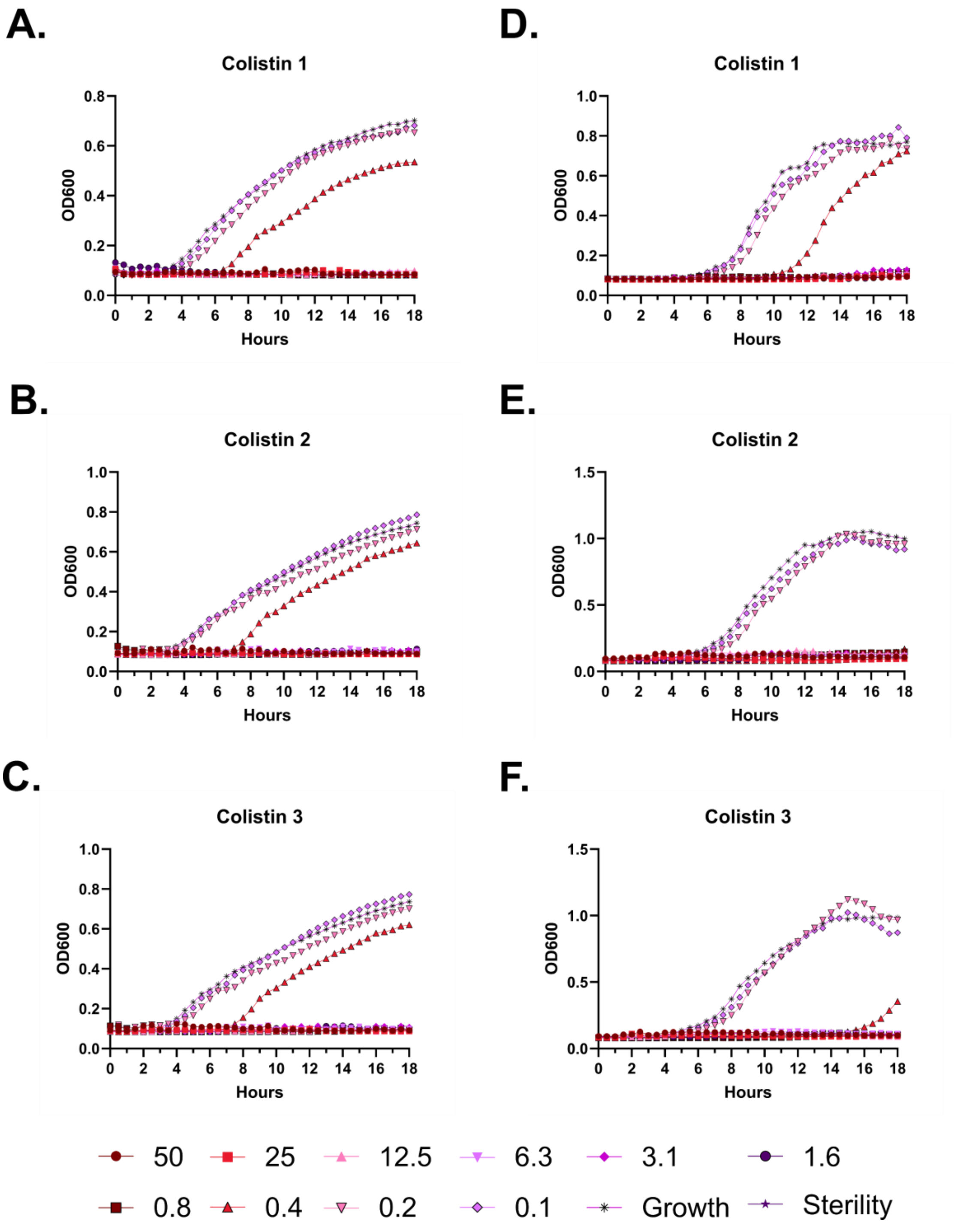
A.
Colistin 1
OD600
Hours
D.
Colistin 1
OD600
Hours
B.
Colistin 2
OD600
Hours
E.
Colistin 2
OD600
Hours
C.
Colistin 3
OD600
Hours
F.
Colistin 3
OD600
Hours
50
25
12.5
6.3
3.1
1.6
0.8
0.4
0.2
0.1
Growth
Sterility

**Figure S105 – Antimicrobial susceptibility testing with colistin. A-C.** Growth of *E. coli* in the presence of the indicated peptide concentrations given in μM as monitored by OD600 at 37 °C over 18 hours. Each panel indicates an independent experiment. **D-F.** As in **A-C**, but with *P. aeruginosa.*

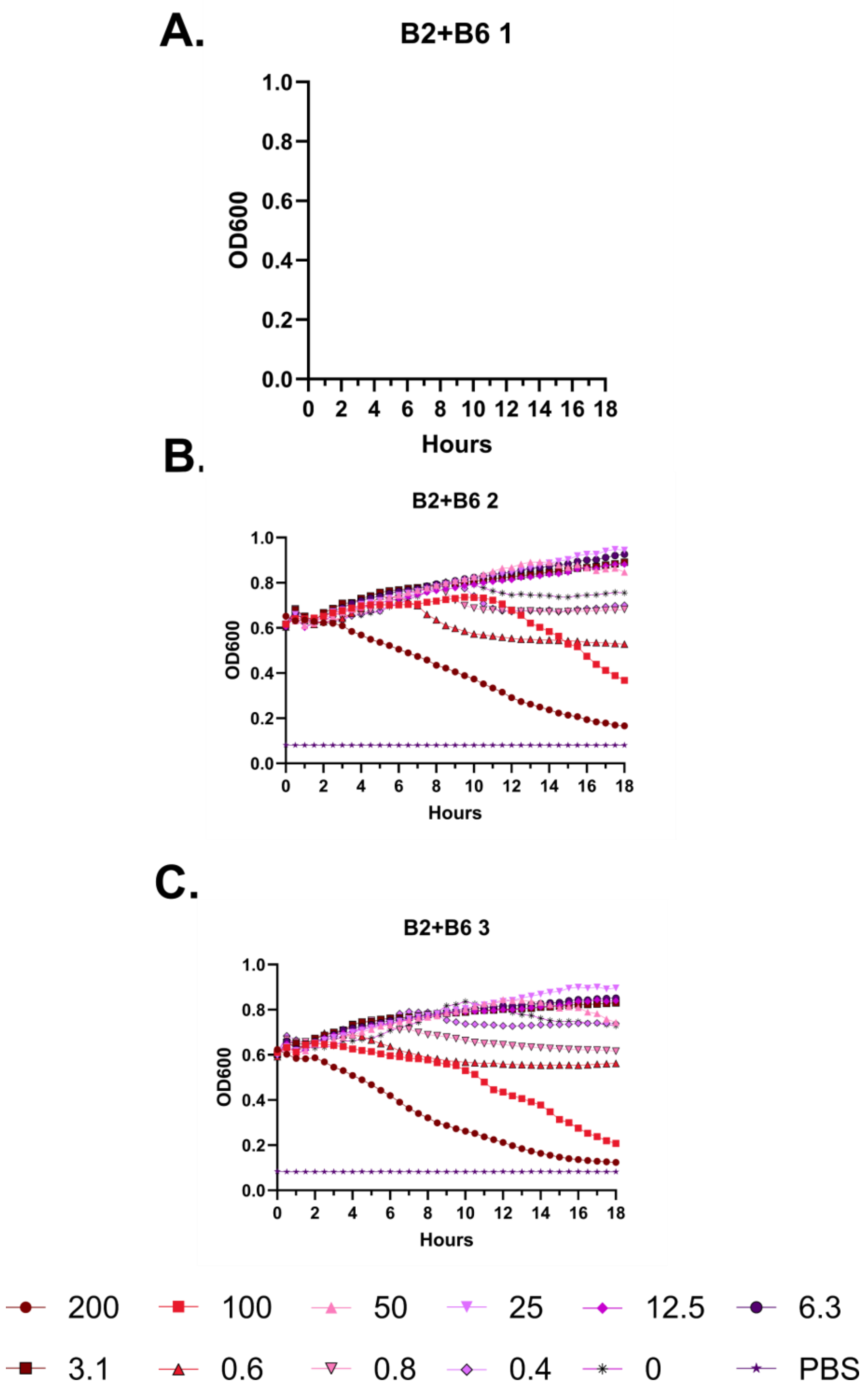
A.
B2+B6 1
1.0
0.8
0.6
0.4
0.2
0.0
OD600
0 2 4 6 8 10 12 14 16 18
Hours
B.
B2+B6 2
OD600
Hours
C.
B2+B6 3
OD600
Hours
200
100
50
25
12.5
6.3
3.1
0.6
0.8
0.4
0
PBS

**Figure S106 – Hemolysis over time with FF-14 B2+B6. A-C.** Each of three independent experiments showing trends in OD600 over time on incubation of defibrinated sheep blood with the indicated concentrations of peptide in μM at 37 °C. Decreasing OD600 indicates increasing hemolysis. Blank panels were part of a triplicate series that was not completed.

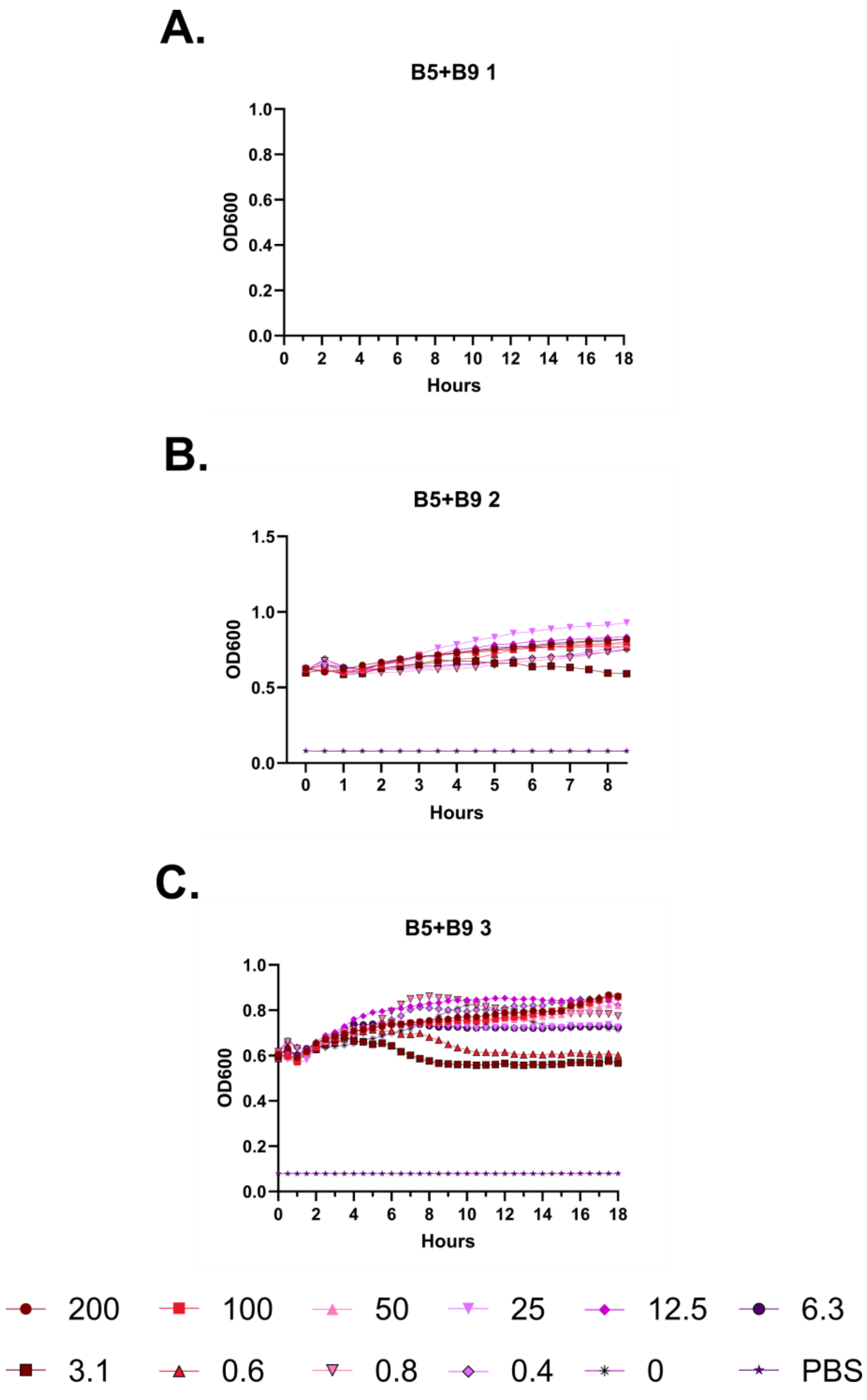
A.
B5+B9 1
OD600
Hours
B.
B5+B9 2
OD600
Hours
C.
B5+B9 3
OD600
Hours
200
100
50
25
12.5
6.3
3.1
0.6
0.8
0.4
0
PBS

**Figure S107 – Hemolysis over time with FF-14 B5+B9. A-C.** Each of three independent experiments showing trends in OD600 over time on incubation of defibrinated sheep blood with the indicated concentrations of peptide in μM at 37 °C. Decreasing OD600 indicates increasing hemolysis. Blank panels were part of a triplicate series that was not completed.

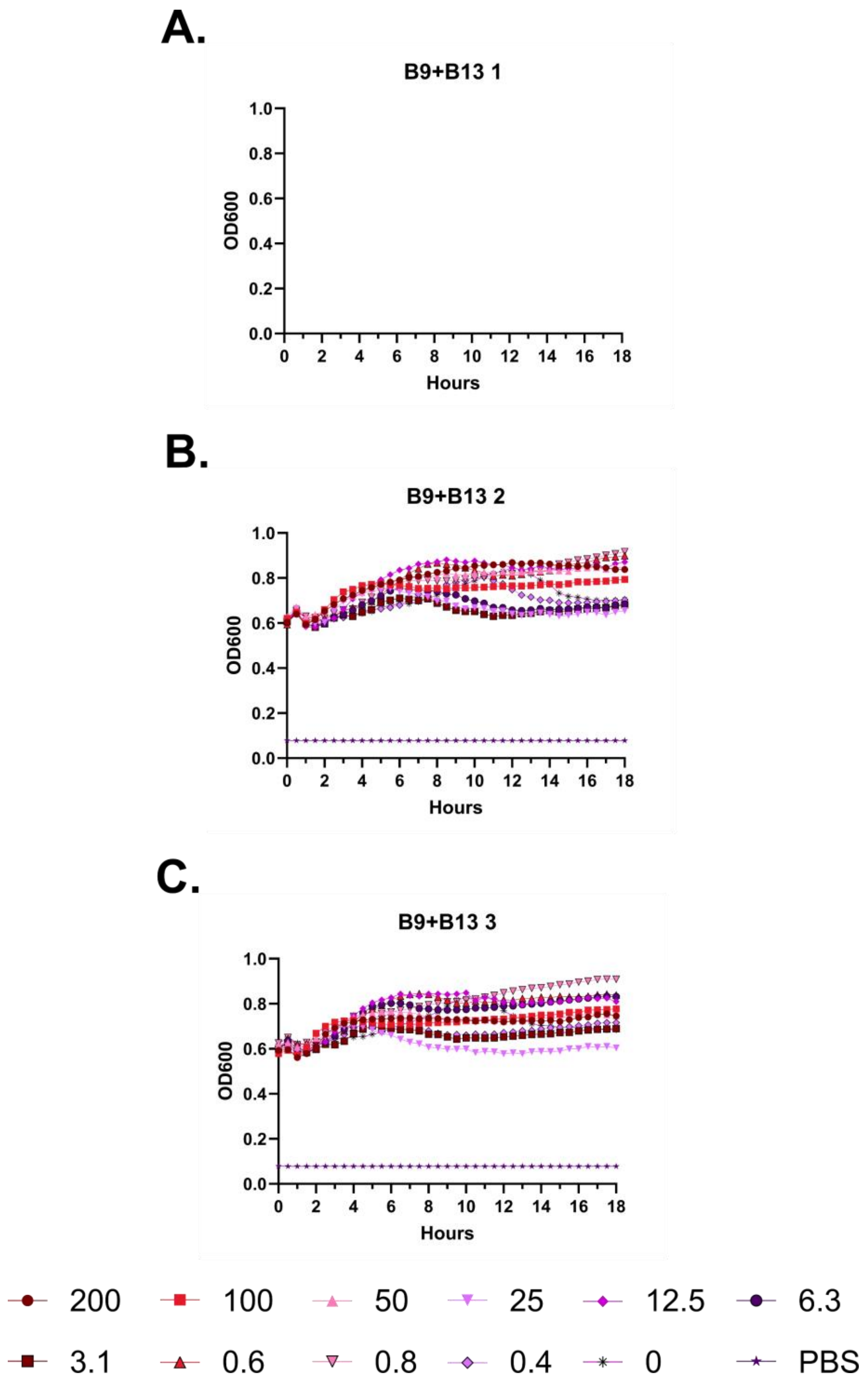
A.
B9+B13 1
OD600
Hours
B.
B9+B13 2
OD600
Hours
C.
B9+B13 3
OD600
Hours
200
100
50
25
12.5
6.3
3.1
0.6
0.8
0.4
0
PBS

**Figure S108 – Hemolysis over time with FF-14 B9+B13. A-C.** Each of three independent experiments showing trends in OD600 over time on incubation of defibrinated sheep blood with the indicated concentrations of peptide in μM at 37 °C. Decreasing OD600 indicates increasing hemolysis. Blank panels were part of a triplicate series that was not completed.

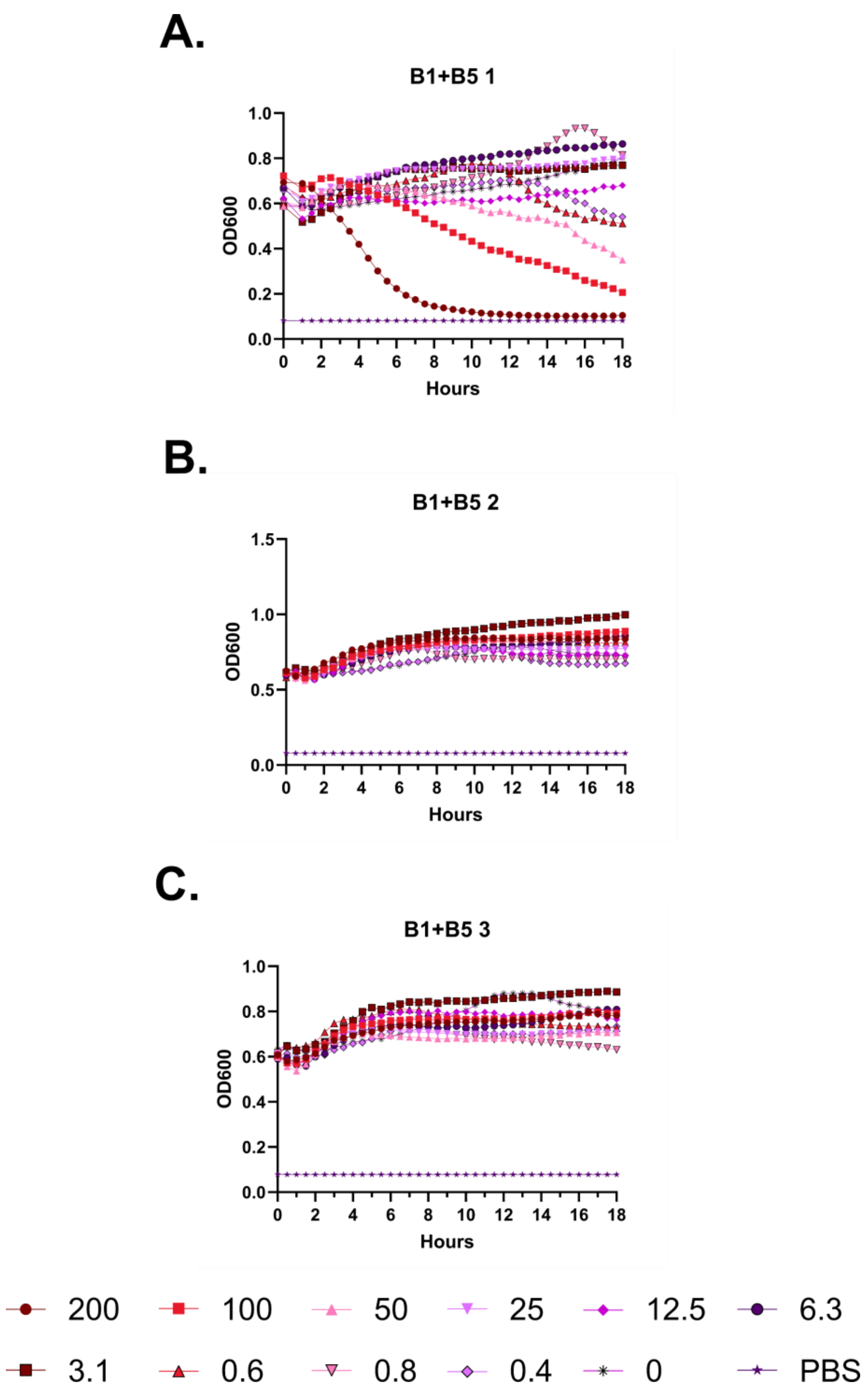
A.
B1+B5 1
OD600
Hours
0.0
0.2
0.4
0.6
0.8
1.0
0 2 4 6 8 10 12 14 16 18
B.
B1+B5 2
OD600
Hours
0.0
0.5
1.0
1.5
0 2 4 6 8 10 12 14 16 18
C.
B1+B5 3
OD600
Hours
0.0
0.2
0.4
0.6
0.8
1.0
0 2 4 6 8 10 12 14 16 18
200
100
50
25
12.5
6.3
3.1
0.6
0.8
0.4
0
PBS

**Figure S109 – Hemolysis over time with FF-14 B1+B5. A-C.** Each of three independent experiments showing trends in OD600 over time on incubation of defibrinated sheep blood with the indicated concentrations of peptide in μM at 37 °C. Decreasing OD600 indicates increasing hemolysis. Blank panels were part of a triplicate series that was not completed.

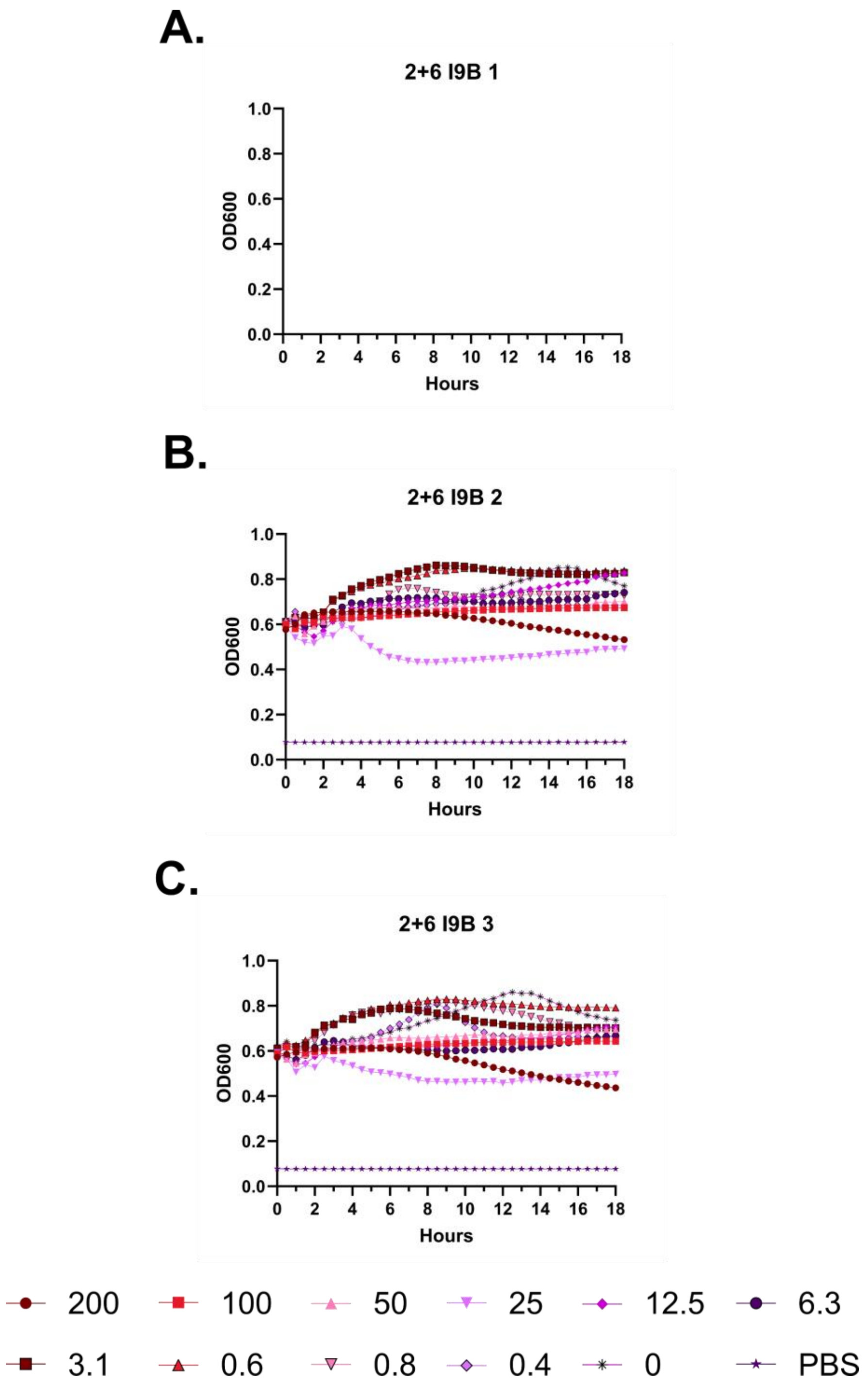
A.
2+6 I9B 1
OD600
Hours
B.
2+6 I9B 2
OD600
Hours
C.
2+6 I9B 3
OD600
Hours
200
100
50
25
12.5
6.3
3.1
0.6
0.8
0.4
0
PBS

**Figure S110 – Hemolysis over time with 2+6 I9B. A-C.** Each of three independent experiments showing trends in OD600 over time on incubation of defibrinated sheep blood with the indicated concentrations of peptide in μM at 37 °C. Decreasing OD600 indicates increasing hemolysis. Blank panels were part of a triplicate series that was not completed.

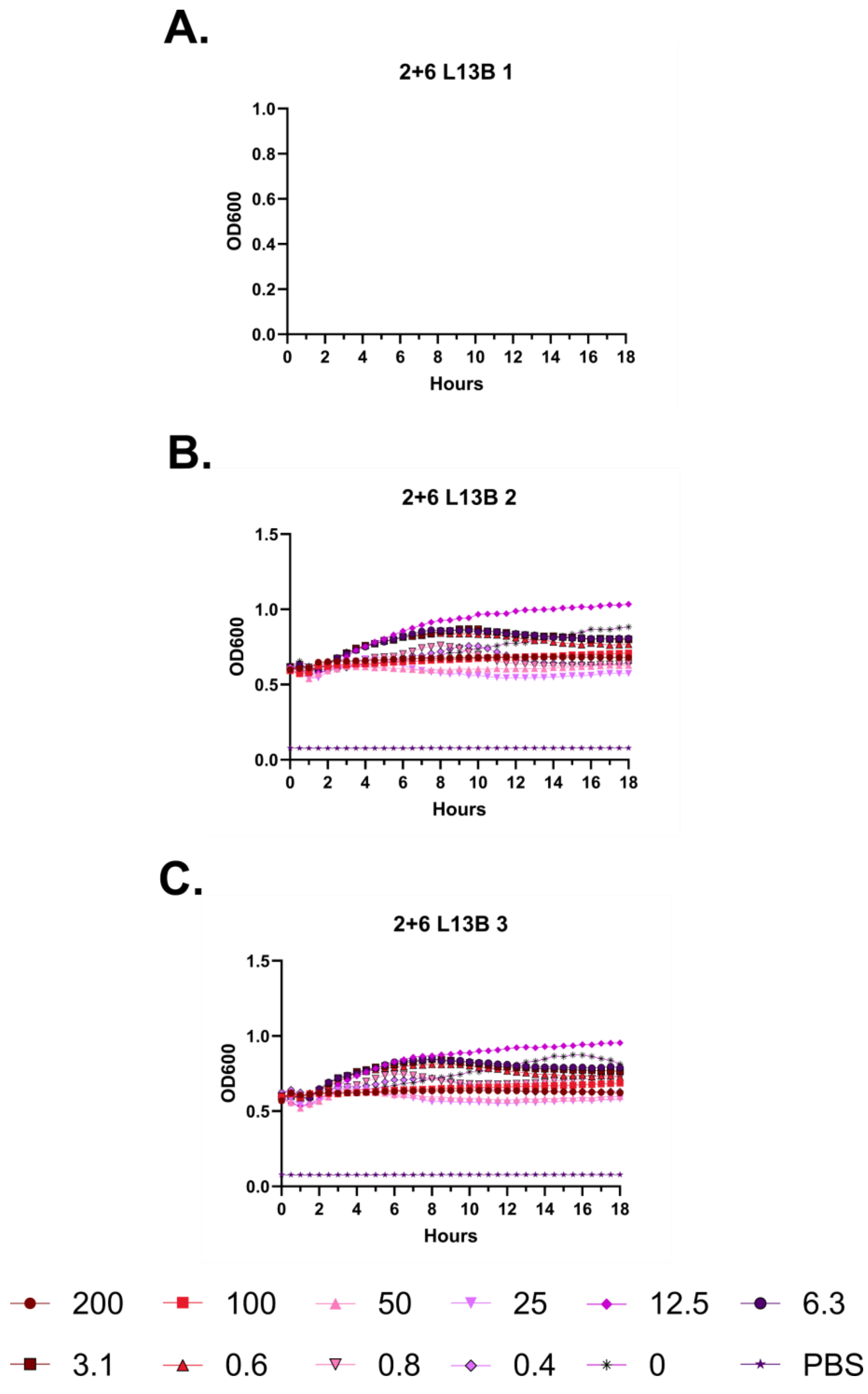
A.
2+6 L13B 1
OD600
Hours
B.
2+6 L13B 2
OD600
Hours
C.
2+6 L13B 3
OD600
Hours
200
100
50
25
12.5
6.3
3.1
0.6
0.8
0.4
0
PBS

**Figure S111 – Hemolysis over time with 2+6 L13B. A-C.** Each of three independent experiments showing trends in OD600 over time on incubation of defibrinated sheep blood with the indicated concentrations of peptide in μM at 37 °C. Decreasing OD600 indicates increasing hemolysis. Blank panels were part of a triplicate series that was not completed.

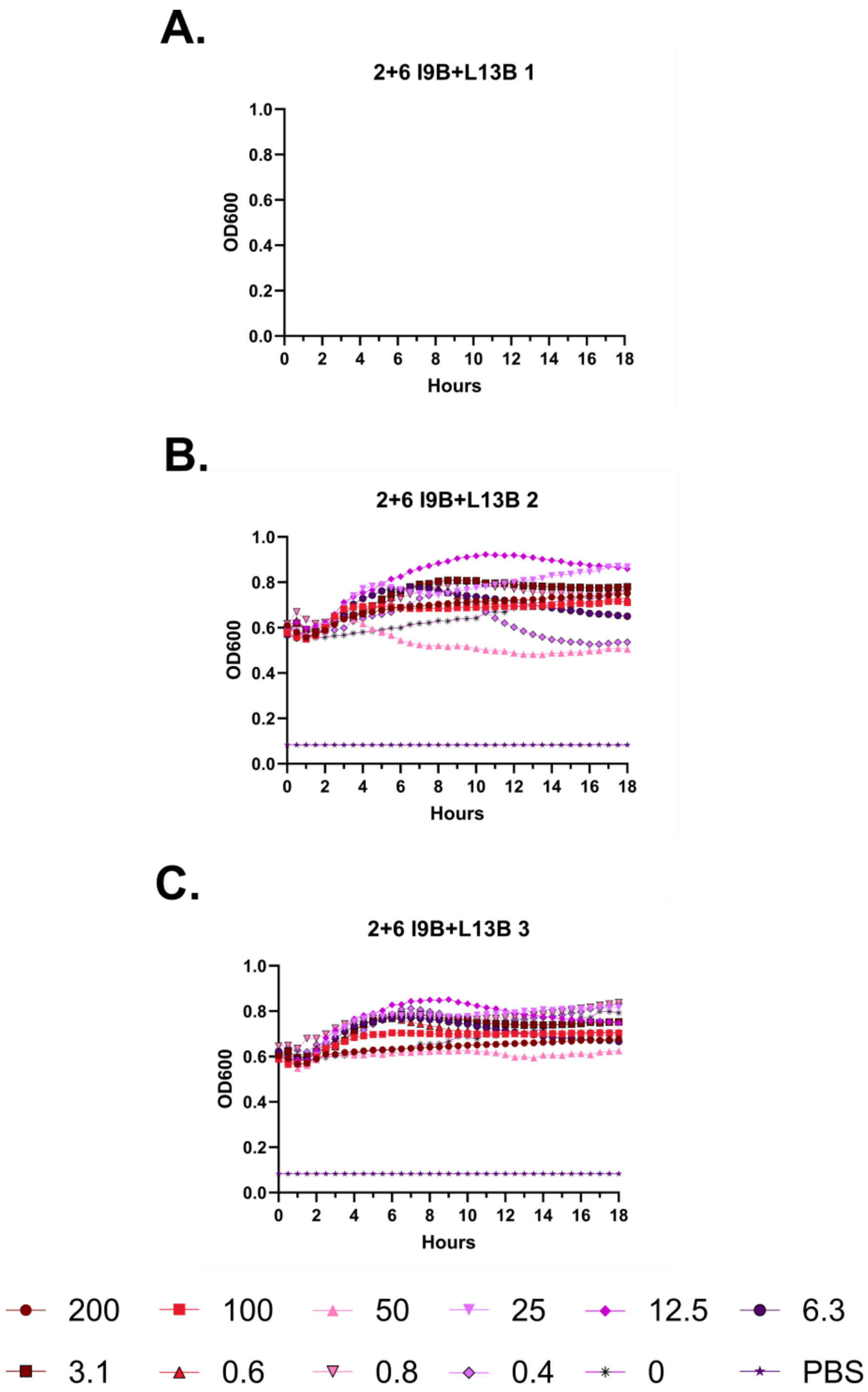
A.
2+6 I9B+L13B 1
B.
2+6 I9B+L13B 2
C.
2+6 I9B+L13B 3
OD600
Hours
1.0
0.8
0.6
0.4
0.2
0.0
0
2
4
6
8
10
12
14
16
18
200
100
50
25
12.5
6.3
3.1
0.6
0.8
0.4
0
PBS

**Figure S112 – Hemolysis over time with 2+6 I9B+L13B. A-C.** Each of three independent experiments showing trends in OD600 over time on incubation of defibrinated sheep blood with the indicated concentrations of peptide in μM at 37 °C. Decreasing OD600 indicates increasing hemolysis. Blank panels were part of a triplicate series that was not completed.

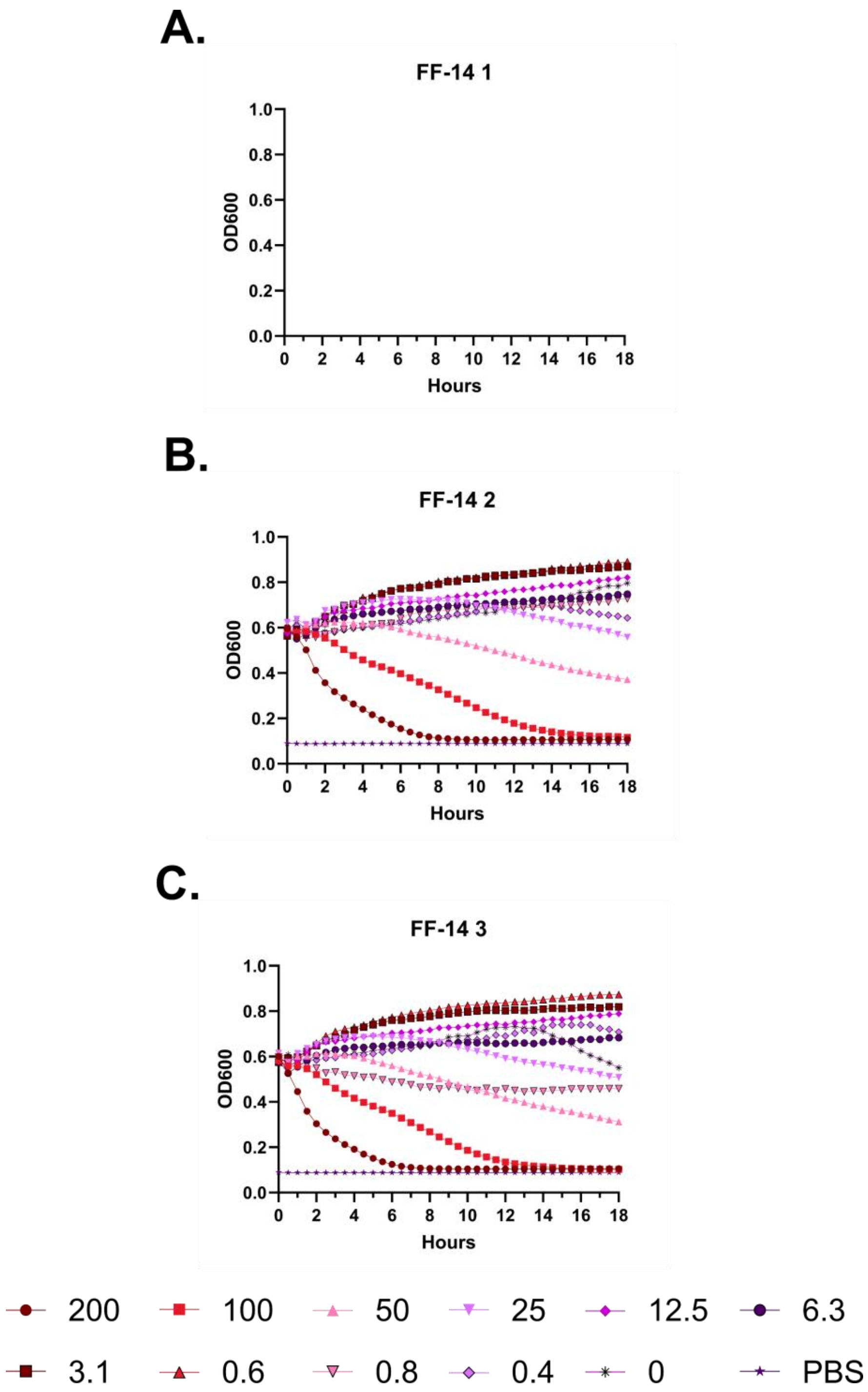

A.
FF-14 1
OD600
Hours
B.
FF-14 2
OD600
Hours
C.
FF-14 3
OD600
Hours
200
100
50
25
12.5
6.3
3.1
0.6
0.8
0.4
0
PBS

**Figure S113 – Hemolysis over time with FF-14. A-C.** Each of three independent experiments showing trends in OD600 over time on incubation of defibrinated sheep blood with the indicated concentrations of peptide in μM at 37 °C. Decreasing OD600 indicates increasing hemolysis. Blank panels were part of a triplicate series that was not completed.

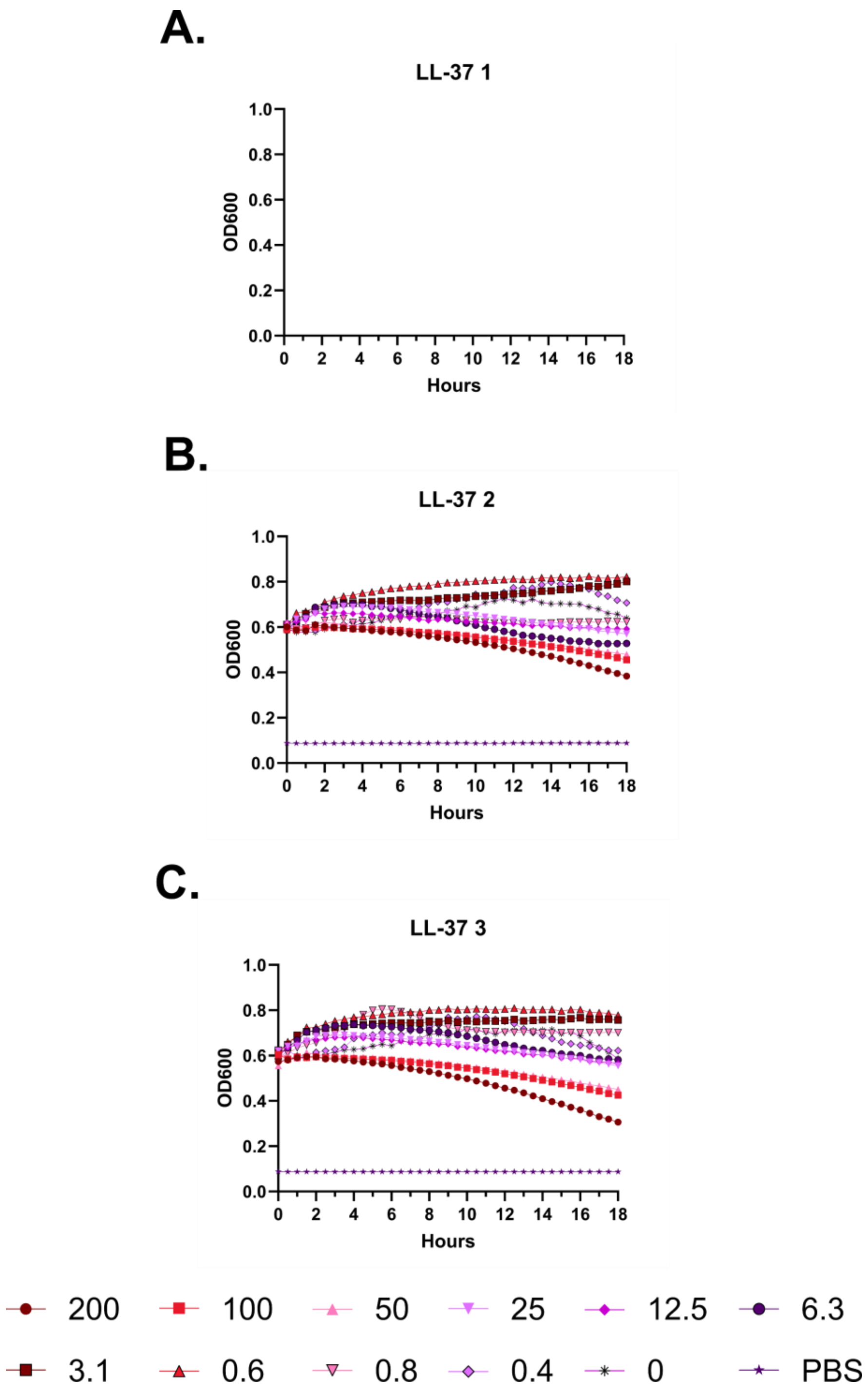

A.
LL-37 1
OD600
Hours
B.
LL-37 2
OD600
Hours
C.
LL-37 3
OD600
Hours
200
100
50
25
12.5
6.3
3.1
0.6
0.8
0.4
0
PBS

**Figure S114 – Hemolysis over time with LL-37. A-C.** Each of three independent experiments showing trends in OD600 over time on incubation of defibrinated sheep blood with the indicated concentrations of peptide in µM at 37 °C. Decreasing OD600 indicates increasing hemolysis. Blank panels were part of a triplicate series that was not completed.

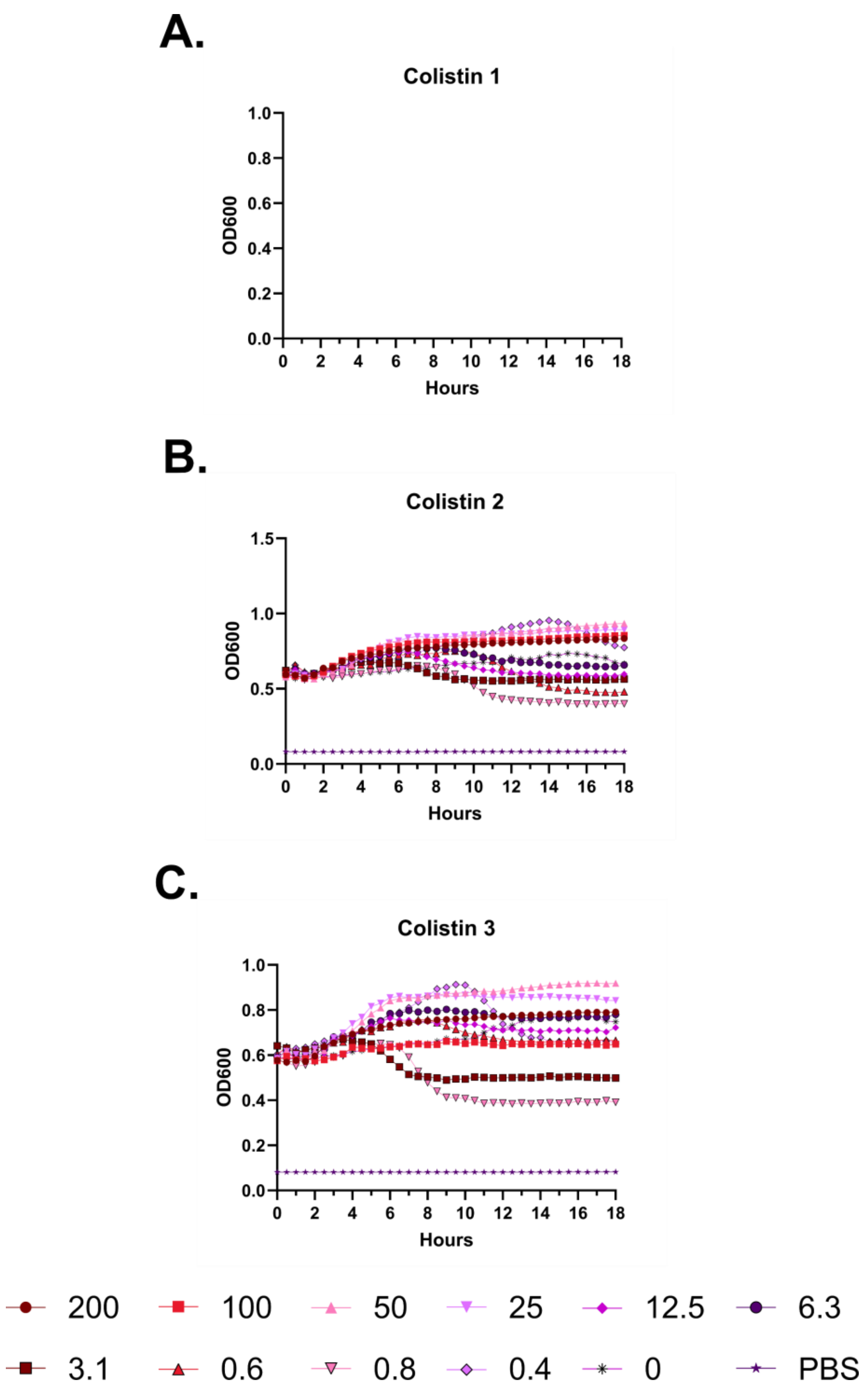
A.
Colistin 1
OD600
Hours
B.
Colistin 2
OD600
Hours
C.
Colistin 3
OD600
Hours
200
100
50
25
12.5
6.3
3.1
0.6
0.8
0.4
0
PBS

**Figure S115 – Hemolysis over time with colistin. A-C.** Each of three independent experiments showing trends in OD600 over time on incubation of defibrinated sheep blood with the indicated concentrations of peptide in μM at 37 °C. Decreasing OD600 indicates increasing hemolysis. Blank panels were part of a triplicate series that was not completed.

**A.**

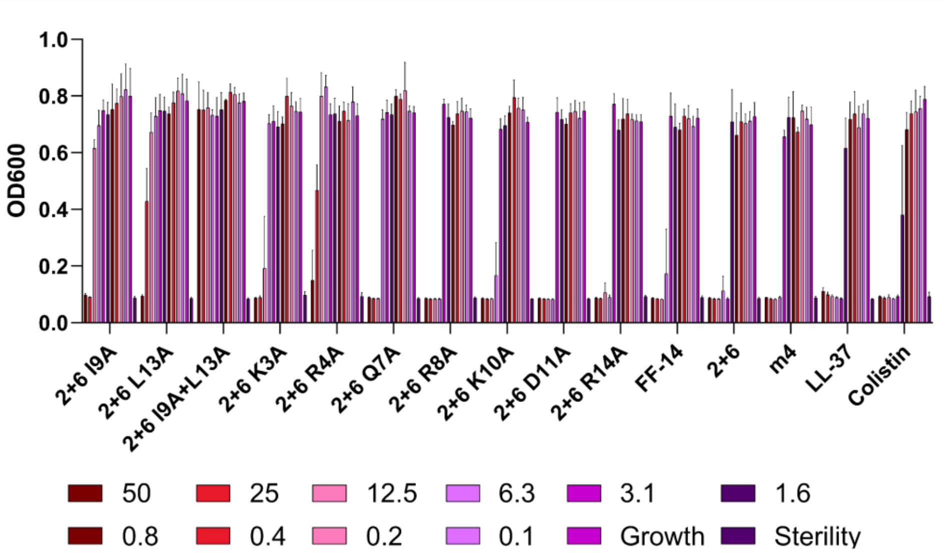
OD600
1.0
0.8
0.6
0.4
0.2
0.0
2+6 I9A
2+6 L13A
2+6 I9A+L13A
2+6 K3A
2+6 R4A
2+6 Q7A
2+6 R8A
2+6 K10A
2+6 D11A
2+6 R14A
FF-14
2+6
m4
LL-37
Colistin
50
25
12.5
6.3
3.1
1.6
0.8
0.4
0.2
0.1
Growth
Sterility

**B.**

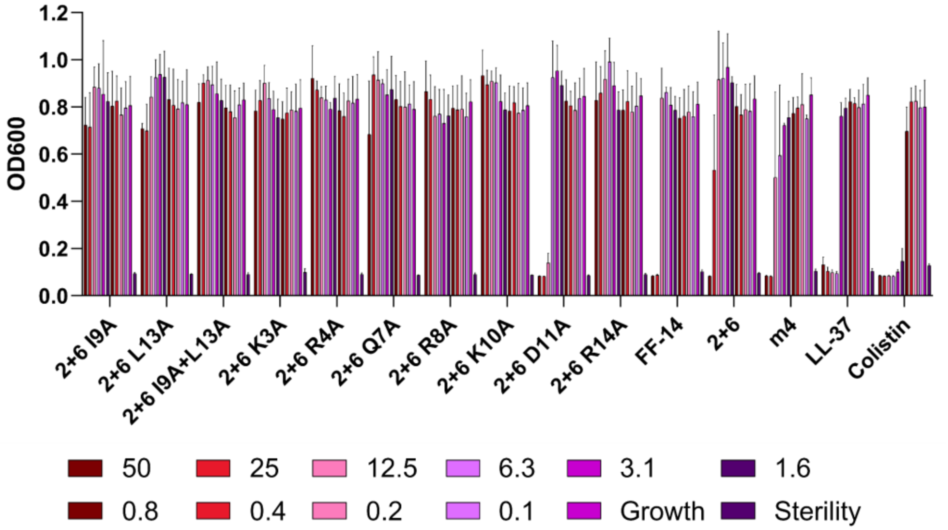
OD600
1.2
1.0
0.8
0.6
0.4
0.2
0.0
2+6 I9A
2+6 L13A
2+6 I9A+L13A
2+6 K3A
2+6 R4A
2+6 Q7A
2+6 R8A
2+6 K10A
2+6 D11A
2+6 R14A
FF-14
2+6
m4
LL-37
Colistin
50
25
12.5
6.3
3.1
1.6
0.8
0.4
0.2
0.1
Growth
Sterility

**C.**

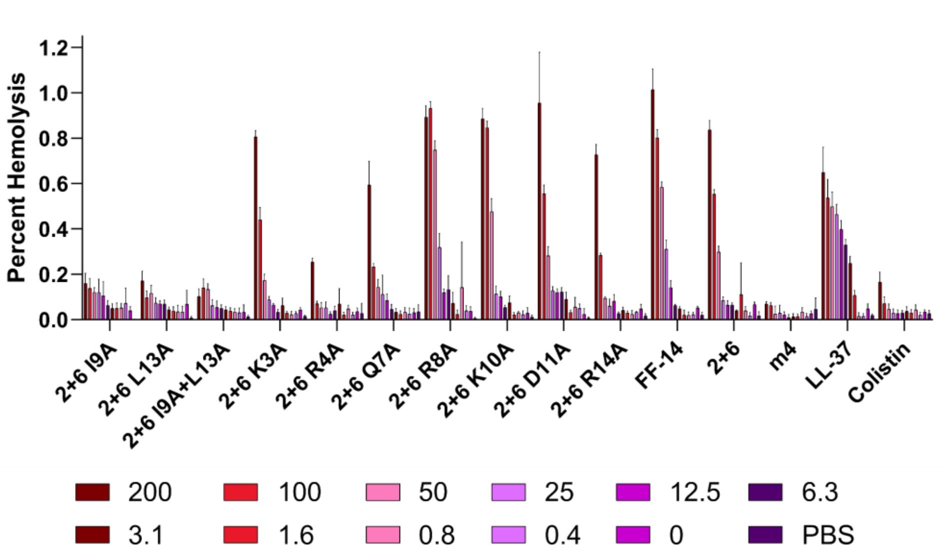
Percent Hemolysis
1.2
1.0
0.8
0.6
0.4
0.2
0.0
2+6 I9A
2+6 L13A
2+6 I9A+L13A
2+6 K3A
2+6 R4A
2+6 Q7A
2+6 R8A
2+6 K10A
2+6 D11A
2+6 R14A
FF-14
2+6
m4
LL-37
Colistin
200
100
50
25
12.5
6.3
3.1
1.6
0.8
0.4
0
PBS

**Figure S116 – Histogram depiction of data from Figure 5. A.** Endpoint activity of each peptide against *E. coli*; measures are raw absorbance at 600 nm at 18 hours. **B.** Endpoint activity of each peptide against *P. aeruginosa*. **C.** Hemolytic activity of each peptide; measures are absorbance at 414 nm scaled to a freeze-thaw positive control set at 100% hemolysis. Concentrations throughout are shown are in μM.

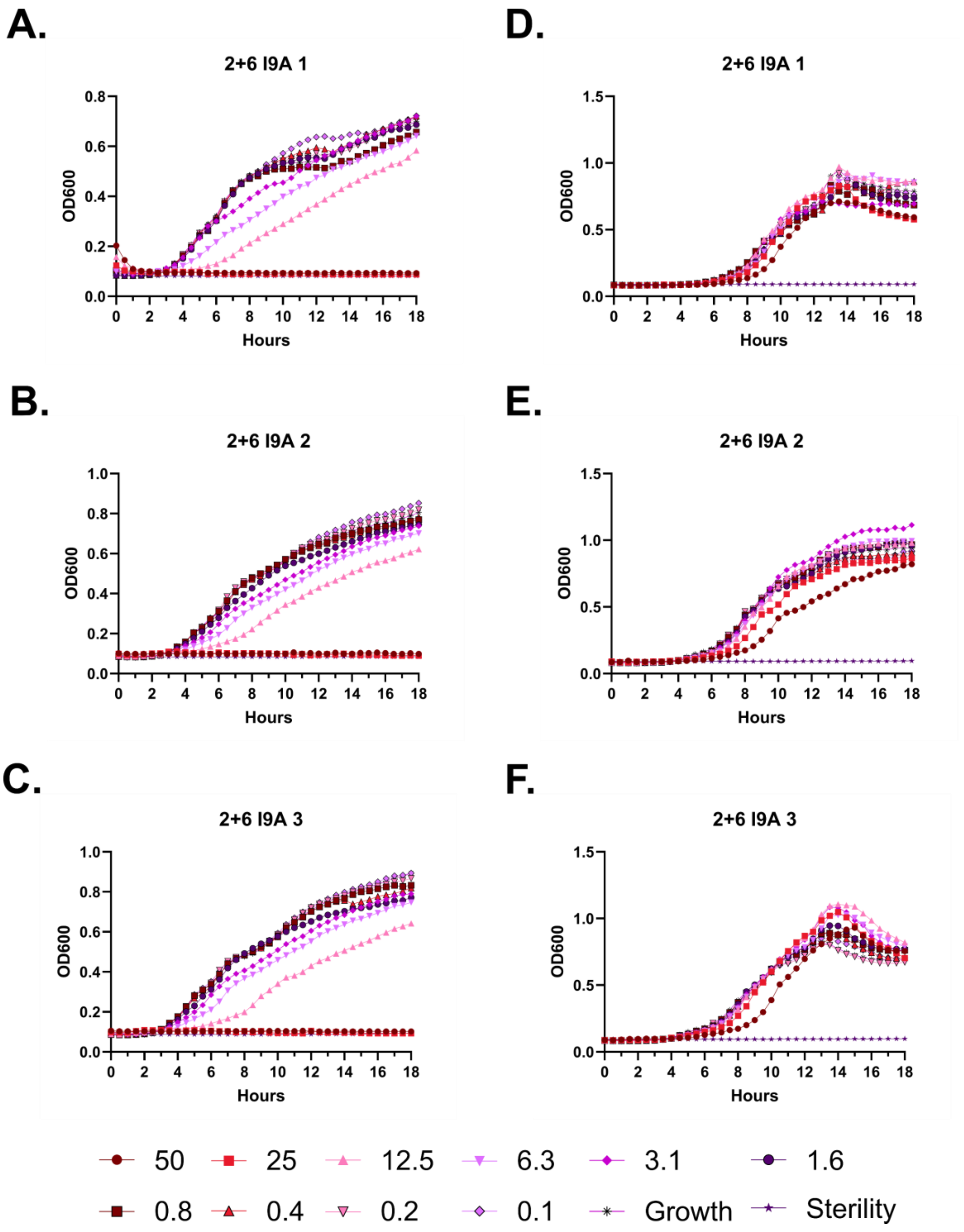

A.
2+6 I9A 1
D.
2+6 I9A 1
B.
2+6 I9A 2
E.
2+6 I9A 2
C.
2+6 I9A 3
F.
2+6 I9A 3
OD600
Hours
50
25
12.5
6.3
3.1
1.6
0.8
0.4
0.2
0.1
Growth
Sterility

**Figure S117 – Antimicrobial susceptibility testing with 2+6 I9A. A-C.** Growth of *E. coli* in the presence of the indicated peptide concentrations given in μM as monitored by OD600 at 37 °C over 18 hours. Each panel indicates an independent experiment. **D-F.** As in **A-C**, but with *P. aeruginosa.*

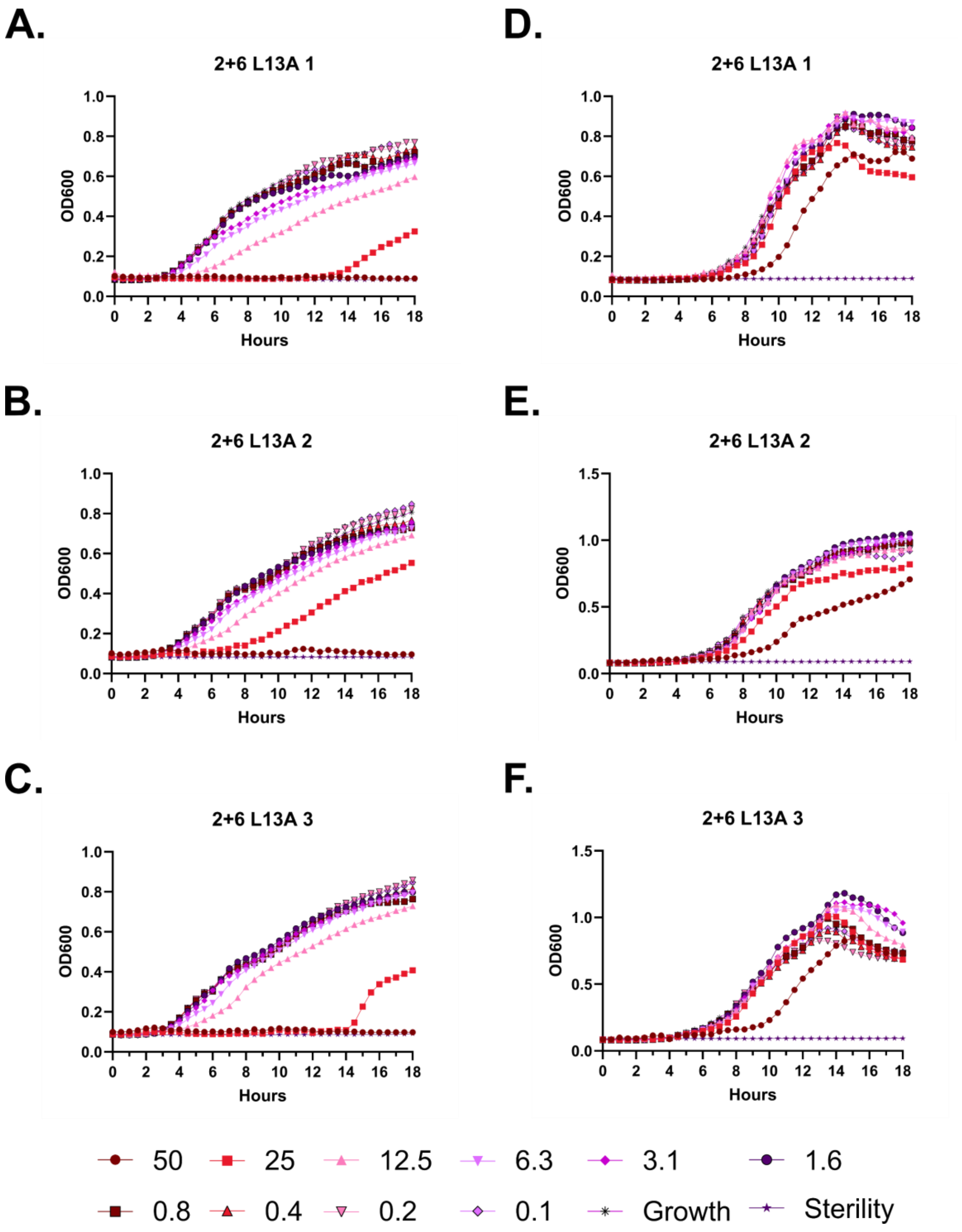
A.
2+6 L13A 1
OD600
Hours
B.
2+6 L13A 2
OD600
Hours
C.
2+6 L13A 3
OD600
Hours
D.
2+6 L13A 1
OD600
Hours
E.
2+6 L13A 2
OD600
Hours
F.
2+6 L13A 3
OD600
Hours
50
25
12.5
6.3
3.1
1.6
0.8
0.4
0.2
0.1
Growth
Sterility

**Figure S118 – Antimicrobial susceptibility testing with 2+6 L13A. A-C.** Growth of *E. coli* in the presence of the indicated peptide concentrations given in μM as monitored by OD600 at 37 °C over 18 hours. Each panel indicates an independent experiment. **D-F.** As in **A-C**, but with *P. aeruginosa.*

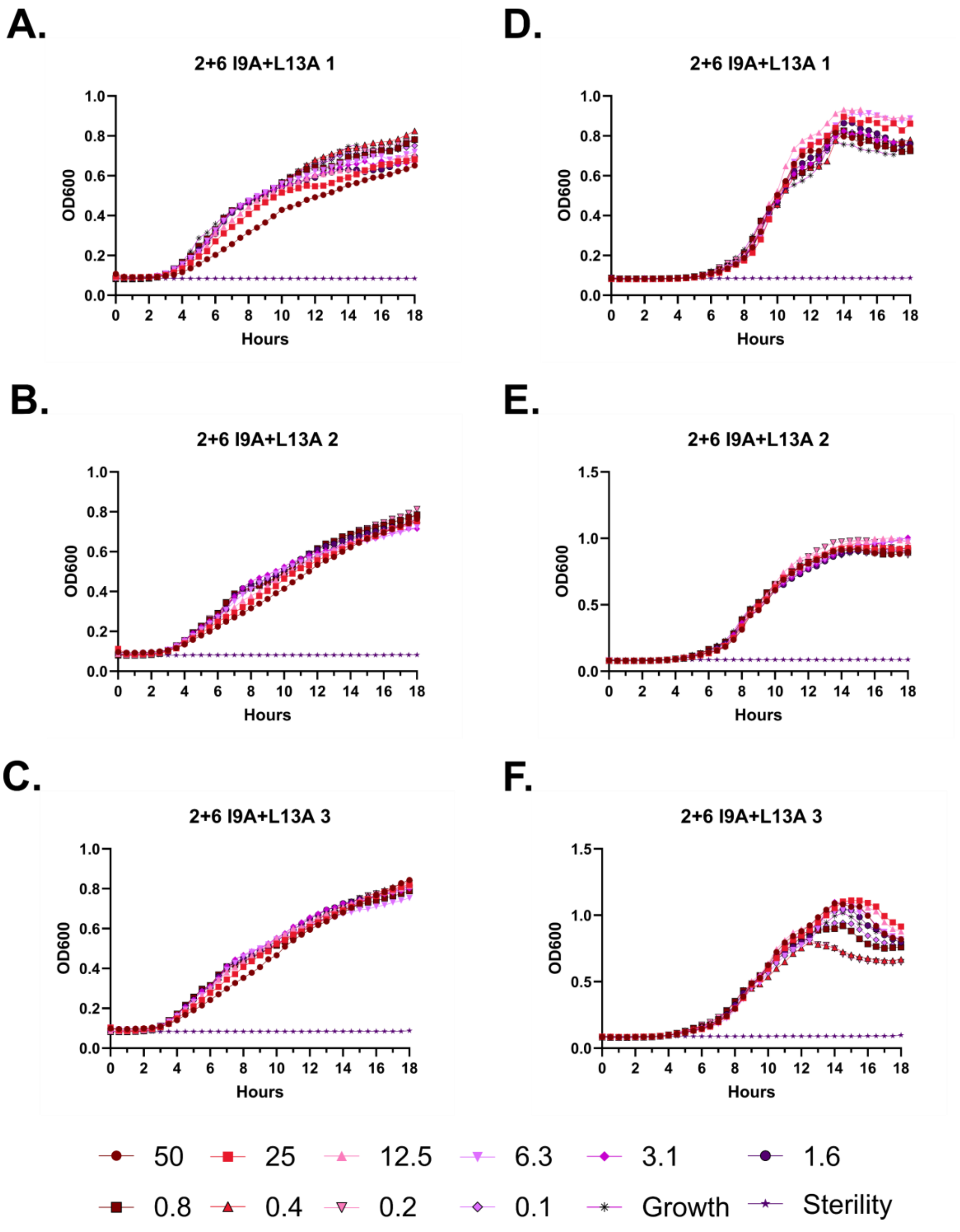
A.
2+6 I9A+L13A 1
B.
2+6 I9A+L13A 2
C.
2+6 I9A+L13A 3
D.
2+6 I9A+L13A 1
E.
2+6 I9A+L13A 2
F.
2+6 I9A+L13A 3
OD600
Hours
50
25
12.5
6.3
3.1
1.6
0.8
0.4
0.2
0.1
Growth
Sterility

**Figure S119 – Antimicrobial susceptibility testing with 2+6 I9A+L13A. A-C.** Growth of *E. coli* in the presence of the indicated peptide concentrations given in μM as monitored by OD600 at 37 °C over 18 hours. Each panel indicates an independent experiment. **D-F.** As in **A-C**, but with *P. aeruginosa.*

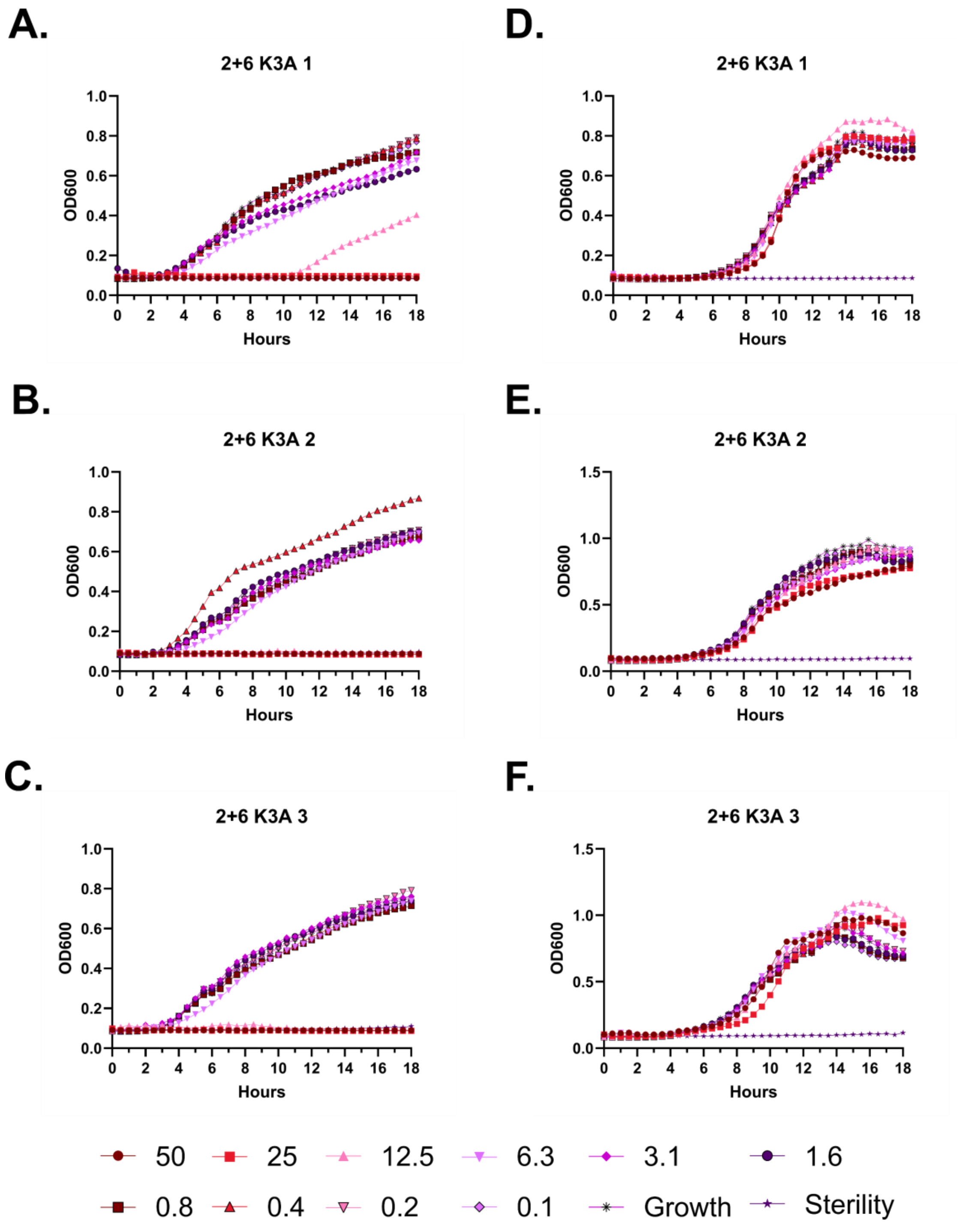
A.
2+6 K3A 1
OD600
Hours
D.
2+6 K3A 1
OD600
Hours
B.
2+6 K3A 2
OD600
Hours
E.
2+6 K3A 2
OD600
Hours
C.
2+6 K3A 3
OD600
Hours
F.
2+6 K3A 3
OD600
Hours
50
25
12.5
6.3
3.1
1.6
0.8
0.4
0.2
0.1
Growth
Sterility

**Figure S120 – Antimicrobial susceptibility testing with 2+6 K3A. A-C.** Growth of *E. coli* in the presence of the indicated peptide concentrations given in μM as monitored by OD600 at 37 °C over 18 hours. Each panel indicates an independent experiment. **D-F.** As in **A-C**, but with *P. aeruginosa.*

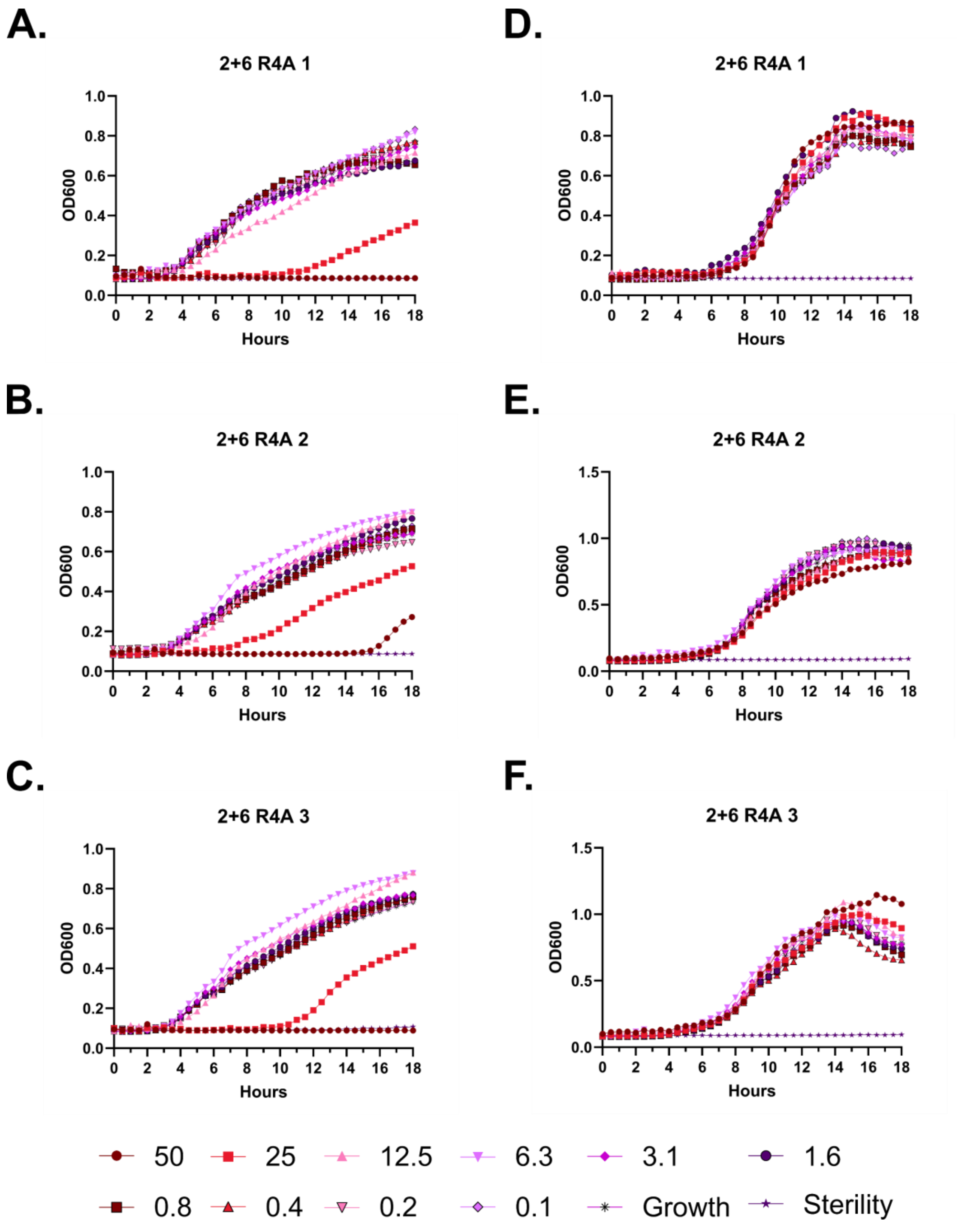

A.
2+6 R4A 1
D.
2+6 R4A 1
B.
2+6 R4A 2
E.
2+6 R4A 2
C.
2+6 R4A 3
F.
2+6 R4A 3
OD600
Hours
50
25
12.5
6.3
3.1
1.6
0.8
0.4
0.2
0.1
Growth
Sterility

**Figure S121 – Antimicrobial susceptibility testing with 2+6 R4A. A-C.** Growth of *E. coli* in the presence of the indicated peptide concentrations given in μM as monitored by OD600 at 37 °C over 18 hours. Each panel indicates an independent experiment. **D-F.** As in **A-C**, but with *P. aeruginosa.*

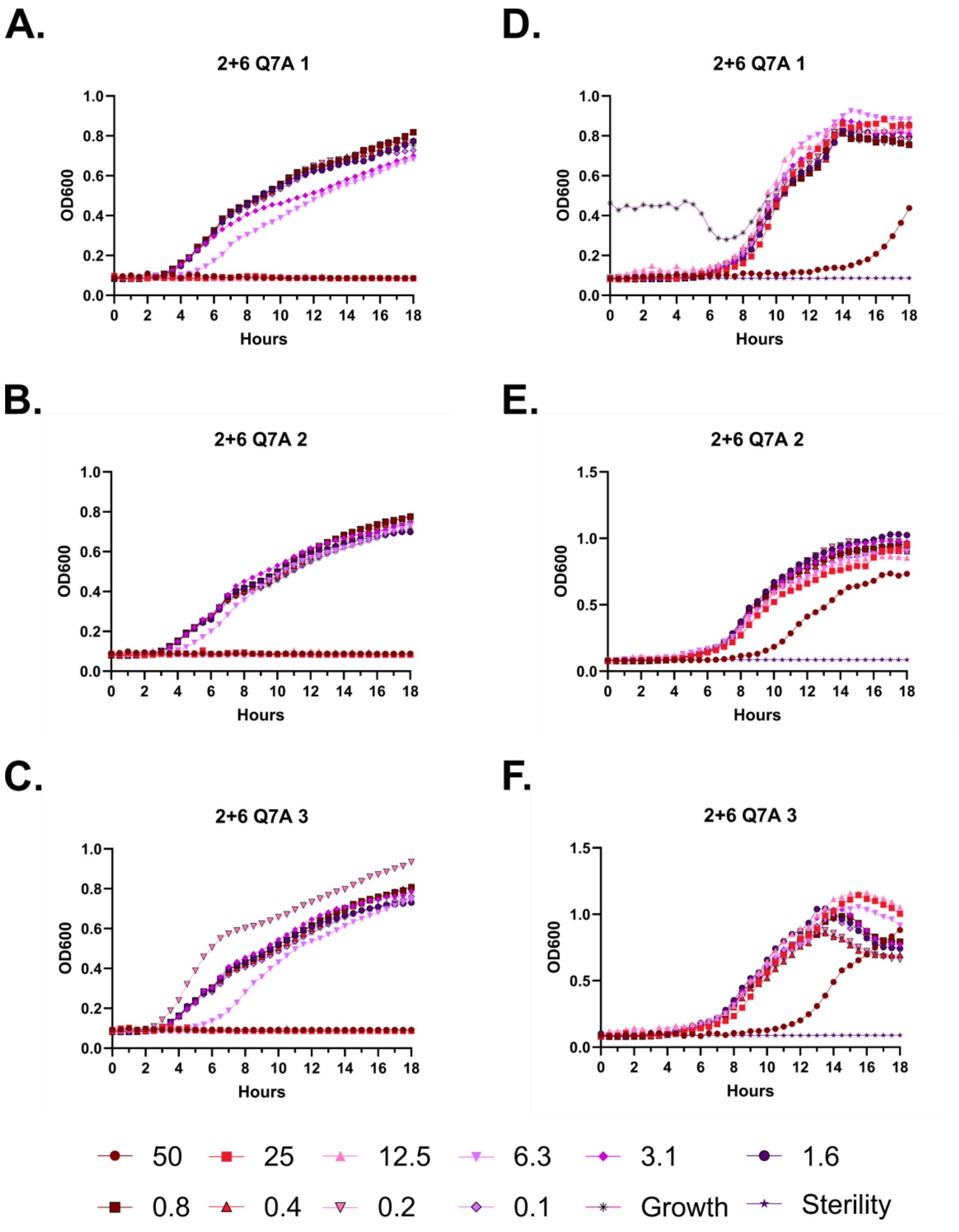

A.
2+6 Q7A 1
OD600
Hours
B.
2+6 Q7A 2
OD600
Hours
C.
2+6 Q7A 3
OD600
Hours
D.
2+6 Q7A 1
OD600
Hours
E.
2+6 Q7A 2
OD600
Hours
F.
2+6 Q7A 3
OD600
Hours
50
25
12.5
6.3
3.1
1.6
0.8
0.4
0.2
0.1
Growth
Sterility

**Figure S122 – Antimicrobial susceptibility testing with 2+6 Q7A. A-C.** Growth of *E. coli* in the presence of the indicated peptide concentrations given in μM as monitored by OD600 at 37 °C over 18 hours. Each panel indicates an independent experiment. **D-F.** As in **A-C**, but with *P. aeruginosa.*

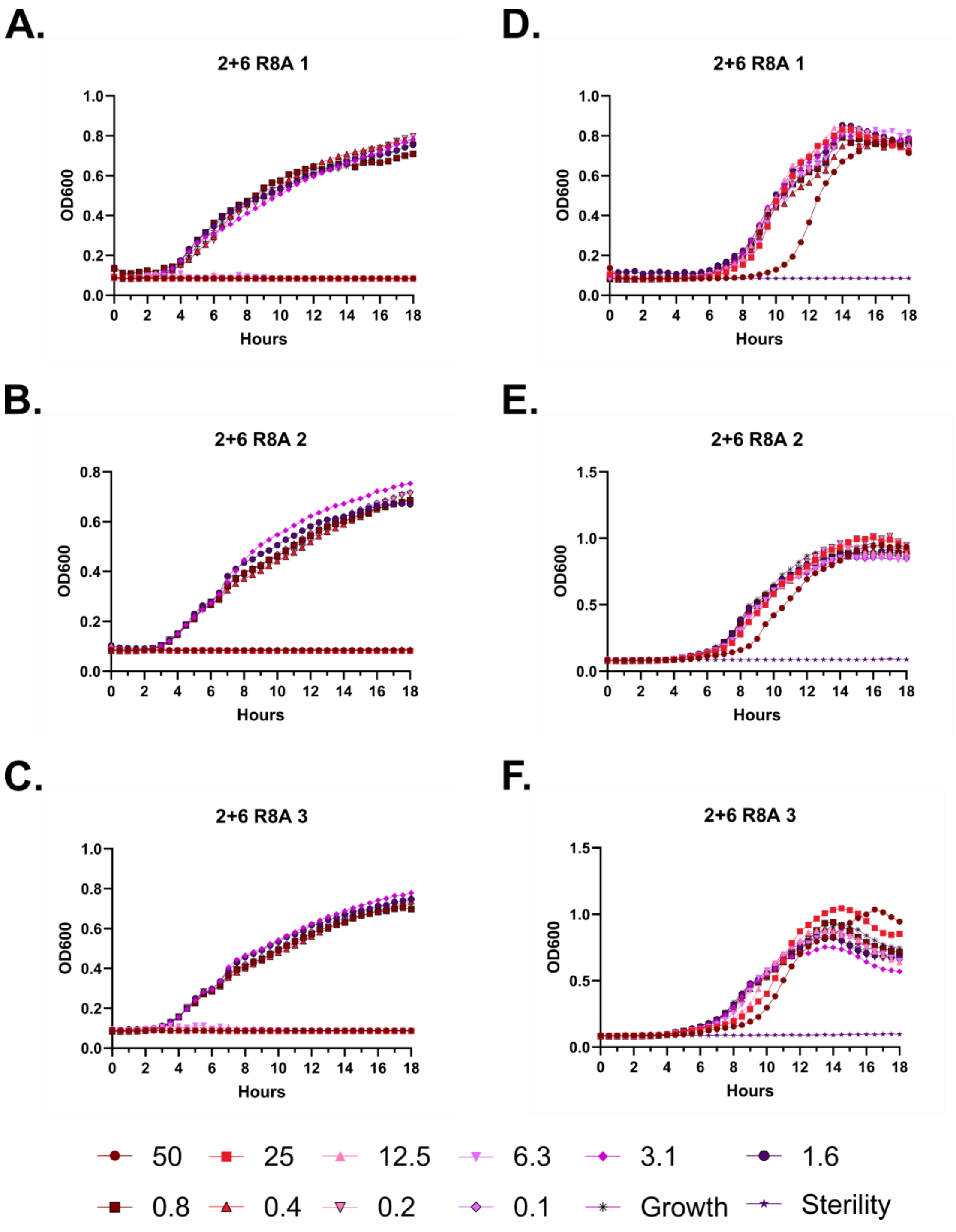
A.
2+6 R8A 1
OD600
Hours
D.
2+6 R8A 1
OD600
Hours
B.
2+6 R8A 2
OD600
Hours
E.
2+6 R8A 2
OD600
Hours
C.
2+6 R8A 3
OD600
Hours
F.
2+6 R8A 3
OD600
Hours
50
25
12.5
6.3
3.1
1.6
0.8
0.4
0.2
0.1
Growth
Sterility

**Figure S123 – Antimicrobial susceptibility testing with 2+6 R8A. A-C.** Growth of *E. coli* in the presence of the indicated peptide concentrations given in μM as monitored by OD600 at 37 °C over 18 hours. Each panel indicates an independent experiment. **D-F.** As in **A-C**, but with *P. aeruginosa.*

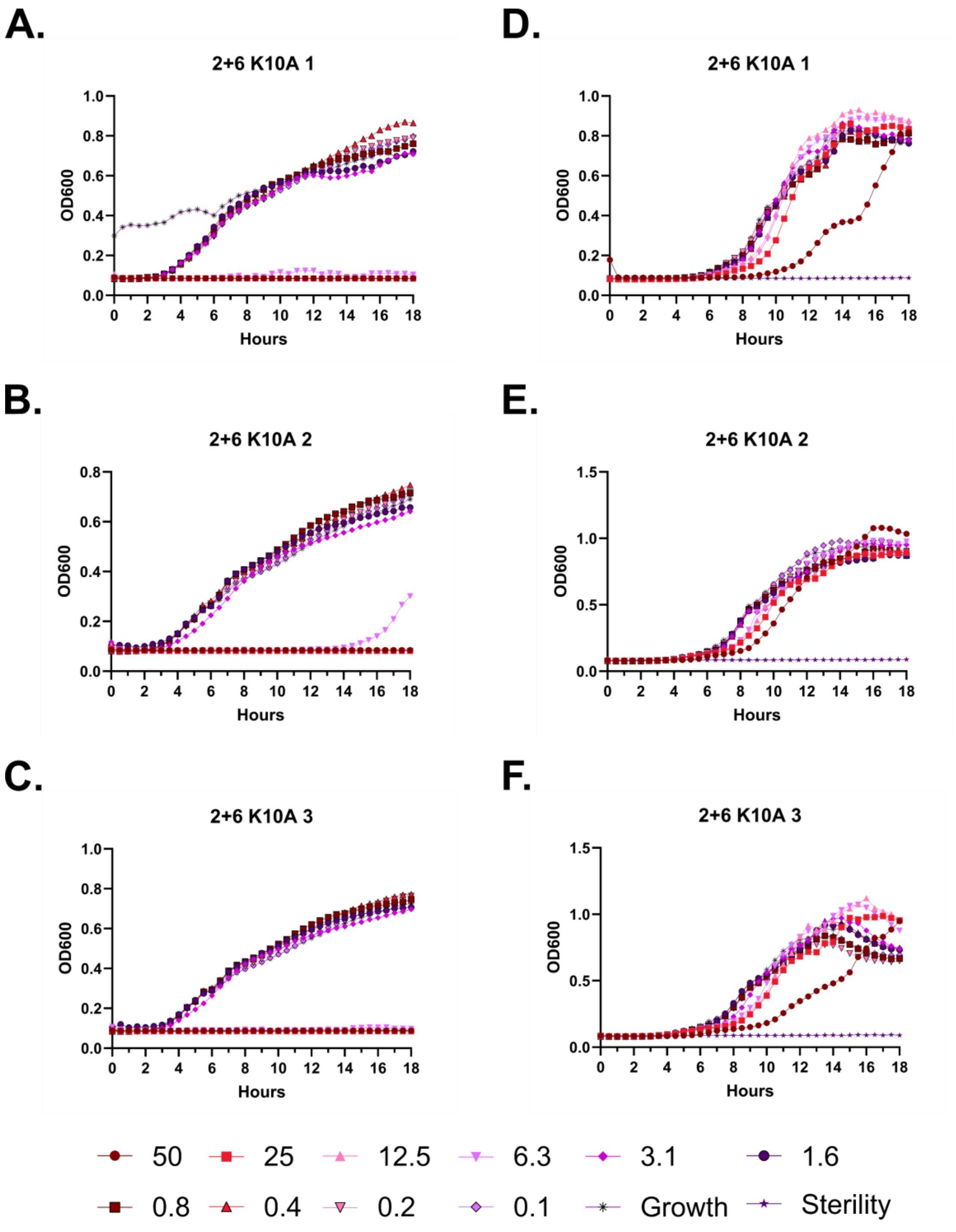
A.
2+6 K10A 1
OD600
Hours
B.
2+6 K10A 2
OD600
Hours
C.
2+6 K10A 3
OD600
Hours
D.
2+6 K10A 1
OD600
Hours
E.
2+6 K10A 2
OD600
Hours
F.
2+6 K10A 3
OD600
Hours
50
25
12.5
6.3
3.1
1.6
0.8
0.4
0.2
0.1
Growth
Sterility

**Figure S124 – Antimicrobial susceptibility testing with 2+6 K10A. A-C.** Growth of *E. coli* in the presence of the indicated peptide concentrations given in μM as monitored by OD600 at 37 °C over 18 hours. Each panel indicates an independent experiment. **D-F.** As in **A-C**, but with *P. aeruginosa.*

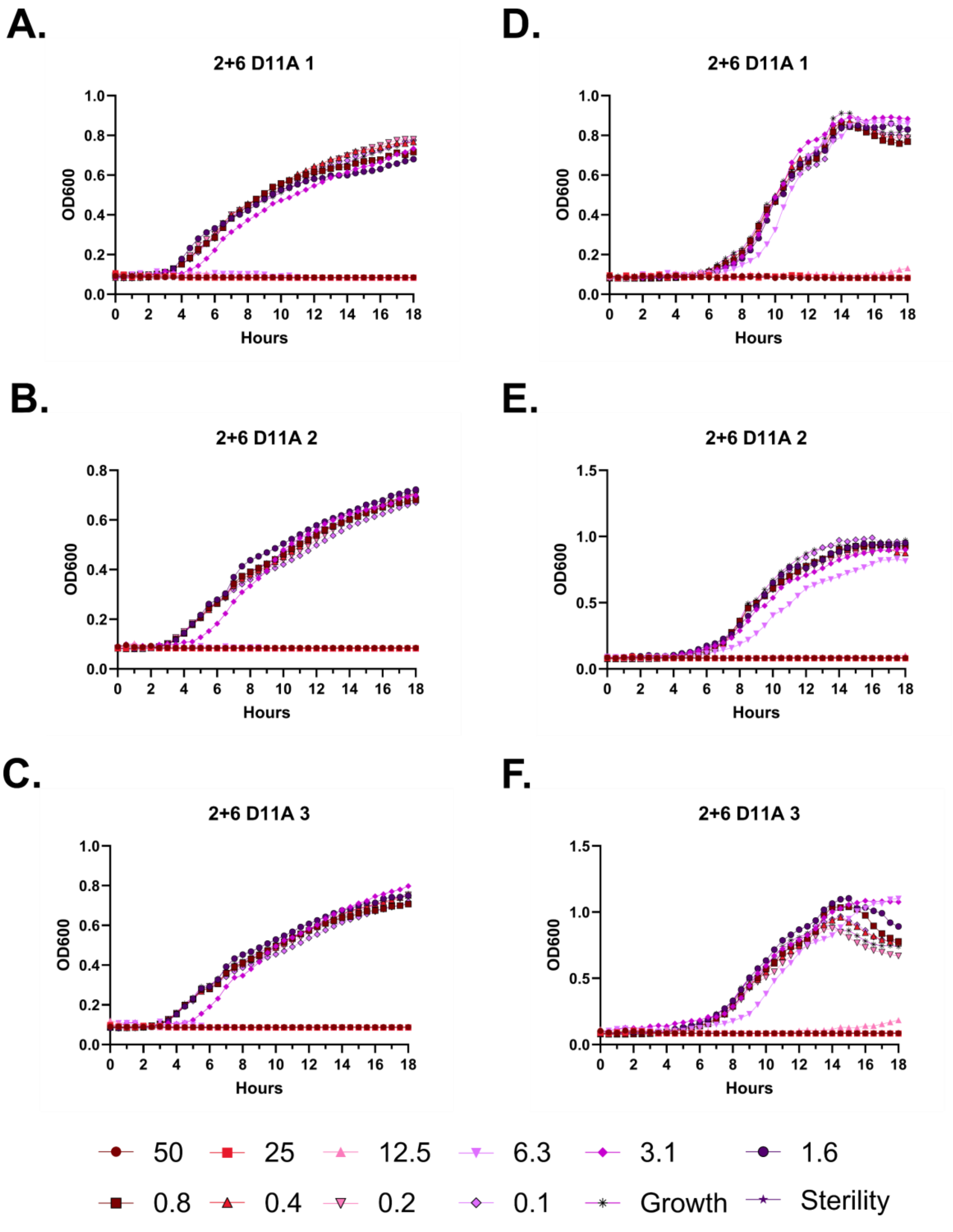

A.
2+6 D11A 1
OD600
Hours
B.
2+6 D11A 2
OD600
Hours
C.
2+6 D11A 3
OD600
Hours
D.
2+6 D11A 1
OD600
Hours
E.
2+6 D11A 2
OD600
Hours
F.
2+6 D11A 3
OD600
Hours
50
25
12.5
6.3
3.1
1.6
0.8
0.4
0.2
0.1
Growth
Sterility

**Figure S125 – Antimicrobial susceptibility testing with 2+6 D11A. A-C.** Growth of *E. coli* in the presence of the indicated peptide concentrations given in μM as monitored by OD600 at 37 °C over 18 hours. Each panel indicates an independent experiment. **D-F.** As in **A-C**, but with *P. aeruginosa.*

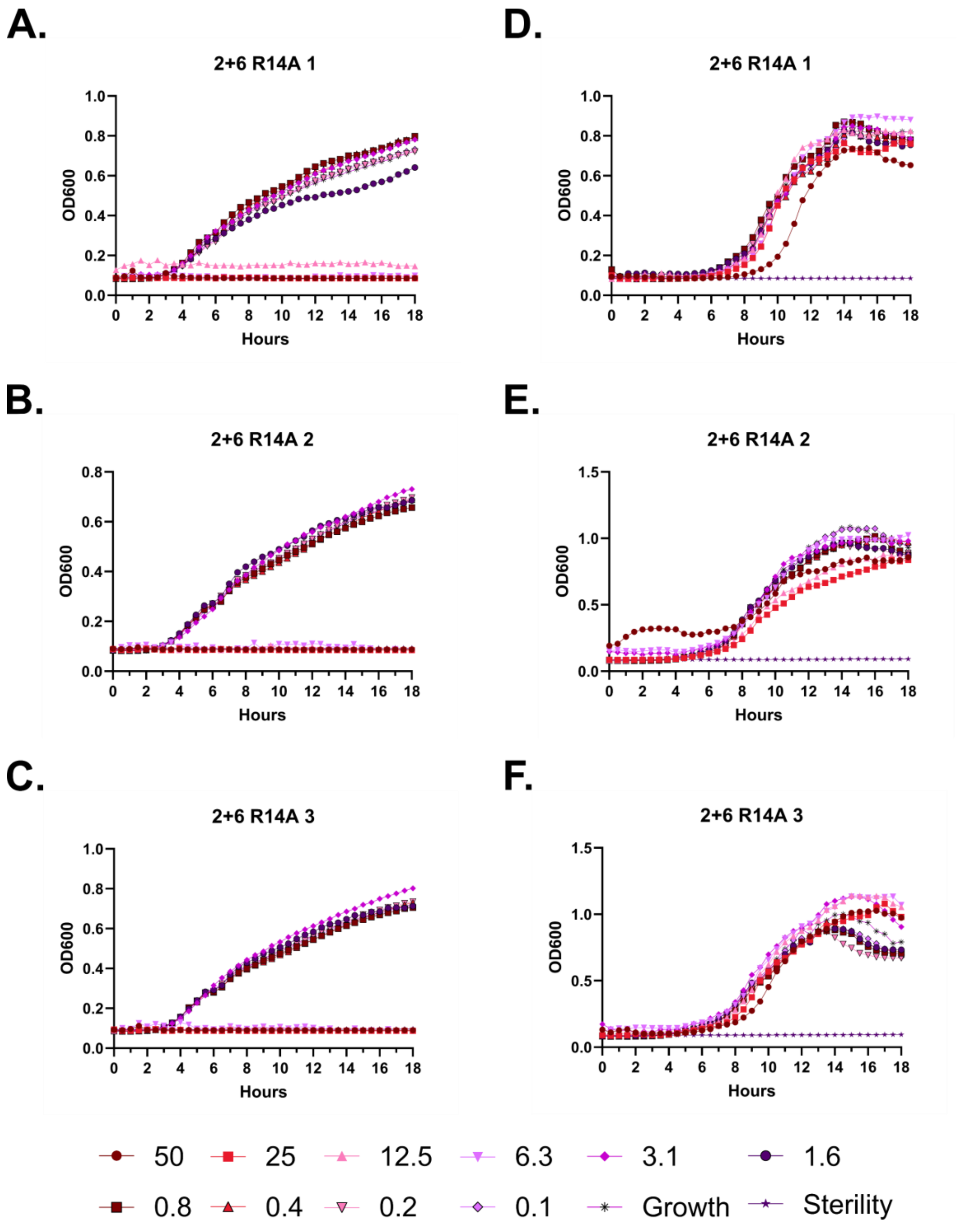
A.
2+6 R14A 1
OD600
Hours
D.
2+6 R14A 1
OD600
Hours
B.
2+6 R14A 2
OD600
Hours
E.
2+6 R14A 2
OD600
Hours
C.
2+6 R14A 3
OD600
Hours
F.
2+6 R14A 3
OD600
Hours
50
25
12.5
6.3
3.1
1.6
0.8
0.4
0.2
0.1
Growth
Sterility

**Figure S126 – Antimicrobial susceptibility testing with 2+6 R14A. A-C.** Growth of *E. coli* in the presence of the indicated peptide concentrations given in μM as monitored by OD600 at 37 °C over 18 hours. Each panel indicates an independent experiment. **D-F.** As in **A-C**, but with *P. aeruginosa.*

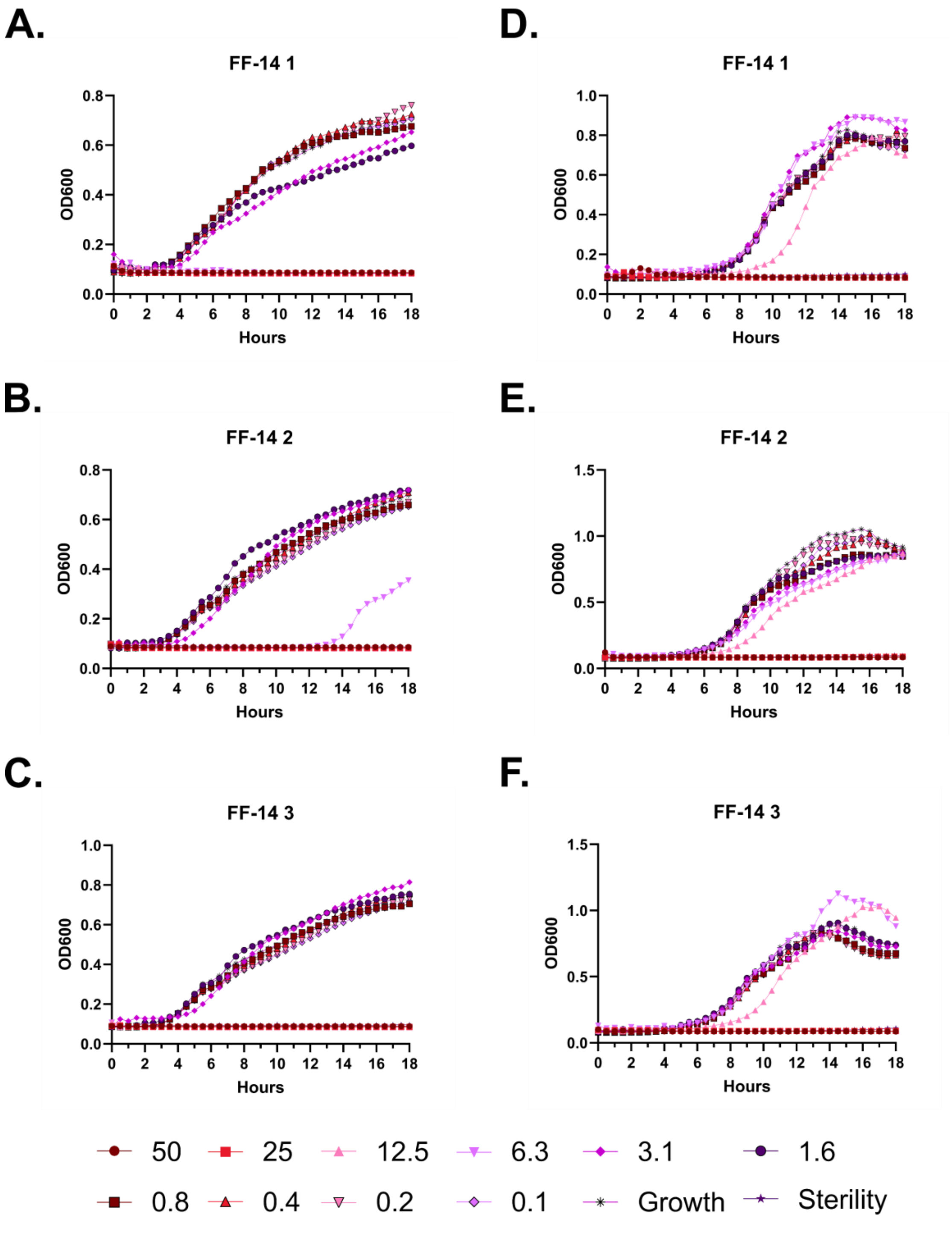

A.
FF-14 1
B.
FF-14 2
C.
FF-14 3
D.
FF-14 1
E.
FF-14 2
F.
FF-14 3
OD600
Hours
50
25
12.5
6.3
3.1
1.6
0.8
0.4
0.2
0.1
Growth
Sterility

**Figure S127 – Antimicrobial susceptibility testing with FF-14. A-C.** Growth of *E. coli* in the presence of the indicated peptide concentrations given in μM as monitored by OD600 at 37 °C over 18 hours. Each panel indicates an independent experiment. **D-F.** As in **A-C**, but with *P. aeruginosa.*

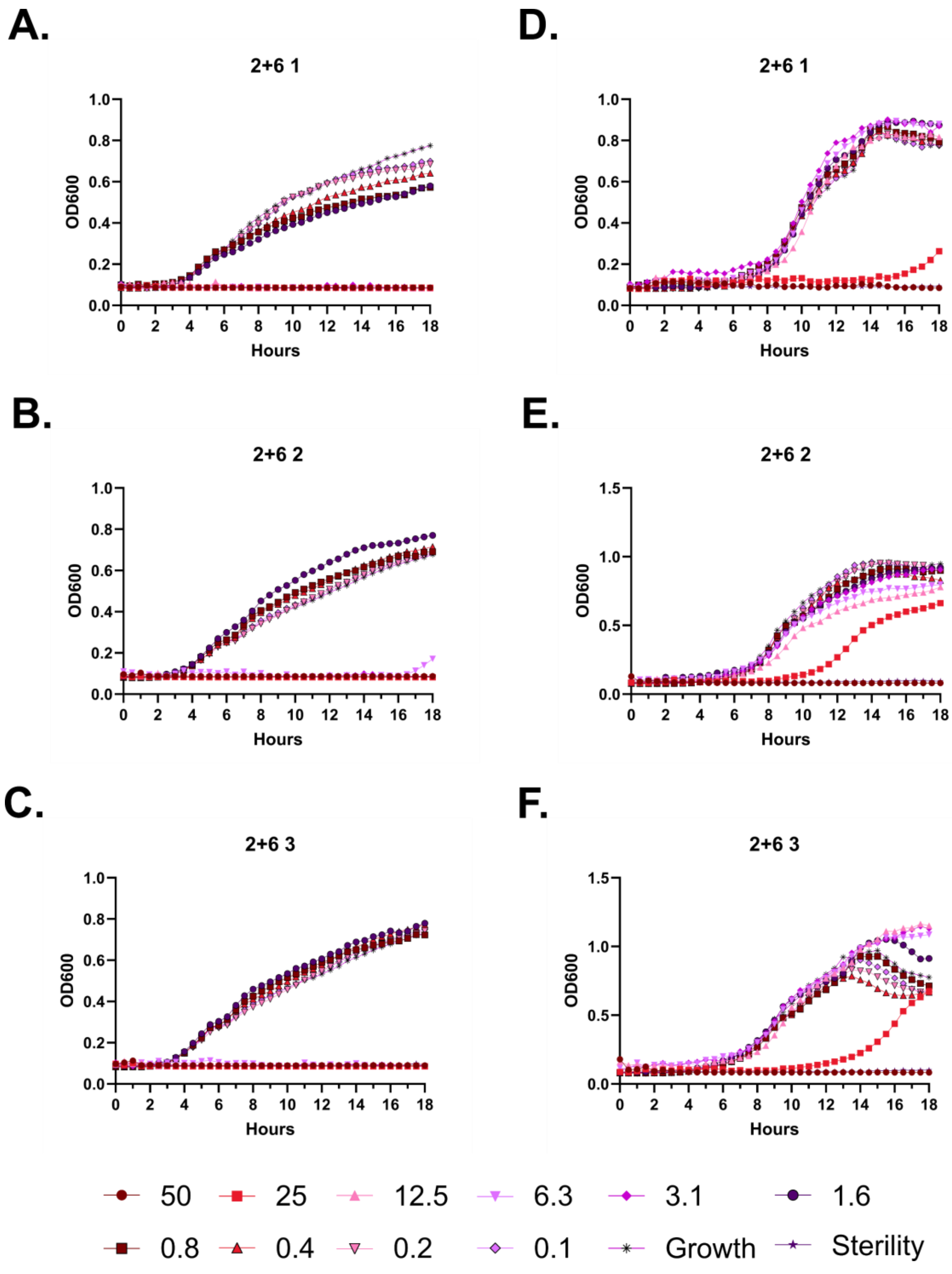
A.
2+6 1
OD600
Hours
D.
2+6 1
OD600
Hours
B.
2+6 2
OD600
Hours
E.
2+6 2
OD600
Hours
C.
2+6 3
OD600
Hours
F.
2+6 3
OD600
Hours
50
25
12.5
6.3
3.1
1.6
0.8
0.4
0.2
0.1
Growth
Sterility

**Figure S128 – Antimicrobial susceptibility testing with FF-14 F2B+V6B. A-C.** Growth of *E. coli* in the presence of the indicated peptide concentrations given in μM as monitored by OD600 at 37 °C over 18 hours. Each panel indicates an independent experiment. **D-F.** As in **A-C**, but with *P. aeruginosa*.

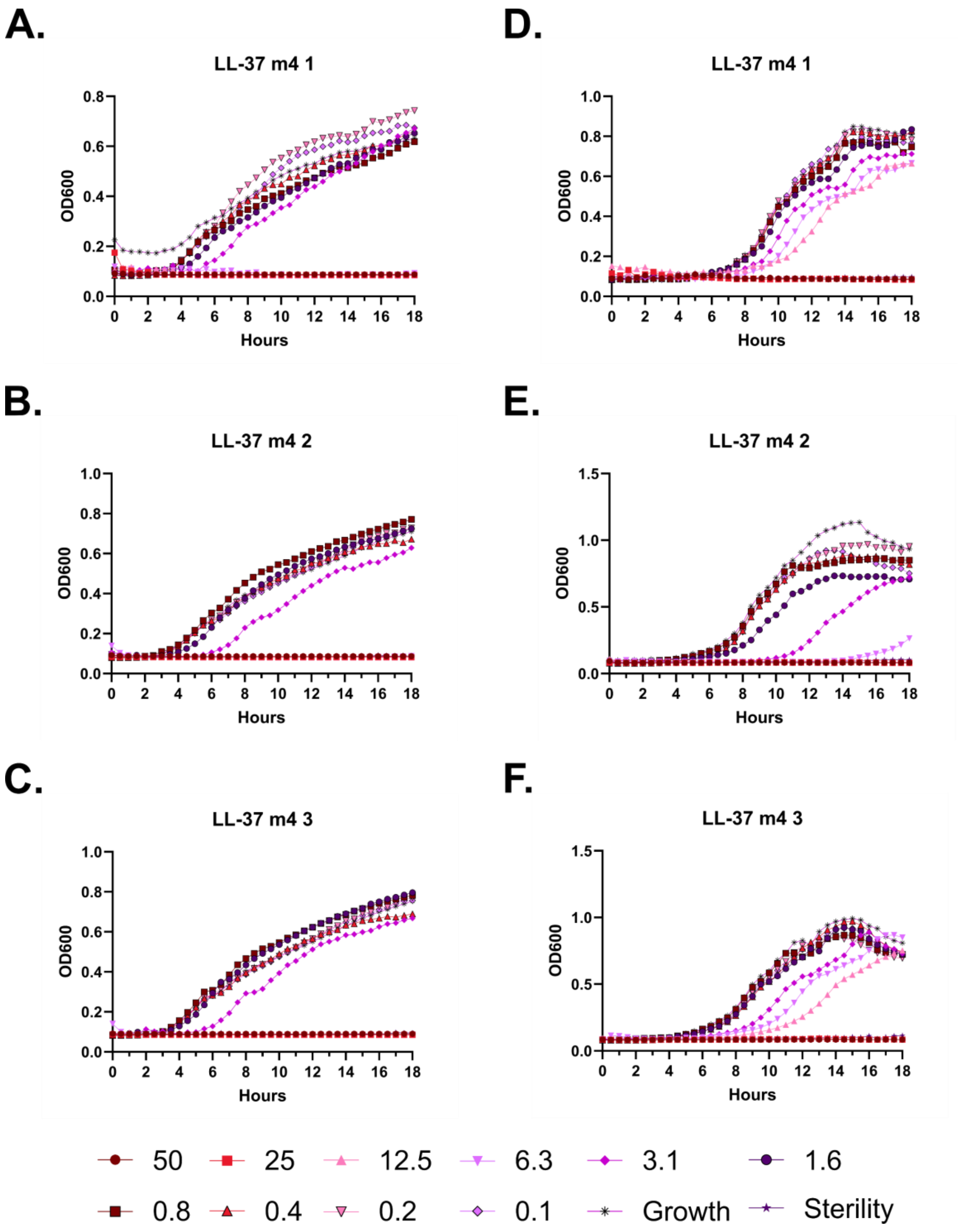
A.
LL-37 m4 1
B.
LL-37 m4 2
C.
LL-37 m4 3
D.
LL-37 m4 1
E.
LL-37 m4 2
F.
LL-37 m4 3
OD600
Hours
50
25
12.5
6.3
3.1
1.6
0.8
0.4
0.2
0.1
Growth
Sterility

**Figure S129 – Antimicrobial susceptibility testing with LL-37 m4. A-C.** Growth of *E. coli* in the presence of the indicated peptide concentrations given in μM as monitored by OD600 at 37 °C over 18 hours. Each panel indicates an independent experiment. **D-F.** As in **A-C**, but with *P. aeruginosa.*

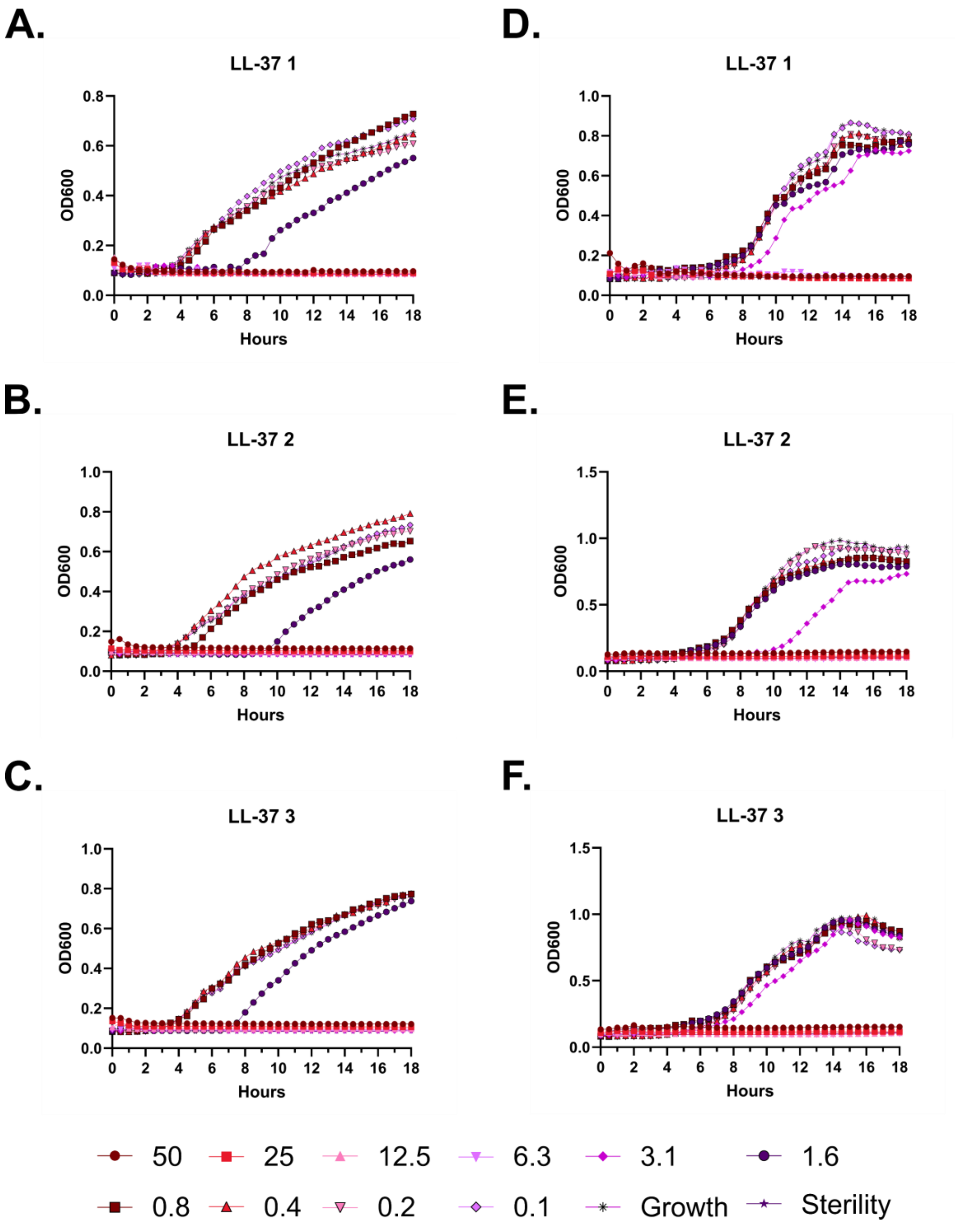
A.
LL-37 1
OD600
Hours
B.
LL-37 2
OD600
Hours
C.
LL-37 3
OD600
Hours
D.
LL-37 1
OD600
Hours
E.
LL-37 2
OD600
Hours
F.
LL-37 3
OD600
Hours
50
25
12.5
6.3
3.1
1.6
0.8
0.4
0.2
0.1
Growth
Sterility

**Figure S130 – Antimicrobial susceptibility testing with LL-37. A-C.** Growth of *E. coli* in the presence of the indicated peptide concentrations given in μM as monitored by OD600 at 37 °C over 18 hours. Each panel indicates an independent experiment. **D-F.** As in **A-C**, but with *P. aeruginosa.*

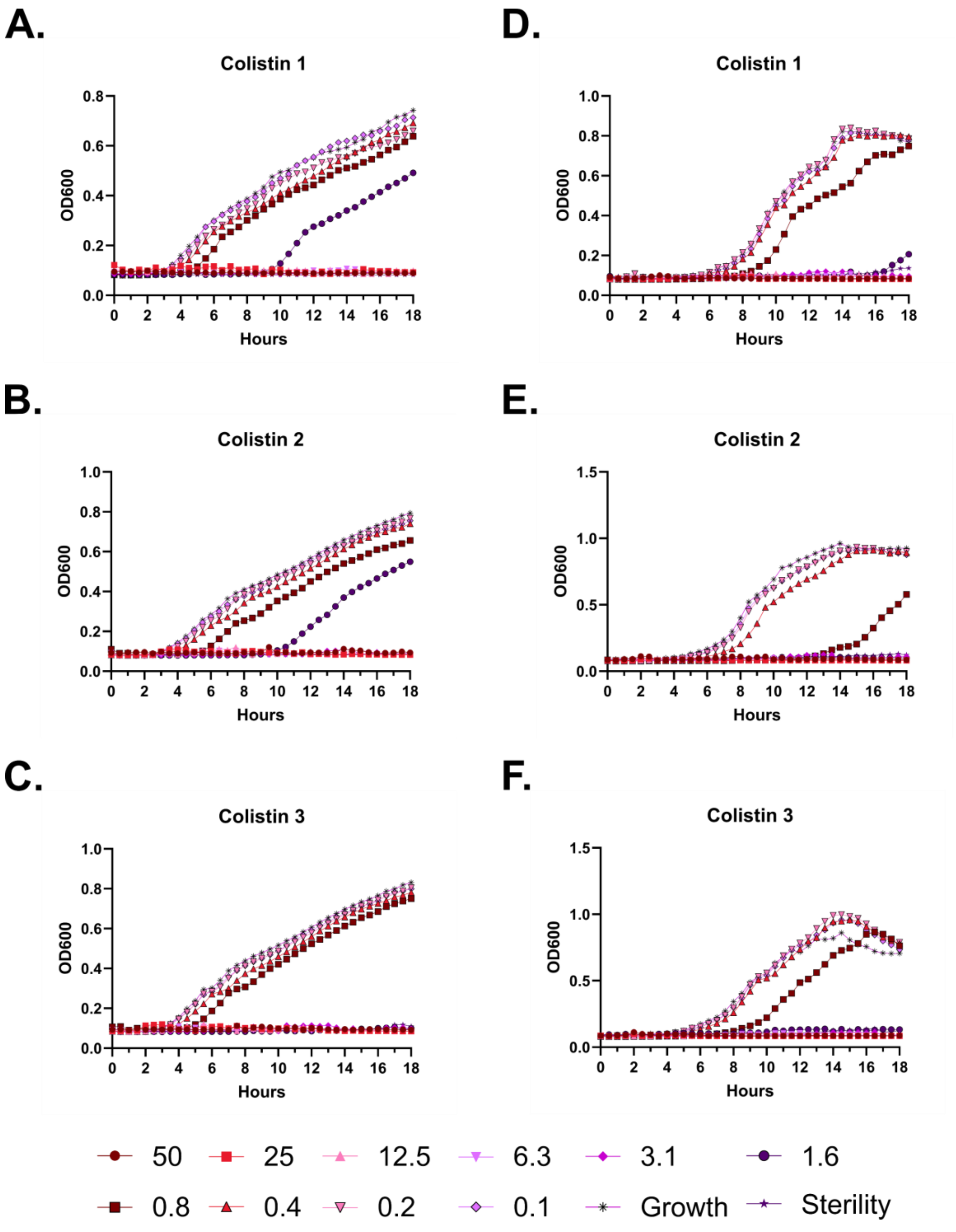
A.
Colistin 1
OD600
Hours
D.
Colistin 1
OD600
Hours
B.
Colistin 2
OD600
Hours
E.
Colistin 2
OD600
Hours
C.
Colistin 3
OD600
Hours
F.
Colistin 3
OD600
Hours
50
25
12.5
6.3
3.1
1.6
0.8
0.4
0.2
0.1
Growth
Sterility

**Figure S131 – Antimicrobial susceptibility testing with colistin. A-C.** Growth of *E. coli* in the presence of the indicated peptide concentrations given in µM as monitored by OD600 at 37 °C over 18 hours. Each panel indicates an independent experiment. **D-F.** As in **A-C**, but with *P. aeruginosa.*

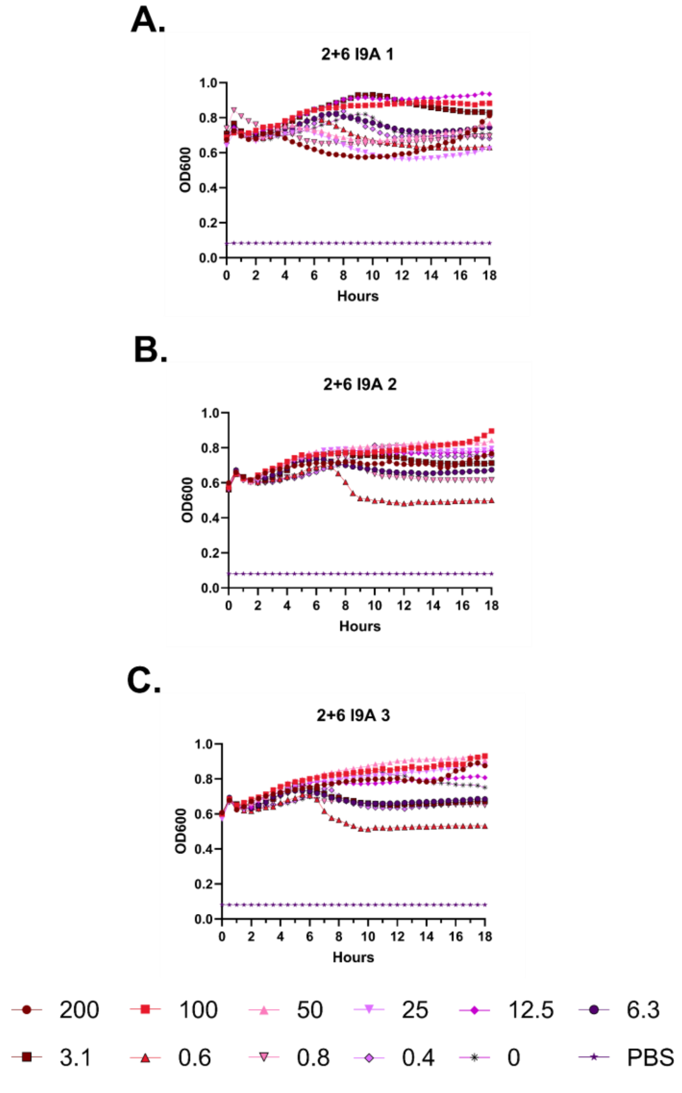
A.
2+6 I9A 1
OD600
Hours
B.
2+6 I9A 2
OD600
Hours
C.
2+6 I9A 3
OD600
Hours
200
100
50
25
12.5
6.3
3.1
0.6
0.8
0.4
0
PBS

**Figure S132 – Hemolysis over time with 2+6 I9A. A-C.** Each of three independent experiments showing trends in OD600 over time on incubation of defibrinated sheep blood with the indicated concentrations of peptide in μM at 37 °C. Decreasing OD600 indicates increasing hemolysis.

**A.**

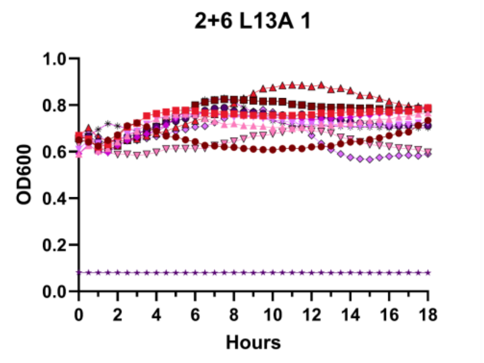
2+6 L13A 1
OD600
1.0
0.8
0.6
0.4
0.2
0.0
0
2
4
6
8
10
12
14
16
18
Hours

**B.**

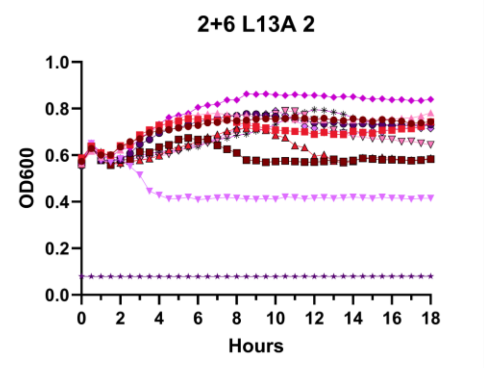
2+6 L13A 2
OD600
1.0
0.8
0.6
0.4
0.2
0.0
0
2
4
6
8
10
12
14
16
18
Hours

**C.**

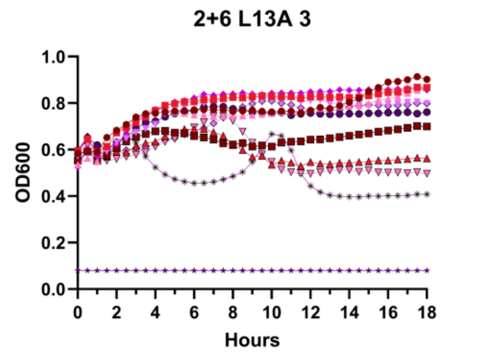
2+6 L13A 3
OD600
1.0
0.8
0.6
0.4
0.2
0.0
0
2
4
6
8
10
12
14
16
18
Hours

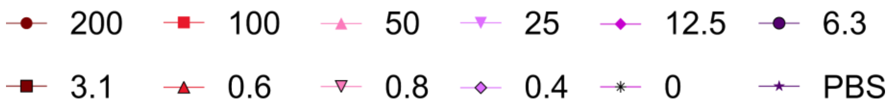
200
100
50
25
12.5
6.3
3.1
0.6
0.8
0.4
0
PBS

**Figure S133 – Hemolysis over time with 2+6 L13A. A-C.** Each of three independent experiments showing trends in OD600 over time on incubation of defibrinated sheep blood with the indicated concentrations of peptide in μM at 37 °C. Decreasing OD600 indicates increasing hemolysis.

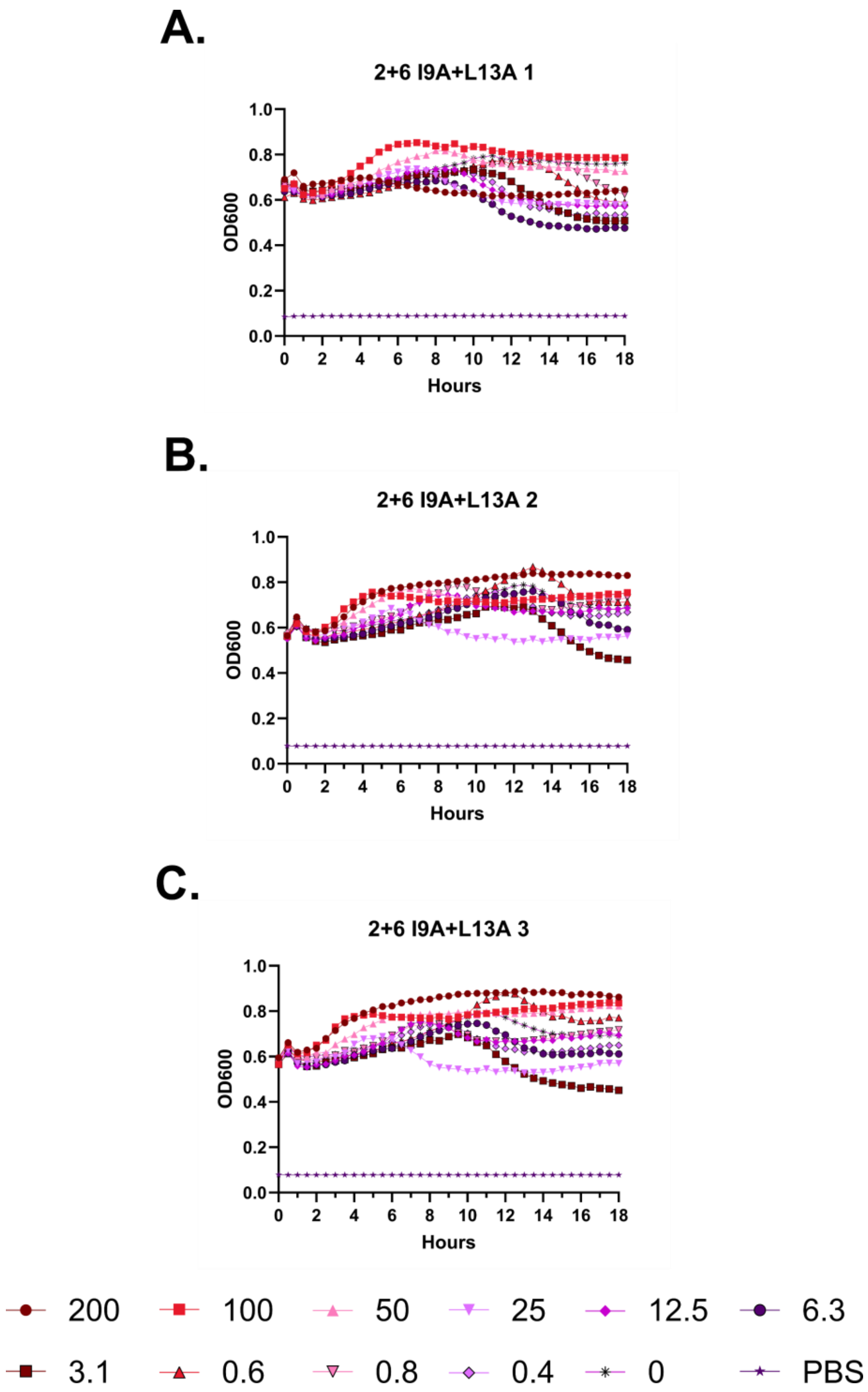
A.
2+6 I9A+L13A 1
B.
2+6 I9A+L13A 2
C.
2+6 I9A+L13A 3
OD600
Hours
1.0
0.8
0.6
0.4
0.2
0.0
0 2 4 6 8 10 12 14 16 18
200
100
50
25
12.5
6.3
3.1
0.6
0.8
0.4
0
PBS

**Figure S134 – Hemolysis over time with 2+6 I9A+L13A. A-C.** Each of three independent experiments showing trends in OD600 over time on incubation of defibrinated sheep blood with the indicated concentrations of peptide in μM at 37 °C. Decreasing OD600 indicates increasing hemolysis.

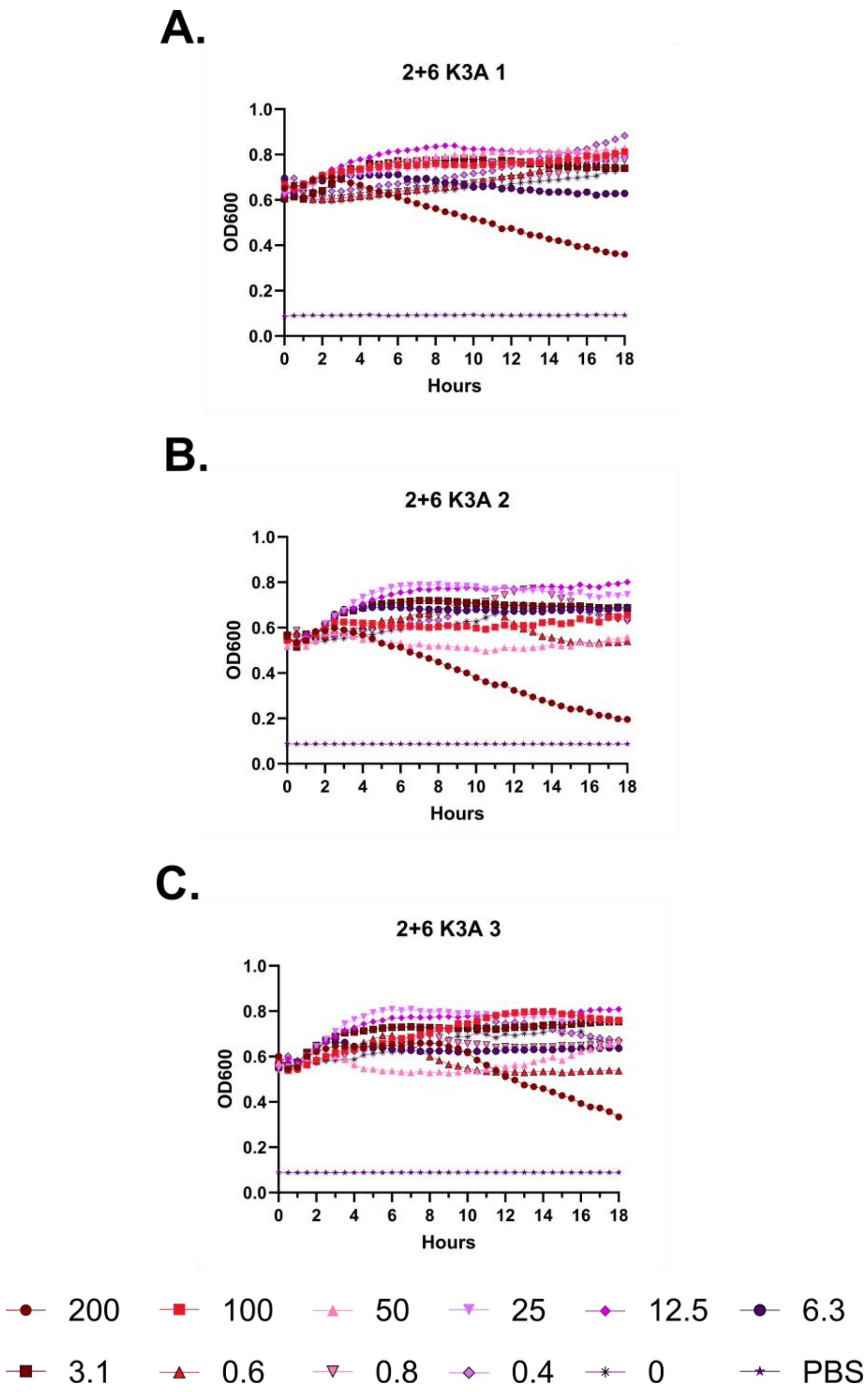
A.
2+6 K3A 1
OD600
Hours
B.
2+6 K3A 2
OD600
Hours
C.
2+6 K3A 3
OD600
Hours
200
100
50
25
12.5
6.3
3.1
0.6
0.8
0.4
0
PBS

**Figure S135 – Hemolysis over time with 2+6 K3A. A-C.** Each of three independent experiments showing trends in OD600 over time on incubation of defibrinated sheep blood with the indicated concentrations of peptide in μM at 37 °C. Decreasing OD600 indicates increasing hemolysis.

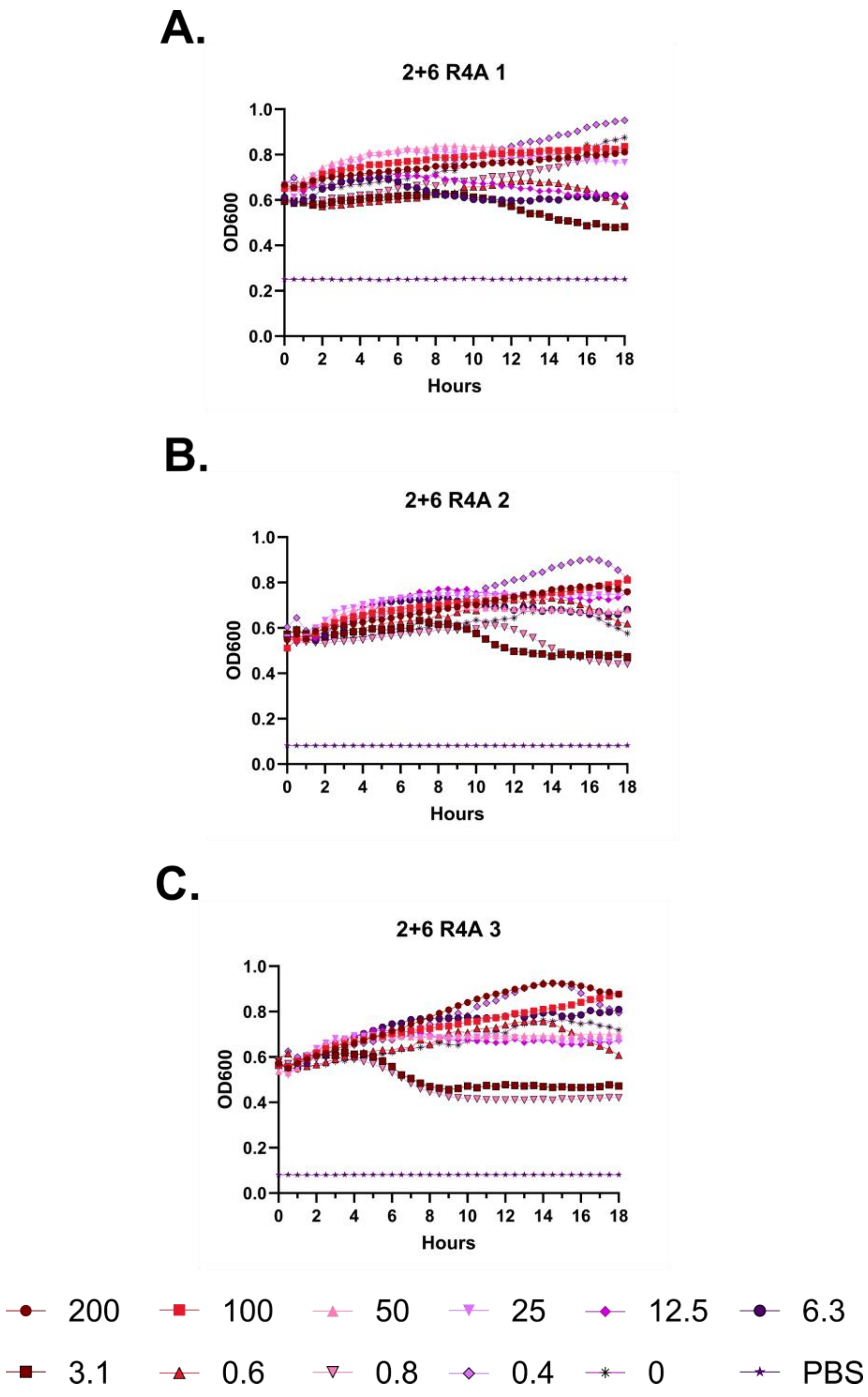
A.
2+6 R4A 1
OD600
Hours
B.
2+6 R4A 2
OD600
Hours
C.
2+6 R4A 3
OD600
Hours
200
100
50
25
12.5
6.3
3.1
0.6
0.8
0.4
0
PBS

**Figure S136 – Hemolysis over time with 2+6 R4A. A-C.** Each of three independent experiments showing trends in OD600 over time on incubation of defibrinated sheep blood with the indicated concentrations of peptide in μM at 37 °C. Decreasing OD600 indicates increasing hemolysis.

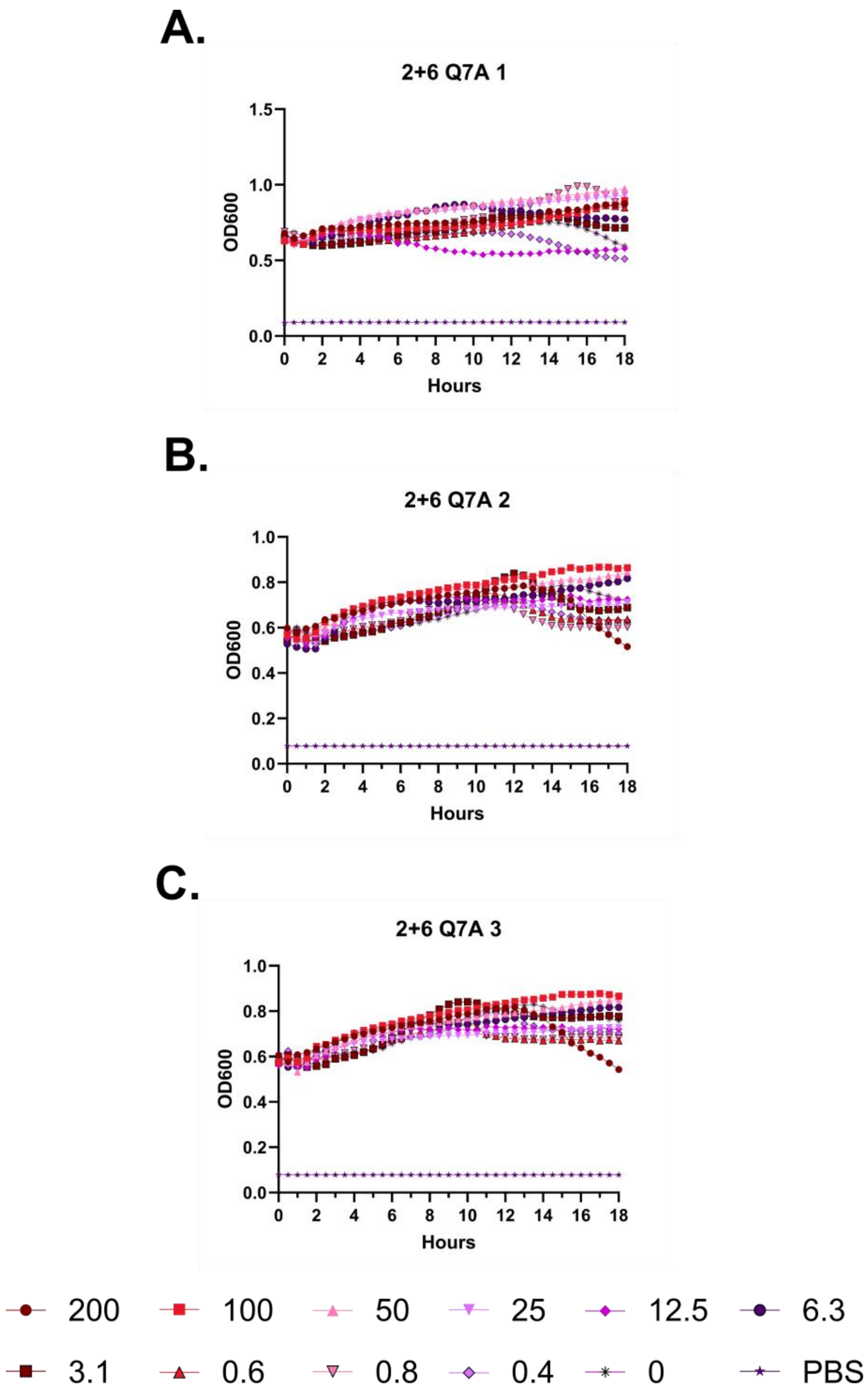
A.
2+6 Q7A 1
OD600
Hours
B.
2+6 Q7A 2
OD600
Hours
C.
2+6 Q7A 3
OD600
Hours
200
100
50
25
12.5
6.3
3.1
0.6
0.8
0.4
0
PBS

**Figure S137 – Hemolysis over time with 2+6 Q7A. A-C.** Each of three independent experiments showing trends in OD600 over time on incubation of defibrinated sheep blood with the indicated concentrations of peptide in μM at 37 °C. Decreasing OD600 indicates increasing hemolysis.

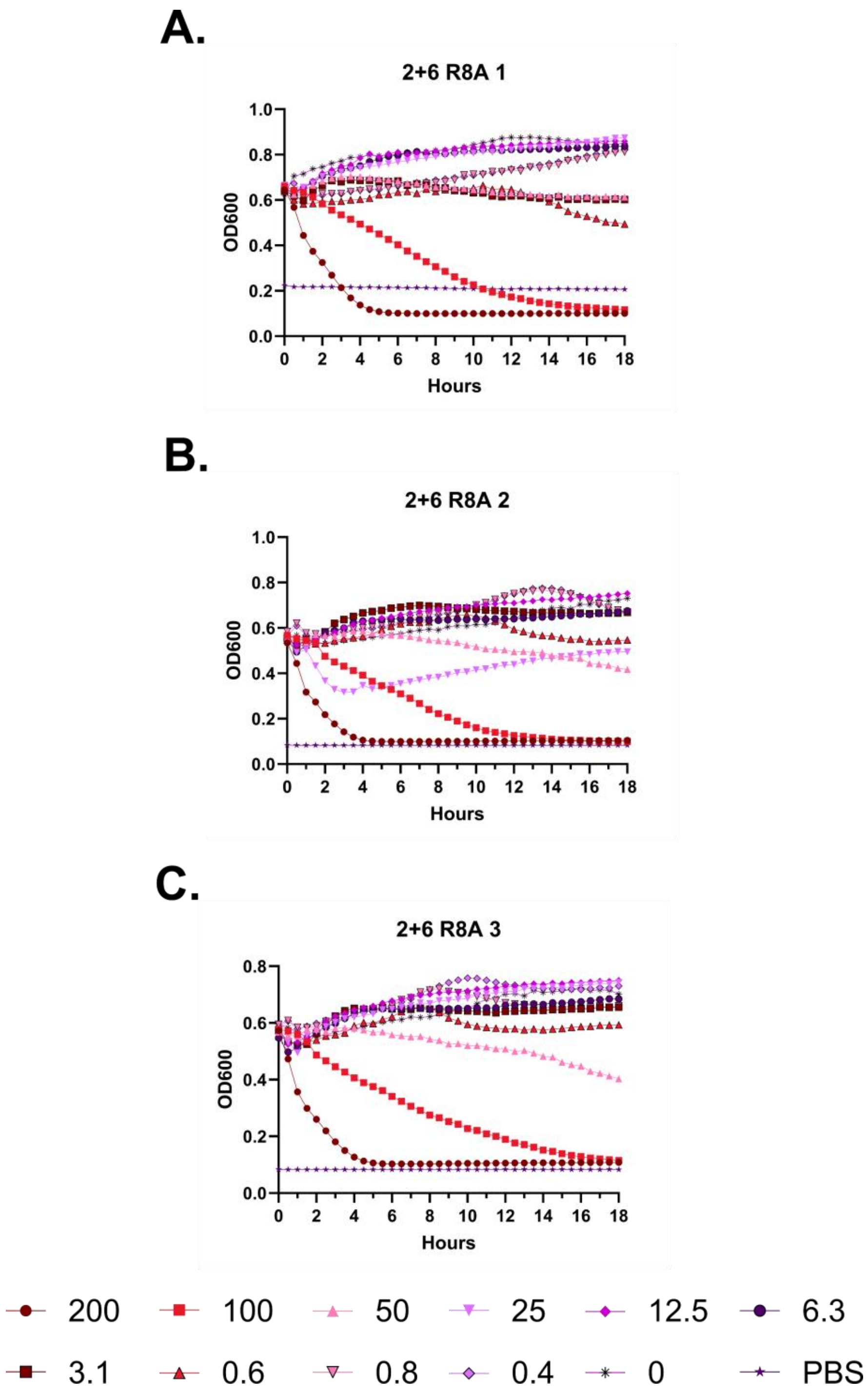
A.
2+6 R8A 1
OD600
Hours
B.
2+6 R8A 2
OD600
Hours
C.
2+6 R8A 3
OD600
Hours
200
100
50
25
12.5
6.3
3.1
0.6
0.8
0.4
0
PBS

**Figure S138 – Hemolysis over time with 2+6 R8A. A-C.** Each of three independent experiments showing trends in OD600 over time on incubation of defibrinated sheep blood with the indicated concentrations of peptide in μM at 37 °C. Decreasing OD600 indicates increasing hemolysis.

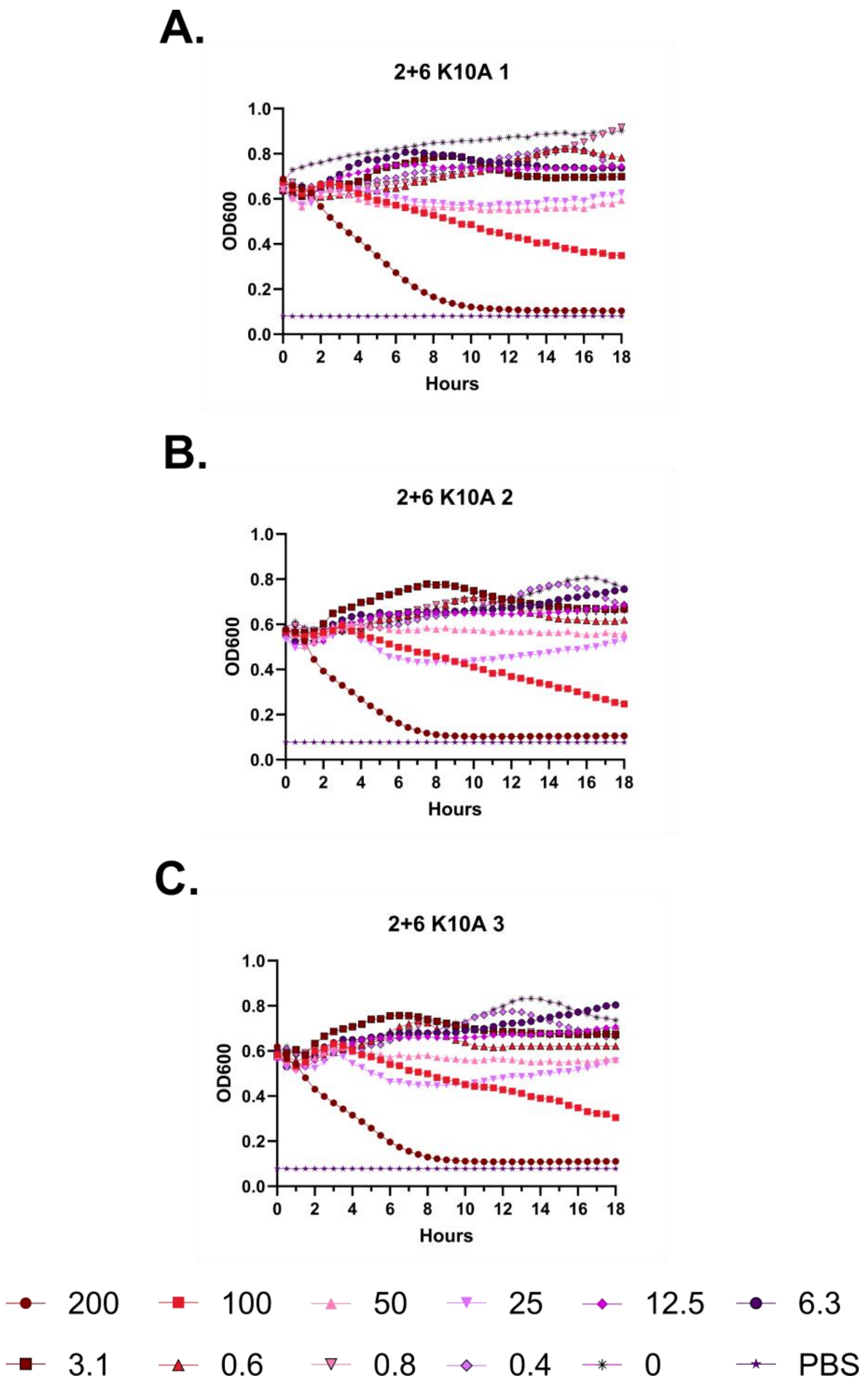
A.
2+6 K10A 1
OD600
Hours
B.
2+6 K10A 2
OD600
Hours
C.
2+6 K10A 3
OD600
Hours
200
100
50
25
12.5
6.3
3.1
0.6
0.8
0.4
0
PBS

**Figure S139 – Hemolysis over time with 2+6 K10A. A-C.** Each of three independent experiments showing trends in OD600 over time on incubation of defibrinated sheep blood with the indicated concentrations of peptide in μM at 37 °C. Decreasing OD600 indicates increasing hemolysis.

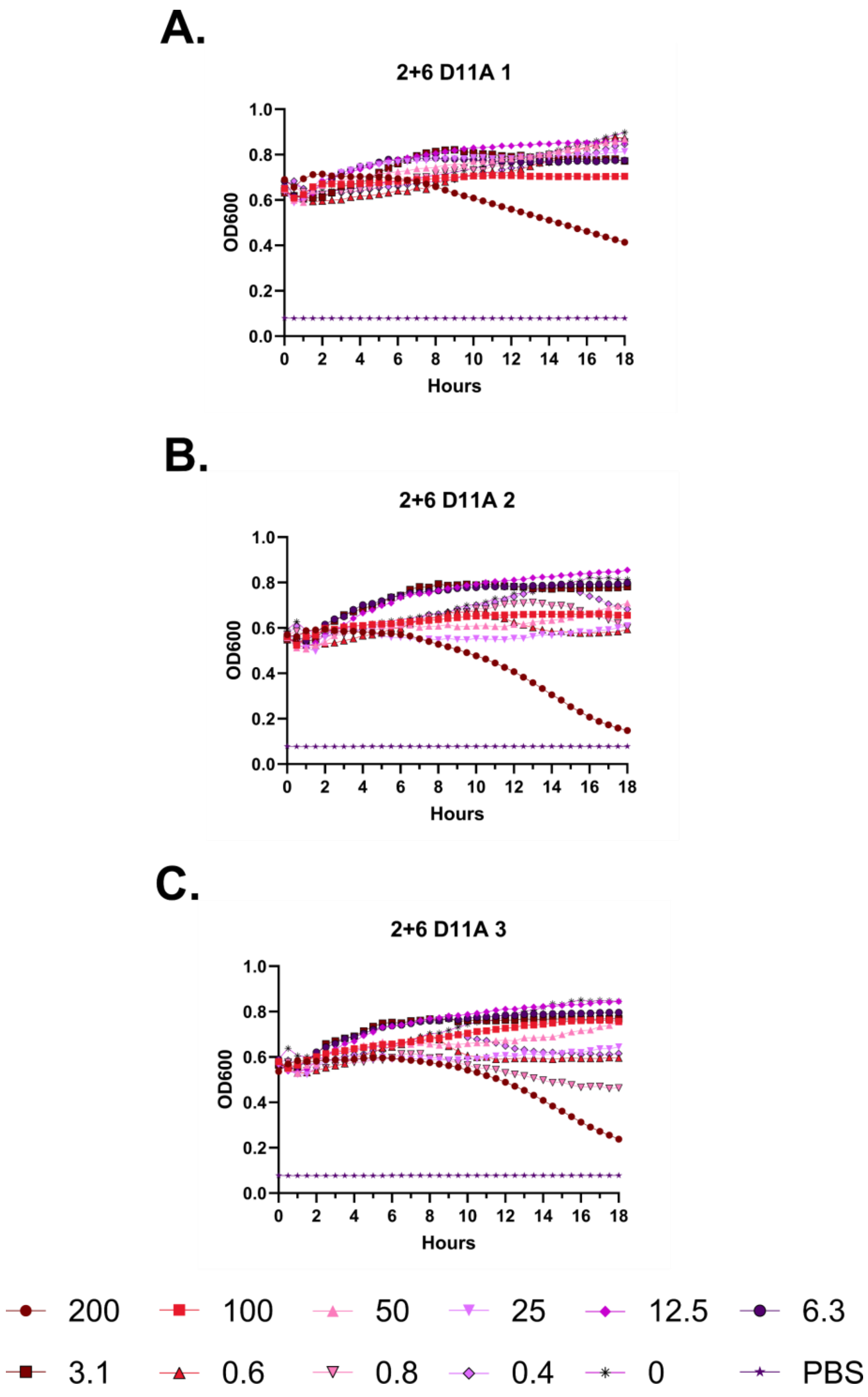
A.
2+6 D11A 1
OD600
Hours
B.
2+6 D11A 2
C.
2+6 D11A 3
200
100
50
25
12.5
6.3
3.1
0.6
0.8
0.4
0
PBS

**Figure S140 – Hemolysis over time with 2+6 D11A. A-C.** Each of three independent experiments showing trends in OD600 over time on incubation of defibrinated sheep blood with the indicated concentrations of peptide in μM at 37 °C. Decreasing OD600 indicates increasing hemolysis.

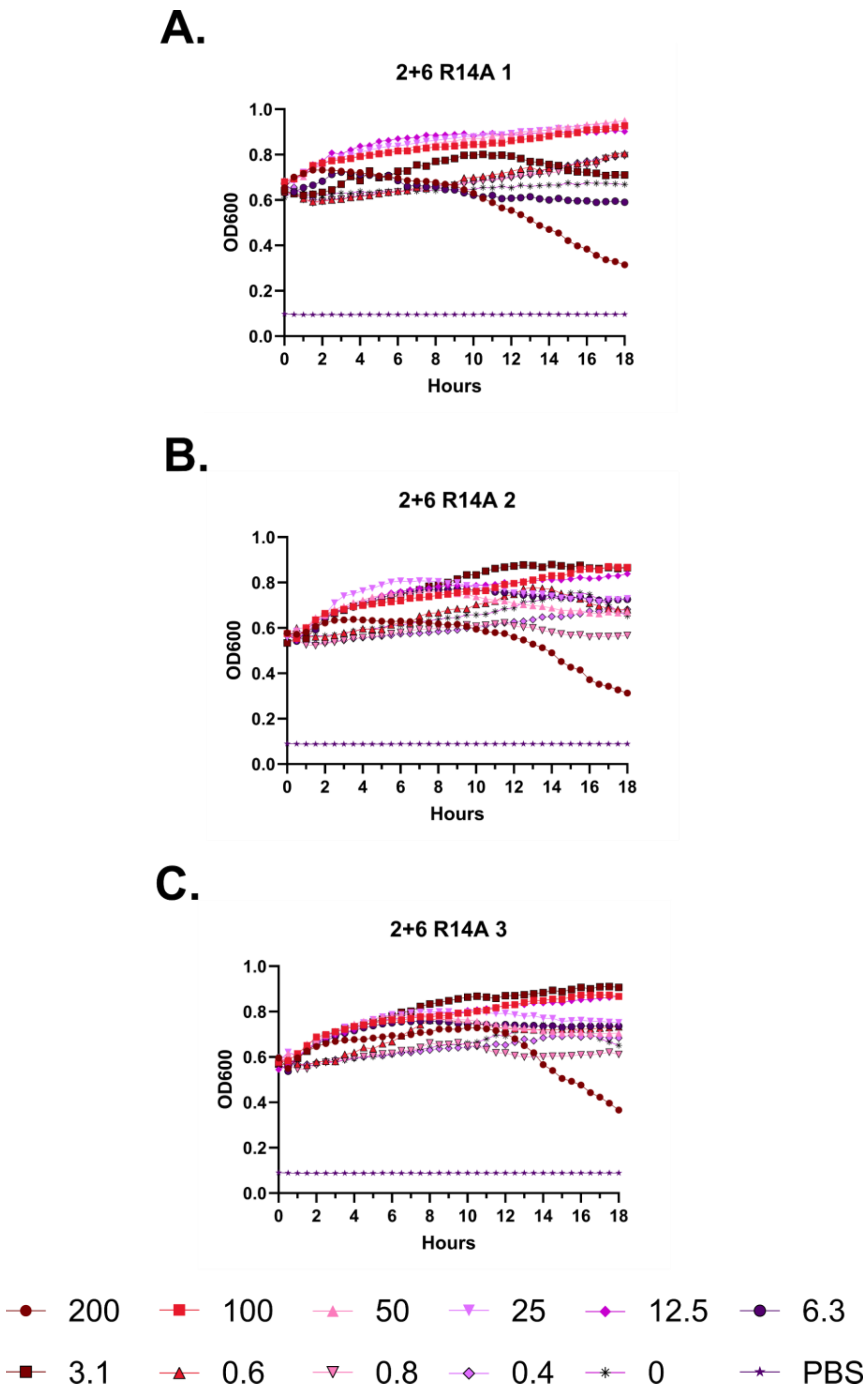
A.
2+6 R14A 1
OD600
Hours
B.
2+6 R14A 2
OD600
Hours
C.
2+6 R14A 3
OD600
Hours
200
100
50
25
12.5
6.3
3.1
0.6
0.8
0.4
0
PBS

**Figure S141 – Hemolysis over time with 2+6 R14A. A-C.** Each of three independent experiments showing trends in OD600 over time on incubation of defibrinated sheep blood with the indicated concentrations of peptide in μM at 37 °C. Decreasing OD600 indicates increasing hemolysis.

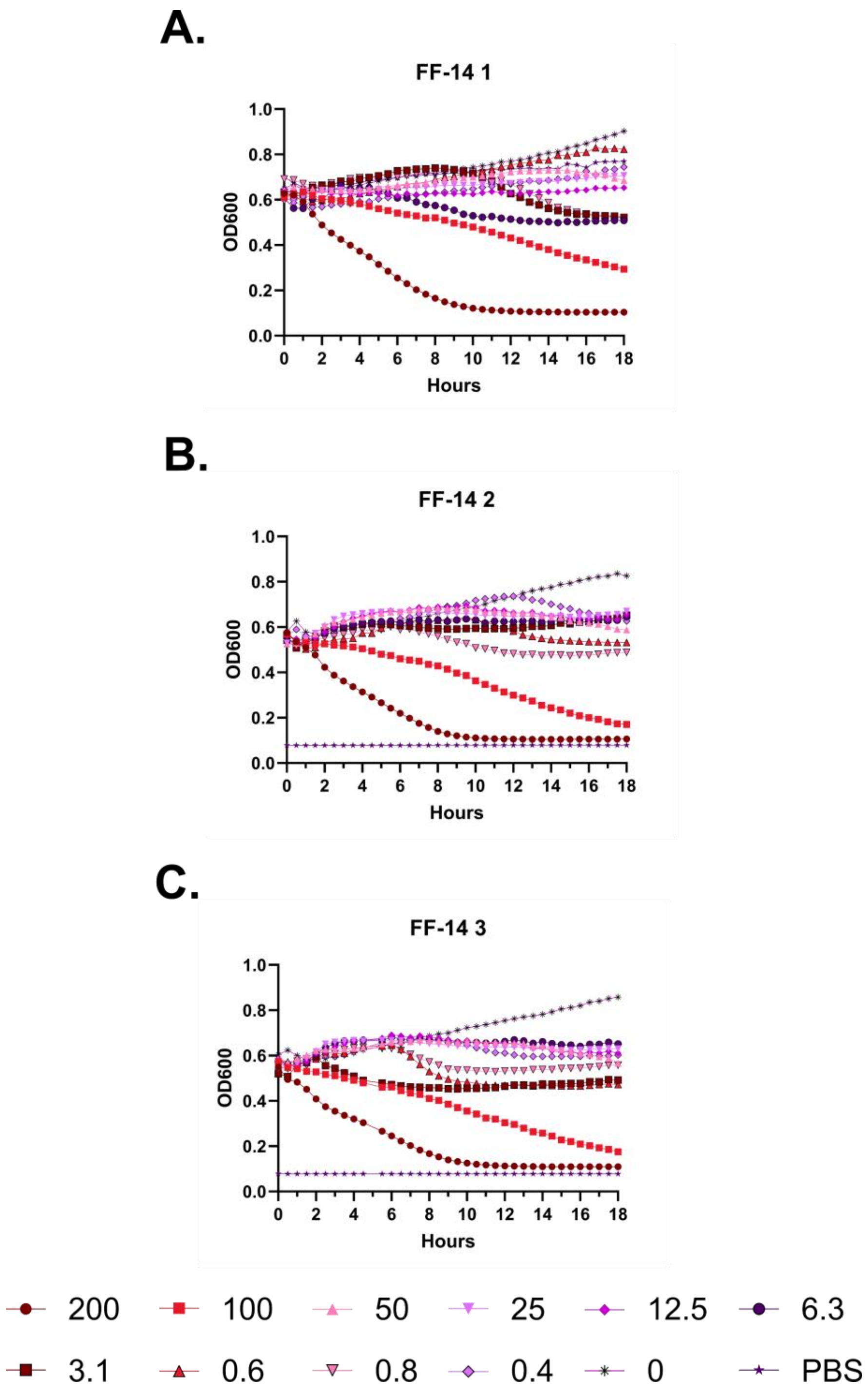
A.
FF-14 1
OD600
Hours
B.
FF-14 2
OD600
Hours
C.
FF-14 3
OD600
Hours
200
100
50
25
12.5
6.3
3.1
0.6
0.8
0.4
0
PBS

**Figure S142 – Hemolysis over time with FF-14. A-C.** Each of three independent experiments showing trends in OD600 over time on incubation of defibrinated sheep blood with the indicated concentrations of peptide in μM at 37 °C. Decreasing OD600 indicates increasing hemolysis.

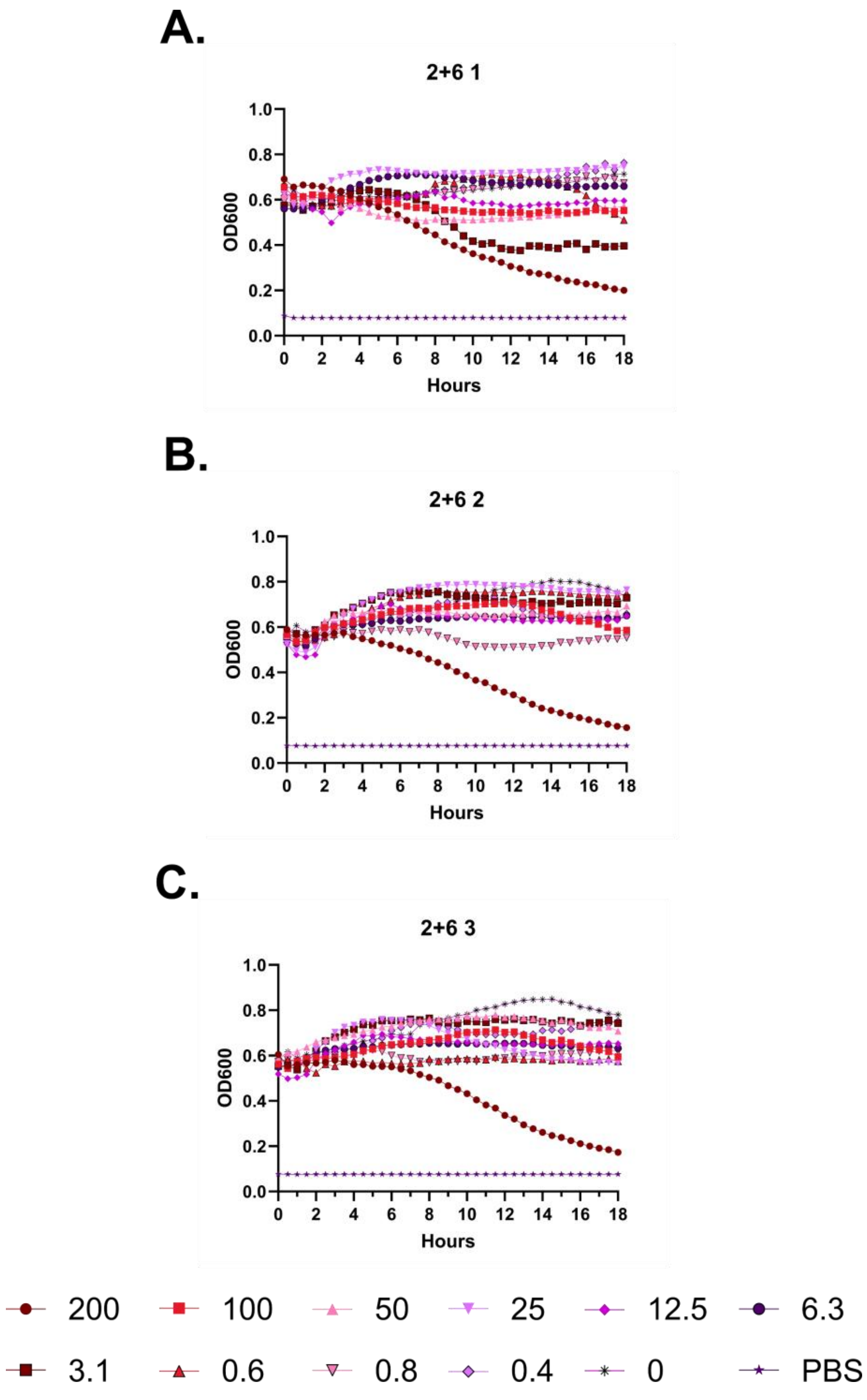
A.
2+6 1
OD600
Hours
B.
2+6 2
OD600
Hours
C.
2+6 3
OD600
Hours
200
100
50
25
12.5
6.3
3.1
0.6
0.8
0.4
0
PBS

**Figure S143 – Hemolysis over time with FF-14 F2B+V6B. A-C.** Each of three independent experiments showing trends in OD600 over time on incubation of defibrinated sheep blood with the indicated concentrations of peptide in μM at 37 °C. Decreasing OD600 indicates increasing hemolysis.

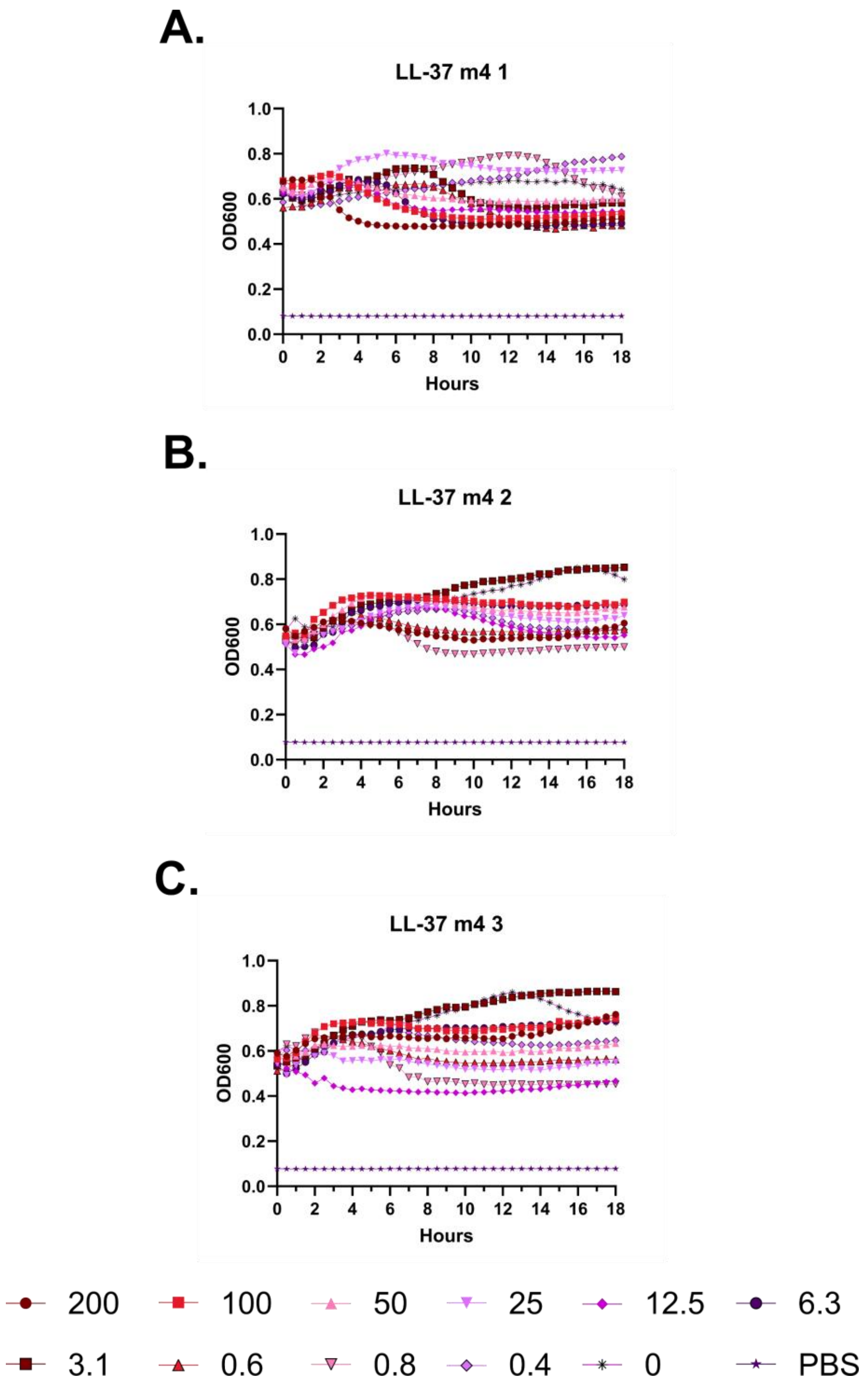
A.
LL-37 m4 1
OD600
Hours
B.
LL-37 m4 2
OD600
Hours
C.
LL-37 m4 3
OD600
Hours
200
100
50
25
12.5
6.3
3.1
0.6
0.8
0.4
0
PBS

**Figure S144 – Hemolysis over time with LL-37 m4. A-C.** Each of three independent experiments showing trends in OD600 over time on incubation of defibrinated sheep blood with the indicated concentrations of peptide in μM at 37 °C. Decreasing OD600 indicates increasing hemolysis.

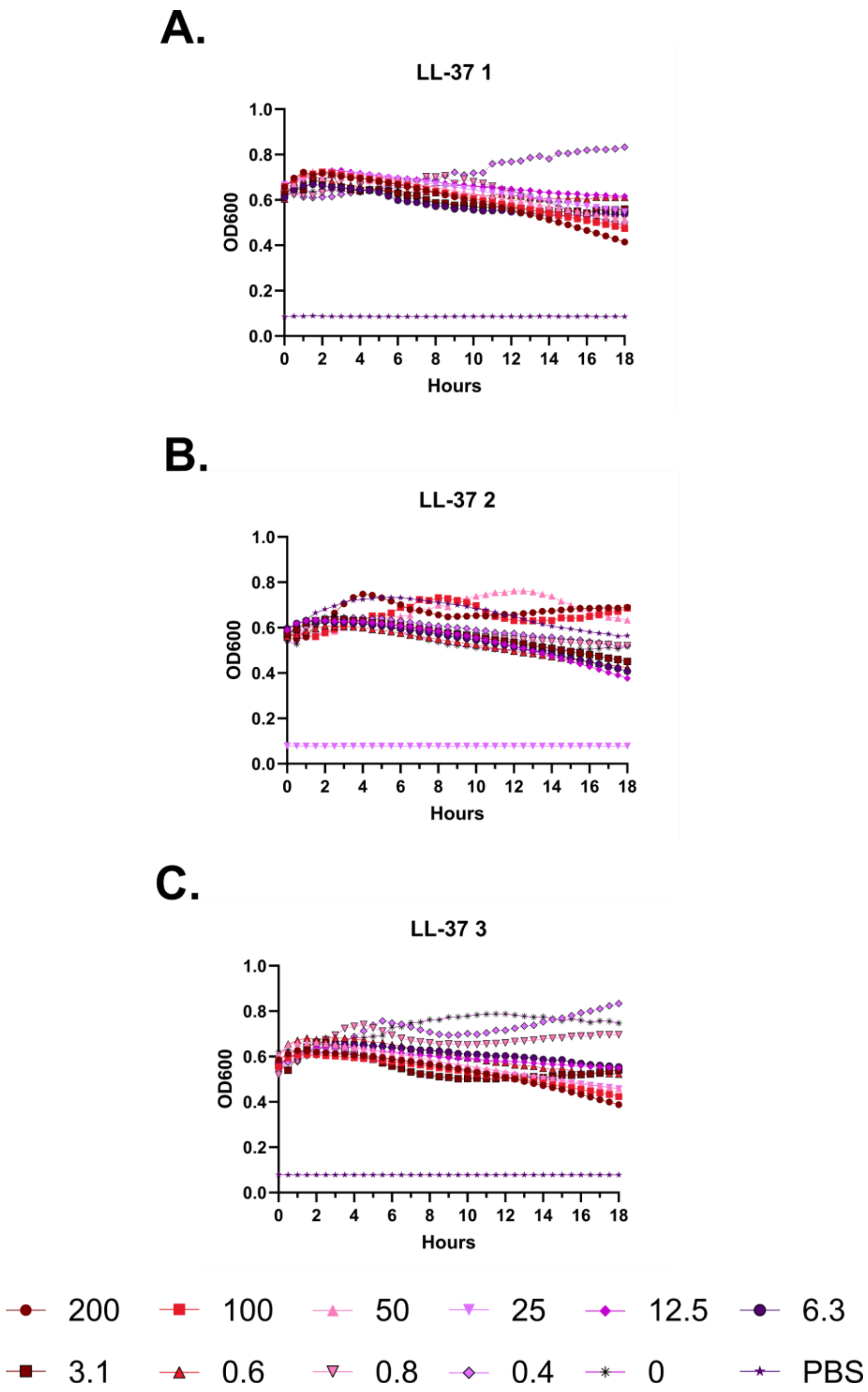
A.
LL-37 1
1.0
0.8
0.6
0.4
0.2
0.0
OD600
0
2
4
6
8
10
12
14
16
18
Hours
B.
LL-37 2
OD600
Hours
C.
LL-37 3
OD600
Hours
200
100
50
25
12.5
6.3
3.1
0.6
0.8
0.4
0
PBS

**Figure S145 – Hemolysis over time with LL-37. A-C.** Each of three independent experiments showing trends in OD600 over time on incubation of defibrinated sheep blood with the indicated concentrations of peptide in μM at 37 °C. Decreasing OD600 indicates increasing hemolysis.

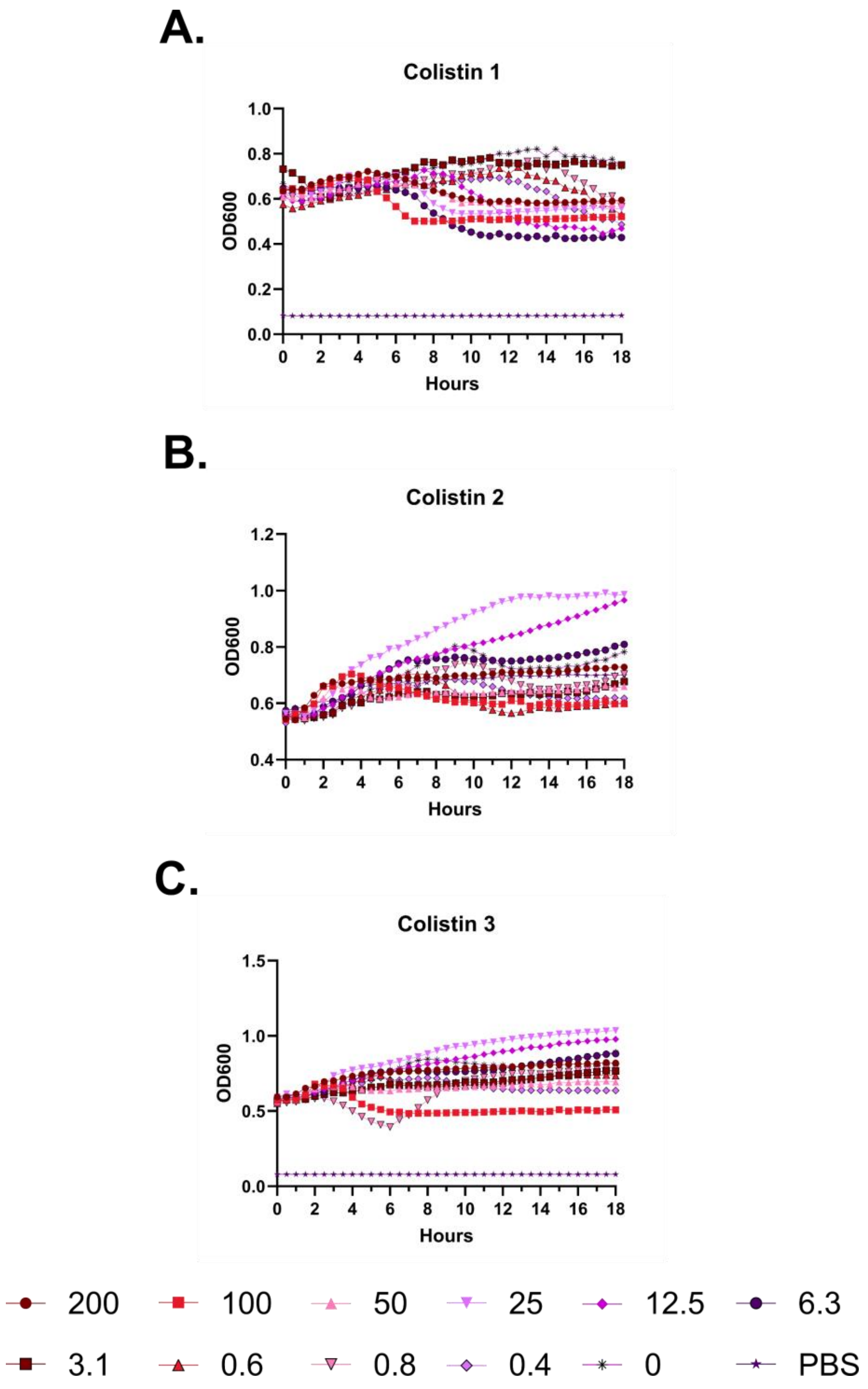
A.
Colistin 1
OD600
Hours
B.
Colistin 2
OD600
Hours
C.
Colistin 3
OD600
Hours
200
100
50
25
12.5
6.3
3.1
0.6
0.8
0.4
0
PBS

**Figure S146 – Hemolysis over time with colistin. A-C.** Each of three independent experiments showing trends in OD600 over time on incubation of defibrinated sheep blood with the indicated concentrations of peptide in μM at 37 °C. Decreasing OD600 indicates increasing hemolysis.

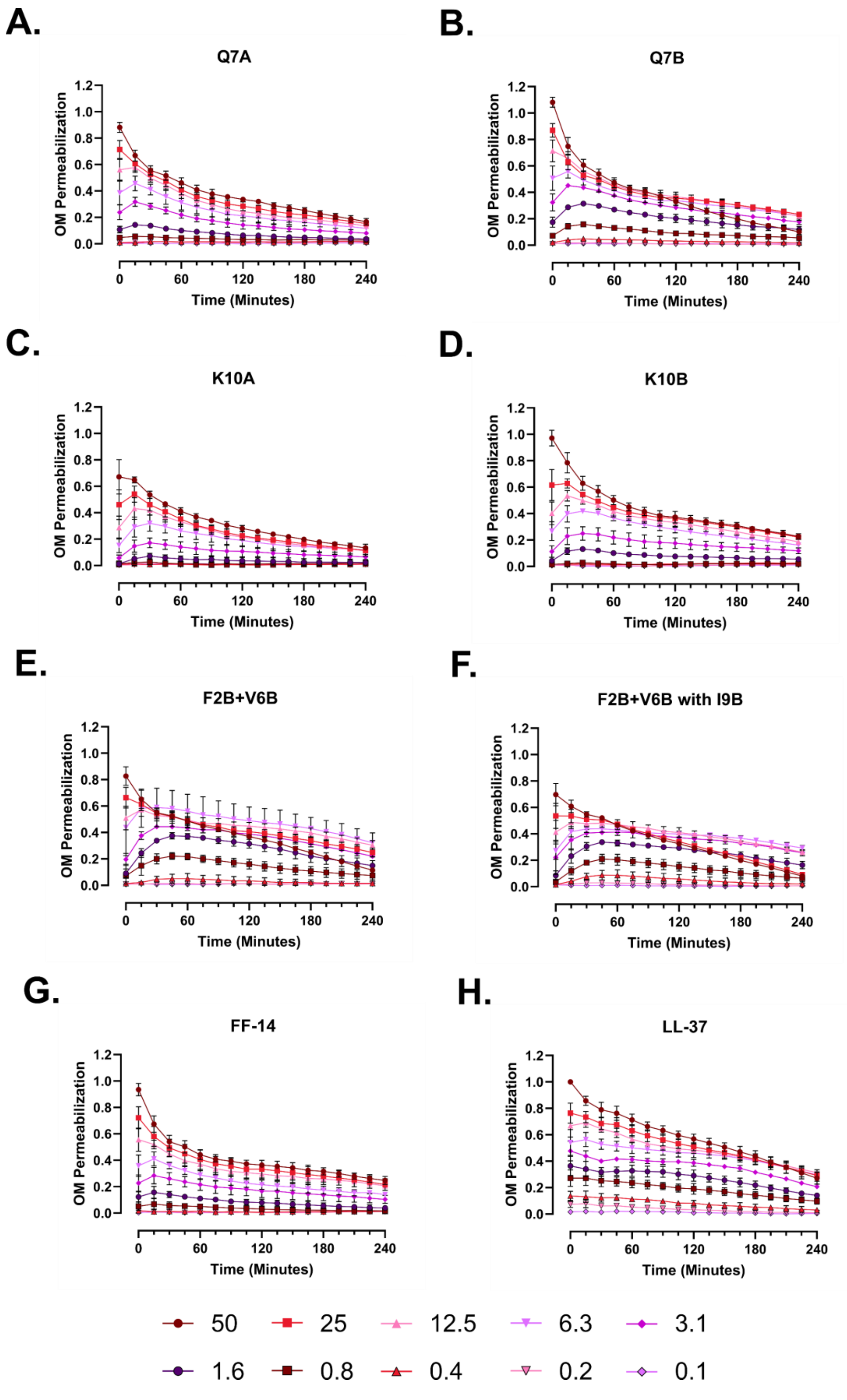

A.
Q7A
B.
Q7B
C.
K10A
D.
K10B
E.
F2B+V6B
F.
F2B+V6B with I9B
G.
FF-14
H.
LL-37
OM Permeabilization
Time (Minutes)
50
25
12.5
6.3
3.1
1.6
0.8
0.4
0.2
0.1

**Figure S147 – *E. coli* outer membrane permeabilization kinetic curves.** Dual membrane permeabilization assays were carried out with NPN and PI stains for the outer and inner membranes, respectively, of *E. coli* (ATCC 25922) in M9+glucose with divalent cation concentrations adjusted to approximate those of unadjusted MHB. **A-H.** Kinetic curves associated with monitoring of outer membrane permeabilization for the indicated peptides over the listed concentration range. Measures represent a fold change over no peptide controls relative to that associated with LL-37, which was set to 100%.

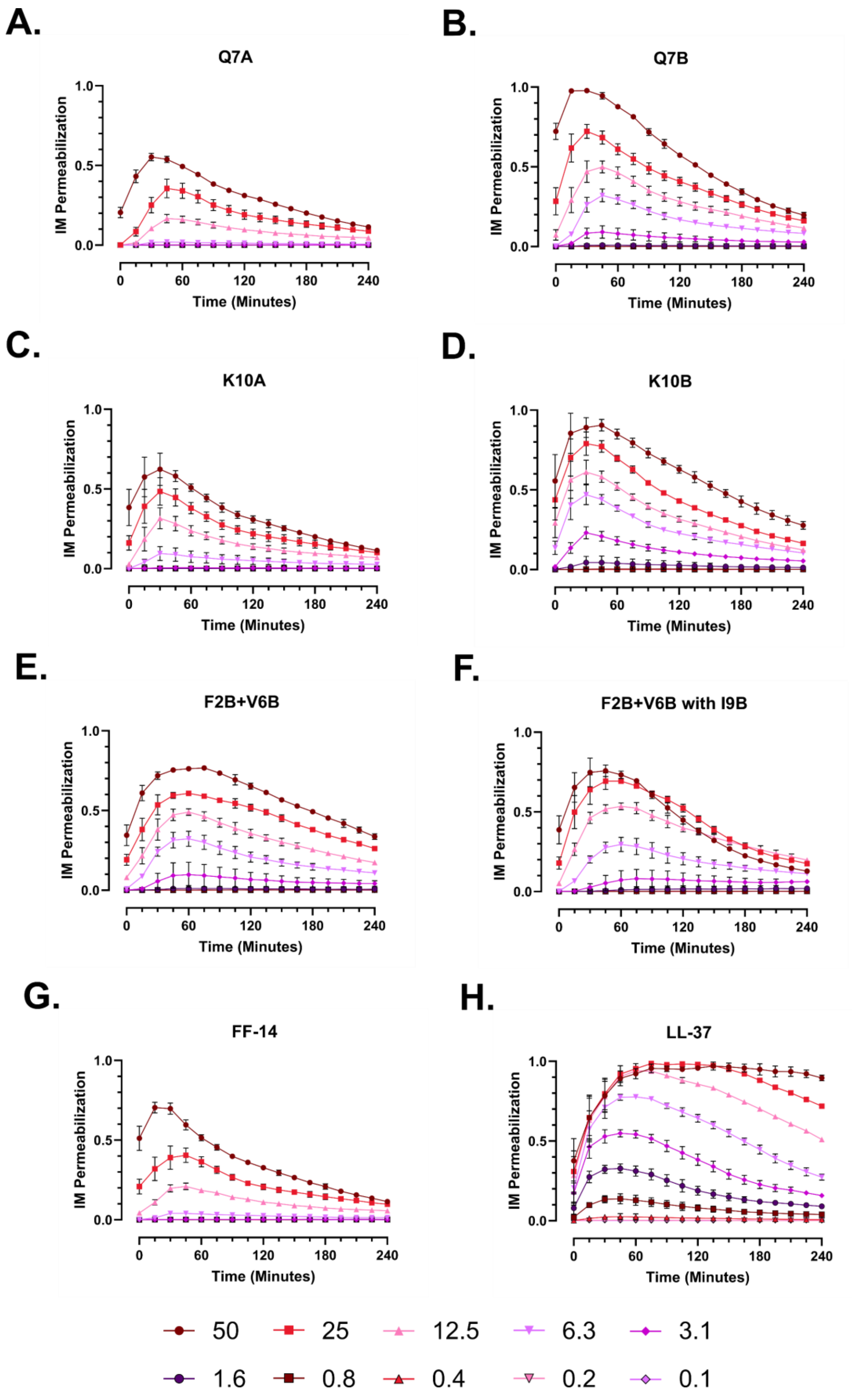

A.
Q7A
B.
Q7B
C.
K10A
D.
K10B
E.
F2B+V6B
F.
F2B+V6B with I9B
G.
FF-14
H.
LL-37
IM Permeabilization
Time (Minutes)
0.0
0.5
1.0
0
60
120
180
240
50
25
12.5
6.3
3.1
1.6
0.8
0.4
0.2
0.1

**Figure S150 – *P. aeruginosa* inner membrane permeabilization kinetic curves.** Dual membrane permeabilization assays were carried out with NPN and PI stains for the outer and inner membranes, respectively, of *P. aeruginosa* (ATCC 27853) in M9+glucose with divalent cation concentrations adjusted to approximate those of unadjusted MHB. **A-H.** Kinetic curves associated with monitoring of inner membrane permeabilization for the indicated peptides over the listed concentration range. Measures represent a fold change over no peptide controls relative to that associated with LL-37, which was set to 100%.

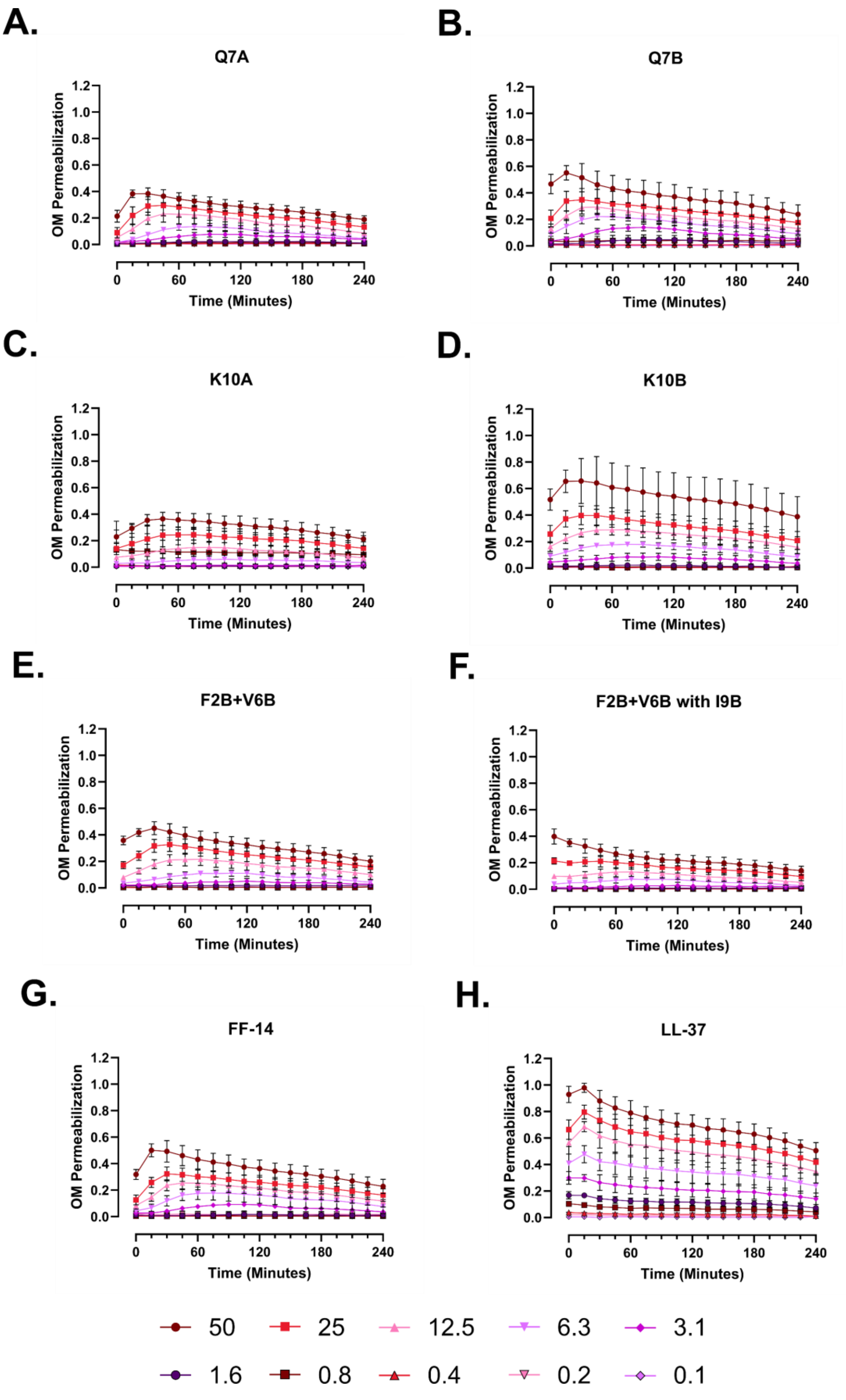
A.
Q7A
B.
Q7B
C.
K10A
D.
K10B
E.
F2B+V6B
F.
F2B+V6B with I9B
G.
FF-14
H.
LL-37
OM Permeabilization
Time (Minutes)
50
25
12.5
6.3
3.1
1.6
0.8
0.4
0.2
0.1

**Figure S149 – *P. aeruginosa* outer membrane permeabilization kinetic curves.** Dual membrane permeabilization assays were carried out with NPN and PI stains for the outer and inner membranes, respectively, of *P. aeruginosa* (ATCC 27853) in M9+glucose with divalent cation concentrations adjusted to approximate those of unadjusted MHB. **A-H.** Kinetic curves associated with monitoring of outer membrane permeabilization for the indicated peptides over the listed concentration range. Measures represent a fold change over no peptide controls relative to that associated with LL-37, which was set to 100%.

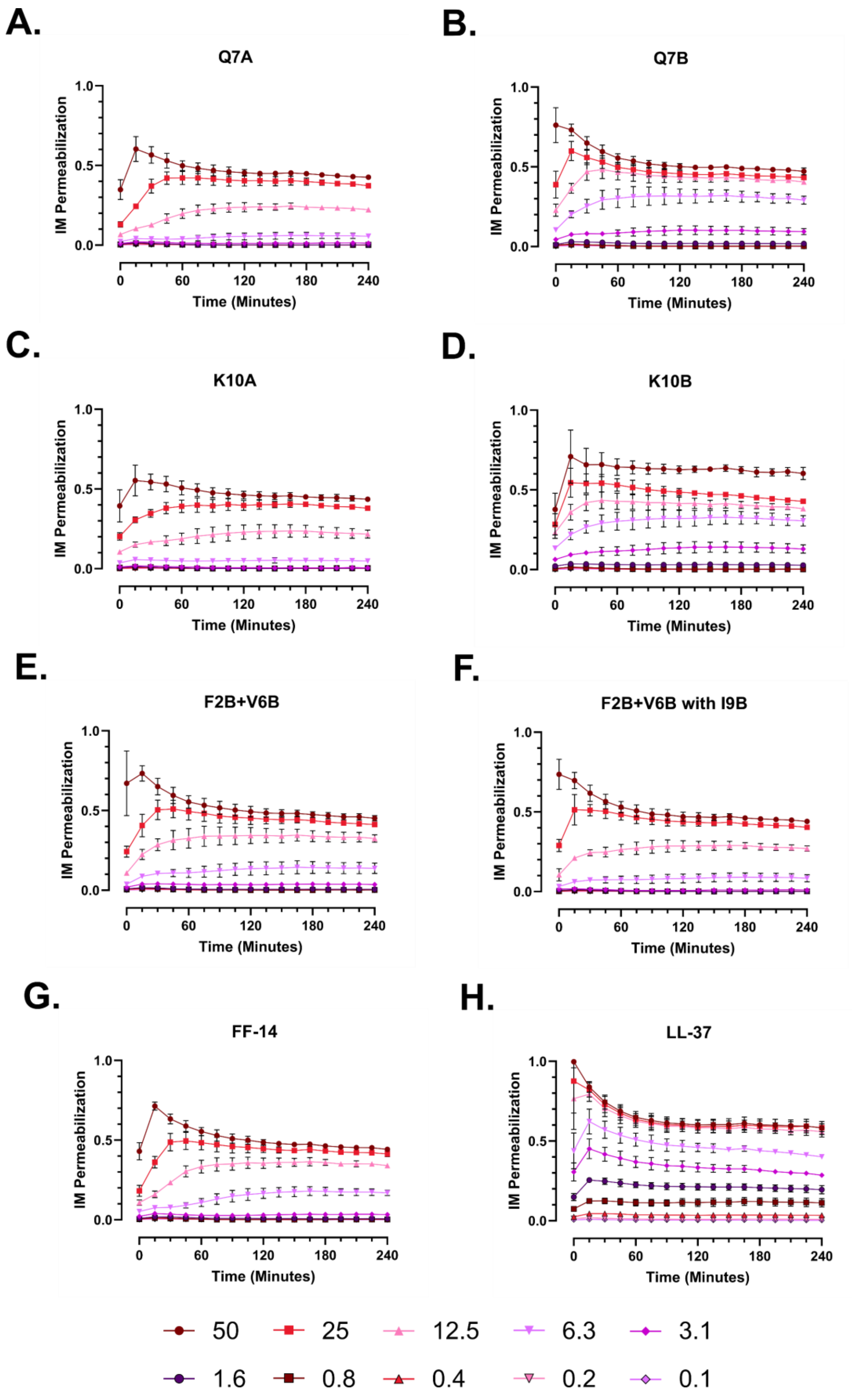

A.
Q7A
B.
Q7B
C.
K10A
D.
K10B
E.
F2B+V6B
F.
F2B+V6B with I9B
G.
FF-14
H.
LL-37
IM Permeabilization
Time (Minutes)
1.0
0.5
0.0
0
60
120
180
240
50
25
12.5
6.3
3.1
1.6
0.8
0.4
0.2
0.1

**Figure S150 – *P. aeruginosa* inner membrane permeabilization kinetic curves.** Dual membrane permeabilization assays were carried out with NPN and PI stains for the outer and inner membranes, respectively, of *P. aeruginosa* (ATCC 27853) in M9+glucose with divalent cation concentrations adjusted to approximate those of unadjusted MHB. **A-H.** Kinetic curves associated with monitoring of inner membrane permeabilization for the indicated peptides over the listed concentration range. Measures represent a fold change over no peptide controls relative to that associated with LL-37, which was set to 100%.

A.

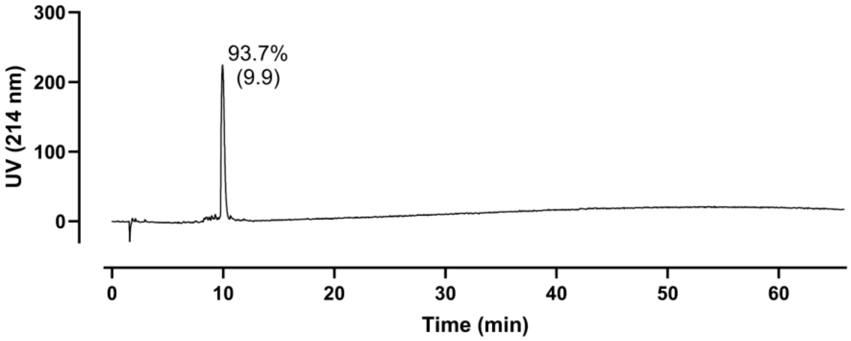

93.7%
(9.9)
UV (214 nm)
Time (min)


B.

C.

D.

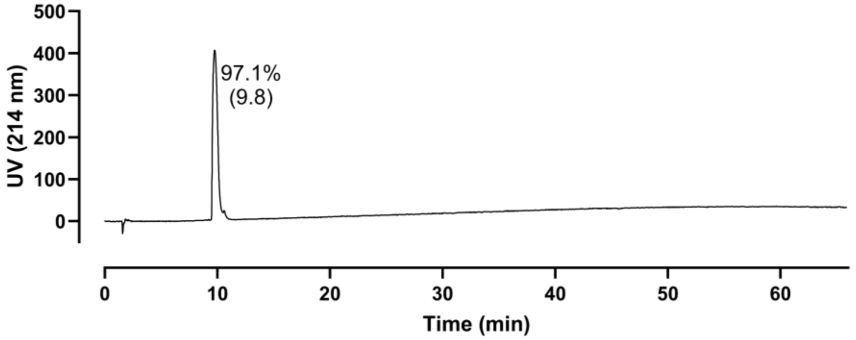

97.1%
(9.8)
UV (214 nm)
Time (min)


E.

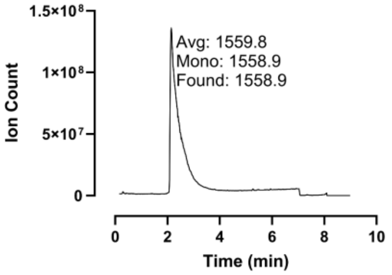

Avg: 1559.8
Mono: 1558.9
Found: 1558.9
Ion Count
Time (min)


F.

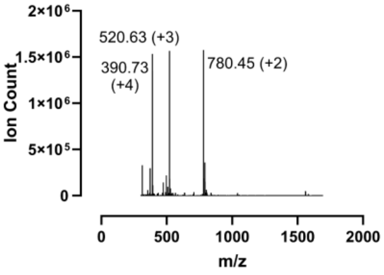

520.63 (+3)
390.73
(+4)
780.45 (+2)
Ion Count
m/z

**Figure S151 – Analytical Data for FF-14 m4. A.** Crude peptide HPLC trace at 214 nm with percent purity reflecting integration of the dominant peak area; retention time is shown in parentheses. **B.** Crude peptide LCMS total ion chromatogram (TIC); not completed. **C.** Mass spectrum at the peak of **B**; not completed. **D.** HPLC of purified peptide. **E.** TIC of purified peptide with the observed mass as well as the calculated average and monoisotopic masses. **F.** Mass spectrum at the peak of **E** with labeling of *m/z* and charge state for selected peaks. Peptides were used at the indicated purity following a single round of purification.

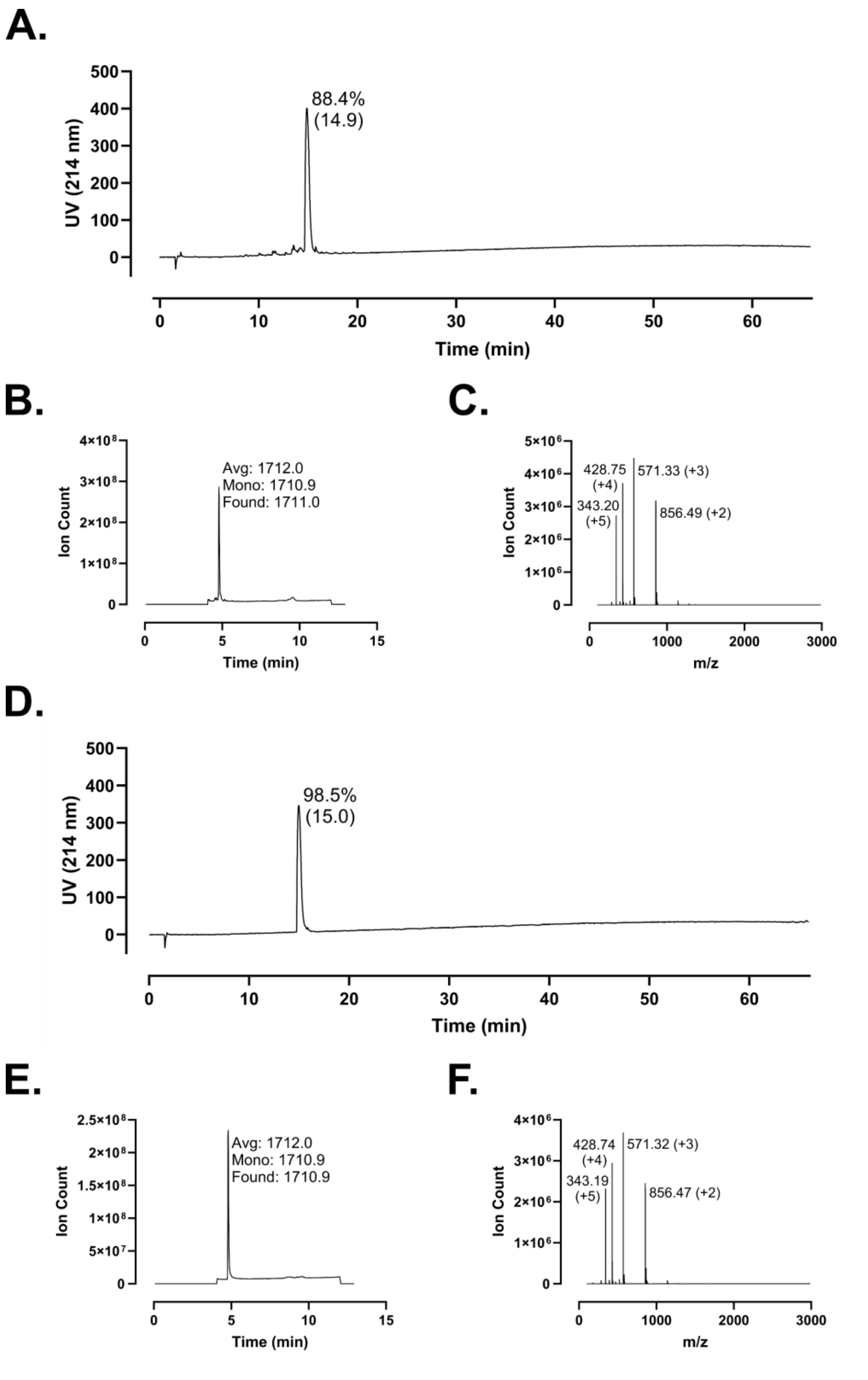
A.
UV (214 nm)
500
400
300
200
100
0
88.4%
(14.9)
0
10
20
30
40
50
60
Time (min)
B.
Ion Count
4×10⁸
3×10⁸
2×10⁸
1×10⁸
0
Avg: 1712.0
Mono: 1710.9
Found: 1711.0
0
5
10
15
Time (min)
C.
Ion Count
5×10⁶
4×10⁶
3×10⁶
2×10⁶
1×10⁶
0
428.75
(+4)
571.33 (+3)
343.20
(+5)
856.49 (+2)
0
1000
2000
3000
m/z
D.
UV (214 nm)
500
400
300
200
100
0
98.5%
(15.0)
0
10
20
30
40
50
60
Time (min)
E.
Ion Count
2.5×10⁸
2×10⁸
1.5×10⁸
1×10⁸
5×10⁷
0
Avg: 1712.0
Mono: 1710.9
Found: 1710.9
0
5
10
15
Time (min)
F.
Ion Count
4×10⁶
3×10⁶
2×10⁶
1×10⁶
0
428.74
(+4)
571.32 (+3)
343.19
(+5)
856.47 (+2)
0
1000
2000
3000
m/z

**Figure S152 – Analytical Data for FF-14 m4-FF. A.** Crude peptide HPLC trace at 214 nm with percent purity reflecting integration of the dominant peak area; retention time is shown in parentheses. **B.** Crude peptide LCMS total ion chromatogram (TIC); observed mass is provided along with the calculated average and monoisotopic masses. **C.** Mass spectrum at the peak of **B**. Peaks are labeled with *m/z* and charge state. **D.** HPLC of purified peptide. **E.** TIC of purified peptide. **F.** Mass spectrum at the peak of **E**. Peptides were used at the indicated purity following a single round of purification.

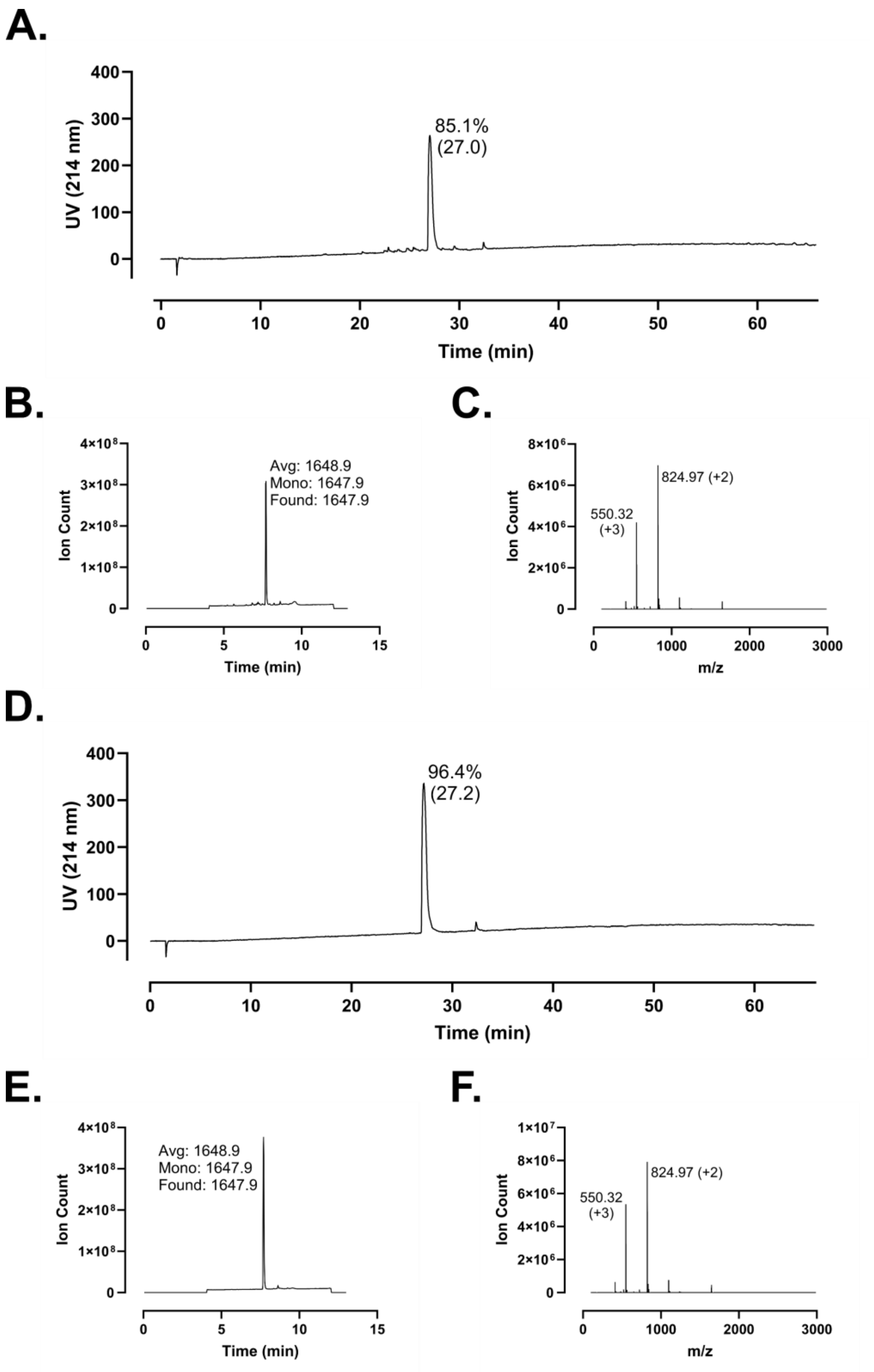

A.
400
300
200
100
0
UV (214 nm)
85.1%
(27.0)
0
10
20
30
40
50
60
Time (min)
B.
4×10⁸
3×10⁸
2×10⁸
1×10⁸
0
Ion Count
Avg: 1648.9
Mono: 1647.9
Found: 1647.9
0
5
10
15
Time (min)
C.
8×10⁶
6×10⁶
4×10⁶
2×10⁶
0
Ion Count
550.32
(+3)
824.97 (+2)
0
1000
2000
3000
m/z
D.
400
300
200
100
0
UV (214 nm)
96.4%
(27.2)
0
10
20
30
40
50
60
Time (min)
E.
4×10⁸
3×10⁸
2×10⁸
1×10⁸
0
Ion Count
Avg: 1648.9
Mono: 1647.9
Found: 1647.9
0
5
10
15
Time (min)
F.
1×10⁷
8×10⁶
6×10⁶
4×10⁶
2×10⁶
0
Ion Count
550.32
(+3)
824.97 (+2)
0
1000
2000
3000
m/z

**Figure S153 – Analytical Data for FF-14 m5. A.** Crude peptide HPLC trace at 214 nm with percent purity reflecting integration of the dominant peak area; retention time is shown in parentheses. **B.** Crude peptide LCMS total ion chromatogram (TIC); observed mass is provided along with the calculated average and monoisotopic masses. **C.** Mass spectrum at the peak of **B**. Peaks are labeled with *m/z* and charge state. **D.** HPLC of purified peptide. **E.** TIC of purified peptide. **F.** Mass spectrum at the peak of **E**. Peptides were used at the indicated purity following a single round of purification.

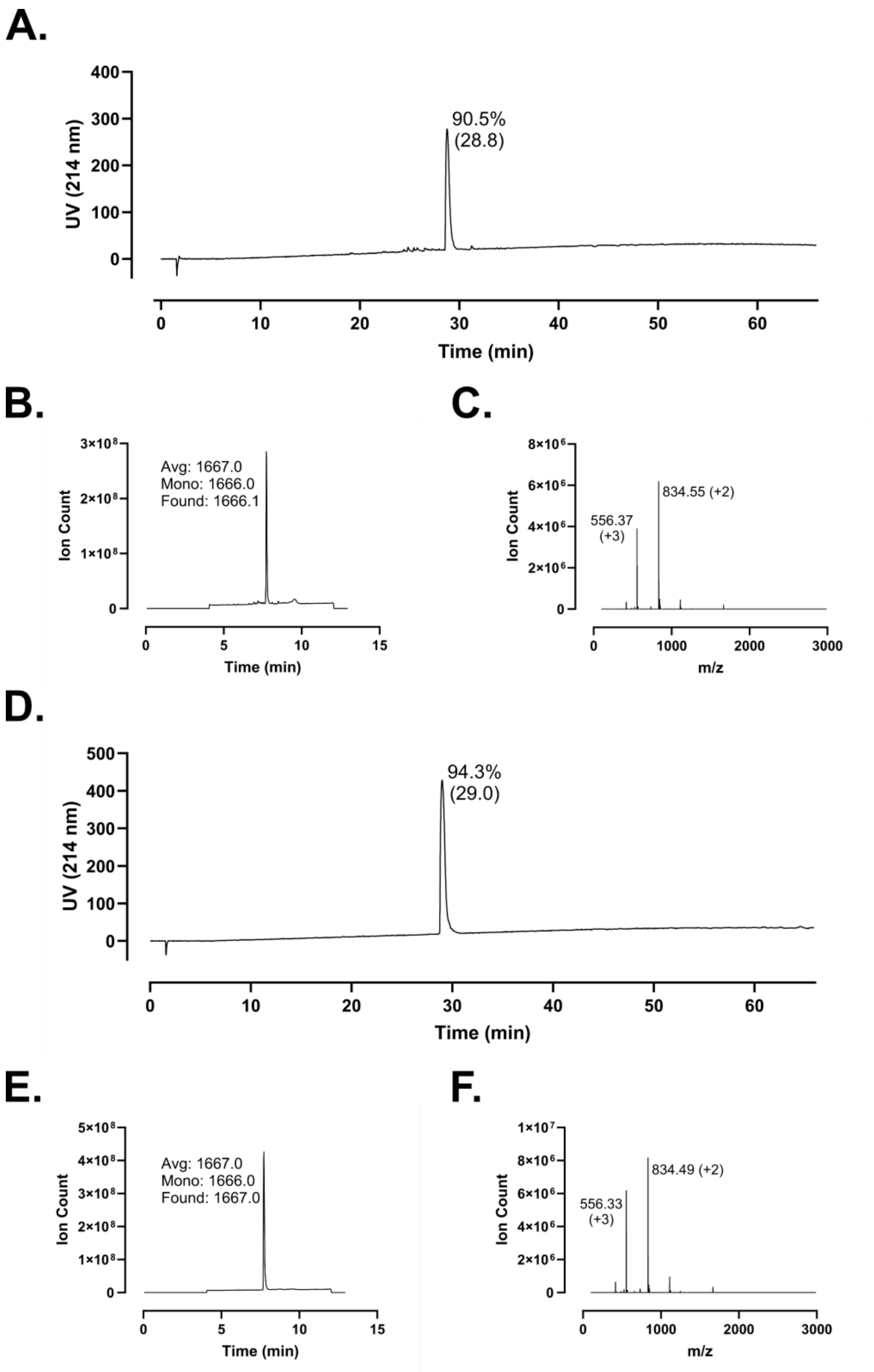
A.
UV (214 nm)
Time (min)
90.5%
(28.8)
B.
Ion Count
Time (min)
Avg: 1667.0
Mono: 1666.0
Found: 1666.1
C.
Ion Count
m/z
556.37
(+3)
834.55 (+2)
D.
UV (214 nm)
Time (min)
94.3%
(29.0)
E.
Ion Count
Time (min)
Avg: 1667.0
Mono: 1666.0
Found: 1667.0
F.
Ion Count
m/z
556.33
(+3)
834.49 (+2)

**Figure S154 – Analytical Data for FF-14 m5-FF. A.** Crude peptide HPLC trace at 214 nm with percent purity reflecting integration of the dominant peak area; retention time is shown in parentheses. **B.** Crude peptide LCMS total ion chromatogram (TIC); observed mass is provided along with the calculated average and monoisotopic masses. **C.** Mass spectrum at the peak of **B**. Peaks are labeled with *m/z* and charge state. **D.** HPLC of purified peptide. **E.** TIC of purified peptide. **F.** Mass spectrum at the peak of **E**. Peptides were used at the indicated purity following a single round of purification.

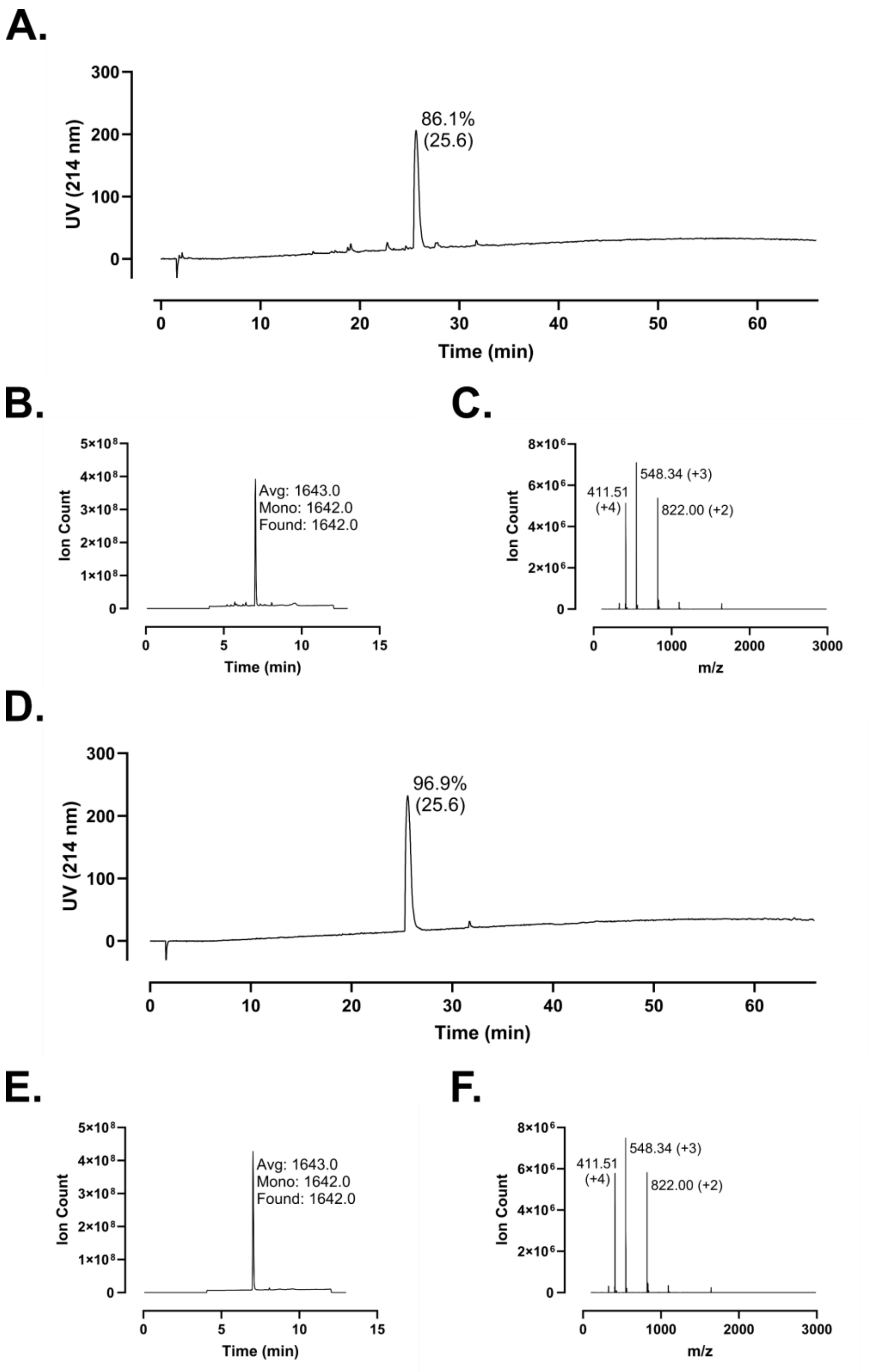
A.
86.1%
(25.6)
UV (214 nm)
Time (min)
B.
Avg: 1643.0
Mono: 1642.0
Found: 1642.0
Ion Count
Time (min)
C.
411.51
(+4)
548.34 (+3)
822.00 (+2)
Ion Count
m/z
D.
96.9%
(25.6)
UV (214 nm)
Time (min)
E.
Avg: 1643.0
Mono: 1642.0
Found: 1642.0
Ion Count
Time (min)
F.
411.51
(+4)
548.34 (+3)
822.00 (+2)
Ion Count
m/z

**Figure S155 – Analytical Data for FF-14 m6. A.** Crude peptide HPLC trace at 214 nm with percent purity reflecting integration of the dominant peak area; retention time is shown in parentheses. **B.** Crude peptide LCMS total ion chromatogram (TIC); observed mass is provided along with the calculated average and monoisotopic masses. **C.** Mass spectrum at the peak of **B**. Peaks are labeled with *m/z* and charge state. **D.** HPLC of purified peptide. **E.** TIC of purified peptide. **F.** Mass spectrum at the peak of **E**. Peptides were used at the indicated purity following a single round of purification.

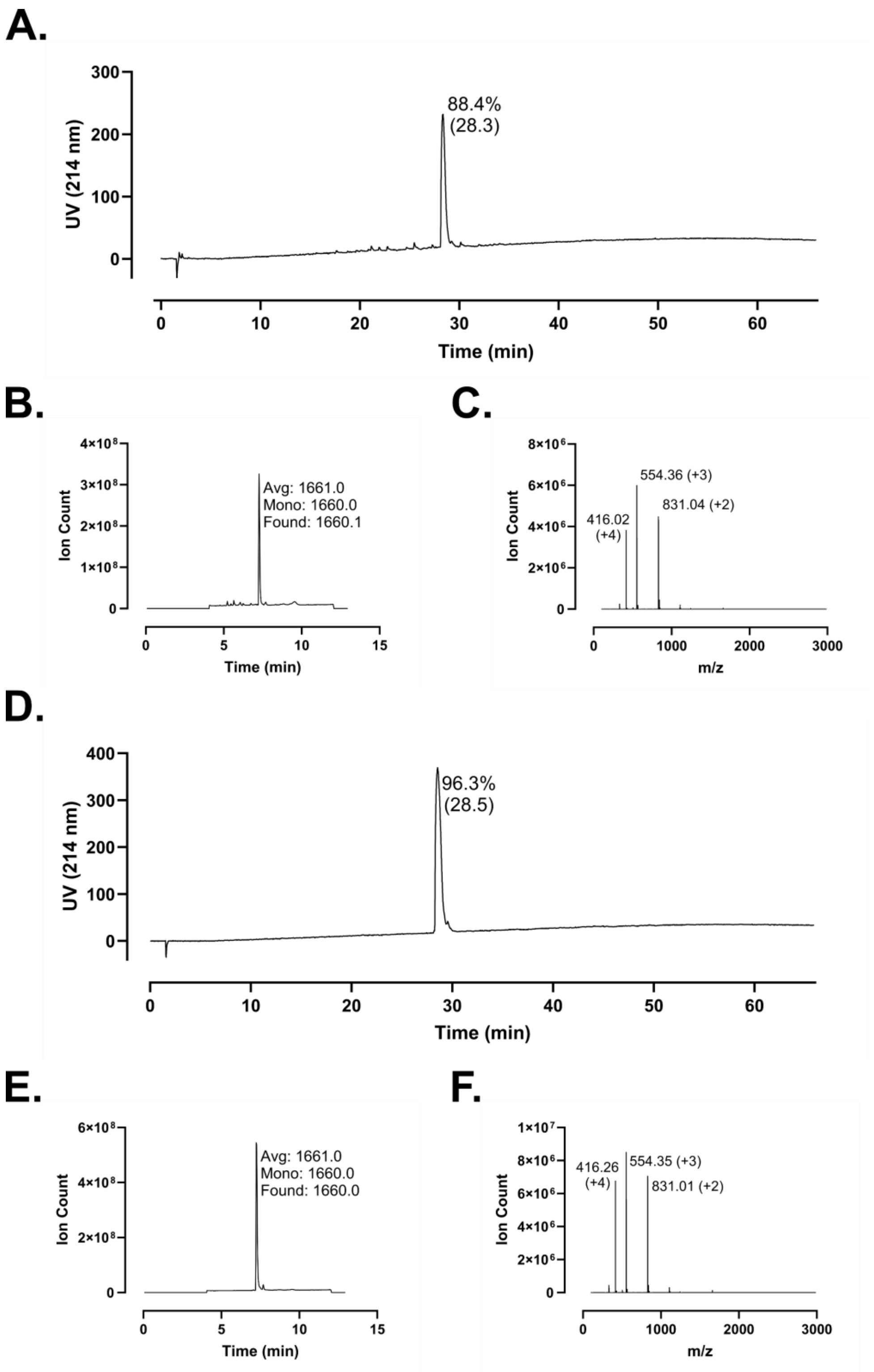
A.
300
200
100
0
UV (214 nm)
88.4%
(28.3)
0
10
20
30
40
50
60
Time (min)
B.
4×10⁸
3×10⁸
2×10⁸
1×10⁸
0
Ion Count
Avg: 1661.0
Mono: 1660.0
Found: 1660.1
0
5
10
15
Time (min)
C.
8×10⁶
6×10⁶
4×10⁶
2×10⁶
0
Ion Count
554.36 (+3)
416.02
(+4)
831.04 (+2)
0
1000
2000
3000
m/z
D.
400
300
200
100
0
UV (214 nm)
96.3%
(28.5)
0
10
20
30
40
50
60
Time (min)
E.
6×10⁸
4×10⁸
2×10⁸
0
Ion Count
Avg: 1661.0
Mono: 1660.0
Found: 1660.0
0
5
10
15
Time (min)
F.
1×10⁷
8×10⁶
6×10⁶
4×10⁶
2×10⁶
0
Ion Count
416.26
(+4)
554.35 (+3)
831.01 (+2)
0
1000
2000
3000
m/z

**Figure S156 – Analytical Data for FF-14 m6-FF. A.** Crude peptide HPLC trace at 214 nm with percent purity reflecting integration of the dominant peak area; retention time is shown in parentheses. **B.** Crude peptide LCMS total ion chromatogram (TIC); observed mass is provided along with the calculated average and monoisotopic masses. **C.** Mass spectrum at the peak of **B**. Peaks are labeled with *m/z* and charge state. **D.** HPLC of purified peptide. **E.** TIC of purified peptide. **F.** Mass spectrum at the peak of **E**. Peptides were used at the indicated purity following a single round of purification.

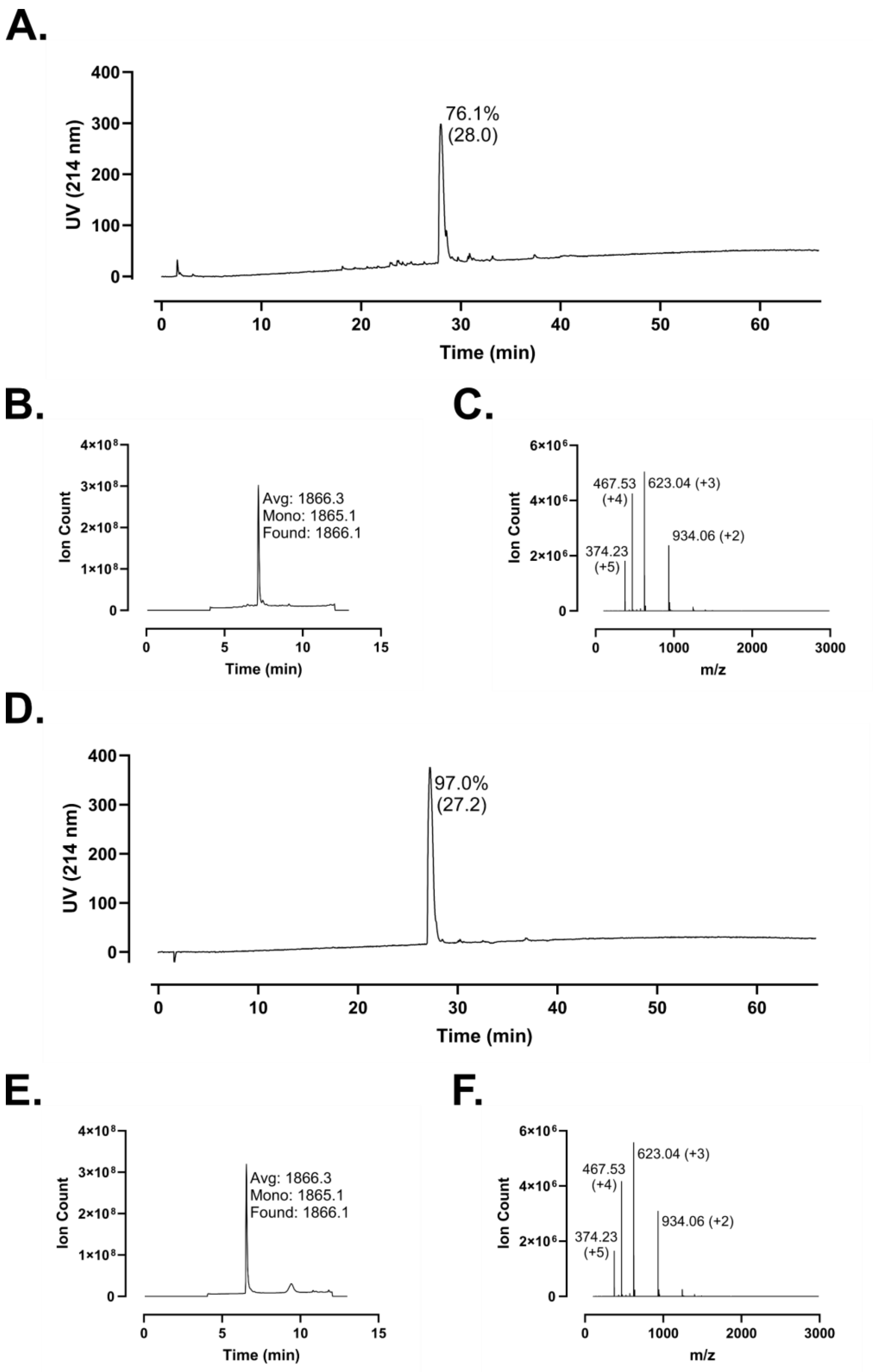
A.
UV (214 nm)
76.1%
(28.0)
Time (min)
B.
Ion Count
Avg: 1866.3
Mono: 1865.1
Found: 1866.1
Time (min)
C.
Ion Count
467.53
(+4)
623.04 (+3)
374.23
(+5)
934.06 (+2)
m/z
D.
UV (214 nm)
97.0%
(27.2)
Time (min)
E.
Ion Count
Avg: 1866.3
Mono: 1865.1
Found: 1866.1
Time (min)
F.
Ion Count
467.53
(+4)
623.04 (+3)
374.23
(+5)
934.06 (+2)
m/z

**Figure S157 – Analytical Data for FF-14. A.** Crude peptide HPLC trace at 214 nm with percent purity reflecting integration of the dominant peak area; retention time is shown in parentheses. **B.** Crude peptide LCMS total ion chromatogram (TIC); observed mass is provided along with the calculated average and monoisotopic masses. **C.** Mass spectrum at the peak of **B**. Peaks are labeled with *m/z* and charge state. **D.** HPLC of purified peptide. **E.** TIC of purified peptide. **F.** Mass spectrum at the peak of **E**. Peptides were used at the indicated purity following a single round of purification. Note: This preparation of FF-14 was used across multiple papers. The analytical characterization shown in this figure is identical to that presented in Reference 34.

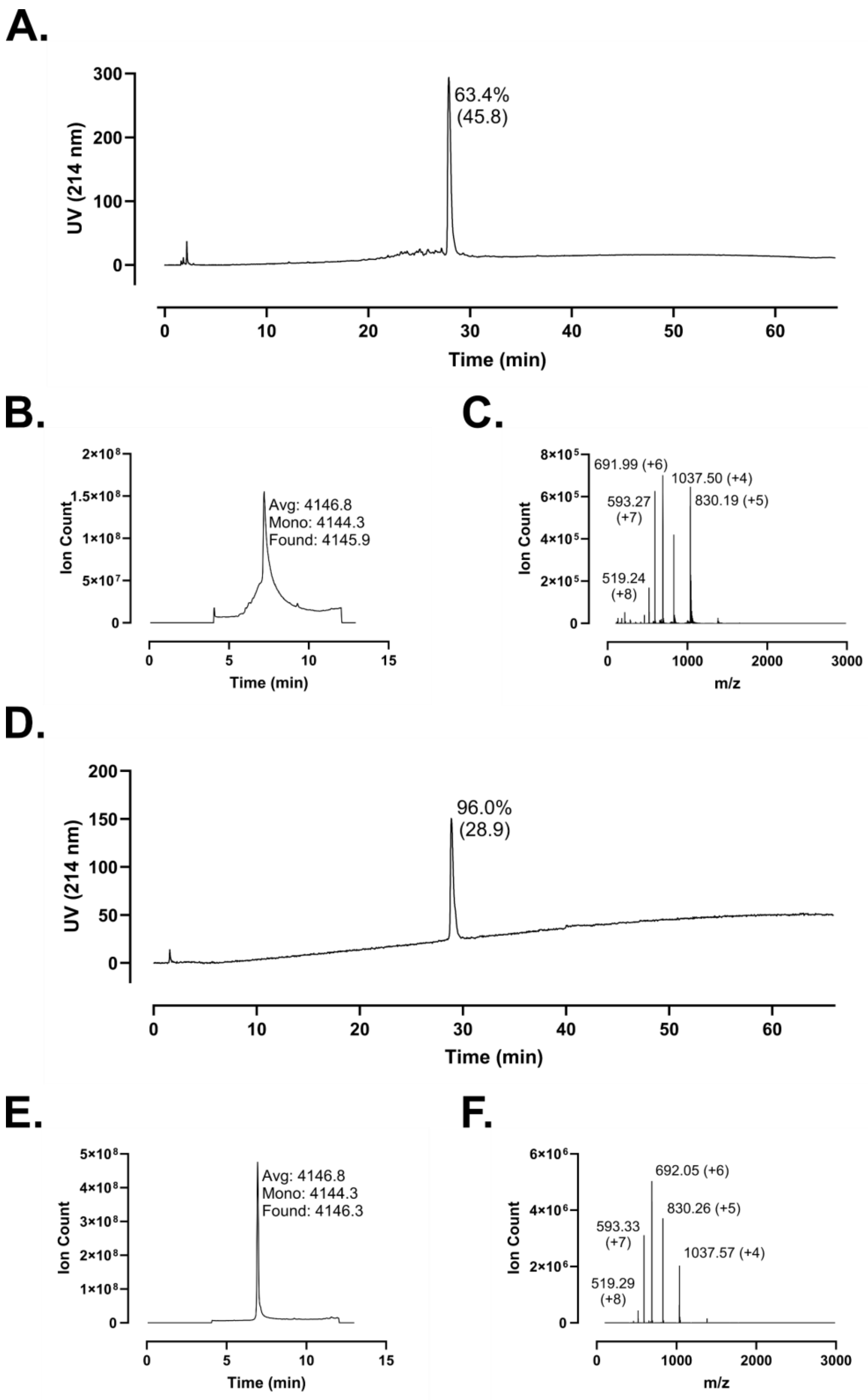
A.
63.4%
(45.8)
UV (214 nm)
Time (min)
B.
Avg: 4146.8
Mono: 4144.3
Found: 4145.9
Ion Count
Time (min)
C.
691.99 (+6)
1037.50 (+4)
593.27
(+7)
830.19 (+5)
519.24
(+8)
Ion Count
m/z
D.
96.0%
(28.9)
UV (214 nm)
Time (min)
E.
Avg: 4146.8
Mono: 4144.3
Found: 4146.3
Ion Count
Time (min)
F.
692.05 (+6)
830.26 (+5)
593.33
(+7)
1037.57 (+4)
519.29
(+8)
Ion Count
m/z

**Figure S158 – Analytical Data for LL-37 m4. A.** Crude peptide HPLC trace at 214 nm with percent purity reflecting integration of the dominant peak area; retention time is shown in parentheses. **B.** Crude peptide LCMS total ion chromatogram (TIC); observed mass is provided along with the calculated average and monoisotopic masses. **C.** Mass spectrum at the peak of **B**. Peaks are labeled with *m/z* and charge state. **D.** HPLC of purified peptide. **E.** TIC of purified peptide. **F.** Mass spectrum at the peak of **E**. Peptides were used at the indicated purity following a single round of purification.

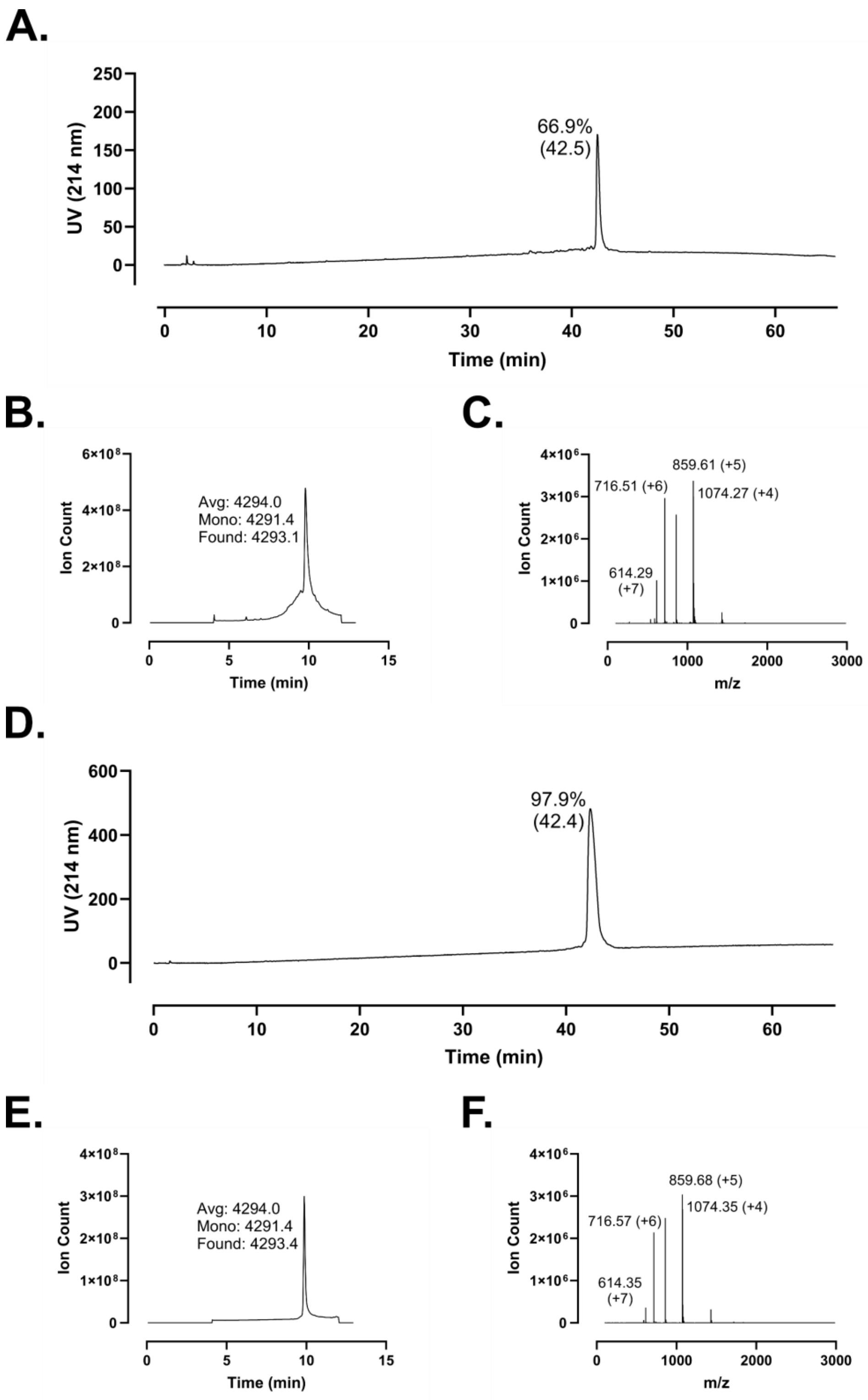
A.
UV (214 nm)
66.9%
(42.5)
Time (min)
B.
Ion Count
Avg: 4294.0
Mono: 4291.4
Found: 4293.1
Time (min)
C.
Ion Count
859.61 (+5)
716.51 (+6)
1074.27 (+4)
614.29
(+7)
m/z
D.
UV (214 nm)
97.9%
(42.4)
Time (min)
E.
Ion Count
Avg: 4294.0
Mono: 4291.4
Found: 4293.4
Time (min)
F.
Ion Count
859.68 (+5)
716.57 (+6)
1074.35 (+4)
614.35
(+7)
m/z

**Figure S159 – Analytical Data for LL-37 m5. A.** Crude peptide HPLC trace at 214 nm with percent purity reflecting integration of the dominant peak area; retention time is shown in parentheses. **B.** Crude peptide LCMS total ion chromatogram (TIC); observed mass is provided along with the calculated average and monoisotopic masses. **C.** Mass spectrum at the peak of **B**. Peaks are labeled with *m/z* and charge state. **D.** HPLC of purified peptide. **E.** TIC of purified peptide. **F.** Mass spectrum at the peak of **E**. Peptides were used at the indicated purity following a single round of purification.

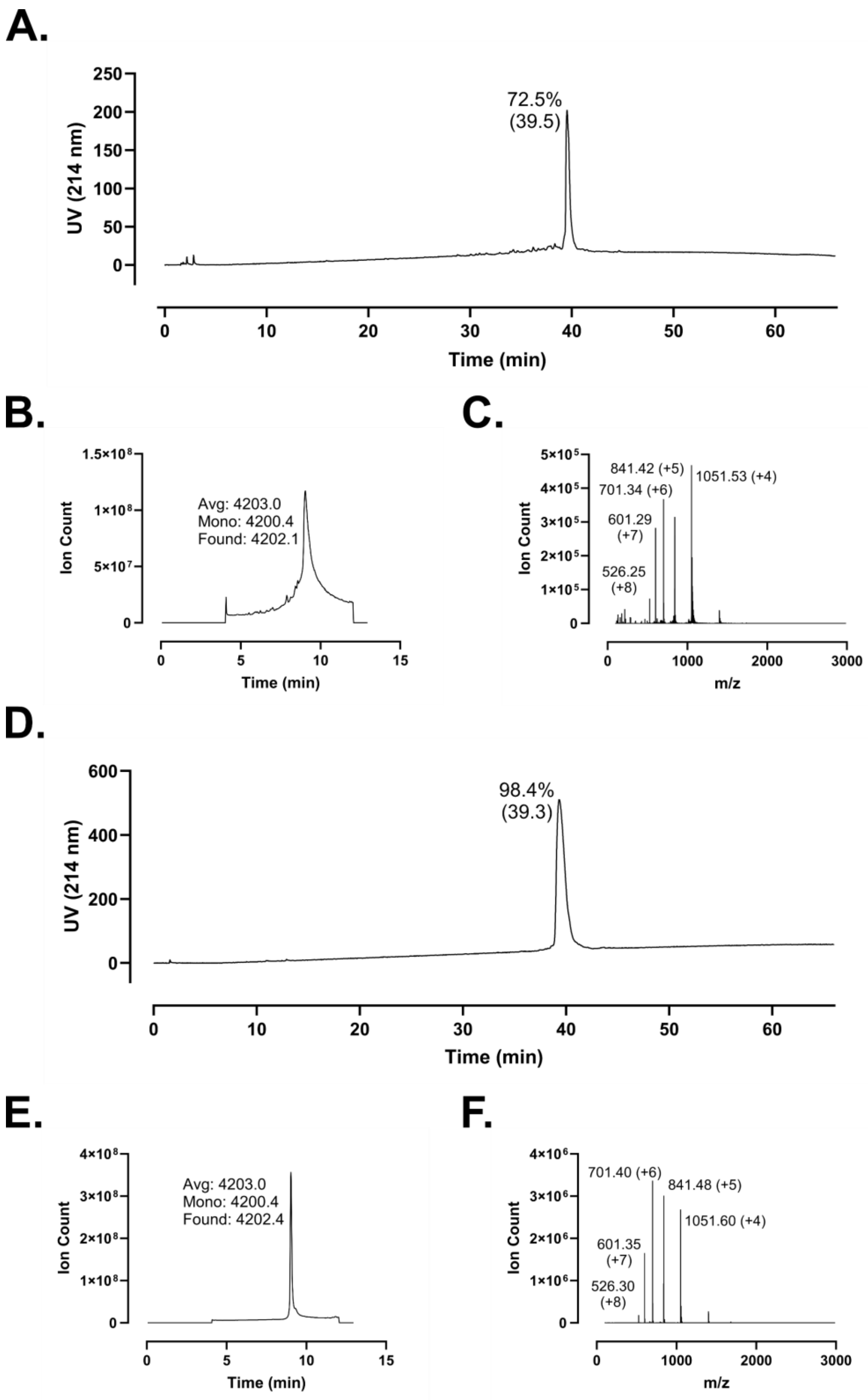
A.
UV (214 nm)
250
200
150
100
50
0
72.5%
(39.5)
0
10
20
30
40
50
60
Time (min)
B.
1.5×10⁸
1×10⁸
5×10⁷
0
Ion Count
Avg: 4203.0
Mono: 4200.4
Found: 4202.1
0
5
10
15
Time (min)
C.
5×10⁵
4×10⁵
3×10⁵
2×10⁵
1×10⁵
0
Ion Count
841.42 (+5)
1051.53 (+4)
701.34 (+6)
601.29
(+7)
526.25
(+8)
0
1000
2000
3000
m/z
D.
UV (214 nm)
600
400
200
0
98.4%
(39.3)
0
10
20
30
40
50
60
Time (min)
E.
4×10⁸
3×10⁸
2×10⁸
1×10⁸
0
Ion Count
Avg: 4203.0
Mono: 4200.4
Found: 4202.4
0
5
10
15
Time (min)
F.
4×10⁶
3×10⁶
2×10⁶
1×10⁶
0
Ion Count
701.40 (+6)
841.48 (+5)
1051.60 (+4)
601.35
(+7)
526.30
(+8)
0
1000
2000
3000
m/z

**Figure S160 – Analytical Data for LL-37 m6. A.** Crude peptide HPLC trace at 214 nm with percent purity reflecting integration of the dominant peak area; retention time is shown in parentheses. **B.** Crude peptide LCMS total ion chromatogram (TIC); observed mass is provided along with the calculated average and monoisotopic masses. **C.** Mass spectrum at the peak of **B**. Peaks are labeled with *m/z* and charge state. **D.** HPLC of purified peptide. **E.** TIC of purified peptide. **F.** Mass spectrum at the peak of **E**. Peptides were used at the indicated purity following a single round of purification.

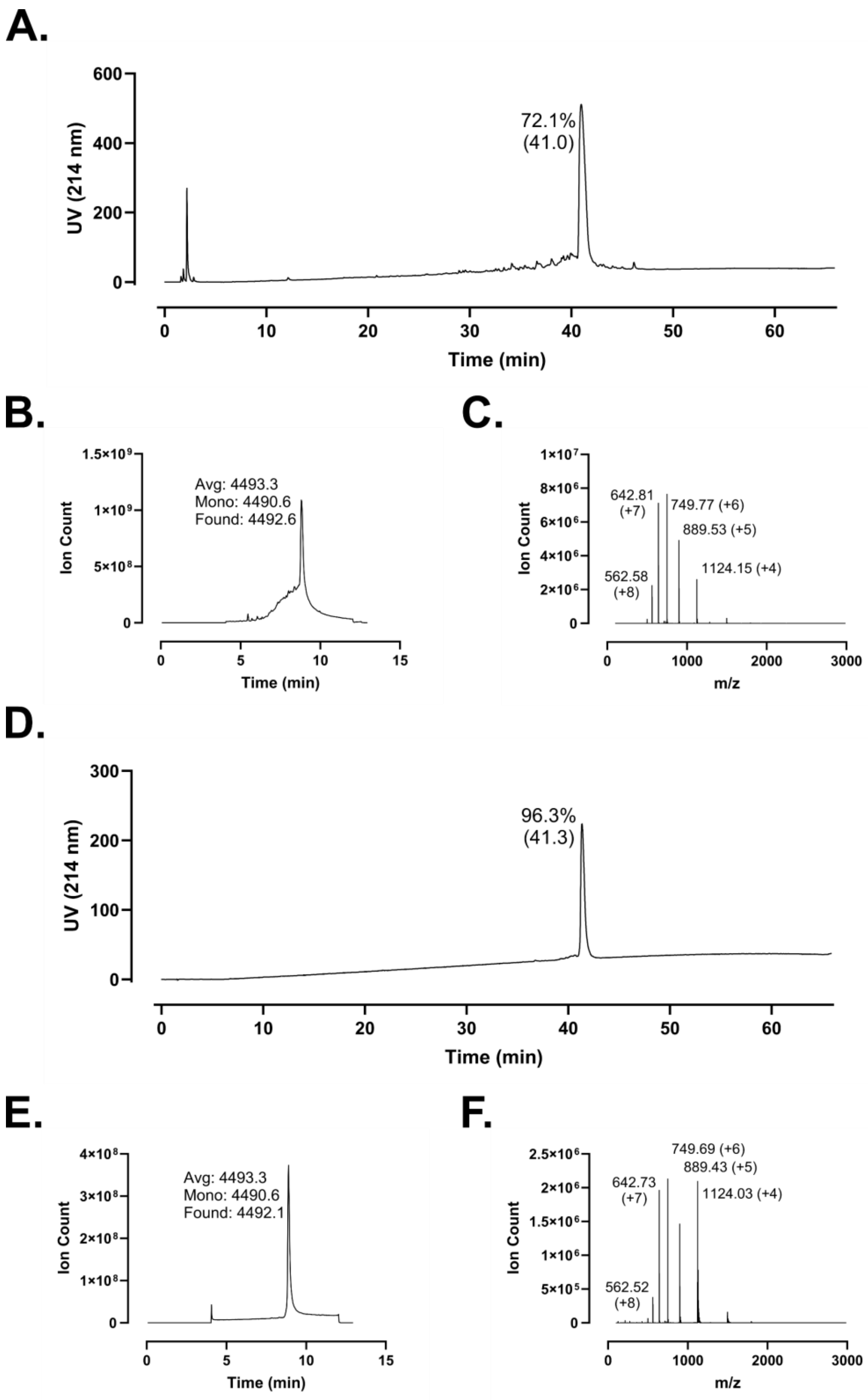
A.
72.1%
(41.0)
UV (214 nm)
Time (min)
B.
Avg: 4493.3
Mono: 4490.6
Found: 4492.6
Ion Count
Time (min)
C.
642.81
(+7)
749.77 (+6)
889.53 (+5)
562.58
(+8)
1124.15 (+4)
Ion Count
m/z
D.
96.3%
(41.3)
UV (214 nm)
Time (min)
E.
Avg: 4493.3
Mono: 4490.6
Found: 4492.1
Ion Count
Time (min)
F.
749.69 (+6)
889.43 (+5)
642.73
(+7)
1124.03 (+4)
562.52
(+8)
Ion Count
m/z

**Figure S161 – Analytical Data for LL-37. A.** Crude peptide HPLC trace at 214 nm with percent purity reflecting integration of the dominant peak area; retention time is shown in parentheses. **B.** Crude peptide LCMS total ion chromatogram (TIC); observed mass is provided along with the calculated average and monoisotopic masses. **C.** Mass spectrum at the peak of **B**. Peaks are labeled with *m/z* and charge state. **D.** HPLC of purified peptide. **E.** TIC of purified peptide. **F.** Mass spectrum at the peak of **E**. Peptides were used at the indicated purity following a single round of purification. Note: This preparation of FF-14 was used across multiple papers. The analytical characterization shown in this figure is identical to that presented in Reference 34.

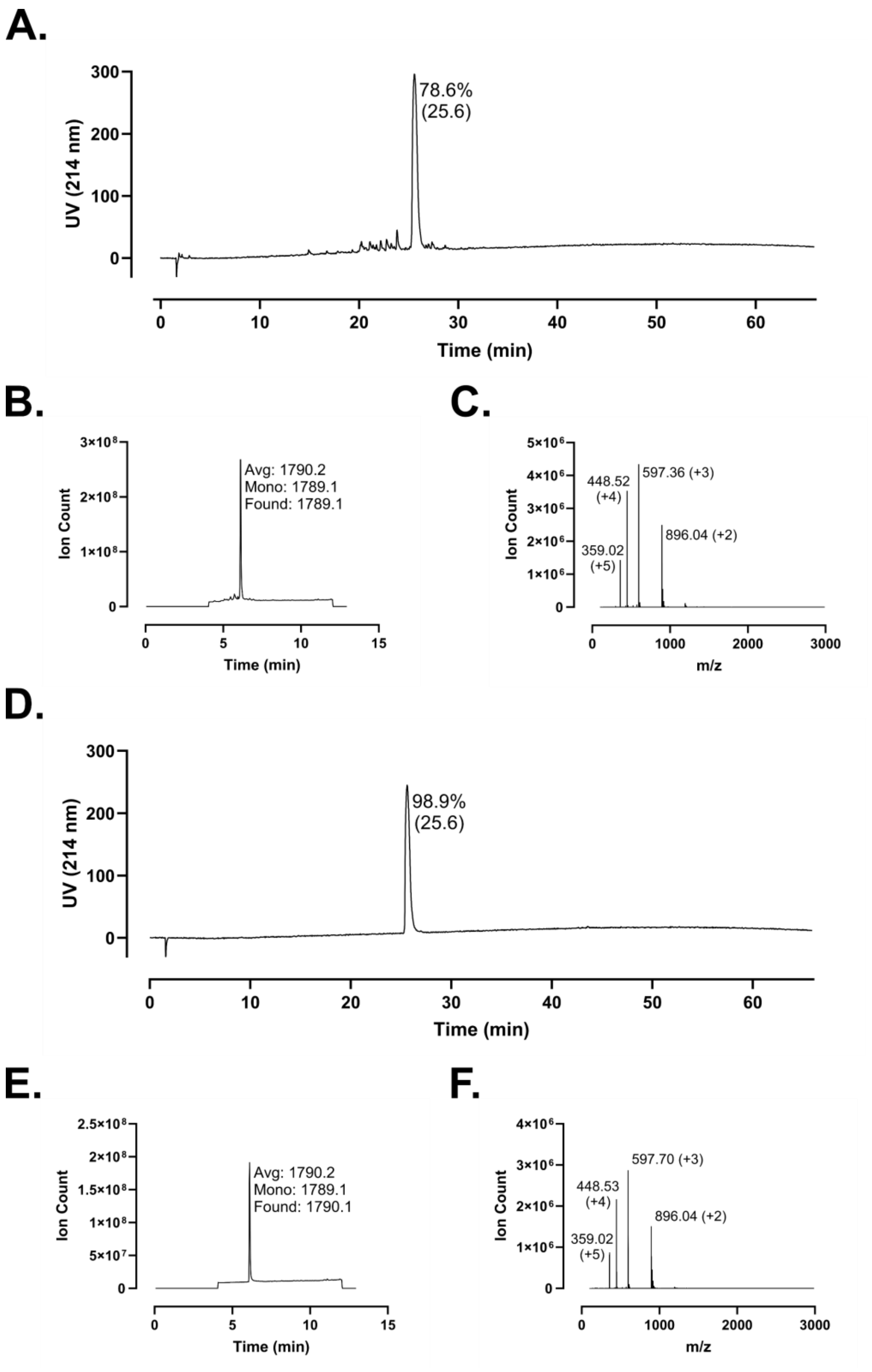
A.
UV (214 nm)
Time (min)
78.6%
(25.6)
B.
Ion Count
Time (min)
Avg: 1790.2
Mono: 1789.1
Found: 1789.1
C.
Ion Count
m/z
448.52
(+4)
597.36 (+3)
359.02
(+5)
896.04 (+2)
D.
UV (214 nm)
Time (min)
98.9%
(25.6)
E.
Ion Count
Time (min)
Avg: 1790.2
Mono: 1789.1
Found: 1790.1
F.
Ion Count
m/z
448.53
(+4)
597.70 (+3)
359.02
(+5)
896.04 (+2)

**Figure S162 – Analytical Data for FF-14 F1A. A.** Crude peptide HPLC trace at 214 nm with percent purity reflecting integration of the dominant peak area; retention time is shown in parentheses. **B.** Crude peptide LCMS total ion chromatogram (TIC); observed mass is provided along with the calculated average and monoisotopic masses. **C.** Mass spectrum at the peak of **B**. Peaks are labeled with *m/z* and charge state. **D.** HPLC of purified peptide. **E.** TIC of purified peptide. **F.** Mass spectrum at the peak of **E**. Peptides were used at the indicated purity following a single round of purification.

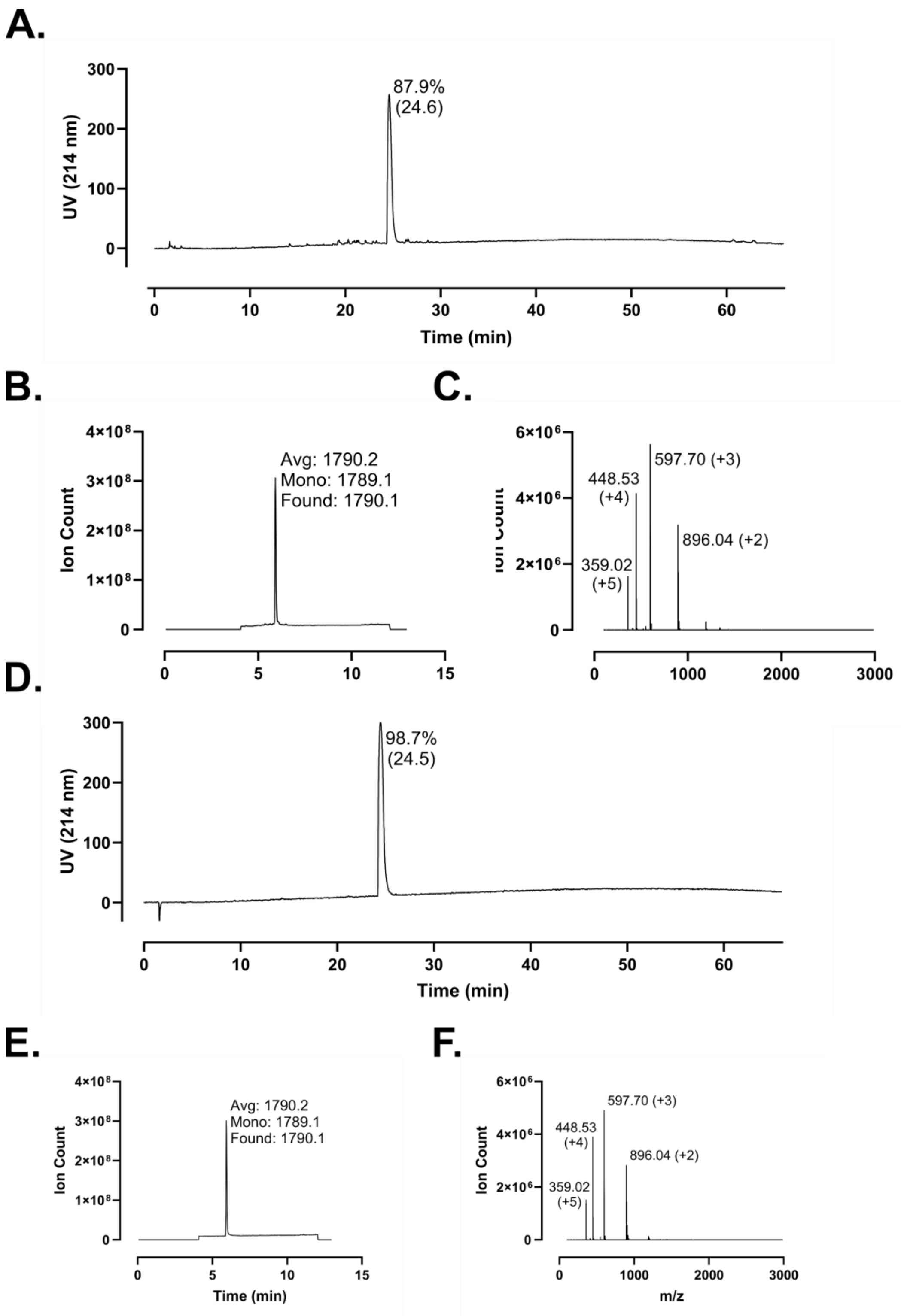
A.
87.9%
(24.6)
UV (214 nm)
Time (min)
B.
Avg: 1790.2
Mono: 1789.1
Found: 1790.1
Ion Count
C.
597.70 (+3)
448.53
(+4)
896.04 (+2)
359.02
(+5)
D.
98.7%
(24.5)
UV (214 nm)
Time (min)
E.
Avg: 1790.2
Mono: 1789.1
Found: 1790.1
Ion Count
Time (min)
F.
597.70 (+3)
448.53
(+4)
896.04 (+2)
359.02
(+5)
Ion Count
m/z

**Figure S163 – Analytical Data for FF-14 F2A. A.** Crude peptide HPLC trace at 214 nm with percent purity reflecting integration of the dominant peak area; retention time is shown in parentheses. **B.** Crude peptide LCMS total ion chromatogram (TIC); observed mass is provided along with the calculated average and monoisotopic masses. **C.** Mass spectrum at the peak of **B**. Peaks are labeled with *m/z* and charge state. **D.** HPLC of purified peptide. **E.** TIC of purified peptide. **F.** Mass spectrum at the peak of **E**. Peptides were used at the indicated purity following a single round of purification.

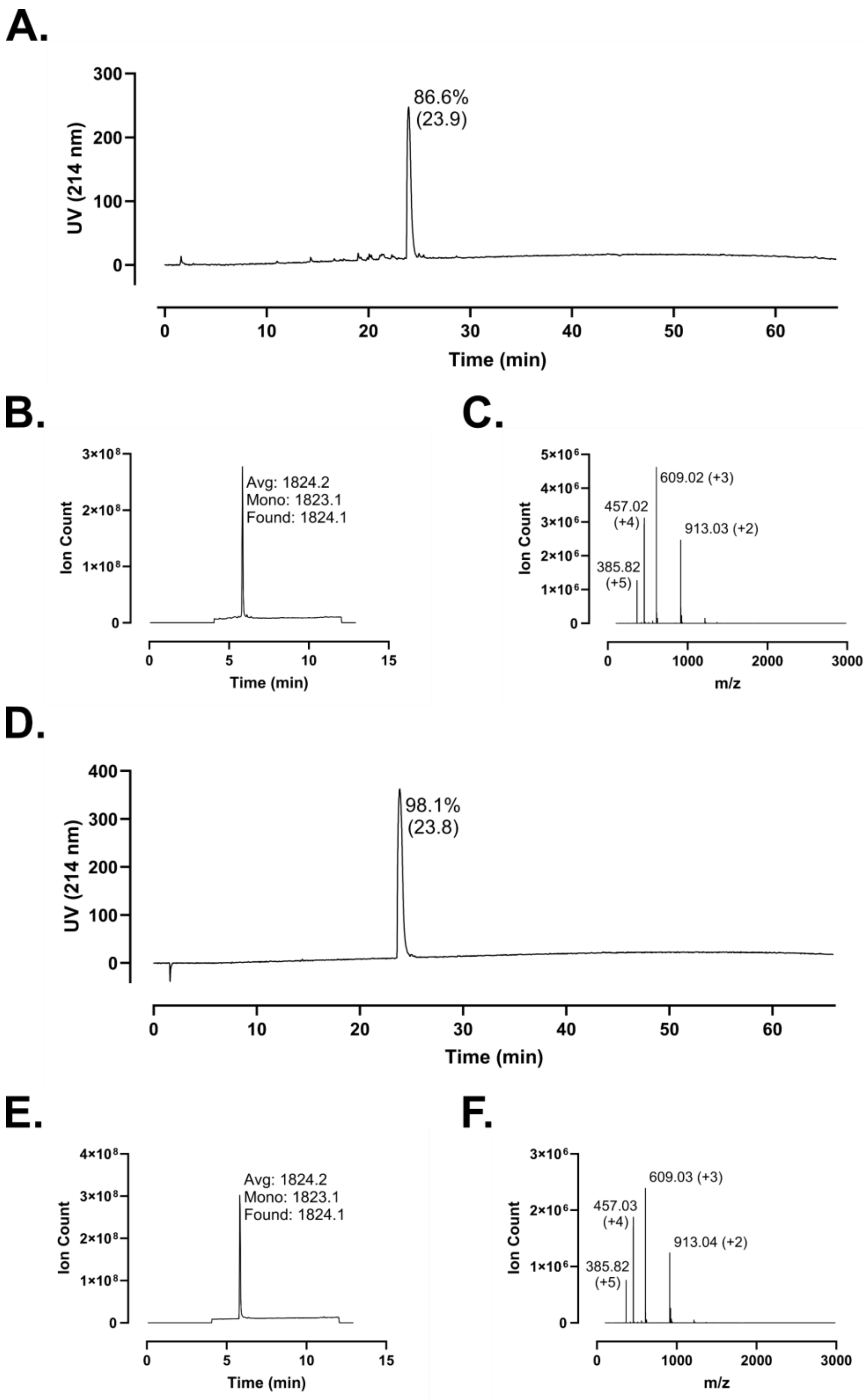

A.
UV (214 nm)
300
200
100
0
86.6%
(23.9)
0
10
20
30
40
50
60
Time (min)
B.
3×10^8
2×10^8
1×10^8
0
Ion Count
Avg: 1824.2
Mono: 1823.1
Found: 1824.1
0
5
10
15
Time (min)
C.
5×10^6
4×10^6
3×10^6
2×10^6
1×10^6
0
Ion Count
609.02 (+3)
457.02
(+4)
913.03 (+2)
385.82
(+5)
0
1000
2000
3000
m/z
D.
UV (214 nm)
400
300
200
100
0
98.1%
(23.8)
0
10
20
30
40
50
60
Time (min)
E.
4×10^8
3×10^8
2×10^8
1×10^8
0
Ion Count
Avg: 1824.2
Mono: 1823.1
Found: 1824.1
0
5
10
15
Time (min)
F.
3×10^6
2×10^6
1×10^6
0
Ion Count
609.03 (+3)
457.03
(+4)
913.04 (+2)
385.82
(+5)
0
1000
2000
3000
m/z

**Figure S164 – Analytical Data for FF-14 I5A. A.** Crude peptide HPLC trace at 214 nm with percent purity reflecting integration of the dominant peak area; retention time is shown in parentheses. **B.** Crude peptide LCMS total ion chromatogram (TIC); observed mass is provided along with the calculated average and monoisotopic masses. **C.** Mass spectrum at the peak of **B**. Peaks are labeled with *m/z* and charge state. **D.** HPLC of purified peptide. **E.** TIC of purified peptide. **F.** Mass spectrum at the peak of **E**. Peptides were used at the indicated purity following a single round of purification.

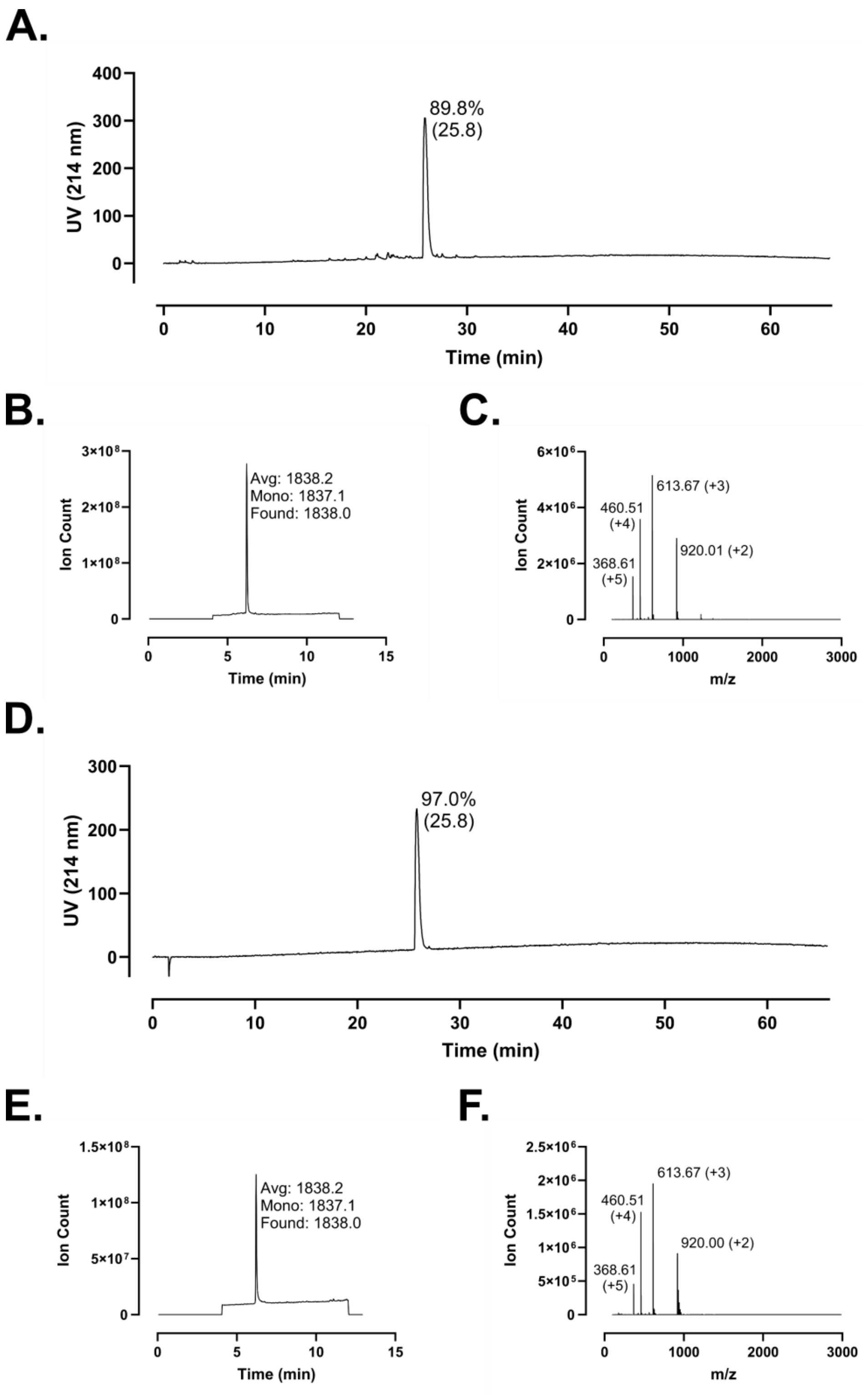
A.
400
300
200
100
0
UV (214 nm)
89.8%
(25.8)
0
10
20
30
40
50
60
Time (min)
B.
3×10⁸
2×10⁸
1×10⁸
0
Ion Count
Avg: 1838.2
Mono: 1837.1
Found: 1838.0
0
5
10
15
Time (min)
C.
6×10⁶
4×10⁶
2×10⁶
0
Ion Count
613.67 (+3)
460.51
(+4)
920.01 (+2)
368.61
(+5)
0
1000
2000
3000
m/z
D.
300
200
100
0
UV (214 nm)
97.0%
(25.8)
0
10
20
30
40
50
60
Time (min)
E.
1.5×10⁸
1×10⁸
5×10⁷
0
Ion Count
Avg: 1838.2
Mono: 1837.1
Found: 1838.0
0
5
10
15
Time (min)
F.
2.5×10⁶
2×10⁶
1.5×10⁶
1×10⁶
5×10⁵
0
Ion Count
613.67 (+3)
460.51
(+4)
920.00 (+2)
368.61
(+5)
0
1000
2000
3000
m/z

**Figure S165 – Analytical Data for FF-14 V6A. A.** Crude peptide HPLC trace at 214 nm with percent purity reflecting integration of the dominant peak area; retention time is shown in parentheses. **B.** Crude peptide LCMS total ion chromatogram (TIC); observed mass is provided along with the calculated average and monoisotopic masses. **C.** Mass spectrum at the peak of **B**. Peaks are labeled with *m/z* and charge state. **D.** HPLC of purified peptide. **E.** TIC of purified peptide. **F.** Mass spectrum at the peak of **E**. Peptides were used at the indicated purity following a single round of purification.

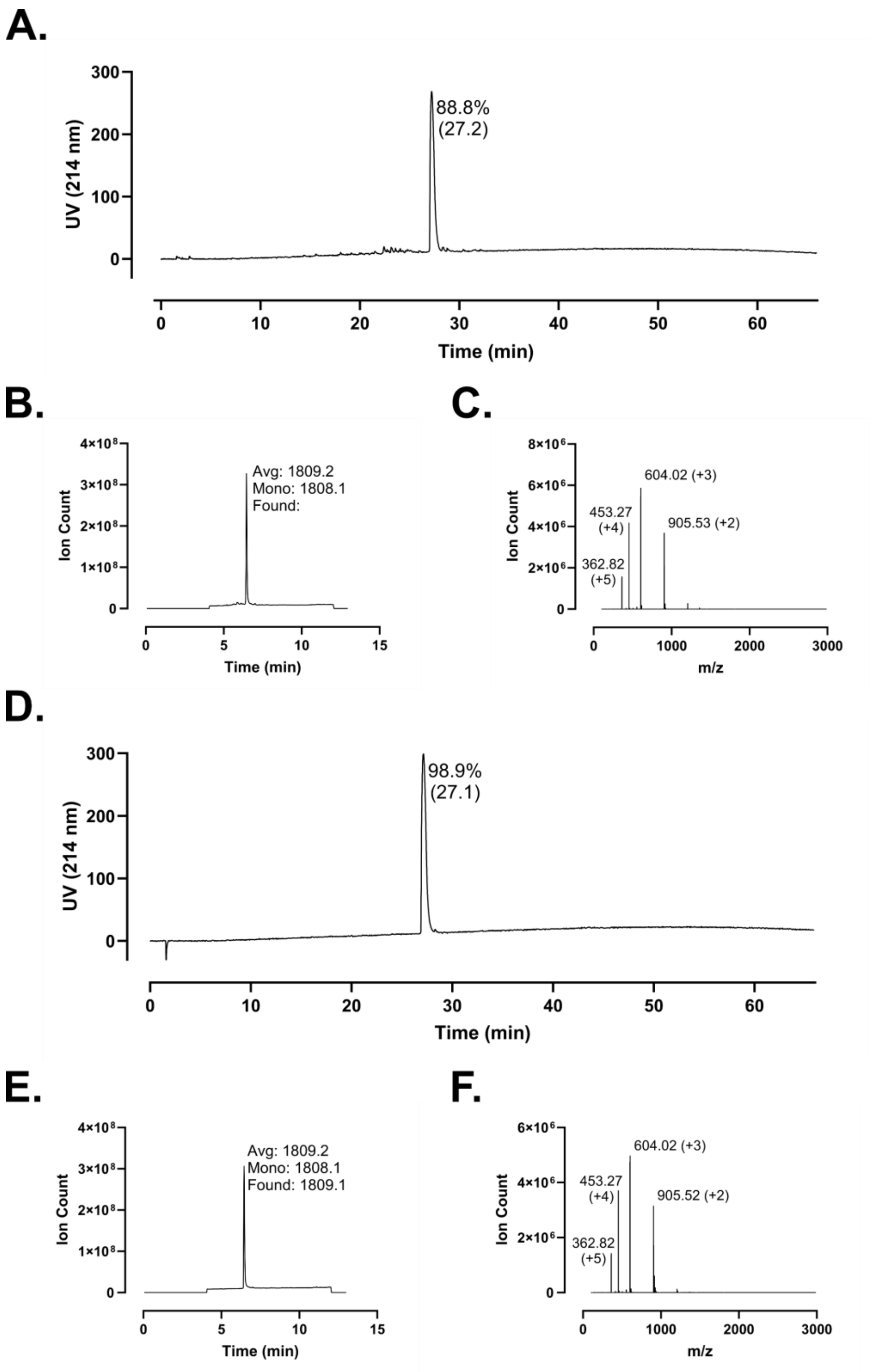
A.
UV (214 nm)
Time (min)
88.8%
(27.2)
B.
Ion Count
Time (min)
Avg: 1809.2
Mono: 1808.1
Found:
C.
Ion Count
m/z
604.02 (+3)
453.27
(+4)
905.53 (+2)
362.82
(+5)
D.
UV (214 nm)
Time (min)
98.9%
(27.1)
E.
Ion Count
Time (min)
Avg: 1809.2
Mono: 1808.1
Found: 1809.1
F.
Ion Count
m/z
604.02 (+3)
453.27
(+4)
905.52 (+2)
362.82
(+5)

**Figure S166 – Analytical Data for FF-14 Q7A. A.** Crude peptide HPLC trace at 214 nm with percent purity reflecting integration of the dominant peak area; retention time is shown in parentheses. **B.** Crude peptide LCMS total ion chromatogram (TIC); observed mass is provided along with the calculated average and monoisotopic masses. **C.** Mass spectrum at the peak of **B**. Peaks are labeled with *m/z* and charge state. **D.** HPLC of purified peptide. **E.** TIC of purified peptide. **F.** Mass spectrum at the peak of **E**. Peptides were used at the indicated purity following a single round of purification.

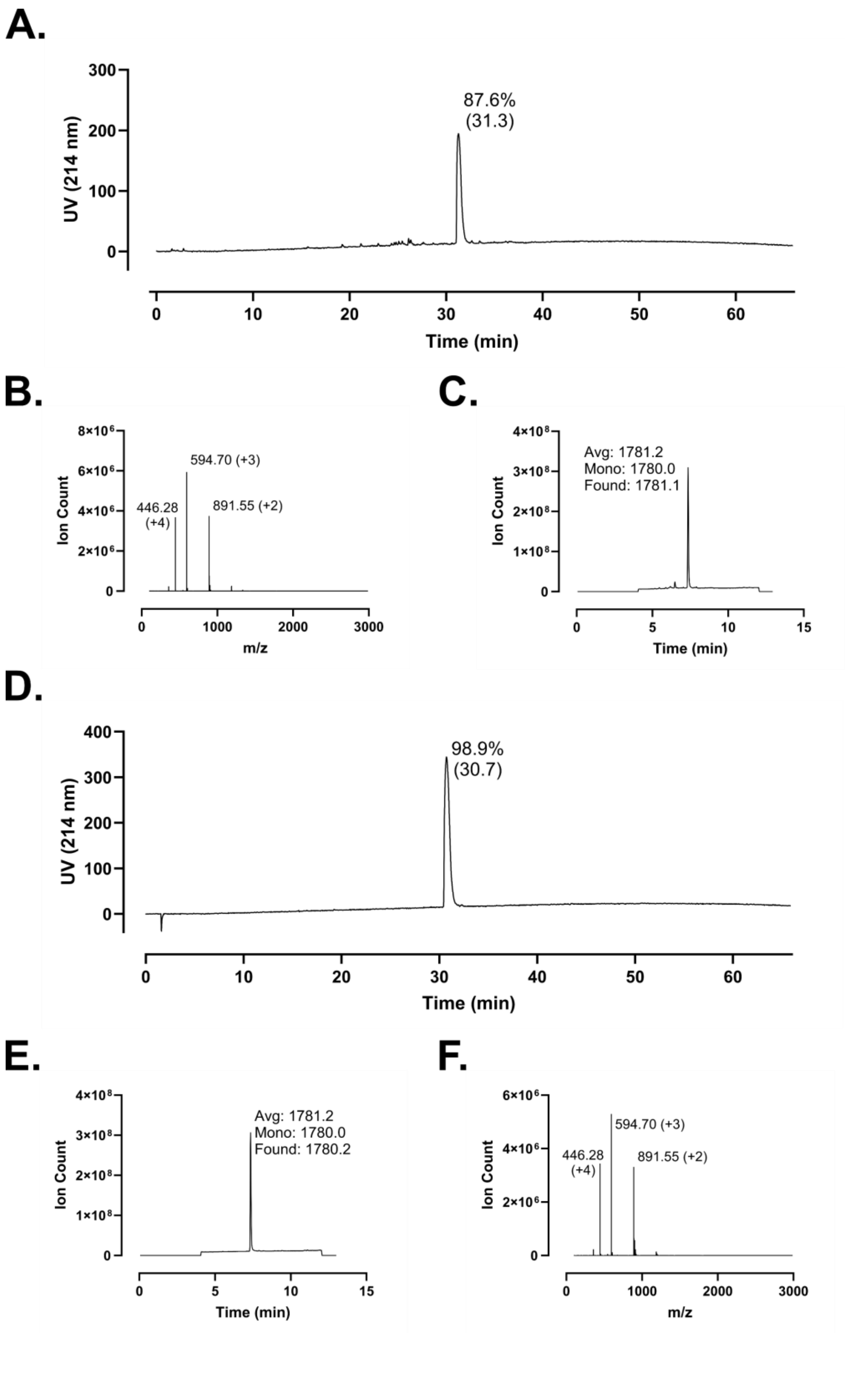
A.
UV (214 nm)
300
200
100
0
87.6%
(31.3)
0
10
20
30
40
50
60
Time (min)
B.
Ion Count
8×10⁶
6×10⁶
4×10⁶
2×10⁶
0
594.70 (+3)
446.28
(+4)
891.55 (+2)
0
1000
2000
3000
m/z
C.
Ion Count
4×10⁸
3×10⁸
2×10⁸
1×10⁸
0
Avg: 1781.2
Mono: 1780.0
Found: 1781.1
0
5
10
15
Time (min)
D.
UV (214 nm)
400
300
200
100
0
98.9%
(30.7)
0
10
20
30
40
50
60
Time (min)
E.
Ion Count
4×10⁸
3×10⁸
2×10⁸
1×10⁸
0
Avg: 1781.2
Mono: 1780.0
Found: 1780.2
0
5
10
15
Time (min)
F.
Ion Count
6×10⁶
4×10⁶
2×10⁶
0
594.70 (+3)
446.28
(+4)
891.55 (+2)
0
1000
2000
3000
m/z

**Figure S167 – Analytical Data for FF-14 R8A. A.** Crude peptide HPLC trace at 214 nm with percent purity reflecting integration of the dominant peak area; retention time is shown in parentheses. **B.** Crude peptide LCMS total ion chromatogram (TIC); observed mass is provided along with the calculated average and monoisotopic masses. **C.** Mass spectrum at the peak of **B**. Peaks are labeled with *m/z* and charge state. **D.** HPLC of purified peptide. **E.** TIC of purified peptide. **F.** Mass spectrum at the peak of **E**. Peptides were used at the indicated purity following a single round of purification.

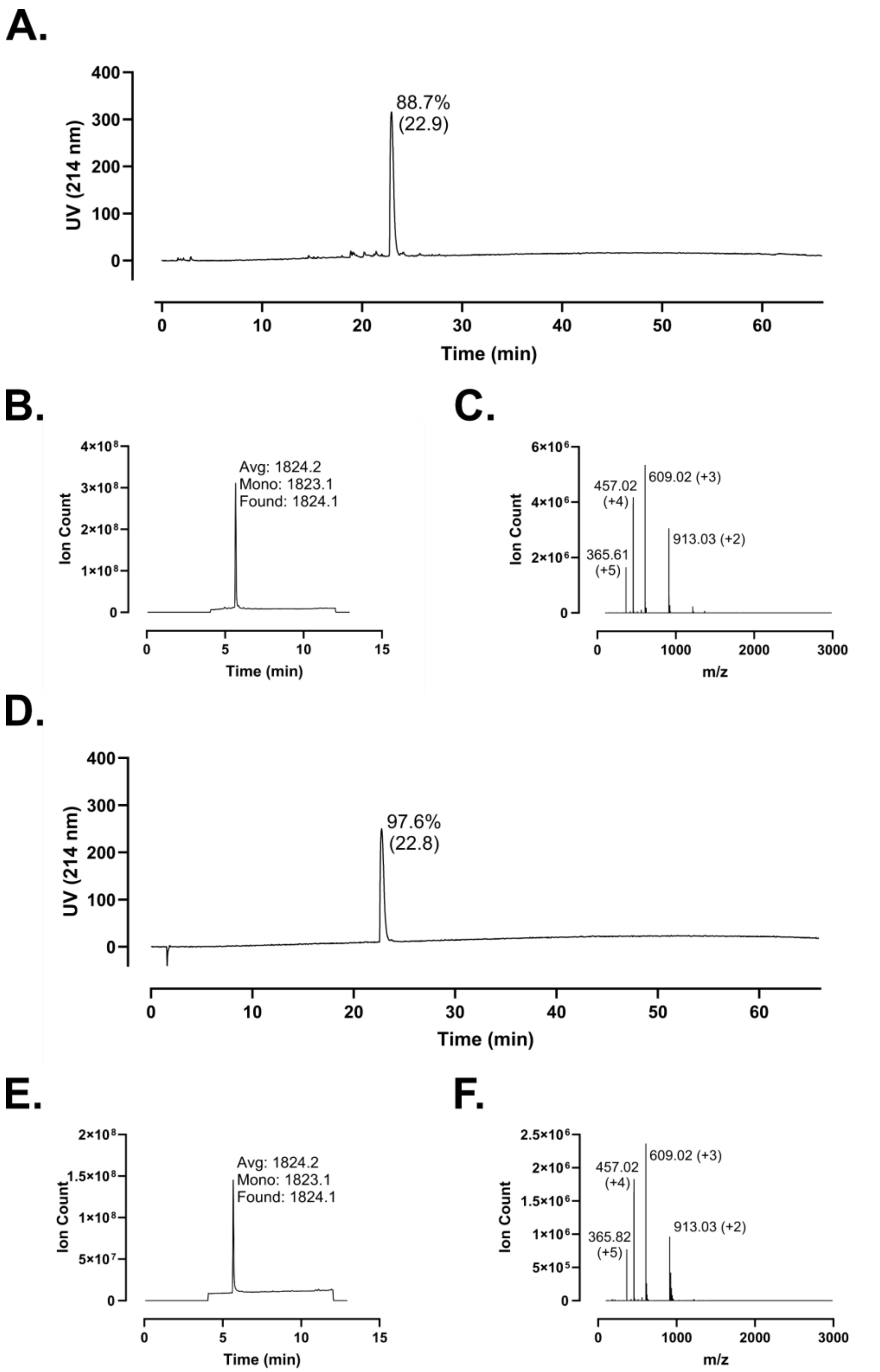
A.
88.7%
(22.9)
UV (214 nm)
Time (min)
B.
Avg: 1824.2
Mono: 1823.1
Found: 1824.1
Ion Count
Time (min)
C.
457.02
(+4)
609.02 (+3)
913.03 (+2)
365.61
(+5)
Ion Count
m/z
D.
97.6%
(22.8)
UV (214 nm)
Time (min)
E.
Avg: 1824.2
Mono: 1823.1
Found: 1824.1
Ion Count
Time (min)
F.
457.02
(+4)
609.02 (+3)
365.82
(+5)
913.03 (+2)
Ion Count
m/z

**Figure S168 – Analytical Data for FF-14 I9A. A.** Crude peptide HPLC trace at 214 nm with percent purity reflecting integration of the dominant peak area; retention time is shown in parentheses. **B.** Crude peptide LCMS total ion chromatogram (TIC); observed mass is provided along with the calculated average and monoisotopic masses. **C.** Mass spectrum at the peak of **B**. Peaks are labeled with *m/z* and charge state. **D.** HPLC of purified peptide. **E.** TIC of purified peptide. **F.** Mass spectrum at the peak of **E**. Peptides were used at the indicated purity following a single round of purification.

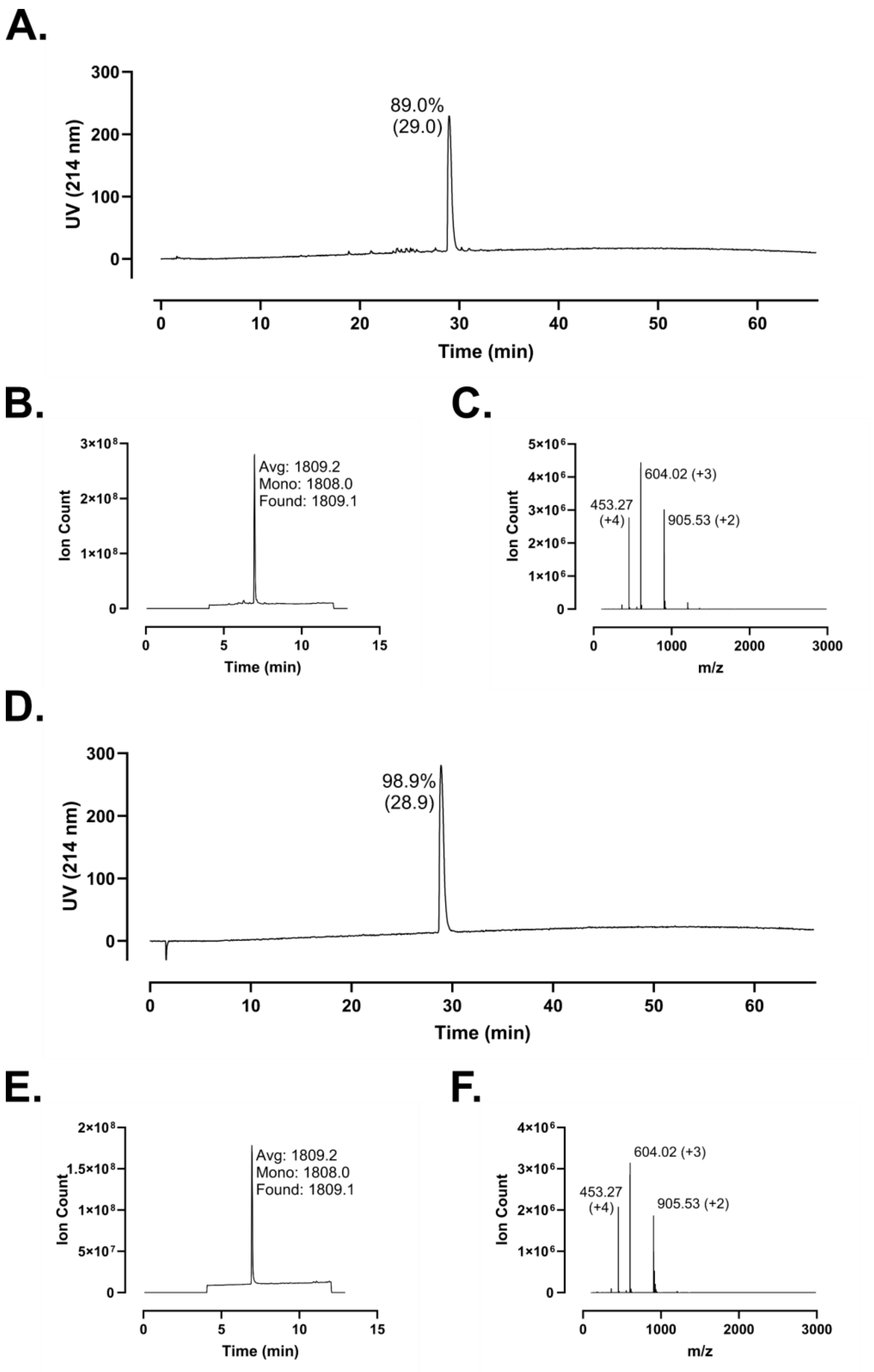
A.
89.0%
(29.0)
UV (214 nm)
Time (min)
B.
Avg: 1809.2
Mono: 1808.0
Found: 1809.1
Ion Count
Time (min)
C.
604.02 (+3)
453.27
(+4)
905.53 (+2)
Ion Count
m/z
D.
98.9%
(28.9)
UV (214 nm)
Time (min)
E.
Avg: 1809.2
Mono: 1808.0
Found: 1809.1
Ion Count
Time (min)
F.
604.02 (+3)
453.27
(+4)
905.53 (+2)
Ion Count
m/z

**Figure S169 – Analytical Data for FF-14 K10A. A.** Crude peptide HPLC trace at 214 nm with percent purity reflecting integration of the dominant peak area; retention time is shown in parentheses. **B.** Crude peptide LCMS total ion chromatogram (TIC); observed mass is provided along with the calculated average and monoisotopic masses. **C.** Mass spectrum at the peak of **B**. Peaks are labeled with *m/z* and charge state. **D.** HPLC of purified peptide. **E.** TIC of purified peptide. **F.** Mass spectrum at the peak of **E**. Peptides were used at the indicated purity following a single round of purification.

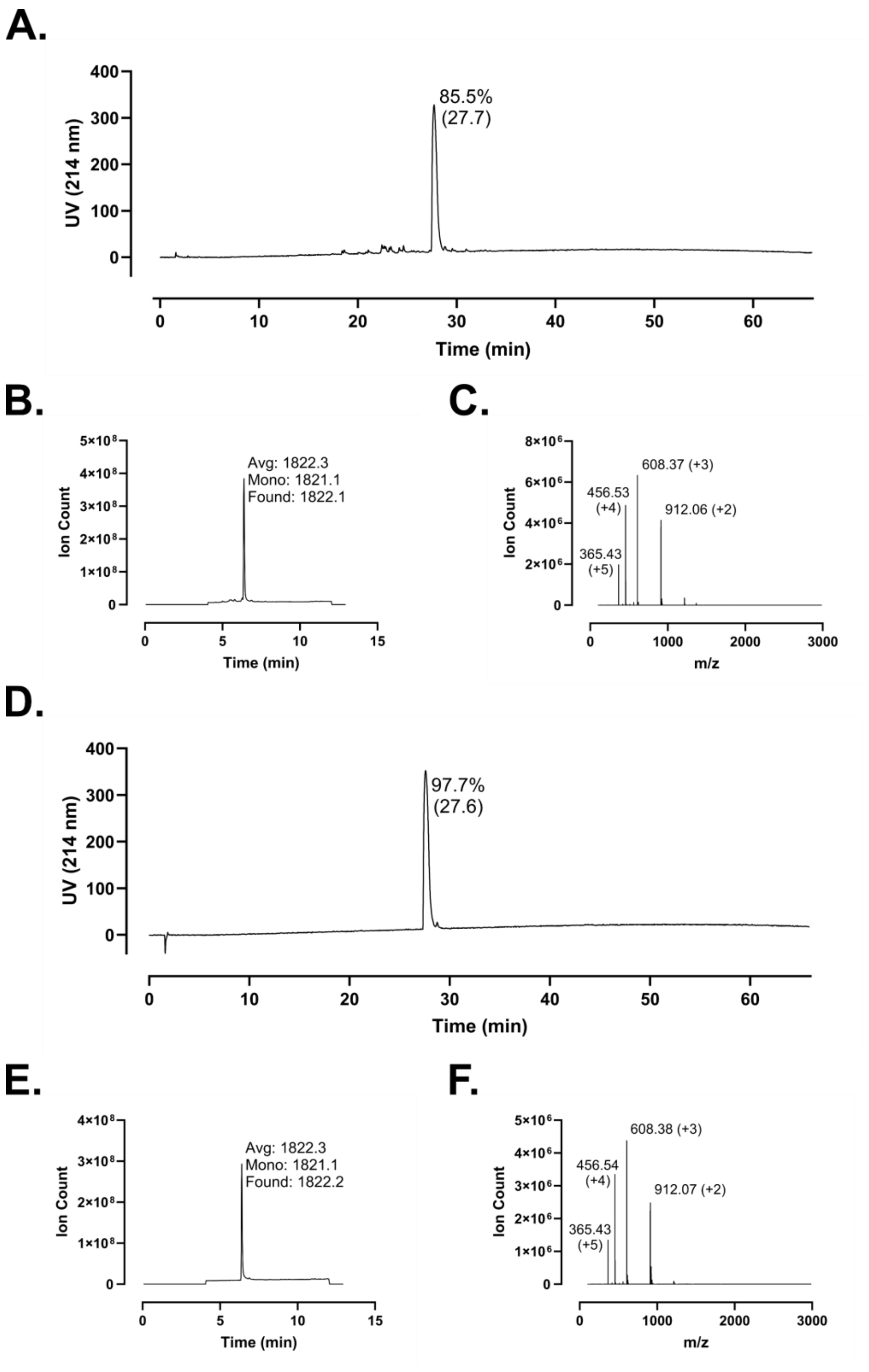
A.
UV (214 nm)
Time (min)
85.5%
(27.7)
B.
Ion Count
Time (min)
Avg: 1822.3
Mono: 1821.1
Found: 1822.1
C.
Ion Count
m/z
608.37 (+3)
456.53
(+4)
912.06 (+2)
365.43
(+5)
D.
UV (214 nm)
Time (min)
97.7%
(27.6)
E.
Ion Count
Time (min)
Avg: 1822.3
Mono: 1821.1
Found: 1822.2
F.
Ion Count
m/z
608.38 (+3)
456.54
(+4)
912.07 (+2)
365.43
(+5)

**Figure S170 – Analytical Data for FF-14 D11A. A.** Crude peptide HPLC trace at 214 nm with percent purity reflecting integration of the dominant peak area; retention time is shown in parentheses. **B.** Crude peptide LCMS total ion chromatogram (TIC); observed mass is provided along with the calculated average and monoisotopic masses. **C.** Mass spectrum at the peak of **B**. Peaks are labeled with *m/z* and charge state. **D.** HPLC of purified peptide. **E.** TIC of purified peptide. **F.** Mass spectrum at the peak of **E**. Peptides were used at the indicated purity following a single round of purification.

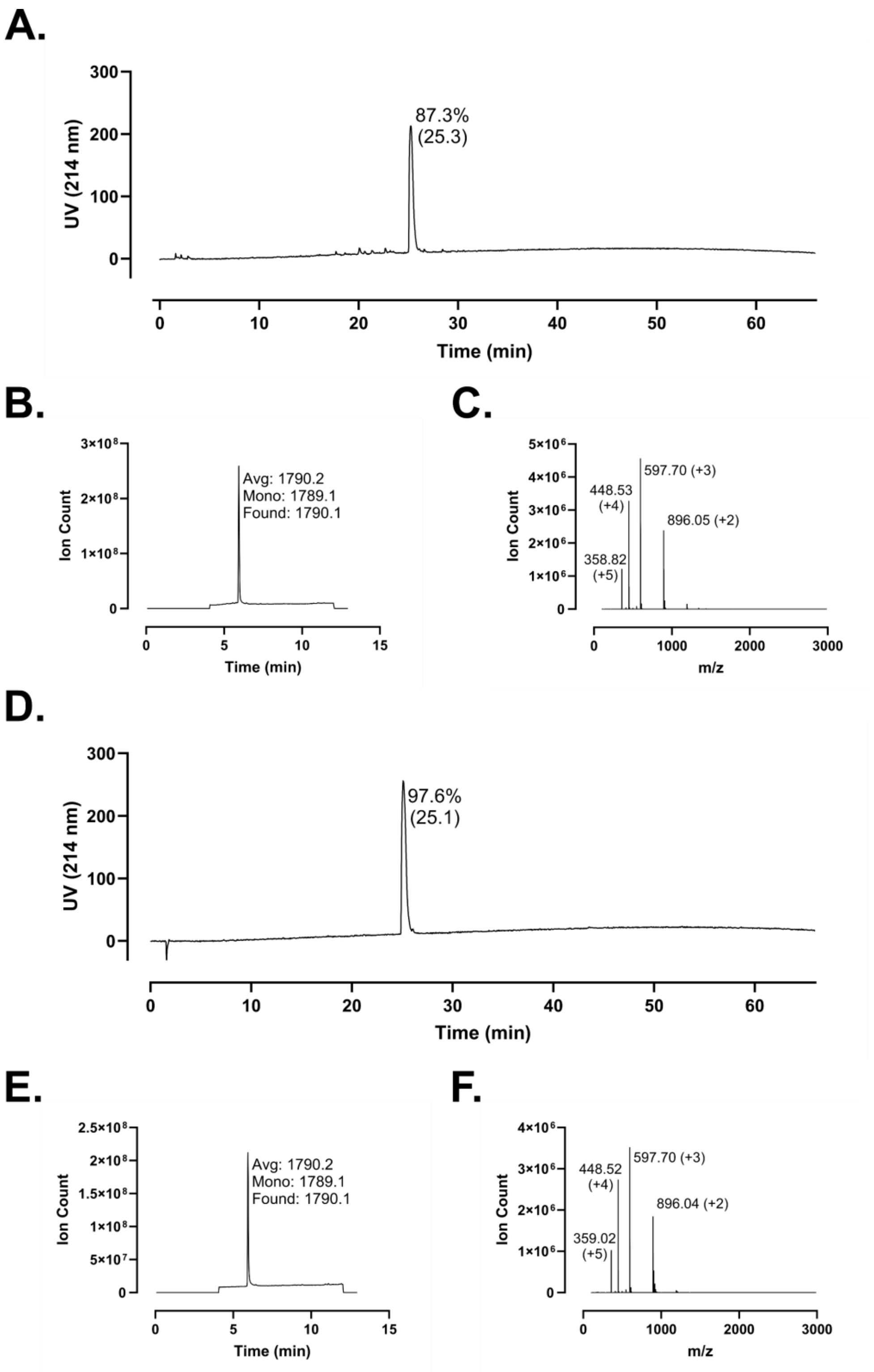
A.
UV (214 nm)
87.3%
(25.3)
Time (min)
B.
Ion Count
Avg: 1790.2
Mono: 1789.1
Found: 1790.1
Time (min)
C.
Ion Count
597.70 (+3)
448.53
(+4)
896.05 (+2)
358.82
(+5)
m/z
D.
UV (214 nm)
97.6%
(25.1)
Time (min)
E.
Ion Count
Avg: 1790.2
Mono: 1789.1
Found: 1790.1
Time (min)
F.
Ion Count
597.70 (+3)
448.52
(+4)
896.04 (+2)
359.02
(+5)
m/z

**Figure S171 – Analytical Data for FF-14 F12A. A.** Crude peptide HPLC trace at 214 nm with percent purity reflecting integration of the dominant peak area; retention time is shown in parentheses. **B.** Crude peptide LCMS total ion chromatogram (TIC); observed mass is provided along with the calculated average and monoisotopic masses. **C.** Mass spectrum at the peak of **B**. Peaks are labeled with *m/z* and charge state. **D.** HPLC of purified peptide. **E.** TIC of purified peptide. **F.** Mass spectrum at the peak of **E**. Peptides were used at the indicated purity following a single round of purification.

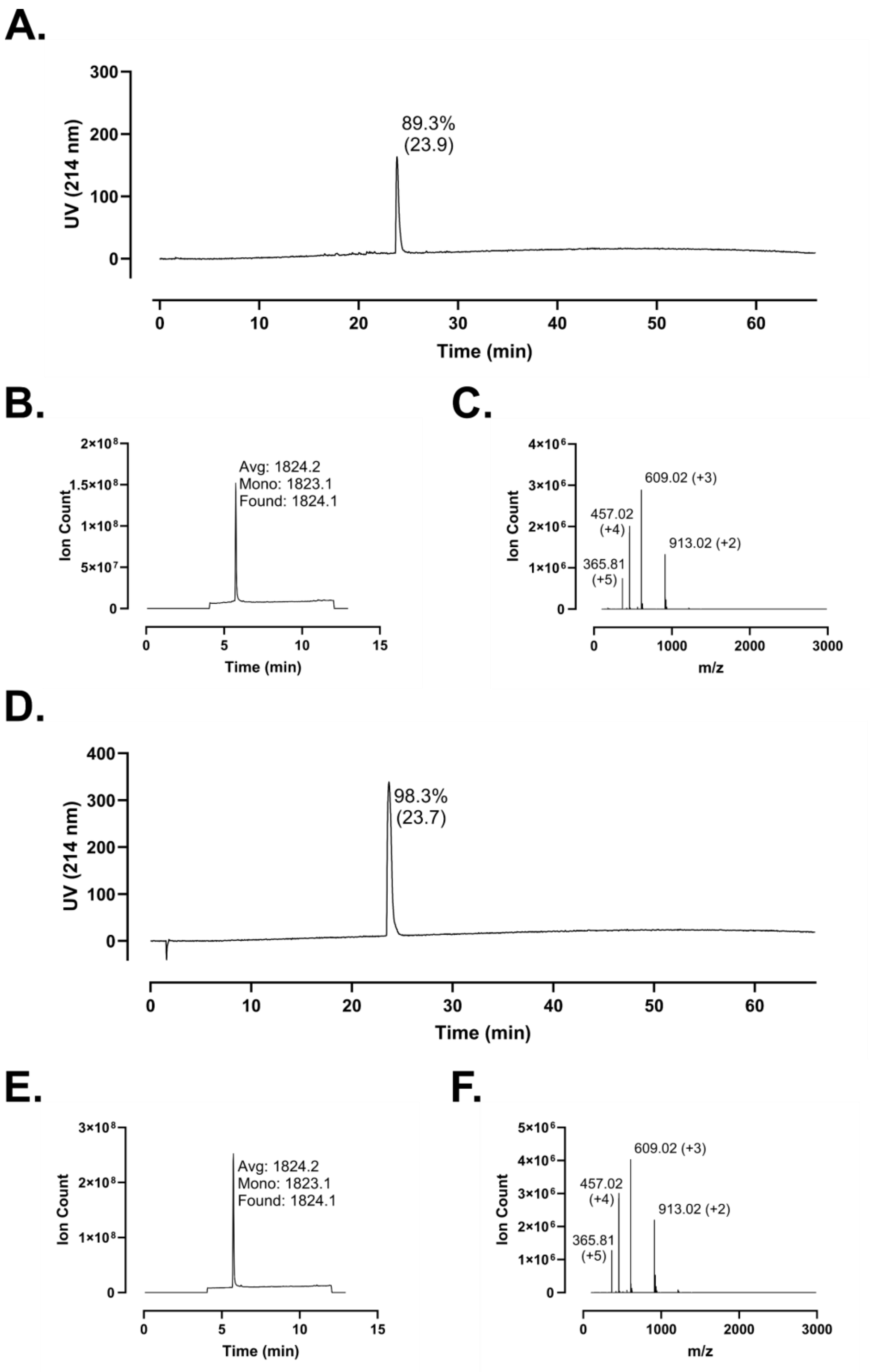
A.
89.3%
(23.9)
UV (214 nm)
Time (min)
B.
Avg: 1824.2
Mono: 1823.1
Found: 1824.1
Ion Count
Time (min)
C.
609.02 (+3)
457.02
(+4)
913.02 (+2)
365.81
(+5)
Ion Count
m/z
D.
98.3%
(23.7)
UV (214 nm)
Time (min)
E.
Avg: 1824.2
Mono: 1823.1
Found: 1824.1
Ion Count
Time (min)
F.
609.02 (+3)
457.02
(+4)
913.02 (+2)
365.81
(+5)
Ion Count
m/z

**Figure S172 – Analytical Data for FF-14 L13A. A.** Crude peptide HPLC trace at 214 nm with percent purity reflecting integration of the dominant peak area; retention time is shown in parentheses. **B.** Crude peptide LCMS total ion chromatogram (TIC); observed mass is provided along with the calculated average and monoisotopic masses. **C.** Mass spectrum at the peak of **B**. Peaks are labeled with *m/z* and charge state. **D.** HPLC of purified peptide. **E.** TIC of purified peptide. **F.** Mass spectrum at the peak of **E**. Peptides were used at the indicated purity following a single round of purification.

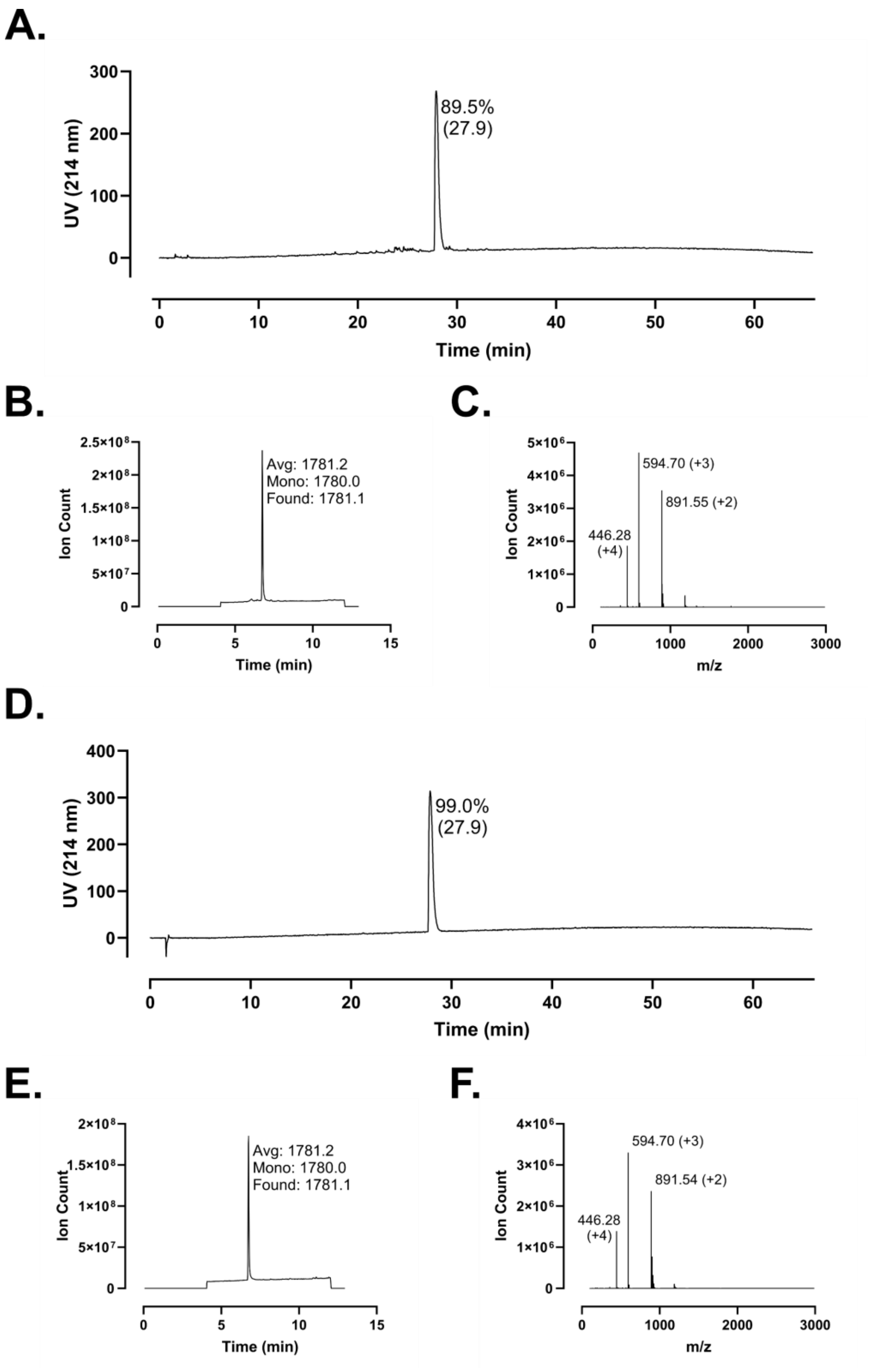
A.
UV (214 nm)
300
200
100
0
89.5%
(27.9)
0
10
20
30
40
50
60
Time (min)
B.
Ion Count
Avg: 1781.2
Mono: 1780.0
Found: 1781.1
Time (min)
C.
Ion Count
594.70 (+3)
891.55 (+2)
446.28
(+4)
m/z
D.
UV (214 nm)
99.0%
(27.9)
Time (min)
E.
Ion Count
Avg: 1781.2
Mono: 1780.0
Found: 1781.1
Time (min)
F.
Ion Count
594.70 (+3)
891.54 (+2)
446.28
(+4)
m/z

**Figure S173 – Analytical Data for FF-14 R14A. A.** Crude peptide HPLC trace at 214 nm with percent purity reflecting integration of the dominant peak area; retention time is shown in parentheses. **B.** Crude peptide LCMS total ion chromatogram (TIC); observed mass is provided along with the calculated average and monoisotopic masses. **C.** Mass spectrum at the peak of **B**. Peaks are labeled with *m/z* and charge state. **D.** HPLC of purified peptide. **E.** TIC of purified peptide. **F.** Mass spectrum at the peak of **E**. Peptides were used at the indicated purity following a single round of purification.

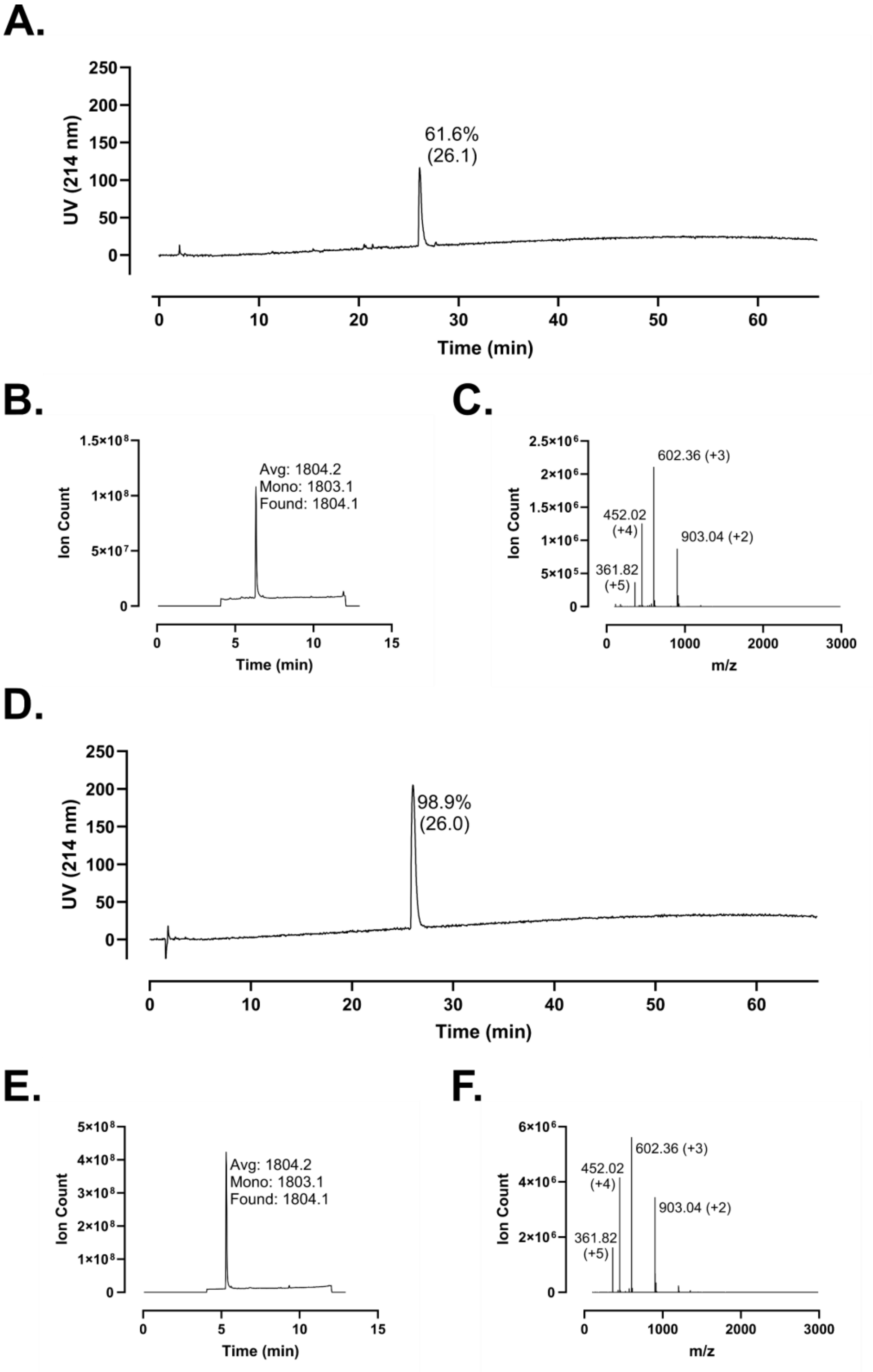
A.
UV (214 nm)
Time (min)
61.6%
(26.1)
B.
Ion Count
Time (min)
Avg: 1804.2
Mono: 1803.1
Found: 1804.1
C.
Ion Count
m/z
602.36 (+3)
452.02
(+4)
903.04 (+2)
361.82
(+5)
D.
UV (214 nm)
Time (min)
98.9%
(26.0)
E.
Ion Count
Time (min)
Avg: 1804.2
Mono: 1803.1
Found: 1804.1
F.
Ion Count
m/z
602.36 (+3)
452.02
(+4)
903.04 (+2)
361.82
(+5)

**Figure S174 – Analytical Data for FF-14 F1B. A.** Crude peptide HPLC trace at 214 nm with percent purity reflecting integration of the dominant peak area; retention time is shown in parentheses. **B.** Crude peptide LCMS total ion chromatogram (TIC); observed mass is provided along with the calculated average and monoisotopic masses. **C.** Mass spectrum at the peak of **B**. Peaks are labeled with *m/z* and charge state. **D.** HPLC of purified peptide. **E.** TIC of purified peptide. **F.** Mass spectrum at the peak of **E**. Peptides were used at the indicated purity following a single round of purification.

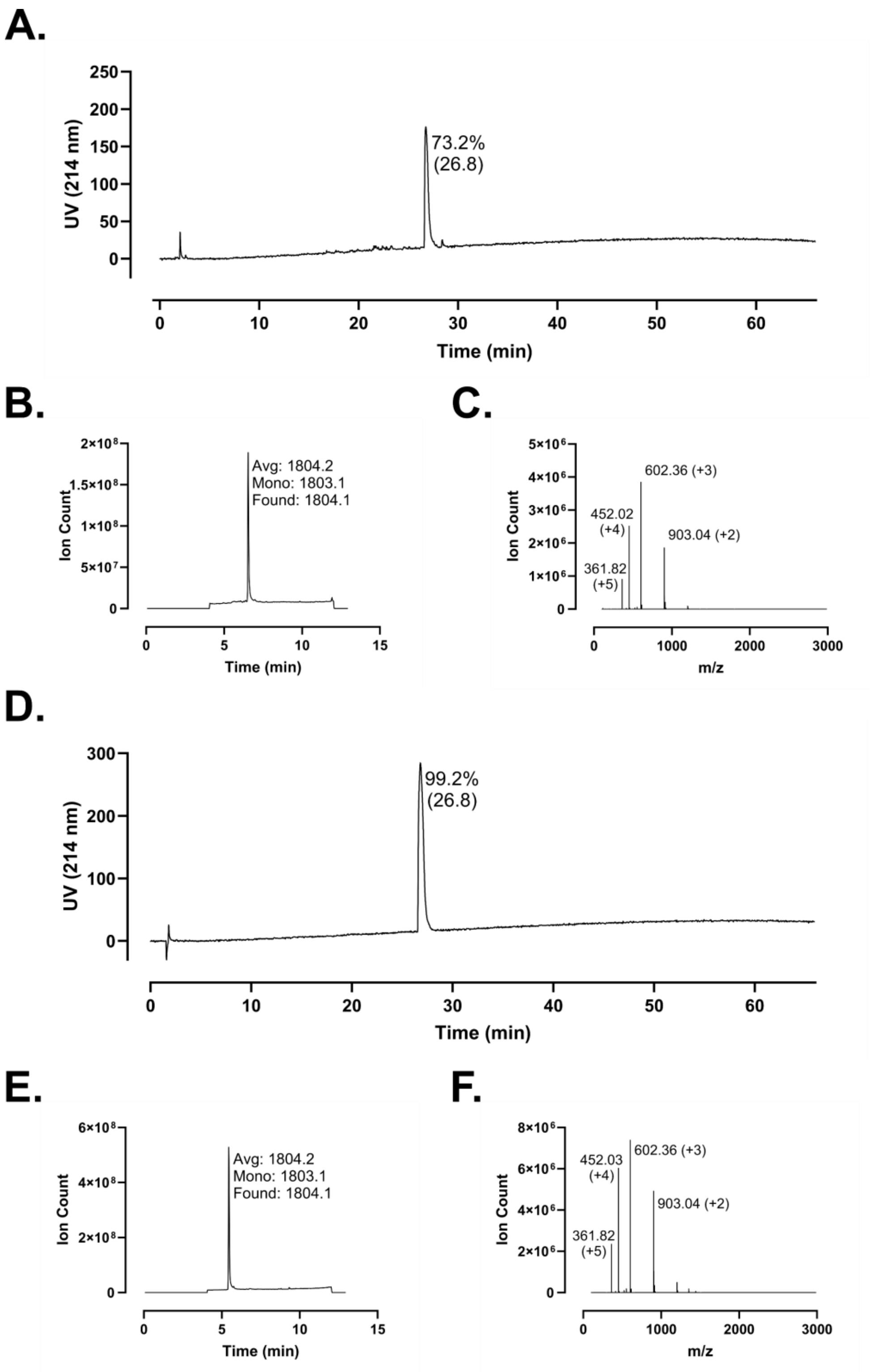
A.
UV (214 nm)
250
200
150
100
50
0
73.2%
(26.8)
Time (min)
0
10
20
30
40
50
60
B.
Ion Count
2×10⁸
1.5×10⁸
1×10⁸
5×10⁷
0
Avg: 1804.2
Mono: 1803.1
Found: 1804.1
Time (min)
0
5
10
15
C.
Ion Count
5×10⁶
4×10⁶
3×10⁶
2×10⁶
1×10⁶
0
602.36 (+3)
452.02
(+4)
903.04 (+2)
361.82
(+5)
m/z
0
1000
2000
3000
D.
UV (214 nm)
300
200
100
0
99.2%
(26.8)
Time (min)
0
10
20
30
40
50
60
E.
Ion Count
6×10⁸
4×10⁸
2×10⁸
0
Avg: 1804.2
Mono: 1803.1
Found: 1804.1
Time (min)
0
5
10
15
F.
Ion Count
8×10⁶
6×10⁶
4×10⁶
2×10⁶
0
452.03
(+4)
602.36 (+3)
903.04 (+2)
361.82
(+5)
m/z
0
1000
2000
3000

**Figure S175 – Analytical Data for FF-14 F2B. A.** Crude peptide HPLC trace at 214 nm with percent purity reflecting integration of the dominant peak area; retention time is shown in parentheses. **B.** Crude peptide LCMS total ion chromatogram (TIC); observed mass is provided along with the calculated average and monoisotopic masses. **C.** Mass spectrum at the peak of **B**. Peaks are labeled with *m/z* and charge state. **D.** HPLC of purified peptide. **E.** TIC of purified peptide. **F.** Mass spectrum at the peak of **E**. Peptides were used at the indicated purity following a single round of purification.

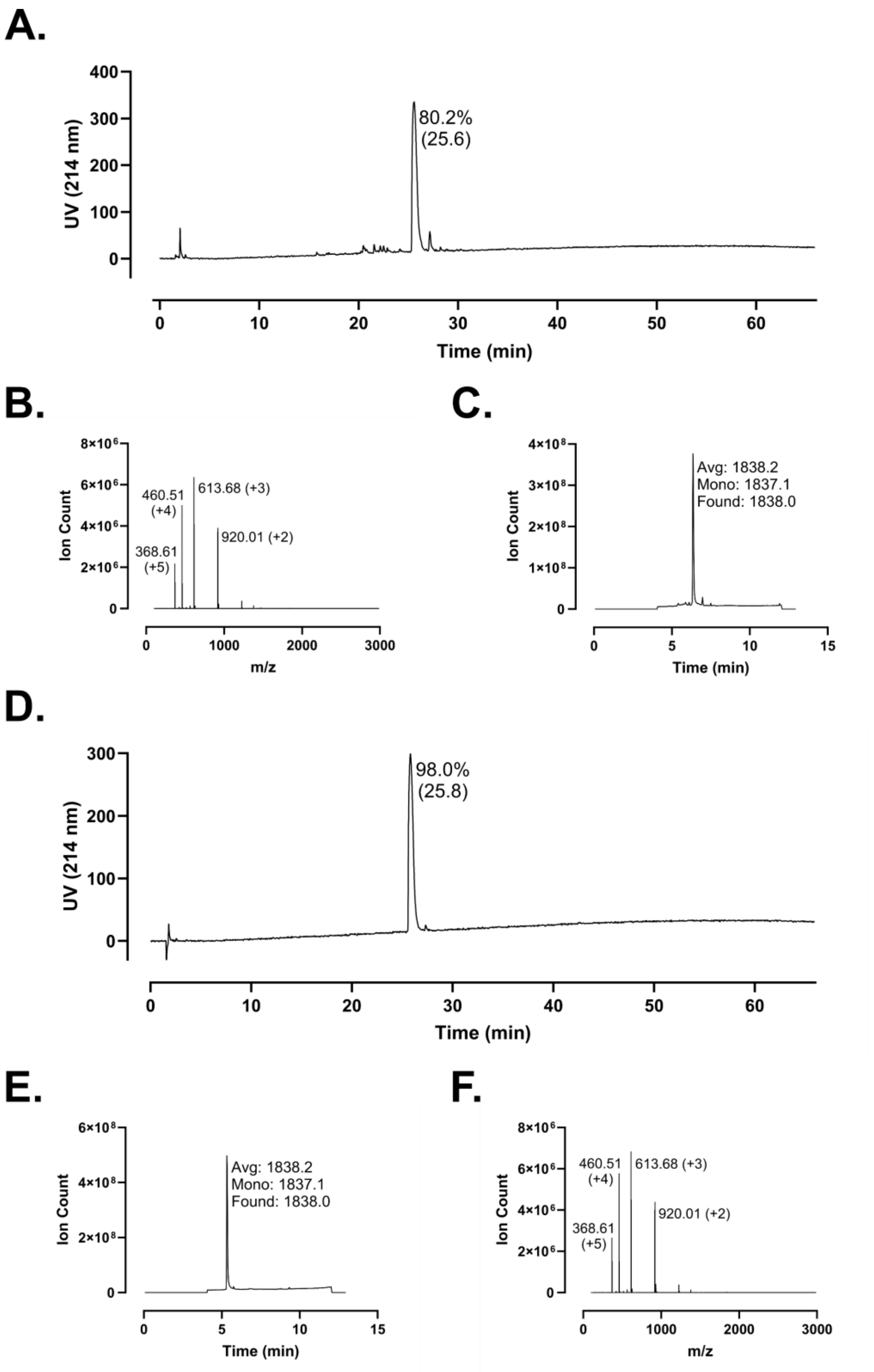
A.
UV (214 nm)
Time (min)
80.2%
(25.6)
B.
Ion Count
m/z
460.51
(+4)
613.68 (+3)
920.01 (+2)
368.61
(+5)
C.
Ion Count
Time (min)
Avg: 1838.2
Mono: 1837.1
Found: 1838.0
D.
UV (214 nm)
Time (min)
98.0%
(25.8)
E.
Ion Count
Time (min)
Avg: 1838.2
Mono: 1837.1
Found: 1838.0
F.
Ion Count
m/z
460.51
(+4)
613.68 (+3)
920.01 (+2)
368.61
(+5)

**Figure S176 – Analytical Data for FF-14 I5B. A.** Crude peptide HPLC trace at 214 nm with percent purity reflecting integration of the dominant peak area; retention time is shown in parentheses. **B.** Crude peptide LCMS total ion chromatogram (TIC); observed mass is provided along with the calculated average and monoisotopic masses. **C.** Mass spectrum at the peak of **B**. Peaks are labeled with *m/z* and charge state. **D.** HPLC of purified peptide. **E.** TIC of purified peptide. **F.** Mass spectrum at the peak of **E**. Peptides were used at the indicated purity following a single round of purification.

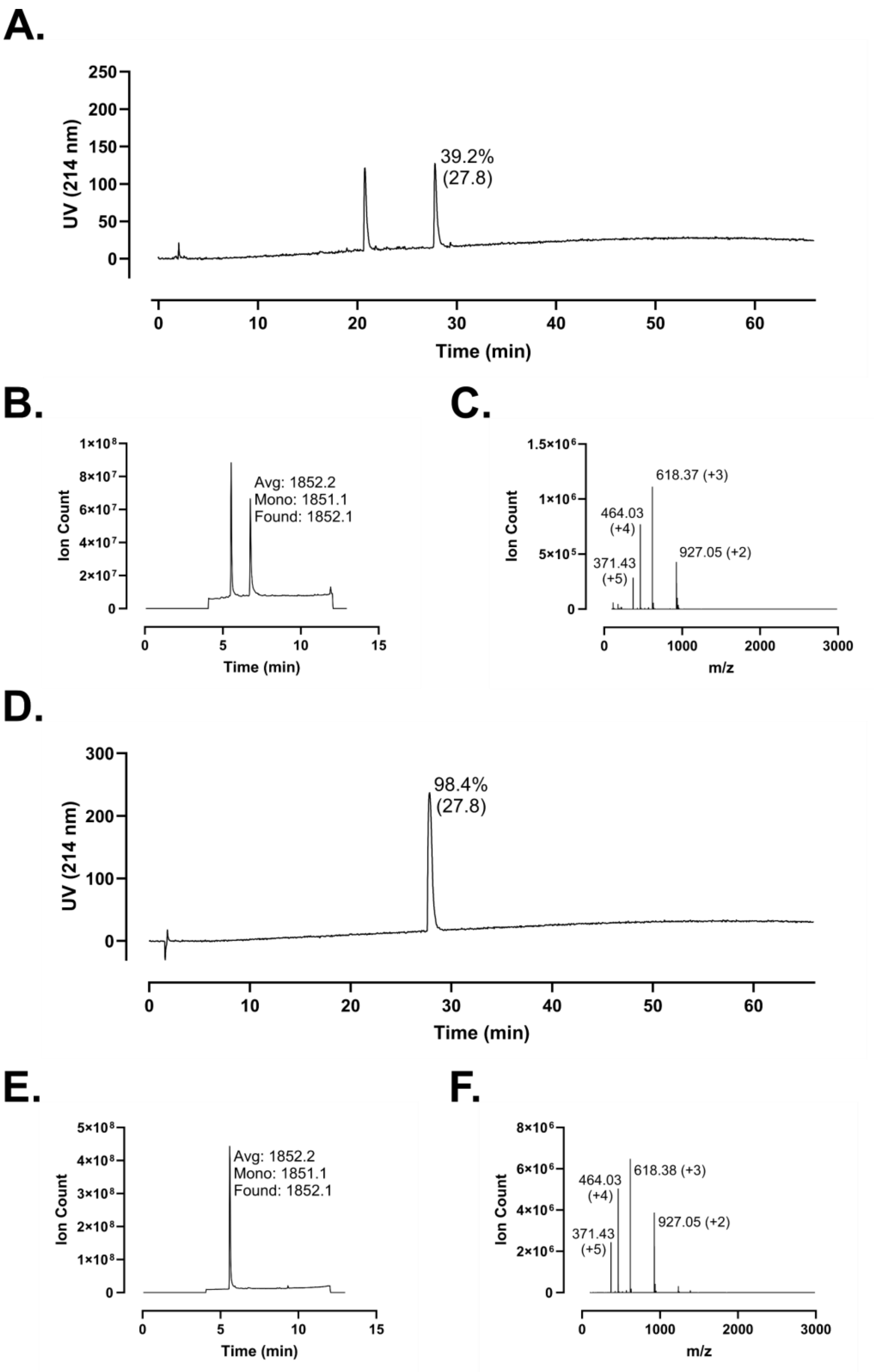
A.
UV (214 nm)
Time (min)
39.2%
(27.8)
B.
Ion Count
Time (min)
Avg: 1852.2
Mono: 1851.1
Found: 1852.1
C.
Ion Count
m/z
618.37 (+3)
464.03
(+4)
371.43
(+5)
927.05 (+2)
D.
UV (214 nm)
Time (min)
98.4%
(27.8)
E.
Ion Count
Time (min)
Avg: 1852.2
Mono: 1851.1
Found: 1852.1
F.
Ion Count
m/z
464.03
(+4)
618.38 (+3)
371.43
(+5)
927.05 (+2)

**Figure S177 – Analytical Data for FF-14 V6B. A.** Crude peptide HPLC trace at 214 nm with percent purity reflecting integration of the dominant peak area; retention time is shown in parentheses. **B.** Crude peptide LCMS total ion chromatogram (TIC); observed mass is provided along with the calculated average and monoisotopic masses. **C.** Mass spectrum at the peak of **B**. Peaks are labeled with *m/z* and charge state. **D.** HPLC of purified peptide. **E.** TIC of purified peptide. **F.** Mass spectrum at the peak of **E**. Peptides were used at the indicated purity following a single round of purification.

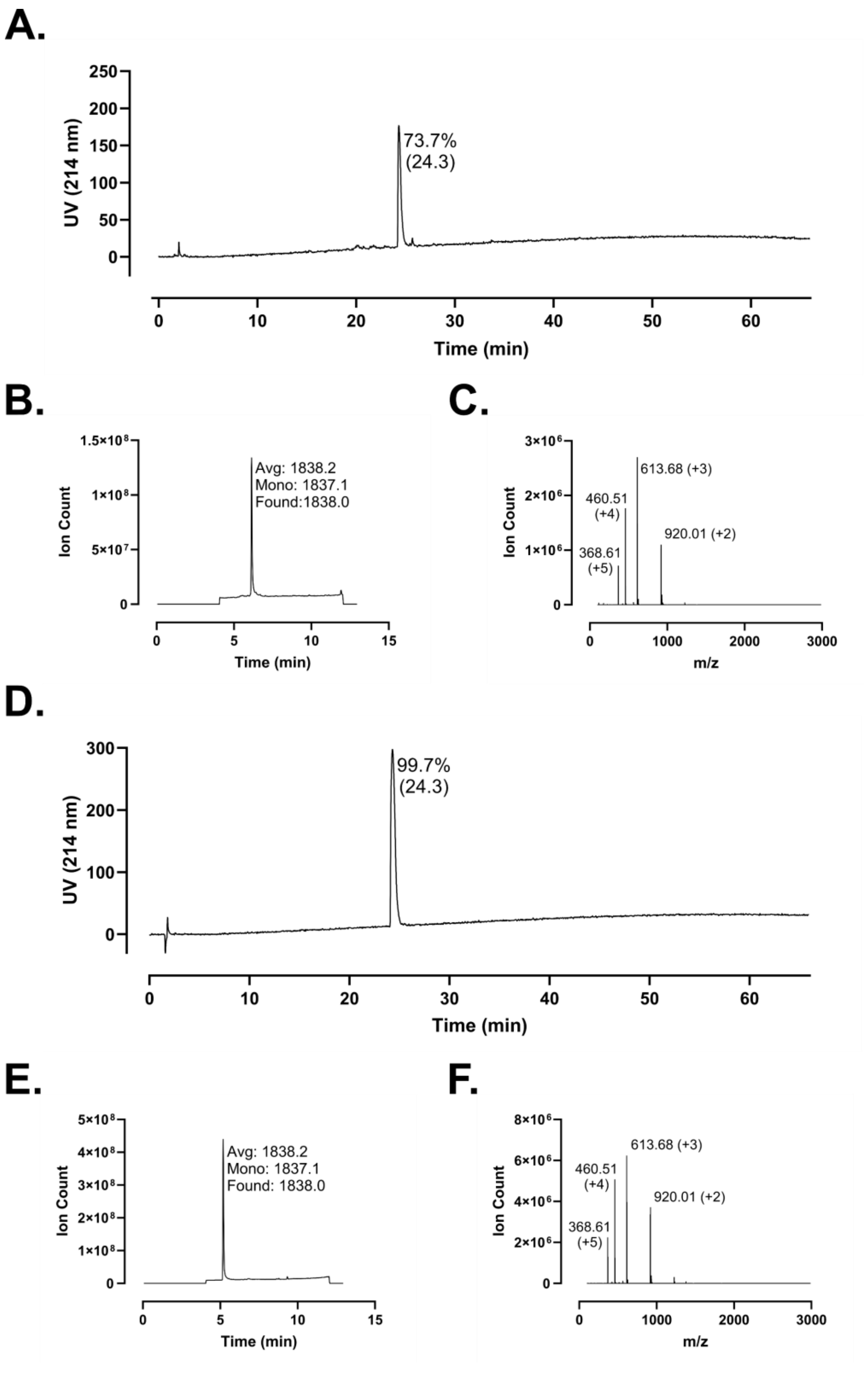
A.
UV (214 nm)
250
200
150
100
50
0
73.7%
(24.3)
0
10
20
30
40
50
60
Time (min)
B.
Ion Count
1.5×10^8
1×10^8
5×10^7
0
Avg: 1838.2
Mono: 1837.1
Found:1838.0
0
5
10
15
Time (min)
C.
Ion Count
3×10^6
2×10^6
1×10^6
0
613.68 (+3)
460.51
(+4)
920.01 (+2)
368.61
(+5)
0
1000
2000
3000
m/z
D.
UV (214 nm)
300
200
100
0
99.7%
(24.3)
0
10
20
30
40
50
60
Time (min)
E.
Ion Count
5×10^8
4×10^8
3×10^8
2×10^8
1×10^8
0
Avg: 1838.2
Mono: 1837.1
Found: 1838.0
0
5
10
15
Time (min)
F.
Ion Count
8×10^6
6×10^6
4×10^6
2×10^6
0
613.68 (+3)
460.51
(+4)
920.01 (+2)
368.61
(+5)
0
1000
2000
3000
m/z

**Figure S178 – Analytical Data for FF-14 Q7B. A.** Crude peptide HPLC trace at 214 nm with percent purity reflecting integration of the dominant peak area; retention time is shown in parentheses. **B.** Crude peptide LCMS total ion chromatogram (TIC); observed mass is provided along with the calculated average and monoisotopic masses. **C.** Mass spectrum at the peak of **B**. Peaks are labeled with *m/z* and charge state. **D.** HPLC of purified peptide. **E.** TIC of purified peptide. **F.** Mass spectrum at the peak of **E**. Peptides were used at the indicated purity following a single round of purification.

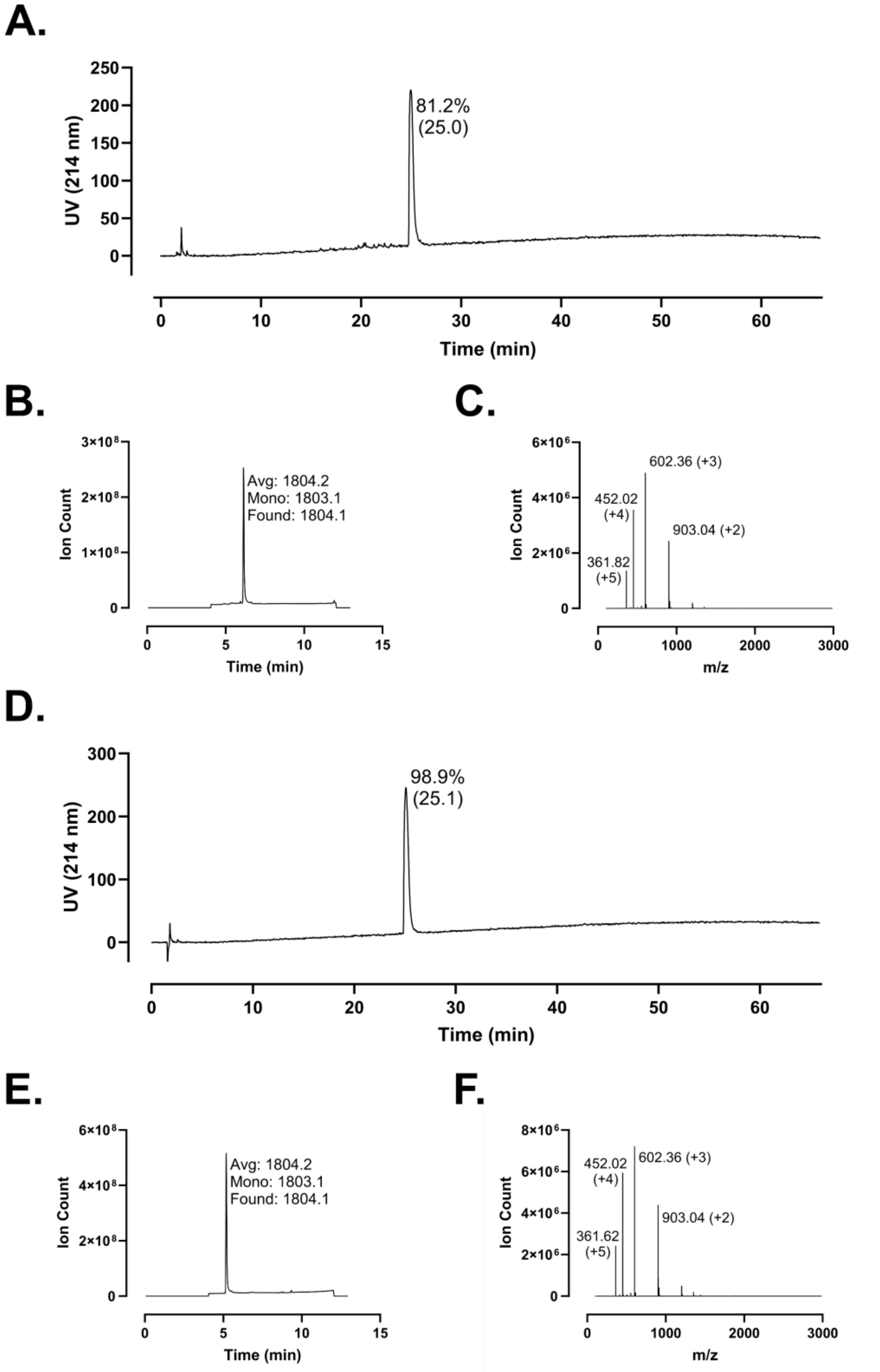
A.
UV (214 nm)
250
200
150
100
50
0
81.2%
(25.0)
0
10
20
30
40
50
60
Time (min)
B.
Ion Count
3×10⁸
2×10⁸
1×10⁸
0
Avg: 1804.2
Mono: 1803.1
Found: 1804.1
0
5
10
15
Time (min)
C.
Ion Count
6×10⁶
4×10⁶
2×10⁶
0
602.36 (+3)
452.02
(+4)
903.04 (+2)
361.82
(+5)
0
1000
2000
3000
m/z
D.
UV (214 nm)
300
200
100
0
98.9%
(25.1)
0
10
20
30
40
50
60
Time (min)
E.
Ion Count
6×10⁸
4×10⁸
2×10⁸
0
Avg: 1804.2
Mono: 1803.1
Found: 1804.1
0
5
10
15
Time (min)
F.
Ion Count
8×10⁶
6×10⁶
4×10⁶
2×10⁶
0
452.02
(+4)
602.36 (+3)
903.04 (+2)
361.62
(+5)
0
1000
2000
3000
m/z

**Figure S179 – Analytical Data for FF-14 R8B. A.** Crude peptide HPLC trace at 214 nm with percent purity reflecting integration of the dominant peak area; retention time is shown in parentheses. **B.** Crude peptide LCMS total ion chromatogram (TIC); observed mass is provided along with the calculated average and monoisotopic masses. **C.** Mass spectrum at the peak of **B**. Peaks are labeled with *m/z* and charge state. **D.** HPLC of purified peptide. **E.** TIC of purified peptide. **F.** Mass spectrum at the peak of **E**. Peptides were used at the indicated purity following a single round of purification.

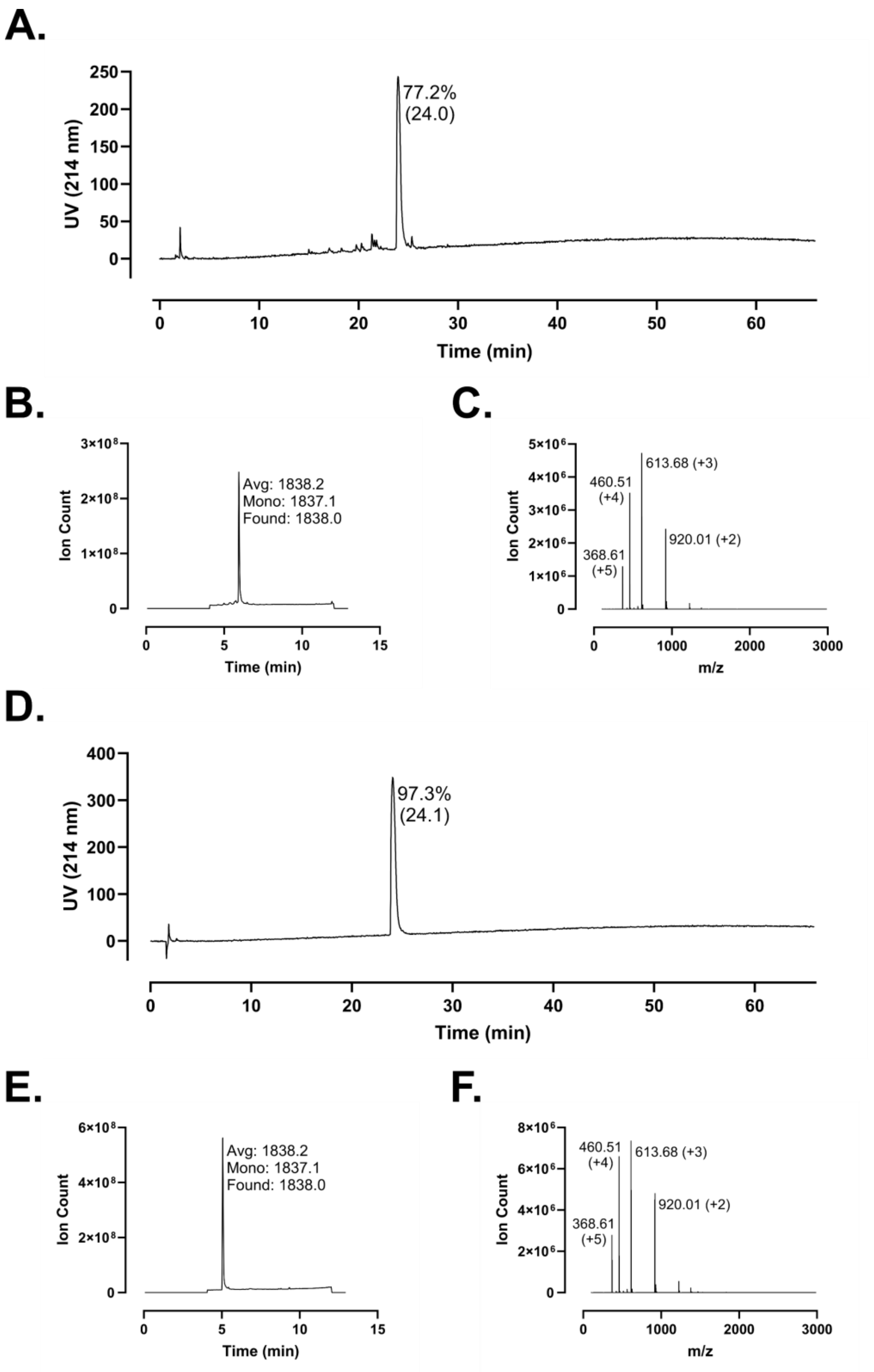
A.
UV (214 nm)
Time (min)
77.2%
(24.0)
B.
Ion Count
Time (min)
Avg: 1838.2
Mono: 1837.1
Found: 1838.0
C.
Ion Count
m/z
460.51
(+4)
613.68 (+3)
368.61
(+5)
920.01 (+2)
D.
UV (214 nm)
Time (min)
97.3%
(24.1)
E.
Ion Count
Time (min)
Avg: 1838.2
Mono: 1837.1
Found: 1838.0
F.
Ion Count
m/z
460.51
(+4)
613.68 (+3)
368.61
(+5)
920.01 (+2)

**Figure S180 – Analytical Data for FF-14 I9B. A.** Crude peptide HPLC trace at 214 nm with percent purity reflecting integration of the dominant peak area; retention time is shown in parentheses. **B.** Crude peptide LCMS total ion chromatogram (TIC); observed mass is provided along with the calculated average and monoisotopic masses. **C.** Mass spectrum at the peak of **B**. Peaks are labeled with *m/z* and charge state. **D.** HPLC of purified peptide. **E.** TIC of purified peptide. **F.** Mass spectrum at the peak of **E**. Peptides were used at the indicated purity following a single round of purification.

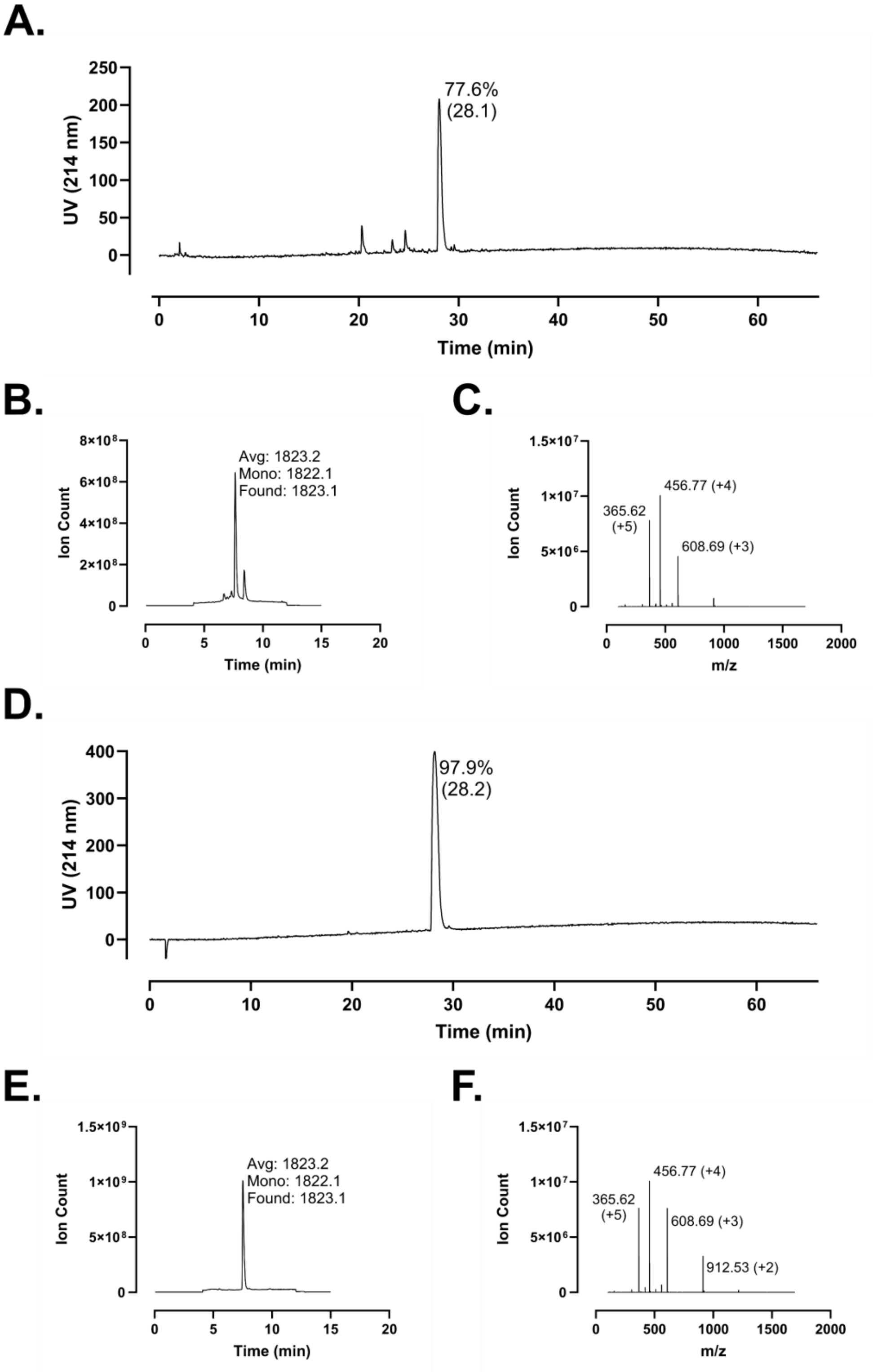
A.
UV (214 nm)
Time (min)
77.6%
(28.1)
B.
Ion Count
Time (min)
Avg: 1823.2
Mono: 1822.1
Found: 1823.1
C.
Ion Count
m/z
365.62
(+5)
456.77 (+4)
608.69 (+3)
D.
UV (214 nm)
Time (min)
97.9%
(28.2)
E.
Ion Count
Time (min)
Avg: 1823.2
Mono: 1822.1
Found: 1823.1
F.
Ion Count
m/z
365.62
(+5)
456.77 (+4)
608.69 (+3)
912.53 (+2)

**Figure S181 – Analytical Data for FF-14 K10B. A.** Crude peptide HPLC trace at 214 nm with percent purity reflecting integration of the dominant peak area; retention time is shown in parentheses. **B.** Crude peptide LCMS total ion chromatogram (TIC); observed mass is provided along with the calculated average and monoisotopic masses. **C.** Mass spectrum at the peak of **B**. Peaks are labeled with *m/z* and charge state. **D.** HPLC of purified peptide. **E.** TIC of purified peptide. **F.** Mass spectrum at the peak of **E**. Peptides were used at the indicated purity following a single round of purification.

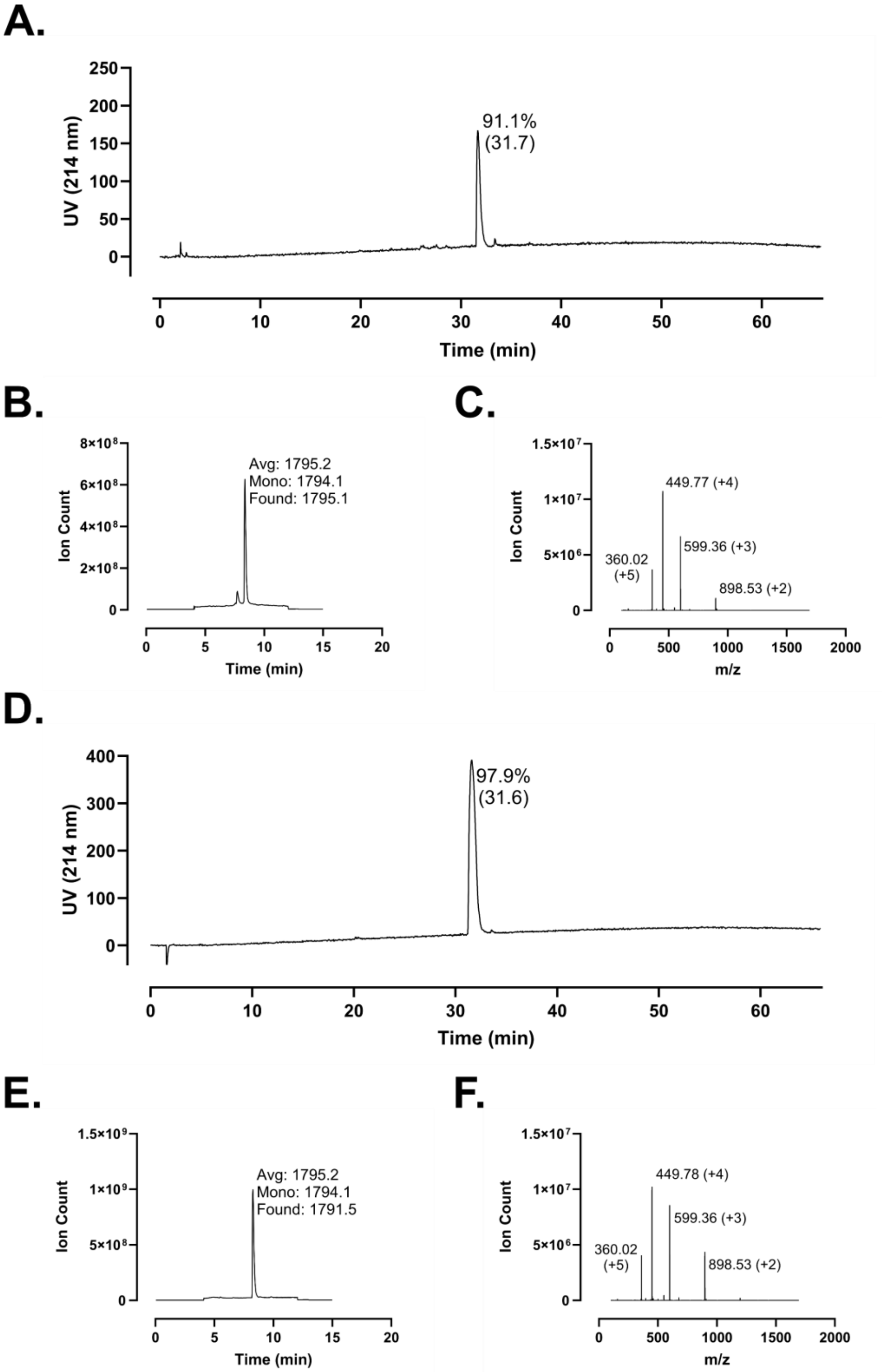
A.
UV (214 nm)
250
200
150
100
50
0
91.1%
(31.7)
0
10
20
30
40
50
60
Time (min)
B.
Ion Count
8×10⁸
6×10⁸
4×10⁸
2×10⁸
0
Avg: 1795.2
Mono: 1794.1
Found: 1795.1
0
5
10
15
20
Time (min)
C.
Ion Count
1.5×10⁷
1×10⁷
5×10⁶
0
449.77 (+4)
599.36 (+3)
360.02
(+5)
898.53 (+2)
0
500
1000
1500
2000
m/z
D.
UV (214 nm)
400
300
200
100
0
97.9%
(31.6)
0
10
20
30
40
50
60
Time (min)
E.
Ion Count
1.5×10⁹
1×10⁹
5×10⁸
0
Avg: 1795.2
Mono: 1794.1
Found: 1791.5
0
5
10
15
20
Time (min)
F.
Ion Count
1.5×10⁷
1×10⁷
5×10⁶
0
449.78 (+4)
599.36 (+3)
360.02
(+5)
898.53 (+2)
0
500
1000
1500
2000
m/z

**Figure S182 – Analytical Data for FF-14 D11B. A.** Crude peptide HPLC trace at 214 nm with percent purity reflecting integration of the dominant peak area; retention time is shown in parentheses. **B.** Crude peptide LCMS total ion chromatogram (TIC); observed mass is provided along with the calculated average and monoisotopic masses. **C.** Mass spectrum at the peak of **B**. Peaks are labeled with *m/z* and charge state. **D.** HPLC of purified peptide. **E.** TIC of purified peptide. **F.** Mass spectrum at the peak of **E**. Peptides were used at the indicated purity following a single round of purification.

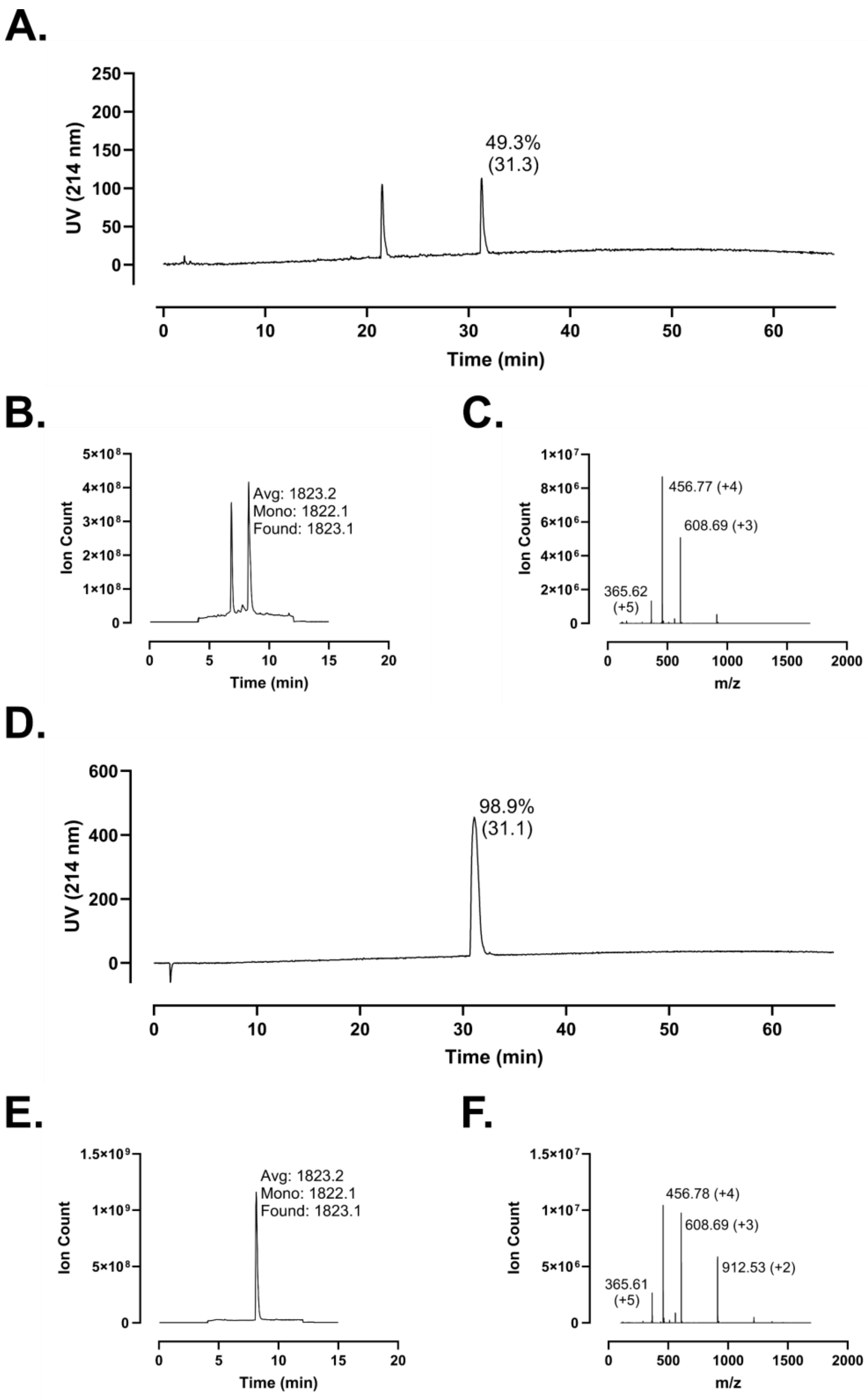
A.
UV (214 nm)
Time (min)
49.3%
(31.3)
B.
Ion Count
Time (min)
Avg: 1823.2
Mono: 1822.1
Found: 1823.1
C.
Ion Count
m/z
456.77 (+4)
608.69 (+3)
365.62
(+5)
D.
UV (214 nm)
Time (min)
98.9%
(31.1)
E.
Ion Count
Time (min)
Avg: 1823.2
Mono: 1822.1
Found: 1823.1
F.
Ion Count
m/z
456.78 (+4)
608.69 (+3)
912.53 (+2)
365.61
(+5)

**Figure S183 – Analytical Data for FF-14 F12B. A.** Crude peptide HPLC trace at 214 nm with percent purity reflecting integration of the dominant peak area; retention time is shown in parentheses. **B.** Crude peptide LCMS total ion chromatogram (TIC); observed mass is provided along with the calculated average and monoisotopic masses. **C.** Mass spectrum at the peak of **B**. Peaks are labeled with *m/z* and charge state. **D.** HPLC of purified peptide. **E.** TIC of purified peptide. **F.** Mass spectrum at the peak of **E**. Peptides were used at the indicated purity following a single round of purification.

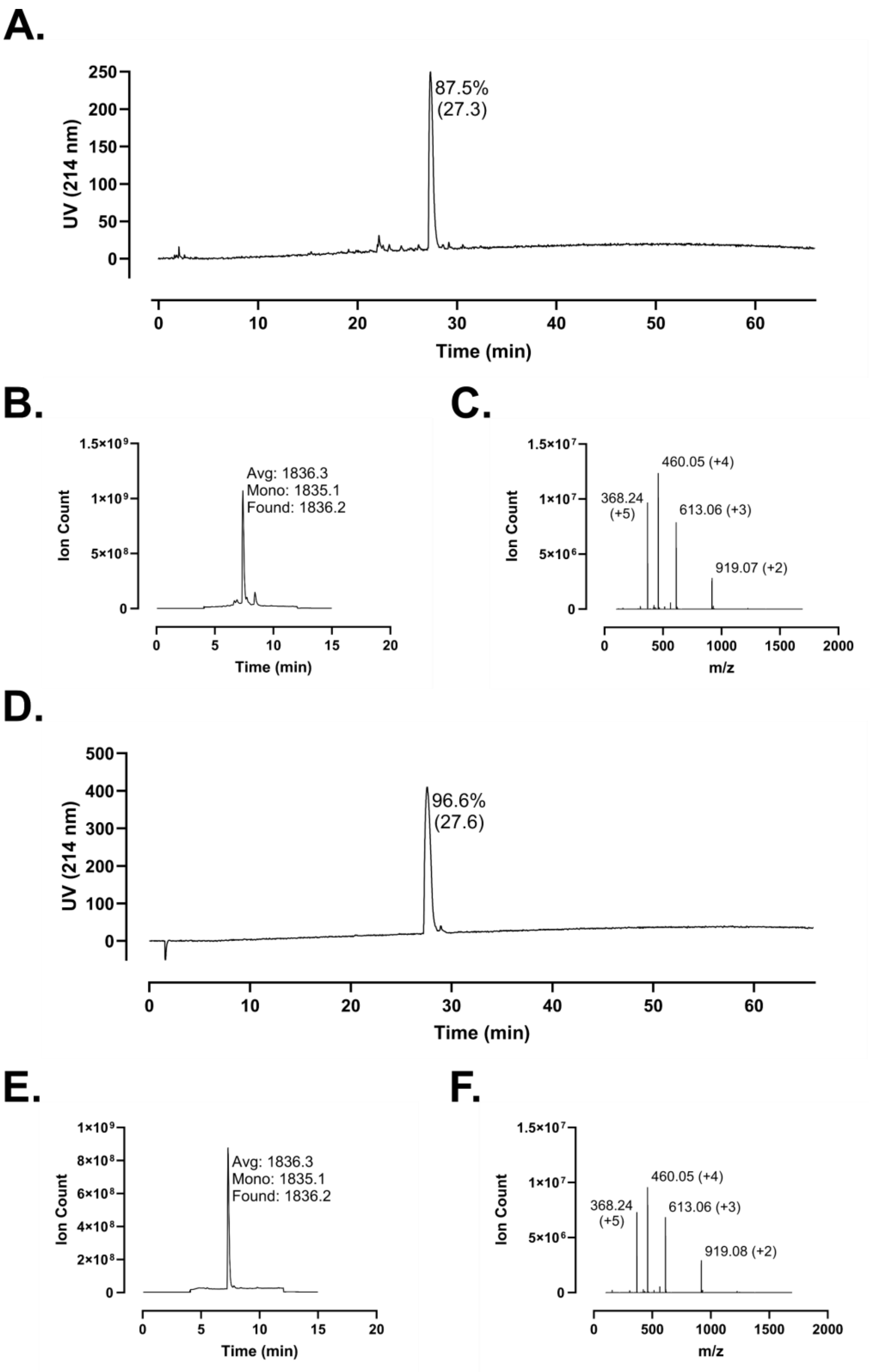
A.
UV (214 nm)
250
200
150
100
50
0
87.5%
(27.3)
0
10
20
30
40
50
60
Time (min)
B.
Ion Count
1.5×10⁹
1×10⁹
5×10⁸
0
Avg: 1836.3
Mono: 1835.1
Found: 1836.2
0
5
10
15
20
Time (min)
C.
Ion Count
1.5×10⁷
1×10⁷
5×10⁶
0
368.24
(+5)
460.05 (+4)
613.06 (+3)
919.07 (+2)
0
500
1000
1500
2000
m/z
D.
UV (214 nm)
500
400
300
200
100
0
96.6%
(27.6)
0
10
20
30
40
50
60
Time (min)
E.
Ion Count
1×10⁹
8×10⁸
6×10⁸
4×10⁸
2×10⁸
0
Avg: 1836.3
Mono: 1835.1
Found: 1836.2
0
5
10
15
20
Time (min)
F.
Ion Count
1.5×10⁷
1×10⁷
5×10⁶
0
368.24
(+5)
460.05 (+4)
613.06 (+3)
919.08 (+2)
0
500
1000
1500
2000
m/z

**Figure S184 – Analytical Data for FF-14 L13B. A.** Crude peptide HPLC trace at 214 nm with percent purity reflecting integration of the dominant peak area; retention time is shown in parentheses. **B.** Crude peptide LCMS total ion chromatogram (TIC); observed mass is provided along with the calculated average and monoisotopic masses. **C.** Mass spectrum at the peak of **B**. Peaks are labeled with *m/z* and charge state. **D.** HPLC of purified peptide. **E.** TIC of purified peptide. **F.** Mass spectrum at the peak of **E**. Peptides were used at the indicated purity following a single round of purification.

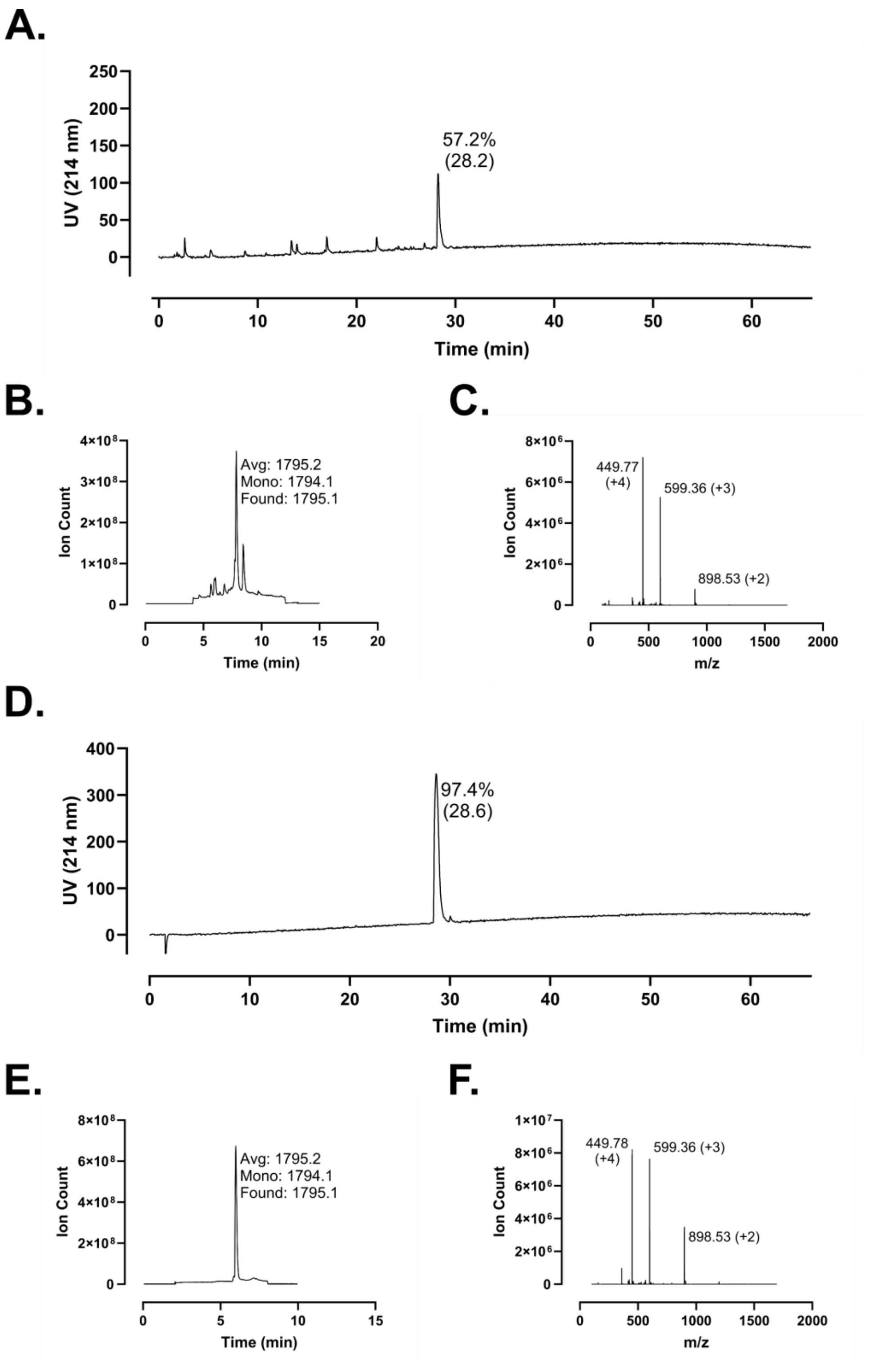
A.
UV (214 nm)
250
200
150
100
50
0
57.2%
(28.2)
0
10
20
30
40
50
60
Time (min)
B.
Ion Count
4×10^8
3×10^8
2×10^8
1×10^8
0
Avg: 1795.2
Mono: 1794.1
Found: 1795.1
0
5
10
15
20
Time (min)
C.
Ion Count
8×10^6
6×10^6
4×10^6
2×10^6
0
449.77
(+4)
599.36 (+3)
898.53 (+2)
0
500
1000
1500
2000
m/z
D.
UV (214 nm)
400
300
200
100
0
97.4%
(28.6)
0
10
20
30
40
50
60
Time (min)
E.
Ion Count
8×10^8
6×10^8
4×10^8
2×10^8
0
Avg: 1795.2
Mono: 1794.1
Found: 1795.1
0
5
10
15
Time (min)
F.
Ion Count
1×10^7
8×10^6
6×10^6
4×10^6
2×10^6
0
449.78
(+4)
599.36 (+3)
898.53 (+2)
0
500
1000
1500
2000
m/z

**Figure S185 – Analytical Data for FF-14 R14B. A.** Crude peptide HPLC trace at 214 nm with percent purity reflecting integration of the dominant peak area; retention time is shown in parentheses. **B.** Crude peptide LCMS total ion chromatogram (TIC); observed mass is provided along with the calculated average and monoisotopic masses. **C.** Mass spectrum at the peak of **B**. Peaks are labeled with *m/z* and charge state. **D.** HPLC of purified peptide. **E.** TIC of purified peptide. **F.** Mass spectrum at the peak of **E**. Peptides were used at the indicated purity following a single round of purification.

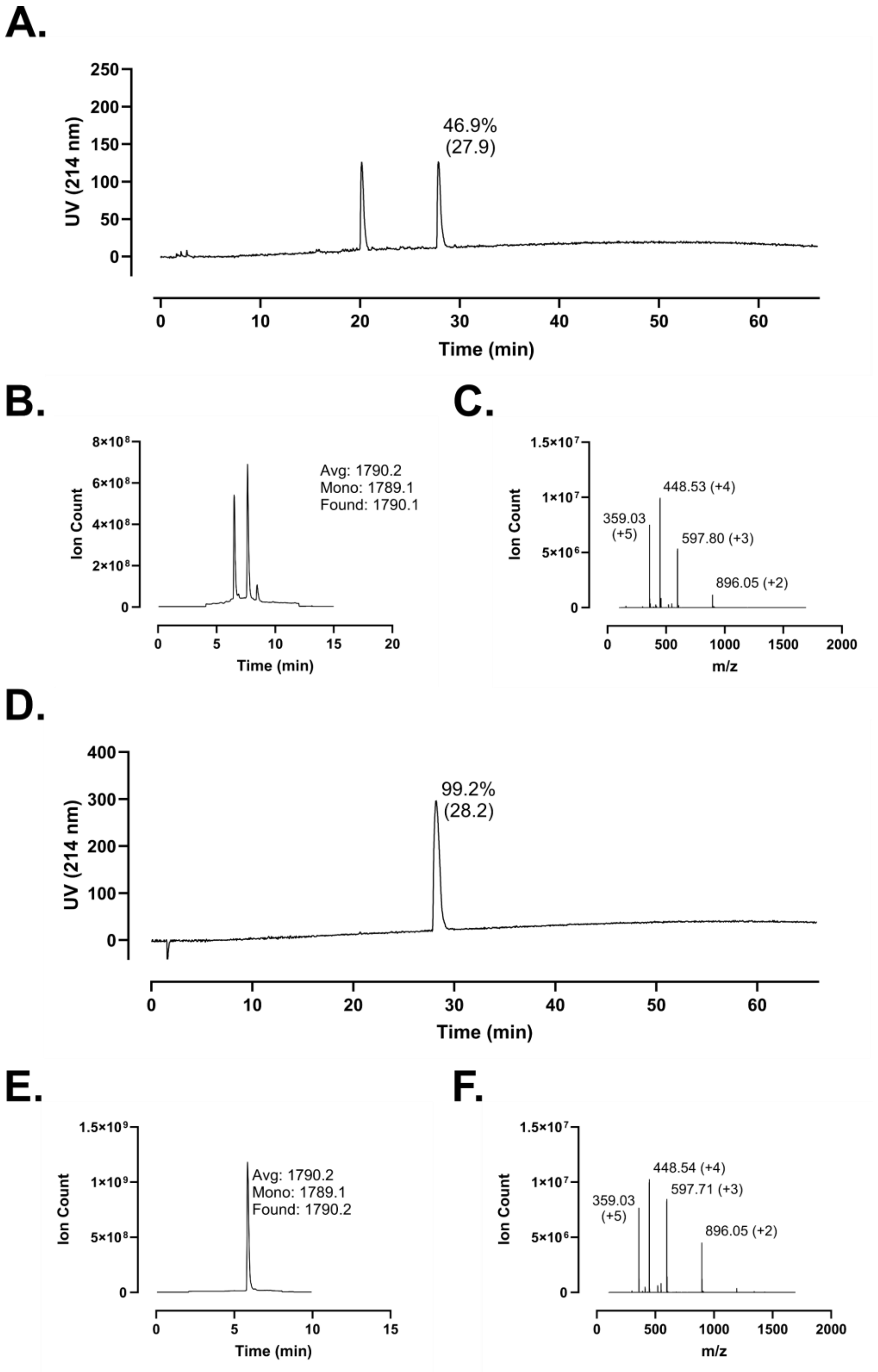
A.
UV (214 nm)
Time (min)
46.9%
(27.9)
B.
Ion Count
Time (min)
Avg: 1790.2
Mono: 1789.1
Found: 1790.1
C.
Ion Count
m/z
448.53 (+4)
359.03
(+5)
597.80 (+3)
896.05 (+2)
D.
UV (214 nm)
Time (min)
99.2%
(28.2)
E.
Ion Count
Time (min)
Avg: 1790.2
Mono: 1789.1
Found: 1790.2
F.
Ion Count
m/z
448.54 (+4)
359.03
(+5)
597.71 (+3)
896.05 (+2)

**Figure S186 – Analytical Data for FF-14 F2B+V6B. A.** Crude peptide HPLC trace at 214 nm with percent purity reflecting integration of the dominant peak area; retention time is shown in parentheses. **B.** Crude peptide LCMS total ion chromatogram (TIC); observed mass is provided along with the calculated average and monoisotopic masses. **C.** Mass spectrum at the peak of **B**. Peaks are labeled with *m/z* and charge state. **D.** HPLC of purified peptide. **E.** TIC from central fraction of purified peptide. **F.** Mass spectrum at the peak of **E**. Peptides were used at the indicated purity following a single round of purification.

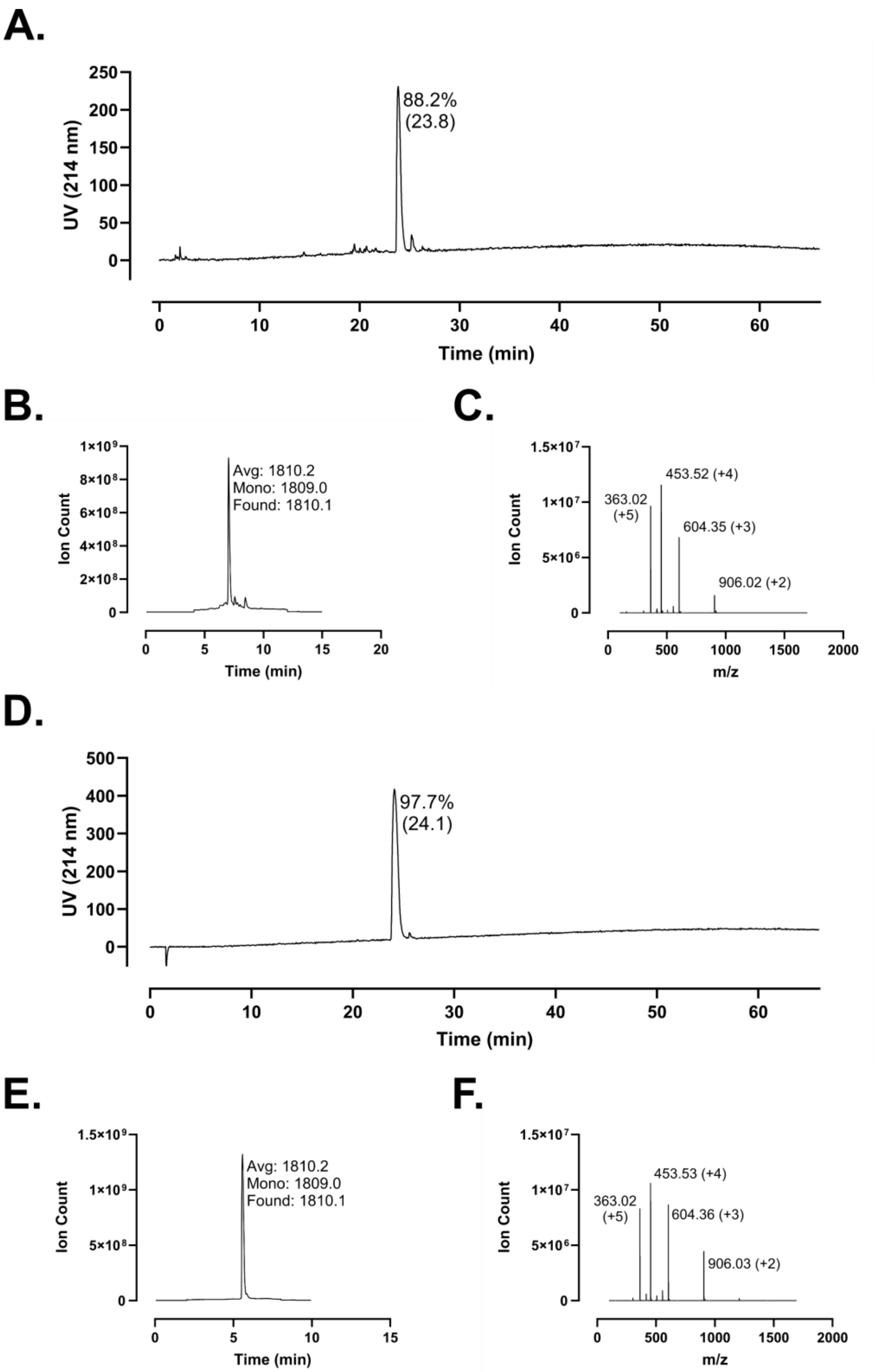
A.
UV (214 nm)
250
200
150
100
50
0
88.2%
(23.8)
0
10
20
30
40
50
60
Time (min)
B.
Ion Count
Avg: 1810.2
Mono: 1809.0
Found: 1810.1
Time (min)
C.
Ion Count
363.02
(+5)
453.52 (+4)
604.35 (+3)
906.02 (+2)
m/z
D.
UV (214 nm)
500
400
300
200
100
0
97.7%
(24.1)
Time (min)
E.
Ion Count
Avg: 1810.2
Mono: 1809.0
Found: 1810.1
Time (min)
F.
Ion Count
363.02
(+5)
453.53 (+4)
604.36 (+3)
906.03 (+2)
m/z

**Figure S187 – Analytical Data for FF-14 I5B+I9B. A.** Crude peptide HPLC trace at 214 nm with percent purity reflecting integration of the dominant peak area; retention time is shown in parentheses. **B.** Crude peptide LCMS total ion chromatogram (TIC); observed mass is provided along with the calculated average and monoisotopic masses. **C.** Mass spectrum at the peak of **B**. Peaks are labeled with *m/z* and charge state. **D.** HPLC of purified peptide. **E.** TIC from central fraction of purified peptide. **F.** Mass spectrum at the peak of **E**. Peptides were used at the indicated purity following a single round of purification

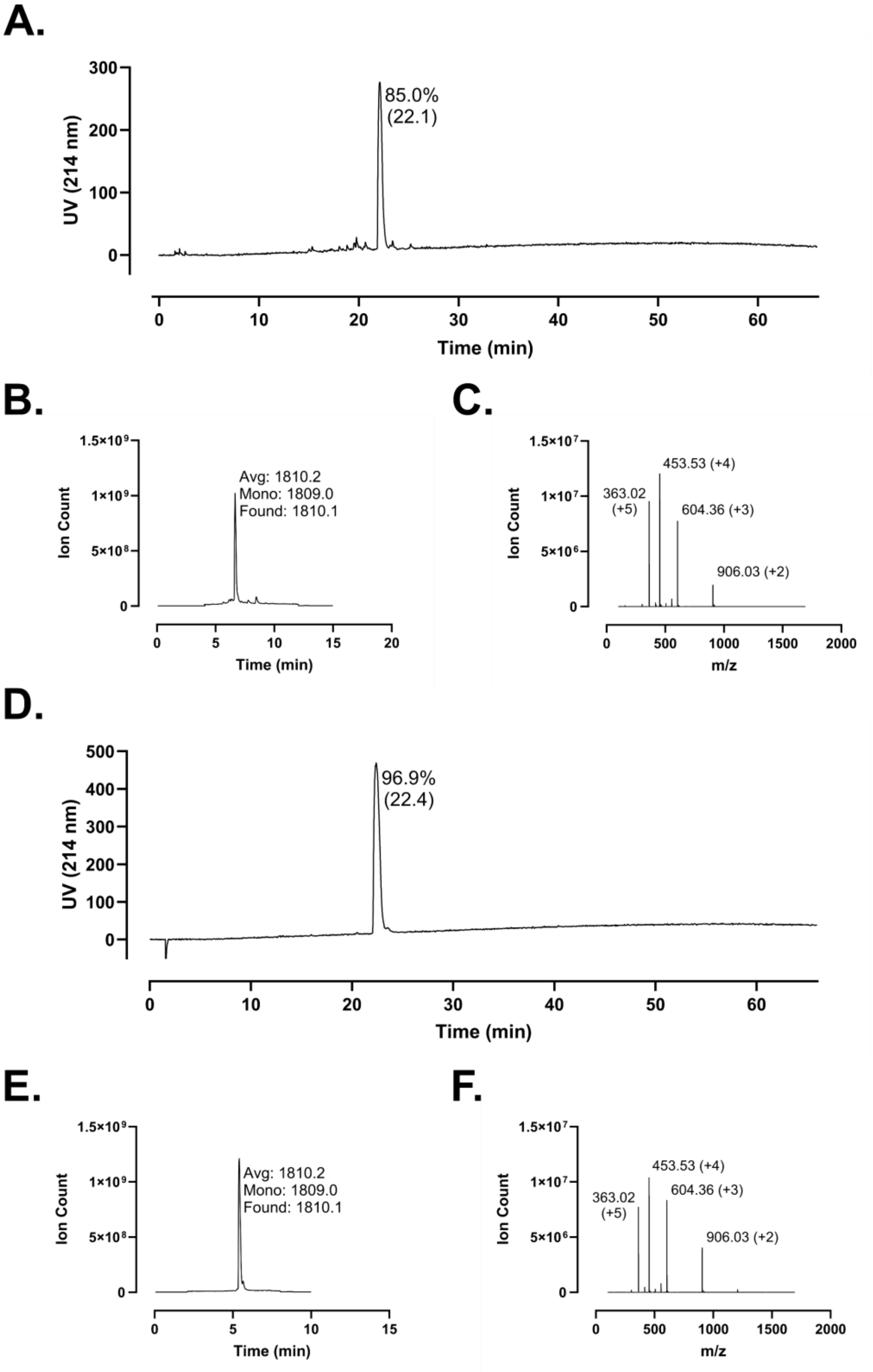
A.
UV (214 nm)
85.0%
(22.1)
Time (min)
B.
Ion Count
Avg: 1810.2
Mono: 1809.0
Found: 1810.1
Time (min)
C.
Ion Count
453.53 (+4)
363.02
(+5)
604.36 (+3)
906.03 (+2)
m/z
D.
UV (214 nm)
96.9%
(22.4)
Time (min)
E.
Ion Count
Avg: 1810.2
Mono: 1809.0
Found: 1810.1
Time (min)
F.
Ion Count
453.53 (+4)
363.02
(+5)
604.36 (+3)
906.03 (+2)
m/z

**Figure S188 – Analytical Data for FF-14 I9B+L13B. A.** Crude peptide HPLC trace at 214 nm with percent purity reflecting integration of the dominant peak area; retention time is shown in parentheses. **B.** Crude peptide LCMS total ion chromatogram (TIC); observed mass is provided along with the calculated average and monoisotopic masses. **C.** Mass spectrum at the peak of **B**. Peaks are labeled with *m/z* and charge state. **D.** HPLC of purified peptide. **E.** TIC from central fraction of purified peptide. **F.** Mass spectrum at the peak of **E**. Peptides were used at the indicated purity following a single round of purification

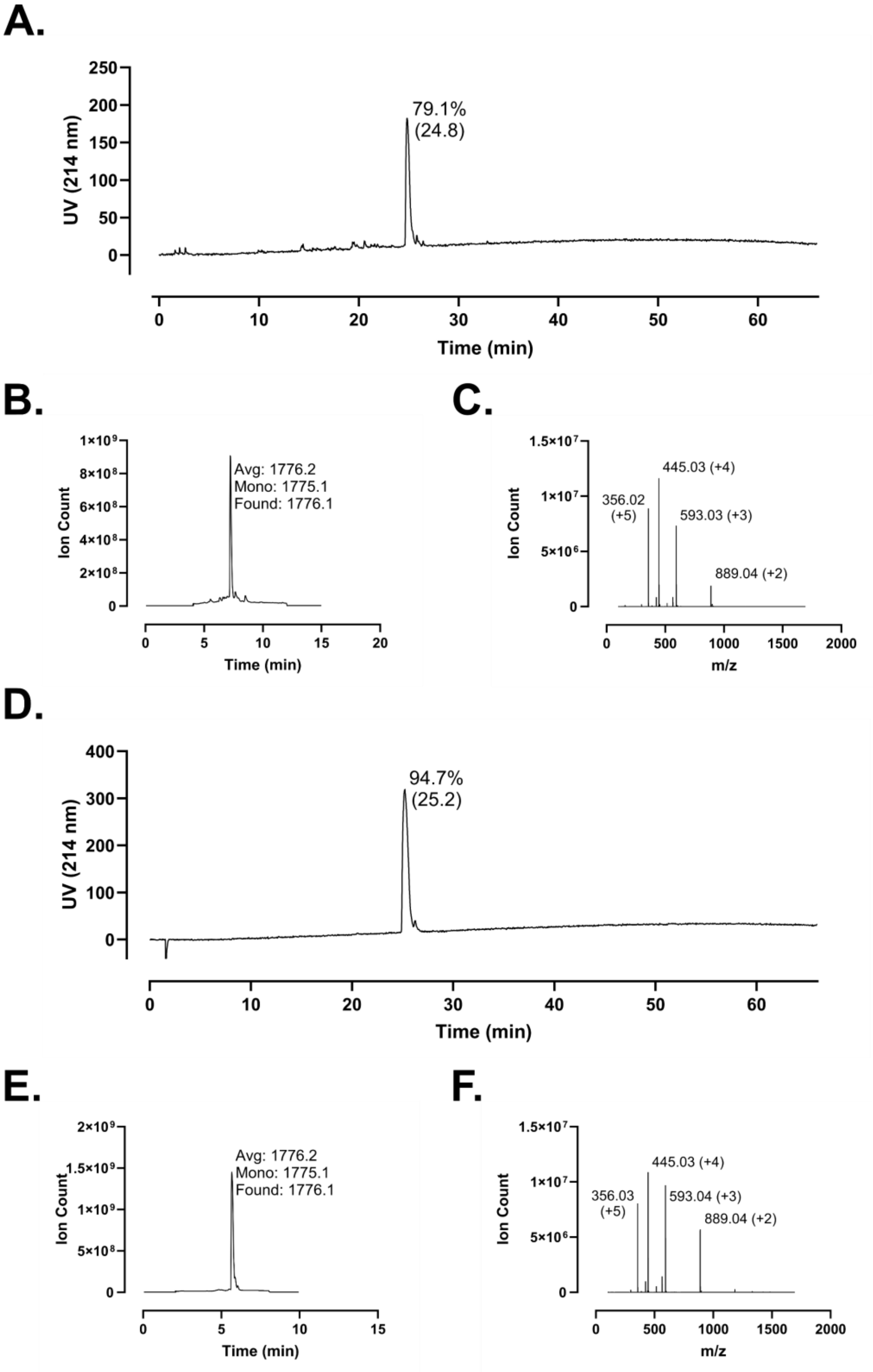
A.
UV (214 nm)
79.1%
(24.8)
Time (min)
B.
Ion Count
Avg: 1776.2
Mono: 1775.1
Found: 1776.1
Time (min)
C.
Ion Count
445.03 (+4)
356.02
(+5)
593.03 (+3)
889.04 (+2)
m/z
D.
UV (214 nm)
94.7%
(25.2)
Time (min)
E.
Ion Count
Avg: 1776.2
Mono: 1775.1
Found: 1776.1
Time (min)
F.
Ion Count
445.03 (+4)
356.03
(+5)
593.04 (+3)
889.04 (+2)
m/z

**Figure S189 – Analytical Data for FF-14 F1B+I5B. A.** Crude peptide HPLC trace at 214 nm with percent purity reflecting integration of the dominant peak area; retention time is shown in parentheses. **B.** Crude peptide LCMS total ion chromatogram (TIC); observed mass is provided along with the calculated average and monoisotopic masses. **C.** Mass spectrum at the peak of **B**. Peaks are labeled with *m/z* and charge state. **D.** HPLC of purified peptide. **E.** TIC from central fraction of purified peptide. **F.** Mass spectrum at the peak of **E**. Peptides were used at the indicated purity following a single round of purification

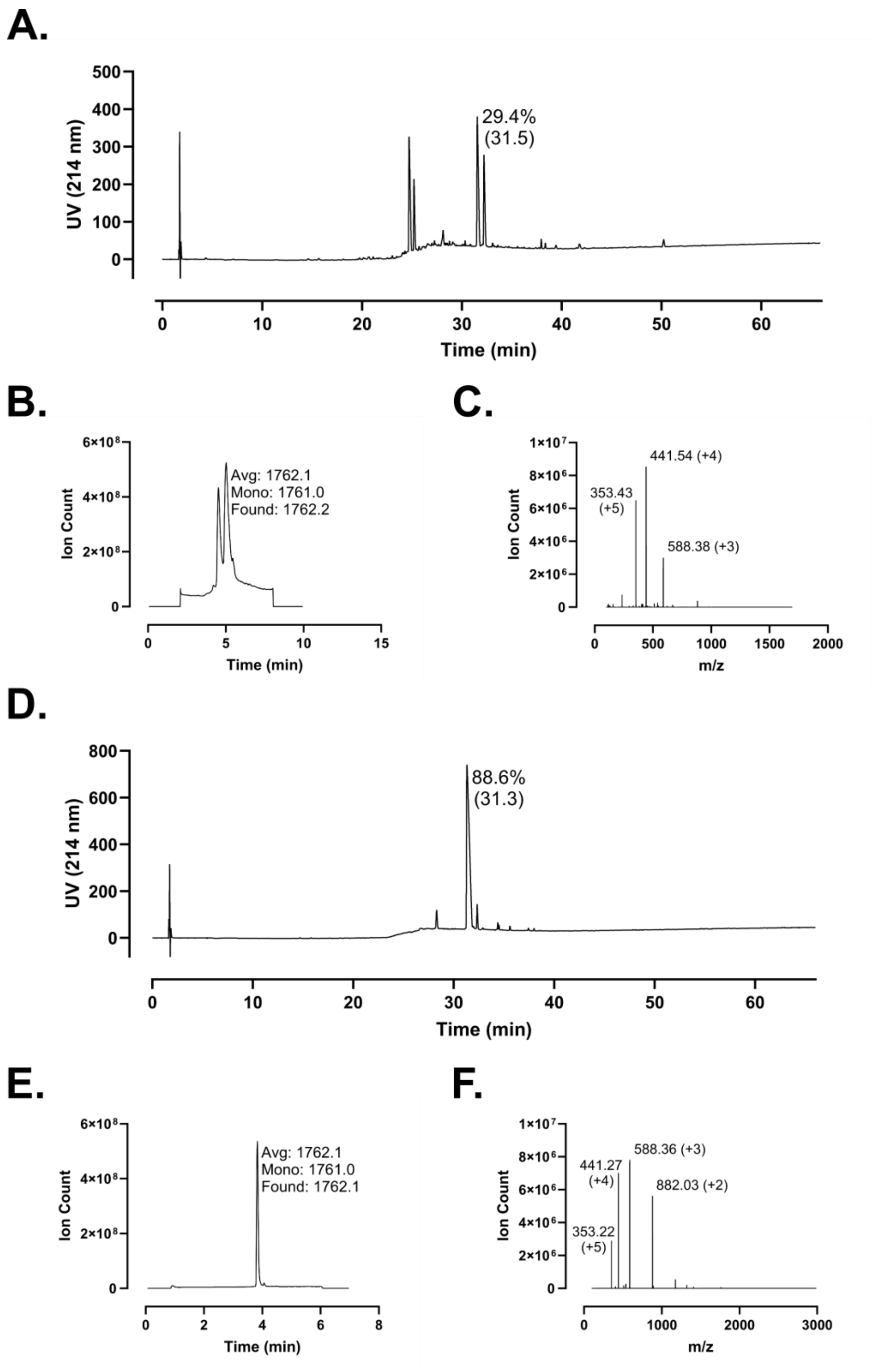
A.
500
400
300
200
100
0
UV (214 nm)
29.4%
(31.5)
0
10
20
30
40
50
60
Time (min)
B.
6×10^8
4×10^8
2×10^8
0
Ion Count
Avg: 1762.1
Mono: 1761.0
Found: 1762.2
0
5
10
15
Time (min)
C.
1×10^7
8×10^6
6×10^6
4×10^6
2×10^6
0
Ion Count
441.54 (+4)
353.43
(+5)
588.38 (+3)
0
500
1000
1500
2000
m/z
D.
800
600
400
200
0
UV (214 nm)
88.6%
(31.3)
0
10
20
30
40
50
60
Time (min)
E.
6×10^8
4×10^8
2×10^8
0
Ion Count
Avg: 1762.1
Mono: 1761.0
Found: 1762.1
0
2
4
6
8
Time (min)
F.
1×10^7
8×10^6
6×10^6
4×10^6
2×10^6
0
Ion Count
441.27
(+4)
588.36 (+3)
882.03 (+2)
353.22
(+5)
0
1000
2000
3000
m/z

**Figure S190 – Analytical Data for FF-14 F2B+V6B+I9B. A.** Crude peptide HPLC trace at 214 nm with percent purity reflecting integration of the dominant peak area; retention time is shown in parentheses. **B.** Crude peptide LCMS total ion chromatogram (TIC); observed mass is provided along with the calculated average and monoisotopic masses. **C.** Mass spectrum at the peak of **B**. Peaks are labeled with *m/z* and charge state. **D.** HPLC of purified peptide. **E.** TIC from central fraction of purified peptide. **F.** Mass spectrum at the peak of **E**. Peptides were used at the indicated purity following a single round of purification

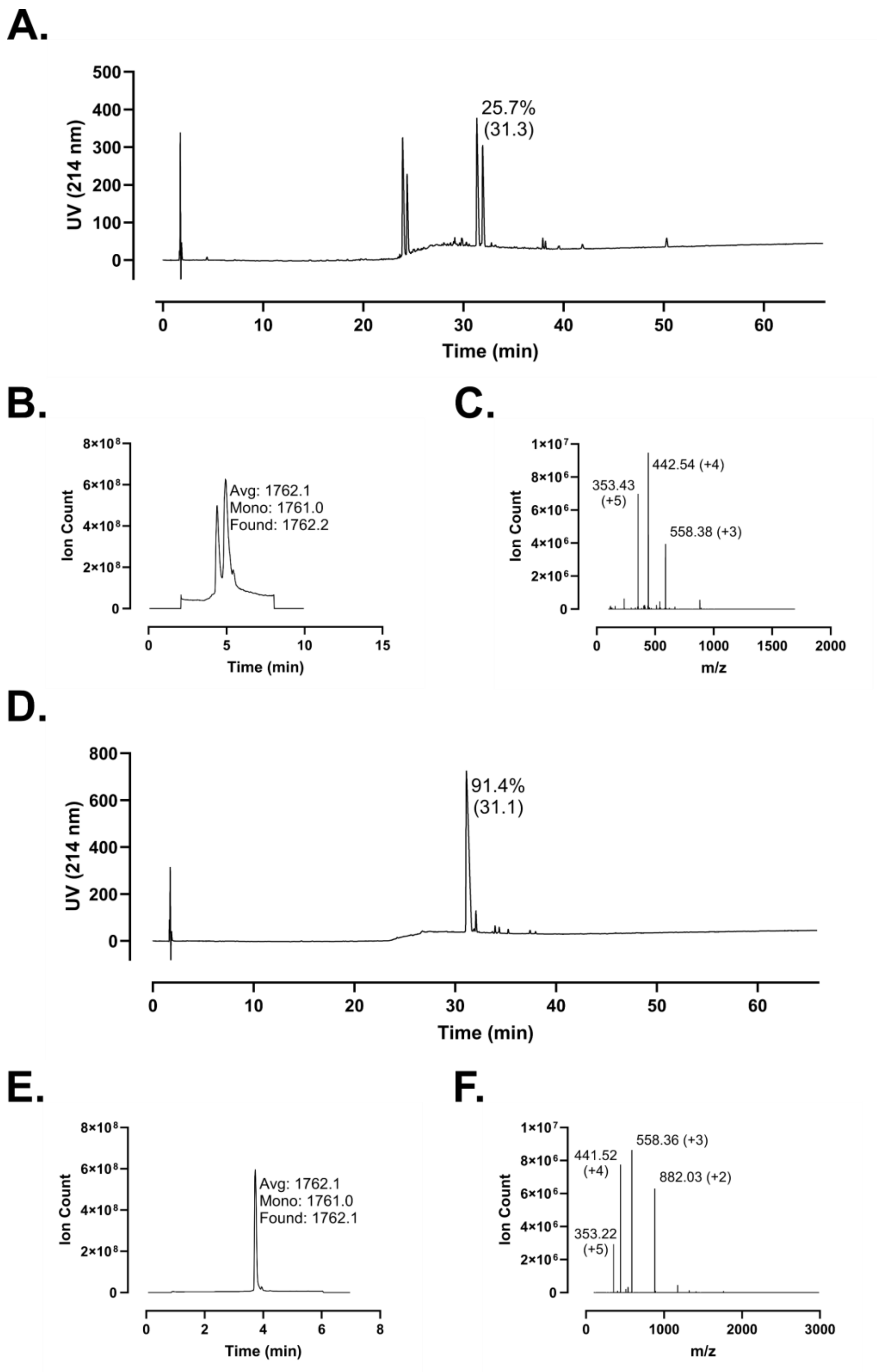
A.
UV (214 nm)
Time (min)
25.7%
(31.3)
B.
Ion Count
Time (min)
Avg: 1762.1
Mono: 1761.0
Found: 1762.2
C.
Ion Count
m/z
353.43
(+5)
442.54 (+4)
558.38 (+3)
D.
UV (214 nm)
Time (min)
91.4%
(31.1)
E.
Ion Count
Time (min)
Avg: 1762.1
Mono: 1761.0
Found: 1762.1
F.
Ion Count
m/z
441.52
(+4)
558.36 (+3)
882.03 (+2)
353.22
(+5)

**Figure S191 – Analytical Data for FF-14 F2B+V6B+L13B. A.** Crude peptide HPLC trace at 214 nm with percent purity reflecting integration of the dominant peak area; retention time is shown in parentheses. **B.** Crude peptide LCMS total ion chromatogram (TIC); observed mass is provided along with the calculated average and monoisotopic masses. **C.** Mass spectrum at the peak of **B**. Peaks are labeled with *m/z* and charge state. **D.** HPLC of purified peptide. **E.** TIC from central fraction of purified peptide. **F.** Mass spectrum at the peak of **E**. Peptides were used at the indicated purity following a single round of purification

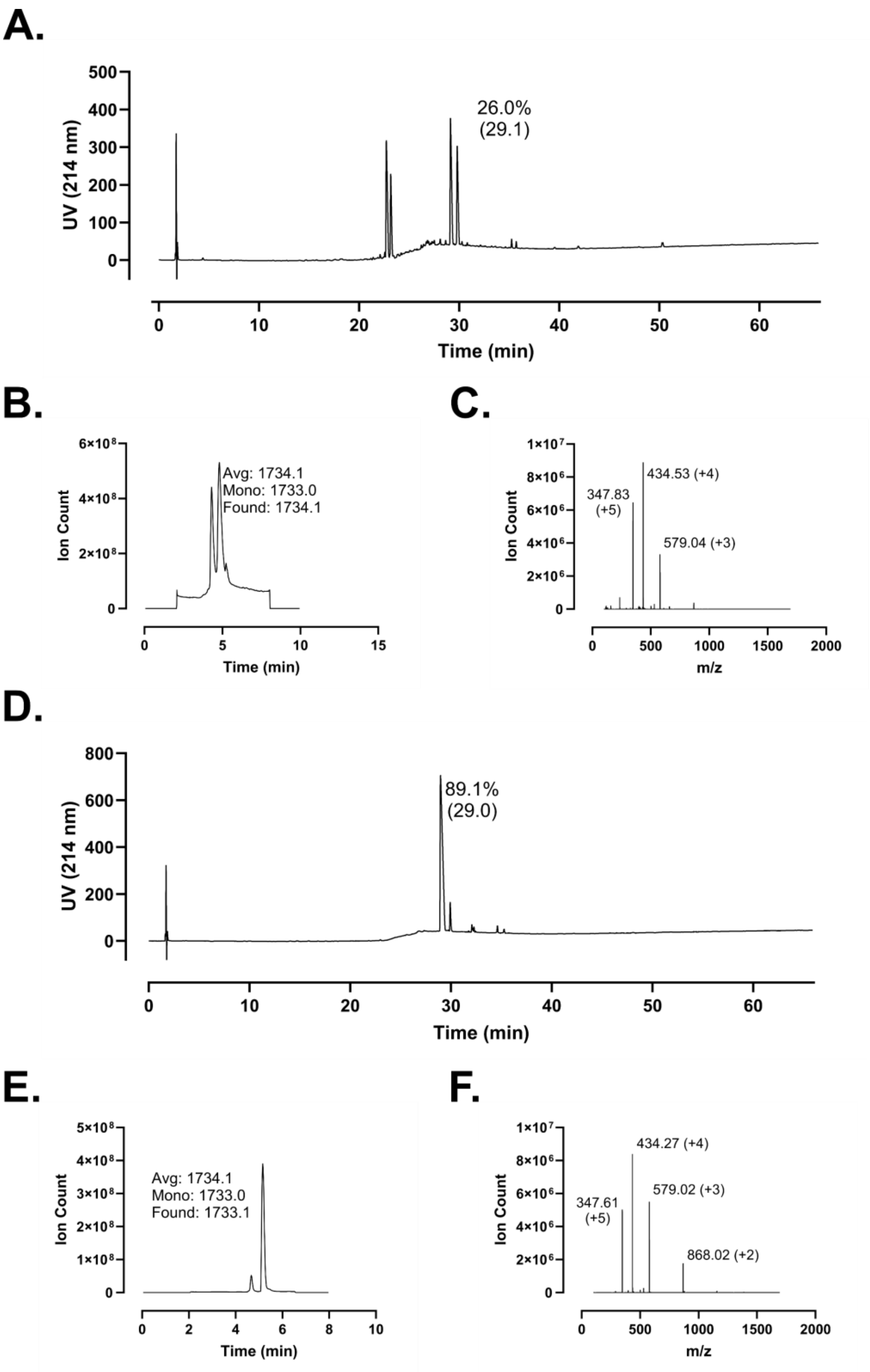
A.
UV (214 nm)
26.0%
(29.1)
Time (min)
B.
Ion Count
Avg: 1734.1
Mono: 1733.0
Found: 1734.1
Time (min)
C.
Ion Count
347.83
(+5)
434.53 (+4)
579.04 (+3)
m/z
D.
UV (214 nm)
89.1%
(29.0)
Time (min)
E.
Ion Count
Avg: 1734.1
Mono: 1733.0
Found: 1733.1
Time (min)
F.
Ion Count
434.27 (+4)
579.02 (+3)
347.61
(+5)
868.02 (+2)
m/z

**Figure S192 – Analytical Data for FF-14 F2B+V6B+I9B+L13B. A.** Crude peptide HPLC trace at 214 nm with percent purity reflecting integration of the dominant peak area; retention time is shown in parentheses. **B.** Crude peptide LCMS total ion chromatogram (TIC); observed mass is provided along with the calculated average and monoisotopic masses. **C.** Mass spectrum at the peak of **B**. Peaks are labeled with *m/z* and charge state. **D.** HPLC of purified peptide. **E.** TIC from central fraction of purified peptide. **F.** Mass spectrum at the peak of **E**. Peptides were used at the indicated purity following a single round of purification

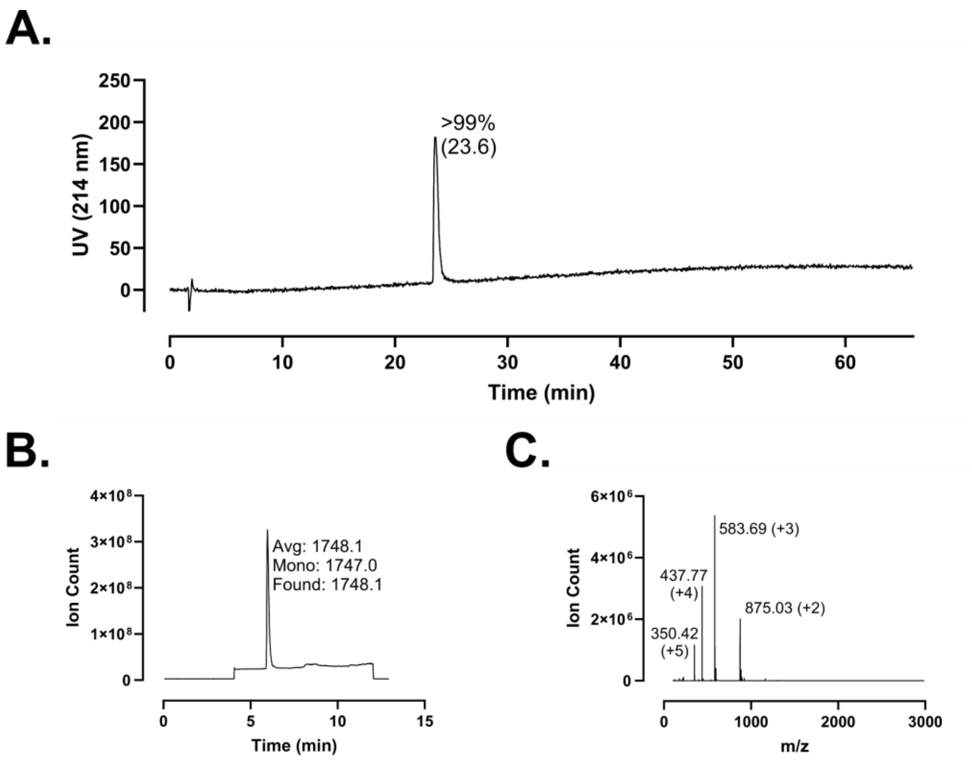
A.
UV (214 nm)
250
200
150
100
50
0
>99%
(23.6)
0
10
20
30
40
50
60
Time (min)
B.
Ion Count
4×10⁸
3×10⁸
2×10⁸
1×10⁸
0
Avg: 1748.1
Mono: 1747.0
Found: 1748.1
0
5
10
15
Time (min)
C.
Ion Count
6×10⁶
4×10⁶
2×10⁶
0
583.69 (+3)
437.77
(+4)
875.03 (+2)
350.42
(+5)
0
1000
2000
3000
m/z

**Figure S193 – Analytical Data for FF-14 F2B+V6B I9A. A.** HPLC trace at 214 nm of purified peptide with percent purity reflecting integration of the dominant peak area; retention time is shown in parentheses. **B.** Purified peptide LCMS total ion chromatogram (TIC); observed mass is provided along with the calculated average and monoisotopic masses. **C.** Mass spectrum at the peak of **B**. Peaks are labeled with *m/z* and charge state. Peptides were used at the indicated purity following a single round of purification.

A.

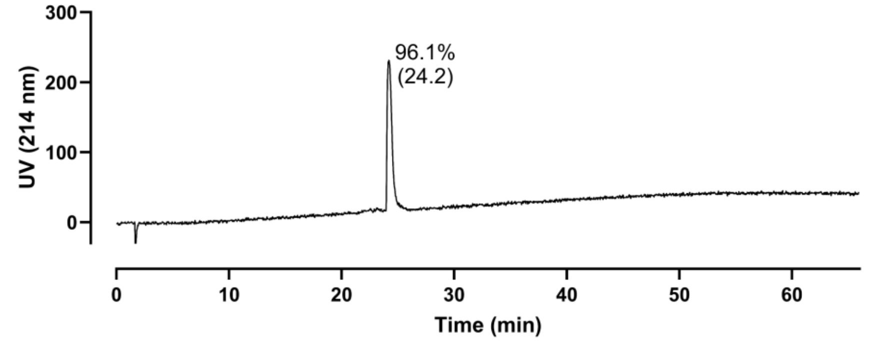

300
200
100
0
UV (214 nm)
96.1%
(24.2)
0
10
20
30
40
50
60
Time (min)


B.

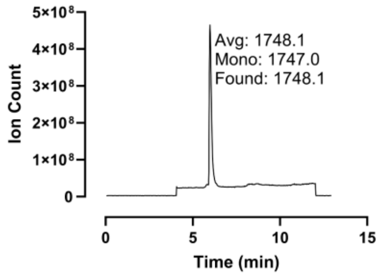

5×10^8
4×10^8
3×10^8
2×10^8
1×10^8
0
Ion Count
Avg: 1748.1
Mono: 1747.0
Found: 1748.1
0
5
10
15
Time (min)


C.

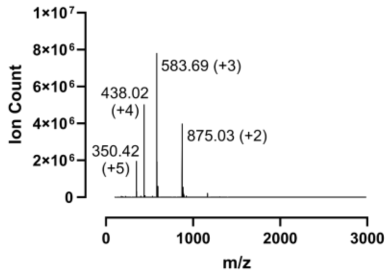

1×10^7
8×10^6
6×10^6
4×10^6
2×10^6
0
Ion Count
583.69 (+3)
438.02
(+4)
875.03 (+2)
350.42
(+5)
0
1000
2000
3000
m/z

**Figure S194 – Analytical Data for FF-14 F2B+V6B L13A. A.** HPLC trace at 214 nm of purified peptide with percent purity reflecting integration of the dominant peak area; retention time is shown in parentheses. **B.** Purified peptide LCMS total ion chromatogram (TIC); observed mass is provided along with the calculated average and monoisotopic masses. **C.** Mass spectrum at the peak of **B**. Peaks are labeled with *m/z* and charge state. Peptides were used at the indicated purity following a single round of purification.

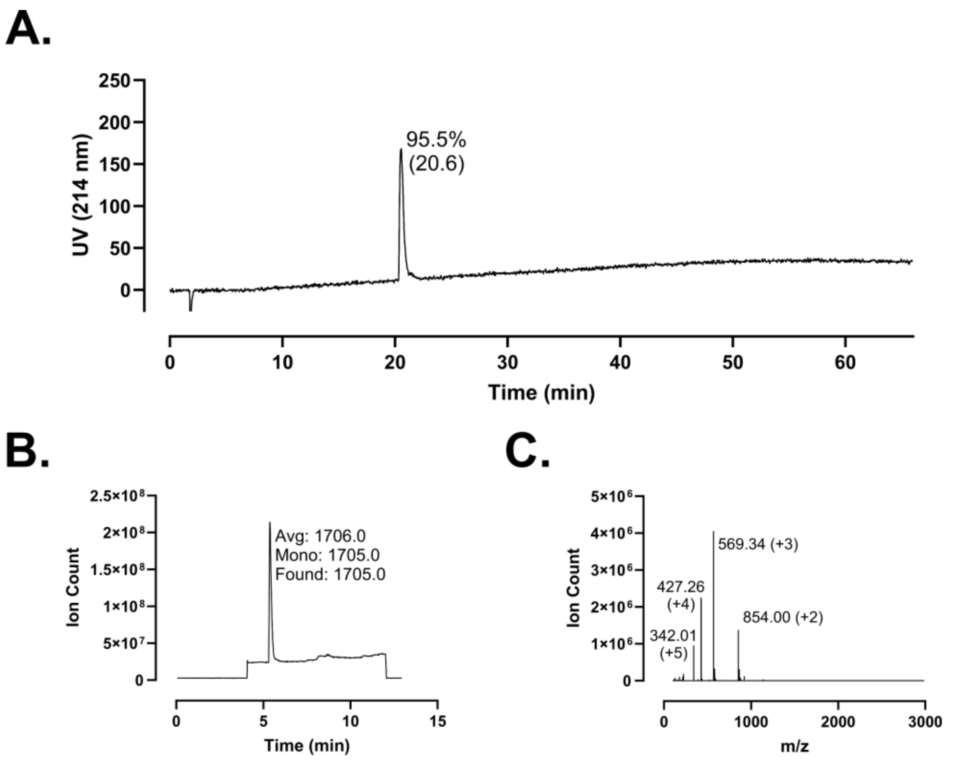
A.
95.5%
(20.6)
UV (214 nm)
Time (min)
B.
Avg: 1706.0
Mono: 1705.0
Found: 1705.0
Ion Count
Time (min)
C.
569.34 (+3)
427.26
(+4)
854.00 (+2)
342.01
(+5)
Ion Count
m/z

**Figure S195 – Analytical Data for FF-14 F2B+V6B I9A+L13A. A.** HPLC trace at 214 nm of purified peptide with percent purity reflecting integration of the dominant peak area; retention time is shown in parentheses. **B.** Purified peptide LCMS total ion chromatogram (TIC); observed mass is provided along with the calculated average and monoisotopic masses. **C.** Mass spectrum at the peak of **B**. Peaks are labeled with *m/z* and charge state. Peptides were used at the indicated purity following a single round of purification.

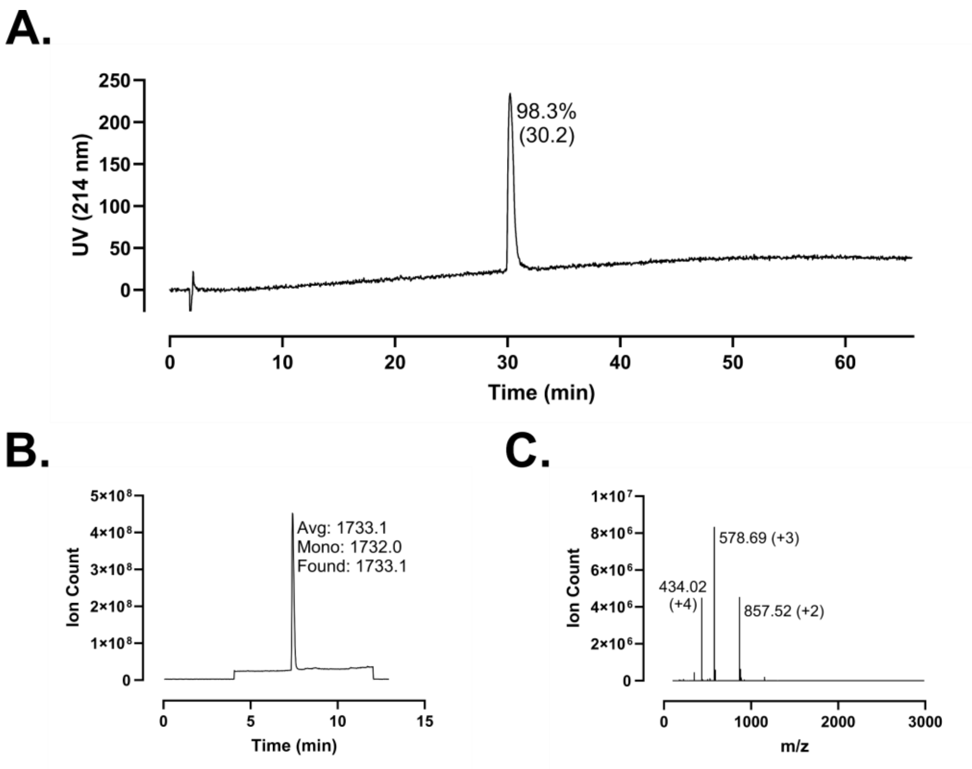
A.
98.3%
(30.2)
UV (214 nm)
Time (min)
B.
Avg: 1733.1
Mono: 1732.0
Found: 1733.1
Ion Count
Time (min)
C.
578.69 (+3)
434.02
(+4)
857.52 (+2)
Ion Count
m/z

**Figure S196 – Analytical Data for FF-14 F2B+V6B K3A. A.** HPLC trace at 214 nm of purified peptide with percent purity reflecting integration of the dominant peak area; retention time is shown in parentheses. **B.** Purified peptide LCMS total ion chromatogram (TIC); observed mass is provided along with the calculated average and monoisotopic masses. **C.** Mass spectrum at the peak of **B**. Peaks are labeled with *m/z* and charge state. Peptides were used at the indicated purity following a single round of purification.

A.

B.

C.

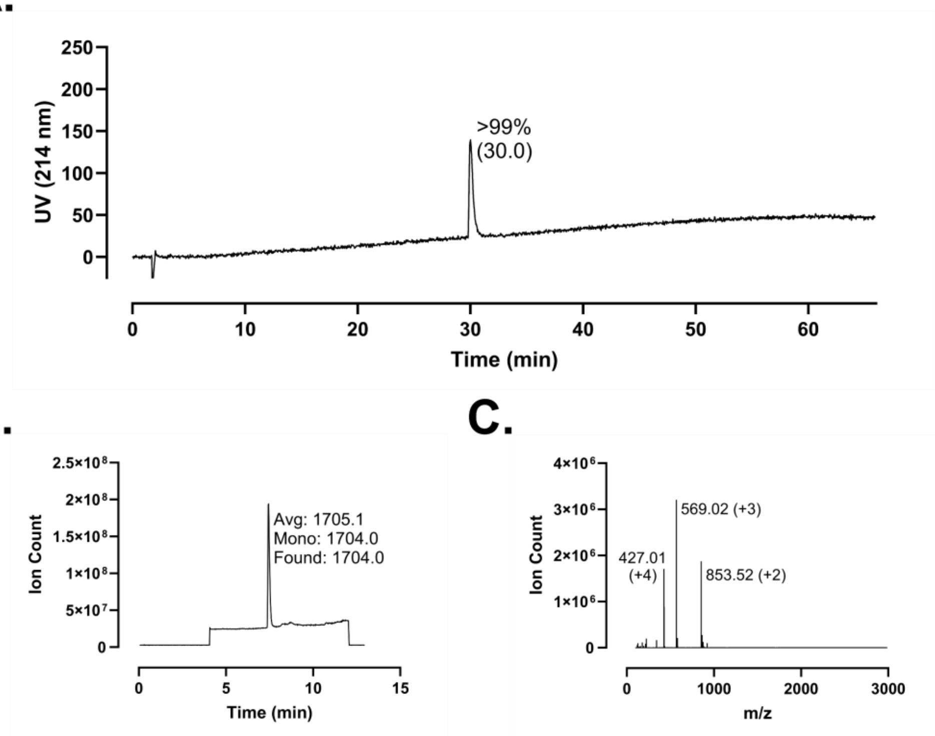

250
200
150
100
50
0
UV (214 nm)
>99%
(30.0)
0
10
20
30
40
50
60
Time (min)
2.5×10^8
2×10^8
1.5×10^8
1×10^8
5×10^7
0
Ion Count
Avg: 1705.1
Mono: 1704.0
Found: 1704.0
0
5
10
15
Time (min)
4×10^6
3×10^6
2×10^6
1×10^6
0
Ion Count
569.02 (+3)
427.01
(+4)
853.52 (+2)
0
1000
2000
3000
m/z

**Figure S197 – Analytical Data for FF-14 F2B+V6B R4A. A.** HPLC trace at 214 nm of purified peptide with percent purity reflecting integration of the dominant peak area; retention time is shown in parentheses. **B.** Purified peptide LCMS total ion chromatogram (TIC); observed mass is provided along with the calculated average and monoisotopic masses. **C.** Mass spectrum at the peak of **B**. Peaks are labeled with *m/z* and charge state. Peptides were used at the indicated purity following a single round of purification.

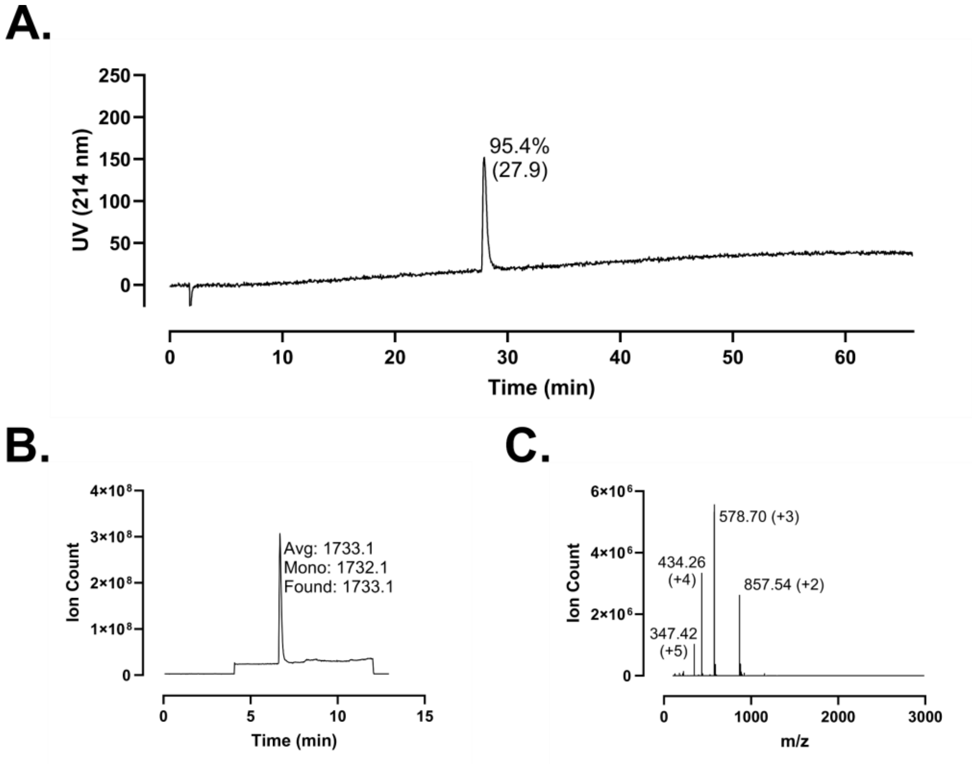
A.
UV (214 nm)
95.4%
(27.9)
Time (min)
B.
Ion Count
Avg: 1733.1
Mono: 1732.1
Found: 1733.1
Time (min)
C.
Ion Count
578.70 (+3)
434.26
(+4)
857.54 (+2)
347.42
(+5)
m/z

**Figure S198 – Analytical Data for FF-14 F2B+V6B Q7A. A.** HPLC trace at 214 nm of purified peptide with percent purity reflecting integration of the dominant peak area; retention time is shown in parentheses. **B.** Purified peptide LCMS total ion chromatogram (TIC); observed mass is provided along with the calculated average and monoisotopic masses. **C.** Mass spectrum at the peak of **B**. Peaks are labeled with *m/z* and charge state. Peptides were used at the indicated purity following a single round of purification.

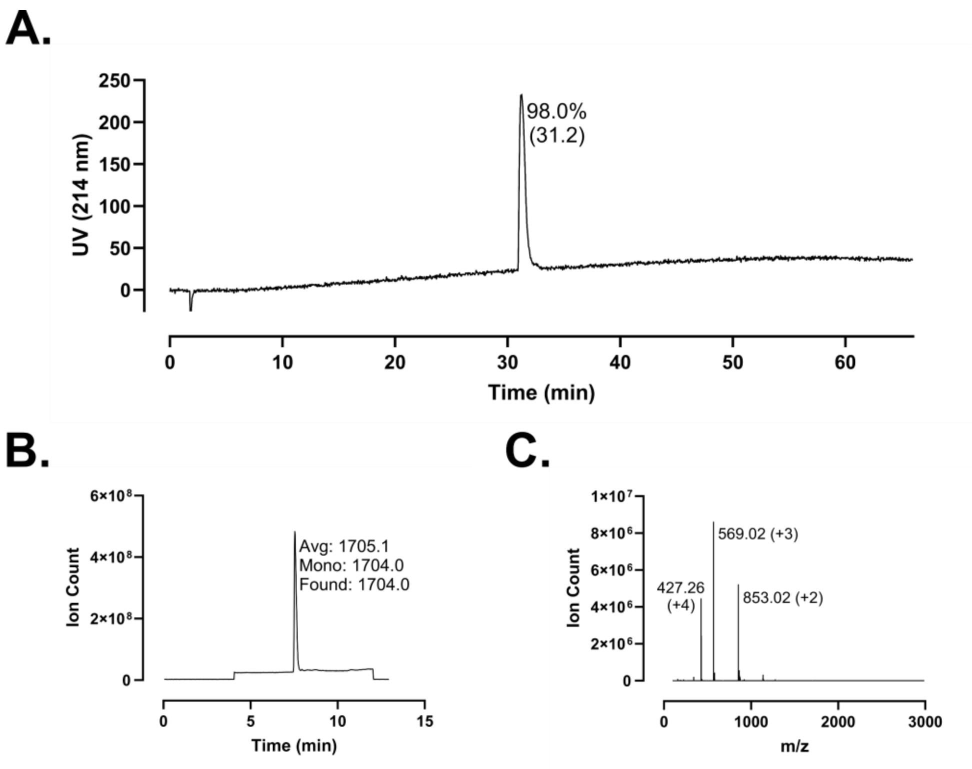
A.
UV (214 nm)
98.0%
(31.2)
Time (min)
B.
Ion Count
Avg: 1705.1
Mono: 1704.0
Found: 1704.0
Time (min)
C.
Ion Count
569.02 (+3)
427.26
(+4)
853.02 (+2)
m/z

**Figure S199 – Analytical Data for FF-14 F2B+V6B R8A. A.** HPLC trace at 214 nm of purified peptide with percent purity reflecting integration of the dominant peak area; retention time is shown in parentheses. **B.** Purified peptide LCMS total ion chromatogram (TIC); observed mass is provided along with the calculated average and monoisotopic masses. **C.** Mass spectrum at the peak of **B**. Peaks are labeled with *m/z* and charge state. Peptides were used at the indicated purity following a single round of purification.

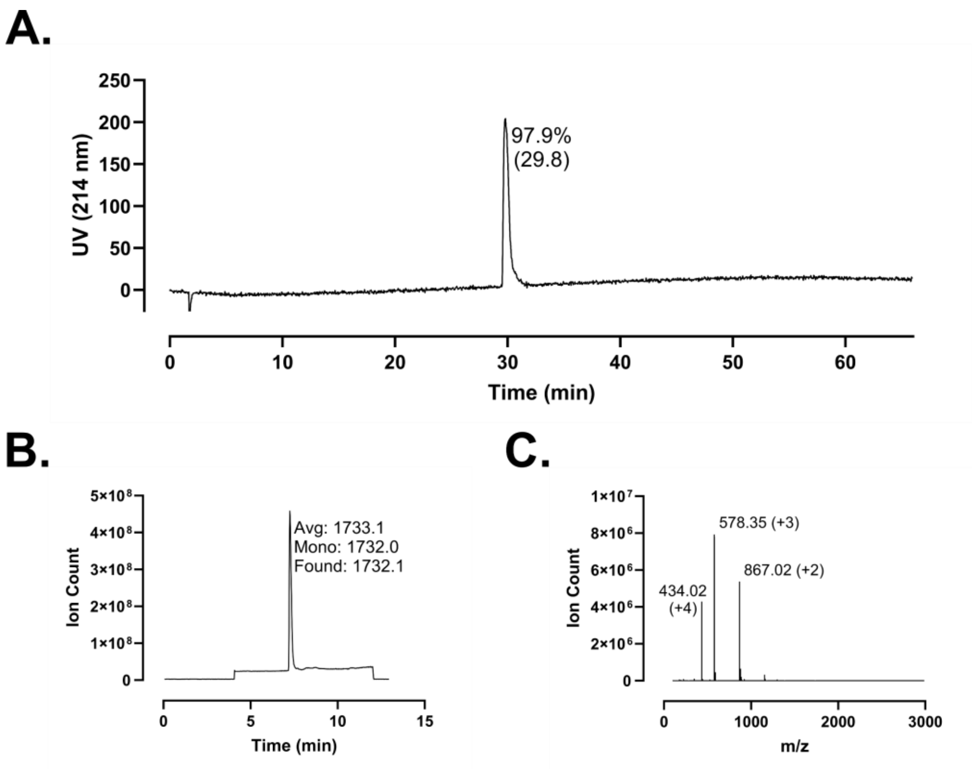
A.
97.9%
(29.8)
UV (214 nm)
Time (min)
B.
Avg: 1733.1
Mono: 1732.0
Found: 1732.1
Ion Count
Time (min)
C.
578.35 (+3)
867.02 (+2)
434.02
(+4)
Ion Count
m/z

**Figure S200 – Analytical Data for FF-14 F2B+V6B K10A. A.** HPLC trace at 214 nm of purified peptide with percent purity reflecting integration of the dominant peak area; retention time is shown in parentheses. **B.** Purified peptide LCMS total ion chromatogram (TIC); observed mass is provided along with the calculated average and monoisotopic masses. **C.** Mass spectrum at the peak of **B**. Peaks are labeled with *m/z* and charge state. Peptides were used at the indicated purity following a single round of purification.

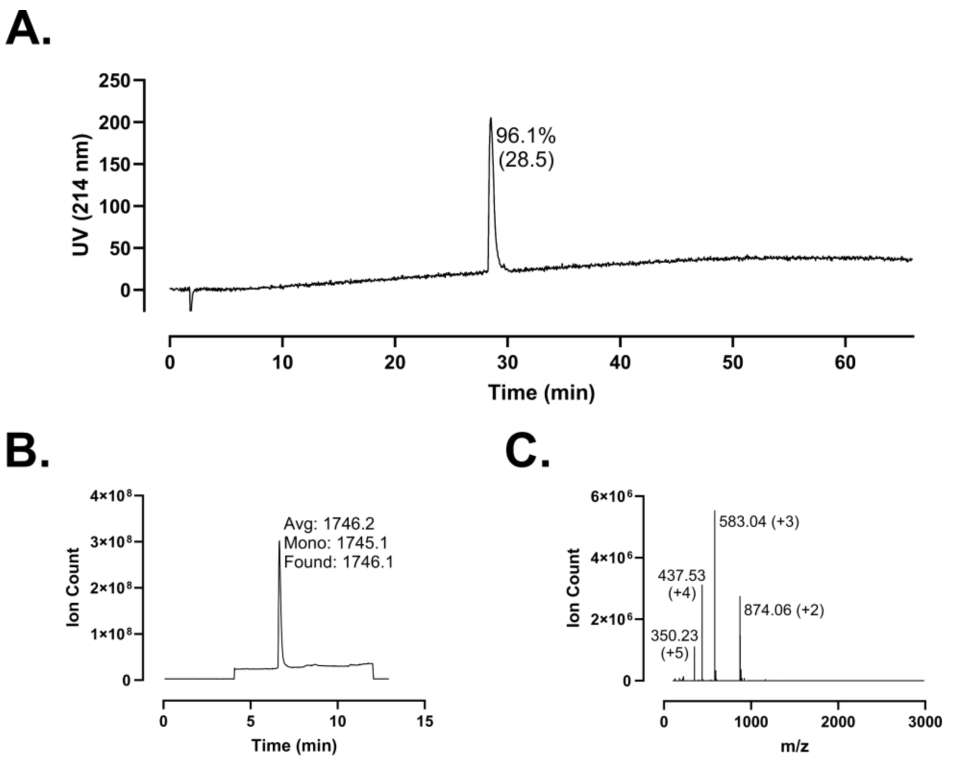
A.
250
200
150
100
50
0
UV (214 nm)
96.1%
(28.5)
0
10
20
30
40
50
60
Time (min)
B.
4×10⁸
3×10⁸
2×10⁸
1×10⁸
0
Ion Count
Avg: 1746.2
Mono: 1745.1
Found: 1746.1
0
5
10
15
Time (min)
C.
6×10⁶
4×10⁶
2×10⁶
0
Ion Count
583.04 (+3)
437.53
(+4)
874.06 (+2)
350.23
(+5)
0
1000
2000
3000
m/z

**Figure S201 – Analytical Data for FF-14 F2B+V6B D11A. A.** HPLC trace at 214 nm of purified peptide with percent purity reflecting integration of the dominant peak area; retention time is shown in parentheses. **B.** Purified peptide LCMS total ion chromatogram (TIC); observed mass is provided along with the calculated average and monoisotopic masses. **C.** Mass spectrum at the peak of **B**. Peaks are labeled with *m/z* and charge state. Peptides were used at the indicated purity following a single round of purification.

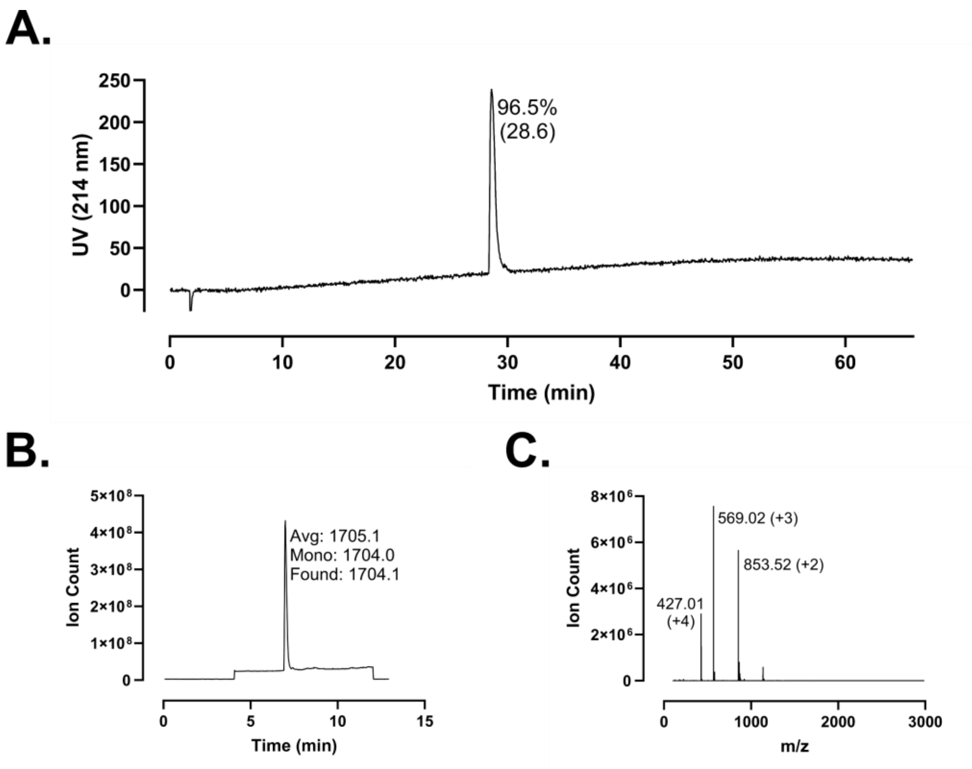
A.
250
200
150
100
50
0
UV (214 nm)
96.5%
(28.6)
0
10
20
30
40
50
60
Time (min)
B.
5×10⁸
4×10⁸
3×10⁸
2×10⁸
1×10⁸
0
Ion Count
Avg: 1705.1
Mono: 1704.0
Found: 1704.1
0
5
10
15
Time (min)
C.
8×10⁶
6×10⁶
4×10⁶
2×10⁶
0
Ion Count
569.02 (+3)
853.52 (+2)
427.01
(+4)
0
1000
2000
3000
m/z

**Figure S202 – Analytical Data for FF-14 F2B+V6B R14A. A.** HPLC trace at 214 nm of purified peptide with percent purity reflecting integration of the dominant peak area; retention time is shown in parentheses. **B.** Purified peptide LCMS total ion chromatogram (TIC); observed mass is provided along with the calculated average and monoisotopic masses. **C.** Mass spectrum at the peak of **B**. Peaks are labeled with *m/z* and charge state. Peptides were used at the indicated purity following a single round of purification.
